# Structure-Uptake Relationship Studies of Oxazolidinones in Gram-negative ESKAPE Pathogens

**DOI:** 10.1101/2022.06.27.497815

**Authors:** Ziwei Hu, Inga V. Leus, Brinda Chandar, Bradley Sherborne, Quentin P. Avila, Valentin V. Rybenkov, Helen I. Zgurskaya, Adam S. Duerfeldt

**Author notes:** Corresponding Authors Helen I. Zgurskaya, Department of Chemistry & Biochemistry, University of Oklahoma, 101 Stephenson Parkway, Norman, OK, 73019., Adam S. Duerfeldt, Department of Medicinal Chemistry, University of Minnesota, 717 Delaware Street SE, Minneapolis, MN, 55455. Contributed equally (Ziwei Hu, Inga V. Leus, Brinda Chandar).

## Abstract

To date, little is known about applicability and/or generality of molecular features and how they impact small molecule permeation into Gram-negative bacteria. Identifying motifs or structural trends that correlate with broad and/or species-specific permeation would enable the rational design of new antibacterials. The clinical success of linezolid for treating Gram-positive infections paired with the high conservation of bacterial ribosomes predicts that if oxazolidinones were engineered to accumulate in Gram-negative bacteria, then this pharmacological class would find broad utility in eradicating infections. Here we report an investigative study of a strategically designed library of oxazolidinones to determine the effects of molecular structure on accumulation and biological activity. *E. coli, A. baumannii*, and *P. aeruginosa* strains with varying degrees of compromise (in efflux and outer membrane) were used to identify motifs that hinder permeation across the outer-membrane and/or enhance efflux susceptibility broadly and specifically between species. The results of this study illustrate that small changes in molecular structure are enough to overcome the efflux and/or permeation issues of this scaffold. Three oxazolidinone analogs (**3e**, **12f**, and **14**) were identified from this study that exhibit activity against all three pathogens assessed, a biological profile not observed for linezolid.

## Introduction

The emergence and widespread prevalence of multidrug-resistant (MDR) bacteria pose a great threat to society.^1–2^ A 2019 report from the Centers for Disease Control and Prevention (CDC) estimates that >2.8 million MDR infections occur annually in the U.S., giving rise to >35,000 deaths.^3^ MDR Gram-negative bacteria (GNB) have been pinpointed as the most urgent threats.^1, 4–5^ In fact, four (*K. pneumoniae*, *A. baumannii*, *P. aeruginosa*, and *Enterobacter spp.*) of the six leading contributors to hospital-acquired infections are GNB.^6^

It is well-established that the encapsulation of the hydrophobic inner membrane by an asymmetric outer membrane (OM) presents a rather formidable barrier to small molecule permeation in GNB. Molecules must either be amphipathic enough to permeate these membranes with contrasting properties or capitalize on porin-mediated entry. Additionally, after a compound successfully crosses the outer or both membranes, it is subject to highly promiscuous efflux pumps.^7–8^ As a result, while potent biochemical inhibitors can often be identified for new targets, developing them into compounds with whole-cell antibacterial activity has proven challenging.^9–11^ A major roadblock to the development of novel antibiotics is our poor understanding of the structural features of small molecules that correlate with GNB permeation and accumulation.

Retrospective efforts to summarize physicochemical properties that coincide with Gram-negative activity^9, 11–12^ have enabled rational approaches to enhance Gram-negative compatibility of compounds in a prospective manner. In fact, predictive models for chemical modification to impart activity against *E. coli* through porin mediated interactions has been achieved^13–14^ and employed for diverse chemotypes.^15–23^ This growing body of work demonstrates that even though difficult, it is feasible to convert Gram-positive specific chemotypes into broader spectrum agents. In addition to characterizing physicochemical properties that influence accumulation in GNB, it is equally important to identify structural motifs that positively or negatively affect accumulation. Identifying such motifs would provide isosteric-like replacements that can be leveraged to enhance accumulation, much like the utilization of motifs to improve pharmacokinetic properties of drugs for oral bioavailability.

Although the basic envelope composition is relatively conserved across GNB, significant variation exists between species and strains,^24^ which limits the generalization and extension of physicochemical predictors determined for *E. coli* to other pathogens. As such, it is crucial to analyze compound behavior across not only species, but also strains, to gain insight into generally applicable “rules” versus those that are more organism specific for each chemotype. While it is worth examining newly discovered chemical scaffolds in these efforts, critical information can also be obtained from well interrogated compound families, as the technology now exists to deconvolute permeation and efflux liabilities that lacked during the original period of chemotype development.

Oxazolidinones, for example, were rationally developed in the 1990s as a new class of synthetic antibiotics that exert their activity by binding to the 23S rRNA of the 50S ribosomal subunit and inhibiting the initiation step of bacterial protein synthesis.^25^ Linezolid (**Figure 1**), first approved in 2000, was the leading branded antibiotic for several resistant Gram-positive infections including methicillin- and vancomycin-resistant staphylococci, vancomycin-resistant enterococci, and penicillin-resistant pneumococci.^26^ However, within 10 years, linezolid resistance was detected in >1% of *S. aureus* clinical isolates.^27^ Linezolid binds to a common antibiotic site on the ribosome and acts by blocking precise positioning of A-site tRNA in the peptidyl transferase center (PTC). It partially shares the site with other antibiotics that inhibit protein synthesis and hence, exhibits overlapping mechanisms of resistance. The major mechanisms of resistance include various mutations in domain V of 23S rRNA and ribosomal proteins L3 (*rplC*) and L4 (*rplD*).^28–29^ In addition, RNA methyltransferases encoded on various mobile elements were found to contribute to linezolid resistance.^30^ Importantly, the pattern of resistance is organism specific in that the obtained mutation sites differ, with little overlap, between different bacterial species.^28, 31^ This specificity is likely related to 1) single-base identity differences in the RNA nucleotides that are more distant from and do not directly contact the bound linezolid molecule and 2) variations in the fitness costs associated with specific mutations in different organisms. The degree of resistance conferred by the mutations in 23S rRNA is not a simple function of the nucleotide-linezolid distance, as mutations at distal nucleotides that do not interact with linezolid directly, in particular, G2576U and G2447U, confer significant resistance.^28, 30^ The only clear non-ribosomal mechanism of resistance in Gram-positive bacteria is related to mutations leading to overproduction of efflux pumps, specifically the LmrS pump in *S. aureus*^32^ and ABC transporter genes in *Streptococcus pneumoniae* ^33^. The contribution of efflux is more significant in Gram-negative bacteria due to its synergistic interactions with the low permeability barrier of the outer membrane.

**Figure 1.**
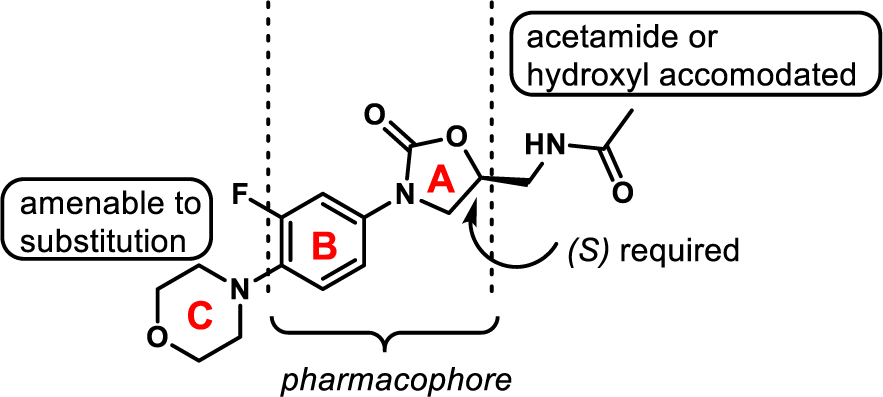
Structure of linezolid and general SAR features of the oxazolidinone chemotype.

The clinical success of linezolid, yet susceptibility to resistance, has inspired pharmaceutical companies and academic groups to develop next-generation oxazolidinones.^34–36^ Thus far, studies have focused mainly on overcoming target-driven linezolid-resistance,^37–39^ exploring the utility in infections of the central nervous system (CNS),^40^ and reducing toxic side effects.^41^ However, the potential of this chemotype for broad-spectrum activity has been underexplored due to the Gram-positive specific activity and inability, thus far, to rationally design GNB active derivatives.^42–45^ High conservation of bacterial ribosomes, however, predict that if oxazolidinones were engineered to accumulate in GNB, then this pharmacological class would find great utility in eradicating infections.^45^ These reasons, paired with the knowledge that the biological activity of this chemotype can be maintained through broad C-ring (**Figure 1**) diversification, establishes the oxazolidinones as a seminal class to interrogate the effect of motif variation on GNB accumulation.

Herein we report the design, synthesis, and evaluation of oxazolidinones distinctly functionalized on the C-ring. Activity against wild type *E. coli*, *P. aeruginosa*, and *A. baumannii*, and corresponding strains with varying degrees of outer membrane and/or efflux pump efficiencies were employed to allow for the deconvolution of structure-uptake relationships (SURs). The activity against *S. aureus* and linezolid-resistant *E. coli* was used to verify target engagement of representative analogs. Motifs that prove problematic to OM permeation and/or efflux for each species were revealed. Three linezolid analogs were identified that exhibit broad spectrum activity against all three pathogens.

## RESULTS AND DISCUSSION

### Molecular Design

An oxazolidinone A-ring containing an (*S*) C-5 one-carbon linked acetamide and an *N*-aryl substituent (B-ring) comprise the bulk of the scaffold required for target engagement (**Figure 1**). A third ring system (C-ring) is appended to the *para*-position of the B-ring. Previous studies clearly demonstrate that the C-ring is the most amenable to modification and that the C-5 acetamide can be replaced with a more synthetically tractable hydroxyl group (**Figure 1**). As such, we aimed to design a library capable of leveraging known chemistry to provide a common intermediate (**2**, **Scheme 1**) that could be exposed to diverse reactants with broad coverage of physicochemical attributes. We recognized however that while the co-crystal structure of linezolid bound to the *S. aureus* 50S (SA50Slin) subunit indicates binding at the PTC, blocking the A-site in an orientation similar to that observed in other ribosome linezolid complexes, a noteworthy difference is apparent. In the SA50Slin complex, the flexible nucleotide U2585 undergoes significant rotation and forms a hydrogen bond with the oxygen of the linezolid morpholine ring, leading to a nonproductive conformation of the PTC.^46^ This is different from other ribosome linezolid complexes including *Deinococcus radiodurans* 50S, the whole ribosome of *Thermus thermophilus*, as well as the *E. coli* 70S structure. Hence, we recognized that alteration of the C-ring could affect target engagement and, by extension, antibacterial activity of linezolid-derivatives in a species-specific manner.

The design of the library was dictated by four factors: 1) the chemistry accessible from a key synthetic intermediate, 2) reactant availability, 3) reagent compatibility with other libraries of interest, and 4) clustering driven reagent selection. To begin the library design (**Figure S1**), we decided to use a single vendor, Enamine, which maintains a catalog of ∼210 million building blocks. To narrow the focus, the catalog was filtered to provide building blocks with ≥250 mg availability. These building blocks were then classified based on the molecular functionality present (e.g., amines, boronates, carboxylic acids, acid chlorides, alkyl halides, etc.), helping to identify available chemistries. Within these classifications, reagents with heavy atom counts ≥15 were removed to provide an inclusive list of reagents available for library generation. Because the targeted key intermediate (**2**, **Scheme 1**) used to synthesize the library congeners would be an aryl halide (or boronic acid/ester), we settled on Suzuki and Sonogashira compatible reactants and filtered the reagents to eliminate any non-compatible compositions. This narrowed the reagent classifications to alkynes, aryl boronic acids/esters, and aryl bromides. Remaining reagents were filtered to exclude structural alerts. At this point, protecting group compatibility was also considered for additional libraries we were targeting. Lastly, reagents were merged and clustered based on fingerprints comprised of AlogP and atom type, with clusters being defined as a group of compounds exhibiting a ≥0.6 Tanimoto score. Samples within each cluster were sorted by increasing size and the smallest reagent was generally selected to represent each cluster. If for any reason the top priority compound was unavailable or otherwise triaged, the next smallest reagent within a cluster was chosen to represent the cluster.

### Synthesis

Analogues **3a-p, 3eb,** and **3oa** were synthesized according to Scheme 1. Treatment of 4-bromo-3-fluoroaniline with benzyl chloroformate under mildly basic conditions afforded intermediate **1**, which was reacted with (*R*)-(-)-glycidyl butyrate to generate the corresponding oxazolidinone **2**. Analogues **3a-p** were obtained following a Suzuki cross-coupling between **2** and various aryl boronic acids/esters. Compound **3eb** was obtained by guanidinylation of compound **3e** followed by deprotection of the resulting carbamate to reveal the free guanidinium salt. Saponification of compound **3o** generated carboxylic acid **3oa**.

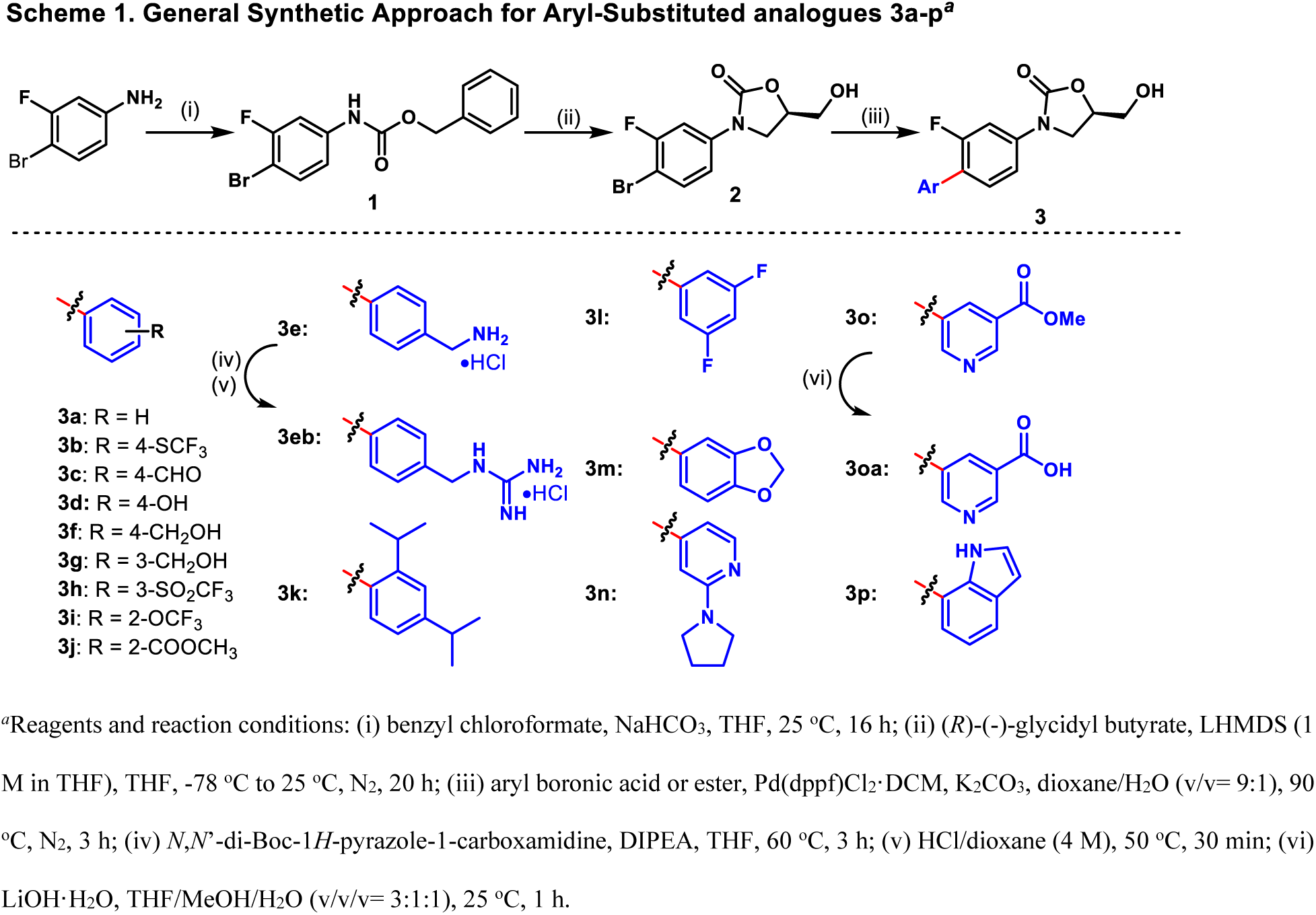

The general synthetic approach for analogues **6a-c** is illustrated in Scheme 2. Unlike the synthesis of **3a-p** the hydroxy group was protected prior to the Suzuki coupling for purification purposes. Briefly, the primary hydroxy group of oxazolidinone **2** was protected as the methoxymethyl ether to give **4**, which was coupled with select aryl boronic acids to provide the protected products **5a-c**. Deprotection of **5a-c** under typical acidic conditions yielded analogues **6a-c**.

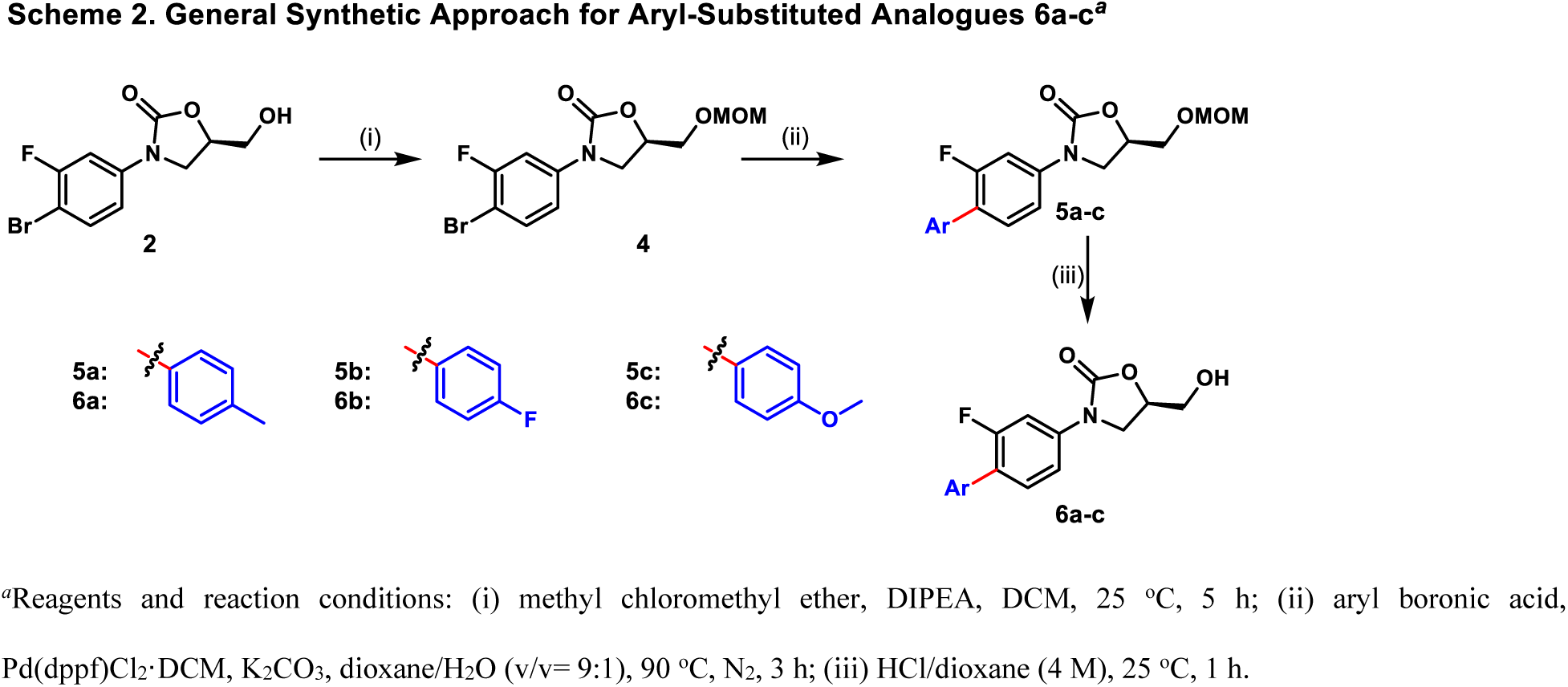

The general synthetic route for analogues **12a-p** and **14** is presented in Scheme 3. Similar to the formation of **2**, treatment of 3-fluoroaniline with benzyl chloroformate under mildly basic conditions followed by base-promoted cyclization employing (*R*)-(-)-glycidyl butyrate afforded oxazolidinone **8**. Regioselective iodination of **8** by treatment with *N-*iodosuccinimide provided intermediate **9,** which was subsequently transformed to boronic ester **10** through a palladium-catalyzed borylation reaction. Suzuki coupling between boronic ester **10** and select aryl bromides yielded compounds **12a-o**. Removal of the Boc protecting group on **12e** under acidic conditions yielded analogue **12ea**. Compound **12p** was generated through a modified Suzuki reaction from the iodo-oxazolidinone **9**. Additionally, the pendant hydroxyl group on **10** could be protected as the methoxymethyl ether to afford **11**, which upon exposure to typical Suzuki conditions, yielded the cyano pyridine **13**. Attempts to isolate the free amine **14** after nitrile reduction proved problematic. Thus, we resorted to a strategy of trapping the free amine as the *t*-butylcarbamate for purification purposes. Once isolated the resulting compound could be treated under acidic conditions to simultaneously cleave the carbamate, methoxymethyl ether, and *t*-butyl group to reveal compound **14.**

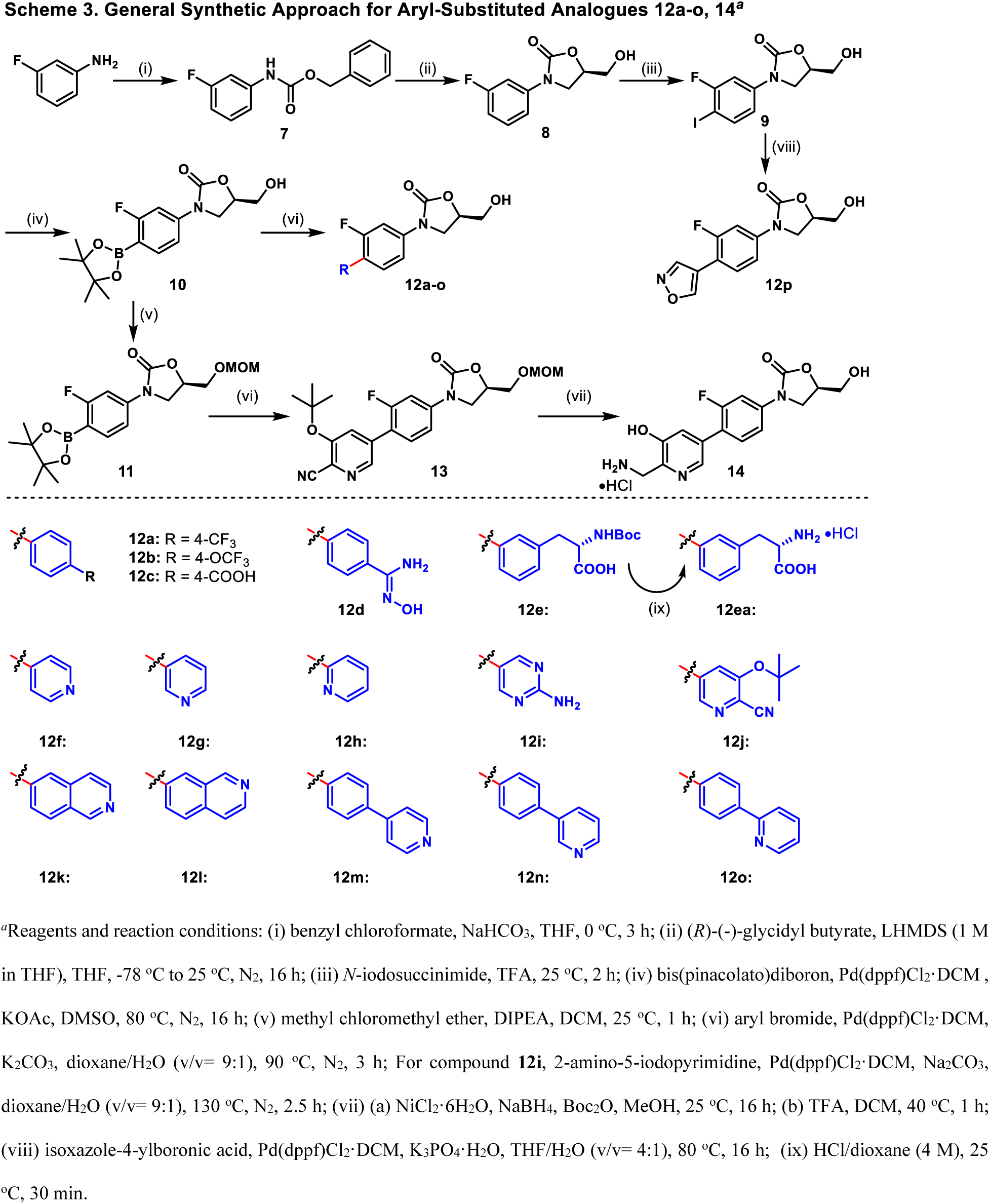

Alkynyl derivatives **15** were generated from **9** through a Sonogashira reaction with select coupling partners as depicted in Scheme 4. In many cases the resulting products were manipulated further to afford additional derivatives. For example, saponification of **15m** provided alcohol **15ma** and hydrochloride salts **15ra-va** were obtained after Boc removal from **15r**-**v** with trifluoracetic acid followed by salt exchange.

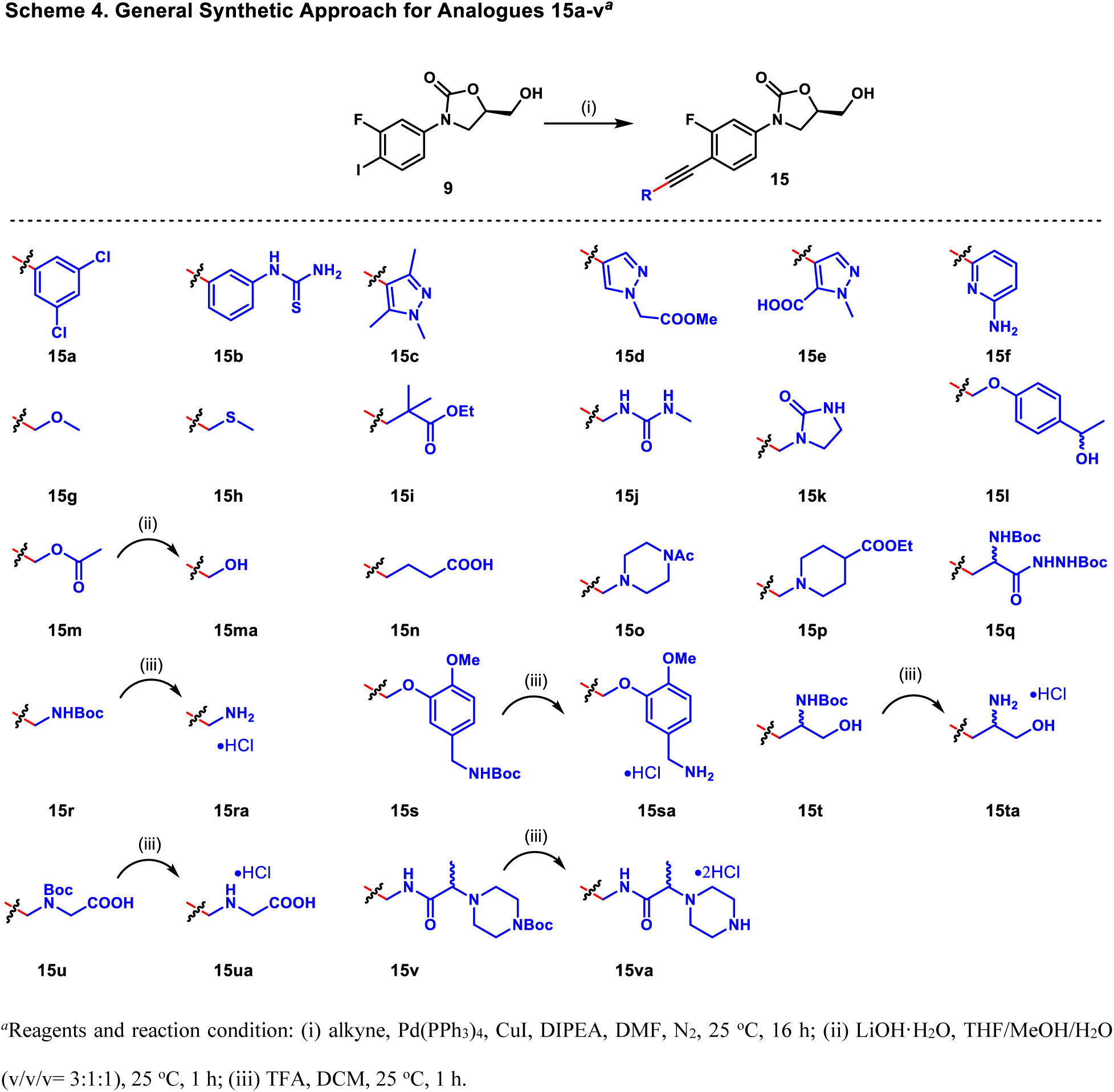

For the last cohort of analogs, the general synthetic route for pyridiniums **17a-l** is shown in Scheme 5. A Suzuki reaction between intermediate **11** and a corresponding heterocyclic bromide provided **16a-l**, which were then reacted with a haloalkane of choice to form the *N*-alkylated pyridiniums. Removal of the methoxy methyl ether afforded final compounds **17a-l**.

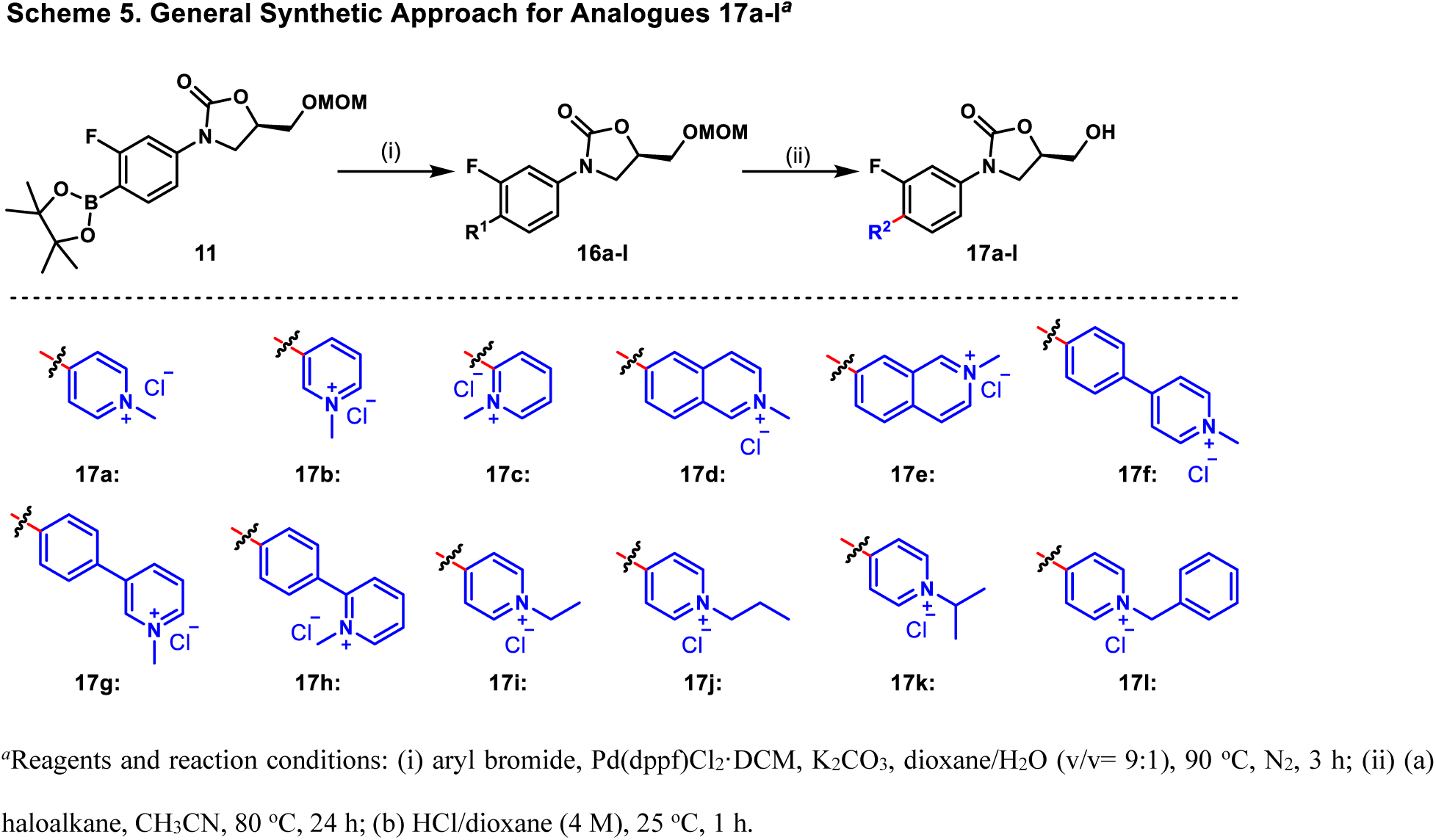

### Antibacterial activity of compounds

We next analyzed activities of the compound series in three GNB, *E. coli, P. aeruginosa* and *A. baumannii.* For all three species, wild type and isogenic mutants lacking efflux pumps (Δ) and/or producing a large recombinant OM pore (Pore) were used to separate the contributions of active efflux and the OM permeability barriers on the activities of compounds.^47^ The MIC (**Table S2**) and IC_50_ values (**Tables 1-4**) were determined first in the defined M9-MOPS medium, a better model (than nutrient rich conditions) for bacterial growth observed *in vivo* during certain infections.^48–50^ The dominant number of compounds had no activity in the wild type strains of GNB (**Tables 1-4** and **Table S2**). In contrast, at least half of the compounds inhibited the growth of the double-compromised (Δ+Pore) strains. With some exceptions, these compounds were active across species, including *S. aureus* (**Table 5**), suggesting that species-specific differences in binding to ribosomes are only minor factors in their activities.

**Table 1.** IC_50_ values of substituted phenyl C-ring derivatives in M9-MOPS medium. Arranged in order of potency against ECWT. All values reported in μM.

| Compound | AB<br>WT | AB<br>Pore | AB<br>$\Delta$ | AB<br>ΔPore | EC<br>WT | EC<br>Pore | EC<br>$\Delta$ | EC<br>ΔPore | PA<br>WT | PA<br>Pore | PA<br>$\Delta$ | PA<br>ΔPore |
| --- | --- | --- | --- | --- | --- | --- | --- | --- | --- | --- | --- | --- |
| 3e | 54.5 | 14.3 | 17.7 | 8.2 | 18.1 | 2.6 | 1.7 | 2.6 | 23.0 | 26.3 | 3.3 | 8.1 |
| 3a | 18.4 | 5.5 | 2.3 | 1.9 | 18.4 | 2.0 | 0.4 | 0.3 | >100 | 13.1 | 2.3 | 1.4 |
| 3m | >100 | >100 | 2.4 | 1.5 | 23.0 | 2.4 | 0.3 | 0.3 | >100 | >100 | 2.4 | 0.3 |
| 6c | >100 | >100 | 0.7 | 0.6 | 23.3 | 3.4 | 0.2 | 0.3 | >100 | >100 | 0.7 | 1.2 |
| 6a | >100 | >100 | 2.3 | 1.6 | 28.0 | 3.2 | 0.3 | 0.3 | >100 | >100 | 2.3 | 3.8 |
| 3b | 63.5 | 23.5 | 9.1 | 8.1 | 35.1 | 9.4 | 6.9 | 4.9 | >100 | >100 | 78.2 | 6.9 |
| 3i | >100 | 59.1 | 33.2 | 24.8 | 42.6 | 46.3 | 9.3 | 21.9 | >100 | >100 | 75.6 | 29.9 |
| 12d | 22.1 | 2.8 | 1.2 | 0.9 | 55.9 | 7.5 | 0.6 | 1.0 | >100 | 63.6 | 5.1 | 1.8 |
| 3c | 18.4 | 16.9 | 1.1 | 1.2 | 66.8 | 5.0 | 0.6 | 0.5 | >100 | 5.9 | 1.1 | 0.02 |
| 3g | 45.8 | 10.8 | 5.4 | 7.6 | 66.9 | 1.5 | 1.5 | 2.0 | >100 | >100 | 15.5 | 13.0 |
| 3f | 45.3 | 5.7 | 4.4 | 4.6 | 67.9 | 17.6 | 3.9 | 3.7 | >100 | >100 | 5.3 | 5.5 |
| 12a | >100 | 41.1 | 8.0 | 3.9 | 88.8 | 35.3 | 4.1 | 2.9 | >100 | >100 | 8.0 | 1.4 |
| 6b | >100 | 89.8 | 8.8 | 9.7 | 89.1 | 22.8 | 4.3 | 2.3 | >100 | 89.8 | 8.8 | 2.2 |
| 3j | >100 | >100 | >100 | >100 | 96.0 | 61.5 | 64.6 | 39.7 | >100 | >100 | >100 | >100 |
| 3k | >100 | 77.9 | 17.3 | 8.2 | 99.1 | 20.4 | 15.9 | 6.6 | >100 | >100 | 49.0 | 22.3 |
| 3d | >100 | 12.9 | 8.4 | 7.3 | >100 | 22.1 | 2.0 | 2.3 | >100 | 27.7 | 8.4 | 5.8 |
| 12c | >100 | 55.5 | 12.8 | 10.2 | >100 | >100 | 54.0 | 2.6 | >100 | >100 | >100 | 8.9 |
| 3h | >100 | >100 | 48.8 | 38.5 | >100 | 21.2 | 19.3 | 12.5 | >100 | >100 | >100 | >100 |
| 12b | >100 | 25.6 | 16.3 | 6.4 | >100 | 76.9 | 17.2 | 18.8 | >100 | >100 | 18.6 | 3.1 |
| 3eb | >100 | 65.9 | 67.8 | 32.5 | >100 | 56.1 | 63.8 | 55.7 | >100 | >100 | 12.0 | 25.7 |
| 3l | >100 | >100 | >100 | >100 | >100 | >100 | >100 | >100 | >100 | >100 | >100 | >100 |
| 12e | >100 | >100 | >100 | >100 | >100 | >100 | >100 | >100 | >100 | >100 | >100 | >100 |
| 12ea | >100 | >100 | >100 | >100 | >100 | >100 | >100 | >100 | >100 | >100 | >100 | >100 |
| Linezolid | 33.4 | 9.2 | 4.9 | 4.0 | >100 | 18.0 | 8.7 | 9.3 | >100 | 19.8 | 24.7 | 1.7 |

**Table 2.** IC_50_ values of substituted pyridyls C-ring derivatives in M9-MOPS medium. Arranged in order of potency against ECWT. All values reported in μM.

| Compound | AB<br>WT | AB<br>Pore | AB<br>$\Delta$ | AB<br>ΔPore | EC<br>WT | EC<br>Pore | EC<br>$\Delta$ | EC<br>ΔPore | PA<br>WT | PA<br>Pore | PA<br>$\Delta$ | PA<br>ΔPore |
| --- | --- | --- | --- | --- | --- | --- | --- | --- | --- | --- | --- | --- |
| 12f | 16.5 | 4.1 | 1.4 | 2.2 | 44.3 | 5.6 | 0.8 | 1.3 | 44.2 | 48.2 | 8.1 | 2.9 |
| 14 | 37.6 | 5.8 | 15.4 | 6.3 | 66.1 | 23.3 | 10.0 | 10.3 | 55.9 | 40.8 | 2.2 | 0.2 |
| 12o | 35.3 | 15.8 | 21.5 | 22.6 | 76.2 | 47.2 | 55.6 | 55.2 | >100 | >100 | >100 | >100 |
| 3p | >100 | >100 | 64.4 | 62.1 | 77.0 | 19.5 | 21.6 | 7.2 | >100 | >100 | >100 | >100 |
| 12i | >100 | >100 | 53.0 | 13.6 | >100 | 2.0 | 5.9 | 1.5 | >100 | >100 | >100 | 33.0 |
| 3n | 71.9 | 40.4 | 72.4 | 49.1 | >100 | 4.0 | 15.4 | 1.5 | >100 | >100 | >100 | >100 |
| 3o | 83.2 | 28.1 | 12.1 | 11.3 | >100 | 4.7 | 2.7 | 2.2 | >100 | >100 | 32.9 | 38.7 |
| 12g | 19.0 | 5.2 | 4.3 | 3.5 | >100 | 14.5 | 2.8 | 3.2 | >100 | 75.3 | 4.7 | 6.0 |
| **12p** | >100 | 39.0 | 83.4 | 73.1 | >100 | 56.1 | 23.6 | 30.6 | >100 | >100 | 22.7 | 32.9 |
| **3oa** | >100 | >100 | >100 | >100 | >100 | >100 | >100 | 42.2 | >100 | >100 | >100 | >100 |
| **12m** | >100 | >100 | >100 | 6.2 | >100 | >100 | >100 | 75.2 | >100 | >100 | 92.1 | 1.0 |
| **12h** | >100 | >100 | >100 | >100 | >100 | >100 | 76.8 | 79.6 | >100 | >100 | 64.2 | 47.4 |
| **12k** | >100 | 35.3 | 5.6 | 3.4 | >100 | >100 | >100 | >100 | >100 | >100 | 67.7 | 37.9 |
| **12l** | >100 | >100 | >100 | >100 | >100 | >100 | >100 | >100 | >100 | >100 | >100 | 0.1 |
| **12n** | >100 | 59.4 | >100 | 20.4 | >100 | >100 | >100 | >100 | >100 | >100 | >100 | >100 |
| **12j** | >100 | >100 | >100 | >100 | >100 | >100 | >100 | >100 | >100 | >100 | >100 | >100 |
| **Linezolid** | 33.4 | 9.2 | 4.9 | 4.0 | >100 | 18.0 | 8.7 | 9.3 | >100 | 19.8 | 24.7 | 1.7 |

**Table 3.** IC_50_ values of substituted pyridinium C-ring derivatives in M9-MOPS medium. Arranged in order of potency against ECWT. All values reported in μM.

| Compound | AB WT | AB Pore | AB $\Delta$ | AB $\Delta$ Pore | EC WT | EC Pore | EC $\Delta$ | EC $\Delta$ Pore | PA WT | PA Pore | PA $\Delta$ | PA $\Delta$ Pore |
| --- | --- | --- | --- | --- | --- | --- | --- | --- | --- | --- | --- | --- |
| **17a** | 59.4 | 13.8 | 23.3 | 13.8 | 13.5 | 12.3 | 10.0 | 11.0 | >100 | 59.7 | 48.8 | 9.4 |
| **17f** | >100 | 10.0 | 64.0 | 10.8 | 15.4 | 1.7 | 0.5 | 0.5 | >100 | >100 | >100 | >100 |
| **17i** | >100 | >100 | >100 | 38.5 | 43.8 | 36.0 | 13.7 | 15.9 | >100 | >100 | 57.4 | 42.3 |
| **17j** | >100 | 19.8 | 89.7 | 19.8 | 43.9 | 28.3 | 12.7 | 16.6 | >100 | >100 | >100 | >100 |
| **17l** | >100 | 15.6 | 23.4 | 13.0 | 51.6 | 23.3 | 15.5 | 15.6 | >100 | 41.7 | 4.6 | 1.1 |
| **17d** | >100 | 20.1 | 29.7 | 27.9 | 63.6 | 30.0 | 4.0 | 11.4 | >100 | >100 | >100 | >100 |
| **17h** | >100 | 13.3 | 20.4 | 10.2 | 72.6 | 13.2 | 3.1 | 4.4 | >100 | 15.9 | 8.4 | 1.2 |
| **17k** | 89.5 | 22.0 | 34.5 | 22.3 | 81.6 | 42.8 | 16.9 | 16.6 | >100 | >100 | 35.1 | 36.6 |
| **17g** | >100 | 13.9 | 29.7 | 12.6 | 98.4 | 7.8 | 1.2 | 2.1 | >100 | 28.1 | 29.7 | 18.1 |
| **17e** | >100 | 27.4 | 48.6 | 9.4 | >100 | 21.0 | 5.6 | 9.6 | >100 | >100 | >100 | >100 |
| **17c** | >100 | >100 | >100 | 33.3 | >100 | >100 | 72.3 | 60.4 | >100 | >100 | >100 | 40.4 |
| **17b** | >100 | >100 | >100 | >100 | >100 | >100 | 70.6 | 88.8 | >100 | >100 | >100 | >100 |
| **Linezolid** | 33.4 | 9.2 | 4.9 | 4.0 | >100 | 18.0 | 8.7 | 9.3 | >100 | 19.8 | 24.7 | 1.7 |

**Table 4.** IC_50_ values of substituted alkynyl derivatives in M9-MOPS medium. Arranged in order of potency against ECWT. All values reported in μM.

| Compound | AB WT | AB Pore | AB $\Delta$ | AB $\Delta$ Pore | EC WT | EC Pore | EC $\Delta$ | EC $\Delta$ Pore | PA WT | PA Pore | PA $\Delta$ | PA $\Delta$ Pore |
| --- | --- | --- | --- | --- | --- | --- | --- | --- | --- | --- | --- | --- |
| **15f** | >100 | 49.8 | 83.9 | 76.7 | 59.0 | 19.2 | 34.2 | 20.5 | >100 | >100 | >100 | >100 |
| **15l** | >100 | >100 | 61.9 | 34.7 | 62.4 | 24.7 | 30.2 | 13.5 | >100 | >100 | >100 | 91.3 |
| **15a** | >100 | >100 | >100 | >100 | 74.8 | 12.8 | 26.1 | 3.6 | >100 | >100 | 64.9 | 16.2 |
| **15g** | >100 | >100 | >100 | >100 | 74.9 | 36.3 | 80.3 | 44.0 | >100 | >100 | >100 | >100 |
| **15h** | >100 | >100 | >100 | >100 | 94.9 | 35.1 | 65.9 | 47.8 | >100 | >100 | >100 | >100 |
| **15c** | >100 | >100 | >100 | >100 | >100 | 1.2 | 86.3 | 3.4 | >100 | >100 | >100 | >100 |
| **15j** | >100 | >100 | >100 | >100 | >100 | >100 | >100 | 11.2 | >100 | >100 | >100 | >100 |
| **15i** | >100 | 60.3 | 44.0 | 30.3 | >100 | 65.3 | 17.8 | 21.9 | >100 | >100 | >100 | 30.1 |
| **15d** | >100 | >100 | >100 | 62.0 | >100 | 52.3 | 88.7 | 36.6 | >100 | >100 | >100 | >100 |
| **15k** | >100 | >100 | >100 | >100 | >100 | >100 | 64.3 | 39.0 | >100 | >100 | >100 | >100 |
| **15m** | >100 | >100 | >100 | >100 | >100 | 44.4 | 72.4 | 39.7 | >100 | >100 | >100 | >100 |
| **15ma** | >100 | >100 | >100 | >100 | >100 | >100 | 84.0 | 43.6 | >100 | >100 | >100 | >100 |
| **15s** | >100 | 42.4 | 72.2 | 28.7 | >100 | >100 | 42.9 | 44.4 | >100 | >100 | >100 | 46.9 |
| **15q** | >100 | >100 | >100 | >100 | >100 | 51.4 | 13.3 | 56.1 | >100 | >100 | >100 | >100 |
| **15v** | >100 | >100 | >100 | >100 | >100 | >100 | 50.9 | 62.6 | >100 | >100 | >100 | >100 |
| **15b** | >100 | >100 | >100 | >100 | >100 | >100 | 85.0 | 64.5 | >100 | >100 | >100 | >100 |
| **15r** | >100 | >100 | >100 | >100 | >100 | 74.3 | 83.4 | 64.7 | >100 | >100 | >100 | >100 |
| **15t** | >100 | >100 | >100 | >100 | >100 | 97.3 | 72.6 | 67.8 | >100 | >100 | >100 | >100 |
| **15e** | >100 | 61.3 | 88.2 | 89.5 | >100 | >100 | >100 | >100 | >100 | >100 | >100 | >100 |
| **15o** | >100 | >100 | >100 | >100 | >100 | >100 | >100 | >100 | >100 | >100 | >100 | >100 |
| **15p** | >100 | >100 | >100 | >100 | >100 | >100 | >100 | >100 | >100 | >100 | >100 | >100 |
| **15n** | >100 | >100 | >100 | >100 | >100 | >100 | >100 | >100 | >100 | >100 | >100 | >100 |
| **15va** | >100 | >100 | >100 | >100 | >100 | >100 | >100 | >100 | >100 | >100 | >100 | >100 |
| **15ta** | >100 | >100 | >100 | >100 | >100 | >100 | >100 | >100 | >100 | >100 | >100 | >100 |
| **15u** | >100 | >100 | >100 | >100 | >100 | >100 | >100 | >100 | >100 | >100 | >100 | >100 |
| **15ua** | >100 | >100 | >100 | >100 | >100 | >100 | >100 | >100 | >100 | >100 | >100 | >100 |
| **15ra** | >100 | >100 | >100 | >100 | >100 | >100 | >100 | >100 | >100 | >100 | >100 | >100 |
| **15sa** | >100 | >100 | >100 | >100 | >100 | >100 | >100 | >100 | >100 | >100 | >100 | >100 |
| **Linezolid** | 33.4 | 9.2 | 4.9 | 4.0 | >100 | 18.0 | 8.7 | 9.3 | >100 | 19.8 | 24.7 | 1.7 |

**Table 5.** MICs of select oxazolidinone analogs in LB medium. All values reported in μM.

| Compound | AB<br>$\Delta\text{Pore}$ | EC<br>$\Delta\text{Pore}$ | PA<br>$\Delta\text{Pore}$ | S. aureus<br>ATCC<br>25923 | SQ110 | SQ110<br>DTC | SQ110<br>DTC<br>G2032A | SQ110<br>DTC<br>C2610A |
| --- | --- | --- | --- | --- | --- | --- | --- | --- |
| **14** | 25 | 50 | 6.25 | 12.5 | >100 | 25 | 100 | 50 |
| **12a** | 6.25 | >100 | 100 | 3.12 | >100 | 25 | >100 | >100 |
| **12b** | 25 | >100 | 100 | 6.25 | >100 | 100 | >100 | >100 |
| **12c** | 50 | 12.5 | >100 | 6.25 | >100 | 25 | 50 | 25 |
| **12f** | 6.25 | 6.25 | 3.125 | 1.56 | >100 | 6.25 | 25 | 6.25 |
| **12g** | 12.5 | 25 | 6.25 | 1.56 | >100 | 12.5 | 50 | 25 |
| **12i** | 50 | 100 | 12.5 | 6.25 | >100 | 25 | 100 | 50 |
| **17a** | 100 | 100 | 100 | >100 | 100 | 50 | >100 | 100 |
| **17f** | 50 | 12.5 | 12.5 | 12.5 | >100 | 3.12 | 25 | 12.5 |
| **17g** | 25 | 25 | 25 | 12.5 | >100 | 12.5 | 50 | 25 |
| **3e** | 25 | 100 | 12.5 | 6.25 | >100 | 50 | 100 | 50 |
| **3f** | 12.5 | 25 | 6.25 | 1.56 | >100 | 12.5 | 50 | 25 |
| Linezolid | 25 | 25 | 12.5 | 6.25 | >100 | 25 | >100 | 50 |

Three IC_50_ ratios: WT/ΔEfflux(Pore) (total barrier ratio, W/PE), WT(Pore)/ΔEfflux(Pore) (efflux impact ratio, P/PE), ΔEfflux/ ΔEfflux(Pore) (OM impact ratio, E/PE) were analyzed by principal component analysis to provide insight into the influence of each barrier (i.e., efflux and porination) on activity for each species. For this PCA, only the 26 compounds that showed a detectable IC_50_ in at least two barrier-variant strains across all three bacteria were used (**Figure 2**). The rest of the compounds had to be excluded from the comparative analysis because the effect of neither efflux nor porination could be quantified in at least one of the species. In most cases for all three bacteria, the effect of efflux inactivation (P/PE ratio) dominated that of porination (E/PE ratio). Thus, any compound specificity observed for each species is primarily defined by the organism’s repertoire of efflux pumps. We further noted that *A. baumannii* was much closer to *E. coli* than *P. aeruginosa*, according to PCA. For both efflux and porination, the first principal component (PC1) was defined by the difference between *P. aeruginosa* and the other two bacteria, whereas the second principal component (PC2) was aligned with the distinction between *E. coli* and *A. baumannii*. Inspection of PCs (**Table S1**) revealed that the PC1 and PC2 in the efflux impact ratio (P/PE) were dominated, respectively, by **14** and **17l**, and by **6a, 6c** and **3m**. Thus, these compounds are the most reflective of the species-specific difference in the efficiency of their permeation barriers. In particular, **14** and **17l** were both identified as efflux liabilities for *P. aeruginosa* but not for *E. coli* or *A. baumannii* (**Figure 4**). For the OM impact most of the difference could be attributed to compounds **3c**, **14** and **LZD** (PC1) and **3l**, **3c, 3e** and **17g** (PC2). As seen in **Figure 3**, permeabilities of compounds **3c, 14**, and **LZD** were greatly affected by the presence of the pore for *P. aeruginosa* but not for *E. coli* or *A. baumannii.* Thus, the distinction of *P. aeruginosa* from the two other bacteria is achieved both through the more efficient efflux and lower transmembrane diffusion. However, the two factors are differentially affected by the compound structures.

**Figure 2.**
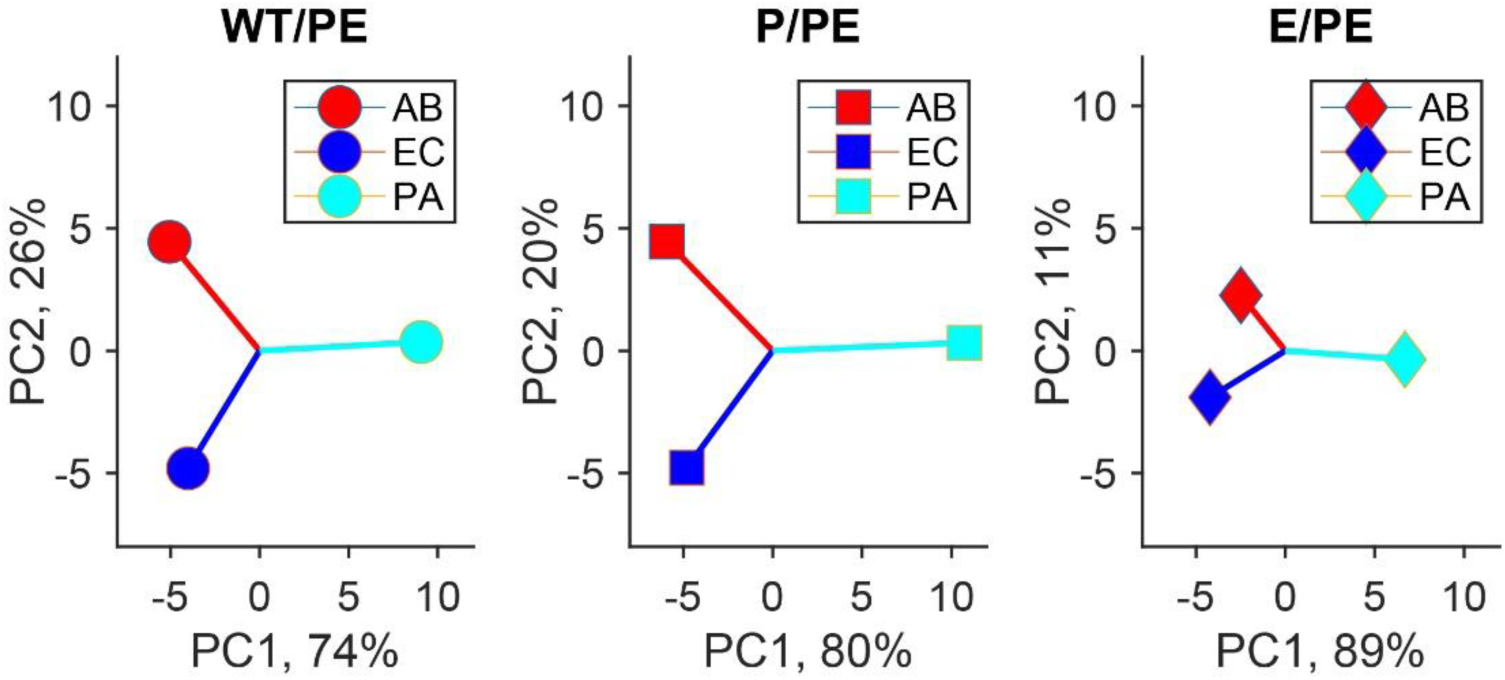
Principal component analysis of the three efflux ratios in *A. baumannii* (AB), *E. coli* (EC) and *P. aeruginosa* (PA).

**Figure 3.**
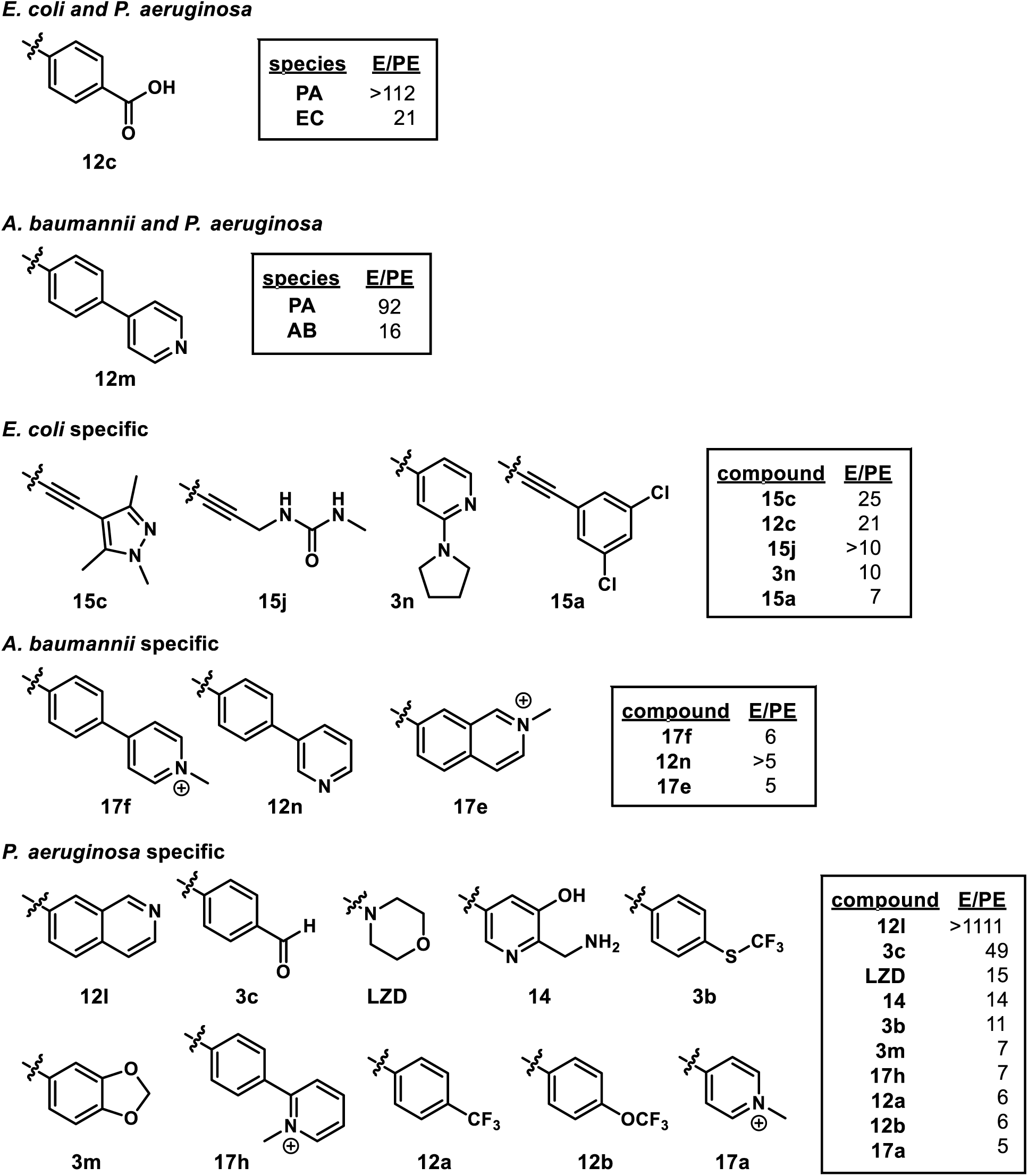
Motifs identified as liabilities to OM permeation. E/PE = Δ/ΔPore IC_50_ ratio for the indicated species.

**Figure 4.**
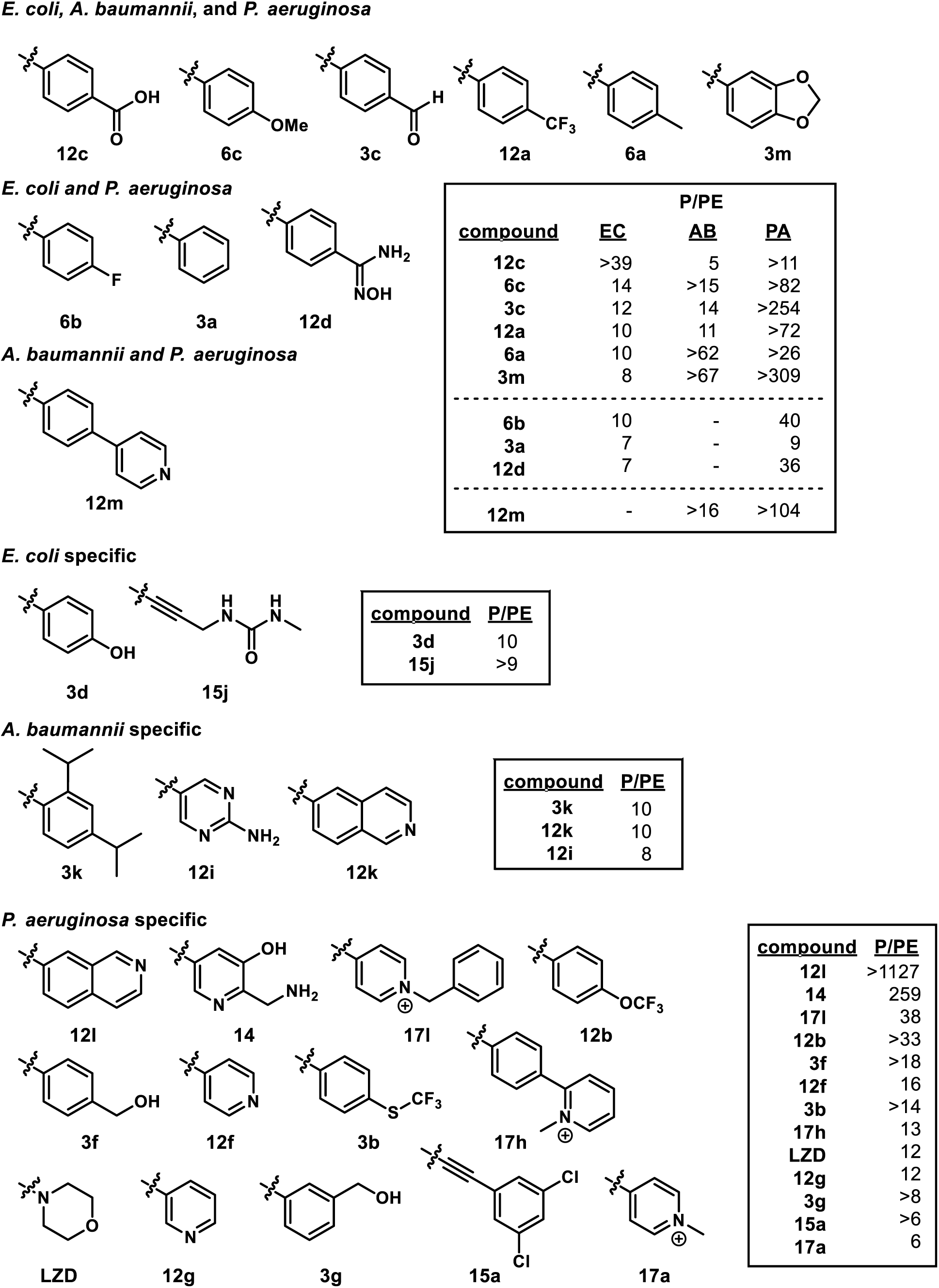
Motifs identified as liabilities to efflux susceptibility. P/PE = Pore/ΔPore IC_50_ ratio for the indicated species.

### Structure-activity relationships

*E. coli* is the best characterized and we will first focus on SAR in this species and then identify species-specific differences for *A. baumannii* and *P. aeruginosa*.

#### Phenyl Series

Among groups **3**, **6**, and **12** with substituted phenyl variants, 15/23 analogs exhibited activity against ECWT (**Table 1**) at the concentrations tested (up to 100 μM). Within this series, 15/23 derivatives exhibited near equal or better activity than LZD against the double compromised ECΔPore strain. Compounds **3j**, **3eb**, **3l**, **12e**, and **12ea** exhibited >5-fold less activity than LZD in the ECΔPore strain, suggesting that the motifs associated with these molecules are likely detrimental to target engagement. Inspection of the corresponding motifs reveal no obvious functionality or trend that may explain target engagement issues. Compounds **3d** and **12c** are interesting in that they exhibited improved activity against the ECΔPore strain but were inactive against ECWT. Efflux susceptibility seems to be the main issue for **3d** whereas E/PE and P/PE IC_50_ ratios for **12c** reveal issues with both OM permeation and efflux susceptibility.

Activity of this series against ABWT decreased to include only 7/23 compounds with **3i**, **3j**, **3k**, **3m**, **6a**-**6c**, and **12a** dropping out from the series that was active against ECWT (**Table 1**). Six compounds (**3a-3c**, **3f**, **3g**, and **12d**) exhibited activity against ECWT and ABWT. Contrary to activity against ECΔPore, only 7/23 analogs exhibited equal or better activity than LZD against the ABΔPore strain. In addition to the analogs identified as problematic for target engagement against *E. coli*, **3h**, and **3i** also prove detrimental in *A. baumannii.* The list of compounds that exhibited activity against the ΔPore strain but not against the WT grew for *A. baumannii* to include **3d**, **3k**, **3m**, **6a-6c**, and **12a**-**12c**. This suggests that the OM and efflux proficiency of *A. baumannii* supersedes that of *E. coli* for this chemotype.

Against *P. aeruginosa*, **3e** was the only compound that exhibited activity against the WT strain (**Table 1**). Within this series 5/23 compounds performed better than LZD against the PAΔPore strain, fewer than both *E. coli* and *A. baumannii.* In addition to the motifs identified that decreased target engagement efficiency against *E. coli* and *A. baumannii*, **3g**, **3k**, and **12c** are included to the list of compounds exhibiting >5-fold decreased activity than LZD against the ΔPore strain. Compound **3c** is interesting to note, as it provided very good potency (20 nM) against PAΔPore, perhaps suggesting additional mechanisms of action. This is a reasonable hypothesis given the presence of a reactive aldehyde.

In summary, from this series, only **3e** exhibit pan activity against *E. coli*, *A. baumanni*, and *P. aeruginosa* WT. No members of this subset that were inactive against ECWT exhibited activity against ABWT or PAWT. SAR trends observed in this series clearly indicate that barrier stringency exhibited between these species can be summarized as *P. aeruginosa* > *A. baumannii* > *E. coli*. Likewise, although the 23S rRNA target of the oxazolidinones is known to be highly conserved, the observation that the list of motifs proving detrimental to target engagement increases from *E. coli* to *A. baumannii* to *P. aeruginosa,* highlights that discernable differences in the target exist and can potentially be exploited for narrowing or broadening activity.

#### Pyridyl Series

Within this series, only 4/16 derivatives (**12f**, **14**, **12o**, and **3p**) were active against ECWT and 6/16 exhibited near equal or better activity than LZD against the ECΔPore strain (**Table 2**). The list of compounds that exhibited >5-fold worse activity against ECΔPore than LZD includes **12h**, **12j**, **12k**, and **12l**-**12o**). It is worth noting that within this group that exhibits less efficient target engagement, motifs include extended pyridyls, comprising both rotatable (**12m-12o**) and fused (**12k** and **12l**) extensions. All analogs of this series that were nearly equal or more potent than LZD against ECΔPore exhibited worse activity against ECWT. Thus, while pyridyls may improve target engagement, they seem to increase OM permeation and efflux liabilities.

Against *A. baumannii*, 6/16 compounds were active against the WT strain with five exhibiting equal or better activity than LZD. However, only 3/16 compounds exhibited near equal or better activity than LZD against ABΔPore. Contrary to *E. coli*, compounds **12m** and **12k** exhibited good activity against ABΔPore. Compound **12n** also exhibited activity against ABΔPore, whereas it was inactive against ECΔPore. These three analogs once again highlight the discernable differences in target engagement between the species. Against *A. baumannii*, in addition to **12h**, **12j**, **12n** and **12o** identified in *E. coli*, derivatives **3n**, **3oa**, **3p**, and **12p** proved to be detrimental to on-target activity.

This series produced two compounds active against PAWT (**12f** and **14**) and three that exhibited better activity than LZD in PAΔPore (**14**, **12m**, and **12l**). Compound **12l** is especially interesting, as it is inactive against ECΔPore and ABΔPore. This suggests either a secondary mechanism of action in *P. aeruginosa* or a special feature of the 23S rRNA binding pocket that can be selectively exploited. LZD exhibits activity against Pore and Δ6 strains, whereas most pyridyl analogs are inactive. This suggests that pyridyls succumb to additive/synergistic properties of OM permeation and efflux.

In summary, this series produced two analogs with pan WT activity (**12f** and **14**) and an additional compound (**12o**) that exhibited activity against ECWT and ABWT. Contrary to the substituted phenyl series, ECWT activity is not a preliminary qualifier for ABWT activity, as three compounds (**3n**, **3o**, and **12g**) exhibited activity against ABWT but are inactive against ECWT. For this series the barrier stringency seems to be summarized as *P. aeruginosa* > *E. coli* > *A. baumannii*, thus highlighting that motif properties can influence species response.

#### Pyridinium Series

Compound series **17** was comprised of various pyridinium motifs. From this series 9/12 analogs were active against ECWT (**Table 3**). Only two derivatives (**17c** and **17b**) were >5-fold less active than LZD against ECΔPore, indicating good complementarity of a pyridinium functionality with the 23S rRNA target. Comparison of the IC_50_ values across all four strains, reveals this series to be less of an issue for OM permeation and efflux susceptibility. This is consistent with previous observations that positively charged motifs may improve accumulation of small molecules, due to porin uptake mechanisms.

Against *A. baumannii*, only two compounds were active against ABWT (**17a** and **17k**) but both were less active than LZD. In addition to **17b** and **17c, 17d**, **17i**, and **17k** exhibited >5-fold lower activity than LZD against ABΔPore. This suggests that the pyridinium motif may not be as complementary to target binding site as it is for *E. coli*. In fact, most analogs in this series are less active than LZD against any of the *A. baumannii* strains.

For *P. aeruginosa*, no analogs within this series were active against WT. In addition to motifs identified for being detrimental to target engagement in *E. coli* and *A. baumannii*, all compounds except **17l** and **17h** can be added to this list for *P. aeruginosa*. This further highlights the decrease in susceptibility of *P. aeruginosa* to modification of the oxazolidinone scaffold for 23S rRNA inhibition.

In summary, no pyridinium analogs were identified that exhibited broad spectrum activity against all three species. While this motif class seems rather beneficial for improving WT activity against *E. coli*, it is detrimental to whole cell activity against *A. baumanni* and *P. aeruginosa*. The trend of motif SAR narrowing from *E. coli* to *A. baumannii* to *P. aeruginosa*, with *P. aeruginosa*, being more stringent for allowable functionality was consistent for this chemotype.

#### Alkynyl Series

This series was the least active with only 5/28 exhibiting activity against ECWT and no compounds being active against ABWT of PAWT (**Table 4**). In fact, although 64% of the compounds in this series exhibited activity against ECΔPore, albeit with lower activity than LZD, only 21% and 14% were active against ABΔPore and PAΔPore, respectively. These results further suggest that the positioning and functionality at this location on the oxazolidinone chemotype is more important for target engagement and the antibacterial activity than typically appreciated. The alkynyl attachment of this series likely extends beyond the binding pocket and is counter-productive to binding and by extension, activity. These results also confirm once against a trend of *E. coli* > *A. > baumannii* > *P. aeruginosa* regarding compatibility with broadened motif functionality.

#### Additional Trends from Cross-Series Comparisons

While interrogation of each subset independently provided insight into each series, cross-series comparisons also provided interesting insight. For example, comparison of IC_50_ values of **3a**, **12f-12h**, and **17a-17c**, elucidates the effect of pyridine inclusion, location, and conversion into a pyridinium on activity (**Tables 1-3**). This series demonstrates that activity against WT and ΔPore strains is relatively maintained against *E. coli* and *A. baumannii* after incorporation of a pyridine with the nitrogen at the *para*-position (**3a** compared to **12f**) and activity against PAWT is gained. Interestingly, if the position of the nitrogen is moved to the 3-position (**12g**) activity against ECWT and PAWT is lost, with both barriers now contributing to loss of activity in *E. coli* and primarily efflux susceptibility becoming the main issue in *P. aeruginosa* (P/PE >> E/PE). Conversion of the pyridine to the pyridinium, however, only improves WT activity against *E. coli*, further illustrating the potential limitation of the benefit that charge incorporation provides with general GNB activity. In fact, inherent charge incorporation was detrimental to ABWT and PAWT activity in every instance in this study.

A second comparison of **3a**, **12f**, **17a**, **17i**-**17l**, IC_50_ values highlights the effect of alkyl size appended to the nitrogen in the pyridinium groups. As mentioned previously, incorporation of a charged pyridinium results in a slight loss in ECΔPore activity but increasing the size of the alkyl group does not continue to decrease activity. However, the activity against WT seems to correlate negatively with size, as groups larger than a methyl demonstrated decreased ECWT activity. Against *A. baumanni* and *P. aeruginosa* smaller or larger substituents exhibited more activity against the ΔPore strains, however, WT activity decreased in both these species following incorporation of a pyridinium.

### Target-based resistance reduces potencies of compounds

The above results show that the generated compounds vary broadly in their potencies against GNB. We next compared the MICs of select compounds against “barrier-less” strains of *E. coli, A. baumannii* and *P. aeruginosa* to those against *S. aureus,* all grown in the nutrient rich LB medium. Our results show consistent differences in MICs between the Gram-negative strains as well as in comparison to *S. aureus* (**Table 5**), albeit compounds seem to be more potent against cells grown in the M9-MOPS medium. In general, *S. aureus* was at least 2-4-fold more susceptible to growth inhibition induced by tested compounds, including the antibiotic linezolid (**Table 5**). The most potent across the species were substituted pyridyls **12f** and **12g** and the substituted benzyl derivative **3f.** Furthermore, these compounds were at least 4-fold more potent than linezolid against *S. aureus*.

We also determined MICs against the *E. coli* SQ110 and SQDTC (ΔTolC) strains carrying only one copy of the rRNA operon and against SQDTC derivatives producing the linezolid-resistant 23S rRNA variants with G2032A or C2610A substitutions.^51^ We found that the SQ110 strain was resistant to all tested compounds (**Table 5**). With a few exceptions, SQ110DTC was more susceptible to compounds than ECΔPore strain, suggesting that 23S rRNA copy number contributes to differences in MICs. However, for all compounds MICs were higher in linezolid resistant strains. Thus, the antibacterial activities of compounds remain specific to 23S rRNA.

### Structure-uptake relationships

We next identified problematic motifs associated with general or species-specific efflux susceptibility and poor OM permeation. All compounds exhibiting measurable IC_50_ values against at least one double-compromised strain were included in the analysis. Ratios for P/PE and E/PE were calculated for each species and strain. Any IC_50_ value >100 for a Δefflux or (Pore) strain was set to 100 to allow P/PE and E/PE ratios for each motif to be calculated. This approach ensures that, if anything, the ratios for compounds with >100 IC_50_ values are underestimated. Ratios exceeding an arbitrary threshold of ≥5 were classified as a motif exhibiting problematic behavior for the corresponding barrier. For example, an E/PE ratio of 10 for a given analog in *E. coli* would identify the corresponding motif as problematic to *E. coli* OM permeation. This ratio analysis was completed for each analog over each of the three species. Motifs associated with poor OM permeation and/or increased efflux susceptibility are shown in **Figure 3** and **Figure 4**, respectively. Interestingly, no motifs were identified that exhibited poor OM permeation across all three species and only a single motif (**3c**) increased efflux susceptibility across all three species. In accordance with previous studies that indicate *P. aeruginosa* to exhibit higher levels of OM and efflux barrier synergy, nearly 50% of the motifs identified as increasing the susceptibility to efflux in this species were also exposed as being problematic for OM permeation (**14**, **17a**, **17h**, **LZD**).

## CONCLUSION

In conclusion, we report the design, synthesis, and evaluation of a library of oxazolidinones against *E. coli*, *P. aeruginosa*, and *A. baumannii*, and their isogenic strains with varying degrees of outer membrane and/or efflux pump deficiencies. The data indicate that evaluated permeation mechanisms do not account for all activity differences between organisms. This could suggest 1) secondary mechanisms of action or different off-target profiles for oxazolidinones in some species;2) tangible differences in target engagement and/or 3) involvement of additional permeation mechanisms such as alternative efflux pumps. While we do observe growth media specific differences in activity, this was expected as growth and metabolic rates of bacteria differ in minimal versus nutrient rich environments. It is important to note however, that the overall trends of activity against certain species were not changed, suggesting that the growth medium does not contribute significantly to species-specific inhibitory differences.

The data clearly indicate that motif manipulation on a chemotype is enough to significantly modulate efflux and permeation. As such, the perception that the efflux and OM permeation issues are a characteristic of the entire chemotype class does not hold true for the oxazolidinone scaffold. Rather the orientation, location, and composition of functional groups are the determinants on chemotype accumulation. In general, *P. aeruginosa* is more divergent from the other two bacteria and the least responsive to oxazolidinone treatment. It would be more challenging to engineer activity against *P. aeruginosa* for this chemotype, as efflux and OM barriers synergize strongly in this organism. *A. baumannii* and *E. coli* are more similar with efflux being the major issue for this chemotype, but certain motifs also face difficulties with OM permeation.

Determination of IC_50_ values revealed three analogs (**3e**, **12f**, and **14**) with a broadened spectrum of activity to include WT *E. coli*, *A. baumanni*, and *P. aeruginosa*. While these three analogs were previously reported to exhibit activity against another ECWT strain (MG1655),^45^ this is the first report of their broadened activity to include two other ESKAPE pathogens *A. baumannii* and *P. aeruginosa*. Of the >50 compounds presented in this study; no single motif was identified that correlated with poor OM permeation across all three GNB species. On the contrary, 6 different motifs were identified that increased efflux susceptibility across all three species (**12c**, **6c**, **3c**, **12a**, **6a**, and **3m**). These observations provide compelling evidence that molecular features that dictate compound accumulation are likely to vary from one species to the next, with OM permeation being the most species specific and efflux susceptibility exhibiting more overlap.

While generating overarching Lipinksi-like rules or guidelines for compound accumulation in Gram-negative bacteria is an enticing goal, we postulate that a more feasible first step may be to identify motifs for several chemotypes that significantly affect accumulation (positively or negatively). Once this has been completed for a variety of chemotypes and across several bacterial species, a clearer picture of general versus chemotype- and species-specific trends is expected to emerge. One can than imagine eventually advancing to a point wherein hierarchical ranking of motifs can be organized to provide a Topliss-like trees^52^ and/or bioisostere-like classifications^53^ to guide accumulation improvement efforts and rational design for Gram-negative active molecules.

## EXPERIMENTAL SECTION

### Bacterial strains and growth conditions

The construction of strains and their properties were described previously.^54–55^ Bacterial strains (**Table 6**) were grown either Luria-Bertani broth (10 g of Bacto tryptone, 5 g of yeast extract, and 5 g of NaCl per liter; pH 7.0), LB agar (LB broth with 15 g of agar per liter) or a minimal M9 medium (1X) supplemented with 50 mM MOPS buffer (pH 7.2) were used for bacterial growth. The M9-MOPS medium was optimized to support the growth and expression of the OM pore in strains of all three species. We previously found that in *E. coli* and *A. baumannii* cells, the expression of the pore was the most consistent under an arabinose-inducible promoter, but in *P. aeruginosa* the most consistent expression was achieved under an IPTG-inducible promoter.^55^ Since glucose acts as a repressor for an arabinose-inducible promoter, this sugar could not be used as a carbon source in these experiments. Therefore, M9-MOPS medium was supplemented with 0.2% xylose (*E. coli, A. baumannii* and *P. aeruginosa*) plus 0.5% sodium citrate (*P. aeruginosa* and *A. baumannii* only) as carbon sources.

**Table 6.** Bacterial strains used in this study.^54–55^

| Name | Strain | Relevant Genotype |
| --- | --- | --- |
| *P. aeruginosa* PAO1 |  |  |
| PA WT | GKCW111 | PAO1 *attTn7::mini-Tn7T-Gm-lacI<sup>q</sup>-pLAC-MCS* |
| PA Pore | GKCW119 | PAO1 *attTn7::mini-Tn7T- Gm<sup>r</sup>-lacI<sup>q</sup>-pLAC- fhuAΔCΔ4L* |
| PA Δ | PAO1116 MCS | PAO1 *ΔmexAB-oprM ΔmexCD-oprJ ΔmexXY ΔmexEF-OprN ΔmexJKL ΔtriABC attTn7::mini-Tn7T- Gm<sup>r</sup>-lacI<sup>q</sup>-pLAC-MCS* |
| PA ΔPore | GKCW120 | PAO1 *ΔmexAB-oprM ΔmexCD-oprJ ΔmexXY ΔmexEF-OprN ΔmexJKL ΔtriABC attTn7::mini-Tn7T- Gm<sup>r</sup>-lacI<sup>q</sup>-pLAC- fhuAΔCΔ4L* |
| *A. baumannii* ATCC17978 |  |  |
| AB WT | JWW19 | ATCC 17978 *attTn7::miniTn7-Tp<sup>r</sup>-araC-P<sub>araBAD</sub>-MCS (pTJ1)* |
| AB Pore | JWW20 | ATCC 17978 *attTn7::miniTn7-Tp<sup>r</sup>-araC-P<sub>araBAD</sub> -fhuA (FhuA<sub>ΔCork</sub>, Δ4Loop, 6His) (pTJ1-FhuA)* |
| AB Δ | IL119 | ATCC 17978 *ΔadeIJK::FRT ΔadeAB::FRT ΔadeFGH::FRT attTn7::miniTn7-Tp<sup>r</sup>-araC-P<sub>araBAD</sub>-MCS (pTJ1)* |
| AB ΔPore | IL123 | ATCC 17978 *ΔadeIJK::FRT ΔadeAB::FRT ΔadeFGH::FRT attTn7::miniTn7-Tp<sup>r</sup>-araC-P<sub>araBAD</sub>-fhuA(FhuA<sub>ΔCorkΔ4Loop, 6His</sub>) (pTJ1-FhuA)* |
| *E. coli* BW25113 |  |  |
| EC WT | GKCW102 | BW25113 *attTn7::mini-Tn7T -Kan<sup>r</sup> -araC-P<sub>araBAD</sub>-MCS* |
| EC Pore | GKCW101 | BW25113 *attTn7::mini-Tn7T-Kan<sup>r</sup> -araC-P<sub>araBAD</sub>-fhuAΔCΔ4L* |
| EC Δ | GKCW104 | BW25113 *ΔtolC-ygiBC attTn7::mini-Tn7T -Kan<sup>r</sup> -araC-P<sub>araBAD</sub>-MCS* |
| EC ΔPore | GKCW103 | BW25113 *ΔtolC-ygiBC attTn7::mini-Tn7T-Kan<sup>r</sup> -araC-P<sub>araBAD</sub>-fhuAΔCΔ4L* |

### Minimum Inhibitory Concentrations

Compounds were dissolved in DMSO (10 mM). The two-fold serial dilution method in 96-well plates was used to determine MICs. *E. coli*, *A. baumannii* and *P. aeruginosa* cultures grown in indicated media were induced with 0.1%, 1% arabinose or 0.1mM IPTG, respectively. The MIC was noted visually after 24 hrs incubation at 37°C and OD_600_ was measured for all 96-well plates followed determination of IC_50_ values as described previously.^56^

### Chemistry

Starting materials, ACS grade methylene chloride (DCM), methanol (MeOH), hexanes, ethyl acetate (EtOAc), acetone, acetonitrile (CH_3_CN), dimethyl formamide (DMF), anhydrous dimethyl sulfoxide (DMSO), dioxane, ethanol, formic acid, anhydrous tetrahydrofuran (THF), tetrahydrofuran (THF), toluene, and trifluoroacetic acid (TFA) were purchased from TCI Chemicals, Oakwood, Alfa Aesar, Fisher Scientific, Enamine, or Sigma-Aldrich. Deionized water was used for all experimental procedures where “water” is indicated. All reactions requiring anhydrous conditions were run under a nitrogen atmosphere. Analytical and preparative thin-layer chromatography (TLC) was performed on silica gel 60 F_254_ plates (Sigma-Aldrich 1.05715). Flash column chromatography was carried out on silica gel (70-230 mesh, SiliCycle). NMR were collected on a 400, 500, and 600 MHz (specified below) Varian VNMRS Direct Drive spectrometer equipped with an indirect detection probe. NMR data were collected at 25 °C unless otherwise indicated. Pulse sequences were used as supplied by Varian VNMRJ 4.2 software. All NMR data were processed in MestReNova. High-resolution mass spectrometry was obtained from and analyzed by the Mass Spectrometry Facility at the University of Minnesota. All compounds evaluated in biological assays were >95% purity based on HPLC and/or NMR.

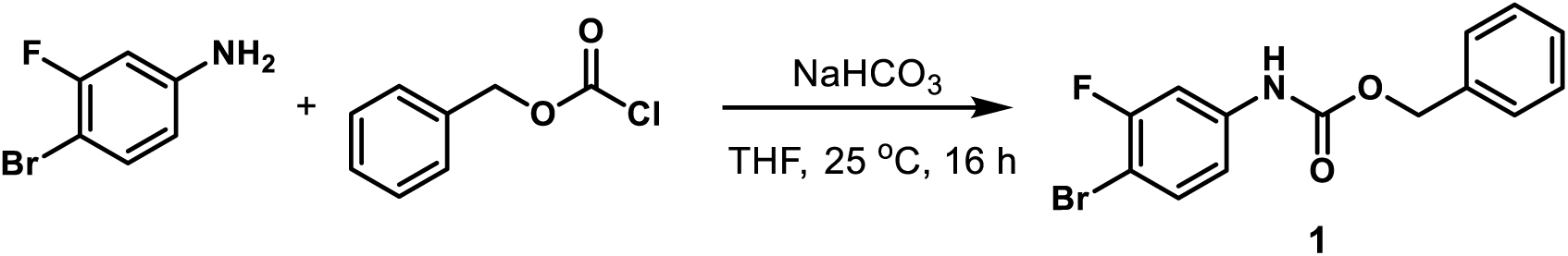

*Benzyl (4-Bromo-3-fluorophenyl)carbamate (**1**).* To a solution of 4-bromo-3-fluoroaniline (3.8 g, 20 mmol) in THF (80 mL) was added sodium bicarbonate (3.36 g, 40 mmol) and the mixture was cooled to 0 °C. Benzyl chloroformate (4.23 mL, 30 mmol) was added dropwise with a syringe. The reaction mixture was stirred at 25 °C. After the reaction was judged to be completed by TLC (16 h), it was quenched with water and extracted with EtOAc three times. The combined organic layers were washed with water and concentrated under reduced pressure by rotary evaporation. The crude residue was purified by flash column chromatography (SiO_2_, eluent gradient 0-20% EtOAc in hexanes) to give compound **1** as a yellow amorphous solid (5.90 g, 91%). ^1^H NMR (300 MHz, Chloroform-*d*) δ 7.71 – 7.31 (m, 7H), 6.93 (dd, *J* = 8.7, 2.5 Hz, 1H), 6.75 (s, 1H), 5.20 (s, 2H).

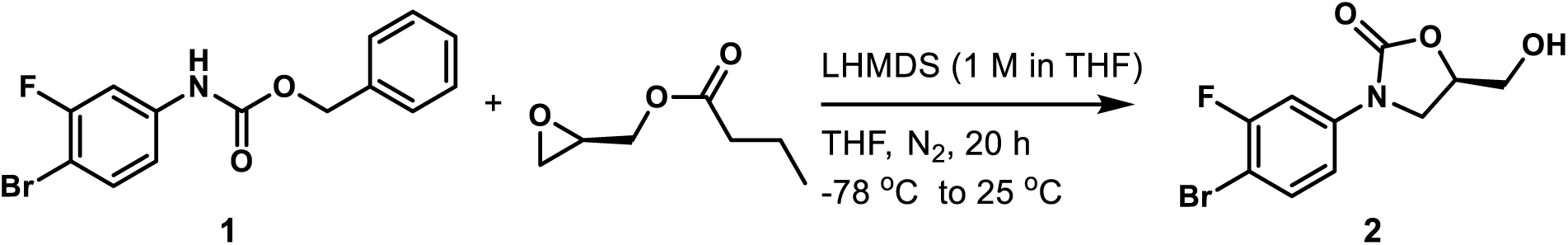

*(R)-3-(4-Bromo-3-fluorophenyl)-5-(hydroxymethyl)oxazolidin-2-one (**2**)*.^43^ Compound **1** (1.62 g, 5 mmol) was dissolved in anhydrous THF (25 mL) and cooled to -78 °C under nitrogen atmosphere. Lithium bis(trimethylsilyl)amide solution (1 M in THF, 4.25 mL, 4.25 mmol) was added to the mixture slowly over a period of 40 min with a syringe. The mixture was stirred at -78 °C for 1 h under a nitrogen atmosphere followed by the addition of (*R*)-(-)-glycidyl butyrate (0.59 mL, 4.25 mmol) dropwise with a syringe at -78 °C. The mixture was stirred at this temperature for an additional 1 h and then gradually warmed to 25 °C. After the reaction was judged to be completed by TLC (20 h), it was quenched with water and extracted with EtOAc three times. The combined organic layers were washed with water and evaporated under reduced pressure by rotary evaporation. The crude reside was purified by flash column chromatography (SiO_2_, eluent gradient 0-100% EtOAc in hexanes) to afford compound **2** as a yellow amorphous solid (1.10 g, 76%). ^1^H NMR (300 MHz, Acetone-*d*_6_) δ 7.74 (dd, *J* = 11.7, 2.7 Hz, 1H), 7.64 (dd, *J* = 8.9, 8.0 Hz, 1H), 7.36 (ddd, *J* = 8.9, 2.7, 1.0 Hz, 1H), 4.96 – 4.70 (m, 1H), 4.38 (dd, *J* = 6.2, 5.6 Hz, 1H), 4.20 (t, *J* = 8.8 Hz, 1H), 4.01 (dd, *J* = 8.8, 6.2 Hz, 1H), 3.90 (ddd, *J* = 12.3, 5.6, 3.4 Hz, 1H), 3.76 (ddd, *J* = 12.3, 6.2, 3.9 Hz, 1H).

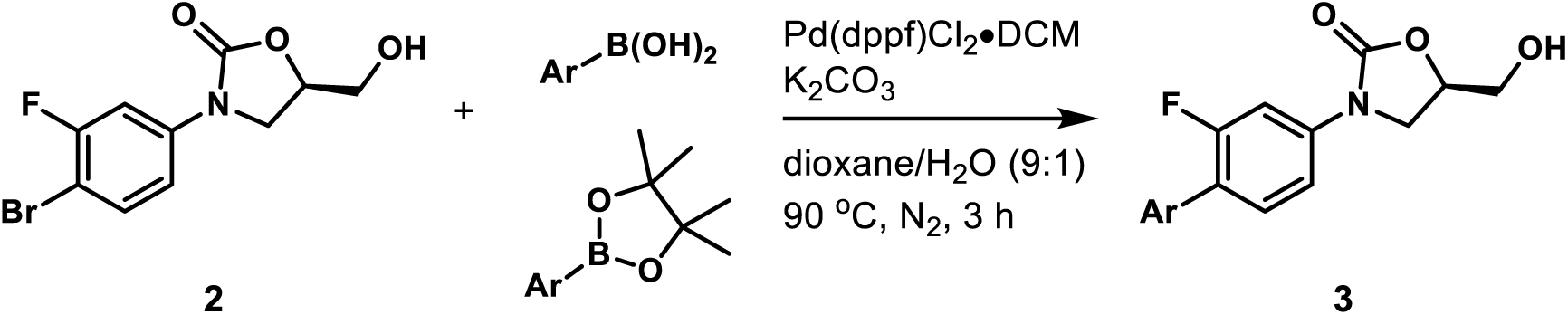

#### General Procedure 1: Synthesis of Heterocyclic Oxazolidinone Analogs 3a-3p.^45^

A mixture of compound **2** (29 mg, 0.1 mmol), aryl boronic acid or ester (0.12 mmol), potassium carbonate (55 mg, 0.4 mmol), and [1,1’-bis(diphenylphosphino)ferrocene]dichloropalladium(II), complex with dichloromethane (8.2 mg, 0.01 mmol) in dioxane/H_2_O (v/v= 9:1, 0.5 mL) was stirred at 90 °C under nitrogen atmosphere for 3 h. The reaction mixture was cooled to room temperature, diluted with EtOAc, washed with water, and concentrated under reduced pressure by rotary evaporation. The crude residue was purified by flash column chromatography (SiO_2_) using an appropriate eluent as described below to afford compound **3**. Various derivatives deviated slightly from this procedure and full descriptions are included as needed for **3a**-**3p**.

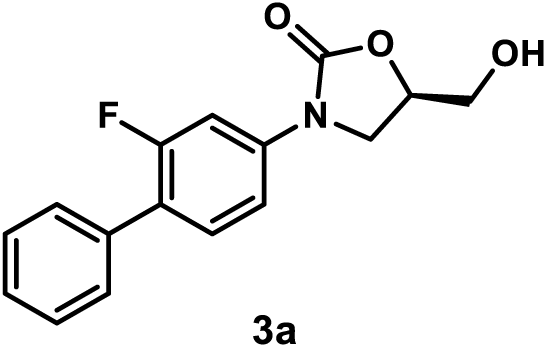

*(R)-3-(2-Fluoro-[1,1’-biphenyl]-4-yl)-5-(hydroxymethyl)oxazolidin-2-one (**3a**)*. Using general procedure 1, employing phenylboronic acid (15 mg, 0.12 mmol), compound **3a** was obtained after flash column chromatography (SiO_2_, eluent gradient 0-100% EtOAc in hexanes) as a white amorphous solid (20 mg, 70%). ^1^H NMR (400 MHz, Acetone-*d*_6_) δ 7.70 (dd, *J* = 13.7, 2.3 Hz, 1H), 7.62 – 7.50 (m, 3H), 7.50 – 7.42 (m, 3H), 7.42 – 7.35 (m, 1H), 4.90 – 4.77 (m, 1H), 4.41 (t, *J* = 5.9 Hz, 1H), 4.24 (t, *J* = 8.8 Hz, 1H), 4.05 (dd, *J* = 8.8, 6.3 Hz, 1H), 3.91 (ddd, *J* = 12.3, 5.9, 3.4 Hz, 1H), 3.78 (ddd, *J* = 12.3, 5.9, 3.9 Hz, 1H). ^13^C NMR (101 MHz, Acetone-*d*_6_) δ 160.5 (d, *J* = 244.5 Hz), 155.3, 140.9 (d, *J* = 11.3 Hz), 136.2 (d, *J* = 1.6 Hz), 131.7 (d, *J* = 5.0 Hz), 129.6 (d, *J* = 3.1 Hz, 2C), 129.4 (2C), 128.4, 124.1 (d, *J* = 14.0 Hz), 114.3 (d, *J* = 3.3 Hz), 106.2 (d, *J* = 29.3 Hz), 74.3, 63.2, 46.9.

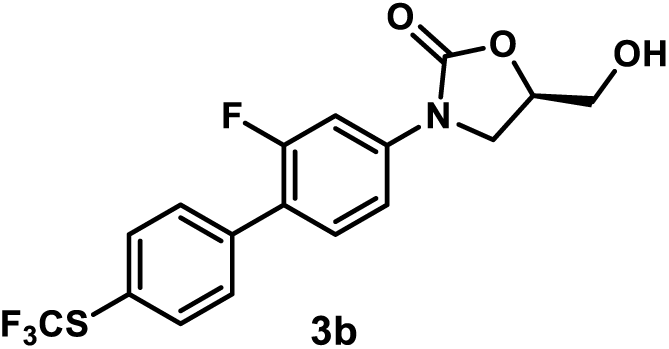

*(R)-3-(2-Fluoro-4’-((trifluoromethyl)thio)-[1,1’-biphenyl]-4-yl)-5-(hydroxymethyl)oxazolidin-2-one (**3b**)*. Using general procedure 1, employing 4,4,5,5-tetramethyl-2-(4-((trifluoromethyl)thio) phenyl)-1,3,2-dioxaborolane (37 mg, 0.12 mmol), compound **3b** was obtained after flash column chromatography (SiO_2_, eluent gradient 0-100% EtOAc in hexanes) as a yellow amorphous solid (26 mg, 67%). ^1^H NMR (400 MHz, Acetone-*d*_6_) δ 7.82 (d, *J* = 8.1 Hz, 2H), 7.77 – 7.69 (m, 3H), 7.61 (t, *J* = 8.7 Hz, 1H), 7.50 (dd, *J* = 8.7, 2.3 Hz, 1H), 4.95 – 4.71 (m, 1H), 4.41 (s, 1H), 4.25 (t, *J* = 8.8 Hz, 1H), 4.06 (dd, *J* = 8.8, 6.2 Hz, 1H), 3.92 (dd, *J* = 12.6, 3.6 Hz, 1H), 3.83 – 3.70 (m, 1H). ^13^C NMR (101 MHz, Acetone-*d*_6_) δ 160.7 (d, *J* = 245.7 Hz), 155.4, 141.9 (d, *J* = 11.3 Hz), 139.5, 137.4 (2C), 131.8 (d, *J* = 4.6 Hz), 131.0 (d, *J* = 3.4 Hz, 2C), 130.9 (q, *J* = 306.9 Hz), 123.6, 122.5 (d, *J* = 13.3 Hz), 114.6 (d, *J* = 3.1 Hz), 106.4 (d, *J* = 29.1 Hz), 74.5, 63.3, 47.0.

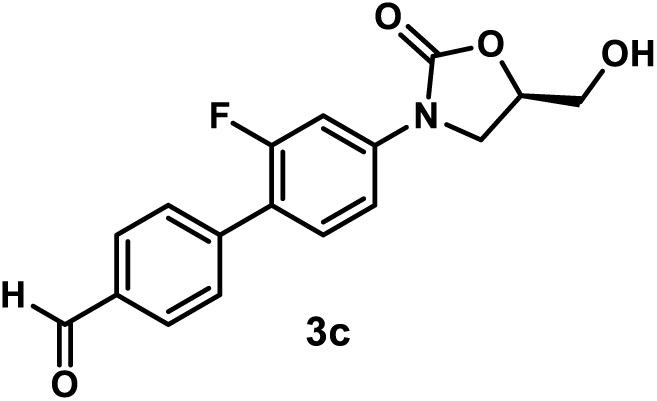

*(R)-2’-Fluoro-4’-(5-(hydroxymethyl)-2-oxooxazolidin-3-yl)-[1,1’-biphenyl]-4-carbaldehyde (**3c**)*. A mixture of compound **2** (58 mg, 0.2 mmol), 4-formylphenylboronic acid (36 mg, 0.24 mmol), potassium carbonate (110 mg, 0.8 mmol), and [1,1’- bis(diphenylphosphino)ferrocene]dichloropalladium(II), complex with dichloromethane (16 mg, 0.02 mmol) in dioxane/H_2_O (v/v= 9:1, 0.5 mL) was stirred at 90 °C under nitrogen atmosphere. After the reaction was judged to be completed by TLC (3 h), it was cooled to room temperature, diluted with EtOAc, washed with water, and concentrated under reduced pressure by rotary evaporation. The crude residue was purified by flash column chromatography (SiO_2,_ eluent gradient 0-100% EtOAc in hexanes) to afford compound **3c** as a yellow amorphous solid (40 mg, 56%). ^1^H NMR (500 MHz, Acetone-*d*_6_) δ 10.09 (s, 1H), 8.01 (d, *J* = 8.3 Hz, 2H), 7.80 (d, *J* = 8.3 Hz, 2H), 7.74 (dd, *J* = 13.8, 2.3 Hz, 1H), 7.63 (t, *J* = 8.8 Hz, 1H), 7.51 (dd, *J* = 8.8, 2.3 Hz, 1H), 4.87 – 4.78 (m, 1H), 4.40 (t, *J* = 5.9 Hz, 1H), 4.25 (t, *J* = 8.9 Hz, 1H), 4.07 (dd, *J* = 8.9, 6.2 Hz, 1H), 3.96 – 3.87 (m, 1H), 3.82 – 3.74 (m, 1H). ^13^C NMR (126 MHz, Acetone-*d*_6_) δ 192.5, 160.6 (d, *J* = 246.0 Hz), 155.3, 142.1 (d, *J* = 1.7 Hz), 141.9 (d, *J* = 11.4 Hz), 136.6, 131.8 (d, *J* = 4.7 Hz), 130.5 (2C), 130.2 (d, *J* = 3.7 Hz, 2C), 122.8 (d, *J* = 13.3 Hz), 114.5 (d, *J* = 3.2 Hz), 106.3 (d, *J* = 29.3 Hz), 74.4, 63.2, 47.0.

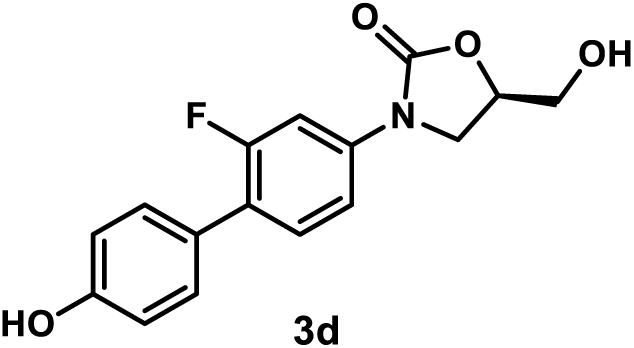

*(R)-3-(2-Fluoro-4’-hydroxy-[1,1’-biphenyl]-4-yl)-5-(hydroxymethyl)oxazolidin-2-one (**3d**)*. A mixture of compound **2** (58 mg, 0.2 mmol), 4-hydroxyphenylboroinc acid (33 mg, 0.24 mmol), potassium carbonate (110 mg, 0.8 mmol), and [1,1’- bis(diphenylphosphino)ferrocene]dichloropalladium(II), complex with dichloromethane (16 mg, 0.02 mmol) in dioxane/H_2_O (v/v= 9:1, 0.5 mL) was stirred at 90 °C under nitrogen atmosphere. After the reaction was judged to be completed by TLC (3 h), it was cooled to room temperature, diluted with EtOAc, washed with water, and concentrated under reduced pressure by rotary evaporation. The crude residue was purified by flash column chromatography (SiO_2,_ eluent gradient 0-100% EtOAc in hexanes) to afford compound **3d** as a yellow amorphous solid (7 mg, 11%). ^1^H NMR (400 MHz, Acetone-*d*_6_) δ 8.48 (s, 1H), 7.66 (dd, *J* = 13.7, 2.3 Hz, 1H), 7.47 (t, *J* = 8.7 Hz, 1H), 7.44 – 7.39 (m, 3H), 6.96 – 6.91 (m, 2H), 4.86 – 4.76 (m, 1H), 4.36 (t, *J* = 5.9 Hz, 1H), 4.22 (t, *J* = 8.8 Hz, 1H), 4.03 (dd, *J* = 8.8, 6.2 Hz, 1H), 3.95 – 3.86 (m, 1H), 3.82 – 3.74 (m, 1H). ^13^C NMR (101 MHz, Acetone-*d*_6_) δ 160.5 (d, *J* = 245.4 Hz), 158.1, 155.4, 140.3, 131.4 (d, *J* = 5.1 Hz), 130.9 (d, *J* = 4.0 Hz, 2C), 127.5 (d, *J* = 2.0 Hz), 124.4 (d, *J* = 14.1 Hz), 116.4 (2C), 116.3, 114.5 (d, *J* = 4.0 Hz), 106.4 (d, *J* = 29.3 Hz), 74.4, 63.4, 47.1.

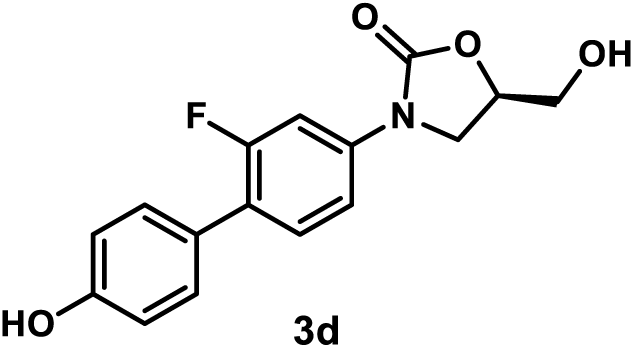

*(R)-3-(4’-(Aminomethyl)-2-fluoro-[1,1’-biphenyl]-4-yl)-5-(hydroxymethyl)oxazolidin-2-one Hydrochloride (**3e**).* A mixture of compound **2** (29 mg, 0.1 mmol), 4-aminomethylphenylboronic acid hydrochloride (23 mg, 0.12 mmol), potassium carbonate (55 mg, 0.4 mmol), and [1,1’- bis(diphenylphosphino)ferrocene]dichloropalladium(II), complex with dichloromethane (8 mg, 0.01 mmol) in dioxane/H_2_O (v/v= 9:1, 0.5 mL) was stirred at 90 °C under nitrogen atmosphere. After the reaction was judged to be completed by TLC (3 h), it was cooled to room temperature and concentrated under reduced pressure with a rotary evaporator. The crude residue was purified by flash column chromatography (SiO_2_, eluent gradient 0-20% MeOH in DCM with 1% ammonia). The appropriate fractions were collected and evaporated under reduced pressure with a rotary evaporator. The resulting residue was dissolved in MeOH (1 mL), charged with the addition of HCl/dioxane (4 M, 50 µL), and concentrated under reduced pressure with a rotary evaporator. The resulting hydrochloride salt was washed with acetone (1 mL) and dried under high vacuum to afford compound **3e** as a white amorphous solid (26 mg, 82%). ^1^H NMR (400 MHz, DMSO-*d*_6_) δ 8.73 (s, 3H), 7.76 – 7.36 (m, 7H), 5.39 (t, *J* = 5.6 Hz, 1H), 4.80 – 4.65 (m, 1H), 4.12 (t, *J* = 8.9 Hz, 1H), 4.04 (s, 2H), 3.92 (dd, *J* = 8.9, 6.0 Hz, 1H), 3.75 – 3.64 (m, 1H), 3.61 – 3.50 (m, 1H). ^13^C NMR (101 MHz, DMSO-*d*_6_) δ 159.1 (d, *J* = 244.7 Hz), 154.4, 139.6 (d, *J* = 11.1 Hz), 134.7, 133.6, 130.9 (d, *J* = 4.6 Hz), 129.4 (2C), 128.7 (d, *J* = 3.0 Hz, 2C), 122.1 (d, *J* = 13.4 Hz), 113.9 (d, *J* = 2.0 Hz), 105.4 (d, *J* = 28.8 Hz), 73.5, 61.5, 46.1, 41.8. MSESI *m/z*: 339.1125 (C_17_H_17_FN_2_O_3_ + Na^+^ requires 339.1115).

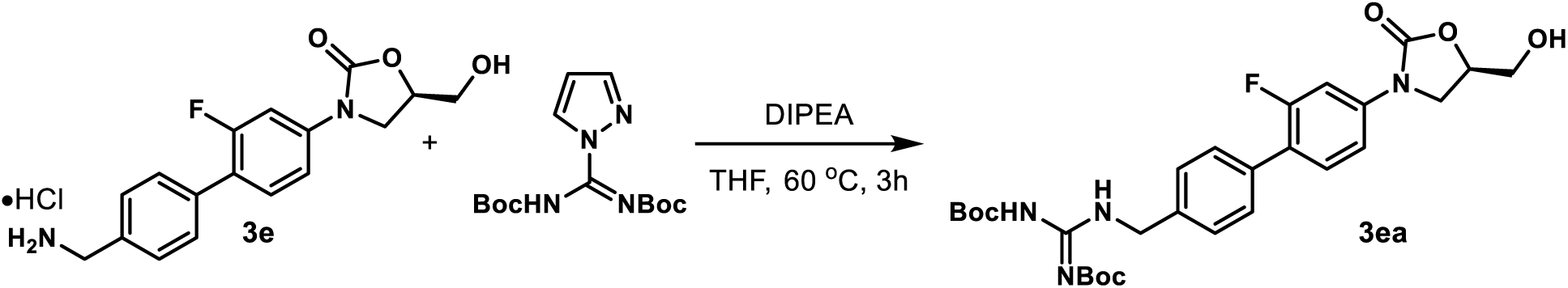

bis-Boc-(*R*)-1-((2’-fluoro-4’-(5-(hydroxymethyl)-2-oxooxazolidin-3-yl)-[1,1’-biphenyl]-4-yl)methyl)guanidine (***3ea***). A mixture of compound **3e** (182 mg, 0.51 mmol), *N*,*N*′-di-Boc-1*H*- pyrazole-1-carboxamidine (240 mg, 0.77 mmol), and *N*,*N*-diisopropylethylamine (266 µL, 1.53 mmol) in THF (10 mL) was stirred at 60 °C. After the reaction was judged to be completed by TLC (3 h), it was cooled to room temperature, diluted with EtOAc, washed with water, and concentrated under reduced pressure with rotary evaporation. The crude residue was purified by flash column chromatography (SiO_2_, eluent gradient 0-75% EtOAc in hexanes) to afford compound **3ea** as a white amorphous solid (74 mg, 26%). ^1^H NMR (400 MHz, Acetone-*d*_6_) δ 11.71 (s, 1H), 8.71 (s, 1H), 7.70 (dd, *J* = 13.7, 2.3 Hz, 1H), 7.59 – 7.42 (m, 6H), 4.86 – 4.76 (m, 1H), 4.70 (d, *J* = 5.8 Hz, 2H), 4.42 (s, 1H), 4.23 (t, *J* = 8.9 Hz, 1H), 4.05 (dd, *J* = 8.9, 6.2 Hz, 1H), 3.96 – 3.85 (m, 1H), 3.82 – 3.73 (m, 1H), 1.51 (s, 9H), 1.44 (s, 9H). ^13^C NMR (101 MHz, Acetone-*d*_6_) δ 164.6, 160.5 (d, *J* = 244.5 Hz), 157.0, 155.3, 153.8, 140.9 (d, *J* = 11.2 Hz), 138.8, 135.2 (d, *J* = 1.5 Hz), 131.6 (d, *J* = 5.0 Hz), 129.8 (d, *J* = 3.1 Hz, 2C), 128.7 (2C), 123.8 (d, *J* = 13.8 Hz), 114.3 (d, *J* = 3.2 Hz), 106.2 (d, *J* = 29.2 Hz), 83.8, 78.9, 74.3, 63.2, 46.9, 44.5, 28.4, 28.1.

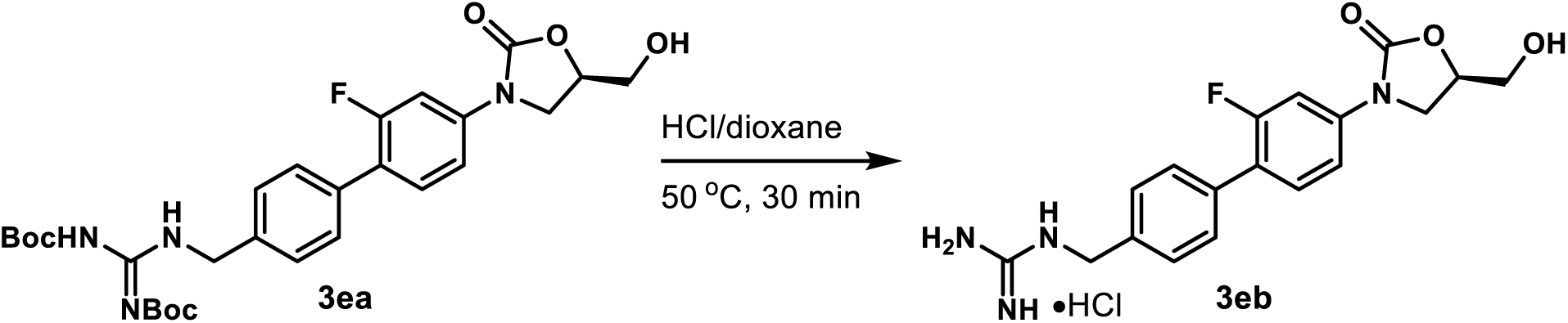

*(R)-1-((2’-Fluoro-4’-(5-(hydroxymethyl)-2-oxooxazolidin-3-yl)-[1,1’-biphenyl]-4-yl)methyl)guanidine Hydrochloride (**3eb**)*. A mixture of **3ea** (28 mg, 0.05 mmol) in HCl/dioxane (4 M, 1 mL) was stirred at 50 °C. After the reaction was judged to be completed by TLC (30 min), its solvent was removed under reduced pressure with rotary evaporation. The residue was washed with acetone (1 mL) and dried under high vacuum to give compound **3eb** as a white amorphous solid (11 mg, 56%). ^1^H NMR (400 MHz, DMSO-*d*_6_) δ 8.31 (t, *J* = 6.2 Hz, 1H), 7.63 (dd, *J* = 13.5, 2.3 Hz, 1H), 7.60 – 7.49 (m, 3H), 7.48 – 7.35 (m, 3H), 5.31 (t, *J* = 5.6 Hz, 1H), 4.79 – 4.69 (m, 1H), 4.45 (d, *J* = 6.2 Hz, 2H), 4.13 (t, *J* = 9.0 Hz, 1H), 3.90 (dd, *J* = 9.0, 6.1 Hz, 1H), 3.69 (ddd, *J* = 12.4, 5.6, 3.3 Hz, 1H), 3.57 (ddd, *J* = 12.4, 5.6, 3.9 Hz, 1H). ^13^C NMR (101 MHz, DMSO-*d*_6_) δ 159.1 (d, *J* = 243.9 Hz), 157.2, 154.4, 139.5 (d, *J* = 11.1 Hz), 136.8, 133.9, 130.9 (d, *J* = 4.4 Hz), 128.8 (2C), 127.5 (2C), 122.3 (d, *J* = 12.9 Hz), 113.8, 105.4 (d, *J* = 28.9 Hz), 73.4, 61.6, 46.0, 43.6. MSESI *m/z*: 359.1532 (C_18_H_19_FN_4_O_3_ + H^+^ requires 359.1514).

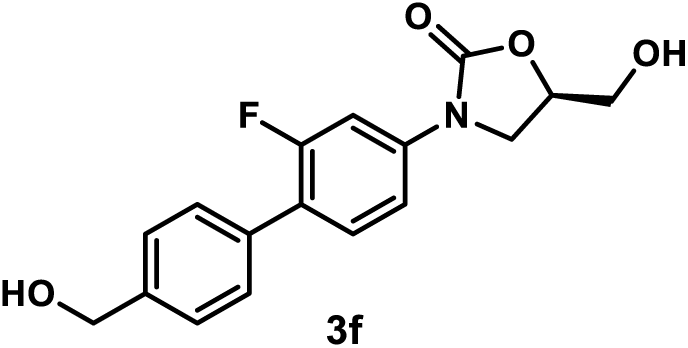

*(R)-3-(2-Fluoro-4’-(hydroxymethyl)-[1,1’-biphenyl]-4-yl)-5-(hydroxymethyl)oxazolidin-2-one (**3f**).* Using general procedure 1, employing 4-(hydroxymethyl)benzeneboronic acid (18 mg, 0.12 mmol), compound **3f** was obtained after flash column chromatography (SiO_2_, eluent gradient 0-10% MeOH in DCM) as a yellow amorphous solid (10 mg, 32%). ^1^H NMR (500 MHz, DMSO- *d*_6_) δ 7.62 (dd, *J* = 13.6, 2.2 Hz, 1H), 7.55 (t, *J* = 8.6 Hz, 1H), 7.52 – 7.48 (m, 2H), 7.44 (dd, *J* = 8.6, 2.2 Hz, 1H), 7.41 (d, *J* = 8.0 Hz, 2H), 5.29 – 5.09 (m, 2H), 4.81 – 4.70 (m, 1H), 4.54 (d, *J* = 5.3 Hz, 2H), 4.13 (t, *J* = 8.9 Hz, 1H), 3.88 (dd, *J* = 8.9, 6.1 Hz, 1H), 3.74 – 3.65 (m, 1H), 3.63 – 3.54 (m, 1H). ^13^C NMR (101 MHz, DMSO-*d*_6_) δ 159.0 (d, *J* = 244.3 Hz), 154.3, 142.0, 139.2 (d, *J* = 11.1 Hz), 133.0, 130.8 (d, *J* = 4.8 Hz), 128.3 (d, *J* = 3.0 Hz, 2C), 126.7 (2C), 122.7 (d, *J* = 13.5 Hz), 113.8 (d, *J* = 3.1 Hz), 105.3 (d, *J* = 29.0 Hz), 73.4, 62.6, 61.6, 46.0. MSESI *m/z*: 340.0949 (C_17_H_16_FNO_4_ + Na^+^ requires 340.0956).

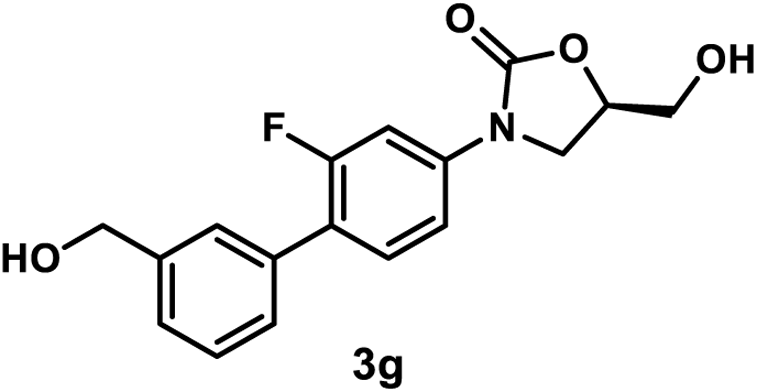

*(R)-3-(2-Fluoro-3’-(hydroxymethyl)-[1,1’-biphenyl]-4-yl)-5-(hydroxymethyl)oxazolidin-2-one (**3g**)*. A mixture of compound **2** (58 mg, 0.2 mmol), 3-(hydroxymethyl)phenylboroinc acid (36 mg, 0.24 mmol), potassium carbonate (110 mg, 0.8 mmol), and [1,1’- bis(diphenylphosphino)ferrocene]dichloropalladium(II), complex with dichloromethane (16 mg, 0.02 mmol) in dioxane/H_2_O (v/v= 9:1, 0.5 mL) was stirred at 90 °C under nitrogen atmosphere. After the reaction was judged to be completed by TLC (3 h), it was cooled to room temperature, diluted with EtOAc, washed with water, and concentrated under reduced pressure by rotary evaporation. The crude residue was purified by flash column chromatography (SiO_2,_ eluent gradient 0-10% MeOH in DCM) to afford compound **3g** as a brown amorphous solid (55 mg, 87%). ^1^H NMR (400 MHz, Methanol-*d*_4_) δ 7.63 (dd, *J* = 13.3, 2.3 Hz, 1H), 7.54 – 7.47 (m, 2H), 7.44 – 7.33 (m, 4H), 4.81 – 4.73 (m, 1H), 4.66 (s, 2H), 4.16 (t, *J* = 9.0 Hz, 1H), 3.97 (dd, *J* = 9.0, 6.4 Hz, 1H), 3.87 (dd, *J* = 12.6, 3.2 Hz, 1H), 3.71 (dd, *J* = 12.6, 4.0 Hz, 1H). ^13^C NMR (101 MHz, Methanol-*d*_4_) δ 161.0 (d, *J* = 246.4 Hz), 156.9, 143.1, 140.6 (d, *J* = 11.1 Hz), 136.7 (d, *J* = 1.0 Hz), 132.0 (d, *J* = 5.1 Hz), 129.6, 128.8 (d, *J* = 3.0 Hz), 128.4 (d, *J* = 3.0 Hz), 127.2, 125.5 (d, *J* = 13.1 Hz), 115.0 (d, *J* = 3.0 Hz), 107.2 (*J* = 29,3 Hz), 75.2, 65.1, 63.3, 47.6.

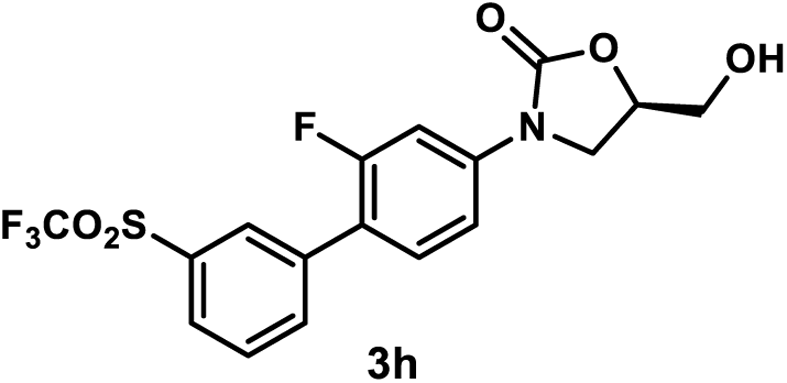

*(R)-3-(2-Fluoro-3’-((trifluoromethyl)sulfonyl)-[1,1’-biphenyl]-4-yl)-5-(hydroxymethyl)oxazolidin-2-one (**3h**)*. Using general procedure 1, employing 4,4,5,5-tetramethyl-2-(3-trifluoromethanesulfonylphenyl)-1,3,2-dioxaborolane (40 mg, 0.12 mmol), compound **3h** was obtained after flash column chromatography (SiO_2_, eluent gradient 0-100% EtOAc in hexanes) as a foamy yellow amorphous solid (34 mg, 81%). ^1^H NMR (400 MHz, Acetone-*d*_6_) δ 8.26 (s, 1H), 8.22 (d, *J* = 7.9 Hz, 1H), 8.14 (d, *J* = 7.9 Hz, 1H), 7.98 (t, *J* = 7.9 Hz, 1H), 7.78 (dd, *J* = 13.9, 2.3 Hz, 1H), 7.69 (t, *J* = 8.8 Hz, 1H), 7.55 (dd, *J* = 8.8, 2.3 Hz, 1H), 4.93 – 4.74 (m, 1H), 4.44 (s, 1H), 4.26 (t, *J* = 8.9 Hz, 1H), 4.08 (dd, *J* = 8.9, 6.2 Hz, 1H), 3.92 (dd, *J* = 12.4, 3.3 Hz, 1H), 3.79 (dd, *J* = 12.4, 3.8 Hz, 1H). ^13^C NMR (101 MHz, Acetone-*d*_6_) δ 159.7 (d, *J* = 245.7 Hz), 154.4, 141.4 (d, *J* = 11.4 Hz), 137.7 (d, *J* = 1.6 Hz), 137.3 (d, *J* = 3.0 Hz), 131.3 (d, *J* = 1.6 Hz), 130.9, 130.9, 130.3 (d, *J* = 4.2 Hz), 129.5, 120.2 (d, *J* = 13.3 Hz), 119.9 (q, *J* = 325.3 Hz), 113.8 (d, *J* = 3.1 Hz), 105.4 (d, *J* = 29.0 Hz), 73.5, 62.2, 46.1.

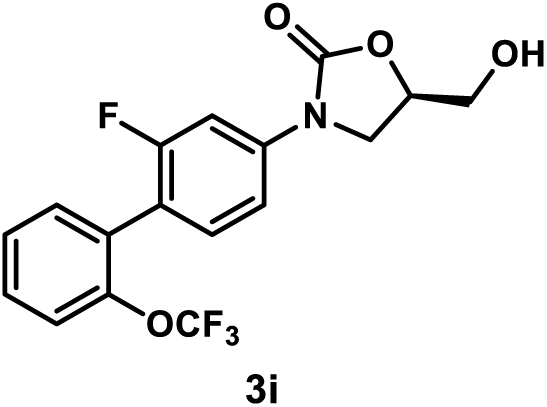

*(R)-3-(2-Fluoro-2’-(trifluoromethoxy)-[1,1’-biphenyl]-4-yl)-5-(hydroxymethyl)oxazolidin-2-one (**3i**)*. Using general procedure 1, employing 2-(trifluoromethoxy)phenylboronic acid (25 mg, 0.12 mmol), compound **3i** was obtained after flash column chromatography (SiO_2_, eluent gradient 0-100% EtOAc in hexanes) as yellow oil (32 mg, 86%). ^1^H NMR (400 MHz, Acetone-*d*_6_) δ 7.72 (dd, *J* = 12.8, 2.2 Hz, 1H), 7.59 – 7.44 (m, 5H), 7.41 (t, *J* = 8.4 Hz, 1H), 4.98 – 4.75 (m, 1H), 4.44 (t, *J* = 5.9 Hz, 1H), 4.24 (t, *J* = 8.9 Hz, 1H), 4.06 (dd, *J* = 8.9, 6.2 Hz, 1H), 3.99 – 3.86 (m, 1H), 3.84 – 3.73 (m, 1H). ^13^C NMR (101 MHz, Acetone-*d*_6_) δ 160.4 (d, *J* = 244.8 Hz), 155.3, 147.5, 141.8 (d, *J* = 11.2 Hz), 133.1, 132.6 (d, *J* = 4.6 Hz), 130.7, 129.9, 128.2, 121.7 (d, *J* = 1.6 Hz), 121.3 (q, *J* = 256.3 Hz), 119.4 (d, *J* = 16.4 Hz), 113.9 (d, *J* = 3.2 Hz), 105.6 (d, *J* = 28.7 Hz), 74.3, 63.2, 46.9.

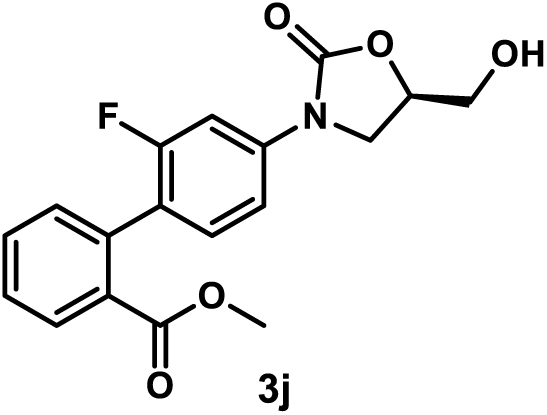

*Methyl (R)-2’-Fluoro-4’-(5-(hydroxymethyl)-2-oxooxazolidin-3-yl)-[1,1’-biphenyl]-2-carboxylate (**3j**)*. Using general procedure 1, employing 2-methoxycarbonylphenylboronic acid (22 mg, 0.12 mmol), compound **3j** was obtained after flash column chromatography (SiO_2_, eluent gradient 0-100% EtOAc in hexanes) followed by preparative TLC (eluent 100% EtOAc) as yellow oil (33 mg, 94%). ^1^H NMR (400 MHz, acetone) δ 7.93 (d, *J* = 7.6 Hz, 1H), 7.74 – 7.59 (m, 2H), 7.52 (t, *J* = 7.6 Hz, 1H), 7.45 – 7.31 (m, 3H), 4.89 – 4.77 (m, 1H), 4.46 (s, 1H), 4.23 (t, *J* = 8.8 Hz, 1H), 4.04 (dd, *J* = 8.8, 6.2 Hz, 1H), 3.91 (dd, *J* = 12.4, 3.3 Hz, 1H), 3.79 (dd, *J* = 12.4, 3.8 Hz, 1H), 3.68 (s, 3H).^13^C NMR (101 MHz, Acetone-*d*_6_) δ 168.2, 160.3 (d, *J* = 242.6 Hz), 155.3, 141.0 (d, *J* = 11.1 Hz), 136.7, 132.7, 132.3, 132.1, 131.5 (d, *J* = 5.0 Hz), 130.7, 128.7, 124.3 (d, *J* = 16.1 Hz), 113.7 (d, *J* = 3.1 Hz), 105.2 (d, *J* = 28.9 Hz), 74.3, 63.2, 52.2, 46.9.

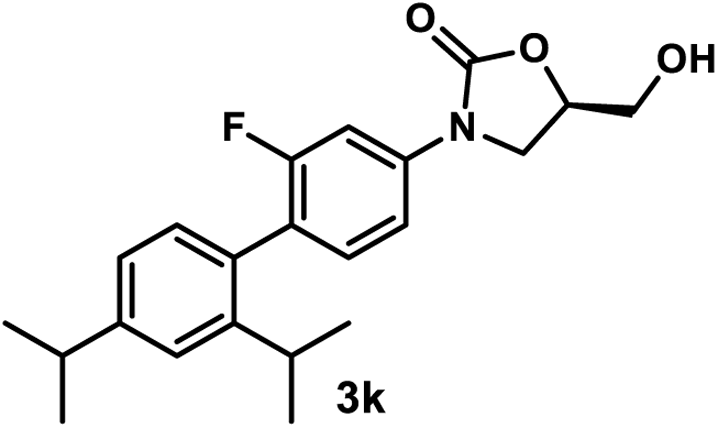

*(R)-3-(2-Fluoro-2’,4’-diisopropyl-[1,1’-biphenyl]-4-yl)-5-(hydroxymethyl)oxazolidin-2-one (**3k**)*. Using general procedure 1, employing [2,4-bis(propan-2-yl)phenyl]boronic acid (25 mg, 0.12 mmol), compound **3k** was obtained after flash column chromatography (SiO_2_, eluent gradient 0-100% EtOAc in hexanes) as a white amorphous solid (33 mg, 89%). ^1^H NMR (400 MHz, Acetone-*d*_6_) δ 7.67 (dd, *J* = 12.3, 2.3 Hz, 1H), 7.43 (dd, *J* = 8.5, 2.3 Hz, 1H), 7.32 (s, 1H), 7.27 (t, *J* = 8.5 Hz, 1H), 7.13 (d, *J* = 7.9, 1H), 7.06 (d, *J* = 7.9 Hz, 1H), 4.91 – 4.71 (m, 1H), 4.40 (t, *J* = 5.9 Hz, 1H), 4.24 (t, *J* = 8.8 Hz, 1H), 4.05 (dd, *J* = 8.8, 6.2 Hz, 1H), 3.92 (ddd, *J* = 12.4, 5.9, 3.5 Hz, 1H), 3.79 (ddd, *J* = 12.4, 5.9, 4.0 Hz, 1H), 2.96 (p, *J* = 7.0 Hz, 1H), 2.82 – 2.78 (m, 1H), 1.28 (d, *J* = 7.0 Hz, 6H), 1.25 – 1.01 (m, 6H). ^1^H NMR (400 MHz, acetone) δ 7.67 (dd, *J* = 12.3, 2.3 Hz, 1H), 7.43 (dd, *J* = 8.5, 2.2 Hz, 1H), 7.32 (d, *J* = 1.9 Hz, 1H), 7.27 (t, *J* = 8.5 Hz, 1H), 7.13 (dd, *J* = 7.8, 1.8 Hz, 1H), 7.06 (d, *J* = 7.8 Hz, 1H), 4.89 – 4.75 (m, 1H), 4.40 (t, *J* = 5.9 Hz, 1H), 4.24 (t, *J* = 8.8 Hz, 1H), 4.05 (dd, *J* = 8.8, 6.2 Hz, 1H), 3.92 (ddd, *J* = 12.4, 5.9, 3.5 Hz, 1H), 3.79 (ddd, *J* = 12.4, 5.9, 4.0 Hz, 1H), 2.96 (p, *J* = 6.9 Hz, 1H), 2.84 – 2.77 (m, 1H), 1.28 (d, *J* = 6.9 Hz, 6H), 1.22 – 1.05 (m, 6H). ^13^C NMR (101 MHz, Acetone-*d*_6_) δ 160.6 (d, *J* = 241.5 Hz), 155.4, 149.8, 148.0, 140.9 (d, *J* = 10.7 Hz), 132.9 (d, *J* = 5.0 Hz), 132.5, 131.2, 124.6 (d, *J* = 17.6 Hz), 124.4 (d, *J* = 5.2 Hz), 113.9 (d, *J* = 3.2 Hz), 105.7 (d, *J* = 29.1 Hz), 74.4, 63.3, 47.1, 35.0, 31.1, 24.5.

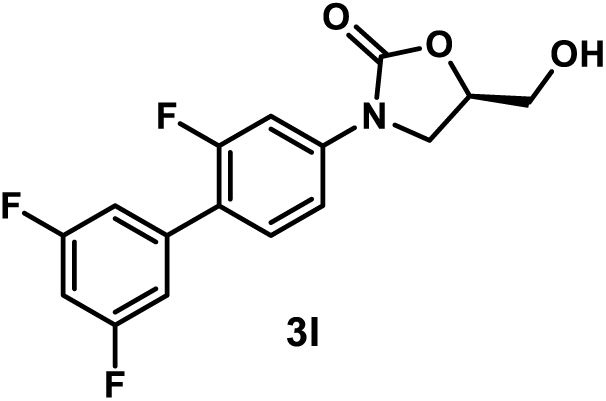

*(R)-5-(Hydroxymethyl)-3-(2,3’,5’-trifluoro-[1,1’-biphenyl]-4-yl)oxazolidin-2-one (**3l**)*. A mixture of compound **2** (58 mg, 0.2 mmol), 3,5-difluorophenylboroinc acid (38 mg, 0.24 mmol), potassium carbonate (110 mg, 0.8 mmol), and [1,1’-bis(diphenylphosphino)ferrocene]dichloropalladium(II), complex with dichloromethane (16 mg, 0.02 mmol) in dioxane/H_2_O (v/v= 9:1, 0.5 mL) was stirred at 90 °C under nitrogen atmosphere. After the reaction was judged to be completed by TLC (3 h), it was cooled to room temperature, diluted with EtOAc, washed with water, and concentrated under reduced pressure by rotary evaporation. The crude residue was purified by flash column chromatography (SiO_2,_ eluent gradient 0-100% EtOAc in hexanes) to afford compound **3l** as a yellow amorphous solid (48 mg, 66%). ^1^H NMR (300 MHz, acetone) δ 7.73 (dd, *J* = 13.9, 2.3 Hz, 1H), 7.62 (t, *J* = 8.8 Hz, 1H), 7.53 – 7.44 (m, 1H), 7.34 – 7.18 (m, 2H), 7.05 (tt, *J* = 9.2, 2.3 Hz, 1H), 4.95 – 4.71 (m, 1H), 4.47 – 4.37 (m, 1H), 4.25 (t, *J* = 8.8 Hz, 1H), 4.06 (dd, *J* = 8.8, 6.2 Hz, 1H), 3.99 – 3.85 (m, 1H), 3.84 – 3.67 (m, 1H). ^13^C NMR (101 MHz, Acetone-*d*_6_) δ 163.9 (dd, *J* = 246.4, 13.1 Hz, 2C), 160.5 (d, *J* = 246.4 Hz), 155.3, 142.0 (d, *J* = 12.1 Hz), 139. 7 (td, *J* = 10.1, 1.0 Hz), 131.6 (d, *J* = 5.1 Hz), 114.5 (d, *J* = 3.0 Hz), 112.5 (dd, *J* = 26.3, 3.0 Hz, 2C), 112.5 (d, *J* = 12.1, 3.0 Hz), 106.3 (d, *J* = 29.3 Hz), 103.5 (t, J = 26.3 Hz), 74.4, 63.2, 47.0.

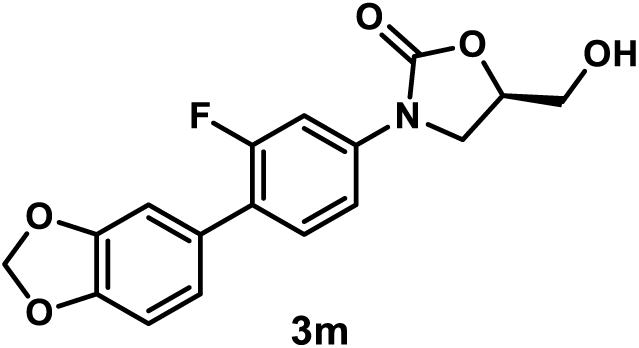

*(R)-3-(4-(Benzo[d][1,3]dioxol-5-yl)-3-fluorophenyl)-5-(hydroxymethyl)oxazolidin-2-one (**3m**)*. A mixture of compound **2** (58 mg, 0.2 mmol), 3,4-(methylenedioxy)phenylboronic acid (40 mg, 0.24 mmol), potassium carbonate (110 mg, 0.8 mmol), and [1,1’-bis(diphenylphosphino)ferrocene]dichloropalladium(II), complex with dichloromethane (16 mg, 0.02 mmol) in dioxane/H_2_O (v/v= 9:1, 0.5 mL) was stirred at 90 °C under nitrogen atmosphere. After the reaction was judged to be completed by TLC (3 h), it was cooled to room temperature, diluted with EtOAc, washed with water, and concentrated under reduced pressure by rotary evaporation. The crude residue was purified by flash column chromatography (SiO_2,_ eluent gradient 0-67% EtOAc in hexanes) to afford compound **3m** as a brown amorphous solid (48 mg, 64%). ^1^H NMR (300 MHz, Acetone-*d*_6_) δ 7.67 (dd, *J* = 13.7, 2.2 Hz, 1H), 7.55 – 7.35 (m, 2H), 7.12 – 7.01 (m, 2H), 6.99 – 6.83 (m, 1H), 6.05 (s, 2H), 4.93 – 4.68 (m, 1H), 4.39 (t, *J* = 5.9 Hz, 1H), 4.22 (t, *J* = 8.9 Hz, 1H), 4.04 (dd, *J* = 8.9, 6.2 Hz, 1H), 3.91 (ddd, *J* = 12.3, 5.9, 3.4 Hz, 1H), 3.78 (ddd, *J* = 12.3, 5.9, 3.9 Hz, 1H). ^13^C NMR (101 MHz, Acetone-*d*_6_) δ 160.5 (d, *J* = 245.4 Hz), 155.4, 148.9, 148.3, 140.7 (d, *J* = 11.1 Hz), 131.6 (d, *J* = 5.1 Hz), 130.1 (d, *J* = 1.0 Hz), 124.1 (d, *J* = 14.1 Hz), 123.4 (d, *J* = 3.0 Hz), 114.4 (d, *J* = 3.0 Hz), 110.0 (d, *J* = 4.0 Hz), 109.3, 106.4 (d, *J* = 29.3 Hz), 102.4, 74.4, 63.3, 47.1.

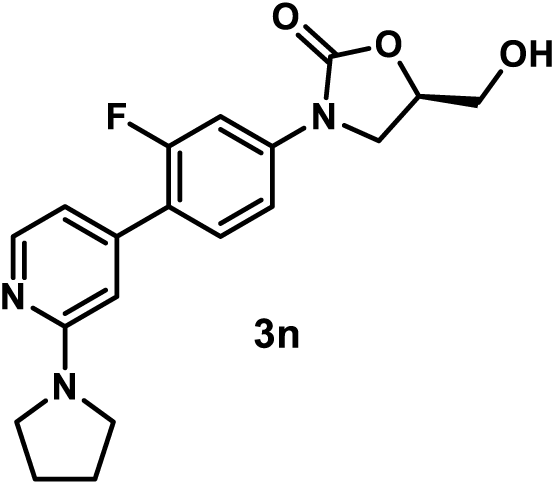

*(R)-3-(3-Fluoro-4-(2-(pyrrolidin-1-yl)pyridin-4-yl)phenyl)-5-(hydroxymethyl)oxazolidin-2-one (**3n**)*. Using general procedure 1, employing 2-(pyrrolidino)pyridine-4-boronic acid pinacol ester (33 mg, 0.12 mmol), compound **3n** was obtained after flash column chromatography (SiO_2_, eluent gradient 0-10% MeOH in DCM) followed by washing with acetone (1 mL) as a gray amorphous solid (29 mg, 81%). 1H NMR (600 MHz, Acetone-d6) δ 8.12 (d, *J* = 5.2 Hz, 1H), 7.70 (dd, *J* = 13.7, 2.2 Hz, 1H), 7.60 (t, *J* = 8.7 Hz, 1H), 7.51 – 7.44 (m, 1H), 6.70 (d, *J* = 5.2 Hz, 1H), 6.55 (s, 1H), 4.91 – 4.75 (m, 1H), 4.39 (t, *J* = 5.9 Hz, 1H), 4.24 (t, *J* = 8.7 Hz, 1H), 4.05 (dd, *J* = 8.7, 6.2 Hz, 1H), 3.95 – 3.85 (m, 1H), 3.82 – 3.72 (m, 1H), 3.63 – 3.37 (m, 4H), 2.02 – 1.88 (m, 4H). ^13^C NMR (101 MHz, Acetone-*d*_6_) δ 160.9 (d, *J* = 246.0 Hz), 158.9, 155.4, 149.3, 144.6 (d, *J* = 1.0 Hz), 141.8 (d, *J* = 11.4 Hz), 131.6 (d, *J* = 4.9 Hz), 122.9 (d, *J* = 13.2 Hz), 114.5 (d, *J* = 3.1 Hz), 112.1 (d, *J* = 3.4 Hz), 106.6, 106.4 (d, *J* = 26.9 Hz), 74.5, 63.3, 47.4, 47.1, 26.3. MSESI *m/z*: 358.1572 (C_19_H_20_FN_3_O_3_ + H^+^ requires 358.1561).

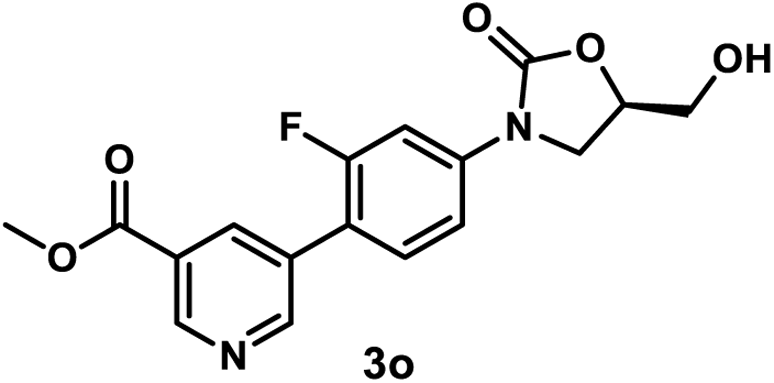

*Methyl (R)-5-(2-Fluoro-4-(5-(hydroxymethyl)-2-oxooxazolidin-3-yl)phenyl)nicotinate (**3o**)*. Using general procedure 1, employing methyl 5-(4,4,5,5-tetramethyl-1,3,2-dioxaborolan-2-yl)nicotinate (32 mg, 0.12 mmol), compound **3o** was obtained after flash column chromatography (SiO_2_, eluent gradient 0-10% MeOH in DCM) as a yellow amorphous solid (27 mg, 77%). ^1^H NMR (600 MHz, Acetone-*d*_6_) δ 9.11 (d, *J* = 1.9 Hz, 1H), 8.99 (d, *J* = 1.9 Hz, 1H), 8.46 (s, 1H), 7.79 (dd, *J* = 13.7, 2.3 Hz, 1H), 7.69 (t, *J* = 8.7 Hz, 1H), 7.55 (dd, *J* = 8.7, 2.3 Hz, 1H), 4.90 – 4.75 (m, 1H), 4.40 (t, *J* = 5.9 Hz, 1H), 4.27 (t, *J* = 8.8 Hz, 1H), 4.08 (dd, *J* = 8.8, 6.1 Hz, 1H), 3.96 (s, 3H), 3.95 – 3.89 (m, 1H), 3.82 – 3.73 (m, 1H). ^13^C NMR (151 MHz, Acetone-*d*_6_) δ 166.3, 160.9 (d, *J* = 245.6 Hz), 155.4, 154.0 (d, *J* = 3.6 Hz), 150.1, 142.4 (d, *J* = 11.3 Hz), 137.3 (d, *J* = 3.5 Hz), 132.2 (d, *J* = 1.5 Hz), 131.9 (d, *J* = 4.2 Hz), 127.0, 119.8 (d, *J* = 13.9 Hz), 114.9 (d, *J* = 3.4 Hz), 106.5 (d, *J* = 28.8 Hz), 74.6, 63.3, 53.0, 47.2. MSESI *m/z*: 369.0856 (C_17_H_15_FN_2_O_5_ + Na^+^ requires 369.0857).

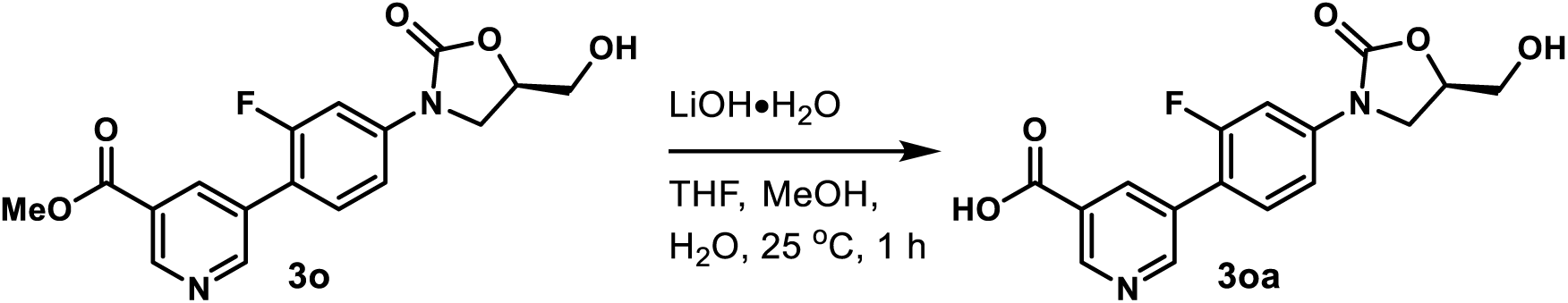

*(R)-5-(2-Fluoro-4-(5-(hydroxymethyl)-2-oxooxazolidin-3-yl)phenyl)nicotinic Acid (**3oa**)*. Lithium hydroxide monohydrate (42 mg, 1 mmol) was added to a solution of **3o** (35 mg, 0.1 mmol) in THF (1.5 mL), MeOH (0.5 mL), and H_2_O (0.5 mL). The mixture was stirred at 25 °C. After the reaction was judged to be completed by TLC (1 h), it was acidified to pH = 1 with the addition of formic acid and concentrated under reduced pressure by rotary evaporation. The crude residue was purified by flash column chromatography (SiO_2_, eluent 10% MeOH in DCM then 10% MeOH in DCM with 1% formic acid) to give compound **3oa** as a yellow amorphous solid (19 mg, 56%). ^1^H NMR (400 MHz, DMSO-*d*_6_) δ 9.05 (s, 1H), 8.92 (s, 1H), 8.37 (s, 1H), 8.21 (s, 1H), 7.79 – 7.63 (m, 2H), 7.55 – 7.41 (m, 1H), 4.81 – 4.70 (m, 1H), 4.15 (t, *J* = 8.9 Hz, 1H), 3.90 (dd, *J* = 8.9, 5.9 Hz, 1H), 3.70 (dd, *J* = 12.4, 3.2 Hz, 1H), 3.58 (dd, *J* = 12.4, 3.9 Hz, 1H). ^13^C NMR (101 MHz, DMSO-*d*_6_) δ 163.5, 159.3 (d, *J* = 245.2 Hz), 154.4, 151.9 (d, *J* = 3.6 Hz), 149.2, 140.5 (d, *J* = 11.4 Hz), 136.3, 131.0 (d, *J* = 4.1 Hz), 130.3, 128.1, 118.6 (d, *J* = 13.4 Hz), 114.0 (d, *J* = 3.0 Hz), 105.3 (d, *J* = 28.4 Hz), 73.5, 61.7, 46.0. MSESI *m/z*: 355.0733 (C_16_H_13_FN_2_O_5_ + Na^+^ requires 355.0701).

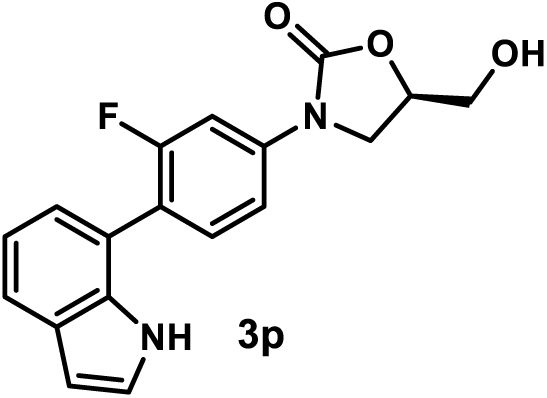

*(R)-3-(3-Fluoro-4-(1H-indol-7-yl)phenyl)-5-(hydroxymethyl)oxazolidin-2-one (**3p**)*. Using general procedure 1, employing 7-(4,4,5,5-tetramethyl-1,3,2-dioxaborolan-2-yl)-1*H*-indole (29 mg, 0.12 mmol), compound **3p** was obtained after flash column chromatography (SiO_2_, eluent gradient 0-100% EtOAc in hexanes) as a brown amorphous solid (25 mg, 76%). ^1^H NMR (400 MHz, Acetone-*d*_6_) δ 10.16 (s, 1H), 7.74 (dd, *J* = 13.0, 2.2 Hz, 1H), 7.63 (t, *J* = 4.5 Hz, 1H), 7.55 (t, *J* = 8.4 Hz, 1H), 7.45 (dd, *J* = 8.4, 2.2 Hz, 1H), 7.34 (t, *J* = 2.6 Hz, 1H), 7.19 – 6.98 (m, 2H), 6.56 (t, *J* = 2.6 Hz, 1H), 5.04 – 4.75 (m, 1H), 4.44 (s, 1H), 4.25 (t, *J* = 8.8 Hz, 1H), 4.07 (dd, *J* = 8.8, 6.1 Hz, 1H), 3.93 (dd, *J* = 12.4, 3.3 Hz, 1H), 3.79 (dd, *J* = 12.4, 3.8 Hz, 1H). ^13^C NMR (101 MHz, Acetone-*d*_6_) δ 160.8 (d, *J* = 244.3 Hz), 155.4, 141.0 (d, *J* = 11.0 Hz), 135.3, 132.6 (d, *J* = 5.6 Hz), 129.6, 126.1, 123.4, 122.1 (d, *J* = 16.4 Hz), 121.1, 120.2, 120.2, 114.4 (d, *J* = 3.1 Hz), 106.3 (d, *J* = 28.8 Hz), 102.7, 74.3, 63.2, 47.1.

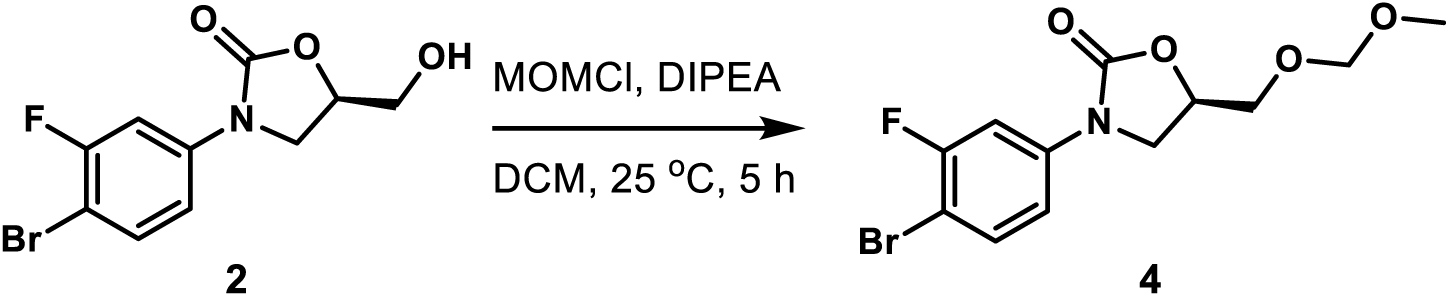

*(R)-3-(4-Bromo-3-fluorophenyl)-5-((methoxymethoxy)methyl)oxazolidin-2-one (**4**)*. To a solution of compound **2** (300 mg, 1.04 mmol) in DCM (5 mL), *N*,*N*-diisopropylethylamine (0.54 mL, 3.11 mmol) and methyl chloromethyl ether (0.24 mL, 3.11 mmol) were added. The reaction mixture was stirred at 25 °C. After the reaction was judged to be completed by TLC (5 h), it was diluted with EtOAc, washed with water, and concentrated under reduced pressure by rotary evaporation. The crude residue was purified by flash column chromatography (SiO_2_, eluent gradient 0-67% EtOAc in hexanes) to afford compound **4** as a white amorphous solid (296 mg, 85%). ^1^H NMR (300 MHz, Acetone-*d*_6_) δ 7.74 (dd, *J* = 11.6, 2.6 Hz, 1H), 7.64 (dd, *J* = 8.9, 8.0 Hz, 1H), 7.37 (ddd, *J* = 8.9, 2.6, 1.0 Hz, 1H), 4.95 (dddd, *J* = 9.0, 6.2, 4.4, 3.5 Hz, 1H), 4.66 (s, 2H), 4.26 (t, *J* = 9.0 Hz, 1H), 4.00 (dd, *J* = 9.0, 6.2 Hz, 1H), 3.86 (dd, *J* = 11.3, 3.5 Hz, 1H), 3.80 (dd, *J* = 11.3, 4.4 Hz, 1H), 3.32 (s, 3H). ^13^C NMR (101 MHz, Acetone-*d*_6_) δ 159.7 (d, *J* = 243.1 Hz), 155.0, 141.2 (d, *J* = 10.0 Hz), 134.3 (d, *J* = 1.8 Hz), 115.6 (d, *J* = 3.4 Hz), 106.9 (d, *J* = 28.2 Hz), 102.1 (d, *J* = 21.1 Hz), 97.3, 72.7, 68.6, 55.4, 47.4.

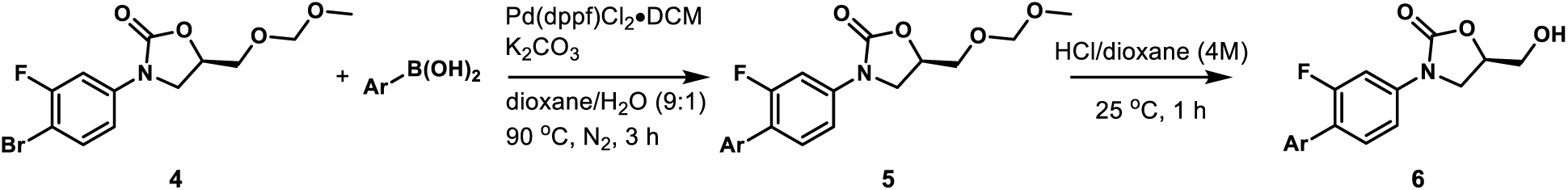

#### General Procedure 2: Synthesis of Heterocyclic Oxazolidinone Analogs 6a-6c

A mixture of compound **4** (67 mg, 0.2 mmol), aryl boronic acid (0.24 mmol), potassium carbonate (110 mg, 0.8 mmol), and [1,1’-bis(diphenylphosphino)ferrocene]dichloropalladium(II), complex with dichloromethane (16 mg, 0.02 mmol) in dioxane/H_2_O (v/v= 9:1, 0.5 mL) was stirred at 90 °C under nitrogen atmosphere. After the reaction was judged to be completed by TLC (3 h), it was cooled to room temperature, diluted with EtOAc, washed with water, and concentrated under reduced pressure by rotary evaporation. The crude residue was purified by flash column chromatography (SiO_2_, eluent gradient 0-40% EtOAc in hexanes) to afford compound **5**. Compound **5** (25 mg) was added to HCl/dioxane (4 M, 1 mL) and the mixture was stirred at 25 °C. After the reaction was judged to be completed by TLC (1 h), its solvent was evaporated (the residue was purified by flash column chromatography (SiO_2_) if necessary) to afford compound **6**.

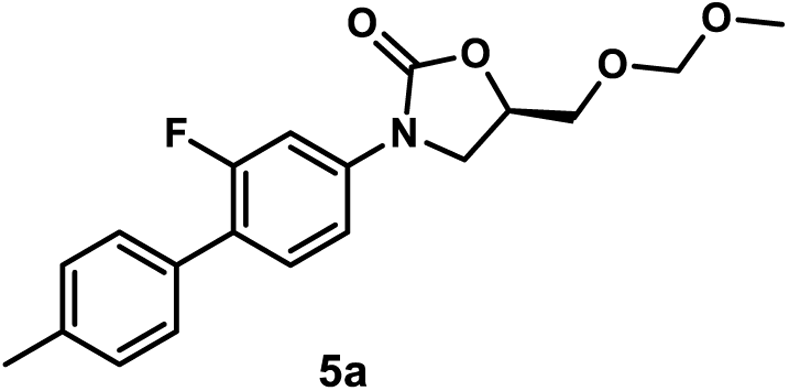

*(R)-3-(2-Fluoro-4’-methyl-[1,1’-biphenyl]-4-yl)-5-((methoxymethoxy)methyl)oxazolidin-2-one (**5a**)*. Using general procedure 2, employing 4-methylphenylboronic acid (33 mg, 0.24 mmol), compound **5a** was obtained as a yellow amorphous solid (68 mg, 98%). ^1^H NMR (300 MHz, Chloroform-*d*) δ 7.51 (dd, *J* = 12.8, 2.3 Hz, 1H), 7.47 – 7.38 (m, 3H), 7.34 (dd, *J* = 8.6, 2.3 Hz, 1H), 7.28 – 7.21 (m, 2H), 4.92 – 4.74 (m, 1H), 4.69 (s, 2H), 4.33 – 4.04 (m, 1H), 3.96 (dd, *J* = 8.7, 6.2 Hz, 1H), 3.85 (dd, *J* = 11.1, 4.2 Hz, 1H), 3.78 (dd, *J* = 11.1, 4.1 Hz, 1H), 3.39 (s, 3H), 2.40 (s, 3H).

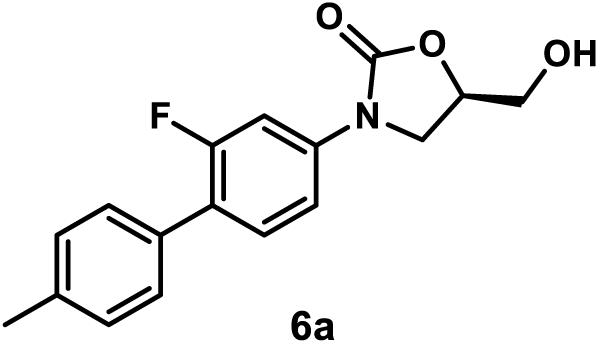

*(R)-3-(2-Fluoro-4’-methyl-[1,1’-biphenyl]-4-yl)-5-(hydroxymethyl)oxazolidin-2-one (**6a**).* Using general procedure 2, employing **5a** (25 mg, 0.072 mmol), compound **6a** was obtained (without flash column chromatography) as a yellow amorphous solid (22 mg, 100%). ^1^H NMR (300 MHz, Methanol-*d*_4_) δ 7.61 (dd, *J* = 13.4, 2.2 Hz, 1H), 7.50 – 7.32 (m, 4H), 7.25 (d, *J* = 7.9 Hz, 2H), 4.83 – 4.71 (m, 1H), 4.16 (t, *J* = 8.9 Hz, 1H), 3.96 (dd, *J* = 8.9, 6.4 Hz, 1H), 3.87 (dd, *J* = 12.5, 3.2 Hz, 1H), 3.71 (dd, *J* = 12.5, 4.0 Hz, 1H), 2.38 (s, 3H). ^13^C NMR (101 MHz, Methanol-*d*_4_) δ 161.0 (d, *J* = 245.2 Hz), 156.9, 140.3 (d, *J* = 11.0 Hz), 138.5, 133.7, 131.8 (d, *J* = 5.0 Hz), 130.2 (2C), 129.7 (d, *J* = 3.2 Hz, 2C), 125.5 (d, *J* = 13.8 Hz), 115.0 (d, *J* = 3.4 Hz), 107.2 (d, *J* = 29.3 Hz), 75.2, 63.3, 47.6, 21.2.

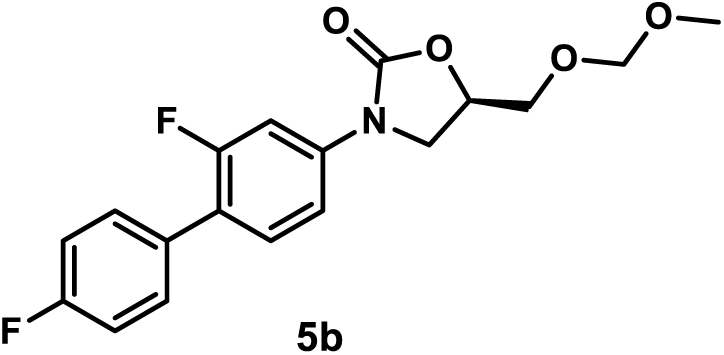

*(R)-3-(2,4’-Difluoro-[1,1’-biphenyl]-4-yl)-5-((methoxymethoxy)methyl)oxazolidin-2-one (***5b**). A mixture of compound **4** (50 mg, 0.15 mmol), 4-fluorophenylboronic acid (25 mg, 0.18 mmol), potassium carbonate (83 mg, 0.6 mmol), and [1,1’-bis(diphenylphosphino)ferrocene]dichloropalladium(II), complex with dichloromethane (12 mg, 0.015 mmol) in dioxane/H_2_O (v/v= 9:1, 0.4 mL) was stirred at 90 °C under nitrogen atmosphere for 2 h. The mixture was cooled to room temperature, diluted with EtOAc, washed with water, and concentrated under reduced pressure by rotary evaporation. The crude residue was purified by flash column chromatography (SiO_2_, eluent gradient 0-40% EtOAc in hexanes) to afford compound **5b** as a yellow amorphous solid (45 mg, 85%). ^1^H NMR (300 MHz, CDCl_3_) δ 7.63 – 7.32 (m, 5H), 7.21 – 7.08 (m, 2H), 4.93 – 4.76 (m, 1H), 4.69 (s, 2H), 4.10 (t, *J* = 8.8 Hz, 1H), 3.97 (dd, *J* = 8.8, 6.2 Hz, 1H), 3.91 – 3.73 (m, 2H), 3.39 (s, 3H).

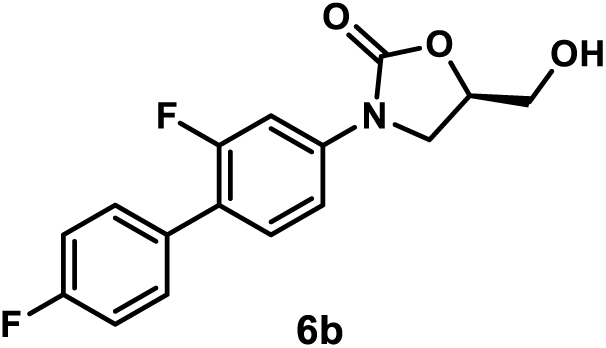

*(R)-3-(2,4’-Difluoro-[1,1’-biphenyl]-4-yl)-5-(hydroxymethyl)oxazolidin-2-one (**6b**)*. Using general procedure 2, employing **5b** (25 mg, 0.072 mmol), compound **6b** was obtained after flash column chromatography (SiO_2_, eluent gradient 0-100% EtOAc in hexanes) as a white amorphous solid (11 mg, 50%). ^1^H NMR (400 MHz, Acetone-*d*_6_) δ 7.70 (dd, *J* = 13.7, 2.2 Hz, 1H), 7.65 – 7.57 (m, 2H), 7.57 – 7.50 (m, 1H), 7.46 (dd, *J* = 8.6, 2.2 Hz, 1H), 7.35 – 7.11 (m, 2H), 4.93 – 4.76 (m, 1H), 4.52 – 4.31 (m, 1H), 4.23 (t, *J* = 9.0 Hz, 1H), 4.11 – 3.98 (m, 1H), 3.96 – 3.85 (m, 1H), 3.82 – 3.71 (m, 1H). ^13^C NMR (101 MHz, Acetone-*d*_6_) δ 163.2 (d, *J* = 245.2 Hz), 160.4 (d, *J* = 244.2 Hz), 155.3, 141.1 (d, *J* = 11.1 Hz), 132. 5 (dd, *J* = 3.0, 1.0 Hz), 131.6 (d, *J* = 3.8 Hz, 2C), 131.5 (d, *J* = 3.1 Hz), 123. 1 (d, *J* = 13.9 Hz), 116.2 (d, *J* = 21.6 Hz, 2C), 114.4 (d, *J* = 3.2 Hz), 106.3 (d, *J* = 29.1 Hz), 74.3, 63.2, 47.0.

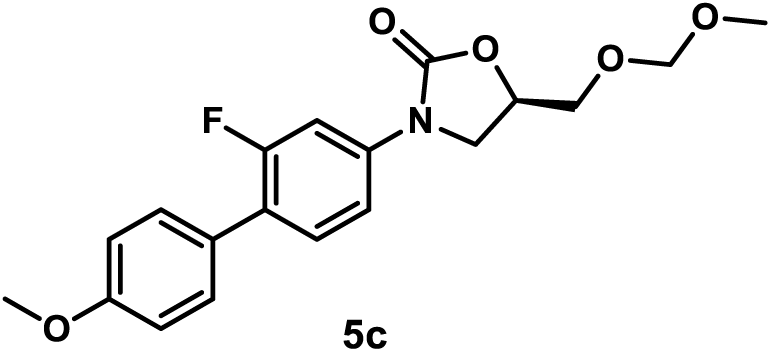

*(R)-3-(2-Fluoro-4’-methoxy-[1,1’-biphenyl]-4-yl)-5-((methoxymethoxy)methyl)oxazolidin-2-one (**5c**)*. Using general procedure 2, employing 4-methoxyphenylboronic acid (36 mg, 0.24 mmol), compound **5c** was obtained as a yellow amorphous solid (66 mg, 90%). ^1^H NMR (300 MHz, Chloroform-*d*) δ 7.57 – 7.30 (m, 5H), 6.98 (d, *J* = 8.7 Hz, 2H), 4.83 (ddt, *J* = 8.6, 6.2, 4.2 Hz, 1H), 4.69 (s, 2H), 4.10 (t, *J* = 8.6 Hz, 1H), 3.96 (dd, *J* = 8.6, 6.2 Hz, 1H), 3.93 – 3.63 (m, 5H), 3.39 (s, 3H).

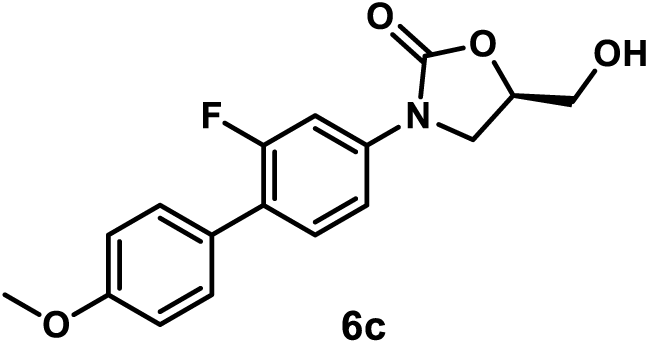

*(R)-3-(2-Fluoro-4’-methoxy-[1,1’-biphenyl]-4-yl)-5-(hydroxymethyl)oxazolidin-2-one (**6c**).* Using general procedure 2, employing **5c** (25 mg, 0.069 mmol), compound **6c** was obtained (without flash column chromatography) as a yellow amorphous solid (22 mg, 100%). ^1^H NMR (300 MHz, Acetone-*d*_6_) δ 7.67 (dd, *J* = 13.7, 2.3 Hz, 1H), 7.55 – 7.38 (m, 4H), 7.09 – 6.98 (m, 2H), 4.91 – 4.72 (m, 1H), 4.39 (dd, *J* = 6.2, 5.6 Hz, 1H), 4.22 (t, *J* = 8.9 Hz, 1H), 4.03 (dd, *J* = 8.9, 6.3 Hz, 1H), 3.95 – 3.87 (m, 1H), 3.84 (s, 3H), 3.83 – 3.73 (m, 1H). ^13^C NMR (101 MHz, Acetone-*d*_6_) δ 160.4 (d, *J* = 245.4 Hz), 160.3, 155.3, 140.4 (d, *J* = 11.1 Hz), 131.4 (d, *J* = 5.0 Hz), 130.7 (d, *J* = 3.1 Hz, 2C), 128.4 (d, *J* = 1.3 Hz), 124.0 (d, *J* = 13.7 Hz), 114.9 (2C), 114.4 (d, *J* = 3.3 Hz), 106.3 (d, *J* = 29.3 Hz), 74.3, 63.2, 55.6, 47.0. MSESI *m/z*: 340.0943 (C_17_H_16_FNO_4_ + Na^+^ requires 340.0956).

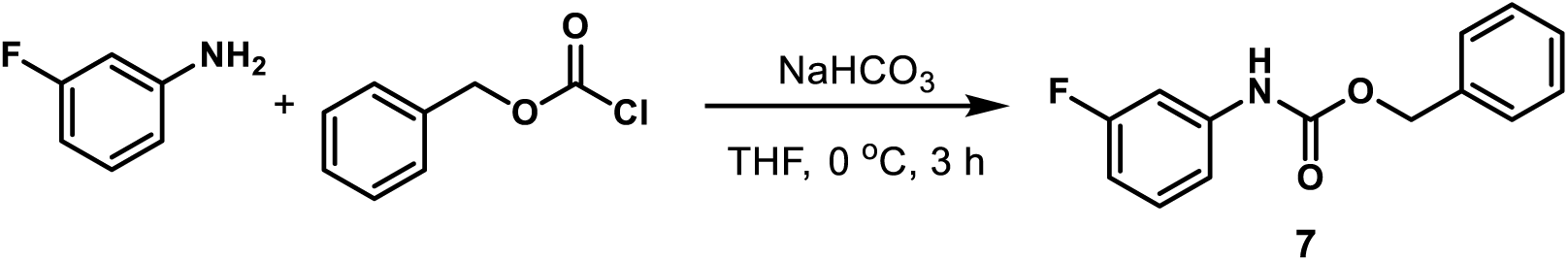

*Benzyl (3-Fluorophenyl)carbamate (**7**)*. A mixture of 3-fluoroaniline (1.92 mL, 20 mmol) and sodium bicarbonate (3.36 g, 40 mmol) was suspended in THF (80 mL) and cooled to 0 °C. Benzyl chloroformate (4.23 mL, 30 mmol) was added dropwise with a syringe. The reaction mixture was stirred at 0 °C. After the reaction was judged to be completed by TLC (3 h), it was quenched with water and extracted with EtOAc three times. The combined organic layers were washed with water and concentrated under reduced pressure by rotary evaporation. The crude residue was purified flash column chromatography (SiO_2_, eluent gradient 0-15% EtOAc in hexanes) to give compound **7** as a white amorphous solid (4.9 g, 99%). ^1^H NMR (300 MHz, Chloroform-*d*) δ 7.47 – 7.29 (m, 5H), 7.29 – 7.16 (m, 1H), 7.01 (ddd, *J* = 8.4, 2.1, 0.9 Hz, 1H), 6.83 – 6.66 (m, 2H), 5.21 (s, 2H).

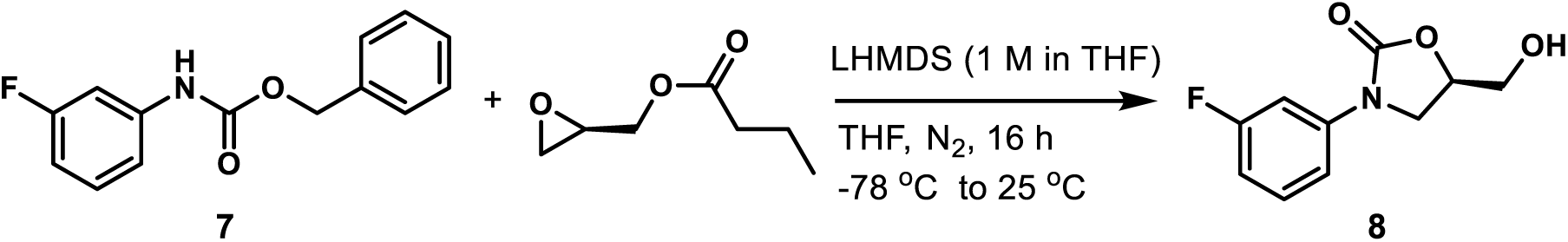

*(R)-3-(3-Fluorophenyl)-5-(hydroxymethyl)oxazolidin-2-one (**8**)*. Compound **7** (4.73 g, 19.27 mmol) was dissolved in anhydrous THF (100 mL) and cooled to -78 °C under nitrogen atmosphere. Lithium bis(trimethylsilyl)amide solution (1 M in THF, 16.38 mL, 16.38 mmol) was added to the mixture slowly over a period of 1 h with a syringe. The mixture was stirred at -78 °C for 1 h under nitrogen atmosphere followed by the addition of (*R*)-(-)-glycidyl butyrate (2.23 mL, 16.38 mmol) dropwise with a syringe at -78 °C. The mixture was stirred at this temperature for an additional 1 h and then gradually warmed to 25 °C. After the reaction was judged to be completed by TLC (16 h), it was quenched with water and evaporated under reduced pressure by rotary evaporation to remove most of THF. The mixture was diluted with water and extracted with EtOAc three times. The combined organic layers were washed with water and evaporated under reduced pressure by rotary evaporation. The crude reside was purified flash column chromatography (SiO_2_, eluent gradient 0-67% EtOAc in hexanes) to afford compound **8** as a white amorphous solid (2.86 g, 70%). ^1^H NMR (400 MHz, Acetone-*d*_6_) δ 7.67 – 7.51 (m, 1H), 7.47 – 7.26 (m, 2H), 6.96 – 6.76 (m, 1H), 4.87 – 4.70 (m, 1H), 4.39 (t, *J* = 5.7 Hz, 1H), 4.25 – 4.09 (m, 1H), 4.04 – 3.95 (m, 1H), 3.94 – 3.83 (m, 1H), 3.81 – 3.71 (m, 1H).

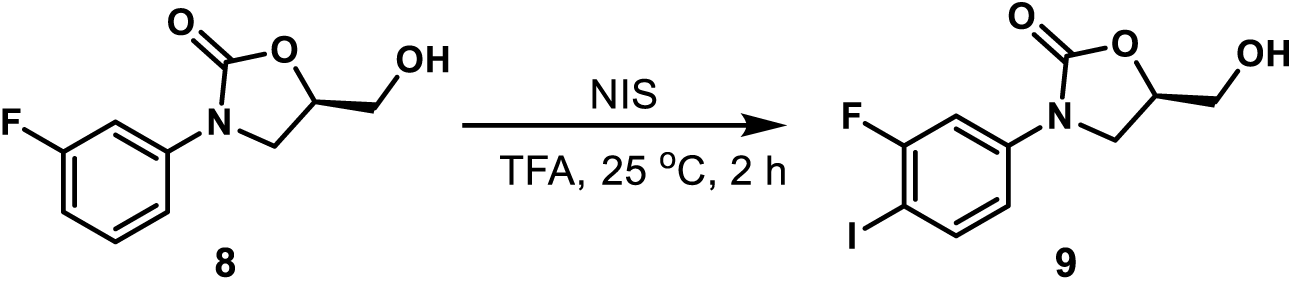

*(R)-3-(3-Fluoro-4-iodophenyl)-5-(hydroxymethyl)oxazolidin-2-one (**9**)*.^43^ *N*-iodosuccinimide (6.31 g, 28.0 mmol) was added to a solution of compound **8** (5.64 g, 26.7 mmol) in TFA (130 mL). The mixture was stirred at 25 °C. After the reaction was judged to be completed by TLC (2 h), it was concentrated under reduced pressure by rotary evaporation. The residue was dissolved in EtOAc and washed with saturated aqueous sodium carbonate until the aqueous phase was basic. The organic layer was evaporated under reduced pressure by rotary evaporation and the residue was purified by flash column chromatography (SiO_2_, eluent gradient 0-100% EtOAc in hexanes) to afford compound **9** as a gray amorphous solid (6.9 g, 77%). ^1^H NMR (400 MHz, Acetone-*d*_6_) δ 7.87 (dd, *J* = 8.8, 7.4 Hz, 1H), 7.67 (dd, *J* = 11.0, 2.5 Hz, 1H), 7.27 (ddd, *J* = 8.8, 2.5, 0.7 Hz, 1H), 4.80 (dddd, *J* = 9.3, 6.2, 4.1, 3.3 Hz, 1H), 4.13 (t, *J* = 9.3 Hz, 1H), 4.03 – 3.84 (m, 2H), 3.80 – 3.70 (m, 1H), 3.33 (t, *J* = 5.9 Hz, 1H).

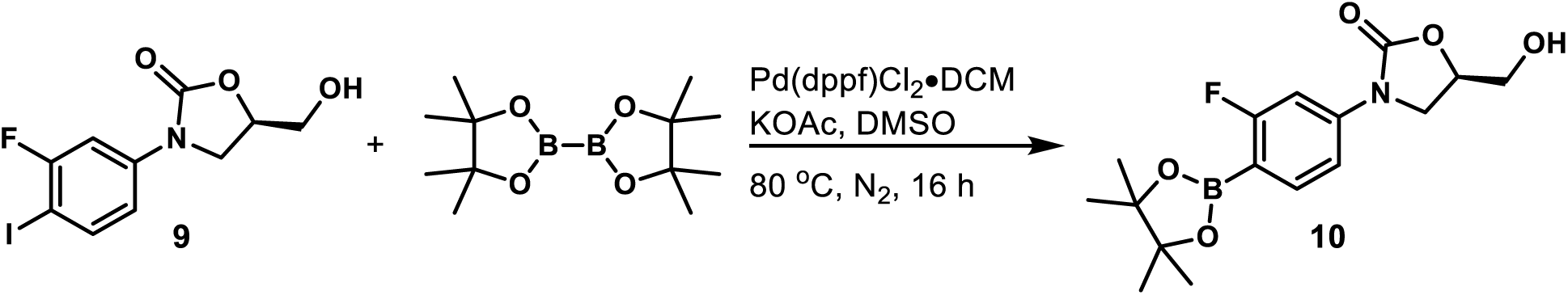

*(R)-3-(3-Fluoro-4-(4,4,5,5-tetramethyl-1,3,2-dioxaborolan-2-yl)phenyl)-5-(hydroxymethyl)oxazolidin-2-one (**10**)*.^43^ To a solution of compound **9** (3.37 g, 10 mmol) in anhydrous DMSO (40 mL), [1,1’-bis(diphenylphosphino)ferrocene]dichloropalladium(II), complex with dichloromethane (408 mg, 0.5 mmol), potassium acetate (4.91 g, 50 mmol), and bis(pinacolato)diboron (5.08 g, 20 mmol) were added. The reaction mixture was stirred at 80 °C under nitrogen atmosphere. After the reaction was judged to be completed by TLC (16 h), it was cooled to room temperature, treated with the addition of MeOH (20 mL), and filtered through Celite. The filtrate was concentrated under reduced pressure by rotary evaporation to remove MeOH, dissolved in EtOAc, washed with water, and evaporated under reduced pressure by rotary evaporation. The crude residue was purified by flash column chromatography (SiO_2_, eluent gradient 0-100% EtOAc in hexanes) to afford compound **10** as a yellow amorphous solid (1.64 g, 49%). ^1^H NMR (400 MHz, Acetone-*d*_6_) δ 7.75 – 7.67 (m, 1H), 7.55 (dd, *J* = 12.2, 2.0 Hz, 1H), 7.37 (dd, *J* = 8.3, 2.0 Hz, 1H), 4.91 – 4.74 (m, 1H), 4.40 (t, *J* = 5.9 Hz, 1H), 4.20 (t, *J* = 8.9 Hz, 1H), 4.01 (dd, *J* = 8.9, 6.2 Hz, 1H), 3.90 (ddd, *J* = 12.3, 5.9, 3.3 Hz, 1H), 3.76 (ddd, *J* = 12.3, 5.9, 4.0 Hz, 1H), 1.33 (s, 12H).

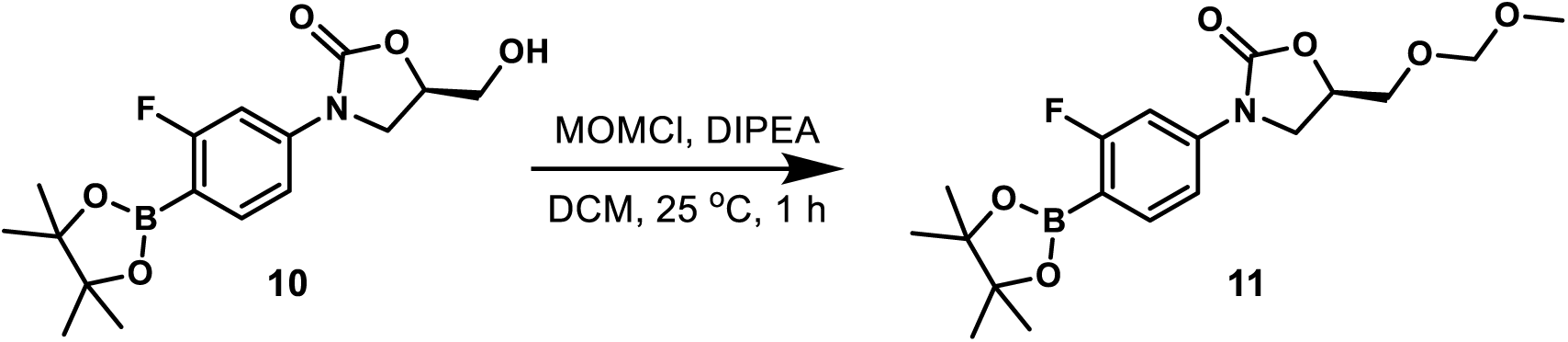

*(R)-3-(3-Fluoro-4-(4,4,5,5-tetramethyl-1,3,2-dioxaborolan-2-yl)phenyl)-5-((methoxymethoxy)methyl)oxazolidin-2-one (**11**)*. To a solution of compound **10** (1.44 g, 4.27 mmol) in DCM (30 mL), *N*,*N*-diisopropylethylamine (2.23 mL, 12.81 mmol) and methyl chloromethyl ether (0.97 mL, 12.81 mmol) were added. The reaction mixture was stirred at 25 °C. After the reaction was judged to be completed by TLC (1 h), it was diluted with EtOAc, washed with water, and concentrated under reduced pressure by rotary evaporation. The crude residue was purified by flash column chromatography (SiO_2_, eluent gradient 0-66% EtOAc in hexanes) to afford compound **11** as a white amorphous solid (940 mg, 58%). ^1^H NMR (400 MHz, Chloroform-*d*) δ 7.72 (dd, *J* = 8.3, 6.9 Hz, 1H), 7.39 (dd, *J* = 11.7, 2.1 Hz, 1H), 7.28 (dd, *J* = 8.3, 2.1 Hz, 1H), 4.88 – 4.75 (m, 1H), 4.67 (s, 2H), 4.07 (t, *J* = 8.8 Hz, 1H), 3.93 (dd, *J* = 8.8, 6.3 Hz, 1H), 3.83 (dd, *J* = 11.2, 4.2 Hz, 1H), 3.76 (dd, *J* = 11.2, 4.1 Hz, 1H), 3.37 (s, 3H), 1.35 (s, 12H).

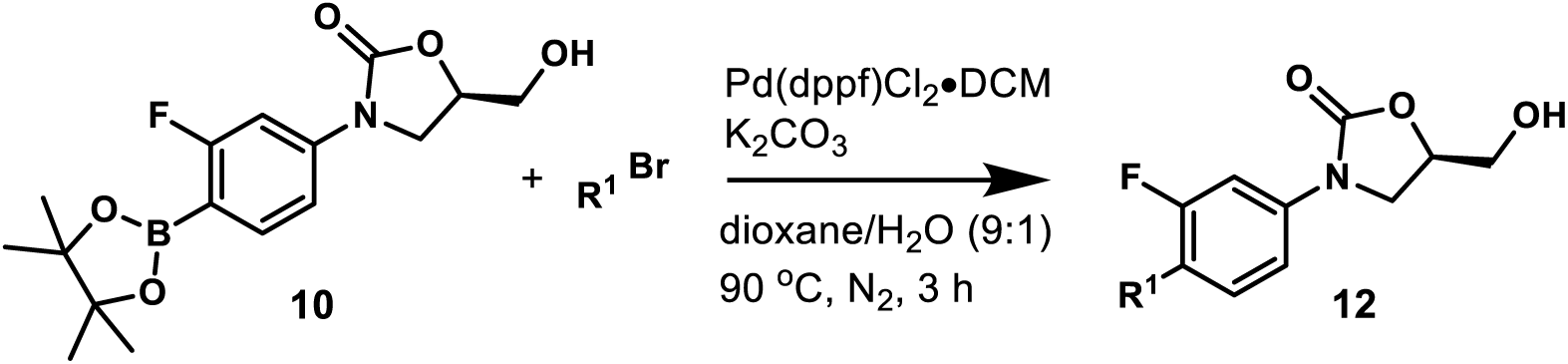

#### General Procedure 3: Synthesis of Heterocyclic Oxazolidinone Analogs 12a-12p

A mixture of compound **10** (34 mg, 0.1 mmol), heterocyclic bromide, potassium carbonate (55 mg, 0.4 mmol), and [1,1’-bis(diphenylphosphino)ferrocene]dichloropalladium(II), complex with dichloromethane (8.2 mg, 0.01 mmol) in dioxane/H_2_O (v/v= 9:1, 0.5 mL) was stirred at 90 °C under nitrogen atmosphere for 3 h. The reaction mixture was cooled to room temperature, diluted with EtOAc, washed with water, and concentrated under reduced pressure by rotary evaporation. The crude residue was purified by flash column chromatography (SiO_2_, eluent gradient 0-10% MeOH in DCM) to afford compound **12**. If not pure, compound **12** was further purified by washing with EtOAc (1 mL).

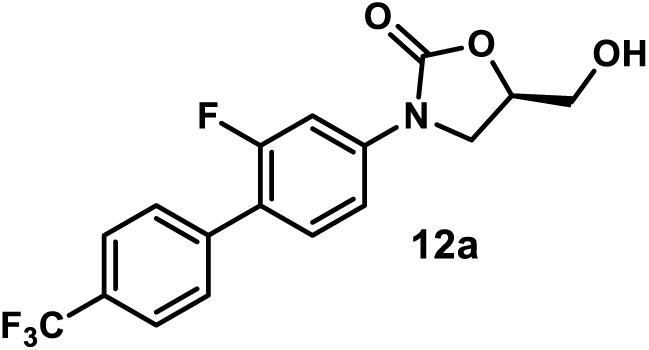

*(R)-3-(2-Fluoro-4’-(trifluoromethyl)-[1,1’-biphenyl]-4-yl)-5-(hydroxymethyl)oxazolidin-2-one (**12a**)*. Using general procedure 3, employing 1-bromo-4-(trifluoromethyl)benzene (45 mg, 0.2 mmol), compound **12a** was obtained by flash column chromatography (SiO_2_, eluent gradient 0-100% EtOAc in hexanes) as a white amorphous solid (16 mg, 45%). ^1^H NMR (400 MHz, Acetone-*d*_6_) δ 7.81 (s, 4H), 7.75 (dd, *J* = 13.8, 2.2 Hz, 1H), 7.62 (t, *J* = 8.6 Hz, 1H), 7.51 (dd, *J* = 8.6, 2.3 Hz, 1H), 4.91 – 4.75 (m, 1H), 4.41 (s, 1H), 4.25 (t, *J* = 8.9 Hz, 1H), 4.07 (dd, *J* = 8.9, 6.2 Hz, 1H), 3.95 – 3.88 (m, 1H), 3.84 – 3.74 (m, 1H).^13^C NMR (101 MHz, Acetone-*d*_6_) δ 160.6 (d, *J* = 245.5 Hz), 155.3, 141.8 (d, *J* = 11.4 Hz), 140.3, 131.8 (d, *J* = 4.5 Hz), 130.3 (d, *J* = 3.4 Hz, 2C), 129.7 (d, *J* = 32.2 Hz), 126.3 (q, *J* = 3.9 Hz, 2C), 125.2 (q, *J* = 253.6 Hz), 122.5 (d, *J* = 13.4 Hz), 114.5 (d, *J* = 3.3 Hz), 106.3 (d, *J* = 29.1 Hz), 74.4, 63.2, 47.0. MSESI *m/z*: 356.0900 (C_17_H_13_F_4_NO_3_ + H^+^ requires 356.0904).

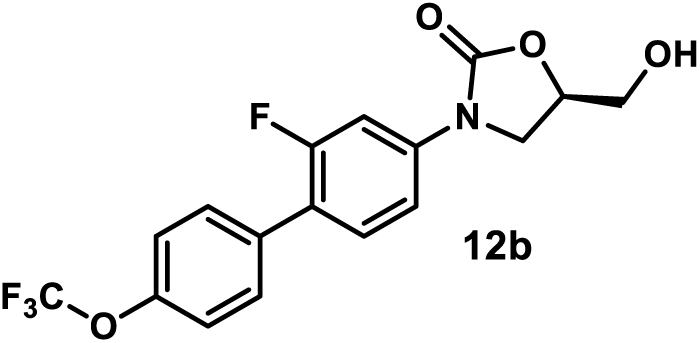

*(R)-3-(2-Fluoro-4’-(trifluoromethoxy)-[1,1’-biphenyl]-4-yl)-5-(hydroxymethyl)oxazolidin-2-one (**12b**)*. Using general procedure 3, employing 1-bromo-4-(trifluoromethoxy)benzene (48 mg, 0.2 mmol), compound **12b** was obtained by flash column chromatography (SiO_2,_ eluent gradient 0-100% EtOAc in hexanes) as a white amorphous solid (17 mg, 46%). ^1^H NMR (400 MHz, Acetone-*d*_6_) δ 7.78 – 7.65 (m, 3H), 7.57 (t, *J* = 8.7 Hz, 1H), 7.48 (dd, *J* = 8.7, 2.3 Hz, 1H), 7.46 – 7.40 (m, 2H), 4.93 – 4.73 (m, 1H), 4.40 (s, 1H), 4.24 (t, *J* = 8.9 Hz, 1H), 4.06 (dd, *J* = 8.9, 6.2 Hz, 1H), 3.95 – 3.87 (m, 1H), 3.81 – 3.75 (m, 1H). ^13^C NMR (101 MHz, Acetone-*d*_6_) δ 160.6 (d, *J* = 245.0 Hz), 155.4, 149.4 (d, *J* = 1.9 Hz), 141.5 (d, *J* = 11.2 Hz), 135.6 (d, *J* = 1.6 Hz), 131.8 (d, *J* = 4.6 Hz), 131.5 (d, *J* = 3.3 Hz, 2C), 122.7 (d, *J* = 13.7 Hz), 122.1 (2C), 121.6 (q, *J* = 255.6 Hz), 114.6 (d, *J* = 3.2 Hz), 106.4 (d, *J* = 29.0 Hz), 74.5, 63.3, 47.1. MSEI *m/z*: 371.0775 (C_17_H_13_F_4_NO_3_ (M^+^) requires 371.0781).

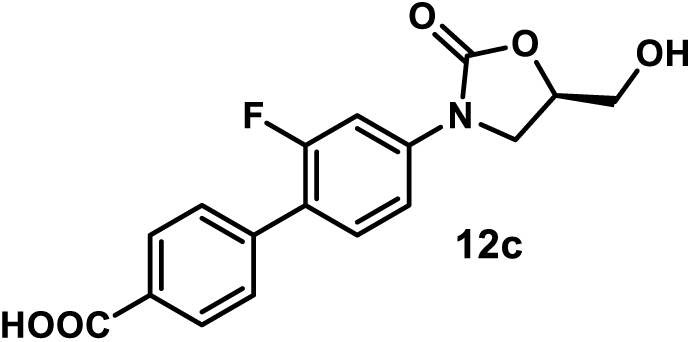

*(R)-2’-Fluoro-4’-(5-(hydroxymethyl)-2-oxooxazolidin-3-yl)-[1,1’-biphenyl]-4-carboxylic Acid (**12c**)*. A mixture of compound **10** (40 mg, 0.12 mmol), 4-bromobenzoic acid (20 mg, 0.1 mmol), potassium carbonate (55 mg, 0.4 mmol), and [1,1’-bis(diphenylphosphino)ferrocene]dichloropalladium(II), complex with dichloromethane (8.2 mg, 0.01 mmol) in dioxane/H_2_O (v/v= 9:1, 0.5 mL) was stirred at 90 °C under nitrogen atmosphere. After the reaction was judged to be completed by TLC (3 h), formic acid (1 mL) was added to acidify the reaction. The mixture was concentrated under reduced pressure by rotary evaporation. The crude residue was purified by flash column chromatography (SiO_2_, eluent gradient 0-15% MeOH in DCM with 1% formic acid). The appropriate fractions were concentrated under reduced pressure by rotary evaporation and the resulting residue was washed with MeOH (2 mL) to give compound **12c** as a white amorphous solid (10 mg, 30%). ^1^H NMR (400 MHz, DMSO-*d*_6_) δ 8.03 (d, *J* = 8.4 Hz, 2H), 7.74 – 7.59 (m, 4H), 7.49 (dd, *J* = 8.6, 2.3 Hz, 1H), 5.27 (s, 1H), 4.86 – 4.68 (m, 1H), 4.14 (t, *J* = 8.9 Hz, 1H), 3.89 (dd, *J* = 8.9, 6.1 Hz, 1H), 3.70 (dd, *J* = 12.4, 3.3 Hz, 1H), 3.57 (dd, *J* = 12.4, 3.9 Hz, 1H). ^13^C NMR (101 MHz, DMSO-*d*_6_) δ 167.1, 159.1 (d, *J* = 245.4 Hz), 154.4, 140.1 (d, *J* = 11.2 Hz), 139.0, 131.0 (d, *J* = 4.5 Hz), 129.8, 129.7 (2C), 128.8 (d, *J* = 3.3 Hz, 2C), 121.6 (d, *J* = 13.0 Hz), 113.9 (d, *J* = 3.0 Hz), 105.4 (d, *J* = 28.6 Hz), 73.5, 61.6, 46.0. MSESI *m/z*: 330.0766 (C_17_H_13_FNO_5_^-^requires 330.0783).

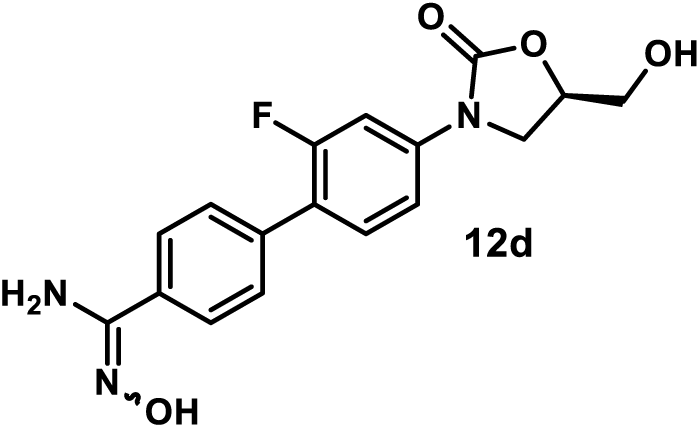

*(R)-2’-Fluoro-N’-hydroxy-4’-(5-(hydroxymethyl)-2-oxooxazolidin-3-yl)-[1,1’-biphenyl]-4-carboximidamide (**12d**)*. A mixture of compound **10** (40 mg, 0.12 mmol), compound **18** (22 mg, 0.1 mmol), potassium carbonate (55 mg, 0.4 mmol), and [1,1’-bis(diphenylphosphino)ferrocene]dichloropalladium(II), complex with dichloromethane (8.2 mg, 0.01 mmol) in dioxane/H_2_O (v/v= 9:1, 0.5 mL) was stirred at 90 °C under a nitrogen atmosphere. After the reaction was judged to be completed by TLC (3 h), it was concentrated under reduced pressure by rotary evaporation. The crude residue was purified by flash column chromatography (SiO_2_, eluent gradient 0-20% MeOH in DCM) and preparative TLC (eluent, 15% MeOH in DCM) to give compound **12d** as a yellow amorphous solid (17 mg, 49%). ^1^H NMR (300 MHz, DMSO-*d*_6_) δ 9.71 (s, 1H), 7.85 – 7.72 (m, 2H), 7.68 – 7.52 (m, 4H), 7.46 (dd, *J* = 8.6, 2.2 Hz, 1H), 5.87 (s, 2H), 5.25 (t, *J* = 5.6 Hz, 1H), 4.80 – 4.68 (m, 1H), 4.14 (t, *J* = 9.0 Hz, 1H), 3.88 (dd, *J* = 9.0, 6.1 Hz, 1H), 3.70 (ddd, *J* = 12.3, 5.6, 3.3 Hz, 1H), 3.57 (ddd, *J* = 12.3, 5.6, 4.0 Hz, 1H). ^13^C NMR (101 MHz, DMSO-*d*_6_) δ 159.1 (d, *J* = 244.8 Hz), 154.4, 150.5, 139.6 (d, *J* = 11.2 Hz), 135.1 (d, *J* = 1.7 Hz), 132.6, 130.8 (d, *J* = 5.0 Hz), 128.3 (d, *J* = 3.3 Hz, 2C), 125.6 (2C), 122.1 (d, *J* = 13.3 Hz), 113.8 (d, *J* = 3.1 Hz), 105.4 (d, *J* = 28.8 Hz), 73.4, 61.6, 46.0. MSESI *m/z*: 346.1187 (C_17_H_16_FN_3_O_4_ + H^+^ requires 346.1198).

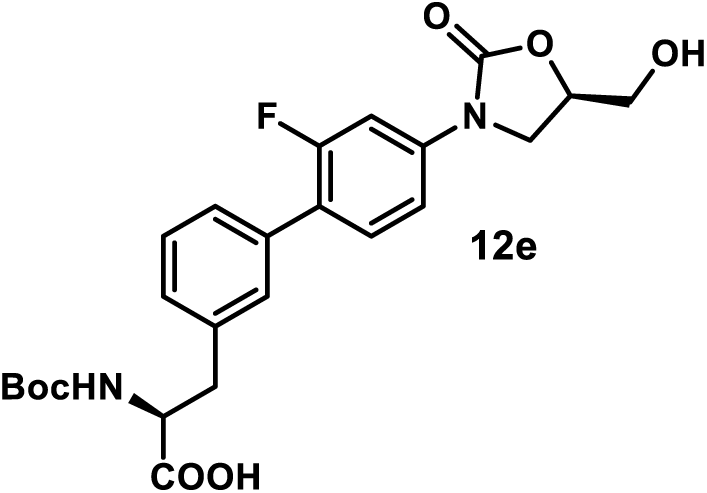

*(S)-2-((tert-Butoxycarbonyl)amino)-3-(2’-fluoro-4’-((R)-5-(hydroxymethyl)-2-oxooxazolidin-3-yl)-[1,1’-biphenyl]-3-yl)propanoic Acid (**12e**)*. A mixture of compound **10** (80 mg, 0.24 mmol), (*S*)-*N*-Boc-3-bromophenylalanine (69 mg, 0.2 mmol), potassium carbonate (110 mg, 0.8 mmol), and [1,1’-bis(diphenylphosphino)ferrocene]dichloropalladium(II), complex with dichloromethane (16 mg, 0.02 mmol) in dioxane/H_2_O (v/v= 9:1, 1.0 mL) was stirred at 90 °C under a nitrogen atmosphere. After the reaction was judged to be completed by TLC (3 h), it was cooled to room temperature, diluted with EtOAc, washed with water, and concentrated under reduced pressure by rotary evaporation. The crude residue was purified by flash column chromatography (SiO_2_, eluent gradient 0-10% MeOH in DCM with 1% formic acid) to afford compound **12e** as a yellow amorphous solid (34 mg, 36%). ^1^H NMR (400 MHz, Acetone-*d*_6_) δ 7.70 (dd, *J* = 13.6, 2.3 Hz, 1H), 7.55 (t, *J* = 8.7 Hz, 1H), 7.50 – 7.42 (m, 3H), 7.39 (t, *J* = 7.5 Hz, 1H), 7.30 (d, *J* = 7.5 Hz, 1H), 6.11 (d, *J* = 8.4 Hz, 1H), 4.95 – 4.71 (m, 1H), 4.57 – 4.42 (m, 1H), 4.41 – 4.35 (m, 1H), 4.22 (t, *J* = 8.9 Hz, 1H), 4.04 (dd, *J* = 8.9, 6.2 Hz, 1H), 3.91 (dd, *J* = 12.3, 3.4 Hz, 1H), 3.78 (dd, *J* = 12.3, 3.9 Hz, 1H), 3.29 (dd, *J* = 13.8, 4.9 Hz, 1H), 3.08 (dd, *J* = 13.8, 8.8 Hz, 1H), 1.34 (s, 9H). ^13^C NMR (101 MHz, Acetone-*d*_6_) δ 173.8, 160.5 (d, *J* = 245.0 Hz), 156.2, 155.3, 140.8 (d, *J* = 11.2 Hz), 138.9, 136.1, 131.7 (d, *J* = 5.0 Hz), 130.6 (d, *J* = 2.7 Hz), 129.4, 129.3, 127.9 (d, *J* = 3.6 Hz), 124.1 (d, *J* = 13.5 Hz), 114.3 (d, *J* = 3.4 Hz), 106.2 (d, *J* = 29.1 Hz), 79.2, 74.3, 63.1, 55.7, 46.9, 38.1, 28.5.

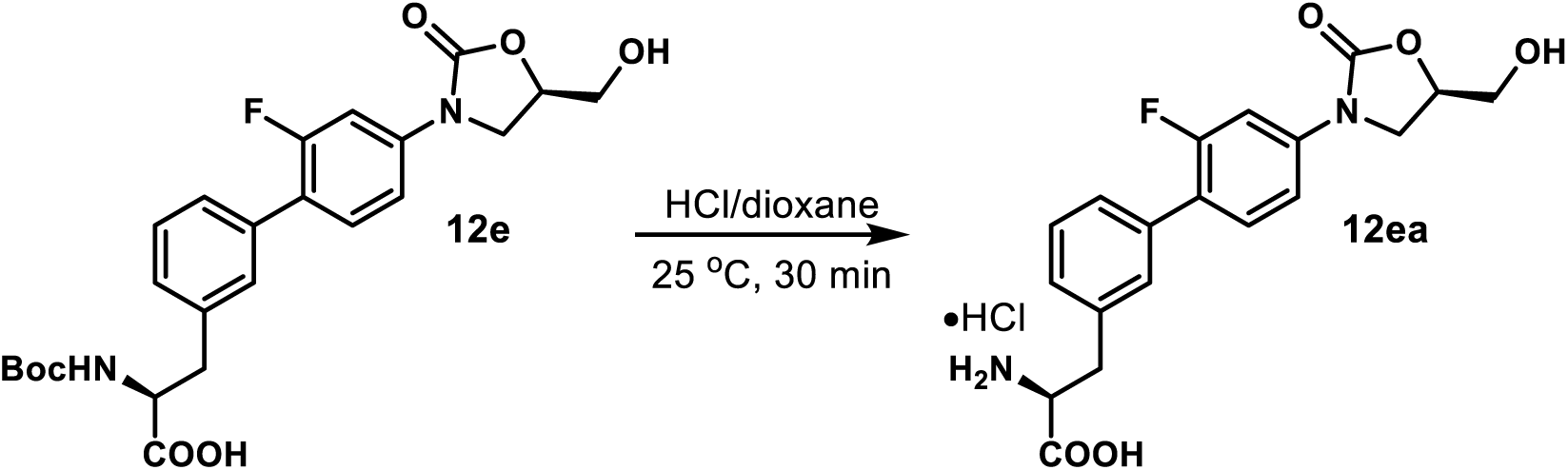

*(S)-2-Amino-3-(2’-fluoro-4’-((R)-5-(hydroxymethyl)-2-oxooxazolidin-3-yl)-[1,1’-biphenyl]-3-yl)propanoic Acid Hydrochloride(**12ea**)*. A mixture of compound **12e** (20 mg, 0.042 mmol) in HCl/dioxane (4 M, 1 mL) was stirred at 25 °C. After the reaction was judged to be completed by TLC (30 min), its solvent was removed under reduced pressure by rotary evaporation. The resulting residue was washed with acetone (1 mL) and dried under high vacuum to give **12ea** as a yellow amorphous solid (16 mg, 91%). ^1^H NMR (400 MHz, DMSO-*d*_6_) δ 8.51 (s, 3H), 7.69 – 7.56 (m, 2H), 7.53 – 7.38 (m, 4H), 7.30 (d, *J* = 7.4 Hz, 1H), 5.33 (s, 1H), 4.81 – 4.67 (m, 1H), 4.22 (t, *J* = 6.1 Hz, 1H), 4.13 (t, *J* = 8.9 Hz, 1H), 3.91 (dd, *J* = 8.9, 6.0 Hz, 1H), 3.69 (dd, *J* = 12.3, 3.3 Hz, 1H), 3.62 – 3.52 (m, 1H), 3.21 (d, *J* = 6.1 Hz, 2H). ^13^C NMR (101 MHz, DMSO-*d*_6_) δ 170.4, 159.0 (d, *J* = 244.9 Hz), 154.4, 139.5 (d, *J* = 11.1 Hz), 135.5, 134.9, 131.0 (d, *J* = 4.8 Hz), 129.8, 128.9, 128.8, 127.6 (d, *J* = 3.2 Hz), 122.5 (d, *J* = 13.1 Hz), 113.8 (d, *J* = 2.5 Hz), 105.4 (d, *J* = 28.8 Hz), 73.4, 61.6, 53.1, 46.0, 35.6. MSESI *m/z*: 375.1360 (C_19_H_19_FN_2_O_5_ + H^+^ requires 375.1351).

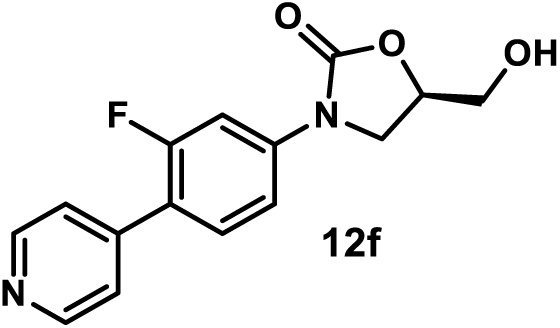

*(R)-3-(3-Fluoro-4-(pyridin-4-yl)phenyl)-5-(hydroxymethyl)oxazolidin-2-one (**12f**)*. Using general procedure 3, employing 4-bromopyridine hydrochloride (25 mg, 0.13 mmol), compound **12f** was obtained as a brown amorphous solid (21 mg, 73%). ^1^H NMR (400 MHz, DMSO-*d*_6_) δ 8.65 (d, *J* = 5.0 Hz, 2H), 7.78 – 7.64 (m, 2H), 7.59 (d, *J* = 5.0 Hz, 2H), 7.54 – 7.47 (m, 1H), 5.27 (t, *J* = 5.5 Hz, 1H), 4.82 – 4.70 (m, 1H), 4.14 (t, *J* = 8.9 Hz, 1H), 3.89 (dd, *J* = 8.9, 6.1 Hz, 1H), 3.74 – 3.65 (m, 1H), 3.63 – 3.52 (m, 1H). ^13^C NMR (101 MHz, DMSO-*d*_6_) δ 159.4 (d, *J* = 247.0 Hz), 154.4, 150.0 (2C), 142.1, 140.9 (d, *J* = 11.4 Hz), 130.8 (d, *J* = 4.5 Hz), 123.1 (d, *J* = 3.8 Hz, 2C), 119.7 (d, *J* = 12.4 Hz), 113.9 (d, *J* = 3.0 Hz), 105.4 (d, *J* = 28.6 Hz), 73.5, 61.6, 46.0. MSESI *m/z*: 289.0965 (C_15_H_13_FN_2_O_3_ + H^+^ requires 289.0983).

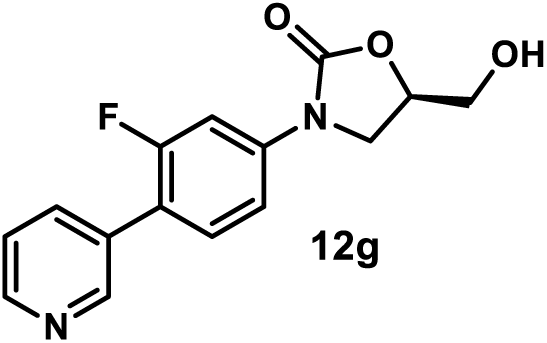

*(R)-3-(3-Fluoro-4-(pyridin-3-yl)phenyl)-5-(hydroxymethyl)oxazolidin-2-one (**12g**).* Using general procedure 3, employing 3-bromopyridine (12.5 µL, 0.13 mmol), compound **12g** was obtained as a white amorphous solid (20 mg, 70%). ^1^H NMR (300 MHz, DMSO-*d*_6_) δ 8.76 (s, 1H), 8.63 – 8.53 (m, 1H), 7.97 (d, *J* = 7.9 Hz, 1H), 7.75 – 7.60 (m, 2H), 7.55 – 7.43 (m, 2H), 5.28 (s, 1H), 4.84 – 4.69 (m, 1H), 4.14 (t, *J* = 8.9 Hz, 1H), 3.89 (dd, *J* = 8.9, 6.1 Hz, 1H), 3.79 – 3.64 (m, 1H), 3.63 – 3.52 (m, 1H). ^13^C NMR (101 MHz, DMSO-*d*_6_) δ 159.2 (d, *J* = 245.0 Hz), 154.4, 149.0 (d, *J* = 3.7 Hz), 148.6, 140.2 (d, *J* = 11.3 Hz), 136.0 (d, *J* = 3.2 Hz), 131.0 (d, *J* = 4.5 Hz), 130.6, 123.7, 119.4 (d, *J* = 13.7 Hz), 114.0 (d, *J* = 3.1 Hz), 105.3 (d, *J* = 28.5 Hz), 73.5, 61.6, 46.0. MSESI *m/z*: 289.0965 (C_15_H_13_FN_2_O_3_ + H^+^ requires 289.0983).

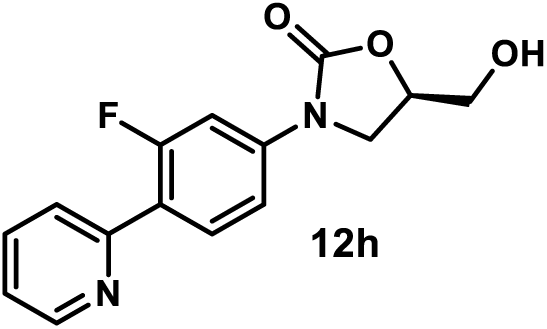

*(R)-3-(3-Fluoro-4-(pyridin-2-yl)phenyl)-5-(hydroxymethyl)oxazolidin-2-one (**12h**)*. Using general procedure 3, employing 2-bromopyridine (13.1 µL, 0.13 mmol), compound **12h** was obtained as a yellow amorphous solid (20 mg, 70%). ^1^H NMR (400 MHz, DMSO-*d*_6_) δ 8.70 (dd, *J* = 5.1, 1.8 Hz, 1H), 8.01 (t, *J* = 8.9 Hz, 1H), 7.96 – 7.85 (m, 1H), 7.83 – 7.75 (m, 1H), 7.65 (dd, *J* = 14.4, 2.2 Hz, 1H), 7.49 (dd, *J* = 8.7, 2.2 Hz, 1H), 7.46 – 7.35 (m, 1H), 5.28 (s, 1H), 4.84 – 4.67 (m, 1H), 4.15 (t, *J* = 8.9 Hz, 1H), 3.89 (dd, *J* = 8.9, 6.1 Hz, 1H), 3.75 – 3.65 (m, 1H), 3.62 – 3.54 (m, 1H). ^13^C NMR (101 MHz, DMSO-*d*_6_) δ 159.9 (d, *J* = 246.9 Hz), 154.4, 152.0 (d, *J* = 2.9 Hz), 149.8, 140.6 (d, *J* = 11.4 Hz), 137.0, 131.1 (d, *J* = 4.5 Hz), 123.8 (d, *J* = 9.5 Hz), 122.7, 121.3 (d, *J* = 11.8 Hz), 113.6 (d, *J* = 2.9 Hz), 105.2 (d, *J* = 29.2 Hz), 73.5, 61.6, 46.0. MSESI *m/z*: 289.0980 (C_15_H_13_FN_2_O_3_ + H^+^ requires 289.0983).

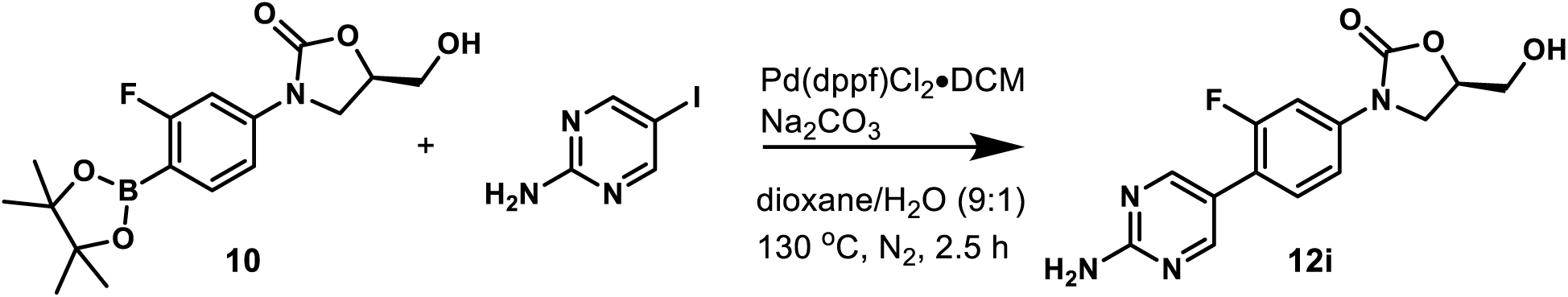

*(R)-3-(4-(2-Aminopyrimidin-5-yl)-3-fluorophenyl)-5-(hydroxymethyl)oxazolidin-2-one (**12i**)*. A mixture of compound **10** (34 mg, 0.1 mmol), 2-amino-5-iodopyrimidine (33 mg, 0.15 mmol), sodium carbonate (23 mg, 0.2 mmol), and [1,1’-bis(diphenylphosphino)ferrocene]dichloropalladium(II), complex with dichloromethane (8 mg, 0.01 mmol) in dioxane/H_2_O (v/v= 9:1, 0.5 mL) was stirred at 130 °C under a nitrogen atmosphere. After the reaction was judged to be completed by TLC (2.5 h), it was cooled to room temperature and concentrated under reduced pressure by rotary evaporation. The resulting residue was purified by flash column chromatography (SiO_2_, eluent gradient 0-10% MeOH in DCM) to afford compound **12i** as a gray amorphous solid (6 mg, 20%). ^1^H NMR (400 MHz, DMSO-*d*_6_) δ 8.47 – 8.33 (m, 2H), 7.62 (dd, *J* = 13.6, 2.3 Hz, 1H), 7.57 (t, *J* = 8.6 Hz, 1H), 7.41 (dd, *J* = 8.6, 2.3 Hz, 1H), 6.87 (s, 2H), 5.26 (s, 1H), 4.81 – 4.64 (m, 1H), 4.12 (t, *J* = 8.9 Hz, 1H), 3.86 (dd, *J* = 8.9, 6.1 Hz, 1H), 3.73 – 3.65 (m, 1H), 3.61 – 3.51 (m, 1H). ^13^C NMR (101 MHz, DMSO-*d*_6_) δ 162.7, 158.9 (d, *J* = 243.5 Hz), 157.3 (d, *J* = 3.9 Hz, 2C), 154.4, 139.1 (d, *J* = 11.1 Hz), 129.7 (d, *J* = 4.9 Hz), 117.7 (d, *J* = 14.2 Hz), 117.0 (d, *J* = 2.2 Hz), 113.9 (d, *J* = 3.1 Hz), 105.4 (d, *J* = 28.5 Hz), 73.4, 61.6, 46.0. MSESI *m/z*: 305.1038 (C_14_H_13_FN_4_O_3_ + H^+^ requires 305.1044).

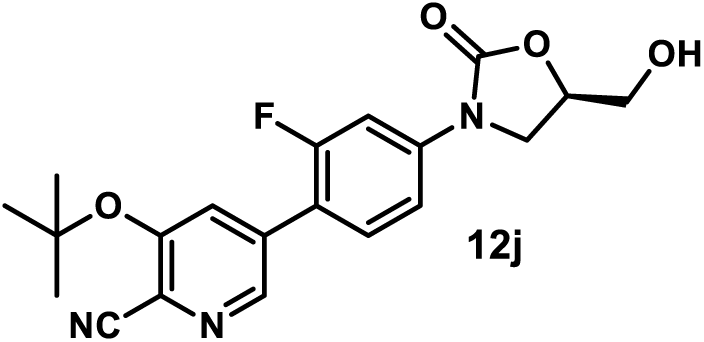

*(R)-3-(tert-Butoxy)-5-(2-fluoro-4-(5-(hydroxymethyl)-2-oxooxazolidin-3-yl)phenyl)picolinonitrile (**12j**)*. A mixture of compound **10** (121 mg, 0.36 mmol), compound **19** (77 mg, 0.3 mmol), potassium carbonate (166 mg, 1.2 mmol), and [1,1’-bis(diphenylphosphino)ferrocene]dichloropalladium(II), complex with dichloromethane (24 mg, 0.03 mmol) in dioxane/H_2_O (v/v= 9:1, 1.5 mL) was stirred at 90 °C under a nitrogen atmosphere. After the reaction was judged to be completed by TLC (3 h), it was cooled to room temperature, diluted with EtOAc, washed with water, and concentrated under reduced pressure by rotary evaporation. The crude residue was purified by flash column chromatography (SiO_2_, eluent gradient 0-100% EtOAc in hexanes) to afford compound **12j** as a yellow amorphous solid (46 mg, 61%). ^1^H NMR (400 MHz, Acetone-*d*_6_) δ 8.59 (d, *J* = 1.6 Hz, 1H), 7.94 (d, *J* = 1.6 Hz, 1H), 7.78 (dd, *J* = 13.9, 2.2 Hz, 1H), 7.71 (t, *J* = 8.8 Hz, 1H), 7.54 (dd, *J* = 8.8, 2.2 Hz, 1H), 4.95 – 4.76 (m, 1H), 4.42 (t, *J* = 5.8 Hz, 1H), 4.26 (t, *J* = 8.8 Hz, 1H), 4.08 (dd, *J* = 8.8, 6.1 Hz, 1H), 3.92 (ddd, *J* = 12.3, 5.8, 3.3 Hz, 1H), 3.78 (ddd, *J* = 12.3, 5.8, 3.7 Hz, 1H), 1.55 (s, 9H). ^13^C NMR (101 MHz, Acetone-*d*_6_) δ 160.8 (d, *J* = 246.4 Hz), 156.4, 155.2, 145.3 (d, *J* = 3.9 Hz), 142.7 (d, *J* = 11.4 Hz), 136.2 (d, *J* = 2.2 Hz), 131.9 (d, *J* = 4.2 Hz), 130.4 (d, *J* = 4.0 Hz), 128.3, 118.8 (d, *J* = 13.4 Hz), 116.9, 114.6 (d, *J* = 3.1 Hz), 106.2 (d, *J* = 28.8 Hz), 84.4, 74.4, 63.1, 46.9, 29.0. MSESI *m/z*: 386.1517 (C_20_H_20_FN_3_O_4_ + H^+^ requires 386.1511).

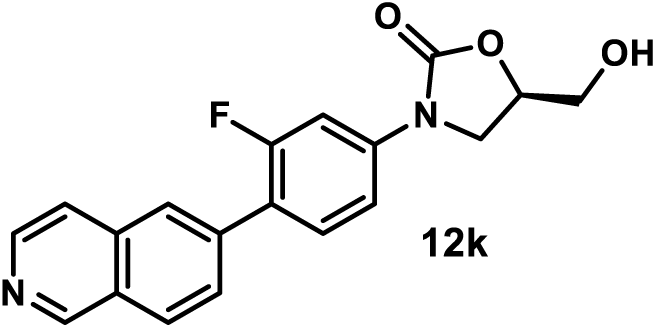

*(R)-3-(3-Fluoro-4-(isoquinolin-6-yl)phenyl)-5-(hydroxymethyl)oxazolidin-2-one (**12k**)*. Using general procedure 3, employing 6-bromoisoquinoline (42 mg, 0.2 mmol), compound **12k** was obtained as a yellow amorphous solid (20 mg, 59%). ^1^H NMR (500 MHz, DMSO-*d*_6_) δ 9.35 (s, 1H), 8.54 (d, *J* = 5.7 Hz, 1H), 8.21 (d, *J* = 8.6 Hz, 1H), 8.15 (s, 1H), 7.96 – 7.85 (m, 2H), 7.80 – 7.64 (m, 2H), 7.60 – 7.47 (m, 1H), 5.27 (s, 1H), 4.83 – 4.70 (m, 1H), 4.17 (t, *J* = 8.9 Hz, 1H), 3.91 (dd, *J* = 8.9, 6.1 Hz, 1H), 3.75 – 3.67 (m, 1H), 3.64 – 3.57 (m, 1H). ^13^C NMR (101 MHz, DMSO-*d*_6_) δ 159.3 (d, *J* = 245.6 Hz), 154.4, 152.1, 143.3, 140.2 (d, *J* = 11.3 Hz), 136.7, 135.3, 131.3 (d, *J* = 4.5 Hz), 128.2 (d, *J* = 3.2 Hz), 127.9, 127.2, 126.0 (d, *J* = 3.4 Hz), 121.8 (d, *J* = 13.2 Hz), 120.5, 113.9 (d, *J* = 3.3 Hz), 105.4 (d, *J* = 28.7 Hz), 73.5, 61.6, 46.0. MSESI *m/z*: 339.1135 (C_19_H_15_FN_2_O_3_ + H^+^ requires 339.1139).

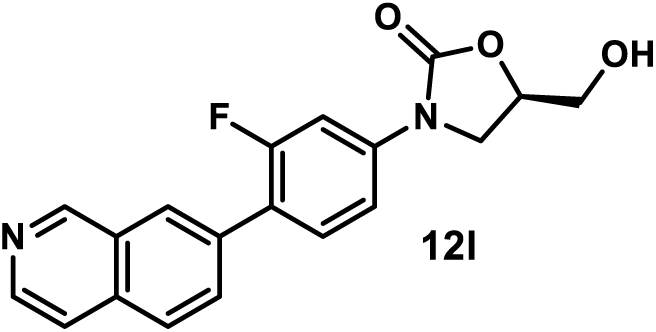

*(R)-3-(3-Fluoro-4-(isoquinolin-7-yl)phenyl)-5-(hydroxymethyl)oxazolidin-2-one (**12l**)*. Using general procedure 3, employing 7-bromoisoquinoline (42 mg, 0.2 mmol), compound **12l** was obtained as a yellow amorphous solid (16 mg, 47%). ^1^H NMR (500 MHz, DMSO-*d*_6_) δ 9.39 (s, 1H), 8.54 (d, *J* = 5.7 Hz, 1H), 8.31 (s, 1H), 8.07 (d, *J* = 8.6 Hz, 1H), 7.98 (d, *J* = 8.6 Hz, 1H), 7.87 (d, *J* = 5.7 Hz, 1H), 7.81 – 7.67 (m, 2H), 7.52 (dd, *J* = 8.5, 2.2 Hz, 1H), 5.27 (s, 1H), 4.96 – 4.71 (m, 1H), 4.17 (t, *J* = 9.0 Hz, 1H), 3.91 (dd, *J* = 9.0, 6.1 Hz, 1H), 3.77 – 3.68 (m, 1H), 3.64 – 3.55 (m, 1H). ^13^C NMR (101 MHz, DMSO-*d*_6_) δ 159.2 (d, *J* = 245.2 Hz), 154.4, 152.7, 143.2, 139.9 (d, *J* = 11.2 Hz), 134.3, 133.6, 131.2 (d, *J* = 2.4 Hz), 131.2, 128.3, 127.1 (d, *J* = 3.3 Hz), 126.8, 121.8 (d, *J* = 13.2 Hz), 120.1, 113.9 (d, *J* = 3.1 Hz), 105.4 (d, *J* = 28.8 Hz), 73.4, 61.6, 46.0. MSESI *m/z*: 339.1130 (C_19_H_15_FN_2_O_3_ + H^+^ requires 339.1139).

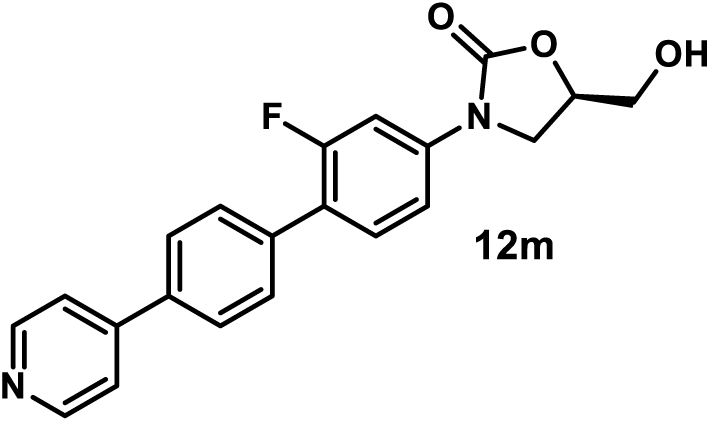

*(R)-3-(2-Fluoro-4’-(pyridin-4-yl)-[1,1’-biphenyl]-4-yl)-5-(hydroxymethyl)oxazolidin-2-one (**12m**)*. Using general procedure 3, employing 4-(4-bromophenyl)pyridine (47 mg, 0.2 mmol), compound **12m** was obtained as a gray amorphous solid (14 mg, 38%). ^1^H NMR (500 MHz, DMSO-*d*_6_) δ 8.66 (d, *J* = 5.1 Hz, 2H), 7.92 (d, *J* = 8.0 Hz, 2H), 7.77 (d, *J* = 5.1 Hz, 2H), 7.71 (d, *J* = 8.0 Hz, 2H), 7.68 – 7.58 (m, 2H), 7.54 – 7.43 (m, 1H), 5.25 (t, *J* = 5.7 Hz, 1H), 4.86 – 4.68 (m, 1H), 4.15 (t, *J* = 9.0 Hz, 1H), 3.90 (dd, *J* = 9.0, 6.1 Hz, 1H), 3.78 – 3.67 (m, 1H), 3.63 – 3.52 (m, 1H). ^13^C NMR (101 MHz, DMSO-*d*_6_) δ 159.1 (d, *J* = 245.2 Hz), 154.4, 150.3 (2C), 146.4, 139.8 (d, *J* = 11.4 Hz), 136.2, 135.5, 130.8 (d, *J* = 4.7 Hz), 129.3 (d, *J* = 3.2 Hz, 2C), 127.1 (2C), 121.8 (d, *J* = 13.1 Hz), 121.1 (2C), 113.9 (d, *J* = 2.9 Hz), 105.4 (d, *J* = 28.7 Hz), 73.4, 61.6, 46.0. MSESI *m/z*: 365.1316 (C_21_H_17_FN_2_O_3_ + H^+^ requires 365.1296).

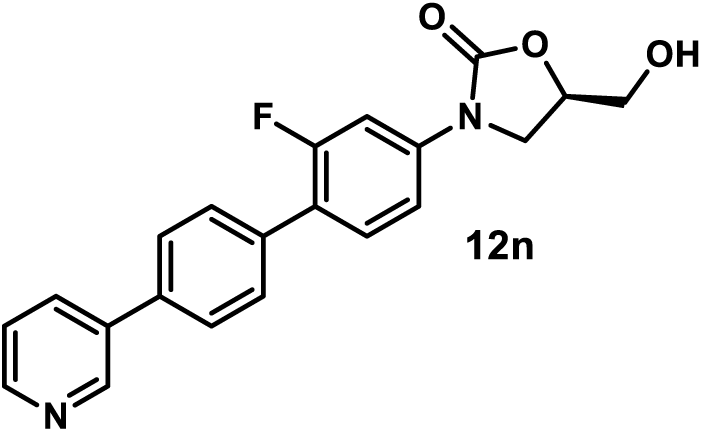

*(R)-3-(2-Fluoro-4’-(pyridin-3-yl)-[1,1’-biphenyl]-4-yl)-5-(hydroxymethyl)oxazolidin-2-one (**12n**)*. Using general procedure 3, employing 3-(4-bromophenyl)pyridine (47 mg, 0.2 mmol), compound **12n** was obtained by flash column chromatography (SiO_2_, eluent gradient 0-6% MeOH in DCM) as a yellow amorphous solid (17 mg, 47%). ^1^H NMR (500 MHz, DMSO-*d*_6_) δ 8.96 (d, *J* = 1.9 Hz, 1H), 8.60 (dd, *J* = 4.7, 1.9 Hz, 1H), 8.14 (dt, *J* = 8.1, 1.9 Hz, 1H), 7.93 – 7.82 (m, 2H), 7.75 – 7.59 (m, 4H), 7.55 – 7.45 (m, 2H), 5.37 – 5.18 (m, 1H), 4.84 – 4.71 (m, 1H), 4.15 (t, *J* = 8.9 Hz, 1H), 3.90 (dd, *J* = 8.9, 6.1 Hz, 1H), 3.77 – 3.66 (m, 1H), 3.63 – 3.54 (m, 1H). ^13^C NMR (101 MHz, DMSO-*d*_6_) δ 159.1 (d, *J* = 244.7 Hz), 154.4, 148.6, 147.6, 139.6 (d, *J* = 11.2 Hz), 136.2, 135.0, 134.4, 134.1, 130.8 (d, *J* = 4.7 Hz), 129.3 (d, *J* = 3.3 Hz, 2C), 127.1 (2C), 123.9, 122.0 (d, *J* = 13.1 Hz), 113.9 (d, *J* = 3.0 Hz), 105.4 (d, *J* = 28.8 Hz), 73.4, 61.6, 46.0. MSESI *m/z*: 365.1279 (C_21_H_17_FN_2_O_3_ + H^+^ requires 365.1296).

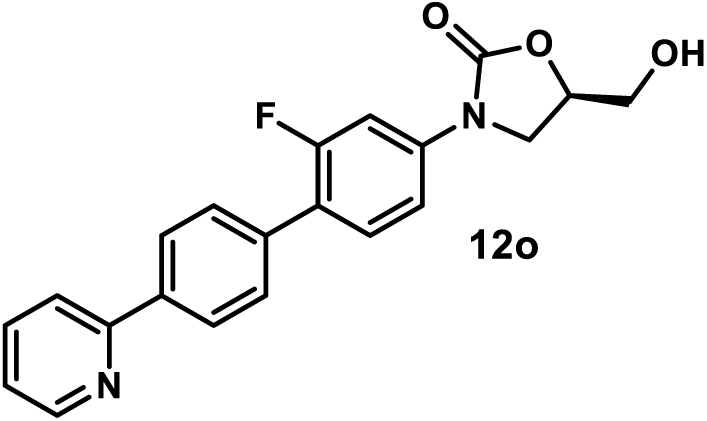

*(R)-3-(2-Fluoro-4’-(pyridin-2-yl)-[1,1’-biphenyl]-4-yl)-5-(hydroxymethyl)oxazolidin-2-one (**12o**)*. Using general procedure 3, employing 2-(4-bromophenyl)pyridine (47 mg, 0.2 mmol), compound **12o** was obtained by flash column chromatography (SiO_2_, eluent gradient 0-100% EtOAc in hexanes) as a yellow amorphous solid (24 mg, 66%). ^1^H NMR (500 MHz, DMSO-*d*_6_) δ 8.75 – 8.63 (m, 1H), 8.20 (d, *J* = 8.4 Hz, 2H), 8.02 (d, *J* = 8.0 Hz, 1H), 7.91 (td, *J* = 8.0, 1.8 Hz, 1H), 7.74 – 7.59 (m, 4H), 7.48 (dd, *J* = 8.6, 2.2 Hz, 1H), 7.38 (ddd, *J* = 8.6, 4.8, 1.0 Hz, 1H), 5.25 (t, *J* = 5.6 Hz, 1H), 4.83 – 4.71 (m, 1H), 4.15 (t, *J* = 8.9 Hz, 1H), 3.90 (dd, *J* = 8.9, 6.1 Hz, 1H), 3.71 (ddd, *J* = 12.4, 5.6, 3.3 Hz, 1H), 3.59 (ddd, *J* = 12.4, 5.6, 4.0 Hz, 1H).^13^C NMR (101 MHz, DMSO-*d*_6_) δ 159.1 (d, *J* = 245.3 Hz), 155.4, 154.4, 149.6, 139.6 (d, *J* = 11.3 Hz), 137.8, 137.3, 135.2 (d, *J* = 1.7 Hz), 130.8 (d, *J* = 4.8 Hz), 128.9 (d, *J* = 3.2 Hz, 2C), 126.7 (2C), 122.7, 122.1 (d, *J* = 13.3 Hz), 120.3, 113.8 (d, *J* = 3.0 Hz), 105.4 (d, *J* = 28.9 Hz), 73.4, 61.6, 46.0. MSESI *m/z*: 365.1307 (C_21_H_17_FN_2_O_3_ + H ^+^ requires 365.1296).

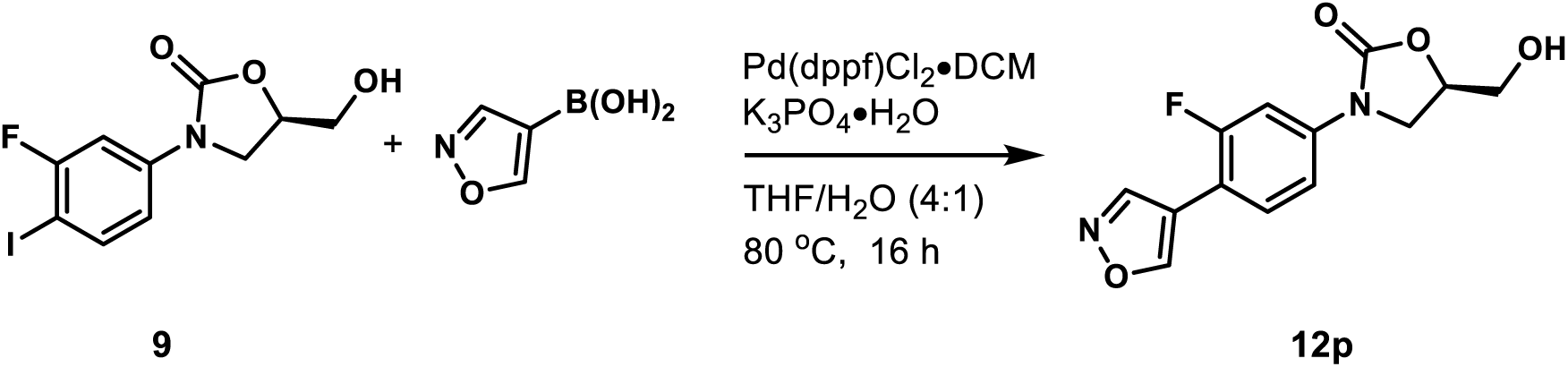

*(R)-3-(3-Fluoro-4-(isoxazol-4-yl)phenyl)-5-(hydroxymethyl)oxazolidin-2-one (**12p**)*.^57^ A mixture of compound **9** (34 mg, 0.1 mmol), isoxazol-4-ylboronic acid (17 mg, 0.15 mmol), potassium phosphate monohydrate (46 mg, 0.2 mmol), and [1,1’-bis(diphenylphosphino)ferrocene]dichloropalladium(II), complex with dichloromethane (8.2 mg, 0.01 mmol) in THF/H_2_O (v/v= 4:1, 2 mL) was stirred at 80 °C. After the reaction was judged to be completed by TLC (16 h), it was cooled to room temperature, diluted with EtOAc, washed with water, and concentrated under reduced pressure by rotary evaporation. The resulting residue was purified by flash column chromatography (SiO_2_, eluent gradient 0-100% EtOAc in hexanes) to afford compound **12p** as a yellow amorphous solid (14 mg, 50%). ^1^H NMR (300 MHz, Acetone-*d*_6_) δ 9.13 (d, *J* = 2.2 Hz, 1H), 8.94 (d, *J* = 2.2 Hz, 1H), 7.85 – 7.66 (m, 2H), 7.51 – 7.41 (m, 1H), 4.82 (ddt, *J* = 9.0, 6.2, 3.7 Hz, 1H), 4.44 (s, 1H), 4.22 (t, *J* = 9.0 Hz, 1H), 4.04 (dd, *J* = 9.0, 6.2 Hz, 1H), 3.95 – 3.85 (m, 1H), 3.83 – 3.71 (m, 1H). ^13^C NMR (101 MHz, Acetone-*d*_6_) δ 160.3 (d, *J* = 245.4 Hz), 156.3 (d, *J* = 8.7 Hz), 155.2, 148.9 (d, *J* = 2.7 Hz), 141.0 (d, *J* = 11.3 Hz), 129.5 (d, *J* = 5.3 Hz), 115.3 (d, *J* = 2.3 Hz), 114.4 (d, *J* = 3.0 Hz), 112.0 (d, *J* = 14.8 Hz), 106.1 (d, *J* = 28.2 Hz), 74.3, 63.1, 46.9. MSESI *m/z*: 279.0787 (C_13_H_11_FN_2_O_4_ + H^+^ requires 279.0776).

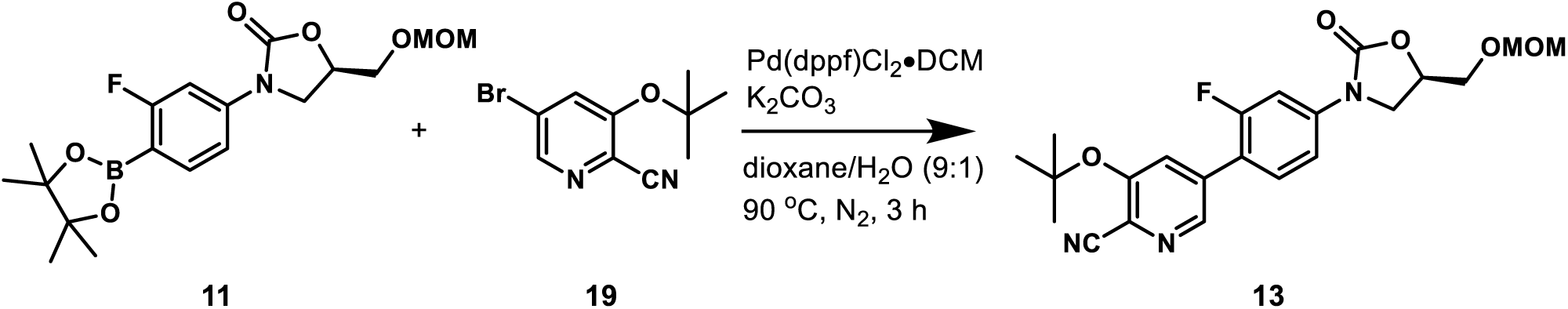

*(R)-3-(tert-Butoxy)-5-(2-fluoro-4-(5-((methoxymethoxy)methyl)-2-oxooxazolidin-3-yl)phenyl)picolinonitrile (**13**)*. A mixture of compound **11** (152 mg, 0.4 mmol), compound **19** (122 mg, 0.48 mmol), potassium carbonate (221 mg, 1.6 mmol), and [1,1’-bis(diphenylphosphino)ferrocene]dichloropalladium(II), complex with dichloromethane (33 mg, 0.04 mmol) in dioxane/H_2_O (v/v= 9:1, 2 mL) was stirred at 90 °C under nitrogen atmosphere. After the reaction was judged to be completed by TLC (3 h), it was cooled to room temperature, diluted with EtOAc, washed with water, and concentrated under reduced pressure by rotary evaporation. The crude residue was purified by flash column chromatography (SiO_2_, eluent gradient 0-100% EtOAc in hexanes) to afford compound **13** as a yellow amorphous solid (154 mg, 90%). ^1^H NMR (400 MHz, acetone) δ 8.59 (t, *J* = 1.6 Hz, 1H), 7.94 (t, *J* = 1.6 Hz, 1H), 7.78 (dd, *J* = 13.8, 2.3 Hz, 1H), 7.71 (t, *J* = 8.6 Hz, 1H), 7.56 (dd, *J* = 8.6, 2.3 Hz, 1H), 5.05 – 4.90 (m, 1H), 4.67 (s, 2H), 4.33 (t, *J* = 9.0 Hz, 1H), 4.07 (dd, *J* = 9.0, 6.1 Hz, 1H), 3.94 – 3.75 (m, 2H), 3.33 (s, 3H), 1.56 (s, 9H).

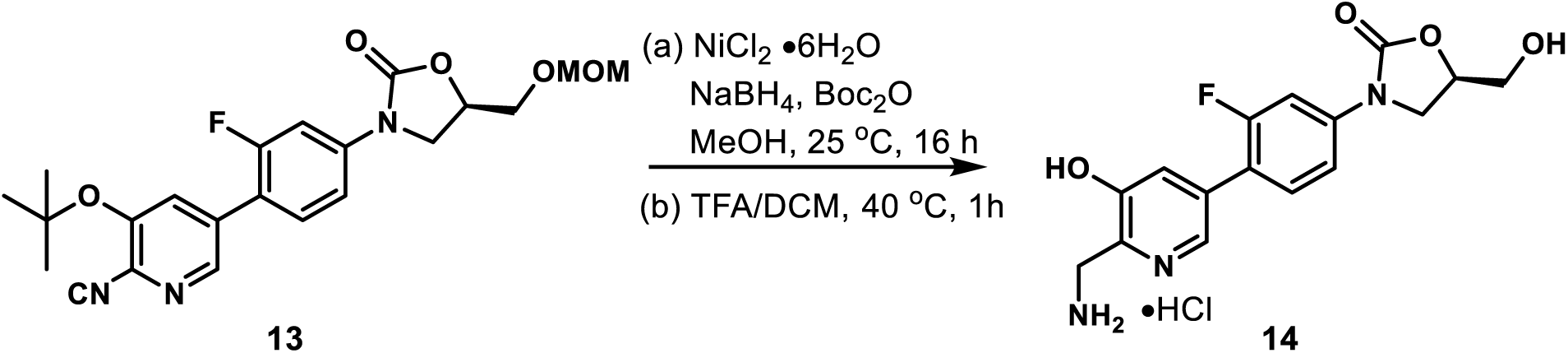

*(R)-3-(4-(6-(Aminomethyl)-5-hydroxypyridin-3-yl)-3-fluorophenyl)-5-(hydroxymethyl)oxazolidin-2-one Hydrochloride (**14**)*. To a solution of compound **13** (86 mg, 0.2 mmol) in methanol (3 mL) in an ice bath were added nickel(II) chloride hexahydrate (24 mg, 0.1 mmol) and di-*tert*-butyl dicarbonate (30% in dioxane, 0.73 mL, 1 mmol). The resulting solution was charged with the addition of sodium borohydride (76 mg, 2 mmol) portion wise and stirred at 25 °C. After the reaction was judged to be completed by TLC (16 h), it was evaporated under reduced pressure by rotary evaporation. The resulting residue was purified by flash column chromatography (SiO_2_, eluent gradient 0-100% EtOAc) to afford compound **13a** as colorless oil (56 mg, 52%). ^1^H NMR (500 MHz, acetone) δ 8.37 (t, *J* = 1.6 Hz, 1H), 7.73 (dd, *J* = 13.6, 2.3 Hz, 1H), 7.66 (t, *J* = 1.6 Hz, 1H), 7.61 (t, *J* = 8.6 Hz, 1H), 7.50 (dd, *J* = 8.6, 2.3 Hz, 1H), 6.20 (s, 1H), 4.97 (dddd, *J* = 9.0, 6.1, 4.5, 3.5 Hz, 1H), 4.67 (s, 2H), 4.44 (d, *J* = 5.0 Hz, 2H), 4.31 (t, *J* = 9.0 Hz, 1H), 4.05 (dd, *J* = 9.0, 6.1 Hz, 1H), 3.87 (dd, *J* = 11.3, 3.5 Hz, 1H), 3.82 (dd, *J* = 11.3, 4.5 Hz, 1H), 1.51 (s, 9H), 1.45 (s, 9H). A solution of compound **13a** (28 mg, 0.05 mmol) in DCM (1 mL) and TFA (0.5 mL) was stirred at 40 °C. After the reaction was judged to be completed by TLC (1 h), its solvent was evaporated under reduced pressure by rotary evaporation. The resulting residue was dissolved in MeOH (1 mL), charged with the addition of HCl/dioxane (4 M, 0.1 mL), and concentrated under reduced pressure by rotary evaporation. The resulting solid was washed with acetone (2 × 1mL) and dried under high vacuum to give **14** as a yellow amorphous solid (12 mg, 66%). ^1^H NMR (400 MHz, CD_3_OD) δ 8.42 (t, *J* = 1.5 Hz, 1H), 7.75 (dd, *J* = 13.6, 2.2 Hz, 1H), 7.69 (t, *J* = 1.5 Hz, 1H), 7.61 (t, *J* = 8.6 Hz, 1H), 7.49 (dd, *J* = 8.6, 2.2 Hz, 1H), 4.84 – 4.76 (m, 1H), 4.37 (s, 2H), 4.19 (t, *J* = 8.9 Hz, 1H), 4.00 (dd, *J* = 8.9, 6.3 Hz, 1H), 3.89 (dd, *J* = 12.6, 3.1 Hz, 1H), 3.72 (dd, *J* = 12.6, 3.8 Hz, 1H). ^13^C NMR (101 MHz, CD_3_OD) δ 161.2 (d, *J* = 246.6 Hz), 156.8, 153.9, 142.4 (d, *J* = 11.3 Hz), 138.5, 138.3 (d, *J* = 3.7 Hz), 135.2, 131.8 (d, *J* = 4.2 Hz), 125.9, 119.8 (d, *J* = 13.7 Hz), 115.3 (d, *J* = 3.1 Hz), 107.1 (d, *J* = 28.9 Hz), 75.3, 63.2, 47.5, 39.3. MSESI *m/z*: 334.1196 (C_16_H_16_FN_3_O_4_ + H^+^ requires 334.1198).

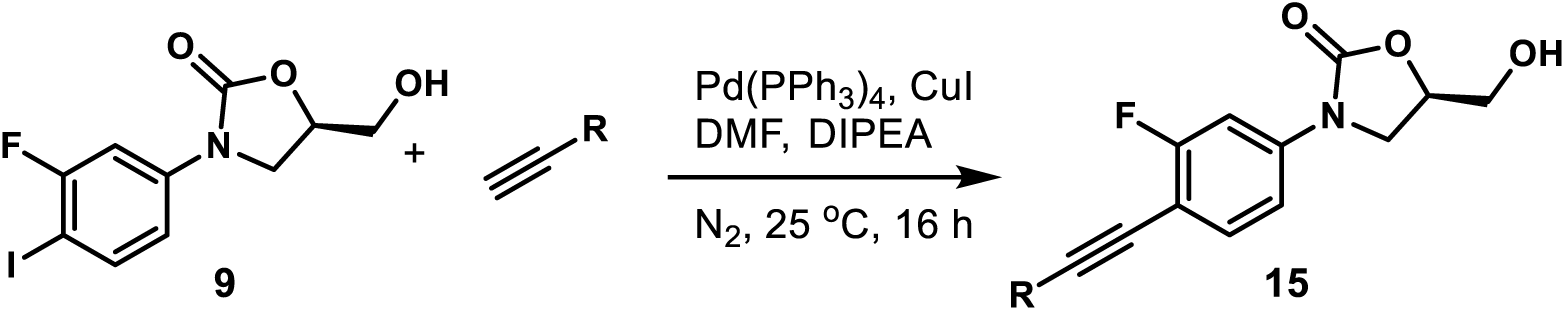

#### General Procedure 4: Synthesis of Alkynyl Oxazolidinone Analogs 15a-15v

A solution of compound **9** (34 mg, 0.1 mmol), alkyne (0.15 mmol), tetrakis(triphenylphosphine)palladium(0) (12 mg, 0.01 mmol), and copper(I) iodide (2 mg, 0.01 mmol) in DMF (1 mL) was evacuated and purged with nitrogen three times. *N*,*N*-diisopropylethylamine (0.5 mL) was added under the protection of nitrogen atmosphere. The mixture was stirred at 25 °C. After the reaction was judged to be completed by TLC (16 h), it was diluted with EtOAc, washed with water three times, and concentrated under reduced pressure by rotary evaporation. The crude residue was purified by flash column chromatography (SiO_2_) to afford compound **15**.

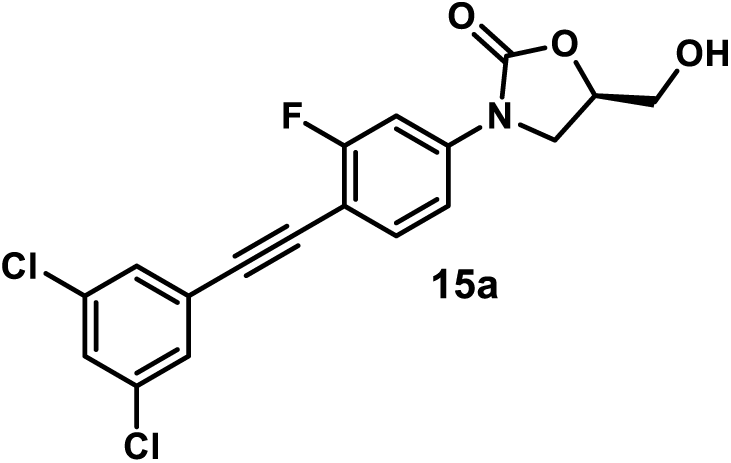

*(R)-3-(4-((3,5-Dichlorophenyl)ethynyl)-3-fluorophenyl)-5-(hydroxymethyl)oxazolidin-2-one (**15a**)*. Using general procedure 4, employing 1,3-dichloro-5-ethynylbenzene (26 mg, 0.15 mmol), compound **15a** was obtained after flash column chromatography (SiO_2_, 0-100% EtOAc in hexanes) as a yellow amorphous solid (34 mg, 89%). ^1^H NMR (400 MHz, Acetone-*d*_6_) δ 7.72 (dd, *J* = 12.5, 2.2 Hz, 1H), 7.62 (t, *J* = 8.4 Hz, 1H), 7.54 (s, 3H), 7.45 (dd, *J* = 8.4, 2.2 Hz, 1H), 4.93 – 4.75 (m, 1H), 4.43 (s, 1H), 4.23 (t, *J* = 8.9 Hz, 1H), 4.05 (dd, *J* = 8.9, 6.3 Hz, 1H), 3.95 – 3.87 (m, 1H), 3.78 (dd, *J* = 12.5, 3.6 Hz, 1H). ^13^C NMR (101 MHz, Acetone-*d*_6_) δ 163.7 (d, *J* = 248.8 Hz), 155.2, 142.7 (d, *J* = 11.0 Hz), 135.9 (2C), 134.9 (d, *J* = 2.4 Hz), 130.5 (2C), 129.6, 127.0, 114.2 (d, *J* = 3.1 Hz), 105.7 (d, *J* = 27.1 Hz), 105.4 (d, *J* = 16.2 Hz), 91.4 (d, *J* = 3.0 Hz), 86.0, 74.5, 63.2, 47.0.

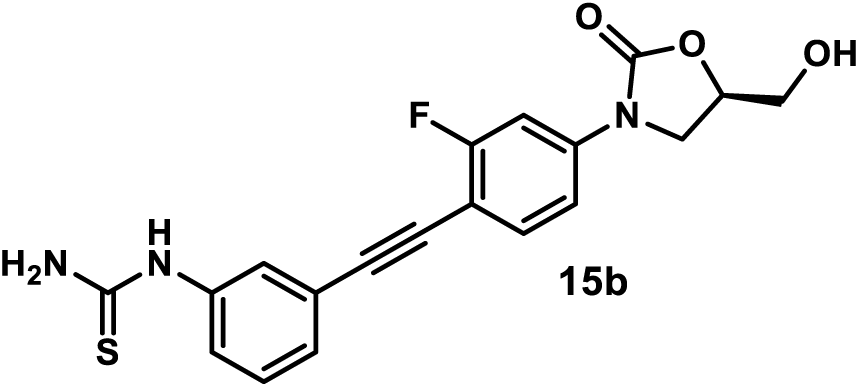

*(R)-1-(3-((2-Fluoro-4-(5-(hydroxymethyl)-2-oxooxazolidin-3-yl)phenyl)ethynyl)phenyl)thiourea (**15b**).* Using general procedure 4, employing (3-ethynylphenyl)thiourea (26 mg, 0.15 mmol), compound **15b** was obtained after flash column chromatography (SiO_2_, 0-15% MeOH in DCM) followed by high-performance liquid chromatography purification as a yellow amorphous solid (17 mg, 44%). ^1^H NMR (400 MHz, DMSO) δ 9.77 (s, 1H), 7.76 – 7.60 (m, 3H), 7.43 (d, *J* = 8.8 Hz, 2H), 7.37 (t, *J* = 7.8 Hz, 1H), 7.31 – 7.26 (m, 1H), 5.21 (t, *J* = 5.6 Hz, 1H), 4.79 – 4.65 (m, 1H), 4.13 (t, *J* = 9.0 Hz, 1H), 3.87 (dd, *J* = 9.0, 6.0 Hz, 2H), 3.75 – 3.63 (m, 1H), 3.64 – 3.51 (m, 1H). ^13^C NMR (101 MHz, DMSO-*d*_6_) δ 181.1, 162.0 (d, *J* = 247.4 Hz), 154.3, 140.7 (d, *J* = 11.1 Hz), 139.7, 133.8 (d, *J* = 2.5 Hz), 129.3, 127.1, 125.2, 123.5, 122.1, 113.5 (d, *J* = 3.0 Hz), 104.7 (d, *J* = 26.8 Hz), 104.6 (d, *J* = 15.7 Hz), 93.5 (d, *J* = 2.02 Hz), 82.5, 73.5, 61.6, 46.0.

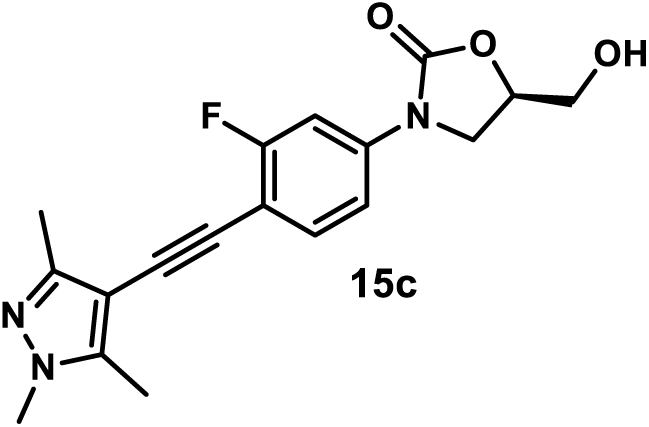

*(R)-3-(3-Fluoro-4-((1,3,5-trimethyl-1H-pyrazol-4-yl)ethynyl)phenyl)-5-(hydroxymethyl)oxazolidin-2-one (**15c**).* Using general procedure 4, employing 4-ethynyl-1,3,5-trimethyl-1*H*-pyrazole (20 mg, 0.15 mmol), compound **15c** was obtained after flash column chromatography (SiO_2_, 0-10% MeOH in DCM) as a white amorphous solid (20 mg, 59%). ^1^H NMR (300 MHz, DMSO-*d*_6_) δ 7.75 – 7.49 (m, 2H), 7.38 (dd, *J* = 8.6, 2.2 Hz, 1H), 5.25 (t, *J* = 5.6 Hz, 1H), 4.84 – 4.53 (m, 1H), 4.11 (t, *J* = 9.0 Hz, 1H), 3.85 (dd, *J* = 9.0, 6.1 Hz, 1H), 3.76 – 3.62 (m, 4H), 3.56 (ddd, *J* = 12.4, 5.6, 3.9 Hz, 1H), 2.30 (s, 3H), 2.18 (s, 3H). ^13^C NMR (151 MHz, DMSO-*d*_6_) δ 161.4 (d, *J* = 246.4 Hz), 154.2, 147.7, 142.0, 139.6 (d, *J* = 10.6 Hz), 133.0, 113.4 (d, *J* = 2.8 Hz), 105.9 (d, *J* = 15.9 Hz), 104.7 (d, *J* = 26.9 Hz), 100.1, 86.7 (d, *J* = 1.5 Hz), 85.2, 73.4, 61.5, 45.9, 36.0, 12.1, 10.0.

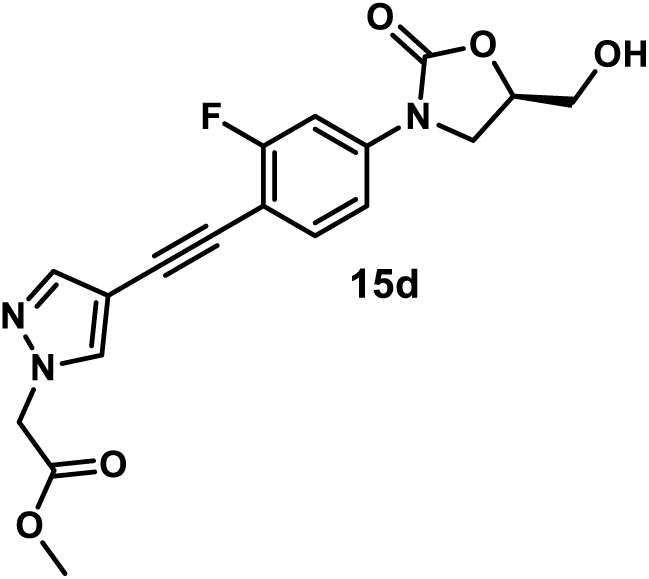

*Methyl (R)-2-(4-((2-Fluoro-4-(5-(hydroxymethyl)-2-oxooxazolidin-3-yl)phenyl)ethynyl)-1H-pyrazol-1-yl)acetate (**15d**)*. Using general procedure 4, employing methyl (4-ethynyl-1*H*-pyrazol-1-yl)acetate (25 mg, 0.15 mmol), compound **15d** was obtained after flash column chromatography (SiO_2_, 0-10% MeOH in DCM) as a white amorphous solid (17 mg, 46%). ^1^H NMR (400 MHz, Acetone-*d*_6_) δ 8.03 (s, 1H), 7.73 – 7.63 (m, 2H), 7.52 (t, *J* = 8.6 Hz, 1H), 7.40 (dd, *J* = 8.6, 2.3 Hz, 1H), 5.09 (s, 2H), 4.89 – 4.72 (m, 1H), 4.44 (s, 1H), 4.22 (t, *J* = 8.9 Hz, 1H), 4.03 (dd, *J* = 8.9, 6.2 Hz, 1H), 3.90 (d, *J* = 12.7 Hz, 1H), 3.81 – 3.64 (m, 4H). ^13^C NMR (101 MHz, Acetone-*d*_6_) δ 168.9, 163.2 (d, *J* = 247.2 Hz), 155.1, 142.5, 141.5 (d, *J* = 10.9 Hz), 135.0, 134.2 (d, *J* = 2.9 Hz), 114.0 (d, *J* = 3.1 Hz), 106.8 (d, *J* = 16.3 Hz), 105.6 (d, *J* = 27.2 Hz), 104.0, 86.0 (d, *J* = 2.9 Hz), 83.5, 74.4, 63.1, 53.5, 52.7, 46.9.

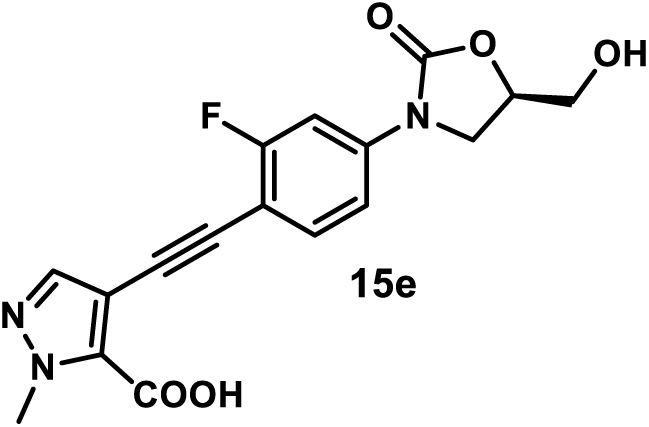

*(R)-4-((2-Fluoro-4-(5-(hydroxymethyl)-2-oxooxazolidin-3-yl)phenyl)ethynyl)-1-methyl-1H-pyrazole-5-carboxylic Acid (**15e**)*. Using general procedure 4, employing 4-ethynyl-1-methyl-1*H*-pyrazole-5-carboxylic acid (23 mg, 0.15 mmol), compound **15e** was obtained after flash column chromatography (SiO_2_, 0-15% MeOH in DCM) followed by high-performance liquid chromatography purification as a gray amorphous solid (11 mg, 42%). ^1^H NMR (400 MHz, DMSO-*d*_6_) δ 7.97 (s, 1H), 7.82 (t, *J* = 8.9 Hz, 1H), 7.70 (dd, *J* = 14.6, 2.3 Hz, 1H), 7.49 (dd, *J* = 8.9, 2.3 Hz, 1H), 7.25 (s, 1H), 5.27 (t, *J* = 5.6 Hz, 1H), 4.82 – 4.70 (m, 1H), 4.21 (s, 3H), 4.13 (t, *J* = 8.9 Hz, 1H), 3.88 (dd, *J* = 8.9, 6.0 Hz, 1H), 3.70 (ddd, *J* = 12.3, 5.6, 3.2 Hz, 1H), 3.57 (ddd, *J* = 12.3, 5.6, 3.9 Hz, 1H). ^13^C NMR (101 MHz, DMSO-*d*_6_) δ 159.0 (d, *J* = 248.9 Hz), 154.3, 153.5, 146.6 (d, *J* = 4.8 Hz), 140.7 (d, *J* = 11.8 Hz), 133.9, 128.7 (d, *J* = 3.4 Hz), 125.9, 124.9, 114.3 (d, *J* = 10.8 Hz), 113.6 (d, *J* = 2.8 Hz), 105.4 (d, *J* = 28.7 Hz), 100.7 (d, *J* = 13.0 Hz), 73.6, 61.6, 46.0, 38.4.

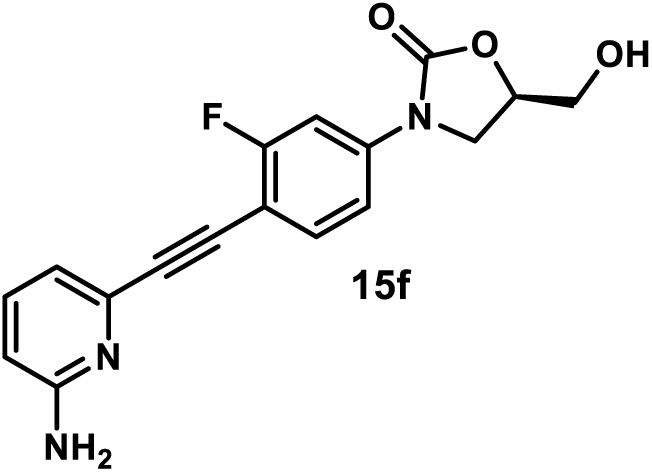

*(R)-3-(4-((6-Aminopyridin-2-yl)ethynyl)-3-fluorophenyl)-5-(hydroxymethyl)oxazolidin-2-one (**15f**)*. Using general procedure 4, employing 6-ethynylpyridin-2-amine (18 mg, 0.15 mmol), compound **15f** was obtained after flash column chromatography (SiO_2_, 0-8% MeOH in DCM) followed by high-performance liquid chromatography purification as a yellow amorphous solid (14 mg, 42%). ^1^H NMR (400 MHz, DMSO-*d*6) δ 7.73 – 7.56 (m, 2H), 7.48 – 7.33 (m, 2H), 6.74 (d, *J* = 7.2 Hz, 1H), 6.46 (d, *J* = 8.4 Hz, 1H), 6.19 (s, 2H), 5.33 – 5.16 (m, 1H), 4.82 – 4.67 (m, 1H), 4.12 (t, *J* = 9.0 Hz, 1H), 3.86 (dd, *J* = 9.0, 6.0 Hz, 1H), 3.72 – 3.64 (m, 1H), 3.61 – 3.52 (m, 1H). ^13^C NMR (101 MHz, DMSO-*d*_6_) δ 162.2 (d, *J* = 247.7 Hz), 159.8, 154.2, 140.8 (d, *J* = 11.1 Hz), 139.8, 137.4, 133.9 (d, *J* = 2.5 Hz), 115.6, 113.5 (d, *J* = 3.0 Hz), 108.8, 104.8 (d, *J* = 26.8 Hz), 104.3 (d, *J* = 15.8 Hz), 94.1 (d, *J* = 2.9 Hz), 79.7, 73.5, 61.6, 46.0.

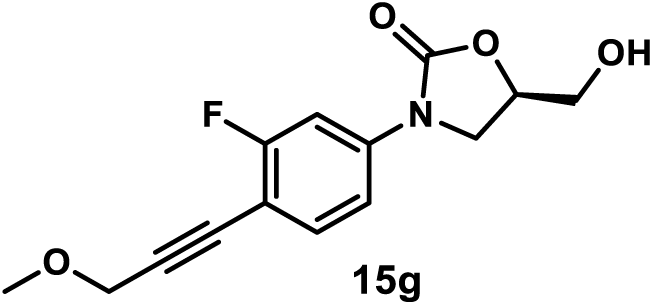

*(R)-3-(3-Fluoro-4-(3-methoxyprop-1-yn-1-yl)phenyl)-5-(hydroxymethyl)oxazolidin-2-one (**15g**)*. Using general procedure 4, employing methyl propargyl ether (12.7 µL, 0.15 mmol), compound **15g** was obtained after flash column chromatography (SiO_2_, 0-100% EtOAc in hexanes) as a white amorphous solid (22 mg, 79%). ^1^H NMR (600 MHz, Acetone-*d*_6_) δ 7.67 (dd, *J* = 12.4, 2.1 Hz, 1H), 7.50 (t, *J* = 8.5 Hz, 1H), 7.38 (dd, *J* = 8.5, 2.1 Hz, 1H), 4.85 – 4.75 (m, 1H), 4.38 (t, *J* = 6.0 Hz, 1H), 4.33 (s, 2H), 4.21 (t, *J* = 8.9 Hz, 1H), 4.02 (dd, *J* = 8.9, 6.2 Hz, 1H), 3.92 – 3.86 (m, 1H), 3.79 – 3.72 (m, 1H), 3.38 (s, 3H). ^13^C NMR (101 MHz, Acetone-*d*_6_) δ 163.7 (d, *J* = 247.8 Hz), 155.1, 142.0 (d, *J* = 10.9 Hz), 134.6 (d, *J* = 2.8 Hz), 113.9 (d, *J* = 3.2 Hz), 105.9 (d, *J* = 16.3 Hz), 105.6 (d, *J* = 27.3 Hz), 90.9 (d, *J* = 3.1 Hz), 79.7, 74.4, 63.1, 60.5, 57.5, 46.9.

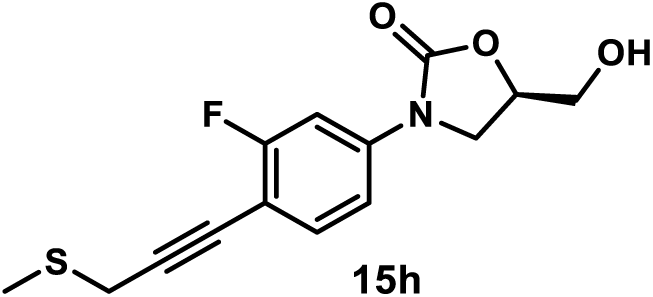

*(R)-3-(3-Fluoro-4-(3-(methylthio)prop-1-yn-1-yl)phenyl)-5-(hydroxymethyl)oxazolidin-2-one (**15h**)*. A solution of compound **9** (34 mg, 0.1 mmol), 3-(methylsulfanyl)-1-propyne (39 mg, 0.3 mmol), tetrakis(triphenylphosphine)palladium(0) (23 mg, 0.02 mmol), and copper(I) iodide (3.8 mg, 0.02 mmol) in DMF (1 mL) was evacuated and purged with nitrogen three times. *N*,*N*-diisopropylethylamine (0.5 mL) was added under the protection of nitrogen atmosphere. The mixture was stirred at 25 °C. After the reaction was judged to be completed by TLC (16 h), it was diluted with EtOAc, washed with water three times, and concentrated under reduced pressure by rotary evaporation. The crude residue was purified by flash column chromatography (SiO_2_, 0-100% EtOAc in hexanes) to afford compound **15** as a yellow amorphous solid (20 mg, 68%). ^1^H NMR (400 MHz, Acetone-*d*_6_) δ 7.65 (dd, *J* = 12.4, 2.3 Hz, 1H), 7.47 (t, *J* = 8.5 Hz, 1H), 7.36 (dd, *J* = 8.5, 2.3 Hz, 1H), 4.89 – 4.73 (m, 1H), 4.41 (t, *J* = 5.9 Hz, 1H), 4.20 (t, *J* = 8.9 Hz, 1H), 4.01 (dd, *J* = 8.9, 6.2 Hz, 1H), 3.90 (ddd, *J* = 12.3, 5.9, 3.3 Hz, 1H), 3.76 (ddd, *J* = 12.3, 5.9, 3.9 Hz, 1H), 3.57 (s, 2H), 2.26 (s, 3H). ^13^C NMR (101 MHz, Acetone-*d*_6_) δ 163.6 (d, *J* = 247.3 Hz), 155.1, 141.6 (d, *J* = 10.9 Hz), 134.5 (d, *J* = 2.9 Hz), 113.9 (d, *J* = 3.1 Hz), 106.4 (d, *J* = 16.2 Hz), 105.5 (d, *J* = 27.3 Hz), 91.2 (d, *J* = 3.1 Hz), 76.3, 74.3, 63.1, 46.9, 22.3, 15.1.

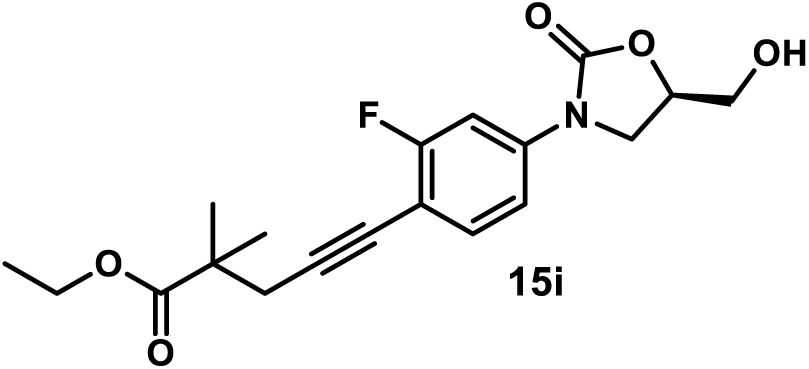

*Ethyl (R)-5-(2-Fluoro-4-(5-(hydroxymethyl)-2-oxooxazolidin-3-yl)phenyl)-2,2-dimethylpent-4-ynoate (**15i**)*. Using general procedure 4, employing ethyl 2,2-dimethyl-4-pentynoate (23 mg, 0.15 mmol), compound **15i** was obtained after flash column chromatography (SiO_2_, 0-100% EtOAc in hexanes) as yellow oil (21 mg, 58%). ^1^H NMR (400 MHz, Acetone-*d*_6_) δ 7.63 (dd, *J* = 12.4, 2.3 Hz, 1H), 7.42 (t, *J* = 8.6 Hz, 1H), 7.34 (dd, *J* = 8.6, 2.3 Hz, 1H), 4.85 – 4.68 (m, 1H), 4.39 (t, *J* = 5.7 Hz, 1H), 4.24 – 4.07 (m, 3H), 4.00 (dd, *J* = 8.9, 6.2 Hz, 1H), 3.93 – 3.81 (m, 1H), 3.82 – 3.69 (m, 1H), 2.70 (s, 2H), 1.31 (s, 6H), 1.23 (t, *J* = 7.1 Hz, 3H). ^13^C NMR (101 MHz, Acetone-*d*_6_) δ 176.5, 163.6 (d, *J* = 247.0 Hz), 155.1, 141.3 (d, *J* = 10.8 Hz), 134.4 (d, *J* = 2.9 Hz), 113.8 (d, *J* = 3.1 Hz), 106.9 (d, *J* = 16.3 Hz), 105.6 (d, *J* = 27.3 Hz), 92.5 (d, *J* = 3.0 Hz), 76.2, 74.3, 63.1, 61.1, 46.9, 43.0, 31.1, 24.9, 14.5.

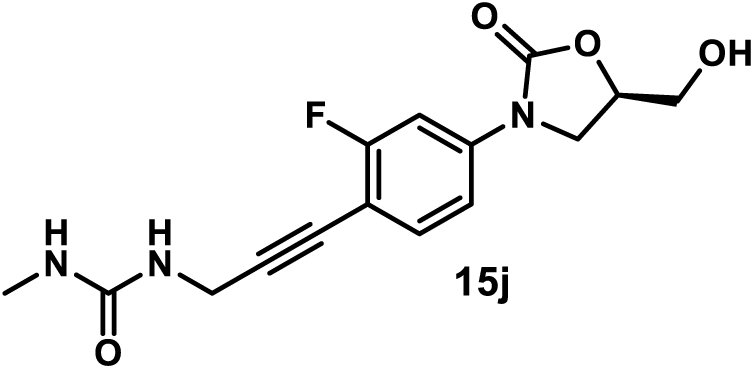

*(R)-1-(3-(2-Fluoro-4-(5-(hydroxymethyl)-2-oxooxazolidin-3-yl)phenyl)prop-2-yn-1-yl)-3-methylurea (**15j**)*. Using general procedure 4, employing *N*-methyl-*N*’-(2-propynyl)urea (17 mg, 0.15 mmol), compound **15j** was obtained after flash column chromatography (SiO_2_, 0-10% MeOH in DCM) as a yellow amorphous solid (7 mg, 22%). ^1^H NMR (400 MHz, DMSO-*d*_6_) δ 7.58 (dd, *J* = 12.3, 2.3 Hz, 1H), 7.49 (t, *J* = 8.6 Hz, 1H), 7.37 (dd, *J* = 8.6, 2.3 Hz, 1H), 6.39 (t, *J* = 5.8 Hz, 1H), 5.92 (q, *J* = 4.6 Hz, 1H), 5.25 (t, *J* = 5.5 Hz, 1H), 4.78 – 4.67 (m, 1H), 4.15 – 3.98 (m, 3H), 3.83 (dd, *J* = 8.9, 6.0 Hz, 1H), 3.73 – 3.62 (m, 1H), 3.60 – 3.51 (m, 1H), 2.56 (d, *J* = 4.6 Hz, 3H). ^13^C NMR (101 MHz, DMSO-*d*_6_) δ 162.2 (d, *J* = 246.9 Hz), 158.1, 154.2, 140.2 (d, *J* = 10.9 Hz), 133.8 (d, *J* = 2.7 Hz), 113.4 (d, *J* = 3.0 Hz), 104.9 (d, *J* = 15.6 Hz), 104.7 (d, *J* = 26.8 Hz), 93.3 (d, *J* = 3.0 Hz), 74.2, 73.5, 61.6, 45.9, 29.7, 26.5.

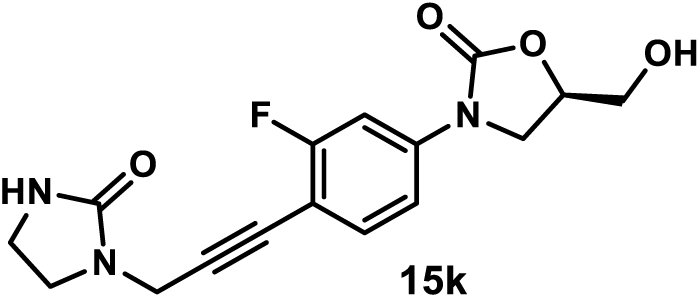

*(R)-3-(3-Fluoro-4-(3-(2-oxoimidazolidin-1-yl)prop-1-yn-1-yl)phenyl)-5-(hydroxymethyl)oxazolidin-2-one (**15k**)*. Using general procedure 4, employing 1-(2-propynyl)-2-imidazolidinone (19 mg, 0.15 mmol), compound **15k** was obtained after flash column chromatography (SiO_2_, 0-10% MeOH in DCM) as a yellow amorphous solid (10 mg, 30%). ^1^H NMR (400 MHz, DMSO-*d*_6_) δ 7.59 (dd, *J* = 12.4, 2.2 Hz, 1H), 7.53 (t, *J* = 8.5 Hz, 1H), 7.37 (dd, *J* = 8.5, 2.2 Hz, 1H), 6.61 (s, 1H), 5.22 (t, *J* = 5.6 Hz, 1H), 4.83 – 4.57 (m, 1H), 4.16 (s, 2H), 4.09 (t, *J* = 9.0 Hz, 1H), 3.84 (dd, *J* = 9.0, 6.0 Hz, 1H), 3.67 (ddd, *J* = 12.5, 5.6, 3.2 Hz, 1H), 3.56 (ddd, *J* = 12.5, 7.9, 5.6 Hz, 1H), 3.50 – 3.37 (m, 2H), 3.30 – 3.15 (m, 2H). ^13^C NMR (101 MHz, DMSO-*d*_6_) δ 162.2 (d, *J* = 247.2 Hz), 161.6, 154.2, 140.4 (d, *J* = 10.9 Hz), 133.8 (d, *J* = 2.6 Hz), 113.4 (d, *J* = 3.0 Hz), 104.6 (d, *J* = 42.8 Hz), 104.5, 89.7 (d, *J* = 2.9 Hz), 76.4, 73.4, 61.5, 45.9, 44.0, 37.2, 33.7.

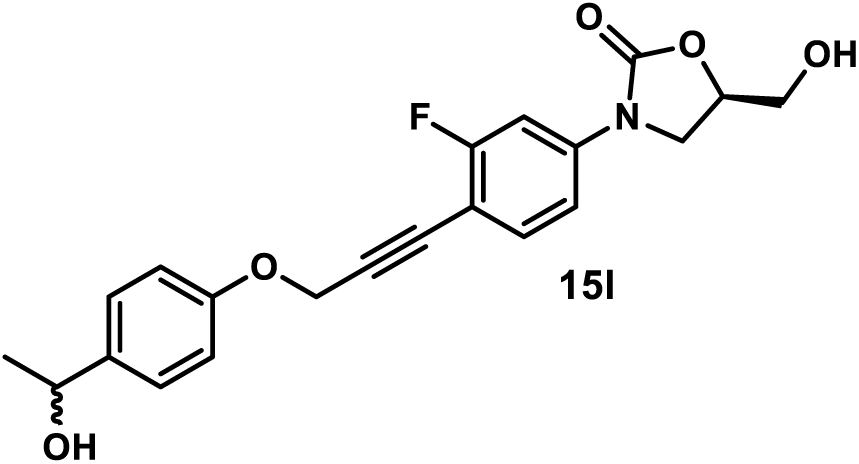

*(5R)-3-(3-Fluoro-4-(3-(4-(1-hydroxyethyl)phenoxy)prop-1-yn-1-yl)phenyl)-5-(hydroxymethyl)oxazolidin-2-one (**15l**)*. Using general procedure 4, employing 1-[4-(2-propynyloxy)phenyl]ethanol (26 mg, 0.15 mmol), compound **15l** was obtained after flash column chromatography (SiO_2_, 0-10% MeOH in DCM) as yellow oil (30 mg, 79%).^1^H NMR (300 MHz, Acetone-*d*_6_) δ 7.66 (dd, *J* = 12.4, 2.2 Hz, 1H), 7.49 (t, *J* = 8.3 Hz, 1H), 7.40 – 7.28 (m, 3H), 7.01 (d, *J* = 8.2 Hz, 2H), 5.02 (s, 2H), 4.88 – 4.71 (m, 2H), 4.42 (t, *J* = 5.8 Hz, 1H), 4.19 (t, *J* = 8.9 Hz, 1H), 4.09 (d, *J* = 4.0 Hz, 1H), 4.01 (dd, *J* = 8.9, 6.2 Hz, 1H), 3.93 – 3.84 (m, 1H), 3.81 – 3.68 (m, 1H), 1.37 (d, *J* = 6.4 Hz, 3H). ^13^C NMR (101 MHz, Acetone-*d*_6_) δ 163.7 (d, *J* = 248.3 Hz), 157.6, • 155.1 , 142.2 (d, *J* = 11.0 Hz), 141.1, 134.7 (d, *J* = 2.8 Hz), 127.3 (2C), 115.3 (2C), 113.9 (d, *J* = 3.1 Hz), 105.5 (d, *J* = 27.2 Hz), 105.5 (d, *J* = 16.2 Hz), 90.1 (d, *J* = 3.0 Hz), 80.4, 74.4, 69.5, 63.1, 57.0, 46.9, 26.2.

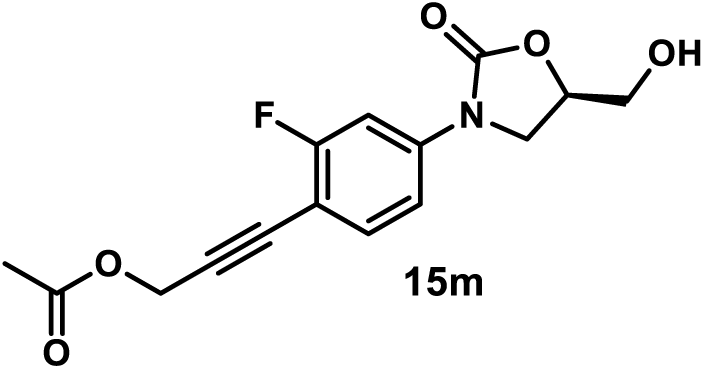

*(R)-3-(2-Fluoro-4-(5-(hydroxymethyl)-2-oxooxazolidin-3-yl)phenyl)prop-2-yn-1-yl Acetate (**15m**)*. Using general procedure 4, employing propargyl acetate (15 mg, 0.15 mmol), compound **15m** was obtained after flash column chromatography (SiO_2_, 0-100% EtOAc in hexanes) as a yellow amorphous solid (20 mg, 65%). ^1^H NMR (400 MHz, Acetone-*d*_6_) δ 7.67 (dd, *J* = 12.5, 2.3 Hz, 1H), 7.50 (t, *J* = 8.3 Hz, 1H), 7.39 (dd, *J* = 8.3, 2.3 Hz, 1H), 4.94 (s, 2H), 4.86 – 4.72 (m, 1H), 4.39 (t, *J* = 5.8 Hz, 1H), 4.21 (t, *J* = 8.9 Hz, 1H), 4.02 (dd, *J* = 8.9, 6.1 Hz, 1H), 3.90 (ddd, *J* = 12.4, 5.8, 3.3 Hz, 1H), 3.77 (ddd, *J* = 12.4, 5.8, 3.8 Hz, 1H), 2.09 (s, 3H). ^13^C NMR (101 MHz, Acetone-*d*_6_) δ 170.4, 163.8 (d, *J* = 248.2 Hz), 155.1, 142.3 (d, *J* = 10.9 Hz), 134.8 (d, *J* = 2.7 Hz), 114.0 (d, *J* = 3.1 Hz), 105.6 (d, *J* = 27.2 Hz), 105.4 (d, *J* = 16.1 Hz), 89.1 (d, *J* = 3.2 Hz), 79.7, 74.4, 63.1, 52.9, 46.9, 20.6.

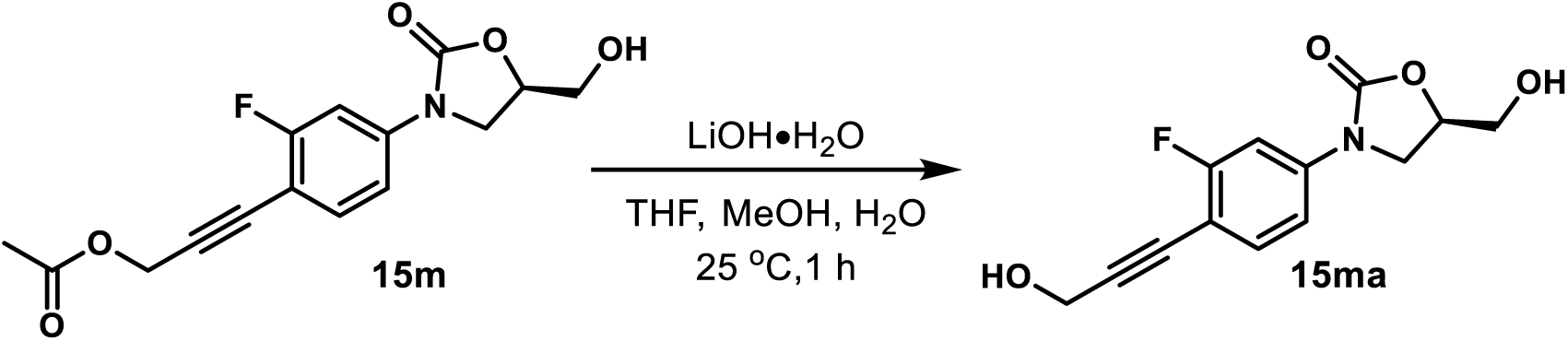

*(R)-3-(3-Fluoro-4-(3-hydroxyprop-1-yn-1-yl)phenyl)-5-(hydroxymethyl)oxazolidin-2-one (**15ma**)*. To a solution of compound **15m** (39 mg, 0.127 mmol) in THF (1.5 mL), MeOH (0.5 mL), and H_2_O (0.5 mL) was added lithium hydroxide monohydrate (53 mg, 1.27 mmol). The reaction mixture was stirred at 25 °C for 1 h, quenched with the addition of water, and extracted with EtOAc four times. The combined organic layers were concentrated under reduced pressure by rotary evaporation. The residue was purified by flash column chromatography (SiO_2_, 0-10% MeOH in DCM) followed by preparative TLC (SiO_2_, 10% MeOH in DCM) to give compound **15ma** as a yellow amorphous solid (16 mg, 47%). ^1^H NMR (400 MHz, Methanol-*d*_4_) δ 7.56 (d, *J* = 12.1 Hz, 1H), 7.42 (t, *J* = 8.6 Hz, 1H), 7.26 (d, *J* = 8.6 Hz, 1H), 4.77 – 4.66 (m, 1H), 4.39 (s, 2H), 4.20 – 4.03 (m, 1H), 3.95 – 3.86 (m, 1H), 3.85 – 3.78 (m, 1H), 3.70 – 3.63 (m, 1H). ^13^C NMR (101 MHz, Methanol-*d*_4_) δ 164.2 (d, *J* = 248.6 Hz), 156.6, 141.6 (d, *J* = 10.8 Hz), 134.9 (d, *J* = 2.8 Hz), 114.4 (d, *J* = 3.2 Hz), 107.4 (d, *J* = 16.1 Hz), 106.4 (d, *J* = 27.3 Hz), 93.7 (d, *J* = 3.0 Hz), 78.4, 75.2, 63.2, 51.2, 47.5.

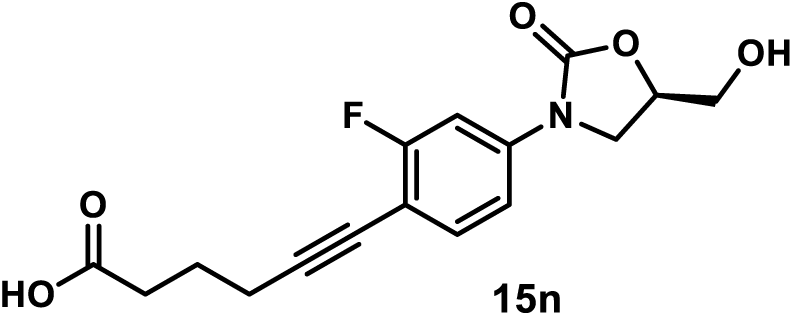

*(R)-6-(2-Fluoro-4-(5-(hydroxymethyl)-2-oxooxazolidin-3-yl)phenyl)hex-5-ynoic Acid (**15n**)*. A solution of compound **9** (34 mg, 0.1 mmol), employing 5-hexynoic acid (17 mg, 0.15 mmol), tetrakis(triphenylphosphine)palladium(0) (12 mg, 0.01 mmol), and copper(I) iodide (2 mg, 0.01 mmol) in CH_3_CN (1 mL) was evacuated and purged with nitrogen three times. *N*,*N*-diisopropylethylamine (0.5 mL) was added under the protection of nitrogen atmosphere. The mixture was stirred at 25 °C. After the reaction was judged to be complete by TLC (16 h), its solvent was concentrated under reduced pressure by rotary evaporation. The residue was diluted with MeOH (10 mL), acidified with formic acid (2 mL), and concentrated under reduced pressure by rotary evaporator. The residue was purified by flash column chromatography (SiO_2_, 10-20% MeOH in DCM with 0.1% formic acid) to give compound **15n** as a yellow amorphous solid (10 mg, 31%). ^1^H NMR (400 MHz, Methanol-*d*_4_) δ 7.51 (dd, *J* = 12.1, 2.3 Hz, 1H), 7.34 (t, *J* = 8.6 Hz, 1H), 7.20 (dd, *J* = 8.6, 2.3 Hz, 1H), 4.75 – 4.63 (m, 1H), 4.06 (t, *J* = 8.9 Hz, 1H), 3.87 (dd, *J* = 8.9, 6.3 Hz, 1H), 3.80 (dd, *J* = 12.5, 3.2 Hz, 1H), 3.64 (dd, *J* = 12.5, 3.9 Hz, 1H), 2.53 – 2.36 (m, 4H), 1.91 – 1.76 (m, 2H). ^13^C NMR (101 MHz, Methanol-*d*_4_) δ 177.4, 164.1 (d, *J* = 247.8 Hz), 156.7, 140.9 (d, *J* = 10.6 Hz), 134.7 (d, *J* = 2.9 Hz), 114.4 (d, *J* = 3.4 Hz), 108.4 (d, *J* = 16.5 Hz), 106.4 (d, *J* = 27.4 Hz), 95.0 (d, *J* = 3.0 Hz), 75.2, 75.0, 63.2, 47.5, 34.2, 25.3, 19.6.

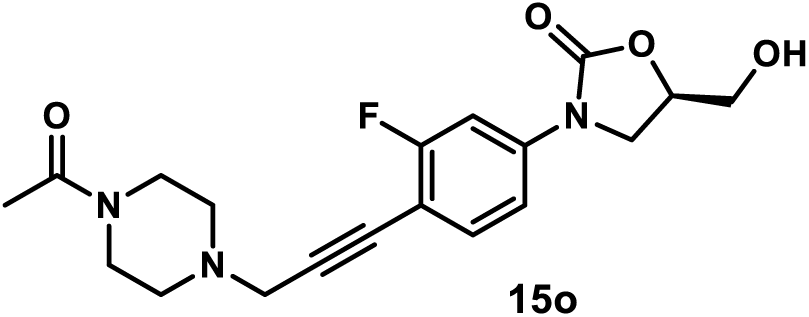

*(R)-3-(4-(3-(4-Acetylpiperazin-1-yl)prop-1-yn-1-yl)-3-fluorophenyl)-5-(hydroxymethyl)oxazolidin-2-one (**15o**)*. Using general procedure 4, employing 1-(4-(prop-2-yn-1-yl)piperazin-1-yl)ethan-1-one (25 mg, 0.15 mmol), compound **15o** was obtained after flash column chromatography (SiO_2_, 0-15% MeOH in DCM) as a white amorphous solid (27 mg, 71%). ^1^H NMR (400 MHz, DMSO-*d*_6_) δ 7.64 (dd, *J* = 12.3, 2.2 Hz, 1H), 7.56 (t, *J* = 8.4 Hz, 1H), 7.41 (dd, *J* = 8.4, 2.1 Hz, 1H), 5.29 (t, *J* = 5.5 Hz, 1H), 4.89 – 4.63 (m, 1H), 4.13 (t, *J* = 9.0 Hz, 1H), 3.88 (dd, *J* = 9.0, 6.1 Hz, 1H), 3.72 (ddd, *J* = 12.4, 5.5, 3.2 Hz, 1H), 3.69 – 3.55 (m, 3H), 3.55 – 3.45 (m, 4H), 2.73 – 2.52 (m, 4H), 2.03 (s, 3H).^13^C NMR (151 MHz, DMSO-*d*_6_) δ 168.1, 162.1 (d, *J* = 246.9 Hz), 154.2, 140.2 (d, *J* = 10.8 Hz), 133.7 (d, *J* = 2.7 Hz), 113.3 (d, *J* = 3.0 Hz), 104.7 (d, *J* = 16.1 Hz), 104.7 (d, *J* = 26.9 Hz), 89.8 (d, *J* = 3.0 Hz), 78.0, 73.4, 61.5, 51.5, 51.0, 46.7, 45.9, 45.5, 40.6, 21.2.

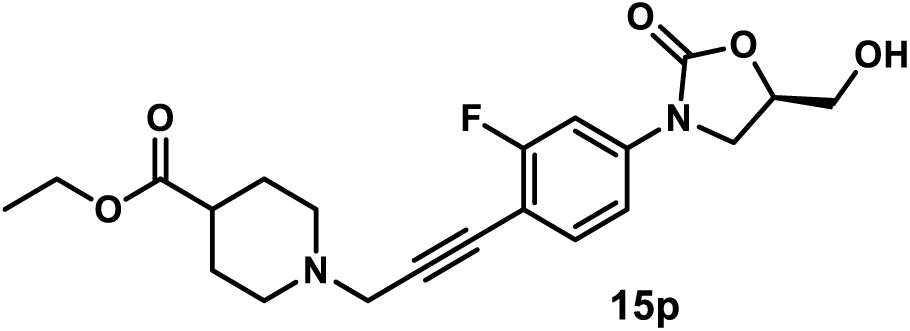

*Ethyl (R)-1-(3-(2-Fluoro-4-(5-(hydroxymethyl)-2-oxooxazolidin-3-yl)phenyl)prop-2-yn-1-yl)piperidine-4-carboxylate (**15p**)*. Using general procedure 4, employing ethyl 1-(prop-2-yn-1-yl) piperidine-4-carboxylate (29 mg, 0.15 mmol), compound **15p** was obtained after flash column chromatography (SiO_2_, 0-15% MeOH in DCM) as a yellow amorphous solid (23 mg, 57%). ^1^H NMR (400 MHz, DMSO-*d*_6_) δ 7.59 (d, *J* = 12.4 Hz, 1H), 7.51 (t, *J* = 8.6 Hz, 1H), 7.36 (d, *J* = 8.6 Hz, 1H), 5.24 (t, *J* = 5.6 Hz, 1H), 4.87 – 4.61 (m, 1H), 4.17 – 4.00 (m, 3H), 3.84 (t, *J* = 7.6 Hz, 1H), 3.67 (dt, *J* = 8.5, 4.1 Hz, 1H), 3.60 – 3.45 (m, 3H), 2.90 – 2.70 (m, 2H), 2.44 – 2.10 (m, 3H), 1.90 – 1.70 (m, 2H), 1.70 – 1.46 (m, 2H), 1.17 (t, *J* = 7.1 Hz, 3H). ^13^C NMR (101 MHz, DMSO-*d*_6_) δ 174.3, 162.1 (d, *J* = 246.9 Hz), 154.2, 140.1 (d, *J* = 10.9 Hz), 133.7 (d, *J* = 2.6 Hz), 113.3 (d, *J* = 3.0 Hz), 104.9 (d, *J* = 16.1 Hz), 104.7 (d, *J* = 27.0 Hz), 90.3 (d, *J* = 3.1 Hz), 77.8, 73.4, 61.6, 59.8, 51.0 (2C), 47.2, 45.9, 39.8, 27.9 (2C), 14.1.

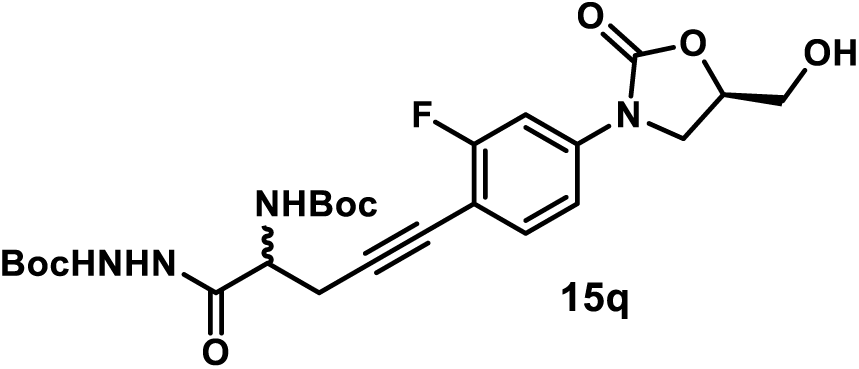

*tert-Butyl 2-(2-((tert-Butoxycarbonyl)amino)-5-(2-fluoro-4-((R)-5-(hydroxymethyl)-2-oxooxazolidin-3-yl)phenyl)pent-4-ynoyl)hydrazine-1-carboxylate (**15q**)*. A solution of compound **9** (34 mg, 0.1 mmol), compound **21** (49 mg, 0.15 mmol), tetrakis(triphenylphosphine)palladium(0) (30 mg, 0.025 mmol), and copper(I) iodide (4 mg, 0.02 mmol) in DMF (1 mL) was evacuated and purged with nitrogen three times. *N*,*N*-diisopropylethylamine (0.5 mL) was added under the protection of nitrogen atmosphere. The mixture was stirred at 50 °C. After the reaction was judged to be completed by TLC (12 h), it was diluted with EtOAc, washed with water three times, and concentrated under reduced pressure by rotary evaporation. The crude residue was purified by flash column chromatography (SiO_2_ eluent gradient 0-100% EtOAc in hexanes) followed by preparative TLC purification (SiO_2_, eluent 100% EtOAc) to afford compound **15q** as a white amorphous solid (32 mg, 59%). ^1^H NMR (400 MHz, Acetone-*d*_6_) δ 9.17 (s, 1H), 7.97 (s, 1H), 7.63 (dd, *J* = 12.3, 2.3 Hz, 1H), 7.47 (t, *J* = 8.5 Hz, 1H), 7.33 (dd, *J* = 8.5, 2.3 Hz, 1H), 6.29 (d, *J* = 8.8 Hz, 1H), 4.87 – 4.74 (m, 1H), 4.55 – 4.29 (m, 2H), 4.19 (t, *J* = 8.9 Hz, 1H), 4.00 (dd, *J* = 8.9, 6.2 Hz, 1H), 3.89 (dd, *J* = 12.3, 3.3 Hz, 1H), 3.76 (dd, *J* = 12.3, 3.8 Hz, 1H), 2.98 (dd, *J* = 17.0, 5.4 Hz, 1H), 2.86 (dd, *J* = 17.0, 8.0 Hz, 1H), 1.51 – 1.34 (m, 18H). ^13^C NMR (101 MHz, Acetone-*d*_6_) δ 170.9, 163.5 (d, *J* = 247.4 Hz), 156.2, 156.1, 155.1, 141.3 (d, *J* = 10.9 Hz), 134.8, 113.7 (d, *J* = 3.2 Hz), 106.8 (d, *J* = 16.0 Hz), 105.4 (d, *J* = 27.4 Hz), 91.3, 80.5, 79.7, 74.3, 63.1, 55.4, 52.9, 46.9, 28.5, 28.4, 24.2.

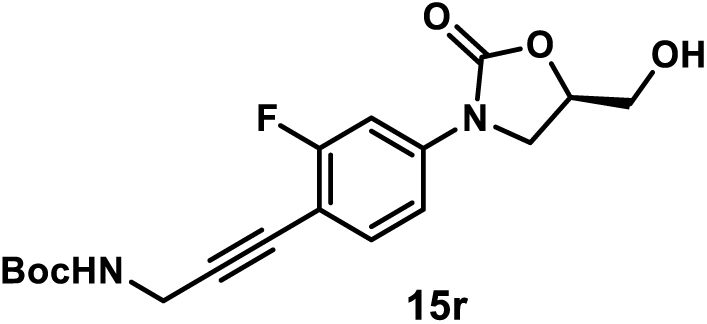

*tert-Butyl (R)-(3-(2-Fluoro-4-(5-(hydroxymethyl)-2-oxooxazolidin-3-yl)phenyl)prop-2-yn-1-yl)carbamate (**15r**)*. A solution of compound **9** (67 mg, 0.2 mmol), **22** (37 mg, 0.24 mmol), tetrakis(triphenylphosphine)palladium(0) (23 mg, 0.02 mmol), and copper(I) iodide (4 mg, 0.02 mmol) in DMF (2 mL) was evacuated and purged with nitrogen three times. *N*,*N*-diisopropylethylamine (1 mL) was added under the protection of nitrogen atmosphere. The mixture was stirred at 50 °C for 3 h, diluted with EtOAc, washed with water three times, and concentrated under reduced pressure by rotary evaporation. The crude residue was purified by flash column chromatography (SiO_2_, eluent gradient 0-10% MeOH in DCM) followed by preparative TLC purification (SiO_2_, eluent 100% EtOAc) to give **15r** as a yellow amorphous solid (64 mg, 88%). ^1^H NMR (400 MHz, Acetone-*d*_6_) δ 7.64 (dd, *J* = 12.4, 2.3 Hz, 1H), 7.45 (t, *J* = 8.5 Hz, 1H), 7.36 (dd, *J* = 8.5, 2.3 Hz, 1H), 6.48 (s, 1H), 4.88 – 4.72 (m, 1H), 4.40 (t, *J* = 5.8 Hz, 1H), 4.20 (t, *J* = 8.9 Hz, 1H), 4.13 (d, *J* = 5.8 Hz, 2H), 4.01 (dd, *J* = 8.9, 6.2 Hz, 1H), 3.89 (ddd, *J* = 12.3, 5.8, 3.2 Hz, 1H), 3.76 (ddd, *J* = 12.3, 5.8, 3.8 Hz, 1H), 1.42 (s, 9H). ^13^C NMR (101 MHz, Acetone-*d*_6_) δ 163.6 (d, *J* = 247.7 Hz), 156.3, 155.1, 141.7 (d, *J* = 10.8 Hz), 134.6 (d, *J* = 2.9 Hz), 113.9 (d, *J* = 3.1 Hz), 106.3 (d, *J* = 16.2 Hz), 105.5 (d, *J* = 27.5 Hz), 92.3, 79.3, 75.6, 74.3, 63.1, 46.9, 31.3, 28.6.

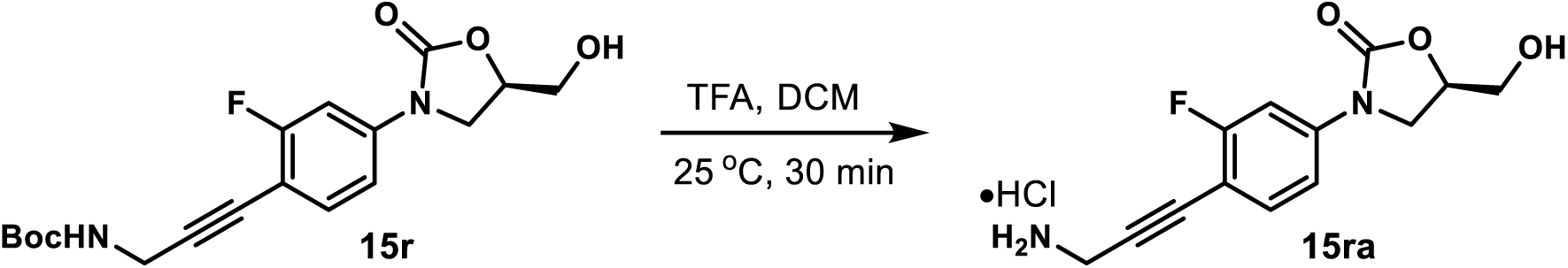

*(R)-3-(4-(3-Aminoprop-1-yn-1-yl)-3-fluorophenyl)-5-(hydroxymethyl)oxazolidin-2-one Hydrochloride (**15ra**)*. A solution of **15r** (8 mg, 0.022 mmol) in TFA (0.2 mL) and DCM (0.6 mL) was stirred at 25 °C. After the reaction was judged to be completed by TLC (30 min), its solvent was removed under reduced pressure by rotary evaporation. The residue was dissolved in MeOH (1 mL), charged with the addition of HCl/dioxane (4 M, 20 µL), and concentrated under reduced pressure by rotary evaporation. The resulting residue was washed with acetone (1 mL) and dried under high vacuum to give compound **15ra** as a yellow amorphous solid (6 mg, 91%). ^1^H NMR (400 MHz, Methanol-*d*_4_) δ 7.65 (dd, *J* = 12.2, 2.3 Hz, 1H), 7.52 (t, *J* = 8.3 Hz, 1H), 7.35 (dd, *J* = 8.3, 2.3 Hz, 1H), 4.84 – 4.68 (m, 1H), 4.14 (t, *J* = 9.0 Hz, 1H), 4.08 (s, 2H), 3.94 (dd, *J* = 9.0, 6.2 Hz, 1H), 3.87 (dd, *J* = 12.6, 3.1 Hz, 1H), 3.69 (dd, *J* = 12.6, 3.8 Hz, 1H). ^13^C NMR (101 MHz, Methanol-*d*_4_) δ 164.5 (d, *J* = 249.4 Hz), 156.6, 142.6 (d, *J* = 10.9 Hz), 135.1 (d, *J* = 2.6 Hz), 114.6 (d, *J* = 3.1 Hz), 106.4 (d, *J* = 27.1 Hz), 105.7 (d, *J* = 16.0 Hz), 86.1 (d, *J* = 2.9 Hz), 81.0, 75.2, 63.1, 47.4, 30.8.

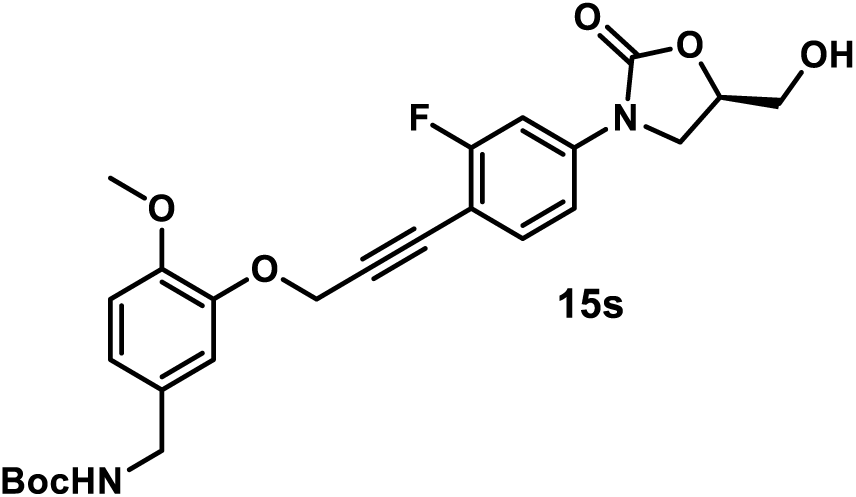

*tert-Butyl (R)-(3-((3-(2-Fluoro-4-(5-(hydroxymethyl)-2-oxooxazolidin-3-yl)phenyl)prop-2-yn-1-yl)oxy)-4-methoxybenzyl)carbamate (**15s**)*. A solution of compound **9** (34 mg, 0.1 mmol), **23** (44 mg, 0.15 mmol), tetrakis(triphenylphosphine)palladium(0) (12 mg, 0.01 mmol), and copper(I) iodide (2 mg, 0.01 mmol) in DMF (1 mL) was evacuated and purged with nitrogen three times. *N*,*N*-diisopropylethylamine (0.5 mL) was added under the protection of nitrogen atmosphere. The mixture was stirred at 50 °C. After the reaction was judged to be completed by TLC (3 h), it was diluted with EtOAc, washed with water three times, and concentrated under reduced pressure by rotary evaporation. The crude residue was purified by flash column chromatography (SiO_2_, eluent gradient 0-100% EtOAc in hexanes) followed by preparative TLC purification (SiO_2_, eluent 100% EtOAc) to give **15s** as a yellow amorphous solid (35 mg, 70%). ^1^H NMR (300 MHz, acetone) δ 7.65 (dd, *J* = 12.4, 2.2 Hz, 1H), 7.51 (t, *J* = 8.7 Hz, 1H), 7.37 (dd, *J* = 8.7, 2.2 Hz, 1H), 7.15 – 7.10 (m, 1H), 6.99 – 6.85 (m, 2H), 6.38 (s, 1H), 5.00 (s, 2H), 4.89 – 4.69 (m, 1H), 4.41 (t, *J* = 5.9 Hz, 1H), 4.27 – 4.12 (m, 3H), 4.01 (dd, *J* = 8.9, 6.2 Hz, 1H), 3.89 (ddd, *J* = 12.4, 5.5, 3.3 Hz, 1H), 3.84 – 3.67 (m, 4H), 1.41 (s, 9H). ^13^C NMR (75 MHz, Acetone-*d*_6_) δ 163.7 (d, *J* = 248.4 Hz), 156.8, 155.1, 150.0, 148.0, 142.1 (d, *J* = 10.9 Hz), 134.9 (d, *J* = 2.2 Hz), 133.6, 122.0, 115.6 (d, *J* = 1.5 Hz), 113.9 (d, J = 2.2 Hz), 112.9, 105.6 (d, *J* = 15.9 Hz), 105. 3 (d, *J* = 2.2 Hz), 90.1 (d, *J* = 2.9 Hz), 80.6, 78.7, 74.4, 63.0, 58.1, 56.13, 56.08, 46.9, 44.4.

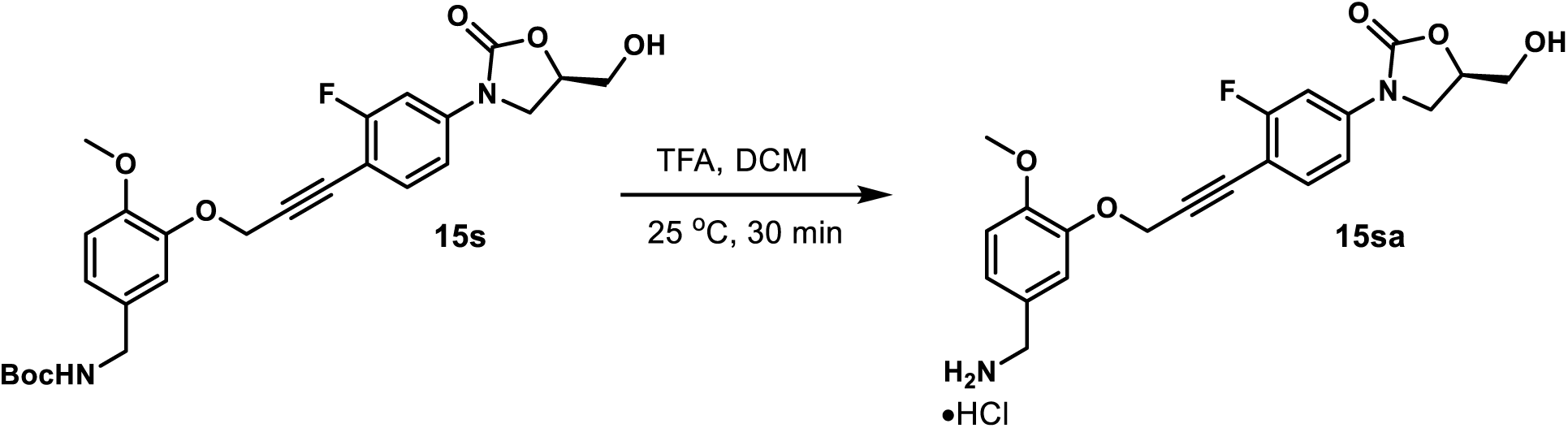

*(R)-3-(4-(3-(5-(Aminomethyl)-2-methoxyphenoxy)prop-1-yn-1-yl)-3-fluorophenyl)-5-(hydroxymethyl)oxazolidin-2-one Hydrochloride (**15sa**)*. A solution of **15s** (30 mg, 0.06 mmol) in TFA (0.3 mL) and DCM (0.9 mL) was stirred at 25 °C. After the reaction was judged to be completed by TLC (30 min), its solvent was removed under reduced pressure by rotary evaporation. The residue was dissolved in MeOH (2 mL), charged with the addition of HCl/dioxane (4 M, 80 µL), and concentrated under reduced pressure by rotary evaporation. The resulting residue was washed with acetone (2 mL) and dried under high vacuum to give compound **15sa** as a yellow amorphous solid (10 mg, 40%). ^1^H NMR (400 MHz, Methanol-*d*_4_) δ 7.62 (dd, *J* = 12.2, 2.2 Hz, 1H), 7.45 (t, *J* = 8.6 Hz, 1H), 7.30 (dd, *J* = 8.6, 2.2 Hz, 1H), 7.24 (d, *J* = 2.0 Hz, 1H), 7.13 – 7.01 (m, 2H), 5.04 (s, 2H), 4.80 – 4.71 (m, 1H), 4.12 (t, *J* = 9.1 Hz, 1H), 4.06 (s, 2H), 3.97 – 3.77 (m, 5H), 3.68 (dd, *J* = 12.5, 3.8 Hz, 1H). ^13^C NMR (101 MHz, Methanol-*d*_4_) δ 164.4 (d, *J* = 248.8 Hz), 156.6, 152.3, 148.6, 142.1 (d, *J* = 10.9 Hz), 135.0 (d, *J* = 2.8 Hz), 126.7, 124.4, 117.4, 114.5 (d, *J* = 3.2 Hz), 113.6, 106.6 (d, *J* = 16.1 Hz), 106.3 (d, *J* = 27.3 Hz), 89.8 (d, *J* = 2.9 Hz), 81.2, 75.2, 63.1, 58.8, 56.5, 47.4, 44.1.

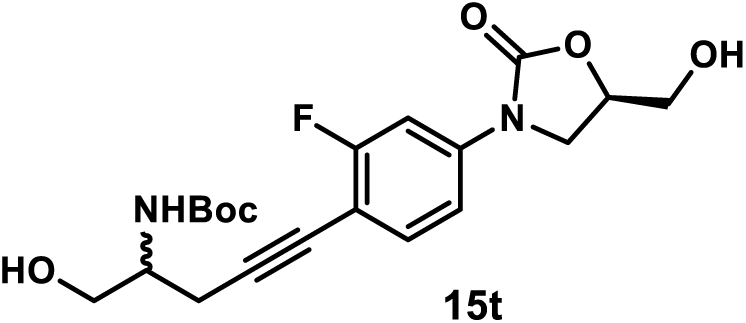

*tert-Butyl (5-(2-Fluoro-4-((R)-5-(hydroxymethyl)-2-oxooxazolidin-3-yl)phenyl)-1-hydroxypent-4-yn-2-yl)carbamate (**15t**)*. A solution of compound **9** (52 mg, 0.155 mmol), **24** (37 mg, 0.186 mmol), tetrakis(triphenylphosphine)palladium(0) (18 mg, 0.0155 mmol), and copper(I) iodide (3 mg, 0.0155 mmol) in DMF (2 mL) was evacuated and purged with nitrogen three times. *N*,*N*-diisopropylethylamine (1 mL) was added under the protection of nitrogen atmosphere. The mixture was stirred at 50 °C. After the reaction was judged to be completed by TLC (5 h), it was diluted with EtOAc, washed with water three times, and concentrated under reduced pressure by rotary evaporation. The crude residue was purified by flash column chromatography (SiO_2_, eluent gradient 0-10% MeOH in DCM) followed by preparative TLC purification (SiO_2_, eluent 100% EtOAc) to give **15t** as a white amorphous solid (34 mg, 54%). ^1^H NMR (400 MHz, Acetone-*d*_6_) δ 7.63 (dd, *J* = 12.4, 2.3 Hz, 1H), 7.44 (t, *J* = 8.6 Hz, 1H), 7.34 (dd, *J* = 8.6, 2.3 Hz, 1H), 5.90 (d, *J* = 8.3 Hz, 1H), 4.89 – 4.75 (m, 1H), 4.40 (t, *J* = 5.8 Hz, 1H), 4.19 (t, *J* = 9.0 Hz, 1H), 4.11 – 3.95 (m, 2H), 3.94 – 3.58 (m, 5H), 2.86 – 2.65 (m, 2H), 1.40 (s, 9H). ^13^C NMR (101 MHz, Acetone-*d*_6_) δ 163.6 (d, *J* = 246.9 Hz), 156.3, 155.1, 141.2 (d, *J* = 10.7 Hz), 134.6 (d, *J* = 2.9 Hz), 113.8 (d, *J* = 3.2 Hz), 107.1 (d, *J* = 16.3 Hz), 105.5 (d, *J* = 27.4 Hz), 92.6 (d, *J* = 3.0 Hz), 78.9, 75.5, 74.3, 63.5, 63.1, 52.7, 46.9, 28.6, 22.7.

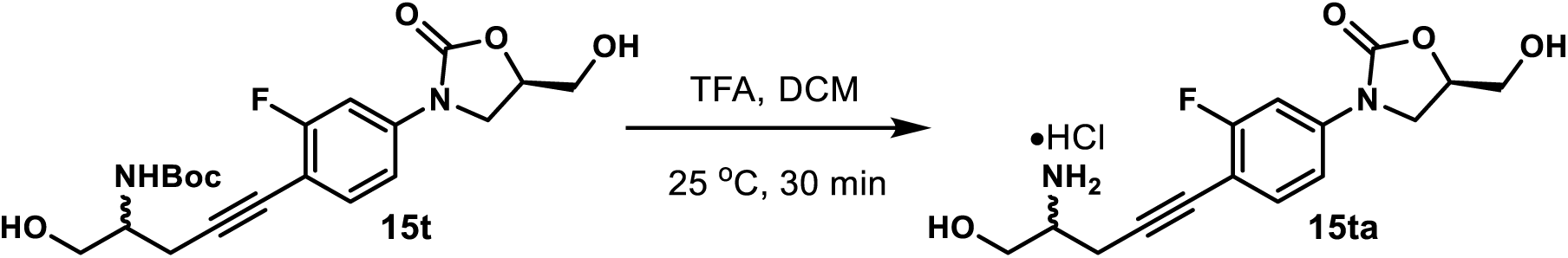

*(5R)-3-(4-(4-Amino-5-hydroxypent-1-yn-1-yl)-3-fluorophenyl)-5-(hydroxymethyl)oxazolidin-2-one Hydrochloride (**15ta**).* A solution of **15t** (16 mg, 0.039 mmol) in TFA (0.2 mL) and DCM (0.6 mL) was stirred at 25 °C. After the reaction was judged to be completed by TLC (30 min), its solvent was removed under reduced pressure by rotary evaporation. The residue was dissolved in MeOH (1 mL), charged with the addition of HCl/dioxane (4 M, 40 µL), and concentrated under reduced pressure by rotary evaporation. The resulting residue was washed with acetone (1 mL) and dried under high vacuum to give compound **15ta** as a yellow amorphous solid (7 mg, 52%). ^1^H NMR (400 MHz, Methanol-*d*_4_) δ 7.61 (dd, *J* = 12.1, 2.3 Hz, 1H), 7.48 (t, *J* = 8.3 Hz, 1H), 7.31 (dd, *J* = 8.3, 2.3 Hz, 1H), 4.83 – 4.71 (m, 1H), 4.13 (t, *J* = 9.0 Hz, 1H), 3.97 – 3.83 (m, 3H), 3.78 (dd, *J* = 11.6, 6.0 Hz, 1H), 3.69 (dd, *J* = 11.6, 3.8 Hz, 1H), 3.56 – 3.42 (m, 1H), 2.89 (dd, *J* = 6.7, 4.1 Hz, 2H). ^13^C NMR (101 MHz, Methanol-*d*_4_) δ 164.4 (d, *J* = 248.4 Hz), 156.6, 141.7 (d, *J* = 10.8 Hz), 134.9 (d, *J* = 2.6 Hz), 114.4 (d, *J* = 3.3 Hz), 107.1 (d, *J* = 16.2 Hz), 106.3 (d, *J* = 27.3 Hz), 88.9 (d, *J* = 3.0 Hz), 77.9, 75.2, 63.1, 61.6, 53.2, 47.5, 21.1.

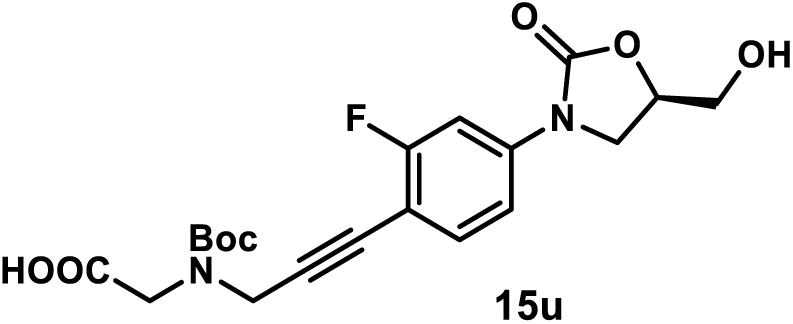

*(R)-N-(tert-Butoxycarbonyl)-N-(3-(2-fluoro-4-(5-(hydroxymethyl)-2-oxooxazolidin-3-yl)phenyl)prop-2-yn-1-yl)glycine (**15u**)*. A solution of compound **9** (82 mg, 0.24 mmol), **25** (62 mg, 0.29 mmol), tetrakis(triphenylphosphine)palladium(0) (28 mg, 0.024 mmol), and copper(I) iodide (5 mg, 0.024 mmol) in CH_3_CN (2 mL) was evacuated and purged with nitrogen three times. *N*,*N*-diisopropylethylamine (1 mL) was added under the protection of nitrogen atmosphere. The mixture was stirred at 50 °C. After the reaction was judged to be completed by TLC (4 h), its solvent was concentrated under reduced pressure by rotary evaporation. The residue was dissolved in acetone, acidified with the addition of formic acid (1 mL), and concentrated under reduced pressure by rotary evaporation. The crude residue was purified by flash column chromatography (SiO_2_, eluent gradient 0-10% MeOH in DCM with 1% formic acid) followed by preparative TLC purification (SiO_2_, eluent 10% MeOH in DCM) to give **15u** as a white amorphous solid (84 mg, 83%). ^1^H NMR (600 MHz, Acetonitrile-*d*_3_) δ 7.54 (dd, *J* = 12.4, 2.3 Hz, 1H), 7.44 (t, *J* = 8.3 Hz, 1H), 7.29 (dd, *J* = 8.3 Hz, 2.3 Hz, 1H), 4.76 – 4.64 (m, 1H), 4.41 – 4.30 (m, 2H), 4.09 – 3.97 (m, 3H), 3.83 (dd, *J* = 8.9, 6.1 Hz, 1H), 3.78 (dd, *J* = 12.5, 3.3 Hz, 1H), 3.64 (dd, *J* = 12.5, 4.2 Hz, 1H), 1.57 – 1.19 (m, 9H). ^13^C NMR (101 MHz, Acetonitrile-*d*_3_) δ 172.5, 163.7 (d, *J* = 247.3 Hz), 155.8, 155.5, 141.6 (d, *J* = 11.0 Hz), 134.8 (d, *J* = 2.0 Hz), 114.2 (d, *J* = 3.2 Hz), 105.8 (d, *J* = 27.2 Hz), 90.4 (d, *J* = 3.3 Hz), 81.3, 81.2, 77.5, 77.3, 74.4, 63.0, 48.6, 48.5, 48.4, 47.1, 39.0, 38.1, 28.4, 28.3.

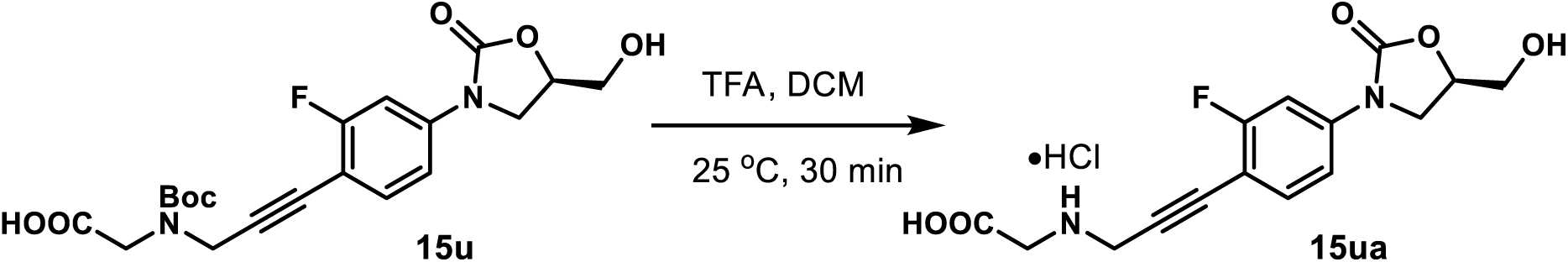

*(R)-(3-(2-Fluoro-4-(5-(hydroxymethyl)-2-oxooxazolidin-3-yl)phenyl)prop-2-yn-1-yl)glycine Hydrochloride (**15ua**)*. A solution of **15u** (10 mg, 0.024 mmol) in TFA (0.2 mL) and DCM (0.6 mL) was stirred at 25 °C. After the reaction was judged to be completed by TLC (30 min), its solvent was removed under reduced pressure by rotary evaporation. The residue was dissolved in MeOH (1 mL), charged with the addition of HCl/dioxane (4 M, 20 µL), and concentrated under reduced pressure by rotary evaporation. The resulting residue was washed with acetone (1 mL) and dried under high vacuum to give compound **15ua** as a yellow amorphous solid (6 mg, 70%). ^1^H NMR (400 MHz, Methanol-*d*_4_) δ 7.67 (dd, *J* = 12.2, 2.2 Hz, 1H), 7.55 (t, *J* = 8.3 Hz, 1H), 7.36 (dd, *J* = 8.3, 2.2 Hz, 1H), 4.84 – 4.69 (m, 1H), 4.28 (s, 2H), 4.14 (t, *J* = 9.0 Hz, 1H), 4.06 (s, 2H), 3.95 (dd, *J* = 9.0, 6.2 Hz, 1H), 3.87 (dd, *J* = 12.6, 3.1 Hz, 1H), 3.70 (dd, *J* = 12.6, 3.8 Hz, 1H). ^13^C NMR (101 MHz, Methanol-*d*_4_) δ 168.7, 164.6 (d, *J* = 249.6 Hz), 156.5, 142.8 (d, *J* = 11.0 Hz), 135.2 (d, *J* = 2.4 Hz), 114.6 (d, *J* = 3.1 Hz), 106.3 (d, *J* = 27.1 Hz), 105.3 (d, *J* = 16.1 Hz), 83.9 (d, *J* = 3.0 Hz), 83.0, 75.2, 63.1, 47.4, 47.3, 38.0.

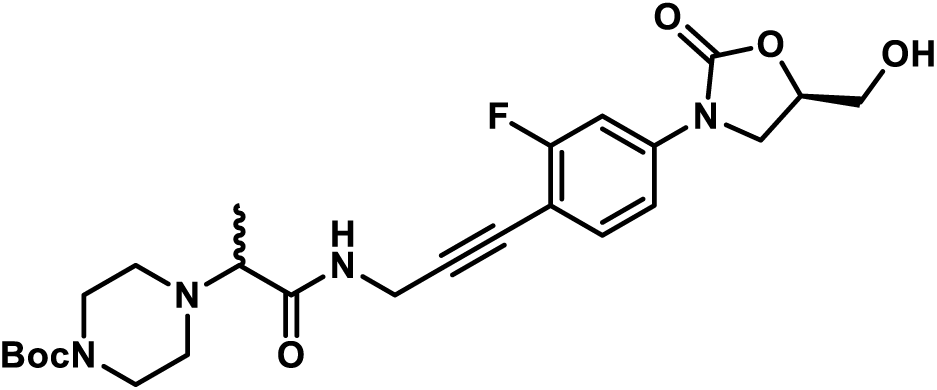

*tert-Butyl 4-(1-((3-(2-Fluoro-4-((R)-5-(hydroxymethyl)-2-oxooxazolidin-3-yl)phenyl)prop-2-yn-1-yl)amino)-1-oxopropan-2-yl)piperazine-1-carboxylate (**15v**).* A solution of compound **9** (34 mg, 0.1 mmol), **26** (35 mg, 0.12 mmol),, tetrakis(triphenylphosphine)palladium(0) (12 mg, 0.01 mmol), and copper(I) iodide (2 mg, 0.01 mmol) in DMF (2 mL) was evacuated and purged with nitrogen three times. *N*,*N*-diisopropylethylamine (1 mL) was added under the protection of nitrogen atmosphere. The mixture was stirred at 50 °C. After the reaction was judged to be completed by TLC (2 h), it was diluted with EtOAc, washed with water three times, and concentrated under reduced pressure by rotary evaporation. The crude residue was purified by flash column chromatography (SiO_2_, eluent gradient 0-10% MeOH in DCM) followed by preparative TLC purification (SiO_2_, eluent 100% EtOAc) to give **15v** as a white amorphous solid (45 mg, 89%). ^1^H NMR (400 MHz, Acetone-*d*_6_) δ 7.93 (t, *J* = 5.7 Hz, 1H), 7.64 (dd, *J* = 12.4, 2.2 Hz, 1H), 7.43 (t, *J* = 8.7 Hz, 1H), 7.35 (dd, *J* = 8.7, 2.2 Hz, 1H), 4.90 – 4.74 (m, 1H), 4.43 (s, 1H), 4.25 (d, *J* = 5.7 Hz, 2H), 4.19 (t, *J* = 8.9 Hz, 1H), 4.00 (dd, *J* = 8.9, 6.2 Hz, 1H), 3.94 – 3.85 (m, 1H), 3.79 – 3.71 (m, 1H), 3.47 – 3.34 (m, 4H), 3.17 (q, *J* = 6.9 Hz, 1H), 2.57 – 2.35 (m, 4H), 1.42 (s, 9H), 1.17 (d, *J* = 6.9 Hz, 3H). ^13^C NMR (101 MHz, Acetone-*d*_6_) δ 173.2, 163.6 (d, *J* = 247.4 Hz), 155.1, 154.9, 141.7 (d, *J* = 10.9 Hz), 134.5 (d, *J* = 2.7 Hz), 113.9 (d, *J* = 3.1 Hz), 106.2 (d, *J* = 16.3 Hz), 105.5 (d, *J* = 27.3 Hz), 92.1 (d, *J* = 3.1 Hz), 79.5, 75.4, 74.3, 64.4, 63.1, 50.2, 46.9, 29.6, 28.5, 11.7.

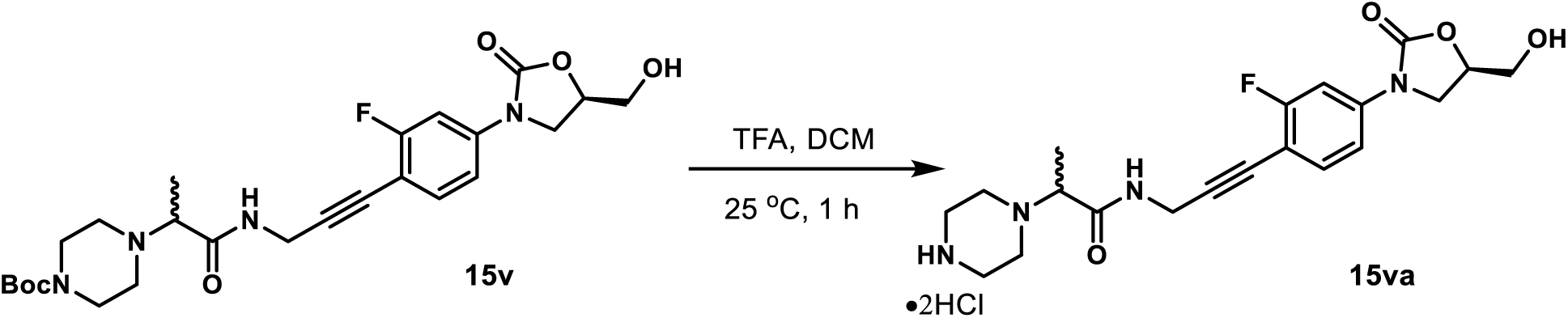

*N-(3-(2-Fluoro-4-((R)-5-(hydroxymethyl)-2-oxooxazolidin-3-yl)phenyl)prop-2-yn-1-yl)-2-(piperazin-1-yl)propenamide Dihydrochloride (**15va**).* A solution of **15v** (40 mg, 0.079 mmol) in TFA (0.8 mL) and DCM (2.4 mL) was stirred at 25 °C. After the reaction was judged to be completed by TLC (1 h), its solvent was removed under reduced pressure by rotary evaporation. The residue was dissolved in MeOH (4 mL), charged with the addition of HCl/dioxane (4 M, 80 µL), and concentrated under reduced pressure by rotary evaporation. The resulting residue was washed with acetone (4 mL) and dried under high vacuum to give compound **15va** as a yellow amorphous solid (22 mg, 58%). ^1^H NMR (400 MHz, Methanol-*d*_4_) δ 7.61 (dd, *J* = 12.2, 2.2 Hz, 1H), 7.44 (t, *J* = 8.3 Hz, 1H), 7.29 (dd, *J* = 8.3, 2.2 Hz, 1H), 4.83 – 4.70 (m, 1H), 4.31 (d, *J* = 2.0 Hz, 2H), 4.12 (t, *J* = 8.9 Hz, 1H), 4.02 (q, *J* = 6.9 Hz, 1H), 3.93 (dd, *J* = 8.9, 6.3 Hz, 1H), 3.86 (dd, *J* = 12.6, 3.0 Hz, 1H), 3.77 – 3.45 (m, 9H), 1.59 (d, *J* = 6.9 Hz, 3H). ^13^C NMR (101 MHz, Methanol-*d*_4_) δ 169.8, 164.3 (d, *J* = 247.9 Hz), 156.6, 141.8 (d, *J* = 11.0 Hz), 134.9 (d, *J* = 2.0 Hz), 114.5 (d, *J* = 3.0 Hz), 106.9 (d, *J* = 16.7 Hz), 106.3 (d, *J* = 27.4 Hz), 90.2 (d, *J* = 3.0 Hz), 77.0, 75.2, 65.1, 63.1, 47.9 (2C), 47.4, 42.7 (2C), 30.5, 14.3.

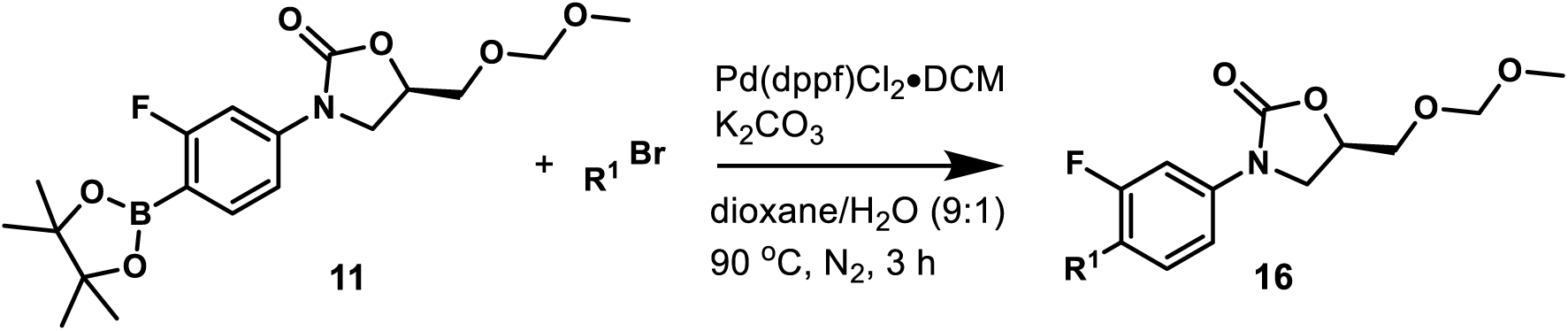

#### General Procedure 5: Synthesis of Heterocyclic MOM-protected Oxazolidinone Analogs 16a-16h

A mixture of compound **11** (76 mg, 0.2 mmol), heterocyclic bromide (0.26 mmol), potassium carbonate (110 mg, 0.8 mmol), and [1,1’-bis(diphenylphosphino)ferrocene]dichloropalladium(II), complex with dichloromethane (16 mg, 0.02 mmol) in dioxane/H_2_O (v/v= 9:1, 1 mL) was stirred at 90 °C under nitrogen atmosphere. After the reaction was judged to be completed by TLC (3 h), it was cooled to room temperature, diluted with EtOAc, washed with water, and concentrated under reduced pressure by rotary evaporation. The crude residue was purified by flash column chromatography (SiO_2_, eluent gradient 0-8% MeOH in DCM) to afford compound **16**.

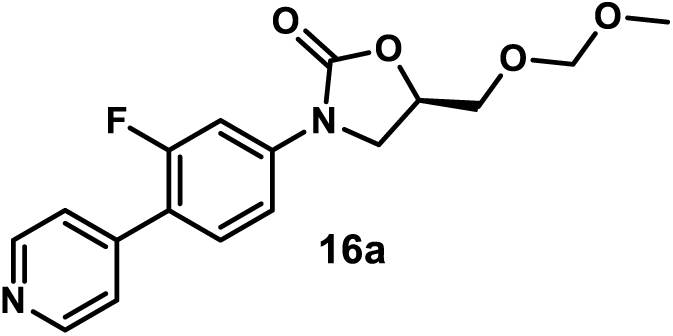

(*R)-3-(3-Fluoro-4-(pyridin-4-yl)phenyl)-5-((methoxymethoxy)methyl)oxazolidin-2-one (**16a**)*. A mixture of compound **11** (125 mg, 0.33 mmol), 4-bromopyridine hydrochloride (83 mg, 0.43 mmol), potassium carbonate (182 mg, 1.32 mmol), and [1,1’-bis(diphenylphosphino)ferrocene]dichloropalladium(II), complex with dichloromethane (27 mg, 0.033 mmol) in dioxane/H_2_O (v/v= 9:1, 2 mL) was stirred at 90 °C under nitrogen atmosphere. After the reaction was judged to be completed by TLC (3 h), it was cooled to room temperature, diluted with EtOAc, washed with water, and concentrated under reduced pressure by rotary evaporation. The crude residue was purified by flash column chromatography (SiO_2_, eluent gradient 0-10% MeOH in DCM) followed by preparative TLC (SiO_2_, eluent gradient 100% EtOAc) to afford compound **16a** as a brown amorphous solid (80 mg, 73%). ^1^H NMR (300 MHz, Acetone-*d*_6_) δ 8.65 (d, *J* = 6.2 Hz, 2H), 7.75 (dd, *J* = 13.9, 2.3 Hz, 1H), 7.67 (t, *J* = 8.7 Hz, 1H), 7.61 – 7.47 (m, 3H), 5.02 – 4.90 (m, 1H), 4.67 (s, 2H), 4.32 (t, *J* = 9.0 Hz, 1H), 4.06 (dd, *J* = 9.0, 6.1 Hz, 1H), 3.93 – 3.70 (m, 2H), 3.33 (s, 3H).

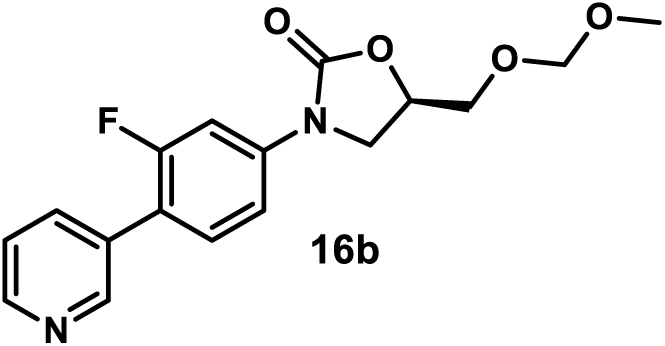

*(R)-3-(3-Fluoro-4-(pyridin-3-yl)phenyl)-5-((methoxymethoxy)methyl)oxazolidin-2-one (**16b**)*. A mixture of compound **11** (67.5 mg, 0.177 mmol), 3-bromopyridine (25 µL, 0.26 mmol), potassium carbonate (110 mg, 0.8 mmol), and [1,1’-bis(diphenylphosphino)ferrocene]dichloropalladium(II), complex with dichloromethane (16 mg, 0.02 mmol) in dioxane/H_2_O (v/v= 9:1, 1 mL) was stirred at 90 °C under nitrogen atmosphere. After the reaction was judged to be completed by TLC (2 h), it was cooled to room temperature, diluted with EtOAc, washed with water, and concentrated under reduced pressure by rotary evaporation. The crude residue was purified by flash column chromatography (SiO_2_, eluent gradient 0-100% EtOAc in hexanes) to afford compound **16b** as colorless oil (42 mg, 71%). ^1^H NMR (300 MHz, Acetone-*d*_6_) δ 8.78 (s, 1H), 8.58 (dd, *J* = 4.8, 1.7 Hz, 1H), 8.03 – 7.89 (m, 1H), 7.75 (dd, *J* = 13.6, 2.3 Hz, 1H), 7.61 (t, *J* = 8.6 Hz, 1H), 7.57 – 7.41 (m, 2H), 5.03 – 4.90 (m, 1H), 4.68 (s, 2H), 4.32 (t, *J* = 9.0 Hz, 1H), 4.06 (dd, *J* = 9.0, 6.1 Hz, 1H), – 3.70 (m, 2H), 3.34 (s, 3H).

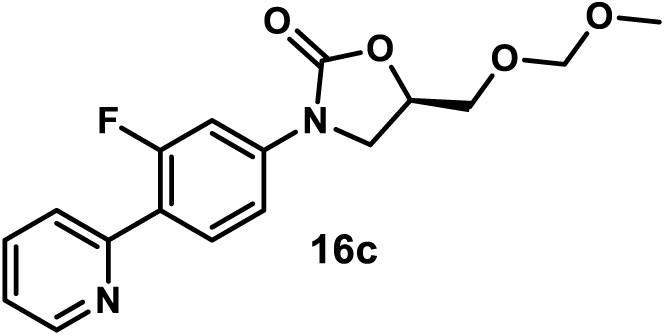

*(R)-3-(3-Fluoro-4-(pyridin-2-yl)phenyl)-5-((methoxymethoxy)methyl)oxazolidin-2-one (**16c**)*. A mixture of compound **11** (67.5 mg, 0.177 mmol), 2-bromopyridine (26.1 µL, 0.26 mmol), potassium carbonate (110 mg, 0.8 mmol), and [1,1’-bis(diphenylphosphino)ferrocene]dichloropalladium(II), complex with dichloromethane (16 mg, 0.02 mmol) in dioxane/H_2_O (v/v= 9:1, 1 mL) was stirred at 90 °C under nitrogen atmosphere After the reaction was judged to be completed by TLC (2 h), it was cooled to room temperature, diluted with EtOAc, washed with water, and concentrated under reduced pressure by rotary evaporation. The crude residue was purified by flash column chromatography (SiO_2_, eluent gradient 0-100% EtOAc in hexanes) to afford compound **16c** as a white amorphous solid (41 mg, 70%). ^1^H NMR (300 MHz, Acetone-*d*_6_) δ 8.70 (dd, *J* = 4.8, 1.5 Hz, 1H), 8.13 (t, *J* = 8.9 Hz, 1H), 7.92 – 7.80 (m, 2H), 7.74 (dd, *J* = 14.4, 2.3 Hz, 1H), 7.49 (dd, *J* = 8.8, 2.3 Hz, 1H), 7.38 – 7.29 (m, 1H), 5.07 – 4.93 (m, 1H), 4.68 (s, 2H), 4.32 (t, *J* = 9.0 Hz, 1H), 4.06 (dd, *J* = 9.0, 6.1 Hz, 1H), 3.96 – 3.74 (m, 2H), 3.33 (s, 3H).

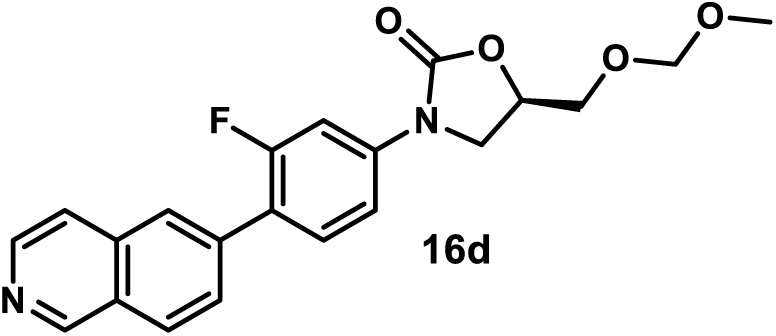

*(R)-3-(3-Fluoro-4-(isoquinolin-6-yl)phenyl)-5-((methoxymethoxy)methyl)oxazolidin-2-one (**16d**).* Using general procedure 5, employing 6-bromoisoquinoline (54 mg), compound **16d** was obtained as a yellow amorphous solid (69 mg, 90%). ^1^H NMR (400 MHz, Acetone-*d*_6_) δ 9.33 (s, 1H), 8.54 (d, *J* = 5.7 Hz, 1H), 8.19 (d, *J* = 8.5 Hz, 1H), 8.13 (s, 1H), 7.89 (dt, *J* = 8.5, 1.8 Hz, 1H), 7.84 (d, *J* = 5.7 Hz, 1H), 7.77 (dd, *J* = 13.8, 2.3 Hz, 1H), 7.71 (t, *J* = 8.6 Hz, 1H), 7.55 (dd, *J* = 8.6, 2.3 Hz, 1H), 5.04 – 4.94 (m, 1H), 4.68 (s, 2H), 4.33 (t, *J* = 8.9 Hz, 1H), 4.07 (dd, *J* = 8.9, 6.1 Hz, 1H), 3.89 (dd, *J* = 11.3, 3.4 Hz, 1H), 3.83 (dd, *J* = 11.3, 4.5 Hz, 1H), 3.34 (s, 3H). ^13^C NMR (101 MHz, Acetone-*d*_6_) δ 160.8 (d, *J* = 245.5 Hz), 155.1, 153.1, 144.5, 141.6 (d, *J* = 11.3 Hz), 138.1 (d, *J* = 1.6 Hz), 136.7, 132.1 (d, *J* = 4.6 Hz), 129.2 (d, *J* = 3.2 Hz), 128.6, 127.0 (d, *J* = 3.3 Hz), 123.3 (d, *J* = 13.5 Hz), 121.3, 114.6 (d, *J* = 3.1 Hz), 106.4 (d, *J* = 29.0 Hz), 97.3, 72.7, 68.7, 55.5, 47.4.

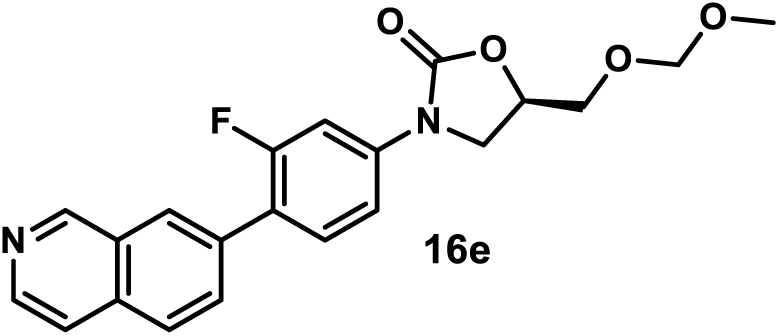

*(R)-3-(3-Fluoro-4-(isoquinolin-7-yl)phenyl)-5-((methoxymethoxy)methyl)oxazolidin-2-one (**16e**).* Using general procedure 5, employing 7-bromoisoquinoline (54 mg), compound **16e** was obtained as a yellow amorphous solid (68 mg, 89%). ^1^H NMR (400 MHz, Acetone-*d*_6_) δ 9.37 (s, 1H), 8.54 (d, *J* = 5.7 Hz, 1H), 8.29 (s, 1H), 8.05 (d, *J* = 8.6 Hz, 1H), 8.01 – 7.96 (m, 1H), 7.82 (d, *J* = 5.7 Hz, 1H), 7.77 (dd, *J* = 13.9, 2.2 Hz, 1H), 7.72 (t, *J* = 8.6 Hz, 1H), 7.55 (dd, *J* = 8.6, 2.2 Hz, 1H), 5.04 – 4.93 (m, 1H), 4.68 (s, 2H), 4.33 (t, *J* = 8.9 Hz, 1H), 4.07 (dd, *J* = 8.9, 6.2 Hz, 1H), 3.89 (dd, *J* = 11.3, 3.4 Hz, 1H), 3.83 (dd, *J* = 11.3, 4.4 Hz, 1H), 3.34 (s, 3H). ^13^C NMR (101 MHz, Acetone-*d*_6_) δ 160.7 (d, *J* = 245.0 Hz), 155.1, 153.6, 144.3, 141.3 (d, *J* = 11.4 Hz), 135.6, 135.2 (d, *J* = 1.7 Hz), 132.1 (d, *J* = 3.3 Hz), 132.0 (d, *J* = 4.8 Hz), 129.7, 128.1 (d, *J* = 3.3 Hz), 127.6, 123.3 (d, *J* = 13.4 Hz), 120.9, 114.6 (d, *J* = 3.1 Hz), 106.3 (d, *J* = 29.1 Hz), 97.3, 72.7, 68.7, 55.4, 47.4.

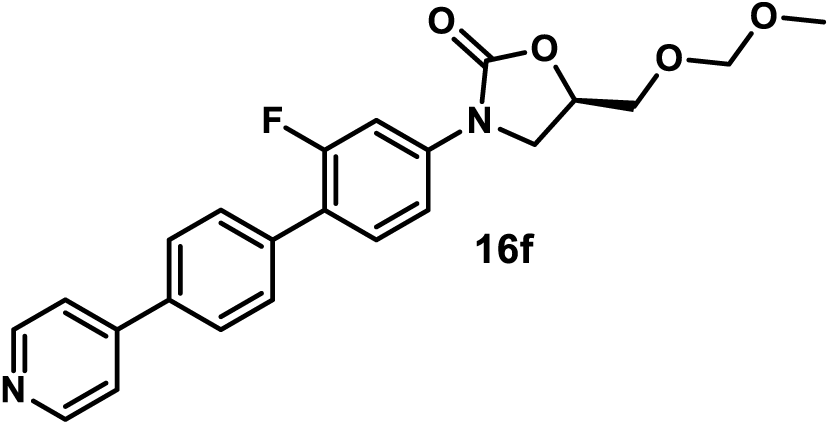

*(R)-3-(2-Fluoro-4’-(pyridin-4-yl)-[1,1’-biphenyl]-4-yl)-5-((methoxymethoxy)methyl)oxazolidin-2-one (**16f**).* Using general procedure 5, employing 4-(4-bromophenyl)pyridine (61 mg), compound **16f** was obtained as a yellow amorphous solid (75 mg, 90%). ^1^H NMR (400 MHz, Acetone-*d*_6_) δ 8.75 – 8.61 (m, 2H), 7.90 (d, *J* = 8.1 Hz, 2H), 7.81 – 7.65 (m, 5H), 7.63 (t, *J* = 8.7 Hz, 1H), 7.51 (dd, *J* = 8.7, 2.2 Hz, 1H), 5.04 – 4.93 (m, 1H), 4.68 (s, 2H), 4.32 (t, *J* = 8.9 Hz, 1H), 4.06 (dd, *J* = 8.9, 6.1 Hz, 1H), 3.88 (dd, *J* = 11.3, 3.4 Hz, 1H), 3.82 (dd, *J* = 11.3, 4.4 Hz, 1H), 3.34 (s, 3H). ^13^C NMR (101 MHz, Acetone-*d*_6_) δ 160.6 (d, *J* = 245.0 Hz), 155.1, 151.3 (2C), 148.0, 141.1 (d, *J* = 11.4 Hz), 137.9, 137.0, 131.6 (d, *J* = 4.8 Hz), 130.3 (d, *J* = 3.4 Hz, 2C), 127.9 (2C), 123.3 (d, *J* = 13.2 Hz), 122.1 (2C), 114.5 (d, *J* = 3.3 Hz), 106.3 (d, *J* = 29.1 Hz), 97.3, 72.6, 68.7, 55.4, 47.4.

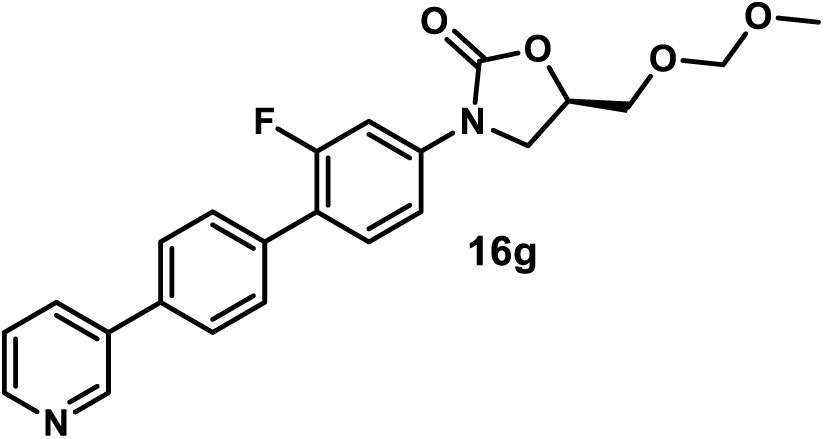

*(R)-3-(2-Fluoro-4’-(pyridin-3-yl)-[1,1’-biphenyl]-4-yl)-5-((methoxymethoxy)methyl)oxazolidin-2-one (**16g**).* Using general procedure 5, employing 3-(4-bromophenyl)pyridine (61 mg), compound **16g** was obtained as a yellow amorphous solid (68 mg, 83%).^1^H NMR (400 MHz, Acetone-*d*_6_) δ 8.94 (d, *J* = 2.4 Hz, 1H), 8.73 – 8.55 (m, 1H), 8.18 – 8.03 (m, 1H), 7.82 (d, *J* = 8.3 Hz, 2H), 7.78 – 7.68 (m, 3H), 7.63 (t, *J* = 8.8 Hz, 1H), 7.57 – 7.44 (m, 2H), 5.03 – 4.93 (m, 1H), 4.68 (s, 2H), 4.32 (t, *J* = 8.9 Hz, 1H), 4.06 (dd, *J* = 8.9, 6.2 Hz, 1H), 3.88 (dd, *J* = 11.3, 3.4 Hz, 1H), 3.83 (dd, *J* = 11.3, 4.4 Hz, 1H), 3.35 (s, 3H). ^13^C NMR (101 MHz, Acetone-*d*_6_) δ 160.6 (d, *J* = 245.2 Hz), 155.1, 149.6, 148.8, 141.0 (d, *J* = 11.3 Hz), 137.8, 136.6, 136.0, 134.8, 131.6 (d, *J* = 4.9 Hz), 130.3 (d, *J* = 3.3 Hz, 2C), 128.0 (2C), 124.6, 123.5 (d, *J* = 13.0 Hz), 114.5 (d, *J* = 3.2 Hz), 106.3 (d, *J* = 29.1 Hz), 97.3, 72.6, 68.7, 55.4, 47.4.

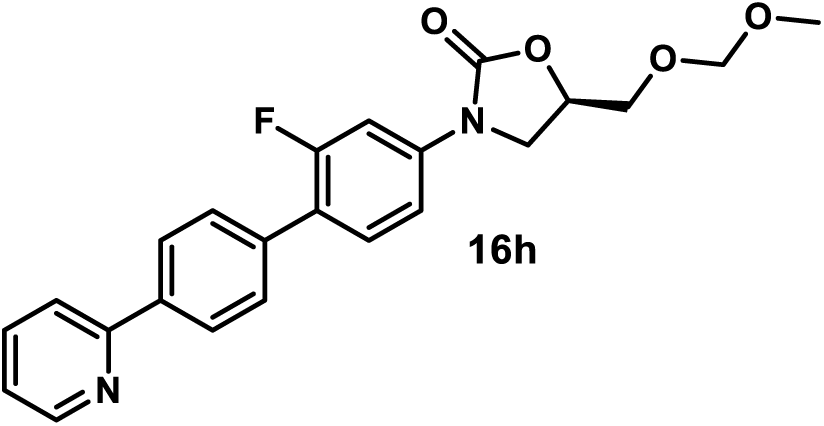

*(R)-3-(2-Fluoro-4’-(pyridin-2-yl)-[1,1’-biphenyl]-4-yl)-5-((methoxymethoxy)methyl)oxazolidin-2-one (**16h**)*. Using general procedure 5, employing 2-(4-bromophenyl)pyridine (61 mg), compound **16h** was obtained after flash column chromatography (SiO_2_, eluent gradient 0-100% EtOAc in hexanes) as a yellow amorphous solid (64 mg, 78%). ^1^H NMR (400 MHz, Acetone-*d*_6_) δ 8.69 (ddd, *J* = 4.7, 1.8, 0.9 Hz, 1H), 8.23 (d, *J* = 8.5 Hz, 2H), 8.03 – 7.97 (m, 1H), 7.89 (td, *J* = 7.7, 1.8 Hz, 1H), 7.77 – 7.67 (m, 3H), 7.63 (t, *J* = 8.8 Hz, 1H), 7.50 (dd, *J* = 8.8, 2.3 Hz, 1H), 7.34 (ddd, *J* = 7.5, 4.8, 1.1 Hz, 1H), 5.04 – 4.92 (m, 1H), 4.68 (s, 2H), 4.31 (t, *J* = 8.9 Hz, 1H), 4.05 (dd, *J* = 8.9, 6.2 Hz, 1H), 3.88 (dd, *J* = 11.3, 3.5 Hz, 1H), 3.82 (dd, *J* = 11.3, 4.5 Hz, 1H), 3.34 (s, 3H). ^13^C NMR (101 MHz, Acetone-*d*_6_) δ 159.7 (d, *J* = 245.2 Hz), 156.2, 154.21, 149.7, 140.1 (d, *J* = 11.2 Hz), 138.4, 136.9, 135.8 (d, *J* = 1.9 Hz), 130.7 (d, *J* = 4.8 Hz), 129.0 (d, *J* = 3.4 Hz, 2C), 126.8, 122.8 (d, *J* = 13.6 Hz), 122.4 (2C), 120.0, 113.6 (d, *J* = 3.2 Hz), 105.4 (d, *J* = 29.3 Hz), 96.4, 71.7, 67.8, 54.5, 46.5.

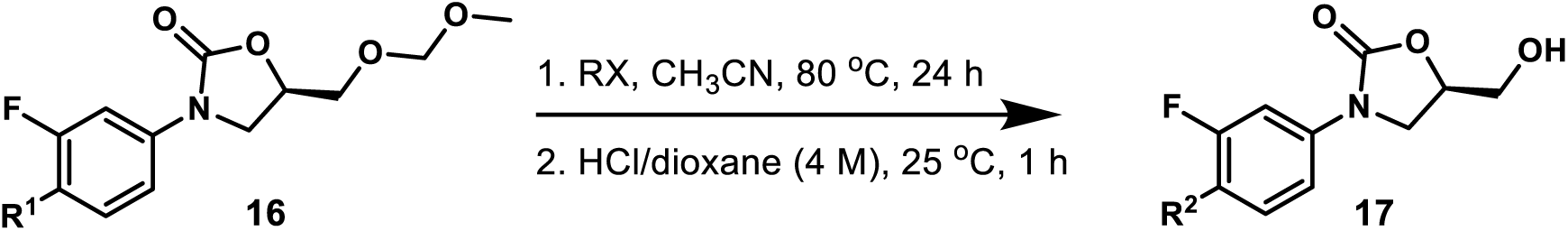

#### General Procedure 6: Synthesis of Positively Charged Oxazolidinone Analogs 17a-17l

A mixture of compound **16** (0.1 mmol) and haloalkane (0.5 mmol) in CH_3_CN (1 mL) was stirred in a sealed tube at 80 °C for 24 h. The reaction mixture was cooled to room temperature, purged with nitrogen flow for 5 min to remove most haloalkane, and concentrated under reduced pressure by rotary evaporation. The resulting residue was charged with the addition of HCl/dioxane (4 M, 1 mL) [if necessary, MeOH (1 mL) was employed as co-solvent]. The reaction mixture was stirred at 25 °C for 1 h, purged with nitrogen flow for 5 minutes to remove most hydrogen chloride, and concentrated under reduced pressure by rotary evaporation. The residue was further purified to give compounds **17** (their purification protocols are reported individually).

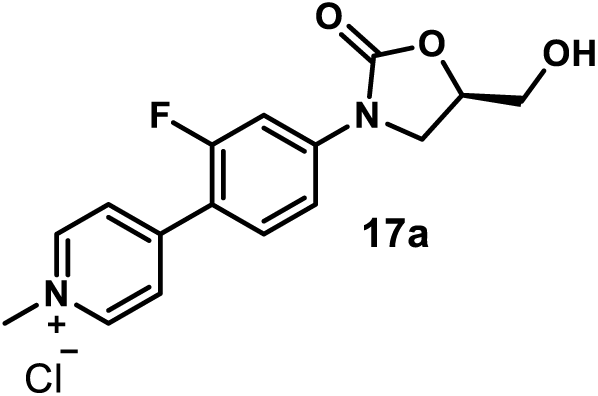

*(R)-4-(2-Fluoro-4-(5-(hydroxymethyl)-2-oxooxazolidin-3-yl)phenyl)-1-methylpyridin-1-ium Chloride (**17a**)*. A mixture of compound **16a** (72 mg, 0.22 mmol) and methyl iodide (30 µL, 0.54 mmol) in CH_3_CN (2 mL) was stirred in a sealed tube at 80 °C for 24 h. The reaction mixture was cooled to room temperature, purged with nitrogen flow for 5 min to remove most of methyl iodide, and concentrated under reduced pressure by rotary evaporation. The resulting residue was charged with the addition of HCl/dioxane (4 M, 2 mL). The reaction mixture was stirred at 25 °C for 1 h, purged with nitrogen flow for 5 minutes to remove most hydrogen chloride, and concentrated under reduced pressure by rotary evaporation. The crude residue was dissolved in methanol (2 mL). The solution was precipitated with the addition of acetone (4 mL) and kept still overnight. The supernatant was removed carefully, and the solid was re-suspended in acetone (4 mL). The acetone supernatant was removed carefully, and the solid was dried under high vacuum to afford **17a** as a yellow amorphous solid (30 mg, 40%). ^1^H NMR (500 MHz, DMSO-*d*_6_) δ 9.05 (d, *J* = 6.6 Hz, 2H), 8.35 (d, *J* = 6.6 Hz, 2H), 7.97 (t, *J* = 8.8 Hz, 1H), 7.77 (dd, *J* = 14.4, 2.2 Hz, 1H), 7.62 (dd, *J* = 8.8, 2.2 Hz, 1H), 5.44 – 5.32 (m, 1H), 4.88 – 4.74 (m, 1H), 4.36 (s, 3H), 4.17 (t, *J* = 9.0 Hz, 1H), 3.97 (dd, *J* = 9.0, 5.9 Hz, 1H), 3.76 – 3.66 (m, 1H), 3.66 – 3.55 (m, 1H). ^13^C NMR (151 MHz, DMSO-*d*_6_) δ 160.2 (d, *J* = 250.4 Hz), 154.3, 149.7, 145.5 (2C), 143.2 (d, *J* = 11.9 Hz), 131.7 (d, *J* = 3.5 Hz), 125.9 (d, *J* = 5.3 Hz, 2C), 116.3 (d, *J* = 11.5 Hz), 114.1 (d, *J* = 2.9 Hz), 105.4 (d, *J* = 28.4 Hz), 73.7, 61.4, 47.2, 46.0. MSESI *m/z*: 303.1145 (C_16_H_16_FN_2_O_3_^+^ requires 303.1139).

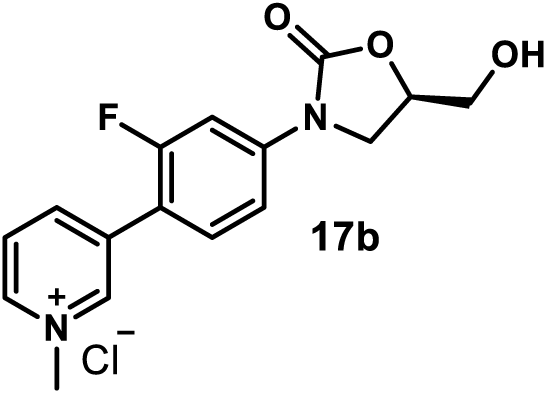

*(R)-3-(2-Fluoro-4-(5-(hydroxymethyl)-2-oxooxazolidin-3-yl)phenyl)-1-methylpyridin-1-ium Chloride (**17b**)*. Using general procedure 6, employing **16b** (33 mg, 0.1 mmol) and methyl iodide (31.1 µL, 0.5 mmol), compound **17b** was obtained as a yellow amorphous solid (10 mg, 30%). Pyridinium **17b** purification protocol: The crude residue was dissolved in methanol (1 mL). The solution was precipitated with the addition of acetone (5 mL) and kept still overnight. The supernatant was removed carefully and the solid was re-suspended in acetone (2 mL). The acetone supernatant was removed carefully and the solid was dried under high vacuum to afford **17b**.^1^H NMR (400 MHz, DMSO-*d*_6_) δ 9.34 (s, 1H), 9.02 (d, *J* = 6.0 Hz, 1H), 8.78 (d, *J* = 8.2 Hz, 1H), 8.22 (dd, *J* = 8.2, 6.0 Hz, 1H), 7.89 – 7.71 (m, 2H), 7.59 (dd, *J* = 8.7, 2.3 Hz, 1H), 5.36 (t, *J* = 5.5 Hz, 1H), 4.85 – 4.72 (m, 1H), 4.43 (s, 3H), 4.17 (t, *J* = 9.0 Hz, 1H), 3.95 (dd, *J* = 9.0, 6.0 Hz, 1H), 3.70 (ddd, *J* = 12.3, 5.5, 3.1 Hz, 1H), 3.58 (ddd, *J* = 12.3, 5.5, 3.7 Hz, 1H). ^13^C NMR (101 MHz, DMSO-*d*_6_) δ 159.3 (d, *J* = 247.2 Hz), 154.4, 144.8 (d, *J* = 3.3 Hz), 144.1, 144.0, 141.9 (d, *J* = 11.6 Hz), 134.1 (d, *J* = 1.3 Hz), 131.3 (d, *J* = 3.6 Hz), 127.5, 115.5 (d, *J* = 12.8 Hz), 114.1 (d, *J* = 2.9 Hz), 105.3 (d, *J* = 27.9 Hz), 73.6, 61.5, 48.2, 46.0. MSESI *m/z*: 303.1122 (C_16_H_16_FN_2_O_3_^+^ requires 303.1139).

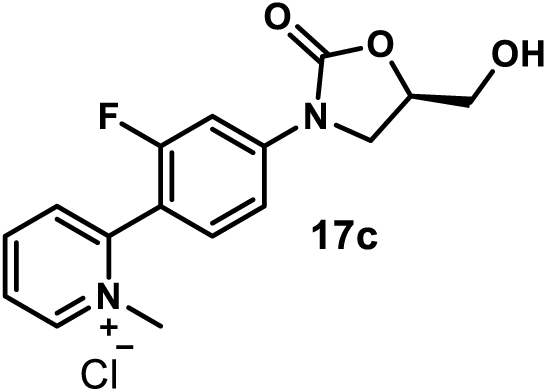

*(R)-2-(2-Fluoro-4-(5-(hydroxymethyl)-2-oxooxazolidin-3-yl)phenyl)-1-methylpyridin-1-ium Chloride (**17c**)*. Using general procedure 5, employing **16c** (33 mg, 0.1 mmol) and methyl iodide (31.1 µL, 0.5 mmol), compound **17c** was obtained as a yellow amorphous solid (5 mg, 15%). Pyridinium **17c** purification protocol: The crude residue was dissolved in methanol (1 mL). The solution was precipitated with the addition of acetone (5 mL) and kept still overnight. The supernatant was removed carefully and the solid was re-suspended in acetone (2 mL). The acetone supernatant was removed carefully and the solid was dried under high vacuum to afford **17c**. ^1^H NMR (400 MHz, Methanol-*d*_4_) δ 9.10 (d, *J* = 6.1 Hz, 1H), 8.70 – 8.59 (m, 1H), 8.20 – 8.05 (m, 2H), 7.90 (dd, *J* = 13.0, 2.1 Hz, 1H), 7.71 – 7.53 (m, 2H), 4.84 – 4.75 (m, 1H), 4.27 – 4.19 (m, 4H), 4.02 (dd, *J* = 9.0, 6.2 Hz, 1H), 3.88 (dd, *J* = 12.6, 3.0 Hz, 1H), 3.71 (dd, *J* = 12.6, 3.7 Hz, 1H). ^13^C NMR (101 MHz, DMSO-*d*_6_) δ 158.8 (d, *J* = 246.7 Hz), 154.3, 149.2, 147.5, 145.6, 143.2 (d, *J* = 11.4 Hz), 131.9 (d, *J* = 2.9 Hz), 130.8, 127.3, 113.8 (d, *J* = 3.0 Hz), 113.3 (d, *J* = 15.0 Hz), 104.9 (d, *J* = 26.8 Hz), 73.6, 61.5, 46.6 (d, *J* = 2.6 Hz), 46.0. MSESI *m/z*: 303.1156 (C_16_H_16_FN O ^+^ requires 303.1139).

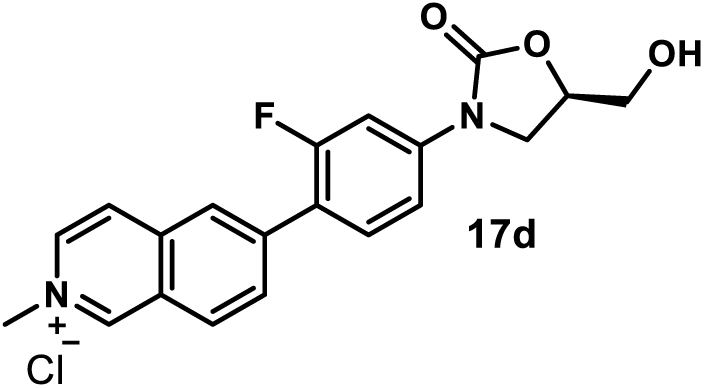

*(R)-6-(2-Fluoro-4-(5-(hydroxymethyl)-2-oxooxazolidin-3-yl)phenyl)-2-methylisoquinolin-2-ium Chloride (**17d**).* Using general procedure 5, employing **16d** (38 mg, 0.1 mmol) and methyl iodide (31.1 µL, 0.5 mmol), employing MeOH (1 mL) as the co-solvent in the deprotection step, compound **17d** was obtained as a yellow amorphous solid (25 mg, 64%). Pyridinium **17d** purification protocol: The crude residue was dissolved in methanol (2 mL), and the solution was added to EtOAc (20 mL). The suspension was kept still overnight. The supernatant was removed carefully. The solid was washed with EtOAc (5 mL) and dried under high vacuum to afford **17d**. ^1^H NMR (400 MHz, Methanol-*d*_4_) δ 9.82 (s, 1H), 8.58 (d, *J* = 6.7 Hz, 1H), 8.53 – 8.42 (m, 3H), 8.25 (dd, *J* = 8.7, 2.0 Hz, 1H), 7.85 – 7.70 (m, 2H), 7.51 (dd, *J* = 8.7, 2.2 Hz, 1H), 4.82 – 4.77 (m, 1H), 4.55 (s, 3H), 4.20 (t, *J* = 8.9 Hz, 1H), 4.01 (dd, *J* = 8.9, 6.3 Hz, 1H), 3.90 (dd, *J* = 12.6, 3.1 Hz, 1H), 3.73 (dd, *J* = 12.6, 3.8 Hz, 1H). ^13^C NMR (101 MHz, Methanol-*d*_4_) δ 161.5 (d, *J* = 248.2 Hz), 156.7, 151.3, 145.4 (d, *J* = 2.0 Hz), 142.9 (d, *J* = 11.5 Hz), 138.9, 137.0, 133.3 (d, *J* = 3.9 Hz), 132.6 (d, *J* = 4.0 Hz), 131.5, 128.0, 127.6 (d, *J* = 4.1 Hz), 127.1, 122.3 (d, *J* = 12.7 Hz), 115.4 (d, *J* = 3.1 Hz), 107.1 (d, *J* = 29.1 Hz), 75.3, 63.1, 48.7, 47.5. MSESI *m/z*: 353.1300 (C_20_H_18_FN O ^+^ requires 353.1296).

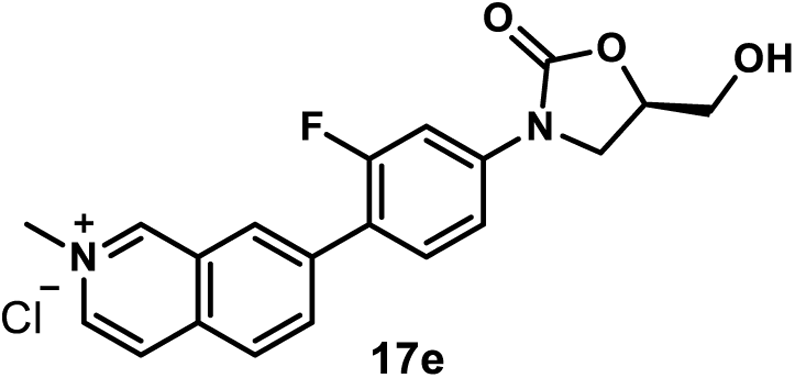

*(R)-7-(2-Fluoro-4-(5-(hydroxymethyl)-2-oxooxazolidin-3-yl)phenyl)-2-methylisoquinolin-2-ium Chloride (**17e**)*. Using general procedure 5, employing **16e** (38 mg, 0.1 mmol) and methyl iodide (31.1 µL, 0.5 mmol), employing MeOH (1 mL) as the co-solvent in the deprotection step, compound **17e** was obtained as a yellow amorphous solid (25 mg, 64%). Pyridinium **17e** purification protocol: The crude residue was dissolved in a mixture of methanol (2 mL) and acetone (4 mL). The solution was precipitated with the addition of EtOAc (10 mL) and kept still overnight. The suspension was centrifuged, and the supernatant was removed carefully. The solid was re-suspended in EtOAc (5 mL), and the suspension was centrifuged. The EtOAc supernatant was removed carefully and the solid was dried under high vacuum to afford **17e**. ^1^H NMR (400 MHz, DMSO-*d*_6_) δ 10.05 (s, 1H), 8.73 (d, *J* = 6.8 Hz, 1H), 8.65 (s, 1H), 8.60 (d, *J* = 6.8 Hz, 1H), 8.44 (s, 2H), 7.91 – 7.71 (m, 2H), 7.59 (d, *J* = 8.8 Hz, 1H), 5.32 (t, *J* = 5.6 Hz, 1H), 4.87 – 4.69 (m, 1H), 4.49 (s, 3H), 4.17 (t, *J* = 9.0 Hz, 1H), 3.98 – 3.88 (m, 1H), 3.78 – 3.67 (m, 1H), 3.65 – 3.55 (m, 1H). ^13^C NMR (101 MHz, DMSO-*d*_6_) δ 159.4 (d, *J* = 246.4 Hz), 154.4, 150.9, 140.9 (d, *J* = 11.6 Hz), 137.1 (d, *J* = 2.3 Hz), 137.0, 136.2, 135.7, 131.4 (d, *J* = 4.1 Hz), 129.1 (d, *J* = 4.2 Hz), 127.8, 127.4, 125.3, 120.2 (d, *J* = 12.8 Hz), 114.1 (d, *J* = 3.0 Hz), 105.4 (d, *J* = 28.4 Hz), 73.5, 61.6, 48.1, 46.0. MSESI *m/z*: 353.1306 (C_20_H_18_FN_2_O_3_^+^ requires 353.1296).

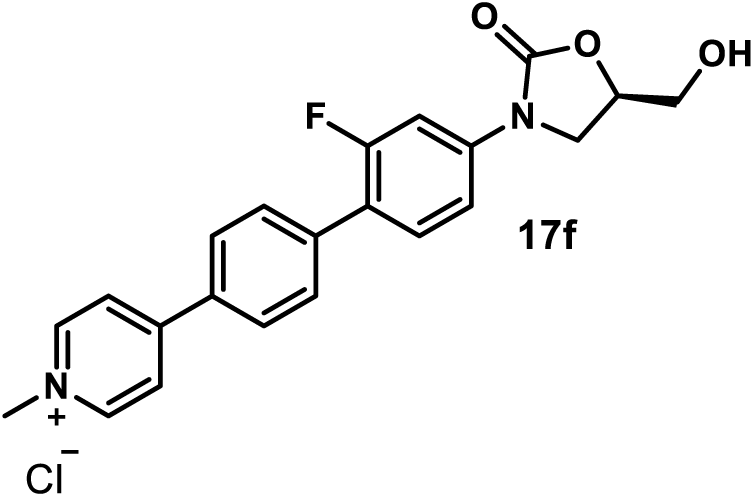

*(R)-4-(2’-Fluoro-4’-(5-(hydroxymethyl)-2-oxooxazolidin-3-yl)-[1,1’-biphenyl]-4-yl)-1-methylpyridin-1-ium Chloride (**17f**).* Using general procedure 5, employing **16f** (41 mg, 0.1 mmol) and methyl iodide (31.1 µL, 0.5 mmol), compound **17f** was obtained as a brown amorphous solid (20 mg, 48%). Pyridinium **17f** purification protocol: The crude residue was suspended in acetone (3 mL), and the suspension was kept still for 30 min. The supernatant was removed carefully and the solid was re-suspended in acetone (3 mL). The acetone supernatant was removed carefully and the solid was dried under high vacuum to afford **17f**. ^1^H NMR (400 MHz, DMSO-*d*_6_) δ 9.05 (d, *J* = 6.6 Hz, 2H), 8.57 (d, *J* = 6.6 Hz, 2H), 8.39 – 8.15 (m, 2H), 7.86 – 7.81 (m, 2H), 7.74 – 7.64 (m, 2H), 7.52 (dd, *J* = 8.6, 2.3 Hz, 1H), 5.31 (s, 1H), 4.86 – 4.69 (m, 1H), 4.34 (s, 3H), 4.15 (t, *J* = 9.0 Hz, 1H), 3.91 (dd, *J* = 9.0, 6.1 Hz, 1H), 3.76 – 3.66 (m, 1H), 3.64 – 3.55 (m, 1H). ^13^C NMR (101 MHz, DMSO-*d*_6_) δ 159.3 (d, *J* = 245.4 Hz), 154.4, 153.6, 145.7 (2C), 140.3 (d, *J* = 11.1 Hz), 138.3 (d, *J* = 1.9 Hz), 132.5, 131.0 (d, *J* = 4.4 Hz), 129.7 (d, *J* = 3.5 Hz, 2C), 128.4 (2C), 124.0 (2C), 121.2 (d, *J* = 13.0 Hz), 114.0 (d, *J* = 3.0 Hz), 105.4 (d, *J* = 28.6 Hz), 73.5, 61.6, 47.1, 46.0. MSESI *m/z*: 379.1418 (C_22_H_20_FN_2_O_3_^+^ requires 379.1452).

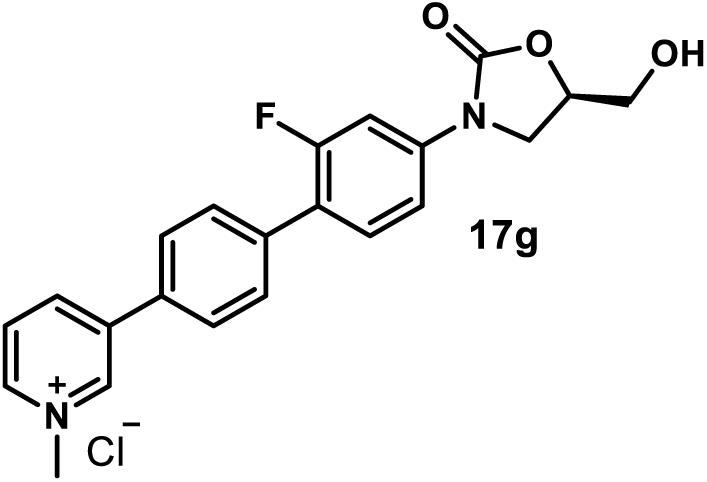

*(R)-3-(2’-Fluoro-4’-(5-(hydroxymethyl)-2-oxooxazolidin-3-yl)-[1,1’-biphenyl]-4-yl)-1-methylpyridin-1-ium Chloride (**17g**).* Using general procedure 5, employing **16g** (41 mg, 0.1 mmol) and methyl iodide (31.1 µL, 0.5 mmol), compound **17g** was obtained as a gray amorphous solid (20 mg, 48%). Pyridinium **17g** purification protocol: The crude residue was suspended in acetone (3 mL), and the suspension was kept still for 30 min. The supernatant was removed carefully and the solid was re-suspended in acetone (3 mL). The acetone supernatant was removed carefully. The solid was suspended in MeOH (2 mL), and the suspension was added to EtOAc (10 mL). The resulting suspension was kept still overnight. The supernatant was removed carefully, and the solid was further washed with EtOAc (5 mL) and then dried under high vacuum to afford **17g**. ^1^H NMR (400 MHz, DMSO-*d*_6_) δ 9.53 (s, 1H), 9.00 (d, *J* = 6.0 Hz, 1H), 8.96 (d, *J* = 8.1 Hz, 1H), 8.23 (dd, *J* = 8.1, 6.0 Hz, 1H), 8.03 (d, *J* = 8.0 Hz, 2H), 7.81 (d, *J* = 8.0 Hz, 2H), 7.73 – 7.60 (m, 2H), 7.50 (dd, *J* = 8.6, 2.2 Hz, 1H), 5.33 (t, *J* = 5.6 Hz, 1H), 4.85 – 4.71 (m, 1H), 4.44 (s, 3H), 4.15 (t, *J* = 9.0 Hz, 1H), 3.92 (dd, *J* = 9.0, 6.1 Hz, 1H), 3.78 – 3.66 (m, 1H), 3.63 – 3.52 (m, 1H). ^13^C NMR (101 MHz, DMSO-*d*_6_) δ 159.2 (d, *J* = 245.1 Hz), 154.4, 143.9, 143.8, 142.0, 140.0 (d, *J* = 11.4 Hz), 138.6, 136.4, 132.3, 131.0 (d, *J* = 4.8 Hz), 129.6 (d, *J* = 3.4 Hz, 2C), 127.73, 127.71 (2C), 121.5 (d, *J* = 13.0 Hz), 113.9 (d, *J* = 3.0 Hz), 105.4 (d, *J* = 28.6 Hz), 73.5, 61.6, 48.1, 46.0. MSESI *m/z*: 379.1468 (C_22_H_20_FN_2_O_3_^+^ requires 379.1452).

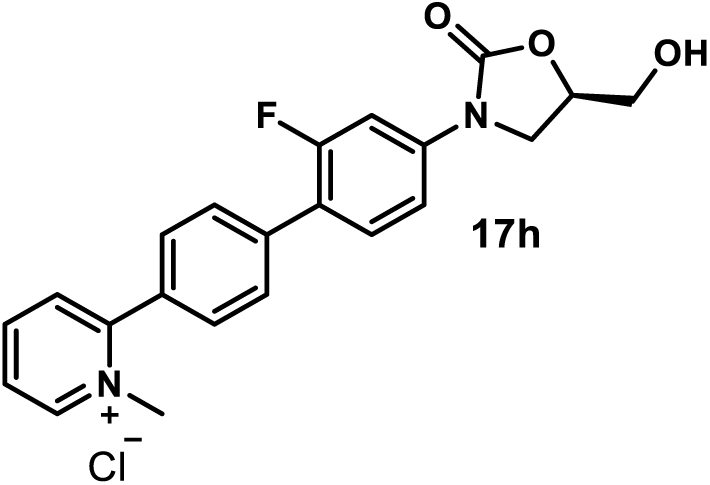

*(R)-2-(2’-Fluoro-4’-(5-(hydroxymethyl)-2-oxooxazolidin-3-yl)-[1,1’-biphenyl]-4-yl)-1-methylpyridin-1-ium Chloride (**17h**).* Using general procedure 5, employing **16h** (41 mg, 0.1 mmol) and methyl iodide (31.1 µL, 0.5 mmol), compound **17h** was obtained as a gray amorphous solid (10 mg, 24%). Pyridinium **17h** purification protocol: The crude residue was suspended in MeOH (2 mL), and the suspension was added to acetone (10 mL). The resulting suspension was kept still overnight. The supernatant was removed carefully, and the solid was further washed with acetone (2 × 10 mL) and then dried under high vacuum to afford **17h**. ^1^H NMR (400 MHz, DMSO-*d*_6_) δ 9.19 (d, *J* = 6.1 Hz, 1H), 8.66 (dd, *J* = 8.6, 7.2 Hz, 1H), 8.33 – 8.10 (m, 2H), 7.93 – 7.76 (m, 4H), 7.74 – 7.63 (m, 2H), 7.53 (dd, *J* = 8.7, 2.3 Hz, 1H), 5.30 (s, 1H), 4.82 – 4.70 (m, 1H), 4.39 – 4.08 (m, 4H), 3.91 (dd, *J* = 8.9, 6.0 Hz, 1H), 3.71 (dd, *J* = 12.4, 3.3 Hz, 1H), 3.58 (dd, *J* = 12.4, 3.8 Hz, 1H). ^13^C NMR (101 MHz, DMSO-*d*_6_) δ 159.2 (d, *J* = 245.5 Hz), 154.7, 154.4, 146.8, 145.5, 140.2 (d, *J* = 11.5 Hz), 137.3, 131.1 (d, *J* = 4.5 Hz), 130.9, 129.9, 129.7 (2C), 129.1 (d, *J* = 3.3 Hz, 2C), 126.8, 121.3 (d, *J* = 13.0 Hz), 114.0 (d, *J* = 3.1 Hz), 105.4 (d, *J* = 28.6 Hz), 73.5, 61.6, 47.2, 46.0. MSESI *m/z*: 379.1451 (C_22_H_20_FN_2_O_3_^+^ requires 379.1452).

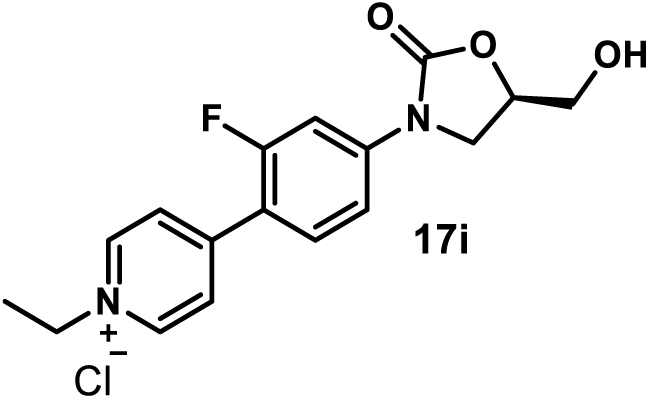

*(R)-1-Ethyl-4-(2-fluoro-4-(5-(hydroxymethyl)-2-oxooxazolidin-3-yl)phenyl)pyridin-1-ium Chloride (**17i**).* Using general procedure 6, employing **16a** (33 mg, 0.1 mmol) and ethyl iodide (40 µL, 0.5 mmol), compound **17i** was obtained as a yellow amorphous solid (15 mg, 42%). Pyridinium **17i** purification protocol: The crude residue was dissolved in MeOH (2 mL), and the solution was added to EtOAc (80 mL). The resulting suspension was kept still overnight. The supernatant was removed carefully, and the solid was further washed with EtOAc (10 mL) and then dried under high vacuum to afford **17i**. ^1^H NMR (500 MHz, DMSO-*d*_6_) δ 9.14 (d, *J* = 6.3 Hz, 2H), 8.37 (d, *J* = 6.3 Hz, 2H), 7.97 (t, *J* = 8.8 Hz, 1H), 7.85 – 7.73 (m, 1H), 7.66 – 7.55 (m, 1H), 5.31 (t, *J* = 5.6 Hz, 1H), 4.88 – 4.74 (m, 1H), 4.64 (q, *J* = 7.2 Hz, 2H), 4.18 (t, *J* = 9.1 Hz, 1H), 3.94 (dd, *J* = 9.1, 5.8 Hz, 1H), 3.78 – 3.66 (m, 1H), 3.62 – 3.54 (m, 1H), 1.57 (t, *J* = 7.2 Hz, 3H). ^13^C NMR (101 MHz, DMSO-*d*_6_) δ 160.2 (d, *J* = 250.6 Hz), 154.3, 150.1 (d, *J* = 1.9 Hz), 144.4 (2C), 143.3 (d, *J* = 11.9 Hz), 131.7 (d, *J* = 3.3 Hz), 126.3 (d, *J* = 5.3 Hz, 2C), 116.3 (d, *J* = 11.3 Hz), 114.1 (d, *J* = 2.8 Hz), 105.4 (d, *J* = 28.5 Hz), 73.7, 61.5, 55.7, 46.0, 16.3. MSESI *m/z*: 317.1291 (C_17_H_18_FN_2_O_3_^+^ requires 317.1296).

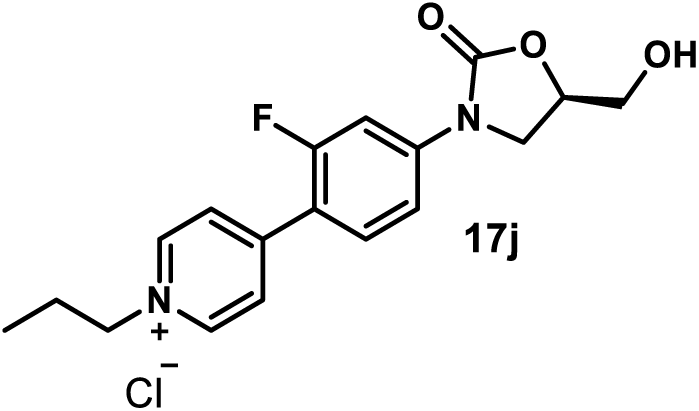

*(R)-4-(2-Fluoro-4-(5-(hydroxymethyl)-2-oxooxazolidin-3-yl)phenyl)-1-propylpyridin-1-ium Chloride (**17j**).* Using general procedure 6, employing **16a** (33 mg, 0.1 mmol) and propyl iodide (48.7 µL, 0.5 mmol), compound **17j** was obtained as a gray amorphous solid (15 mg, 41%). Pyridinium **17j** purification protocol: The crude residue was dissolved in MeOH (2 mL), and the solution was added to EtOAc (60 mL). The resulting suspension was kept still overnight. The supernatant was removed carefully, and the solid was further washed with EtOAc (2 × 20 mL) and then dried under high vacuum to afford **17j**. ^1^H NMR (500 MHz, DMSO-*d*_6_) δ 9.13 (d, *J* = 6.4 Hz, 2H), 8.38 (d, *J* = 6.4 Hz, 2H), 7.99 (t, *J* = 8.8 Hz, 1H), 7.78 (dd, *J* = 14.4, 2.2 Hz, 1H), 7.62 (dd, *J* = 8.8, 2.2 Hz, 1H), 5.31 (s, 1H), 4.89 – 4.75 (m, 1H), 4.58 (t, *J* = 7.3 Hz, 2H), 4.18 (t, *J* = 9.0 Hz, 1H), 3.95 (dd, *J* = 9.0, 5.9 Hz, 1H), 3.71 (dd, *J* = 12.5, 3.2 Hz, 1H), 3.59 (dd, *J* = 12.5, 3.8 Hz, 1H), 2.10 – 1.89 (m, 2H), 0.92 (t, *J* = 7.3 Hz, 3H). ^13^C NMR (101 MHz, DMSO-*d*_6_) δ 160.2 (d, *J* = 250.6 Hz), 154.3, 150.1 (d, *J* = 2.0 Hz), 144.6 (2C), 143.3 (d, *J* = 11.9 Hz), 131.7 (d, *J* = 3.4 Hz), 126.2 (d, *J* = 5.3 Hz, 2C), 116.2 (d, *J* = 11.4 Hz), 114.1 (d, *J* = 2.9 Hz), 105.4 (d, *J* = 28.4 Hz), 73.7, 61.5, 61.4, 46.0, 24.1, 10.3. MSESI *m/z*: 331.1463 (C_18_H_20_FN_2_O_3_^+^ requires 331.1452).

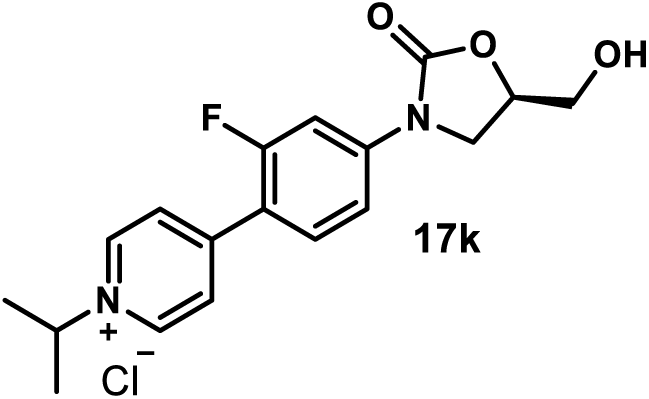

*(R)-4-(2-Fluoro-4-(5-(hydroxymethyl)-2-oxooxazolidin-3-yl)phenyl)-1-isopropylpyridin-1-ium Chloride (**17k**).* Using general procedure 6, employing **16a** (33 mg, 0.1 mmol) and isopropyl iodide (50 µL, 0.5 mmol), compound **17k** was obtained as a yellow amorphous solid (5 mg, 14%). Pyridinium **26k** purification protocol: The crude residue was dissolved in MeOH (1 mL), and the solution was added to EtOAc (80 mL). The resulting suspension was kept still overnight. The supernatant was removed carefully, and the solid was further washed with EtOAc (2 × 20 mL) and then dried under high vacuum to afford **17k**. ^1^H NMR (400 MHz, Methanol-*d*_4_) δ 9.05 (d, *J* = 6.4 Hz, 2H), 8.31 (d, *J* = 6.4 Hz, 2H), 7.99 – 7.86 (m, 1H), 7.83 – 7.75 (m, 1H), 7.55 (dd, *J* = 9.5, 2.1 Hz, 1H), 5.02 (p, *J* = 6.7 Hz, 1H), 4.84 – 4.72 (m, 1H), 4.30 – 4.06 (m, 1H), 4.04 – 3.92 (m, 1H), 3.92 – 3.83 (m, 1H), 3.74 – 3.66 (m, 1H), 1.72 (d, *J* = 6.7 Hz, 6H). ^13^C NMR (101 MHz, Methanol-*d*_4_) δ 162.3 (d, *J* = 251.4 Hz), 156.4, 153.1 (d, *J* = 2.1 Hz), 145.1 (d, *J* = 12.1 Hz), 143.8 (2C), 132.6 (d, *J* = 3.3 Hz), 127.9 (d, *J* = 5.9 Hz, 2C), 117.9 (d, *J* = 11.4 Hz), 115.5 (d, *J* = 3.0 Hz), 107.1 (d, *J* = 29.0 Hz), 75.4, 65.7, 63.1, 47.4, 23.1. MSESI *m/z*: 331.1446 (C_18_H_20_FN_2_O_3_^+^ requires 331.1452).

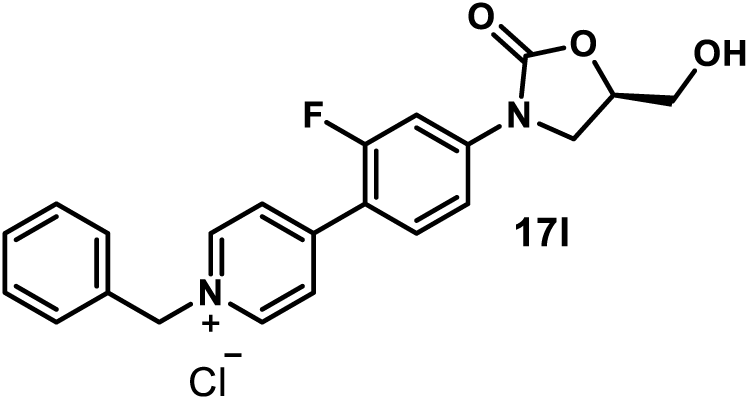

*(R)-1-Benzyl-4-(2-fluoro-4-(5-(hydroxymethyl)-2-oxooxazolidin-3-yl)phenyl)pyridin-1-ium Chloride (**17l**).* Using general procedure 6, employing **16a** (33 mg, 0.1 mmol) and benzyl bromide (59.4 µL, 0.5 mmol), compound **17l** was obtained as a gray amorphous solid (27 mg, 65%). Pyridinium **17l** purification protocol: The crude residue was dissolved in MeOH (2 mL), and the solution was added to EtOAc (15 mL). The resulting suspension was kept still overnight. The supernatant was removed carefully, and the solid was further washed with EtOAc (5 mL) and then dried under high vacuum to afford **17l**. ^1^H NMR (500 MHz, DMSO-*d*_6_) δ 9.23 (d, *J* = 6.5 Hz, 2H), 8.40 (d, *J* = 6.3 Hz, 2H), 7.97 (t, *J* = 8.8 Hz, 1H), 7.77 (dd, *J* = 14.4, 2.2 Hz, 1H), 7.62 (dd, *J* = 8.8, 2.2 Hz, 1H), 7.60 – 7.56 (m, 2H), 7.50 – 7.42 (m, 3H), 5.87 (s, 2H), 5.27 (t, *J* = 5.6 Hz, 1H), 4.84 – 4.69 (m, 1H), 4.17 (t, *J* = 9.1 Hz, 1H), 3.93 (dd, *J* = 9.1, 6.0 Hz, 1H), 3.71 (ddd, *J* = 12.4, 5.6, 3.2 Hz, 1H), 3.58 (ddd, *J* = 12.4, 5.6, 3.8 Hz, 1H). ^13^C NMR (101 MHz, DMSO-*d*_6_) δ 160.3 (d, *J* = 250.8 Hz), 154.2, 150.6 (d, *J* = 2.1 Hz), 144.6 (2C), 143.4 (d, *J* = 12.3 Hz), 134.4, 131.8 (d, *J* = 3.3 Hz), 129.4, 129.3 (2C), 128.8 (2C), 126.6 (d, *J* = 5.5 Hz, 2C), 116.2 (d, *J* = 11.1 Hz), 114.1 (d, *J* = 2.7 Hz), 105.4 (d, *J* = 28.3 Hz), 73.7, 62.6, 61.5, 46.0. MSESI *m/z*: 379.1469 (C_22_H_20_FN O ^+^ requires 379.1452).

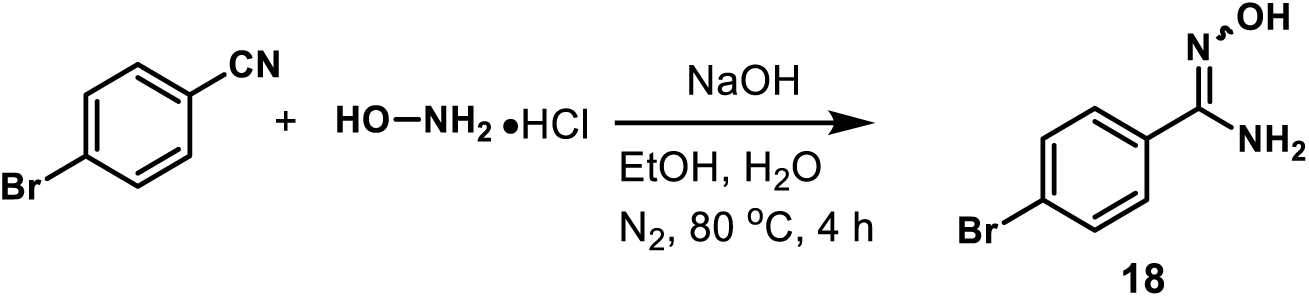

*4- Bromo-N’-hydroxybenzimidamide (**18**).* A mixture of 4-bromobenzonitrile (182 mg, 1 mmol), hydroxylammonium chloride (70 mg, 1 mmol), and sodium hydroxide (40 mg, 1 mmol) in ethanol (2 mL) and water (70 µL) was stirred under nitrogen atmosphere at 80 °C. After the reaction was judged to be completed by TLC (4 h), it was concentrated under reduced pressure by rotary evaporation. The residue was purified by flash column chromatography (SiO_2_, eluent gradient 0-10% MeOH in DCM) to afford compound **18** as a yellow amorphous solid (181 mg, 84%). ^1^H NMR (400 MHz, DMSO) δ 9.73 (s, 1H), 8.20 – 7.05 (m, 4H), 5.86 (s, 2H).

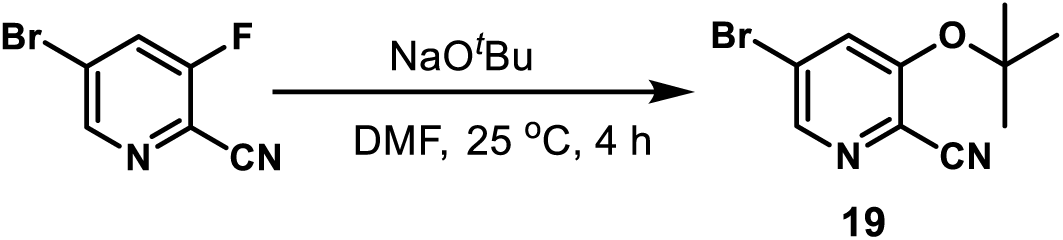

*5- Bromo-3-(tert-butoxy)picolinonitrile (**19**)*. To a solution of 5-bromo-3-fluoropicolinonitrile (1 g, 5 mmol) in anhydrous DMF (25 mL), sodium *tert*-butoxide (625 mg, 6.5 mmol) was added. The reaction mixture was stirred at 25 °C for 4 h, quenched with the addition of water, and extracted with a mixture of hexanes/EtOAc (v/v= 1:2). The organic layer was washed with water three times and evaporated under reduced pressure by rotary evaporation. The crude reside was purified by flash column chromatography (SiO_2_, eluent gradient 0-20% EtOAc in hexanes) to afford compound **19** as yellow oil (419 mg, 33%). ^1^H NMR (400 MHz, Chloroform-*d*) δ 8.41 (d, *J* = 1.9 Hz, 1H), 7.67 (d, *J* = 1.9 Hz, 1H), 1.52 (s, 9H).

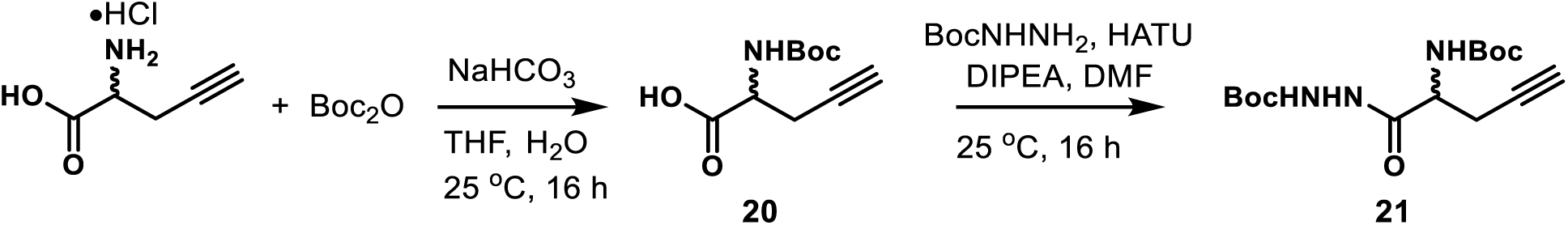

*tert-Butyl 2-(2-((tert-Butoxycarbonyl)amino)pent-4-ynoyl)hydrazine-1-carboxylate (**21**)*.^58^ Sodium bicarbonate (336 mg, 4 mmol) and di-*tert*-butyl dicarbonate (327 mg, 1.5 mmol) were added to a solution of 2-amino-4-pentynoic acid hydrochloride (150 mg, 1 mmol) in THF/H_2_O (v/v = 1:1, 5 mL) at 0 °C. The reaction mixture was stirred at 25 °C for 16 h. The mixture was diluted with Et_2_O and extracted with water three times. The combined aqueous layers were acidified to pH = 4-5 by carefully adding saturated aqueous citric acid in an ice bath and extracted with dichloromethane three times. The combined organic layers were dried over magnesium sulfate and concentrated under reduced pressure by rotary evaporation. The residue **20** was used in the next step without further purification (94 mg, 44%). To a solution of compound **20** (94 mg, 0.44 mmol) in DMF (1.0 mL) were added *tert*-butyl carbazate (87 mg, 0.66 mmol), HATU (335 mg, 0.88 mmol), and *N*,*N*-diisopropylethylamine (0.3 mL). The reaction mixture was stirred at 25 °C for 16 h, quenched with the addition of water, and extracted with EtOAc. The organic layer was washed with water two times as well as brine and evaporated under reduced pressure by rotary evaporation. The crude reside was purified by flash column chromatography (SiO_2_, eluent gradient 0-50% EtOAc in hexanes) to afford compound **21** as a yellow amorphous solid (119 mg, 83%). ^1^H NMR (400 MHz, CDCl_3_) δ 8.48 (s, 1H), 6.70 (s, 1H), 5.44 (d, *J* = 8.5 Hz, 1H), 4.42 (s, 1H), 2.83 – 2.55 (m, 2H), 2.08 (d, *J* = 2.2 Hz, 1H), 1.58 – 1.35 (m, 18H).

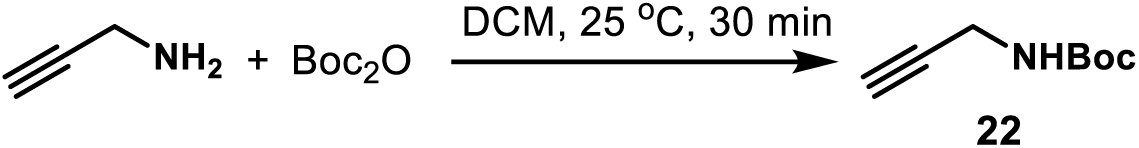

*tert-Butyl Prop-2-yn-1-ylcarbamate (**22**)*. To a solution of propargylamine (110 mg, 2 mmol) in DCM (2 mL) was added a solution of di-*tert*-butyl dicarbonate (458 mg, 2.1 mmol) in DCM (3 mL) dropwise with a syringe at 0 °C. The reaction mixture was stirred at 25 °C for 30 min, concentrated under reduced pressure by rotary evaporation. The crude reside was purified by flash column chromatography (SiO_2_, eluent gradient 0-10% MeOH in DCM) to afford compound **22** as a yellow amorphous solid (260 mg, 84%). ^1^H NMR (400 MHz, CDCl_3_) δ 4.69 (s, 1H), 3.92 (s, 2H), 2.22 (s, 1H), 1.45 (s, 9H).

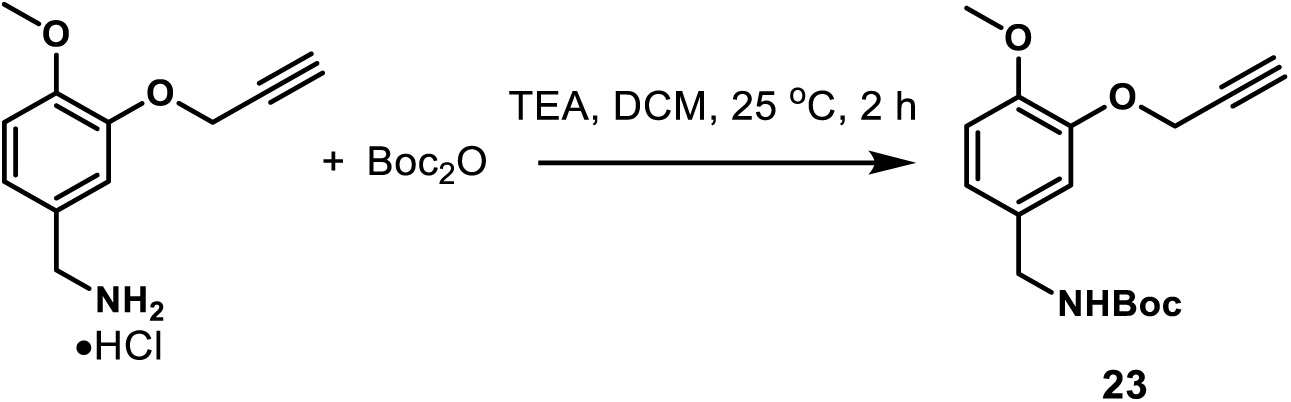

*tert-Butyl (4-Methoxy-3-(prop-2-yn-1-yloxy)benzyl)carbamate (**23**)*. Di-*tert*-butyl dicarbonate (98 mg, 0.45 mmol) was added to a solution of [4-methoxy-3-(2-propynyloxy)phenyl]methanamine hydrochloride (67 mg, 0.3 mmol) and triethylamine (126 µL, 0.9 mmol) in DCM (2 mL). The reaction mixture was stirred at 25 °C for 2 h, quenched with water, and extracted with EtOAc three times. The combined organic layers were evaporated under reduced pressure by rotary evaporation. The crude reside was purified by flash column chromatography (SiO_2_, eluent gradient 0-50% EtOAc in hexanes) to afford compound **23** as a white amorphous solid (64 mg, 73%). ^1^H NMR (400 MHz, CDCl_3_) δ 6.96 (s, 1H), 6.90 – 6.74 (m, 2H), 4.94 – 4.78 (m, 1H), 4.74 (d, *J* = 2.5 Hz, 2H), 4.24 (d, *J* = 5.8 Hz, 2H), 3.84 (s, 3H), 2.50 (t, *J* = 2.5 Hz, 1H), 1.45 (s, 9H).

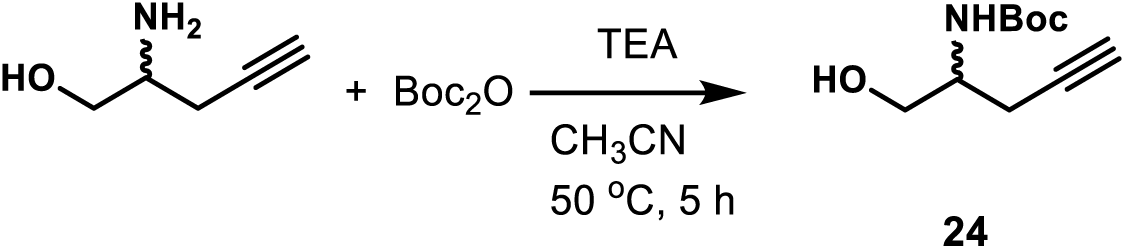

*tert-Butyl (1-Hydroxypent-4-yn-2-yl)carbamate (**24**).* A mixture of 2-aminopent-4-yn-1-ol (20 mg, 0.2 mmol), di-*tert*-butyl dicarbonate (87 mg, 0.4 mmol), and triethylamine (56 µL, 0.4 mmol) in CH_3_CN (1 mL) was stirred at 50 °C for 5 h. The mixture was evaporated under reduced pressure by rotary evaporation and the resulting reside was purified by flash column chromatography (SiO_2_, eluent gradient 0-3% MeOH in DCM) to afford compound **24** as a yellow amorphous solid (39 mg, 99%). ^1^H NMR (400 MHz, acetone) δ 3.95 (s, 1H), 3.77 – 3.50 (m, 2H), 2.60 – 2.30 (m, 3H), 1.77 (s, 1H), 1.40 (s, 9H).

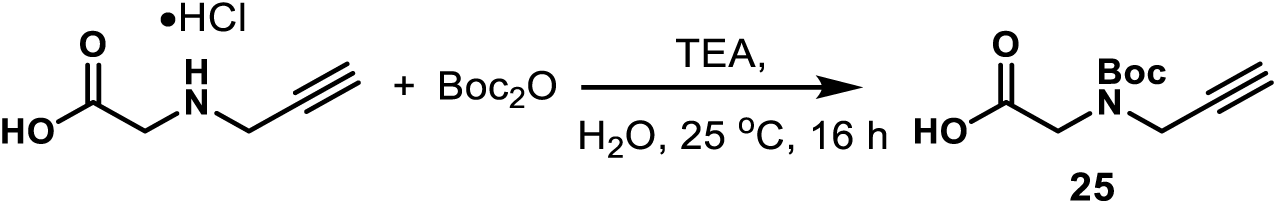

*N-(tert-Butoxycarbonyl)-N-(prop-2-yn-1-yl)glycine (**25**)*.^59^ To a solution of 2-[(prop-2-yn-1-yl) amino]acetic acid hydrochloride (45 mg, 0.3 mmol) in H_2_O (1 mL) were added di-*tert*-butyl dicarbonate (262 mg, 1.2 mmol), and triethylamine (293 µL, 2.1 mmol). The mixture was stirred at 25 °C for 16 h, diluted with water, and washed with hexanes (20 mL) to remove di-*tert*-butyl dicarbonate. The aqueous layer was acidified with the addition of aqueous HCl (1 M) to pH = 3 and extracted with EtOAc three times. The combined organic layers were dried over magnesium sulfate and concentrated under reduced pressure by rotary evaporation to give compound **25** (63 mg, 99%), which was used in the next step without further purification. ^1^H NMR (400 MHz, CDCl_3_) δ 4.43 – 3.89 (m, 4H), 2.27 (s, 1H), 1.66 – 0.87 (m, 9H).

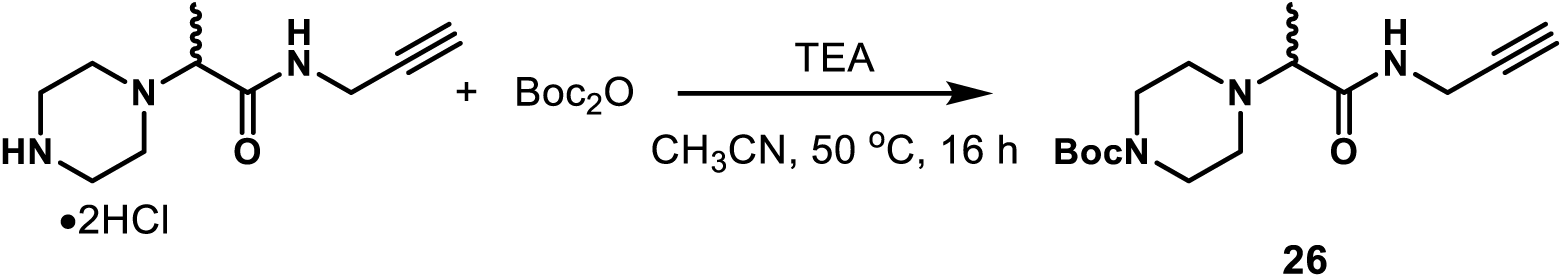

*tert-Butyl 4-(1-Oxo-1-(prop-2-yn-1-ylamino)propan-2-yl)piperazine-1-carboxylate (**26**).* A mixture of 2-(1-piperazinyl)-*N*-(2-propynyl)propanamide dihydrochloride (53 mg, 0.2 mmol), di-*tert*-butyl dicarbonate (174 mg, 0.8 mmol), and triethylamine (111 µL, 0.8 mmol) in CH_3_CN (2 mL) was stirred at 50 °C for 16 h. The mixture was evaporated under reduced pressure by rotary evaporation and the resulting residue was purified by flash column chromatography (SiO_2_, eluent gradient 0-15% MeOH in DCM) to afford compound **26** as a yellow amorphous solid (30 mg, 51%). Compound **26** was not pure but used in the next step without further purification.

## ASSOCIATED CONTENT

The supporting information is available free of charge on the ACS Publications website at DOI: Library design approach, representative HPLC traces, H^1^ and C^13^ NMR spectra for final compounds, table with PC decomposition, MIC values, ratio tables for each species, and comparative activity tables

## Author Contributions

All authors contributed to the writing of the manuscript and have approved its content. Z.H. and Q.P.A. designed, synthesized, purified, and characterized all analogs. I.V.L and B.C. measured antibacterial activities. V.V.R. carried out the PCA analysis. H.I.Z. designed and supervised the microbiological studies and antibacterial analyses. A.S.D. provided design and synthetic insight in library development and analog synthesis. A.S.D., H.I.Z. and V.V.R. interpreted structure-activity and structure-uptake relationships.

## Competing Interests

The authors declare no competing interests.

## ACKNOWLEDGMENTS

This work was supported by the National Institute of Allergy and Infectious Disease of the National Institutes of Health (R01AI136795, H.I.Z., V.V.R., and A.S.D.) The content included in this manuscript does not necessarily reflect the position or the policy of the federal government, and no official endorsement should be inferred. The authors thank Dr. Alexander S. Mankin for sharing *E. coli* strains. The authors gratefully acknowledge the collegiality, collaboration, and scientific discussion provided by the other SPEAR-GN team members.

## For Table of Contents Only

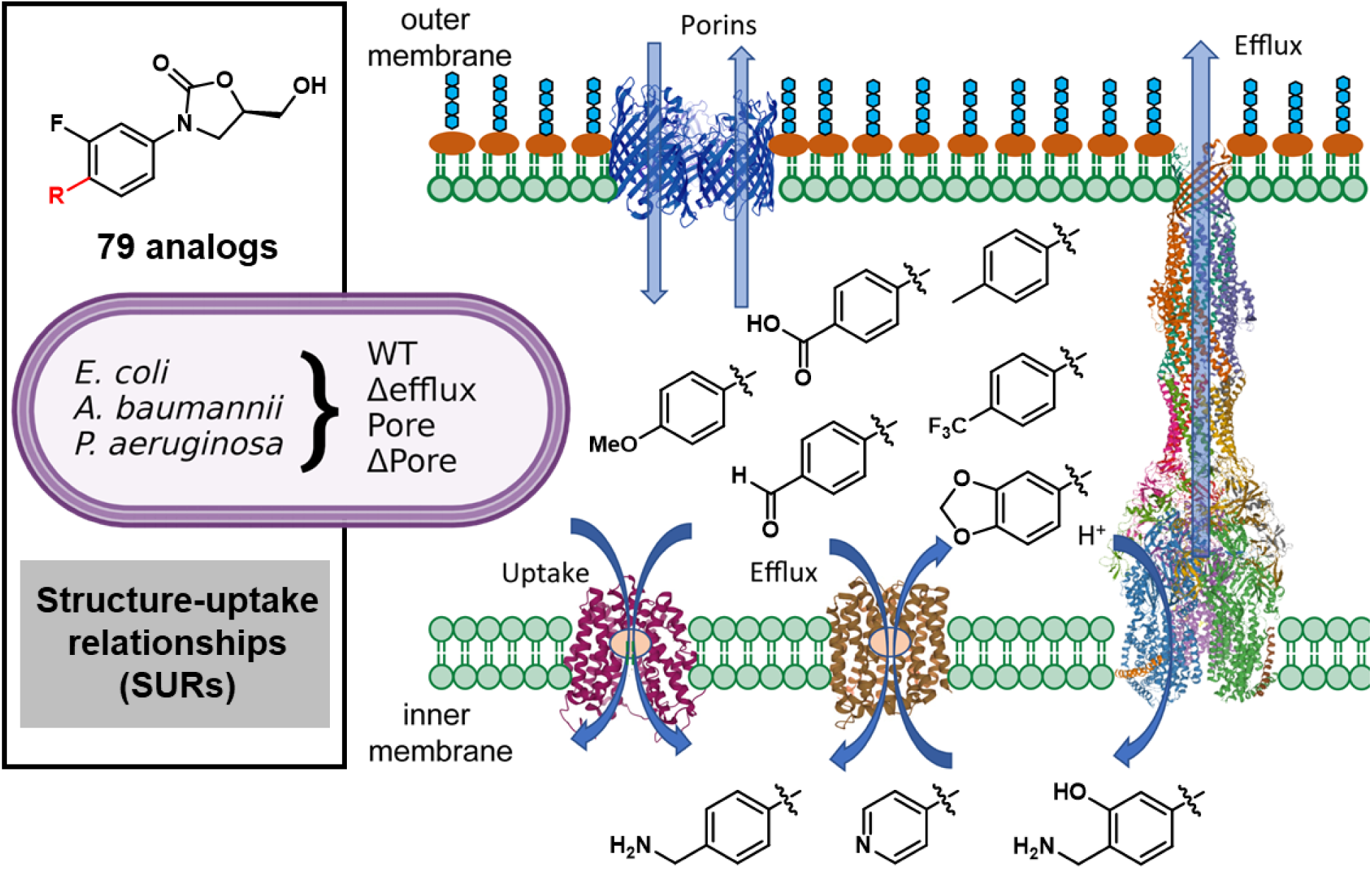

## Notes

### Competing Interest Statement

The authors have declared no competing interest.

## REFERENCES

1. Fischbach, M. A.; Walsh, C. T., Antibiotics for Emerging Pathogens. Science 2009, 325, 1089–1093.

2. Boucher, H. W.; Talbot, G. H.; Bradley, J. S.; Edwards, J. E.; Gilbert, D.; Rice, L. B.; Scheld, M.; Spellberg, B.; Bartlett, J., Bad Bugs, No Drugs: No Eskape! An Update from the Infectious Diseases Society of America. Clin. Infect. Dis. 2009, 48, 1–12.

3. Antibiotic Resistance Threats in the United States. US Centers for Disease Control and Prevention (CDC); 2019.

4. Silver, L. L., Challenges of Antibacterial Discovery. Clin. Microbiol. Rev. 2011, 24, 71–109.

5. Spellberg, B.; Guidos, R.; Gilbert, D.; Bradley, J.; Boucher, H. W.; Scheld, W. M.; Bartlett, J. G.; Edwards, J., Jr.; Infectious Diseases Society of, A., The Epidemic of Antibiotic-Resistant Infections: A Call to Action for the Medical Community from the Infectious Diseases Society of America. Clin. Infect. Dis. 2008, 46, 155–164.

6. Rice, L. B., Federal Funding for the Study of Antimicrobial Resistance in Nosocomial Pathogens: No Eskape. J. Infect. Dis. 2008, 197, 1079–1081.

7. Zgurskaya, H. I.; Lopez, C. A.; Gnanakaran, S., Permeability Barrier of Gram-Negative Cell Envelopes and Approaches to Bypass It. ACS Infect. Dis. 2015, 1, 512–522.

8. Rybenkov, V. V.; Zgurskaya, H. I.; Ganguly, C.; Leus, I. V.; Zhang, Z.; Moniruzzaman, M., The Whole Is Bigger Than the Sum of Its Parts: Drug Transport in the Context of Two Membranes with Active Efflux. Chem. Rev. 2021, 121, 5597–5631.

9. Brown, D. G.; May-Dracka, T. L.; Gagnon, M. M.; Tommasi, R., Trends and Exceptions of Physical Properties on Antibacterial Activity for Gram-Positive and Gram-Negative Pathogens. J. Med. Chem. 2014, 57, 10144–10161.

10. Payne, D. J.; Gwynn, M. N.; Holmes, D. J.; Pompliano, D. L., Drugs for Bad Bugs: Confronting the Challenges of Antibacterial Discovery. Nat. Rev. Drug Discovery 2007, 6, 29–40.

11. Tommasi, R.; Brown, D. G.; Walkup, G. K.; Manchester, J. I.; Miller, A. A., Eskapeing the Labyrinth of Antibacterial Discovery. Nat Rev Drug Discov 2015, 14, 529–542.

12. Moser, R. O. S. a. H. E., Physicochemical Properties of Antibacterial Compounds: Implications for Drug Discovery. J. Med. Chem. 2008, 51, 2871–2879.

13. Richter, M. F.; Hergenrother, P. J., The Challenge of Converting Gram-Positive-Only Compounds into Broad-Spectrum Antibiotics. Annals of the New York Academy of Sciences 2019, 1435, 18–38.

14. Richter, M. F.; Drown, B. S.; Riley, A. P.; Garcia, A.; Shirai, T.; Svec, R. L.; Hergenrother, P. J., Predictive Compound Accumulation Rules Yield a Broad-Spectrum Antibiotic. Nature 2017, 545, 299–304.

15. Lukezic, T.; Fayad, A. A.; Bader, C.; Harmrolfs, K.; Bartuli, J.; Gross, S.; Lesnik, U.; Hennessen, F.; Herrmann, J.; Pikl, S.; Petkovic, H.; Muller, R., Engineering Atypical Tetracycline Formation in Amycolatopsis Sulphurea for the Production of Modified Chelocardin Antibiotics. ACS Chem. Biol. 2019, 14, 468–477.

16. Perlmutter, S. J.; Geddes, E. J.; Drown, B. S.; Motika, S. E.; Lee, M. R.; Hergenrother, P. J., Compound Uptake into *E. Coli* Can Be Facilitated by N-Alkyl Guanidiniums and Pyridiniums. ACS Infect. Dis. 2021, 7, 162–173.

17. Li, Y.; Gardner, J. J.; Fortney, K. R.; Leus, I. V.; Bonifay, V.; Zgurskaya, H. I.; Pletnev, A. A.; Zhang, S.; Zhang, Z. Y.; Gribble, G. W.; Spinola, S. M.; Duerfeldt, A. S., First-Generation Structure-Activity Relationship Studies of 2,3,4,9-Tetrahydro-1h-Carbazol-1-Amines as Cpxa Phosphatase Inhibitors. Bioorg. Med. Chem. Lett. 2019, 29, 1836–1841.

18. Cohen, F.; Aggen, J. B.; Andrews, L. D.; Assar, Z.; Boggs, J.; Choi, T.; Dozzo, P.; Easterday, A. N.; Haglund, C. M.; Hildebrandt, D. J.; Holt, M. C.; Joly, K.; Jubb, A.; Kamal, Z.; Kane, T. R.; Konradi, A. W.; Krause, K. M.; Linsell, M. S.; Machajewski, T. D.; Miroshnikova, O.; Moser, H. E.; Nieto, V.; Phan, T.; Plato, C.; Serio, A. W.; Seroogy, J.; Shakhmin, A.; Stein, A. J.; Sun, A. D.; Sviridov, S.; Wang, Z.; Wlasichuk, K.; Yang, W.; Zhou, X.; Zhu, H.; Cirz, R. T., Optimization of LpxC Inhibitors for Antibacterial Activity and Cardiovascular Safety. ChemMedChem 2019, 14, 1560–1572.

19. Masci, D.; Hind, C.; Islam, M. K.; Toscani, A.; Clifford, M.; Coluccia, A.; Conforti, I.; Touitou, M.; Memdouh, S.; Wei, X.; La Regina, G.; Silvestri, R.; Sutton, J. M.; Castagnolo, D., Switching on the Activity of 1,5-Diaryl-Pyrrole Derivatives against Drug-Resistant Eskape Bacteria: Structure-Activity Relationships and Mode of Action Studies. Eur. J. Med. Chem. 2019, 178, 500–514.

20. Motika, S. E.; Ulrich, R. J.; Geddes, E. J.; Lee, H. Y.; Lau, G. W.; Hergenrother, P. J., Gram-Negative Antibiotic Active through Inhibition of an Essential Riboswitch. J. Am. Chem. Soc. 2020, 142, 10856–10862.

21. Andrews, L. D.; Kane, T. R.; Dozzo, P.; Haglund, C. M.; Hilderbrandt, D. J.; Linsell, M. S.; Machajewski, T.; McEnroe, G.; Serio, A. W.; Wlasichuk, K. B.; Neau, D. B.; Pakhomova, S.; Waldrop, G. L.; Sharp, M.; Pogliano, J.; Cirz, R. T.; Cohen, F., Optimization and Mechanistic Characterization of Pyridopyrimidine Inhibitors of Bacterial Biotin Carboxylase. J. Med. Chem. 2019, 62, 7489–7505.

22. Hu, Y.; Shi, H.; Zhou, M.; Ren, Q.; Zhu, W.; Zhang, W.; Zhang, Z.; Zhou, C.; Liu, Y.; Ding, X.; Shen, H. C.; Yan, S. F.; Dey, F.; Wu, W.; Zhai, G.; Zhou, Z.; Xu, Z.; Ji, Y.; Lv, H.; Jiang, T.; Wang, W.; Xu, Y.; Vercruysse, M.; Yao, X.; Mao, Y.; Yu, X.; Bradley, K.; Tan, X., Discovery of Pyrido[2,3-B]Indole Derivatives with Gram-Negative Activity Targeting Both DNA Gyrase and Topoisomerase Iv. J. Med. Chem. 2020, 63, 9623–9649.

23. Parker, E. N.; Drown, B. S.; Geddes, E. J.; Lee, H. Y.; Ismail, N.; Lau, G. W.; Hergenrother, P. J., Implementation of Permeation Rules Leads to a Fabi Inhibitor with Activity against Gram-Negative Pathogens. Nat. Microbiol. 2020, 5, 67–75.

24. Sohlenkamp, C.; Geiger, O., Bacterial Membrane Lipids: Diversity in Structures and Pathways. FEMS Microbiol. Rev. 2016, 40, 133–159.

25. Leach, K. L.; Swaney, S. M.; Colca, J. R.; McDonald, W. G.; Blinn, J. R.; Thomasco, L. M.; Gadwood, R. C.; Shinabarger, D.; Xiong, L.; Mankin, A. S., The Site of Action of Oxazolidinone Antibiotics in Living Bacteria and in Human Mitochondria. Mol. Cell 2007, 26, 393–402.

26. Pucci, M. J.; Bush, K., Investigational Antimicrobial Agents of 2013. Clin. Microbiol. Rev. 2013, 26, 792–821.

27. Endimiani, A.; Blackford, M.; Dasenbrook, E. C.; Reed, M. D.; Bajaksouszian, S.; Hujer, A. M.; Rudin, S. D.; Hujer, K. M.; Perreten, V.; Rice, L. B.; Jacobs, M. R.; Konstan, M. W.; Bonomo, R. A., Emergence of Linezolid-Resistant *Staphylococcus Aureus* after Prolonged Treatment of Cystic Fibrosis Patients in Cleveland, Ohio. Antimicrob. Agents Chemother. 2011, 55, 1684–1692.

28. Long, K. S.; Vester, B., Resistance to Linezolid Caused by Modifications at Its Binding Site on the Ribosome. Antimicrobial Agents and Chemotherapy 2012, 56, 603–612.

29. Xiong, L.; Kloss, P.; Douthwaite, S.; Andersen, N. M.; Swaney, S.; Shinabarger, D. L.; Mankin, A. S., Oxazolidinone Resistance Mutations in 23s Rrna of Escherichia Coli Reveal the Central Region of Domain V as the Primary Site of Drug Action. J Bacteriol 2000, 182, 5325–5331.

30. Schwarz, S.; Zhang, W.; Du, X. D.; Kruger, H.; Fessler, A. T.; Ma, S.; Zhu, Y.; Wu, C.; Shen, J.; Wang, Y., Mobile Oxazolidinone Resistance Genes in Gram-Positive and Gram-Negative Bacteria. Clin Microbiol Rev 2021, 34, e0018820.

31. Sander, P.; Belova, L.; Kidan, Y. G.; Pfister, P.; Mankin, A. S.; Böttger, E. C., Ribosomal and Non-Ribosomal Resistance to Oxazolidinones: Species-Specific Idiosyncrasy of Ribosomal Alterations. Mol Microbiol 2002, 46, 1295–1304.

32. Floyd, J. L.; Smith, K. P.; Kumar, S. H.; Floyd, J. T.; Varela, M. F., Lmrs Is a Multidrug Efflux Pump of the Major Facilitator Superfamily from Staphylococcus Aureus. Antimicrob Agents Chemother 2010, 54, 5406–5412.

33. Feng, J.; Lupien, A.; Gingras, H.; Wasserscheid, J.; Dewar, K.; Légaré, D.; Ouellette, M., Genome Sequencing of Linezolid-Resistant Streptococcus Pneumoniae Mutants Reveals Novel Mechanisms of Resistance. Genome Research 2009, 19, 1214–1223.

34. Michalska, K.; Karpiuk, I.; Krol, M.; Tyski, S., Recent Development of Potent Analogues of Oxazolidinone Antibacterial Agents. Bioorg. Med. Chem. Lett. 2013, 21, 577–591.

35. Poce, G.; Zappia, G.; Cesare Porretta, G.; Botta, B.; Biava, M., New Oxazolidinone Derivatives as Antibacterial Agents with Improved Activity. Exp. Opin. Ther. Patents 2008, 18, 97–121.

36. Renslo, A. R.; Luehr, G. W.; Gordeev, M. F., Recent Developments in the Identification of Novel Oxazolidinone Antibacterial Agents. Bioorg. Med. Chem. 2006, 14, 4227–4240.

37. Deshmukh, M. S.; Jain, N., Design, Synthesis, and Antibacterial Evaluation of Oxazolidinones with Fused Heterocyclic C-Ring Substructure. ACS Med. Chem. Lett. 2017, 8, 1153–1158.

38. Wu, Y.; Ding, X.; Ding, L.; Zhang, Y.; Cui, L.; Sun, L.; Li, W.; Wang, D.; Zhao, Y., Synthesis and Antibacterial Activity Evaluation of Novel Biaryloxazolidinone Analogues Containing a Hydrazone Moiety as Promising Antibacterial Agents. Eur. J. Med. Chem. 2018, 158, 247–258.

39. Xin, Q.; Fan, H.; Guo, B.; He, H.; Gao, S.; Wang, H.; Huang, Y.; Yang, Y., Design, Synthesis, and Structure-Activity Relationship Studies of Highly Potent Novel Benzoxazinyl-Oxazolidinone Antibacterial Agents. J. Med. Chem. 2011, 54, 7493–7502.

40. Seetharamsingh, B.; Ramesh, R.; Dange, S. S.; Khairnar, P. V.; Singhal, S.; Upadhyay, D.; Veeraraghavan, S.; Viswanadha, S.; Vakkalanka, S.; Reddy, D. S., Design, Synthesis, and Identification of Silicon Incorporated Oxazolidinone Antibiotics with Improved Brain Exposure. ACS Med. Chem. Lett. 2015, 6, 1105–1110.

41. Yang, T.; Chen, G.; Sang, Z.; Liu, Y.; Yang, X.; Chang, Y.; Long, H.; Ang, W.; Tang, J.; Wang, Z.; Li, G.; Yang, S.; Zhang, J.; Wei, Y.; Luo, Y., Discovery of a Teraryl Oxazolidinone Compound (S)-N-((3-(3-Fluoro-4-(4-(Pyridin-2-Yl)-1h-Pyrazol-1-Yl)Phenyl)-2-Oxooxazolidin-5 -Yl)Methyl)Acetamide Phosphate as a Novel Antimicrobial Agent with Enhanced Safety Profile and Efficacies. J. Med. Chem. 2015, 58, 6389–6409.

42. Paulen, A.; Gasser, V.; Hoegy, F.; Perraud, Q.; Pesset, B.; Schalk, I. J.; Mislin, G. L., Synthesis and Antibiotic Activity of Oxazolidinone-Catechol Conjugates against *Pseudomonas Aeruginosa*. Org. Biomol. Chem. 2015, 13, 11567–11579.

43. Spaulding, A.; Takrouri, K.; Mahalingam, P.; Cleary, D. C.; Cooper, H. D.; Zucchi, P.; Tear, W.; Koleva, B.; Beuning, P. J.; Hirsch, E. B.; Aggen, J. B., Compound Design Guidelines for Evading the Efflux and Permeation Barriers of *Escherichia Coli* with the Oxazolidinone Class of Antibacterials: Test Case for a General Approach to Improving Whole Cell Gram-Negative Activity. Bioorg. Med. Chem. Lett. 2017, 27, 5310–5321.

44. Suzuki, H.; Utsunomiya, I.; Shudo, K.; Fukuhara, N.; Iwaki, T.; Yasukata, T., Antibacterial Oxazolidinone Analogues Having a N-Hydroxyacetyl-Substituted Seven-Membered [1,2,5]Triazepane or [1,2,5]Oxadiazepane C-Ring Unit. Eur. J. Med. Chem. 2013, 63, 811–825.

45. Takrouri, K.; Cooper, H. D.; Spaulding, A.; Zucchi, P.; Koleva, B.; Cleary, D. C.; Tear, W.; Beuning, P. J.; Hirsch, E. B.; Aggen, J. B., Progress against *Escherichia Coli* with the Oxazolidinone Class of Antibacterials: Test Case for a General Approach to Improving Whole-Cell Gram-Negative Activity. ACS Infect. Dis. 2016, 2, 405–426.

46. Eyal, Z.; Matzov, D.; Krupkin, M.; Wekselman, I.; Paukner, S.; Zimmerman, E.; Rozenberg, H.; Bashan, A.; Yonath, A., Structural Insights into Species-Specific Features of the Ribosome from the Pathogen *Staphylococcus Aureus*. Proc. Natl. Acad. Sci. U.S.A. 2015, 112, E5805–E5814.

47. Krishnamoorthy, G.; Leus, I. V.; Weeks, J. W.; Wolloscheck, D.; Rybenkov, V. V.; Zgurskaya, H. I., Synergy between Active Efflux and Outer Membrane Diffusion Defines Rules of Antibiotic Permeation into Gram-Negative Bacteria. mBio 2017, 8.

48. Phenotypic Screening. Springer Nature: 2018.

49. Jelsbak, L.; Hartman, H.; Schroll, C.; Rosenkrantz, J. T.; Lemire, S.; Wallrodt, I.; Thomsen, L. E.; Poolman, M.; Kilstrup, M.; Jensen, P. R.; Olsen, J. E., Identification of Metabolic Pathways Essential for Fitness of *Salmonella* Typhimurium in Vivo. PLoS One 2014, 9, e101869.

50. Palmer, K. L.; Mashburn, L. M.; Singh, P. K.; Whiteley, M., Cystic Fibrosis Sputum Supports Growth and Cues Key Aspects of *Pseudomonas Aeruginosa* Physiology. J. Bacteriol. 2005, 187, 5267–5277.

51. Orelle, C.; Carlson, S.; Kaushal, B.; Almutairi, M. M.; Liu, H.; Ochabowicz, A.; Quan, S.; Pham, V. C.; Squires, C. L.; Murphy, B. T.; Mankin, A. S., Tools for Characterizing Bacterial Protein Synthesis Inhibitors. Antimicrob. Agents Chemother. 2013, 57, 5994–6004.

52. Topliss, J. G., Utilization of Operational Schemes for Analog Synthesis in Drug Design. J. Med. Chem. 1972, 15, 1006–1011.

53. Meanwell, N. A., Synopsis of Some Recent Tactical Application of Bioisosteres in Drug Design. J. Med. Chem. 2011, 54, 2529–2591.

54. Cooper, C. J.; Krishnamoorthy, G.; Wolloscheck, D.; Walker, J. K.; Rybenkov, V. V.; Parks, J. M.; Zgurskaya, H. I., Molecular Properties That Define the Activities of Antibiotics in *Escherichia Coli* and *Pseudomonas Aeruginosa*. ACS Infect. Dis. 2018, 4, 1223–1234.

55. Krishnamoorthy, G.; Leus, I. V.; Weeks, J. W.; Wolloscheck, D.; Rybenkov, V. V.; Zgurskaya, H. I., Synergy between Active Efflux and Outer Membrane Diffusion Defines Rules of Antibiotic Permeation into Gram-Negative Bacteria. mBio 2017, 8, e01172–01117.

56. Mehla, J.; Malloci, G.; Mansbach, R.; Lopez, C. A.; Tsivkovski, R.; Haynes, K.; Leus, I. V.; Grindstaff, S. B.; Cascella, R. H.; D’Cunha, N.; Herndon, L.; Hengartner, N. W.; Margiotta, E.; Atzori, A.; Vargiu, A. V.; Manrique, P. D.; Walker, J. K.; Lomovskaya, O.; Ruggerone, P.; Gnanakaran, S.; Rybenkov, V. V.; Zgurskaya, H. I., Predictive Rules of Efflux Inhibition and Avoidance in Pseudomonas Aeruginosa. mBio 2021, 12.

57. Nordqvist, A.; O’Mahony, G.; Friden-Saxin, M.; Fredenwall, M.; Hogner, A.; Granberg, K. L.; Aagaard, A.; Backstrom, S.; Gunnarsson, A.; Kaminski, T.; Xue, Y.; Dellsen, A.; Hansson, E.; Hansson, P.; Ivarsson, I.; Karlsson, U.; Bamberg, K.; Hermansson, M.; Georgsson, J.; Lindmark, B.; Edman, K., Structure-Based Drug Design of Mineralocorticoid Receptor Antagonists to Explore Oxosteroid Receptor Selectivity. ChemMedChem 2017, 12, 50–65.

58. Le, H. T.; Jang, J. G.; Park, J. Y.; Lim, C. W.; Kim, T. W., Antibody Functionalization with a Dual Reactive Hydrazide/Click Crosslinker. Anal. Biochem. 2013, 435, 68–73.

59. Lahasky, S. H.; Serem, W. K.; Guo, L.; Garno, J. C.; Zhang, D., Synthesis and Characterization of Cyclic Brush-Like Polymers by N-Heterocyclic Carbene-Mediated Zwitterionic Polymerization of N-Propargyl N-Carboxyanhydride and the Grafting-to Approach. Macromolecules 2011, 44, 9063–9074.

