## Supplemental Information for "Structure-Uptake Relationship Studies of Oxazolidinones in Gram-negative ESKAPE Pathogens"

*** Contributed equally**
**† Corresponding Authors**

**Figure S1.** Library design approach ........................................ **S1**

Representative HPLC Traces (**3e**, **12f**, and **14**) ........................................ **S2-S4**

$^{1}$H and $^{13}$C NMR spectra of final compounds ........................................ **S5-S83**

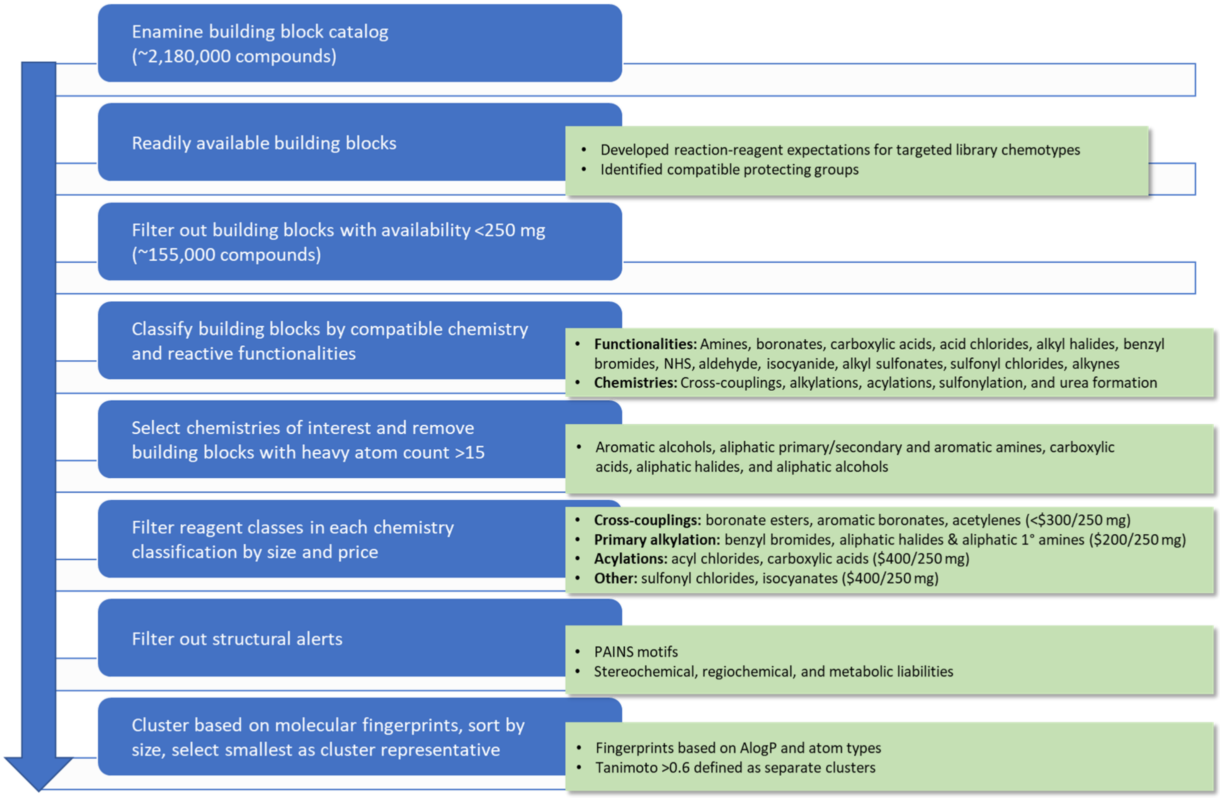


**Figure S1.** Library design approach.

### Representative HPLC Traces

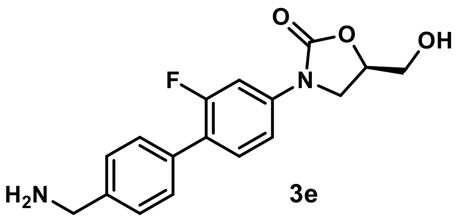


3e

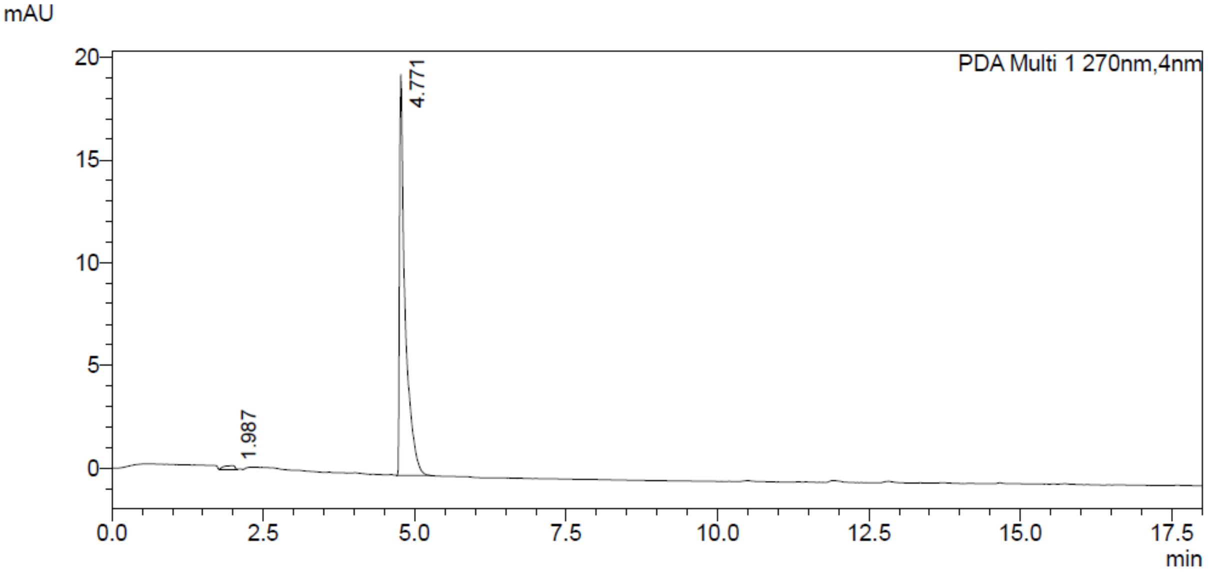


#### <Peak Table>

PDA Ch1 270nm

| Peak# | Ret. Time | Area | Height | Conc. | Unit | Mark | Name |
|---|---|---|---|---|---|---|---|
| 1 | 1.987 | 2193 | 168 | 1.712 | | | |
| 2 | 4.771 | 125906 | 19516 | 98.288 | | | |
| Total | | 128099 | 19684 | | | | |

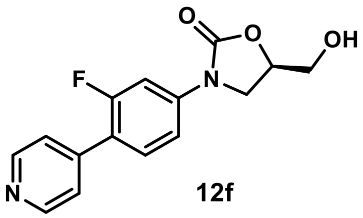

**12f**

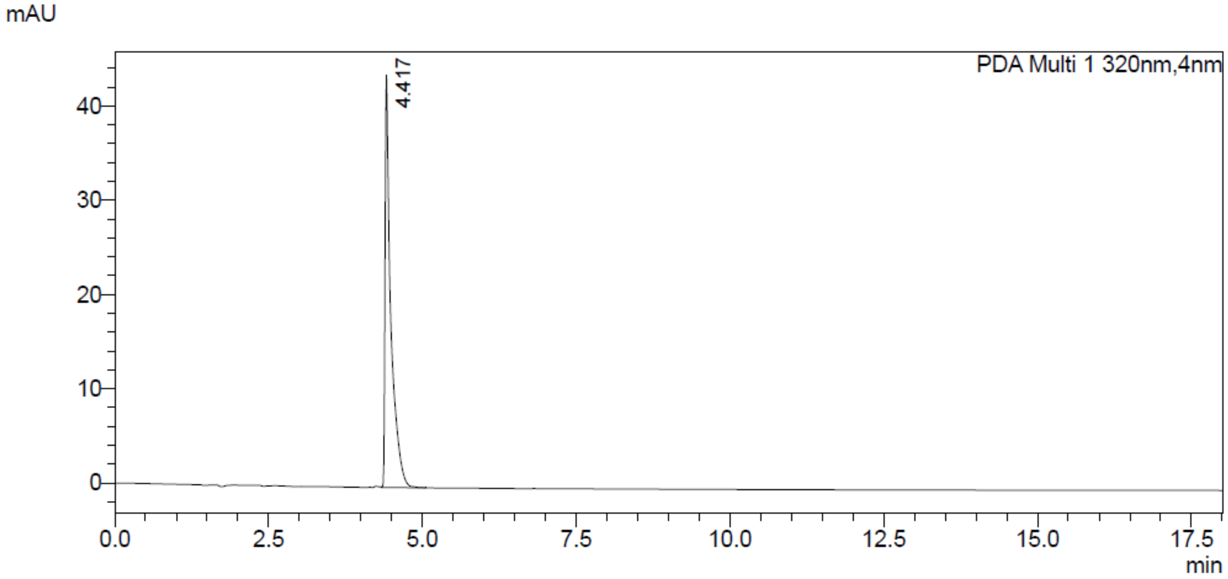


### <Peak Table>

PDA Ch1 320nm

| Peak# | Ret. Time | Area | Height | Conc. | Unit | Mark | Name |
|---|---|---|---|---|---|---|---|
| 1 | 4.417 | 285045 | 43759 | 100.000 | | V | |
| Total | | 285045 | 43759 | | | | |

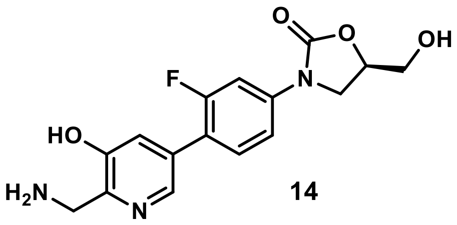


**14**

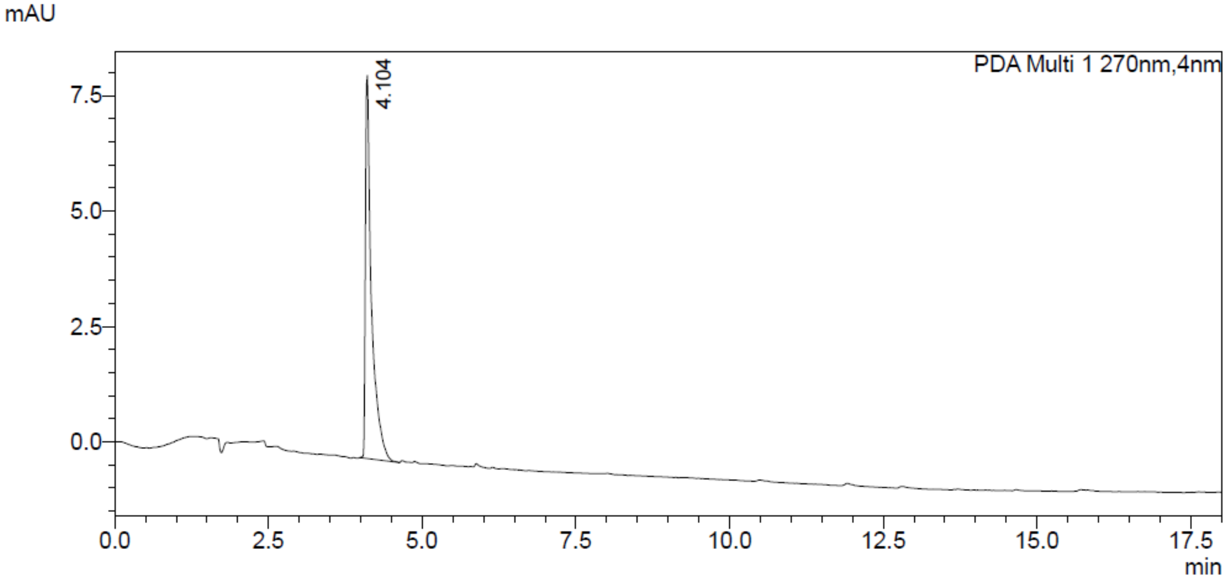


### <Peak Table>

PDA Ch1 270nm

| Peak# | Ret. Time | Area | Height | Conc. | Unit | Mark | Name |
|---|---|---|---|---|---|---|---|
| 1 | 4.104 | 57085 | 8293 | 100.000 | | | |
| Total | | 57085 | 8293 | | | | |

Compound **3a**: $^{1}$H and $^{13}$C NMR Spectrum

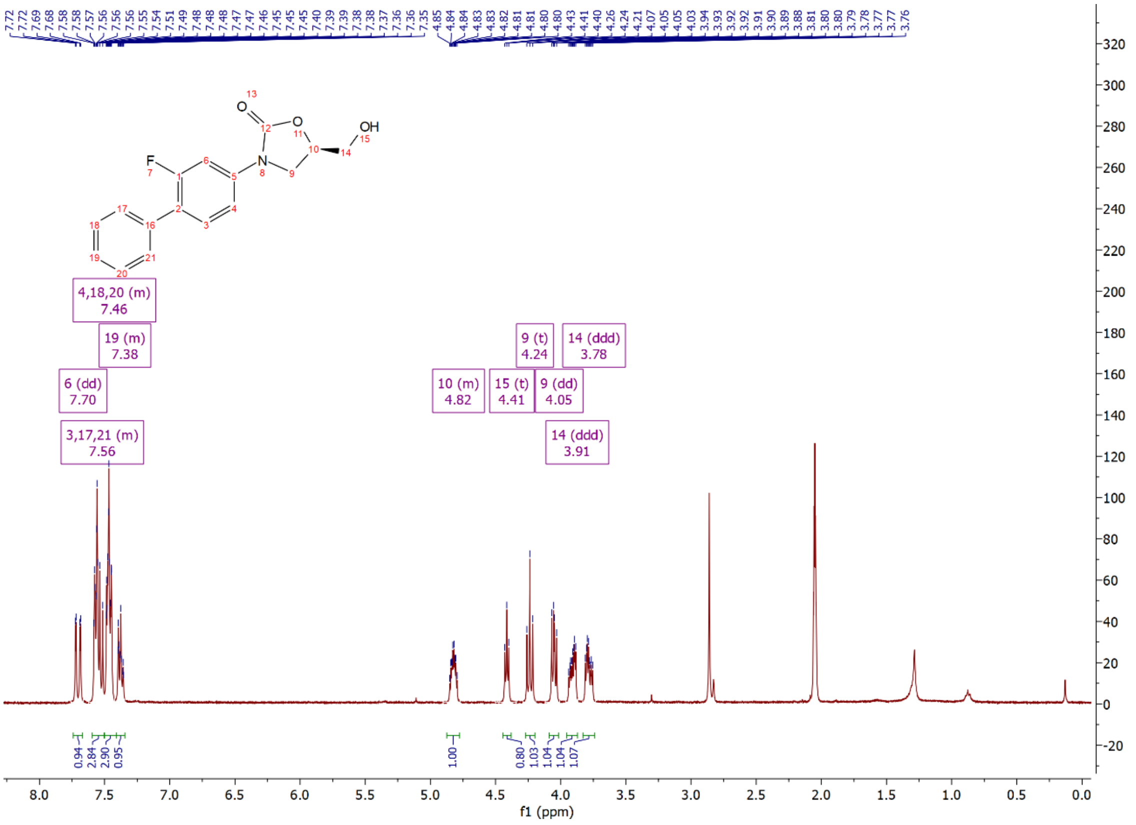

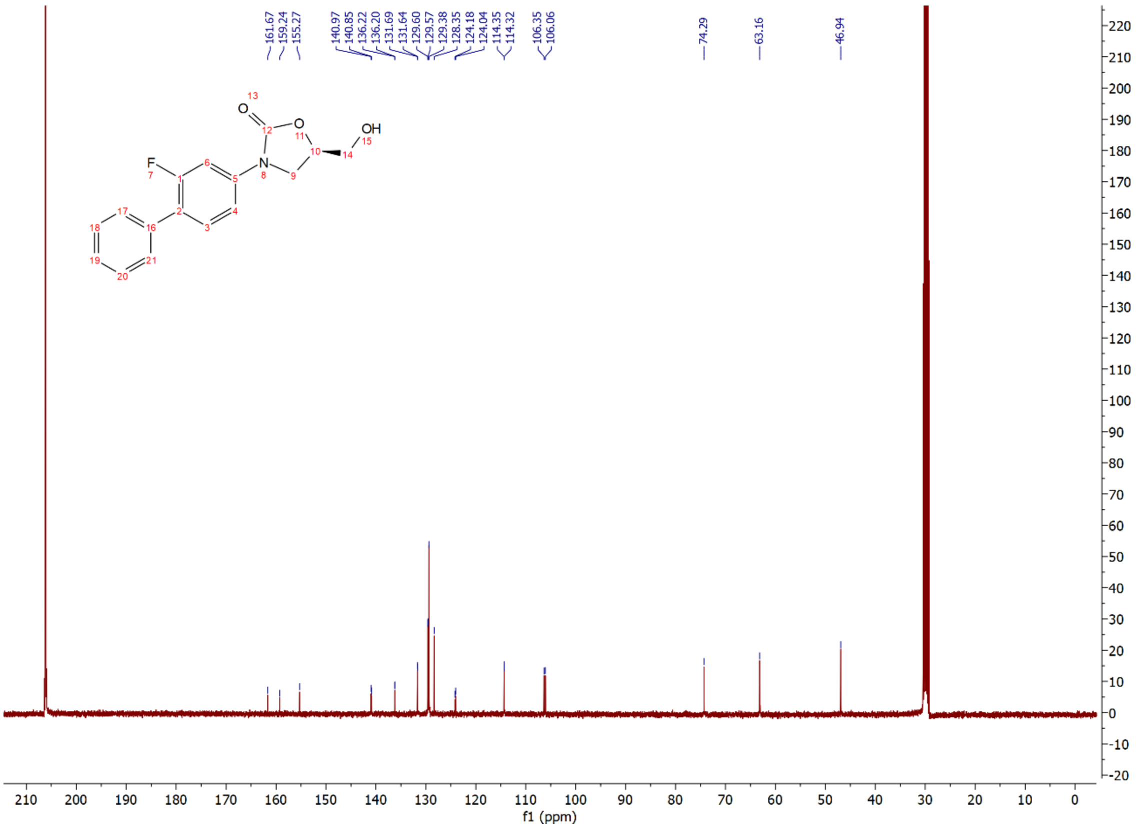

Compound **3b**: $^{1}$H and $^{13}$C NMR Spectrum

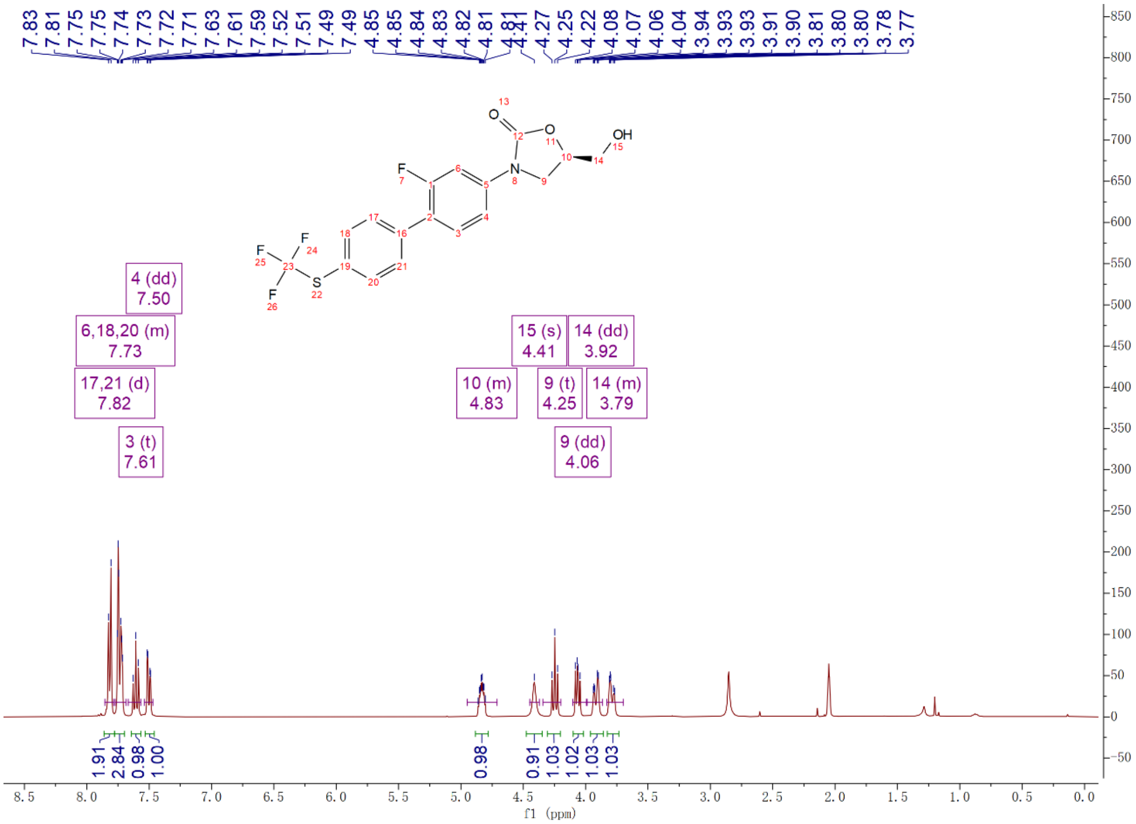

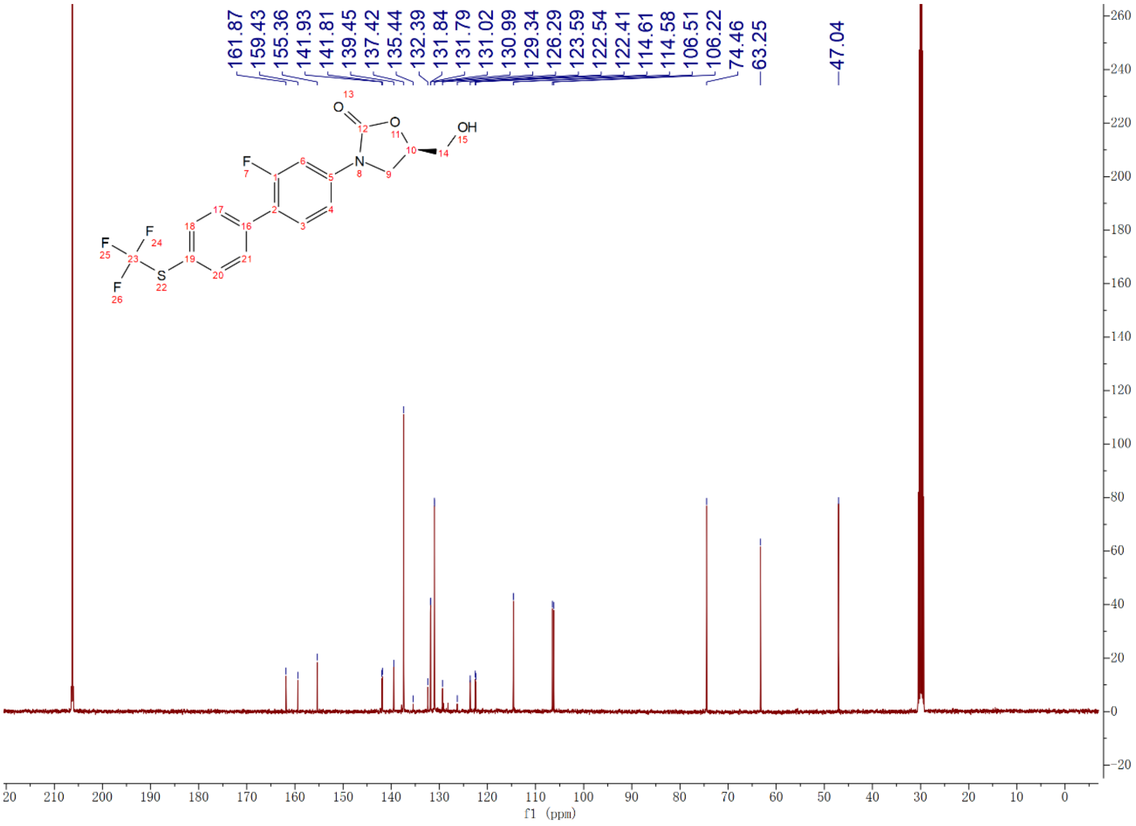

Compound **3c**: $^{1}$H and $^{13}$C NMR Spectrum

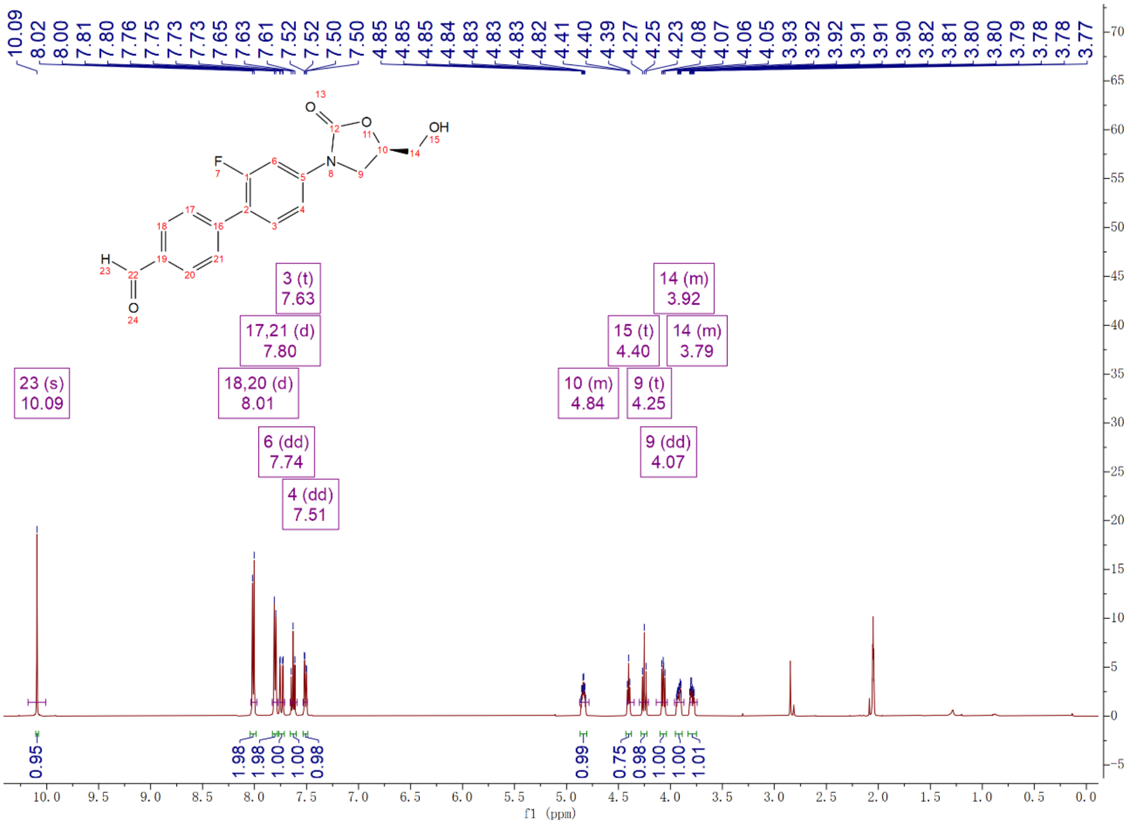


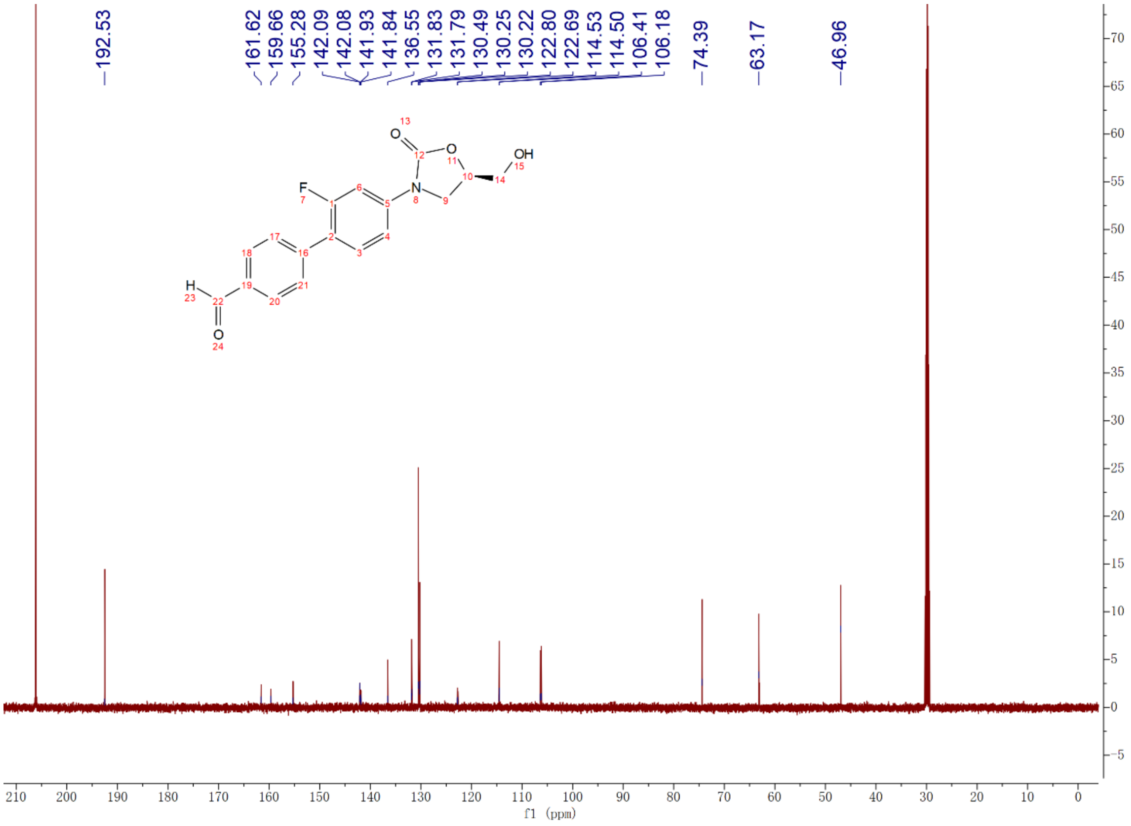

Compound **3d**: $^{1}$H and $^{13}$C NMR Spectrum

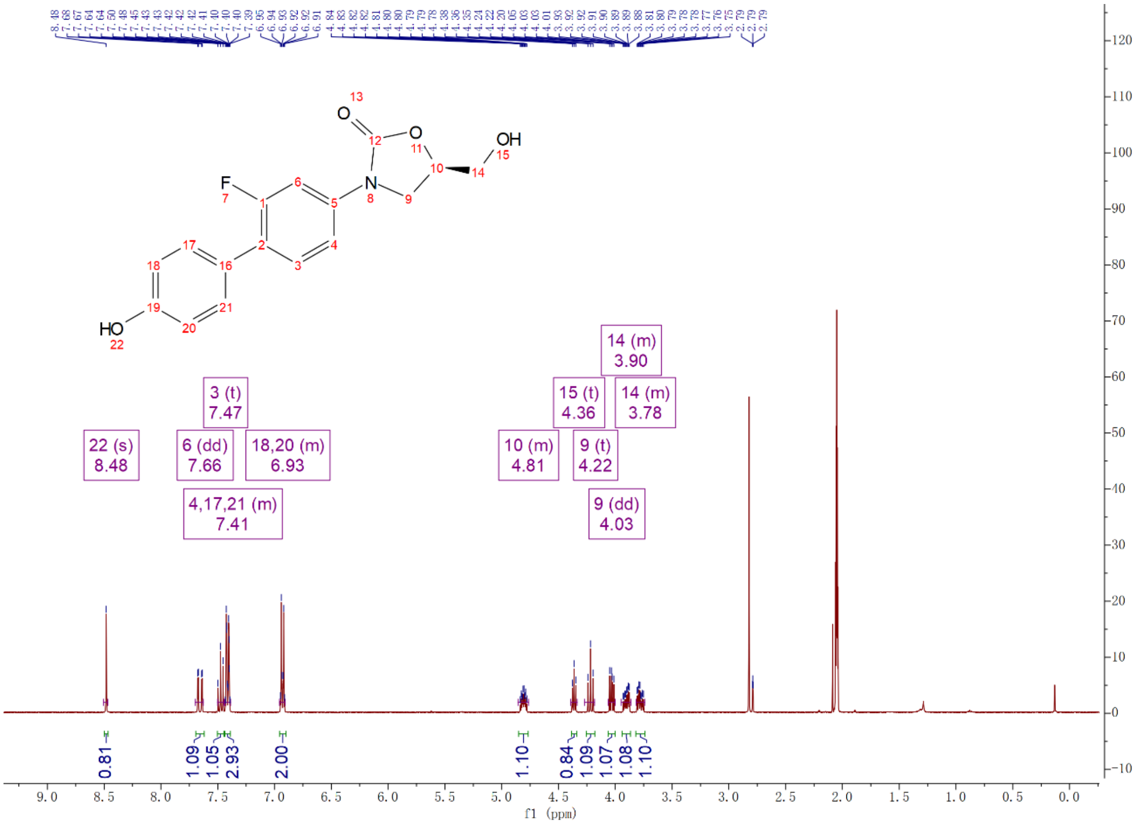


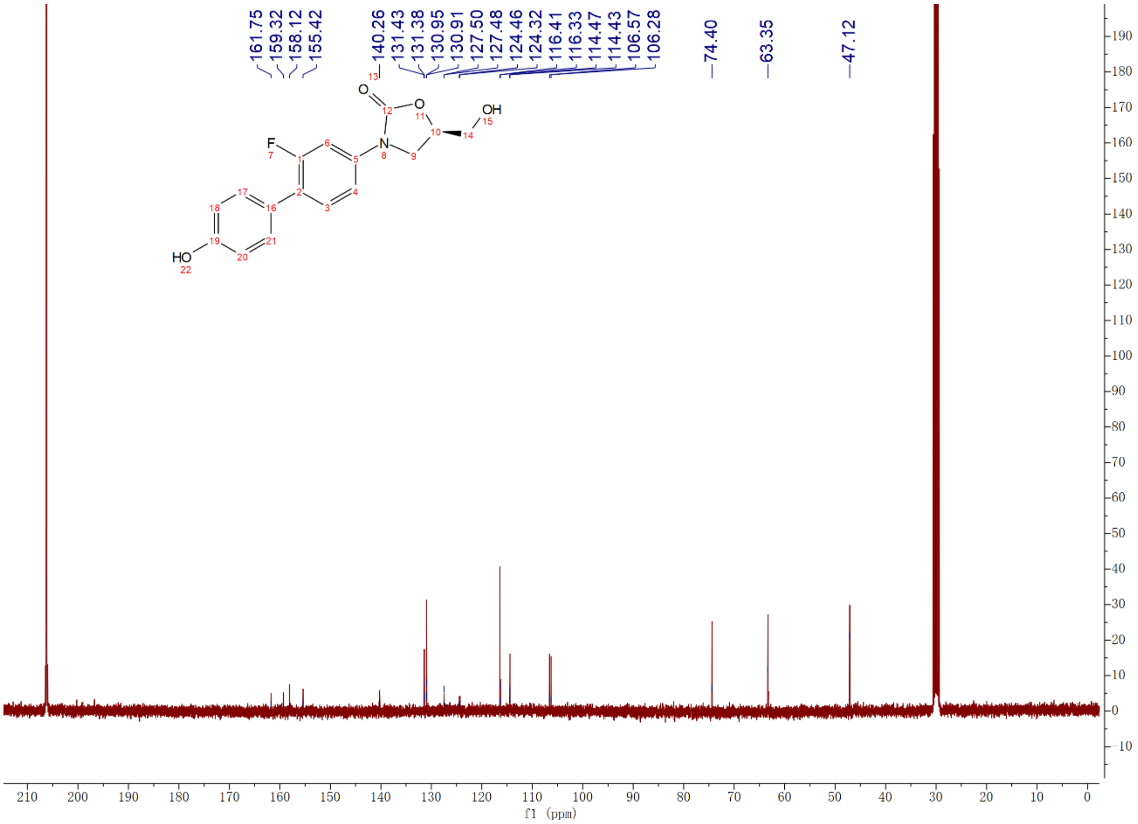

Compound **3e**: $^{1}H$ and $^{13}C$ NMR Spectrum

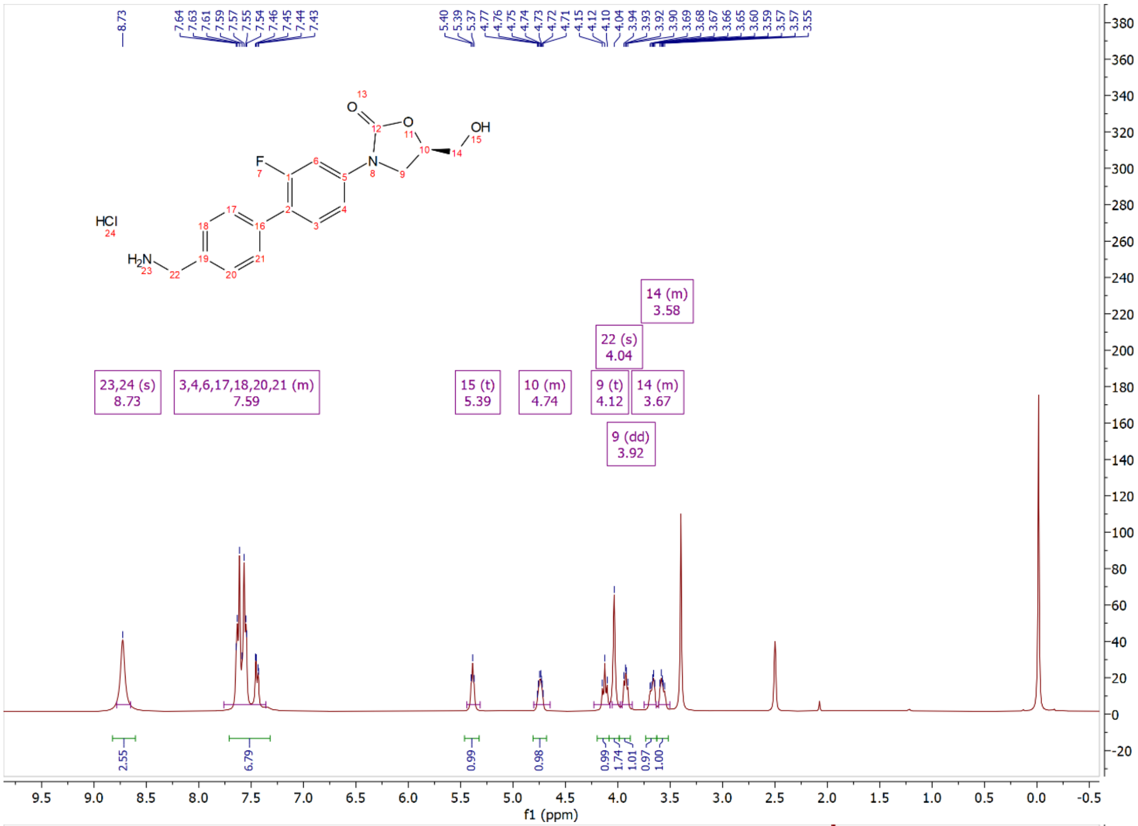

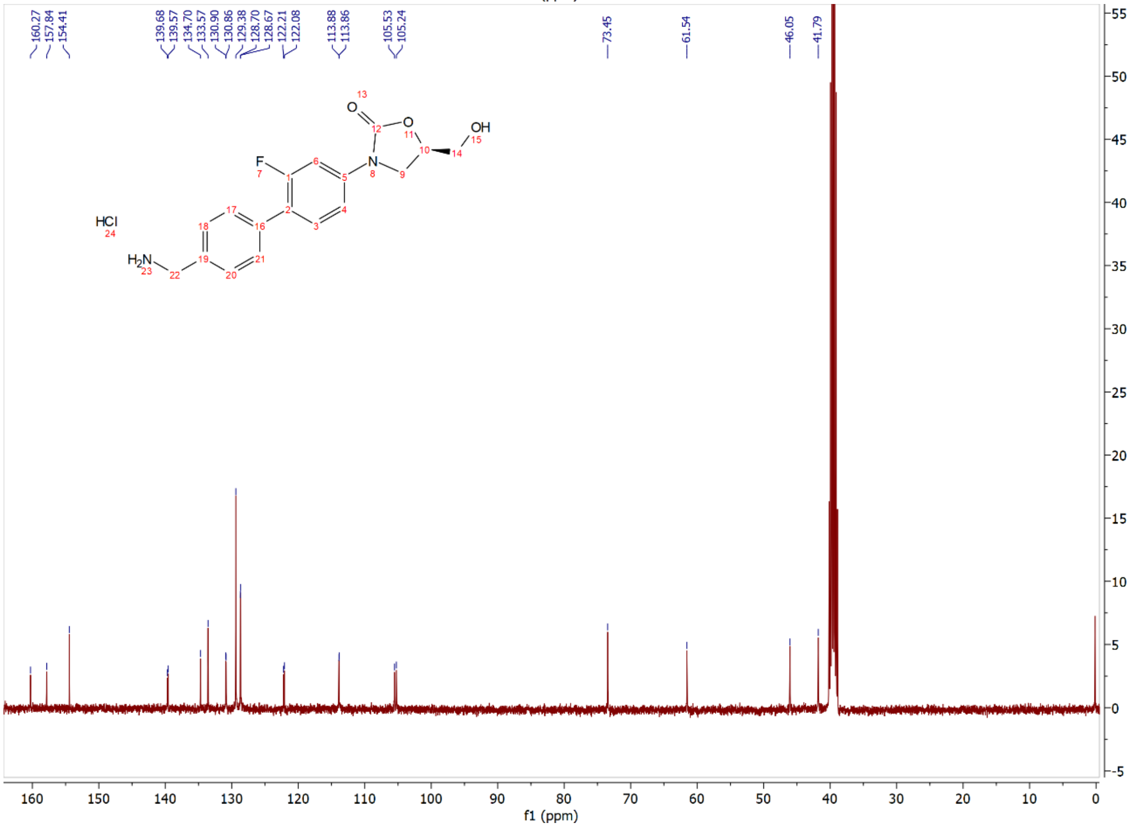

Compound **3eb**: $^{1}$H and $^{13}$C NMR Spectrum

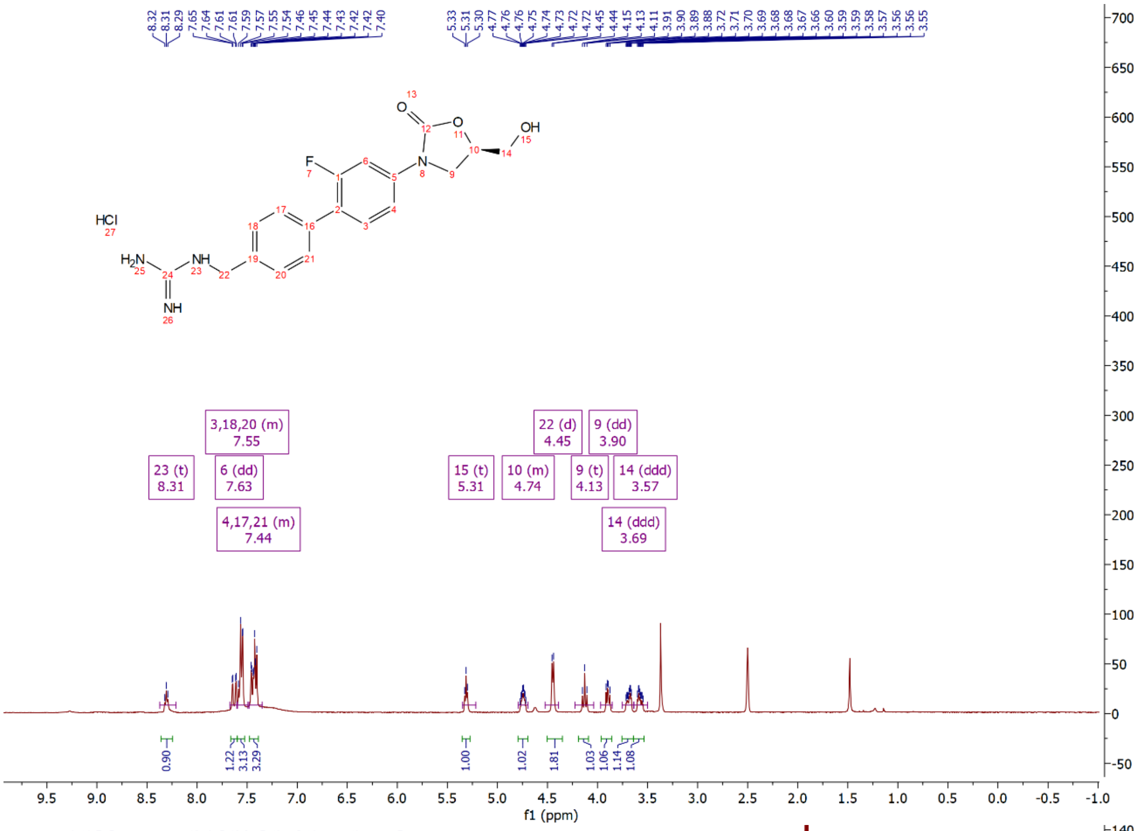


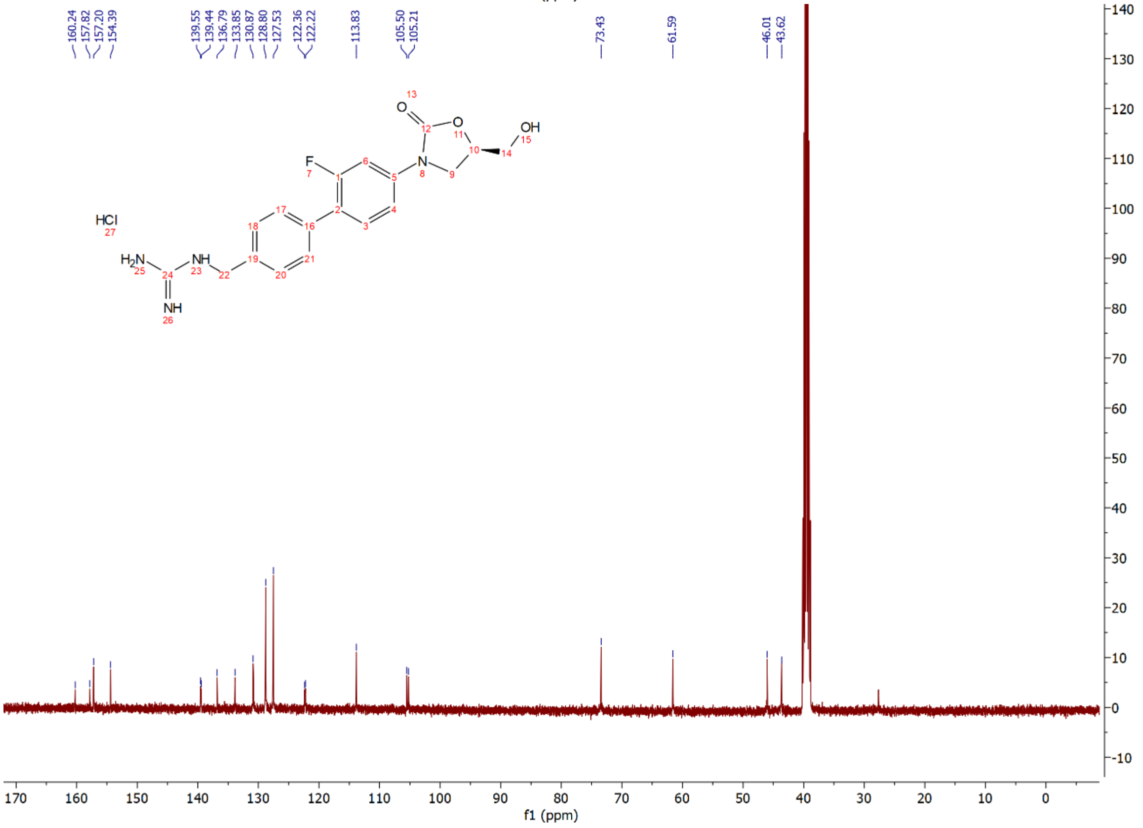

Compound **3f**: $^{1}$H and $^{13}$C NMR Spectrum

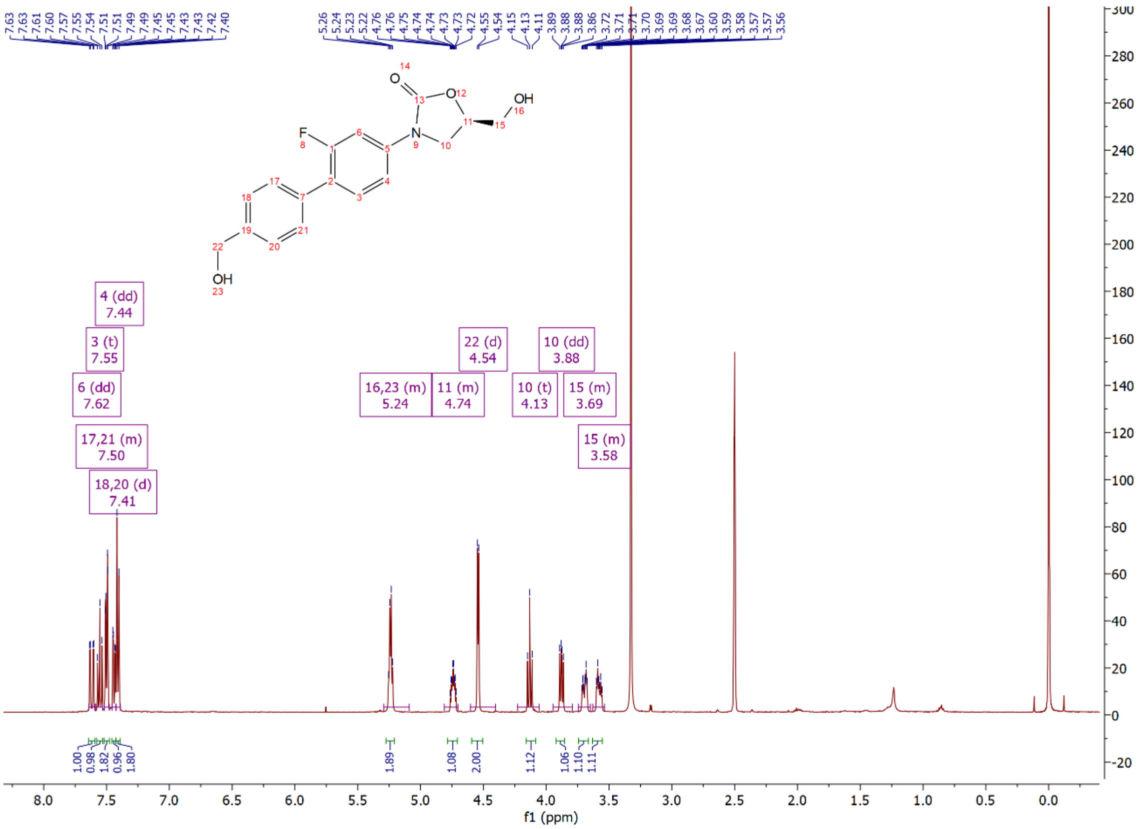

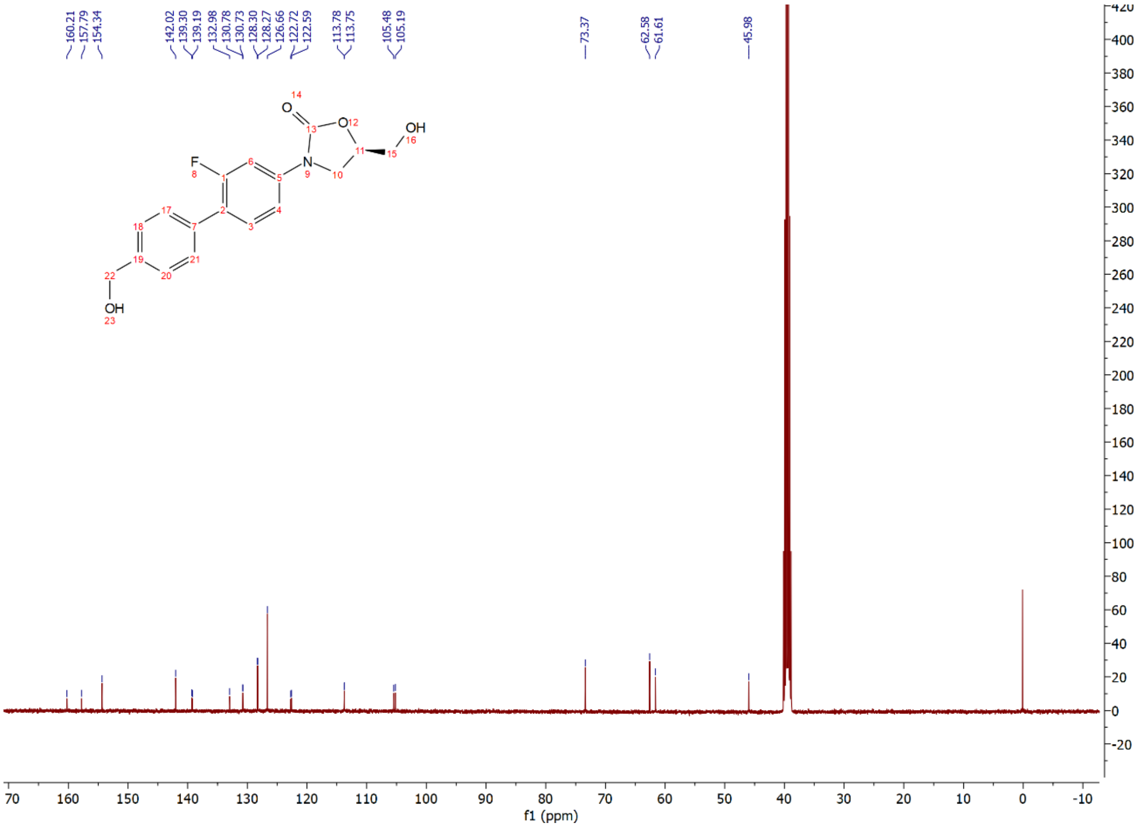

Compound **3g**: $^{1}$H and $^{13}$C NMR Spectrum

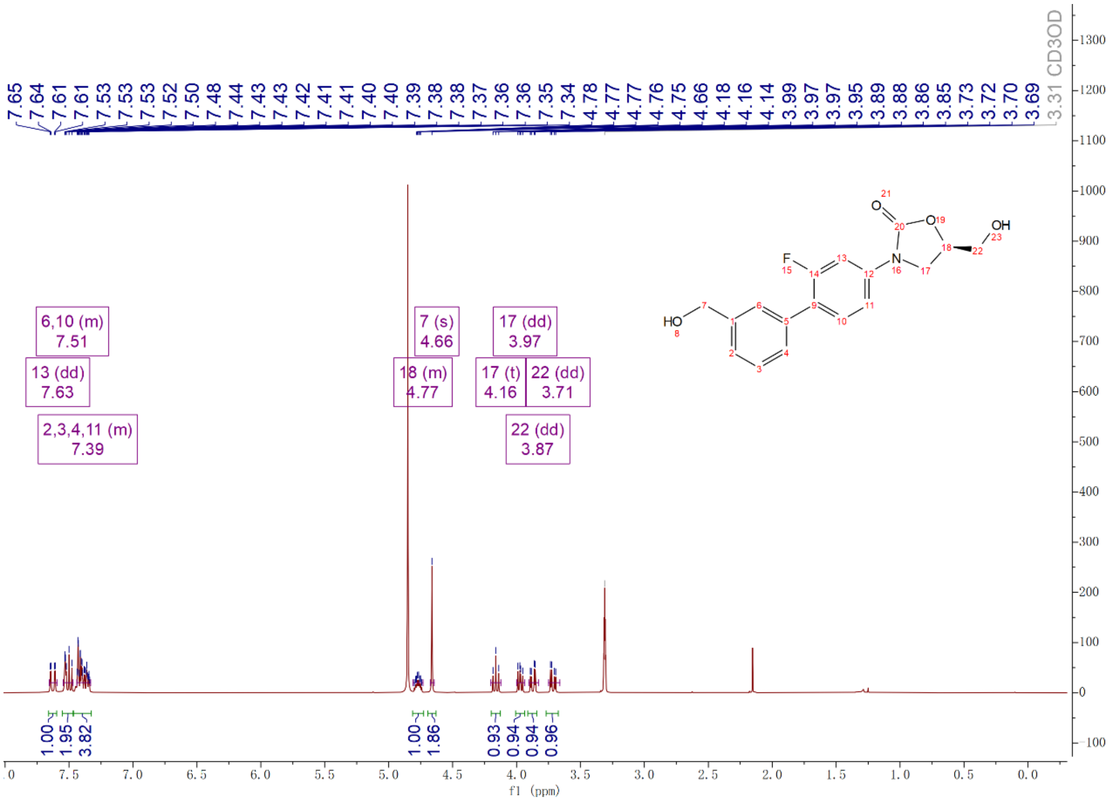

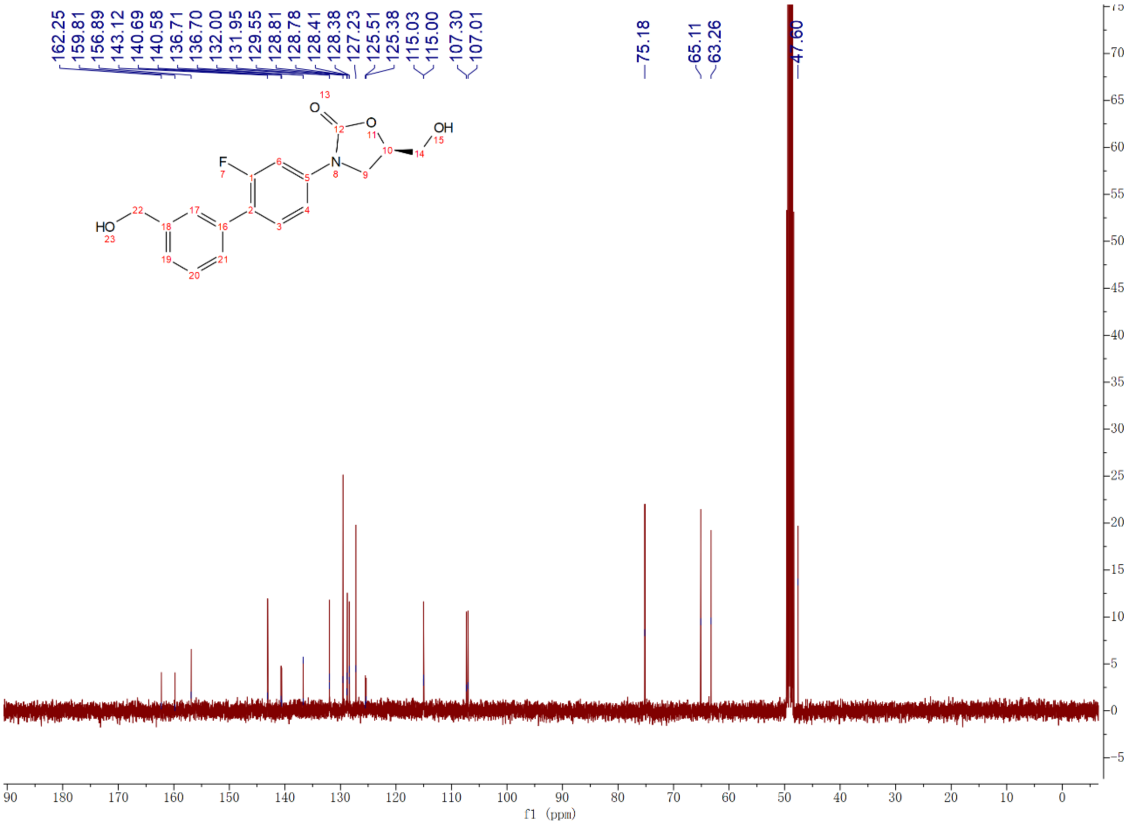

Compound **3h**: $^{1}H$ and $^{13}C$ NMR Spectrum

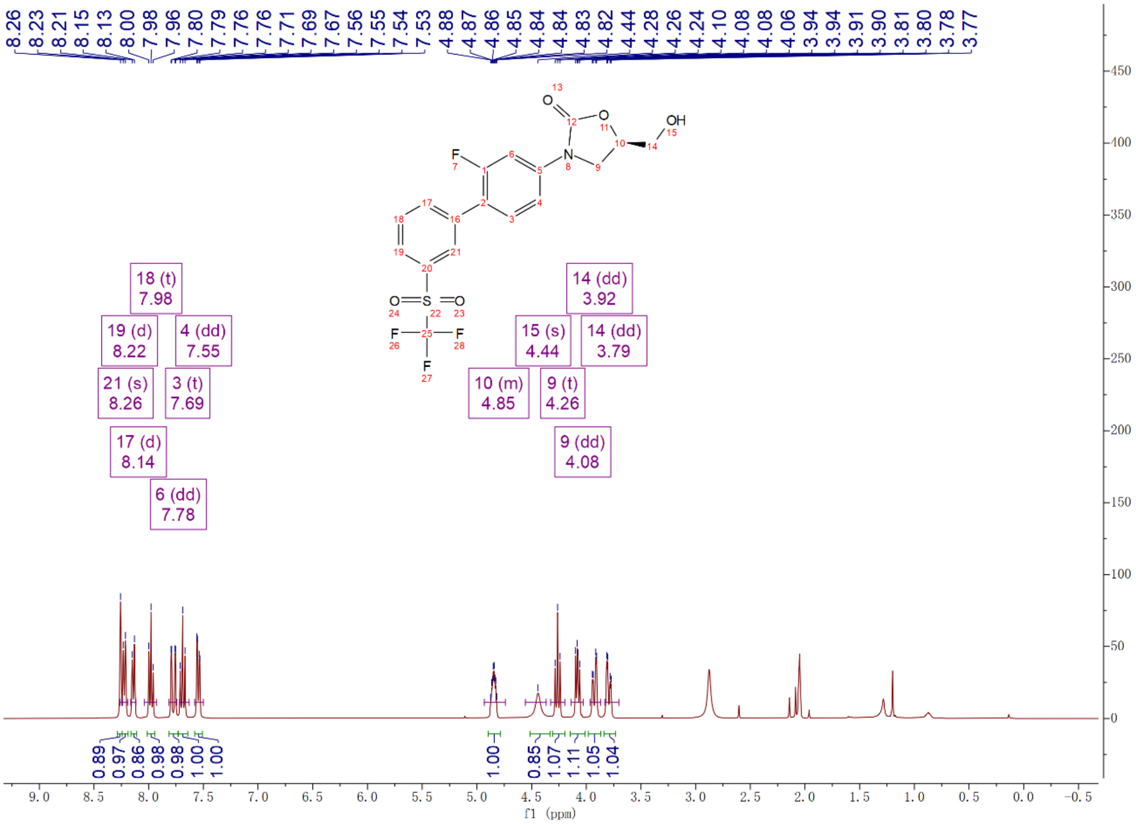

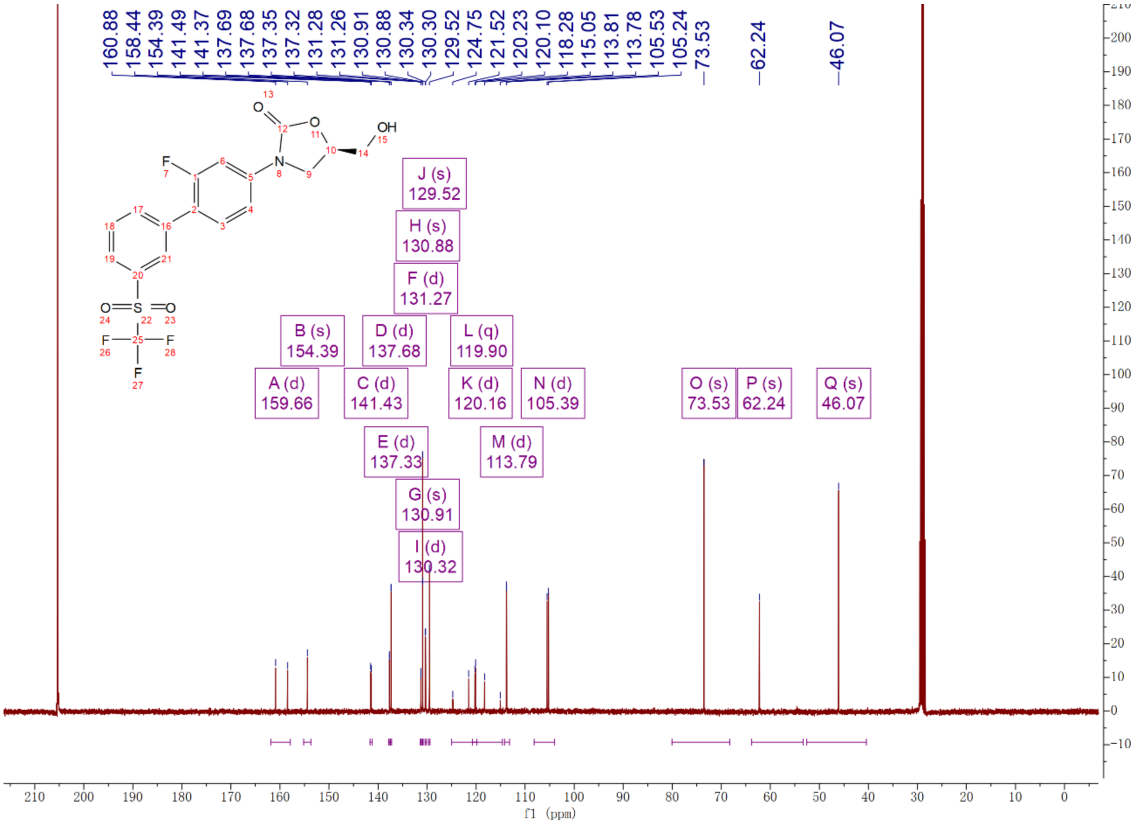

Compound **3i**: $^{1}$H and $^{13}$C NMR Spectrum

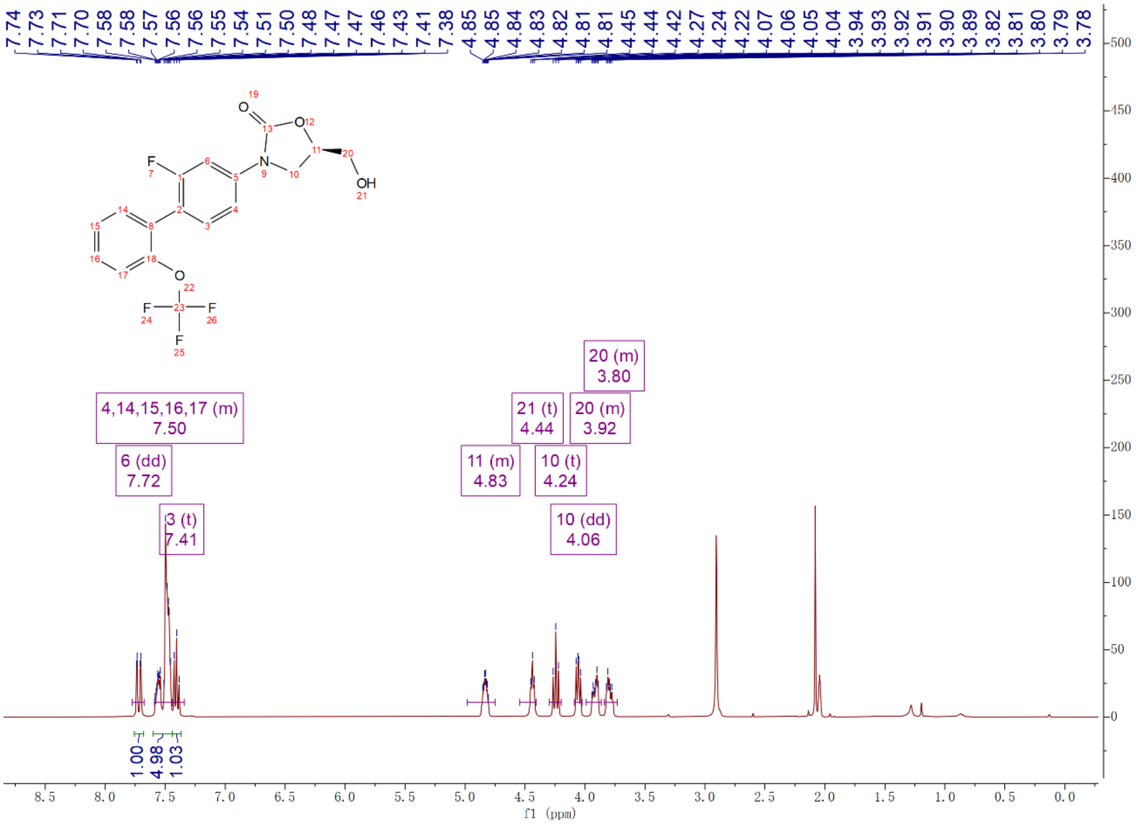

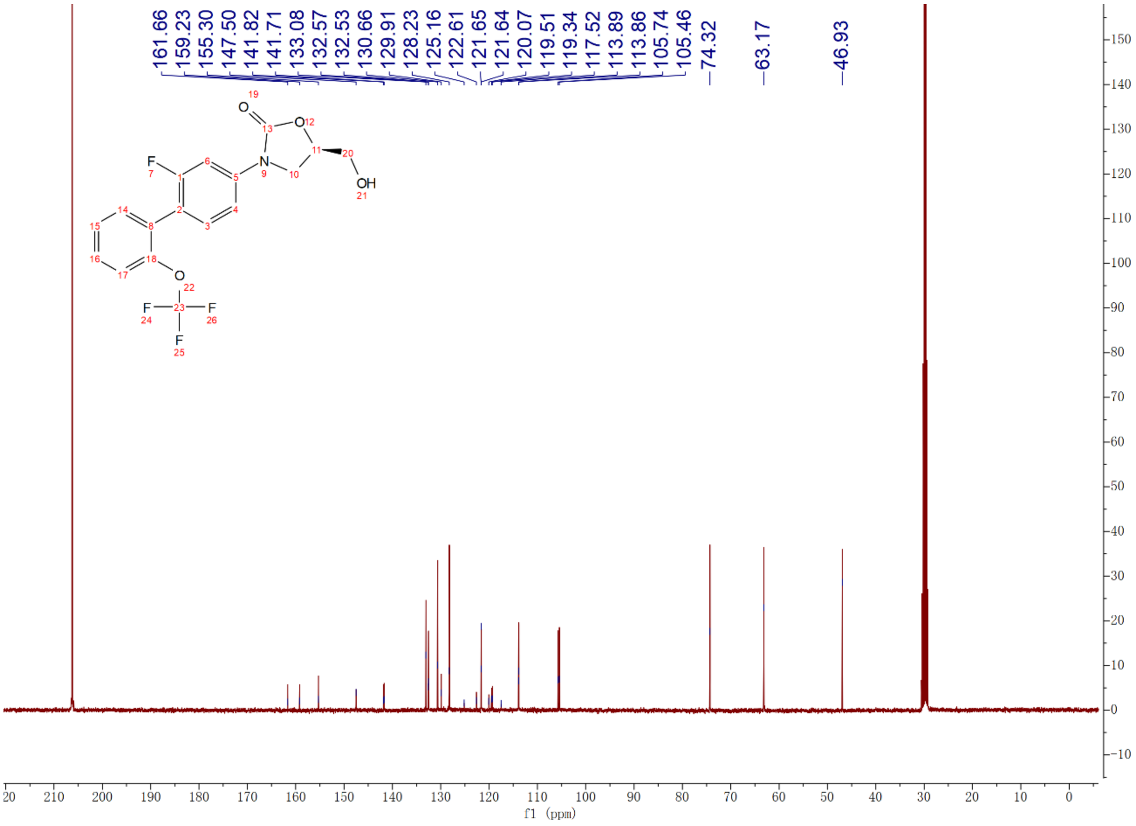

Compound **3j**: [1]H and [13]C NMR Spectrum

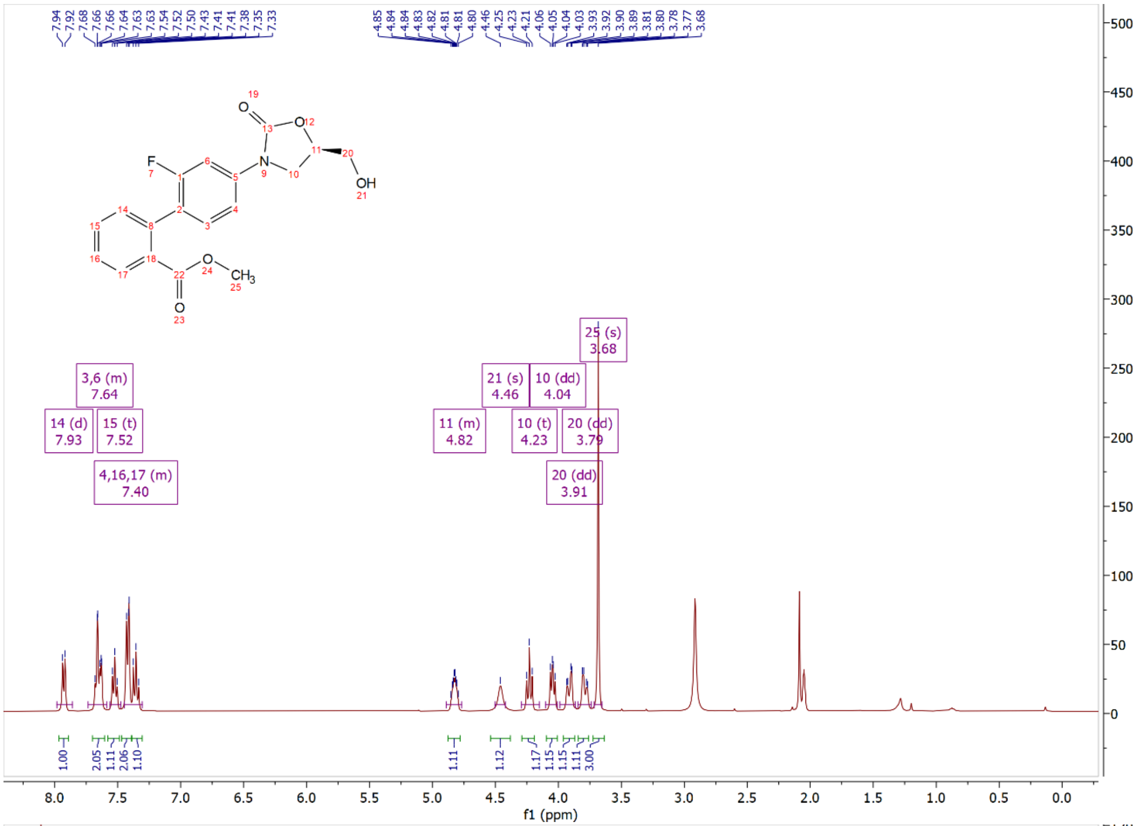

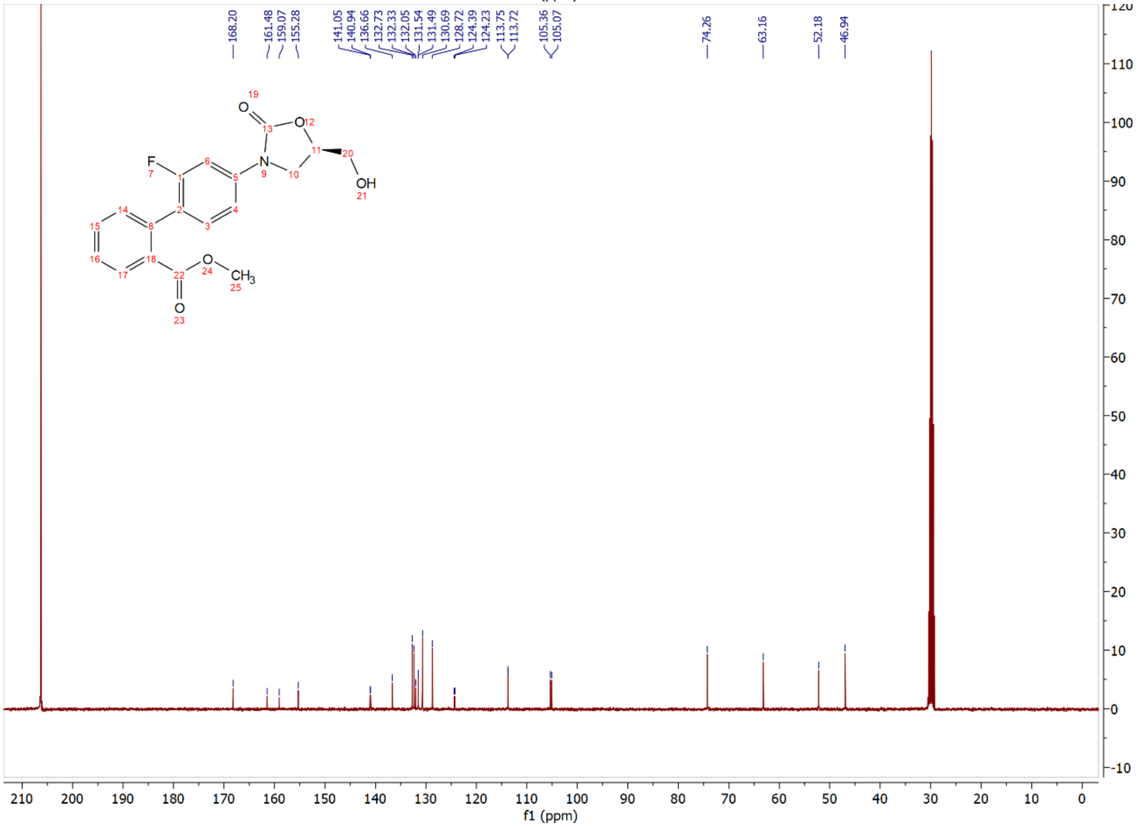

Compound **3k**: $^{1}$H and $^{13}$C NMR Spectrum

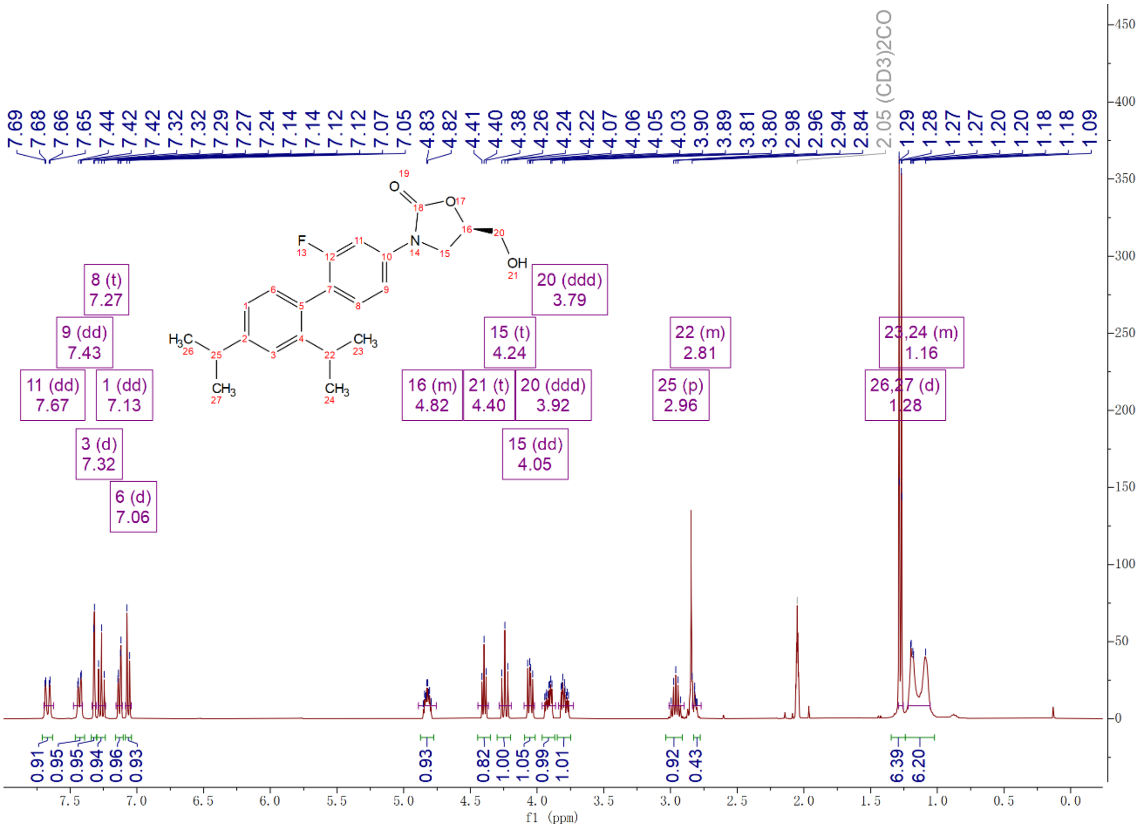

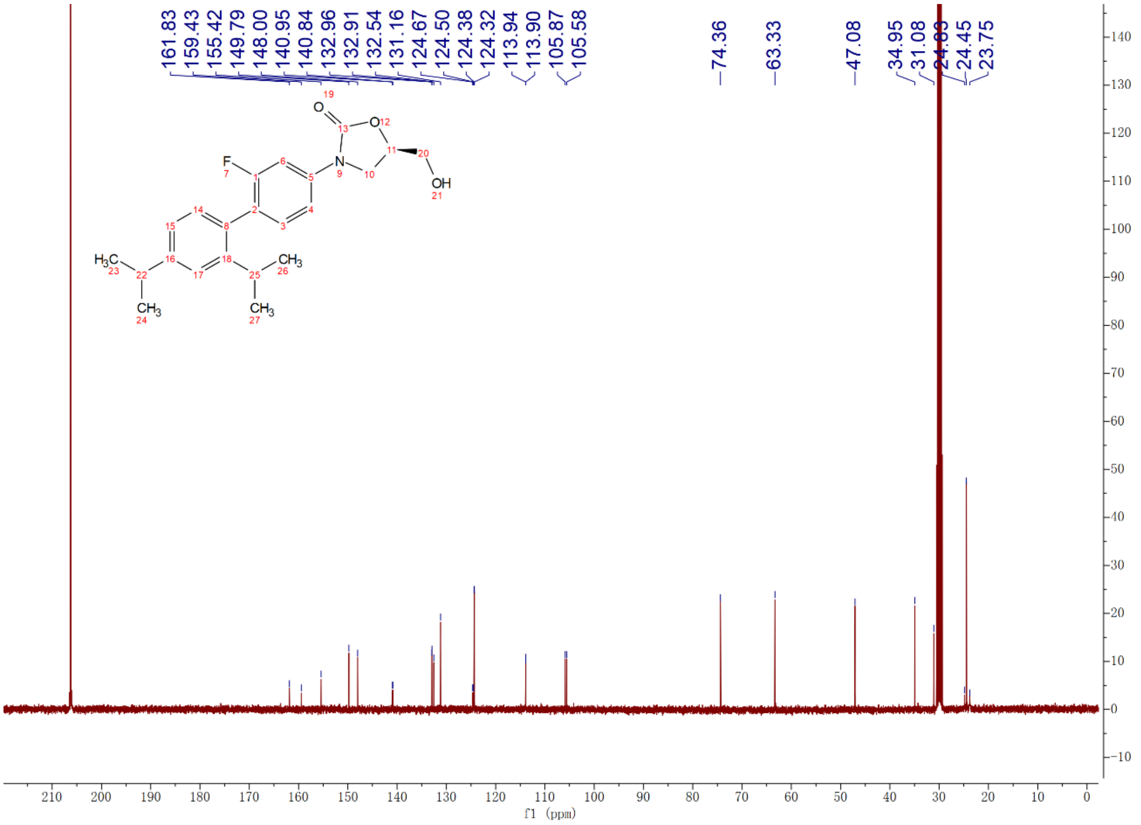

Compound **3l**: $^{1}$H and $^{13}$C NMR Spectrum

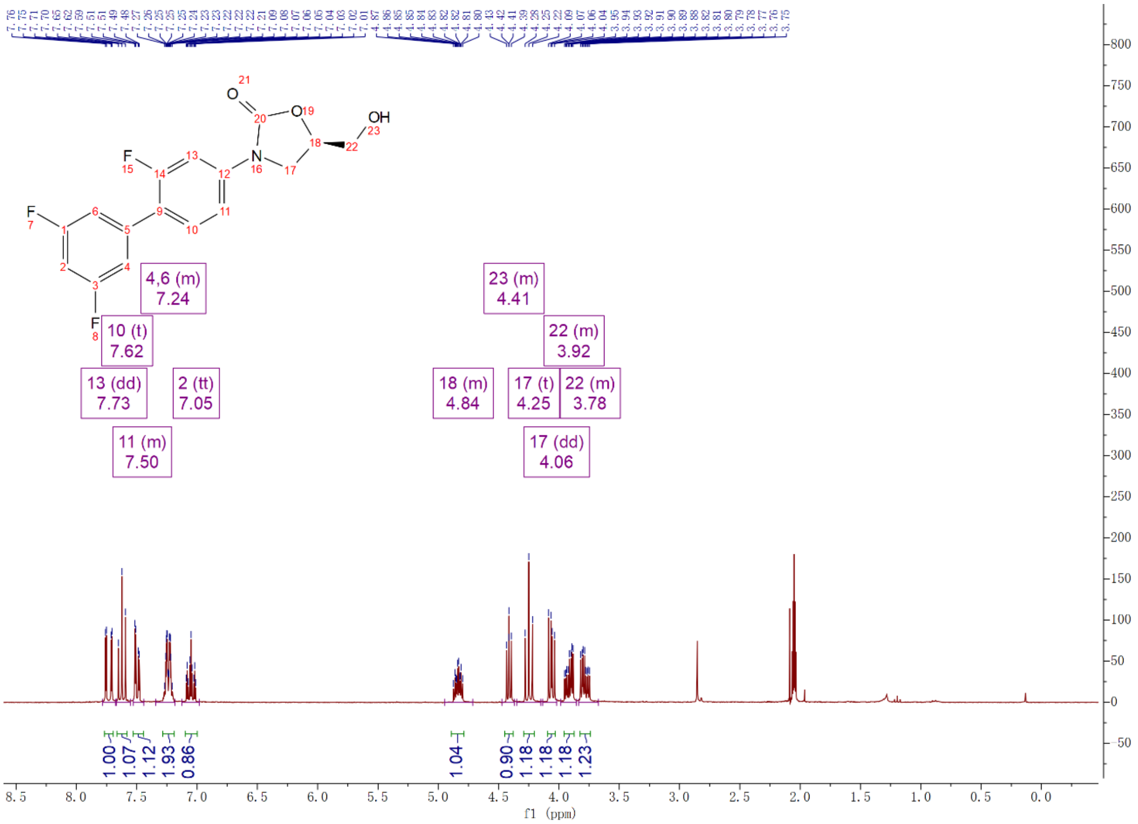


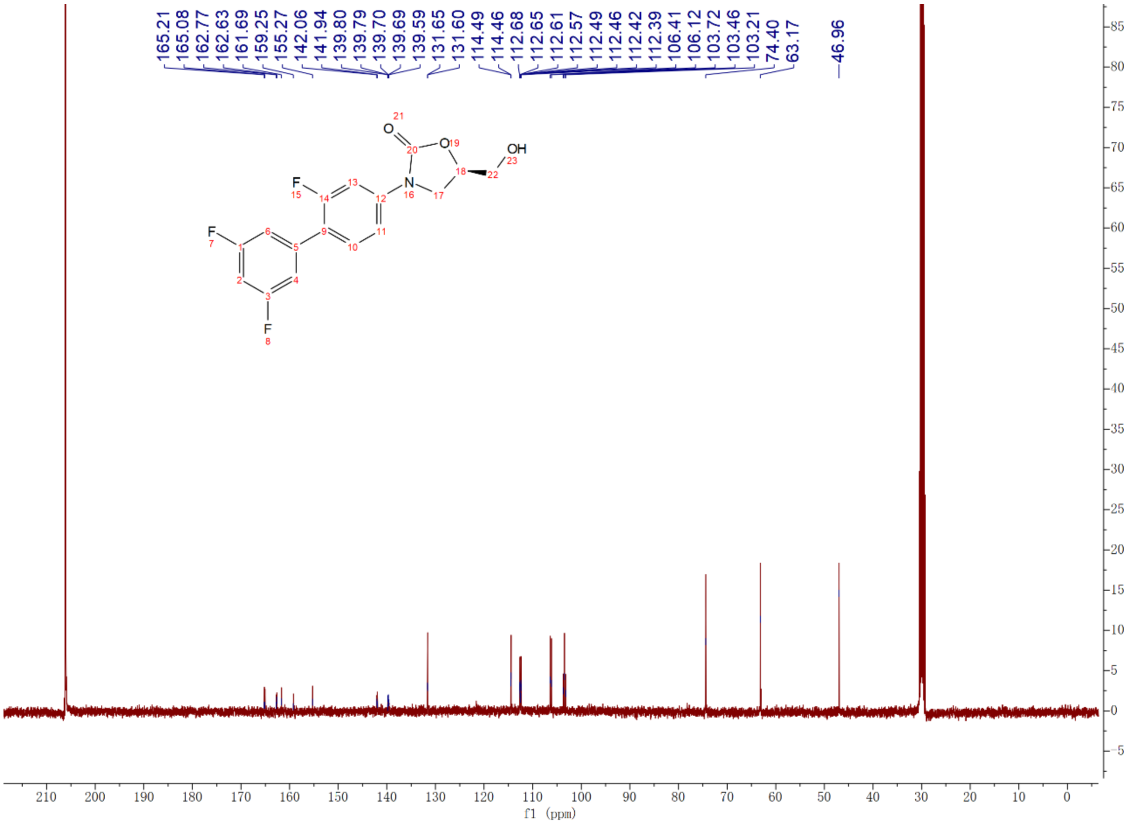

Compound **3m**: $^{1}$H and $^{13}$C NMR Spectrum

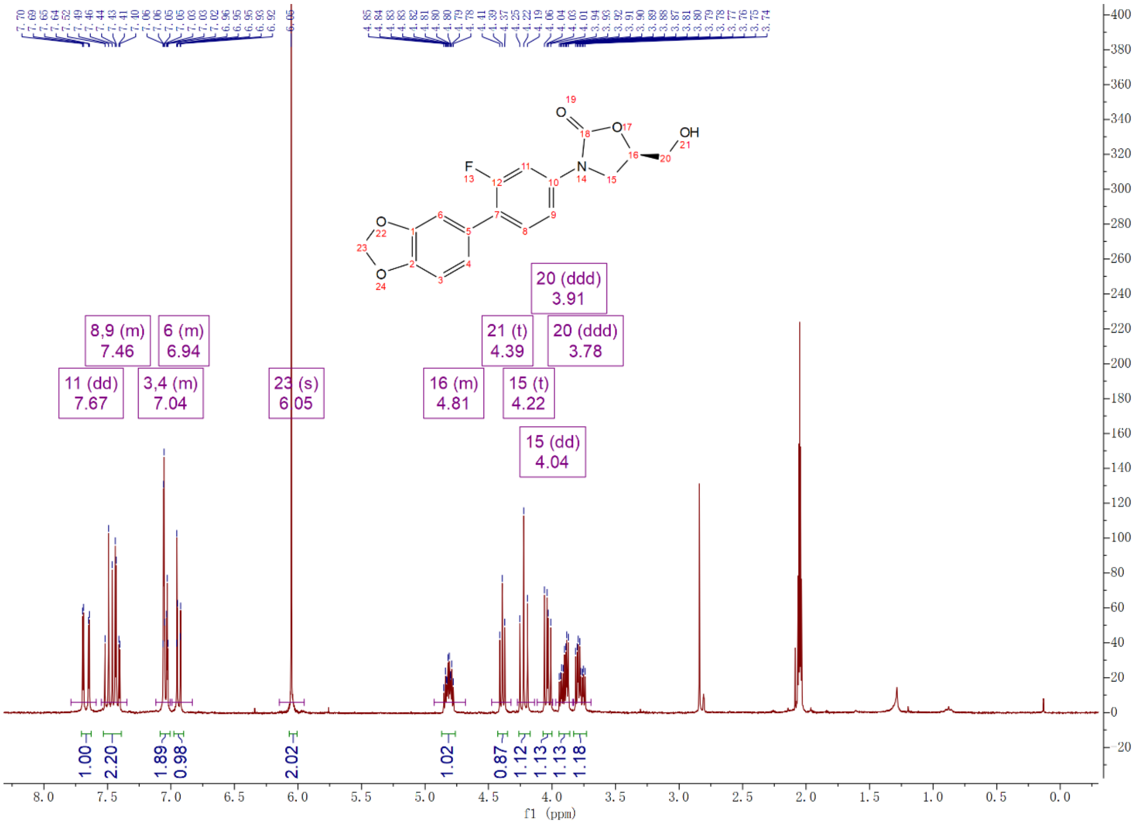

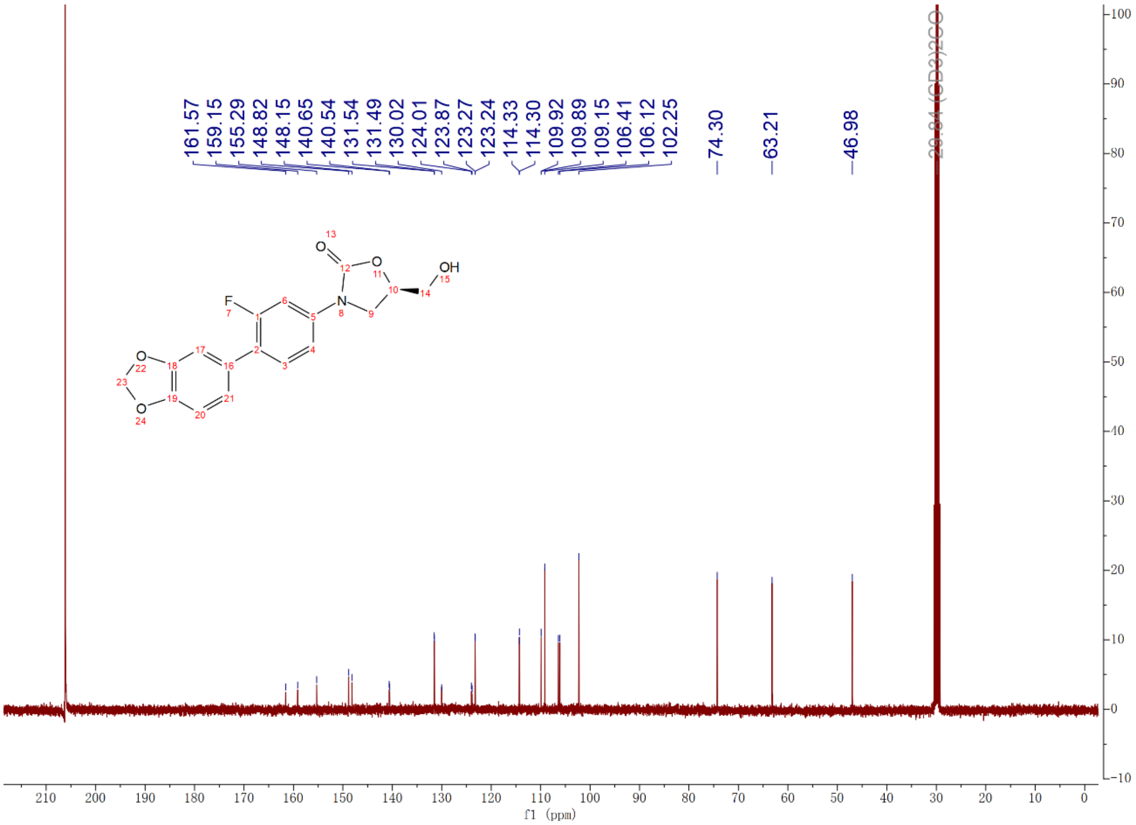

Compound **3n**: $^{1}$H and $^{13}$C NMR Spectrum

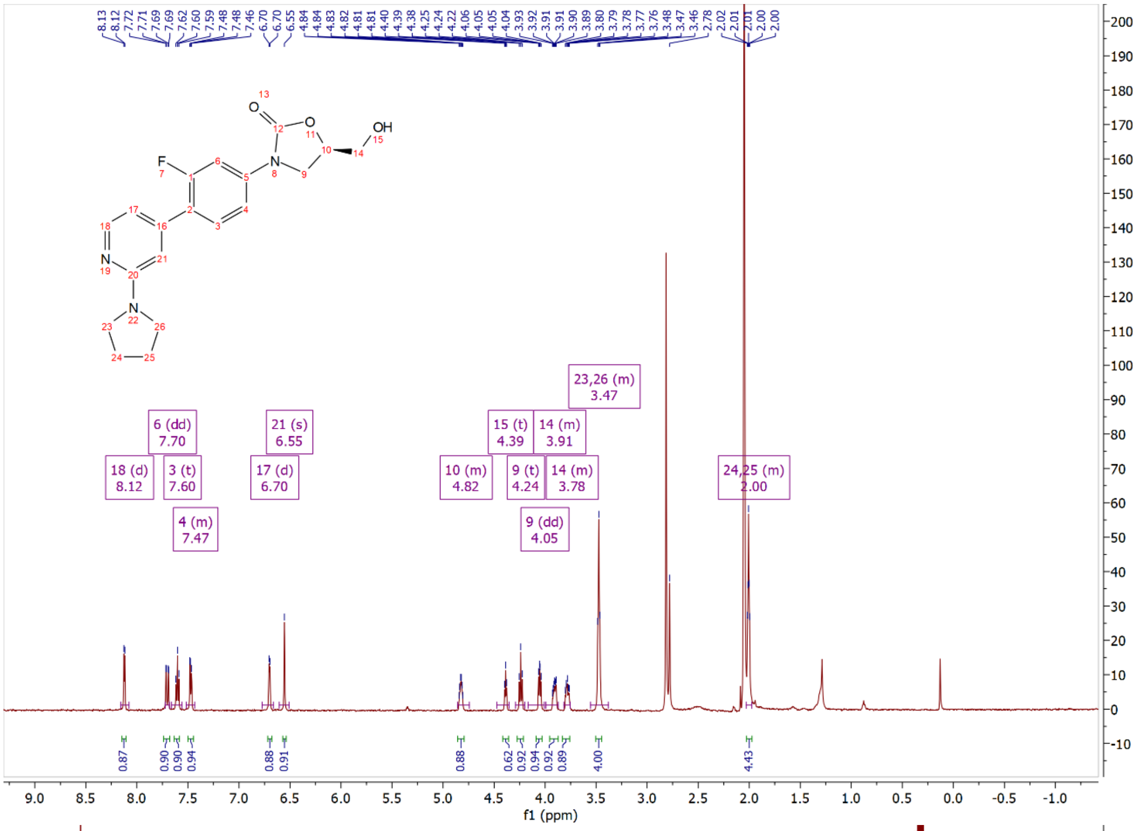


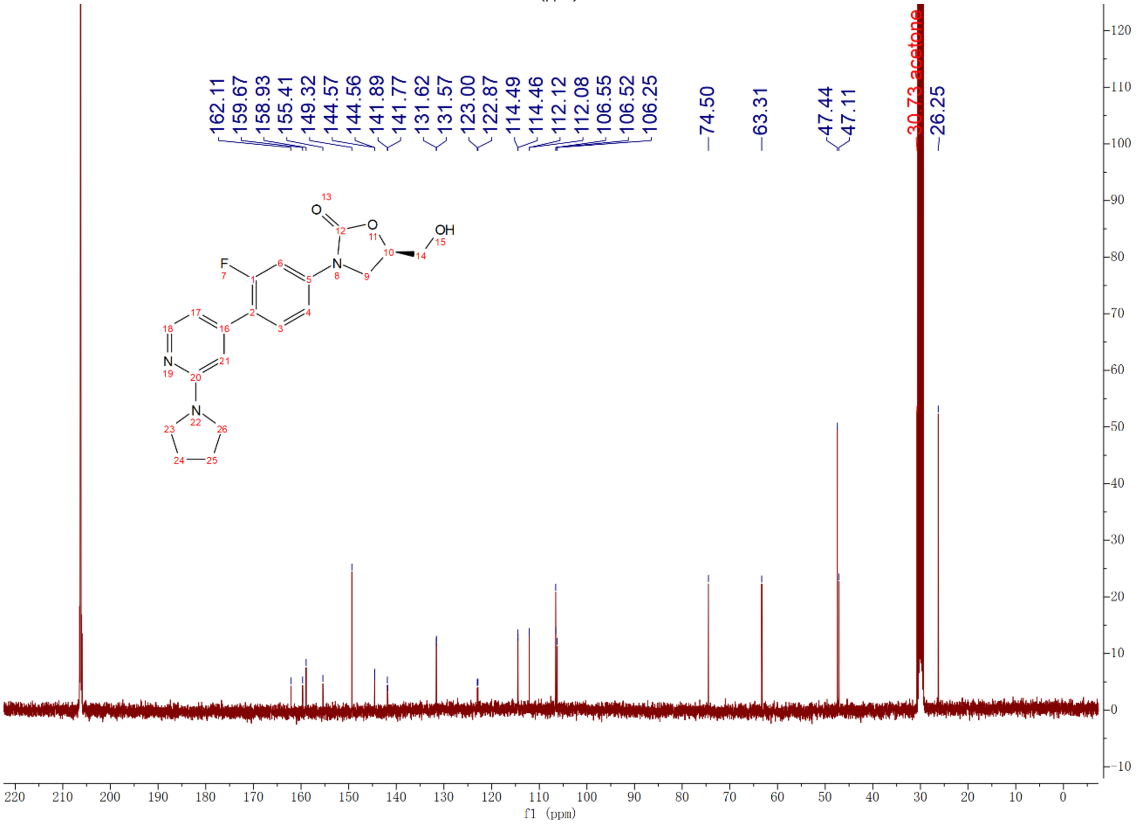

Compound **3o**: $^{1}$H and $^{13}$C NMR Spectrum

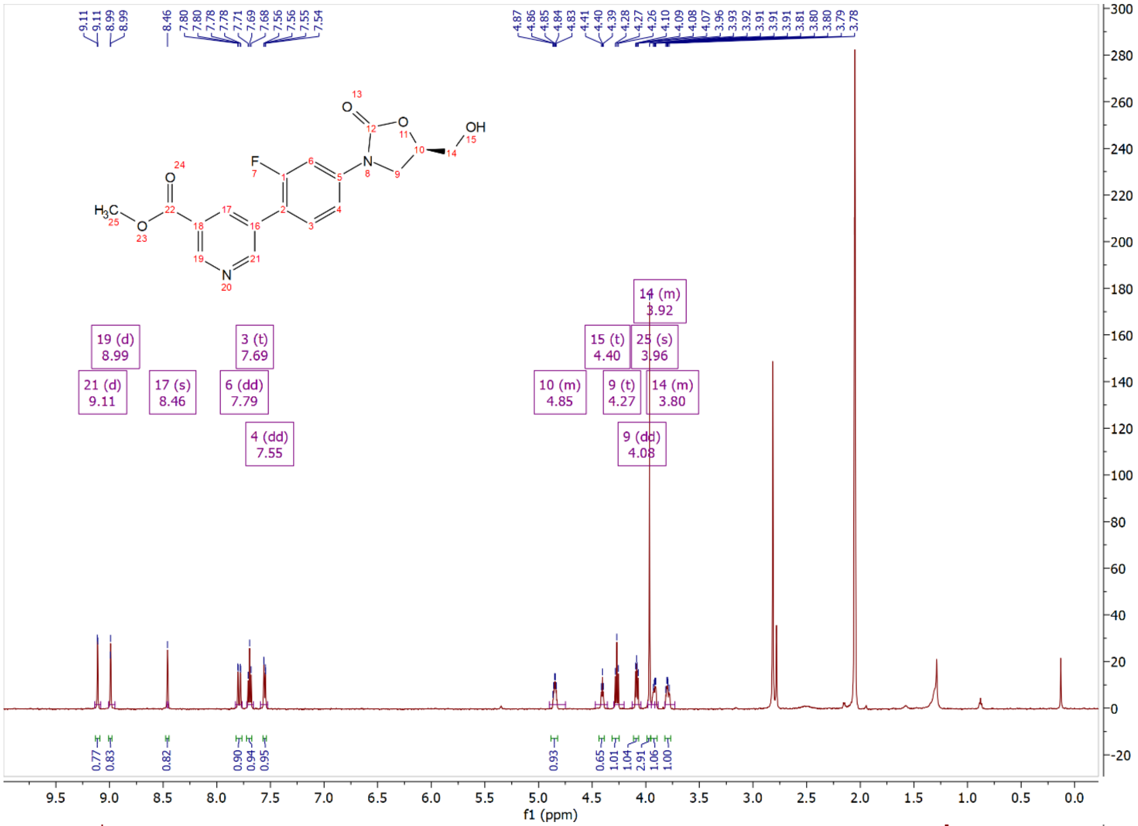

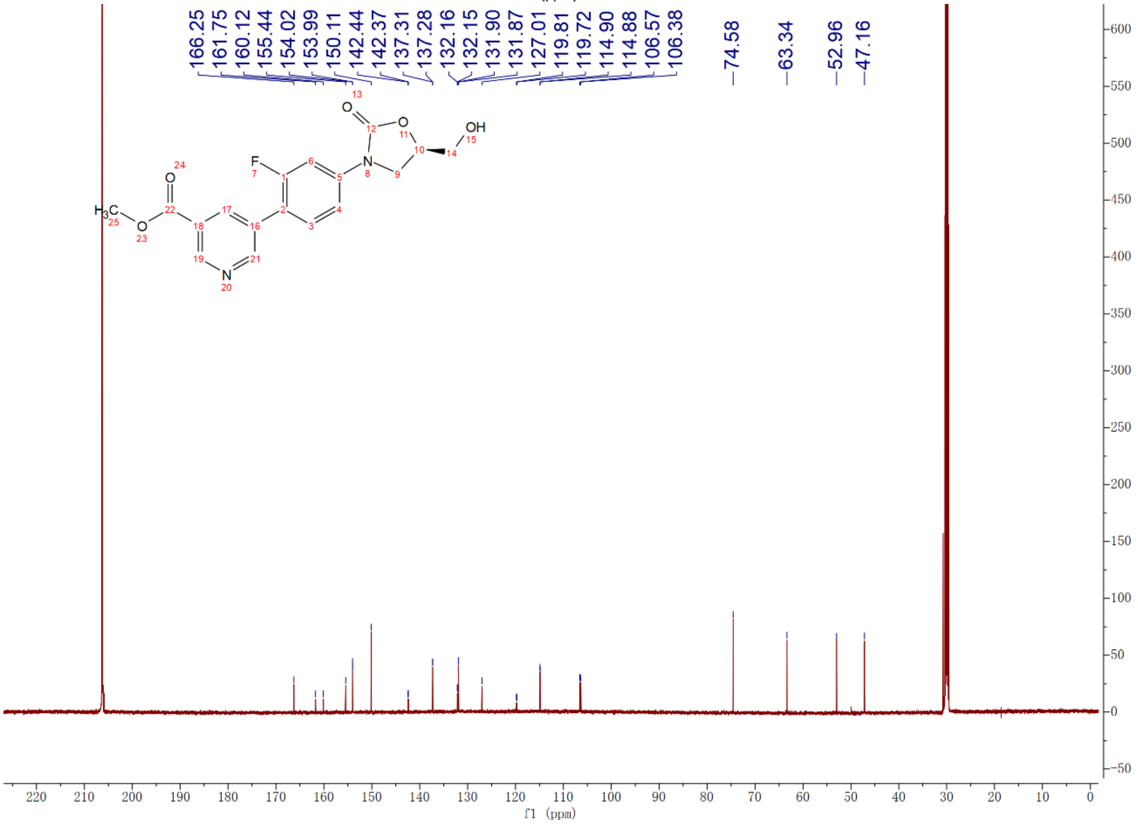

Compound **3oa**: $^{1}$H and $^{13}$C NMR Spectrum

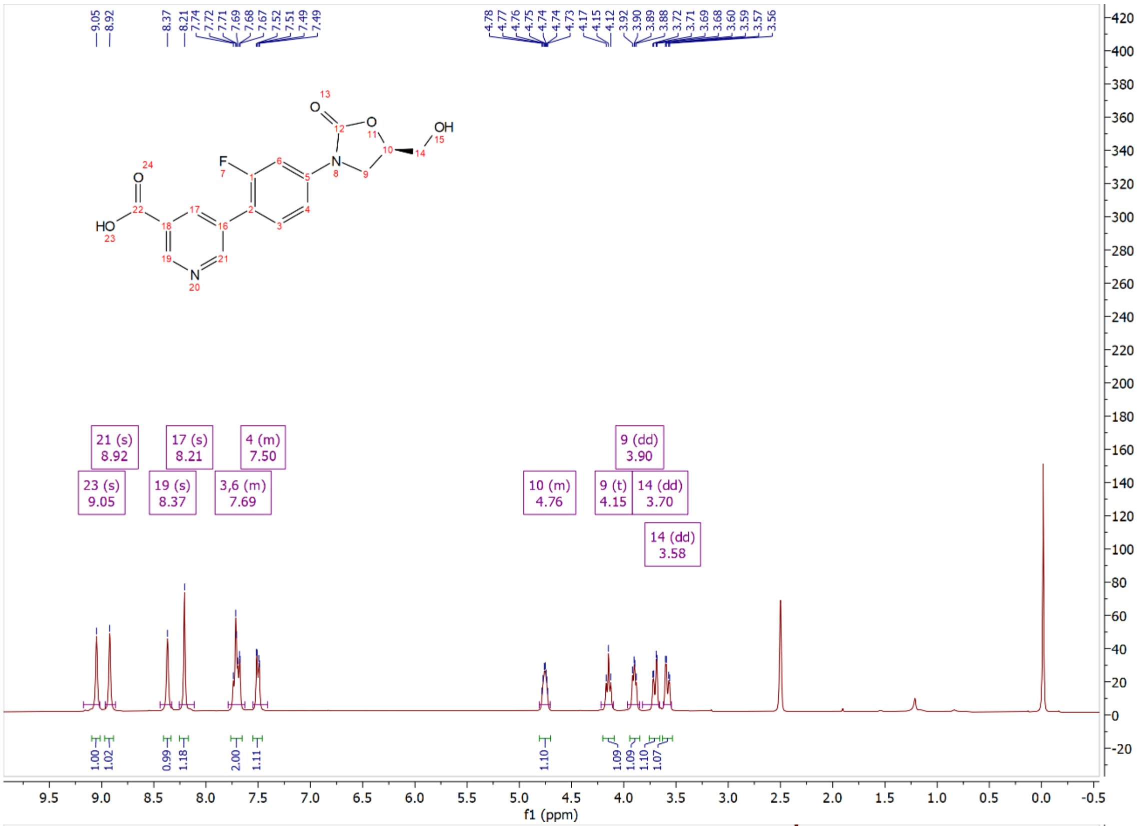

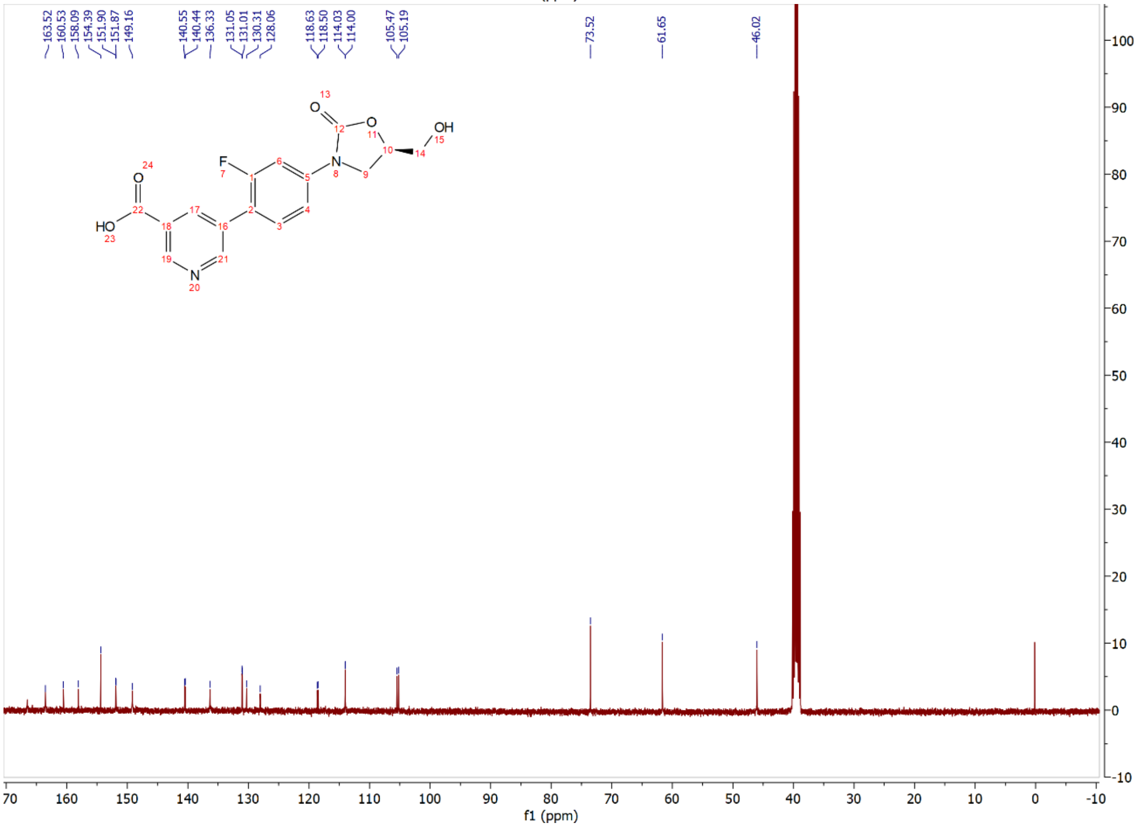

Compound **3p**: $^{1}$H and $^{13}$C NMR Spectrum

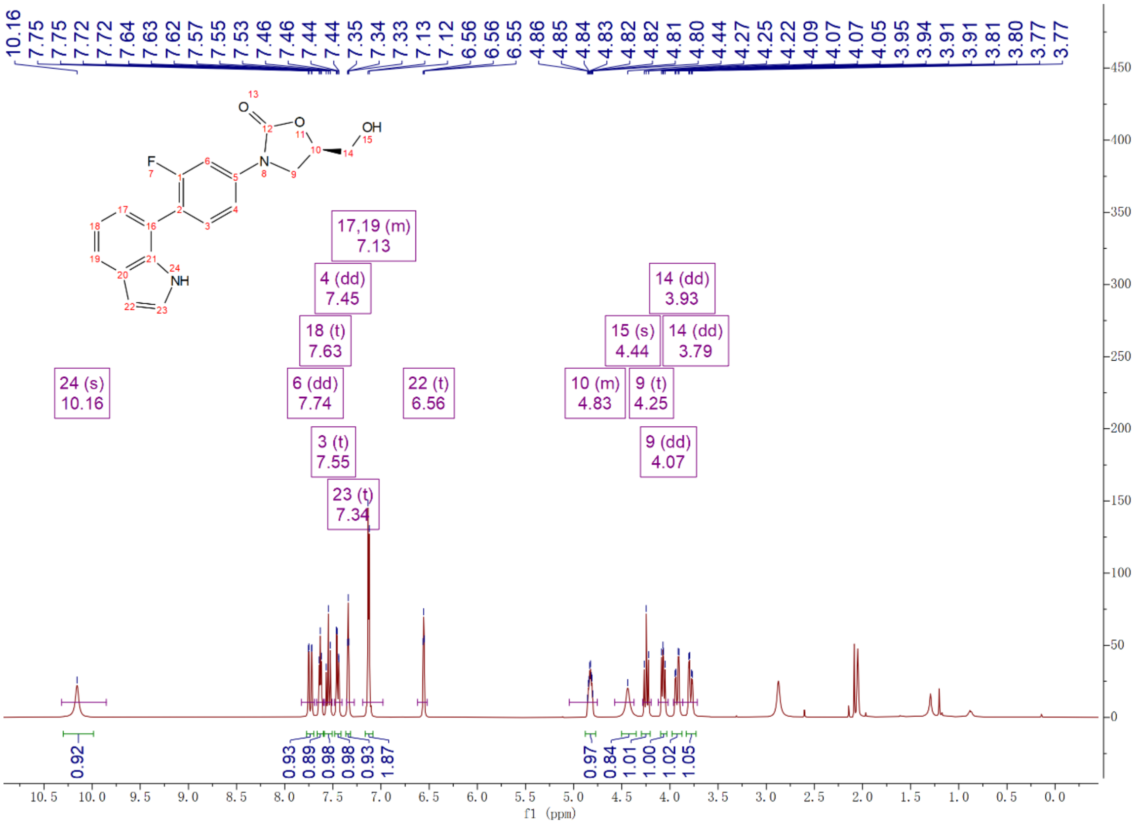


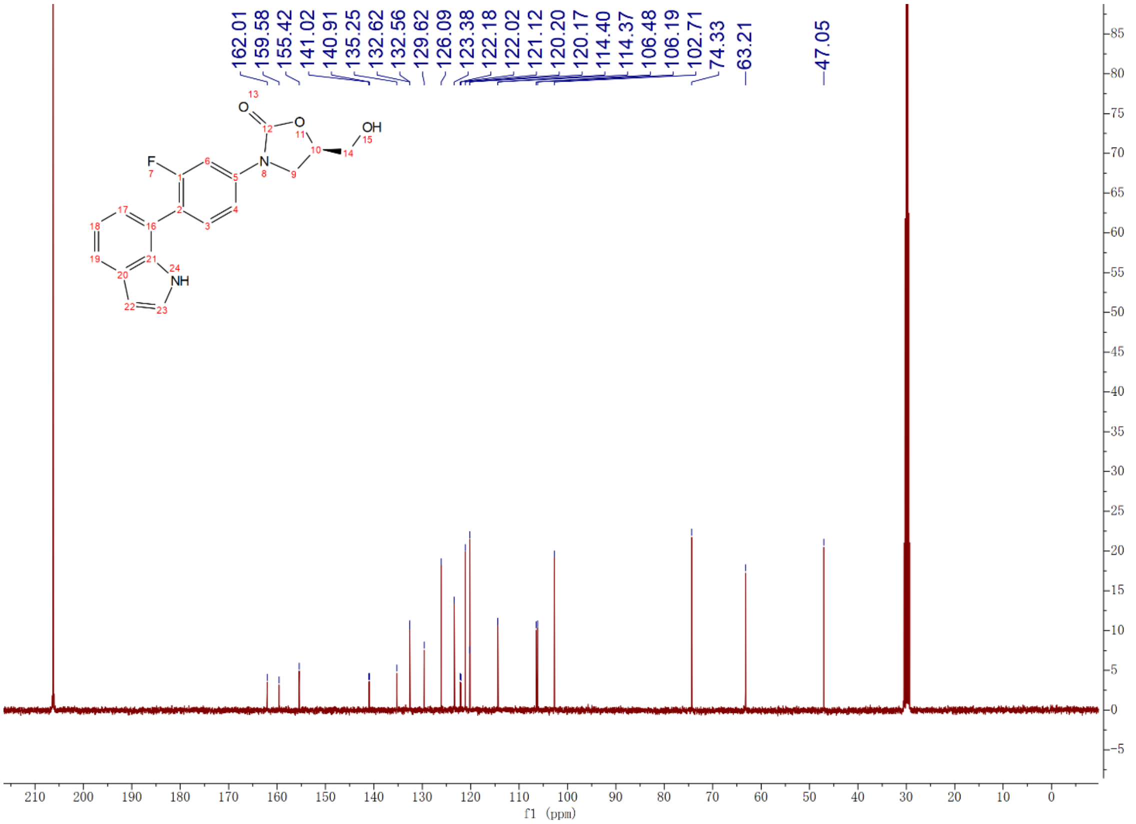

Compound **6a**: $^{1}$H and $^{13}$C NMR Spectrum

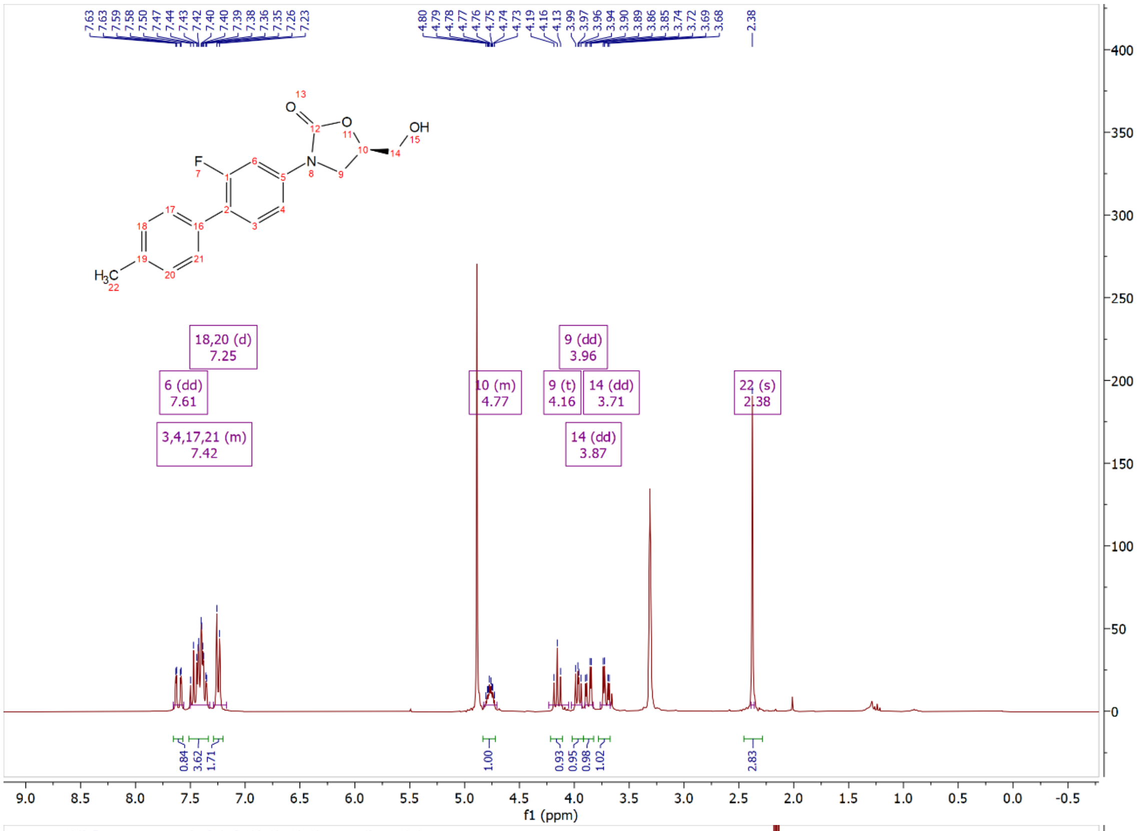

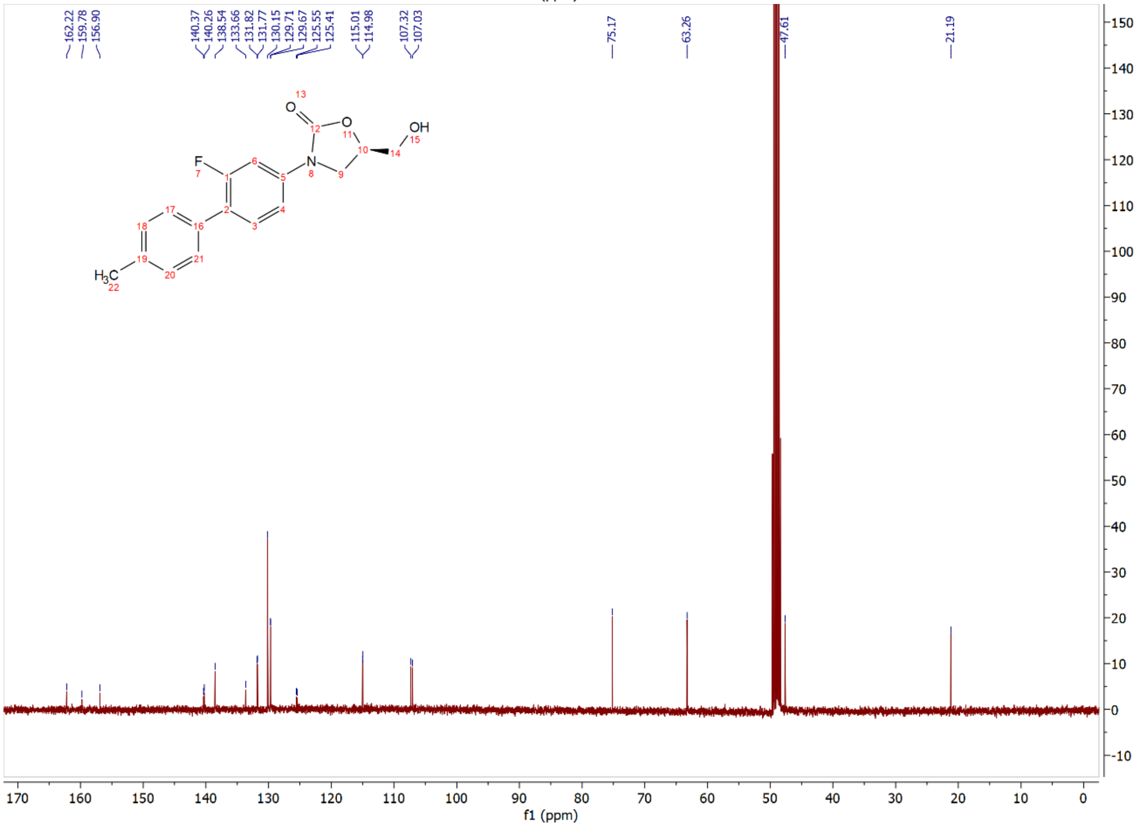

Compound **6b**: $^{1}$H and $^{13}$C NMR Spectrum

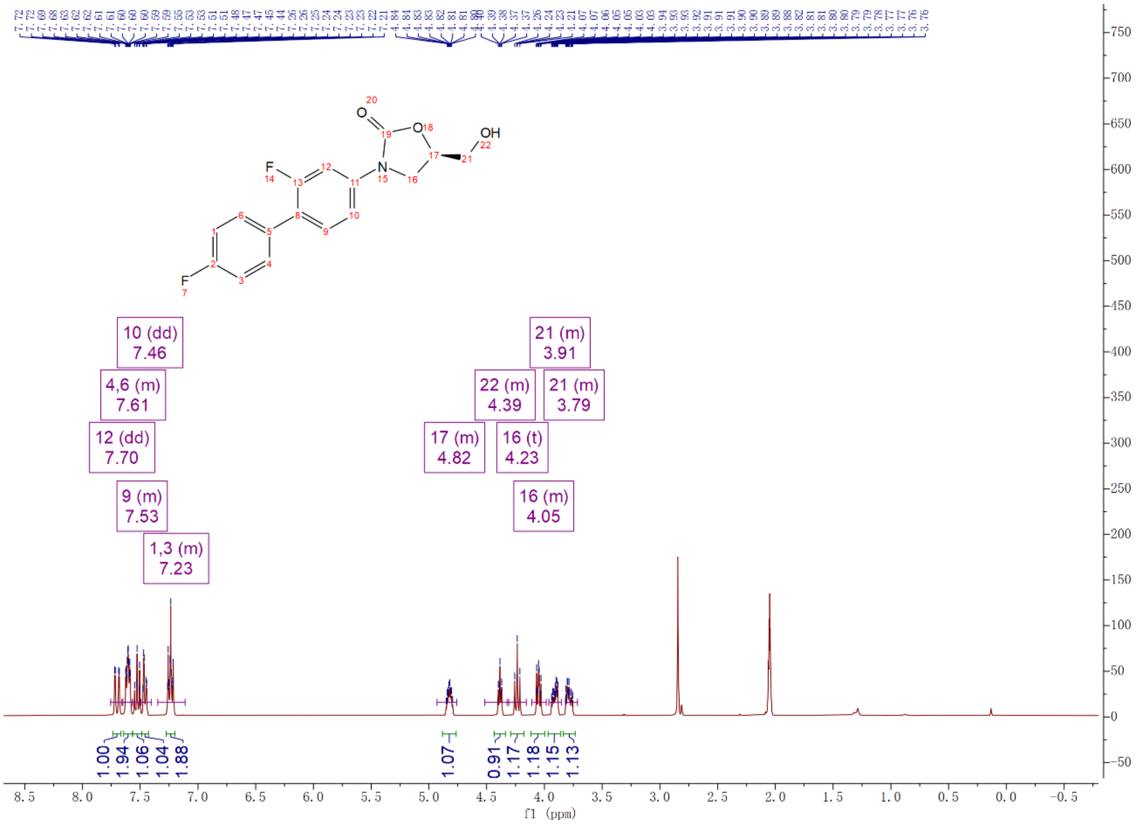

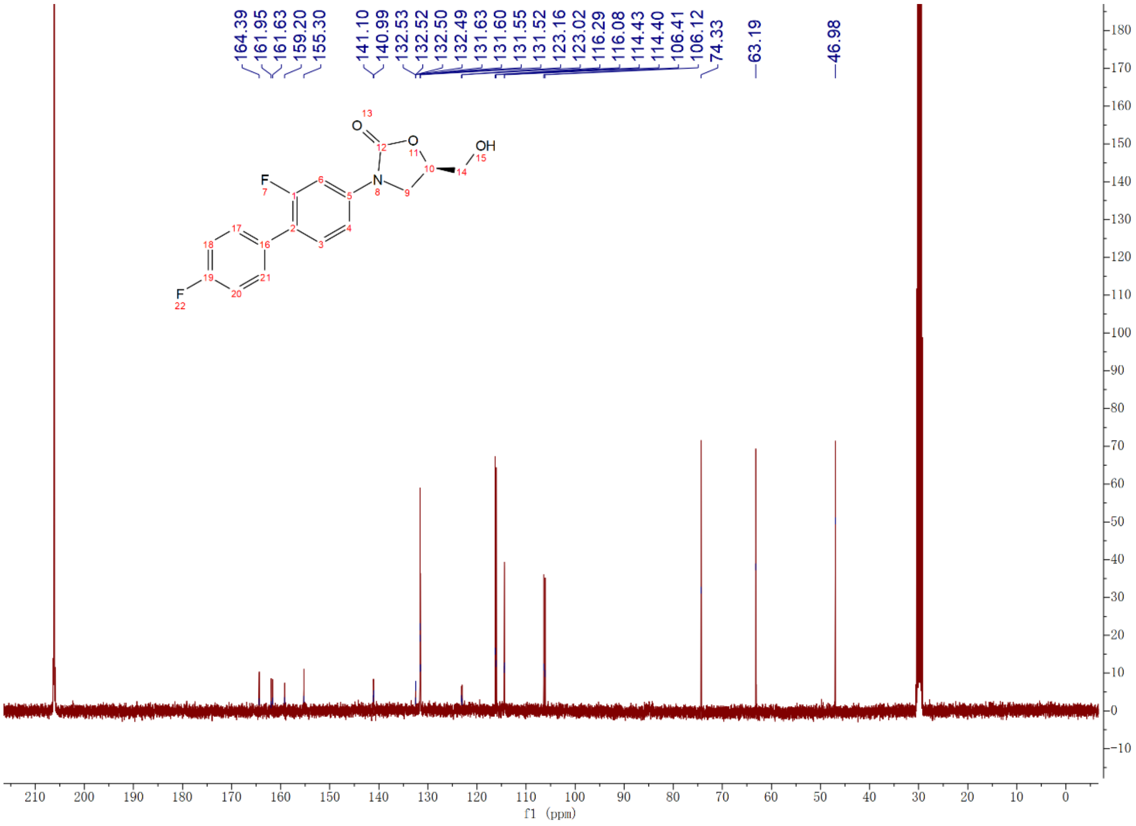

Compound **6c**: $^{1}$H and $^{13}$C NMR Spectrum

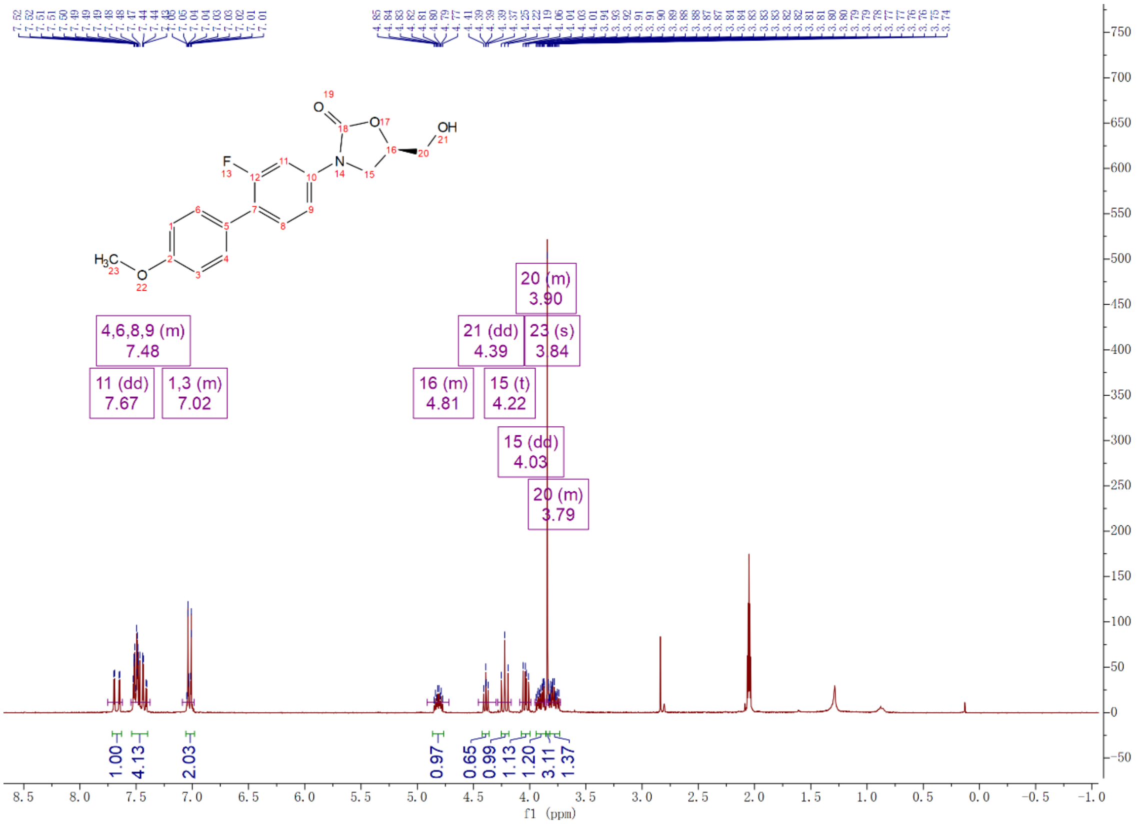

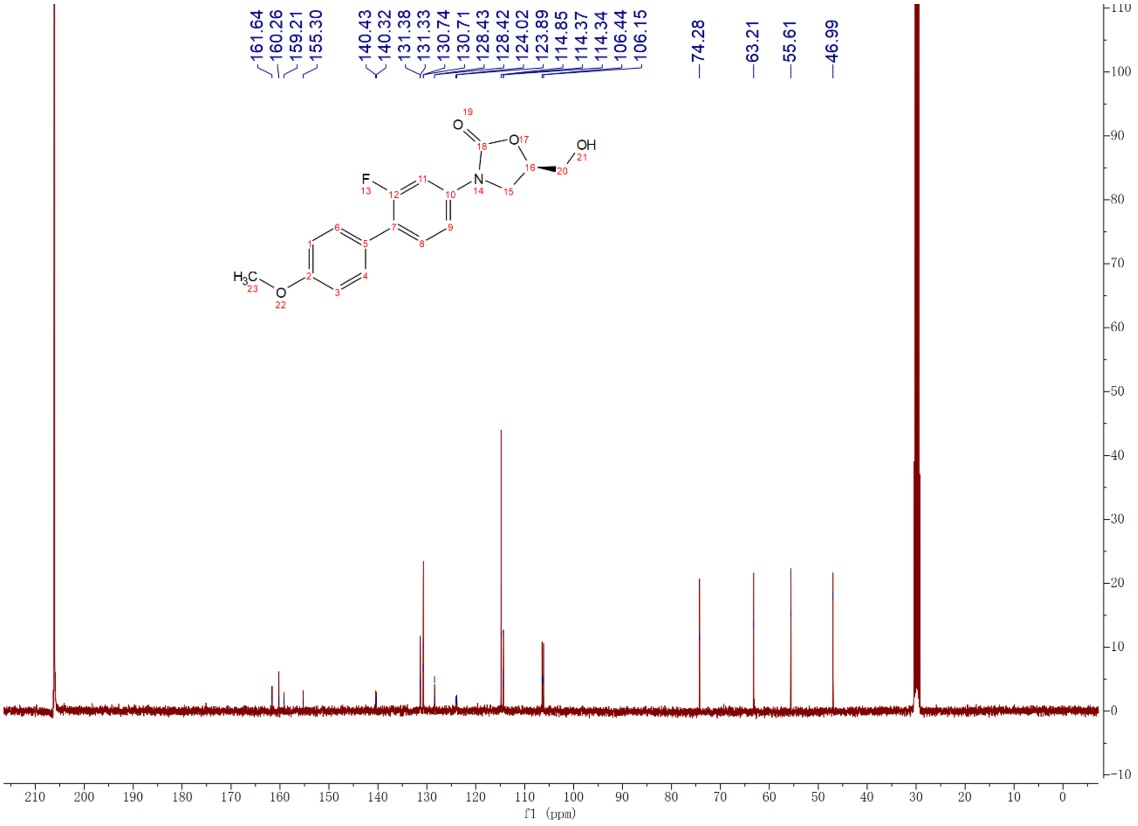

Compound **12a**: $^{1}$H and $^{13}$C NMR Spectrum

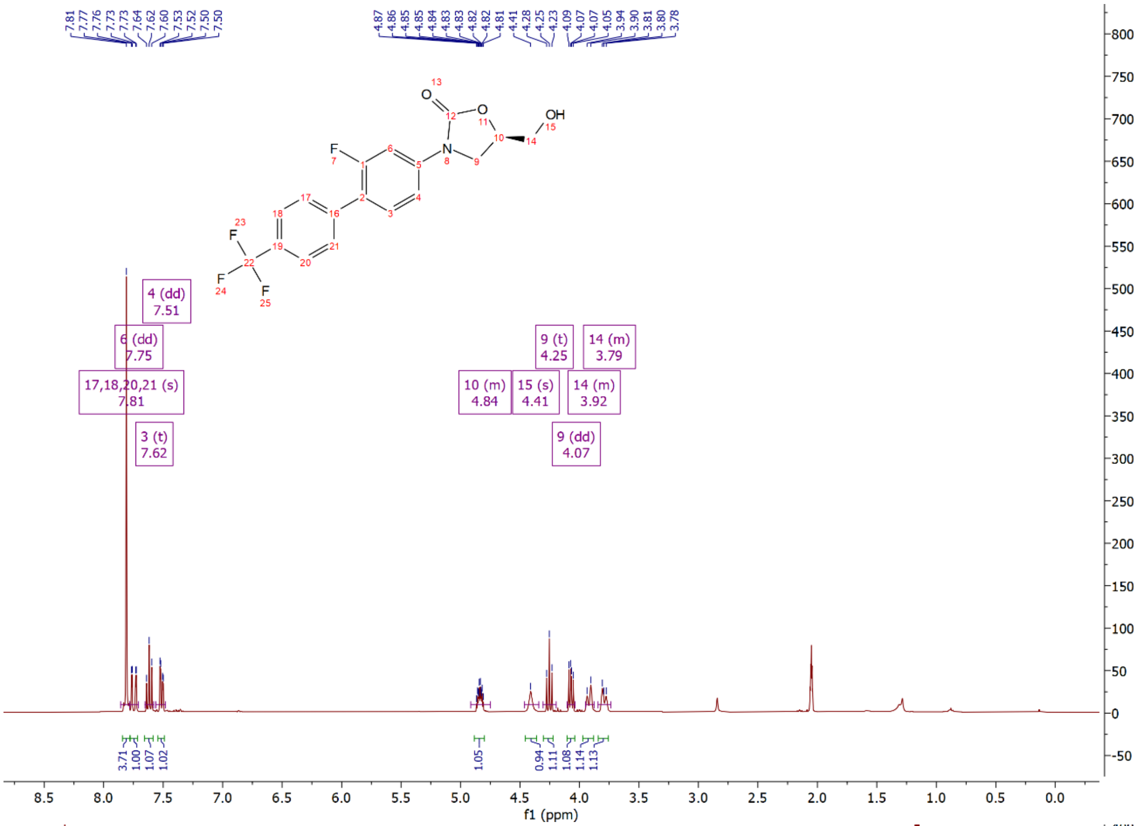


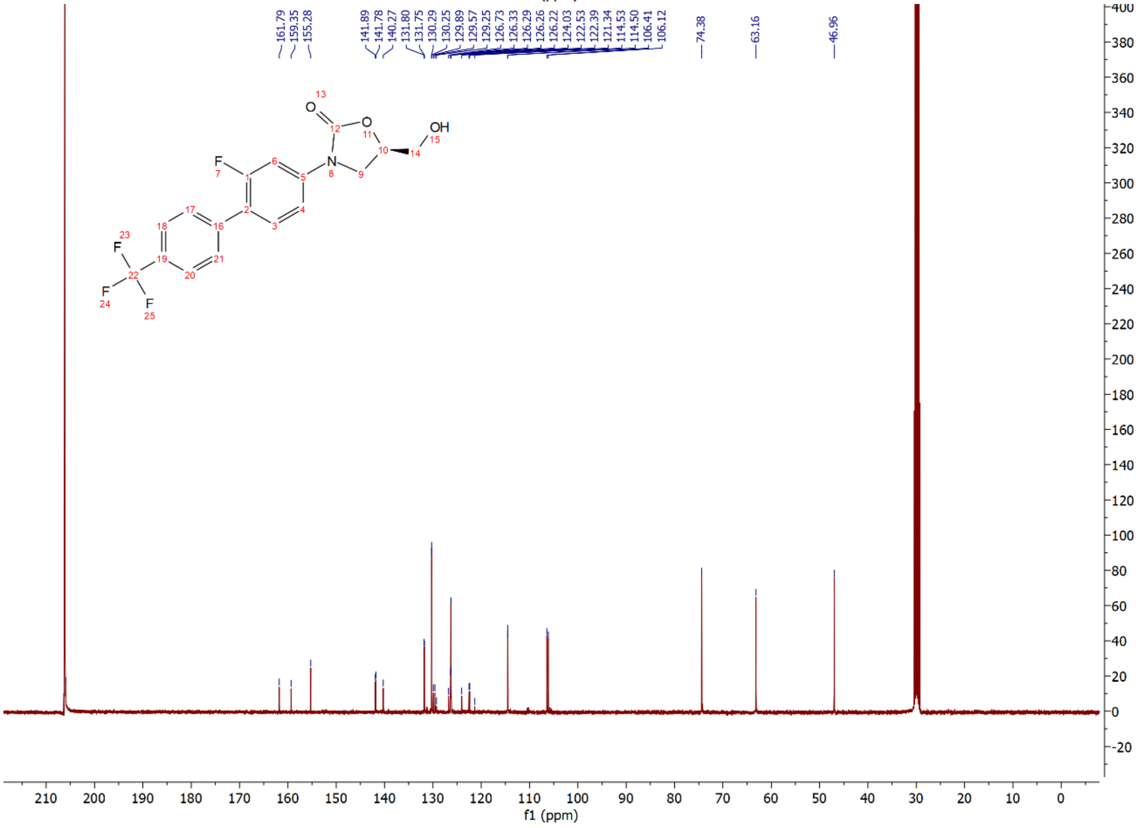

Compound **12b**: $^{1}$H and $^{13}$C NMR Spectrum

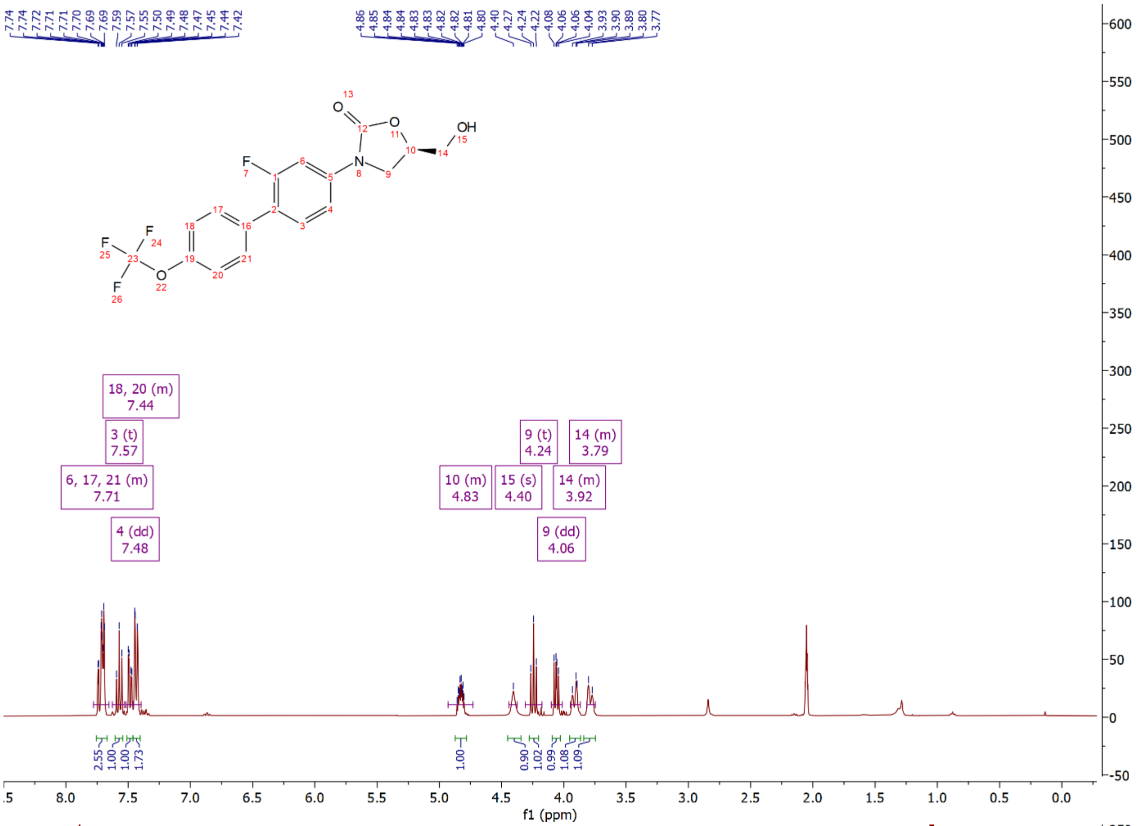

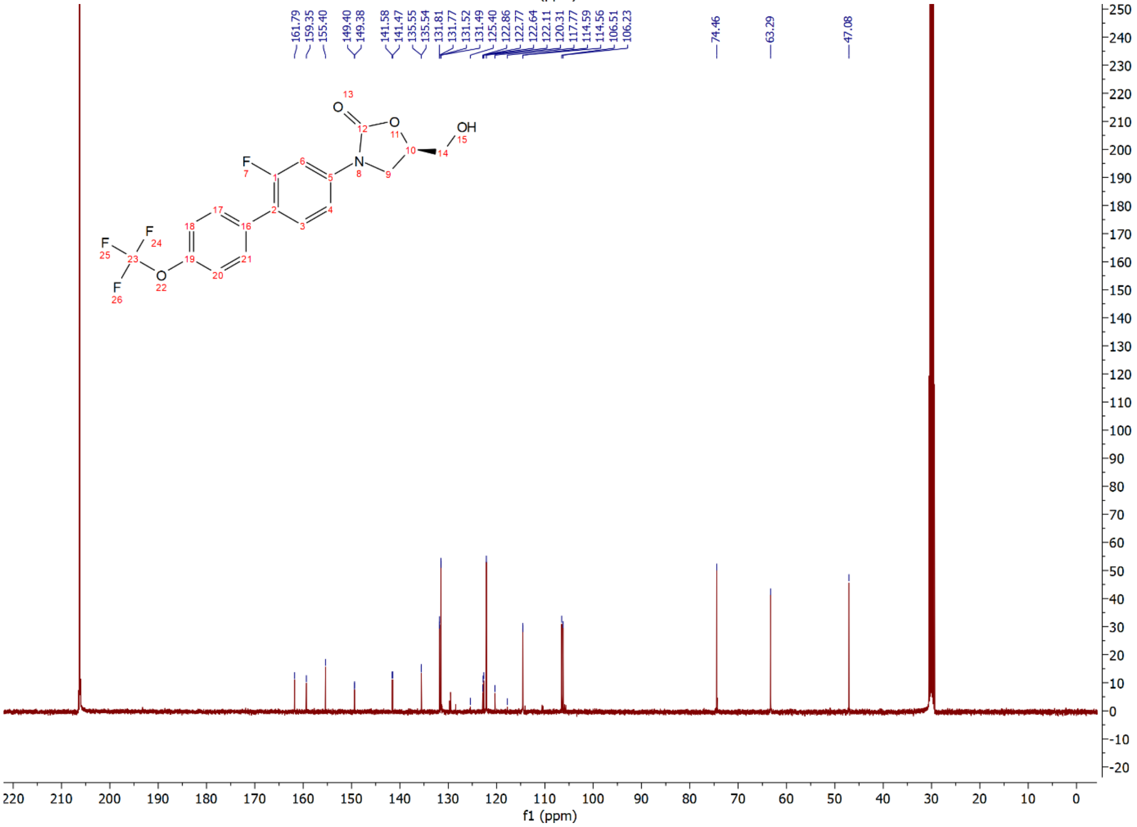

Compound **12c**: $^{1}H$ and $^{13}C$ NMR Spectrum

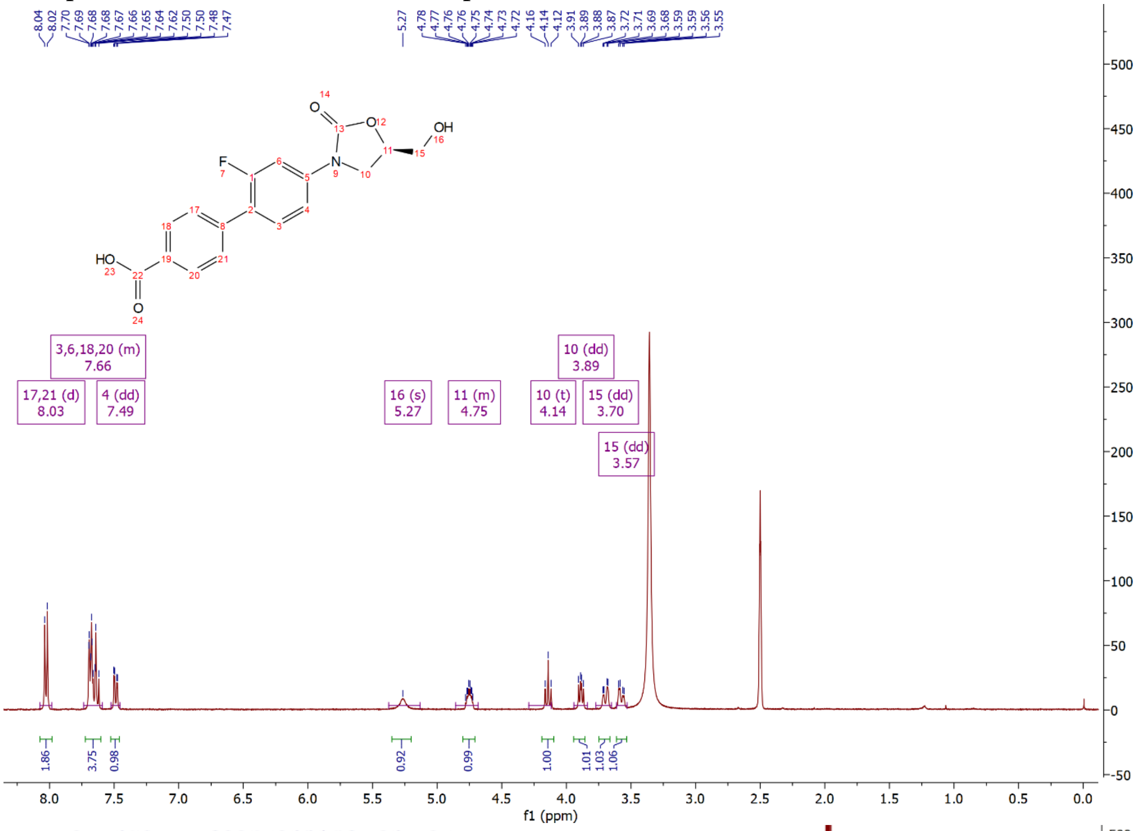


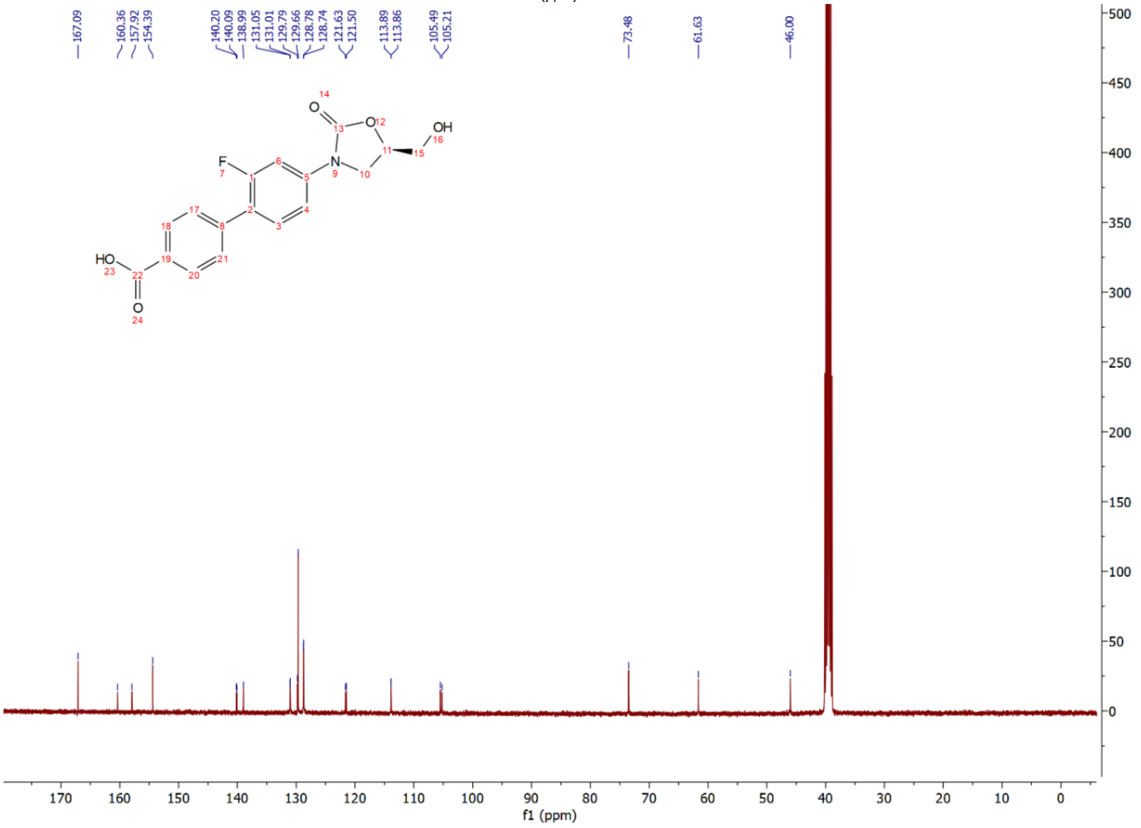

Compound **12d**: $^{1}$H and $^{13}$C NMR Spectrum

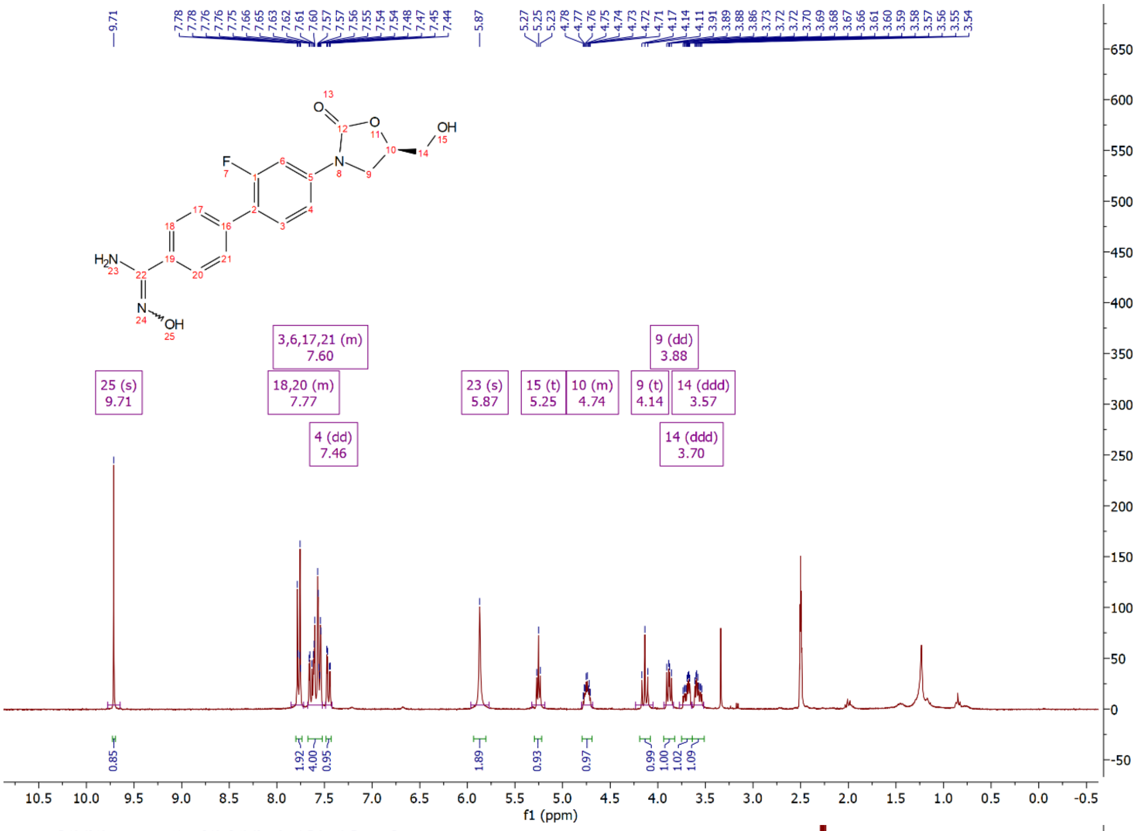

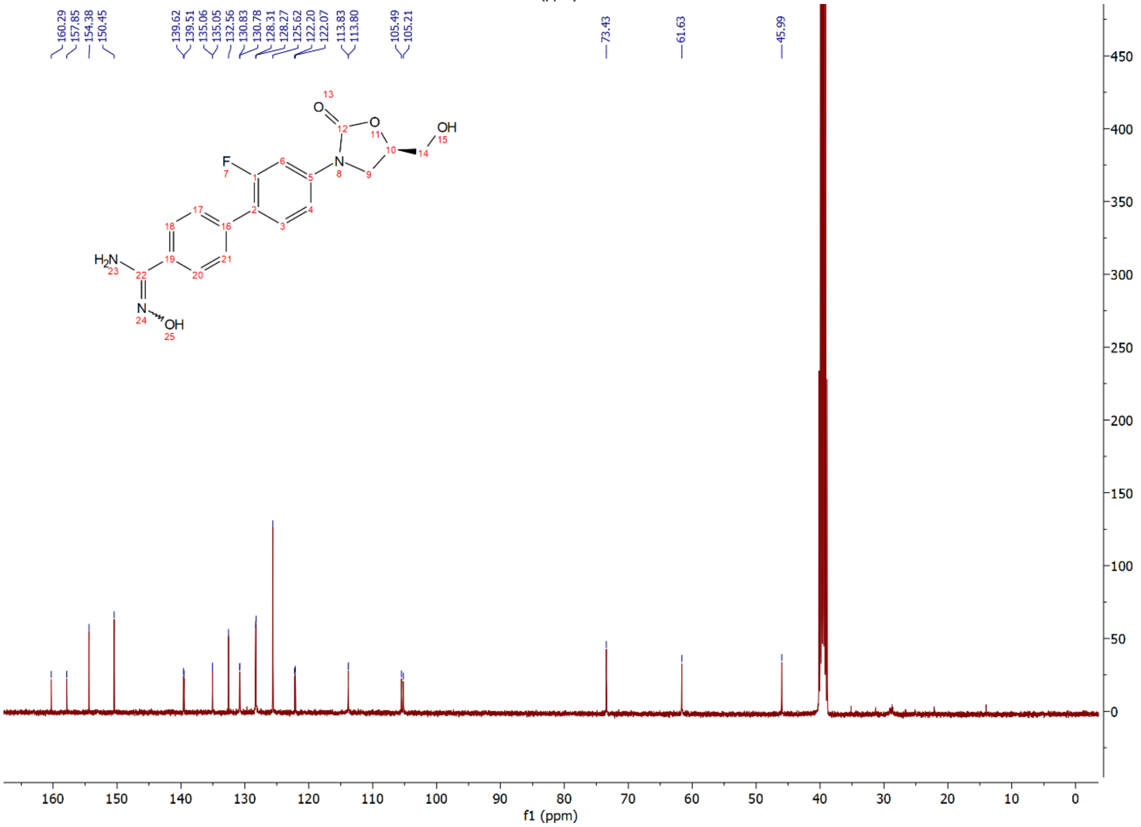

Compound **12e**: $^{1}$H and $^{13}$C NMR Spectrum

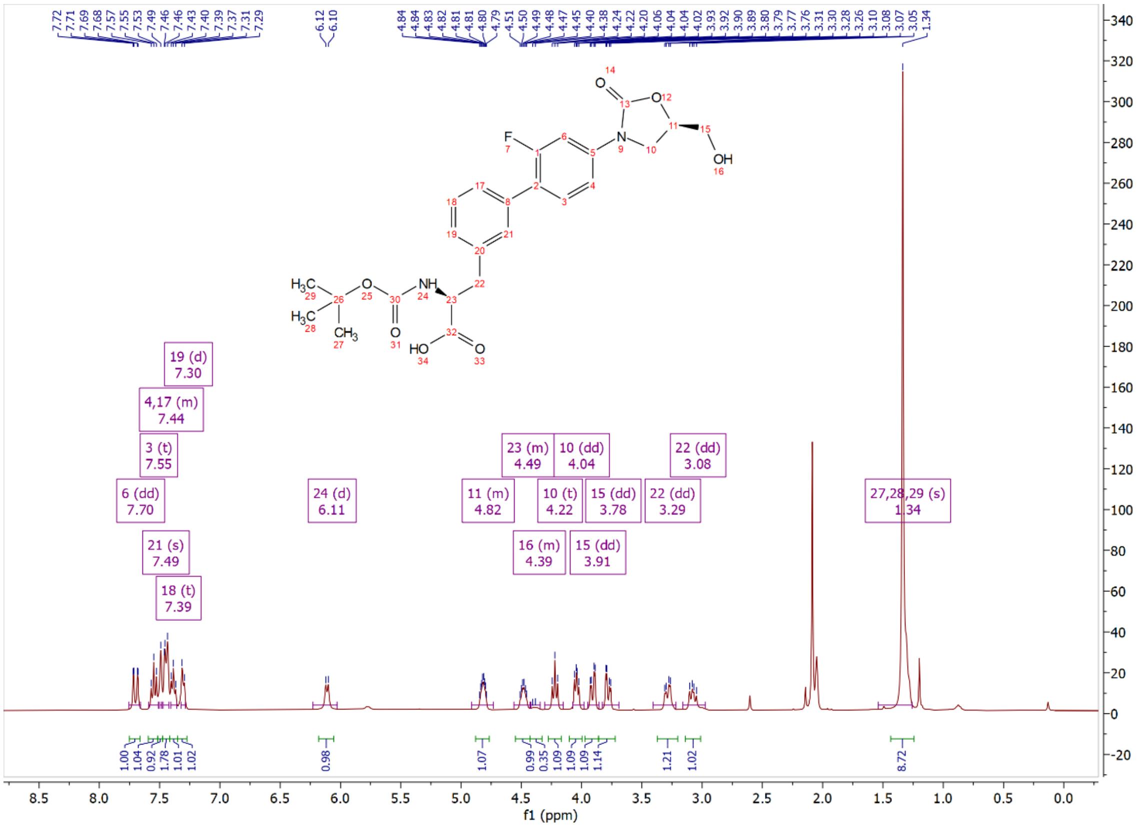


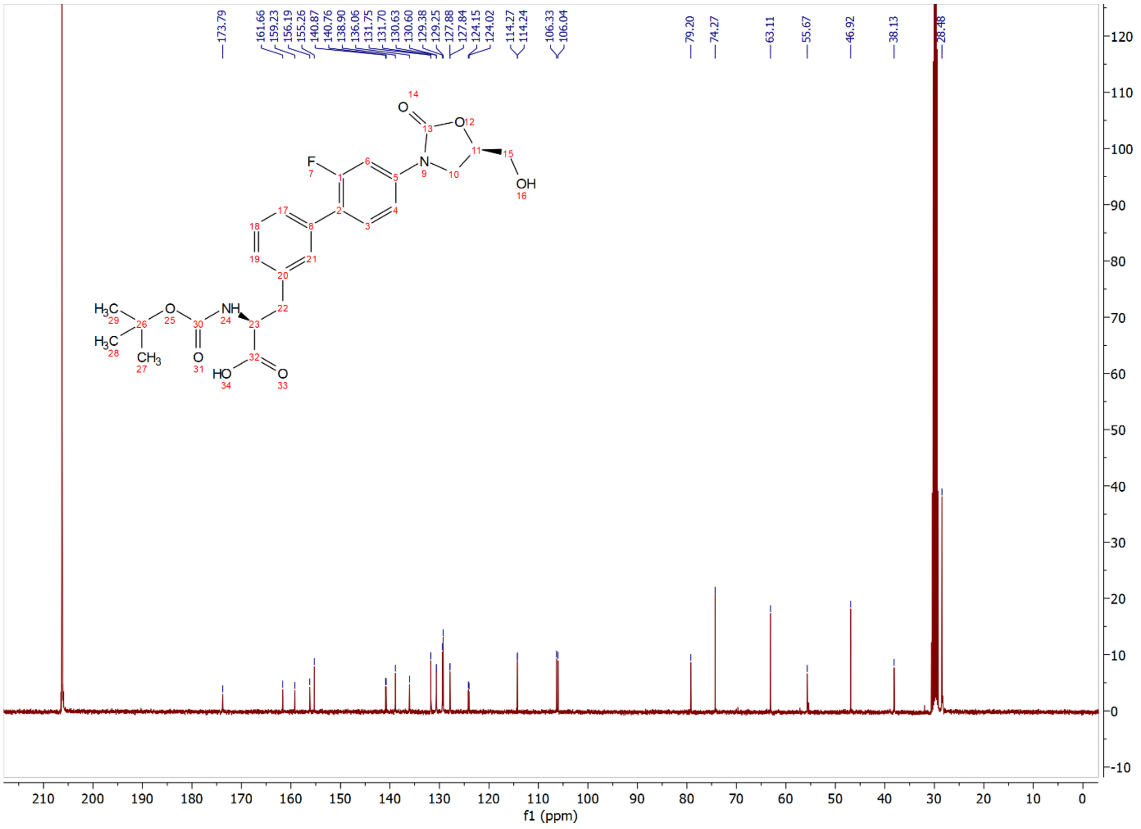

Compound **12ea**: $^{1}$H and $^{13}$C NMR Spectrum

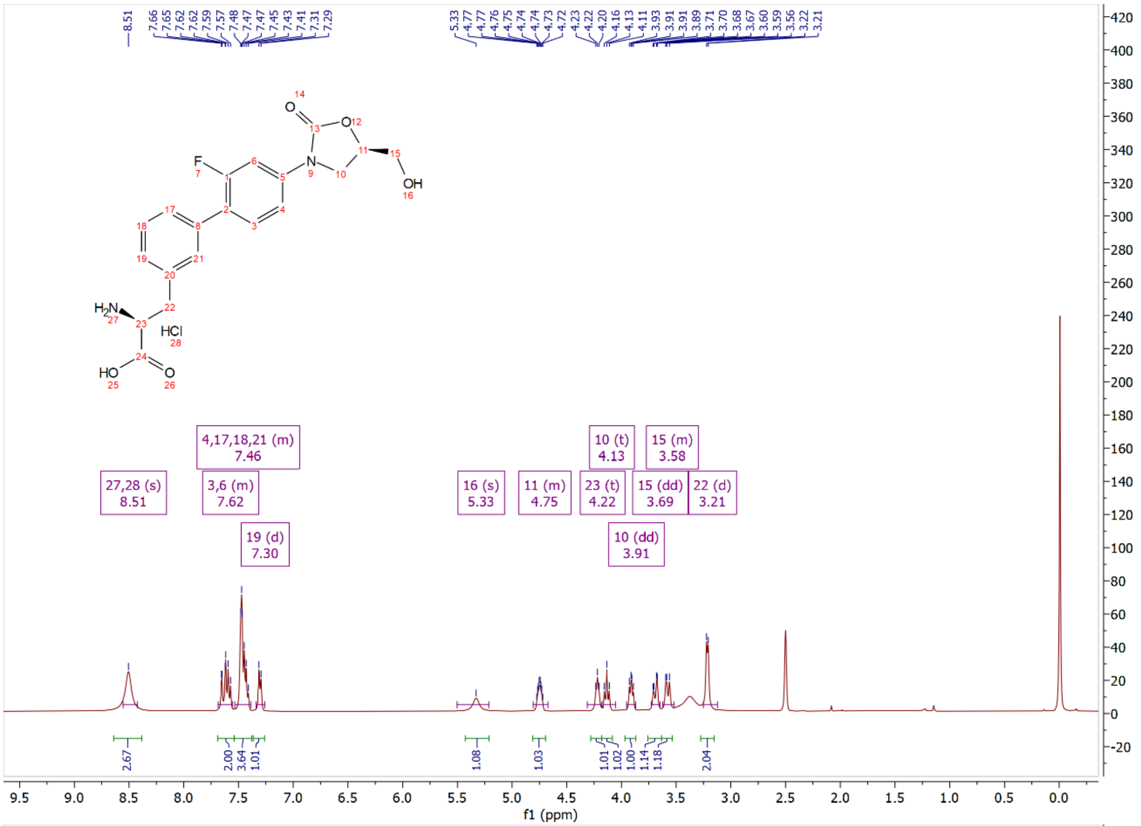


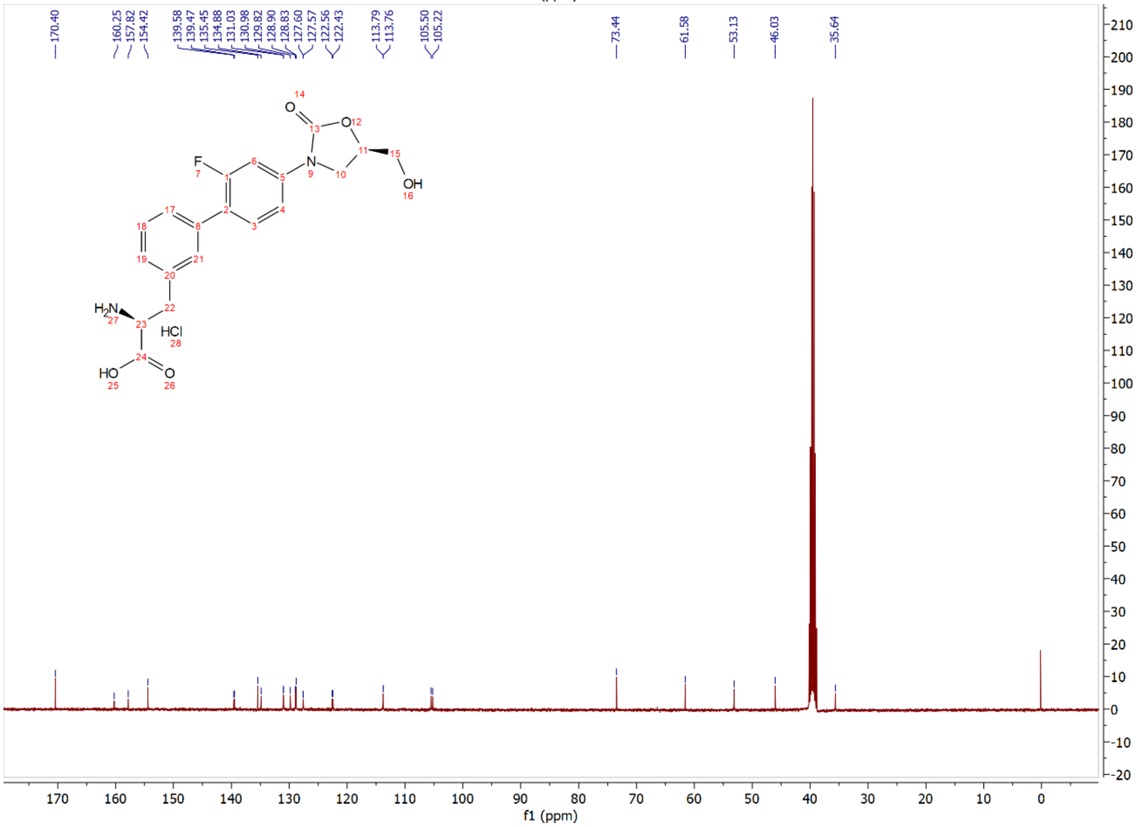

Compound **12f**: $^{1}$H and $^{13}$C NMR Spectrum

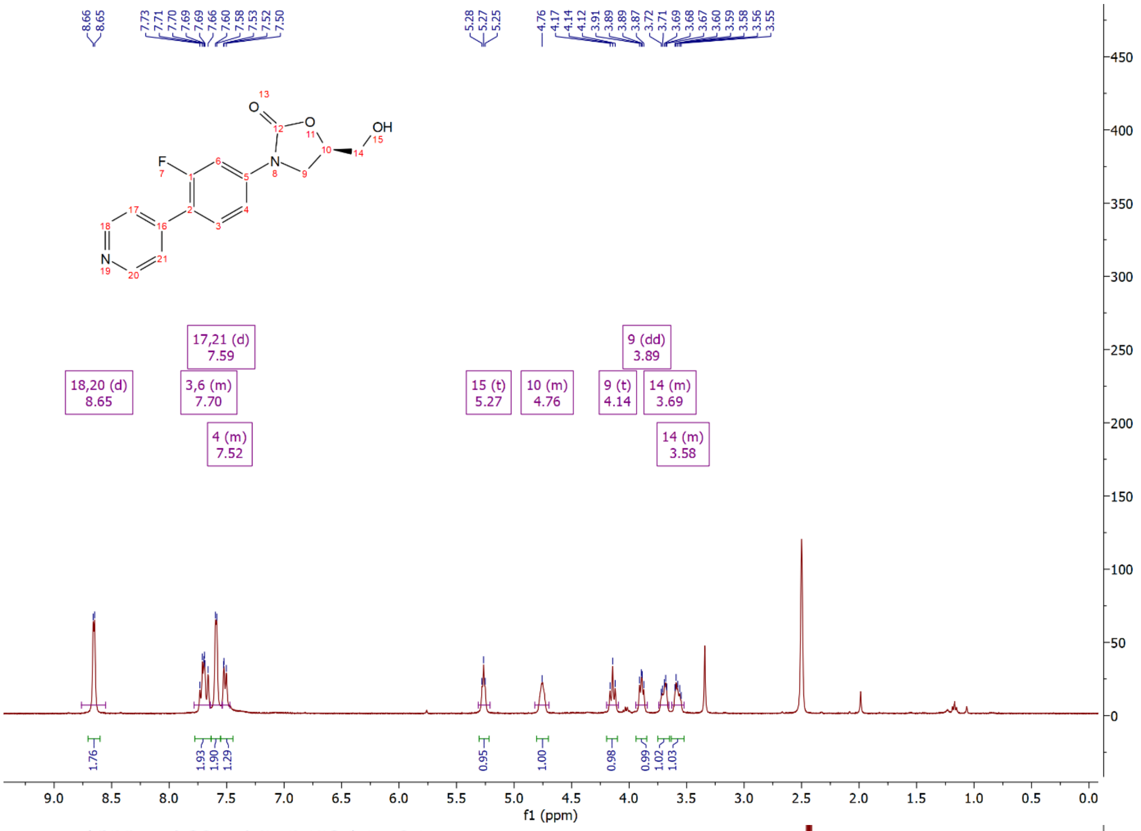

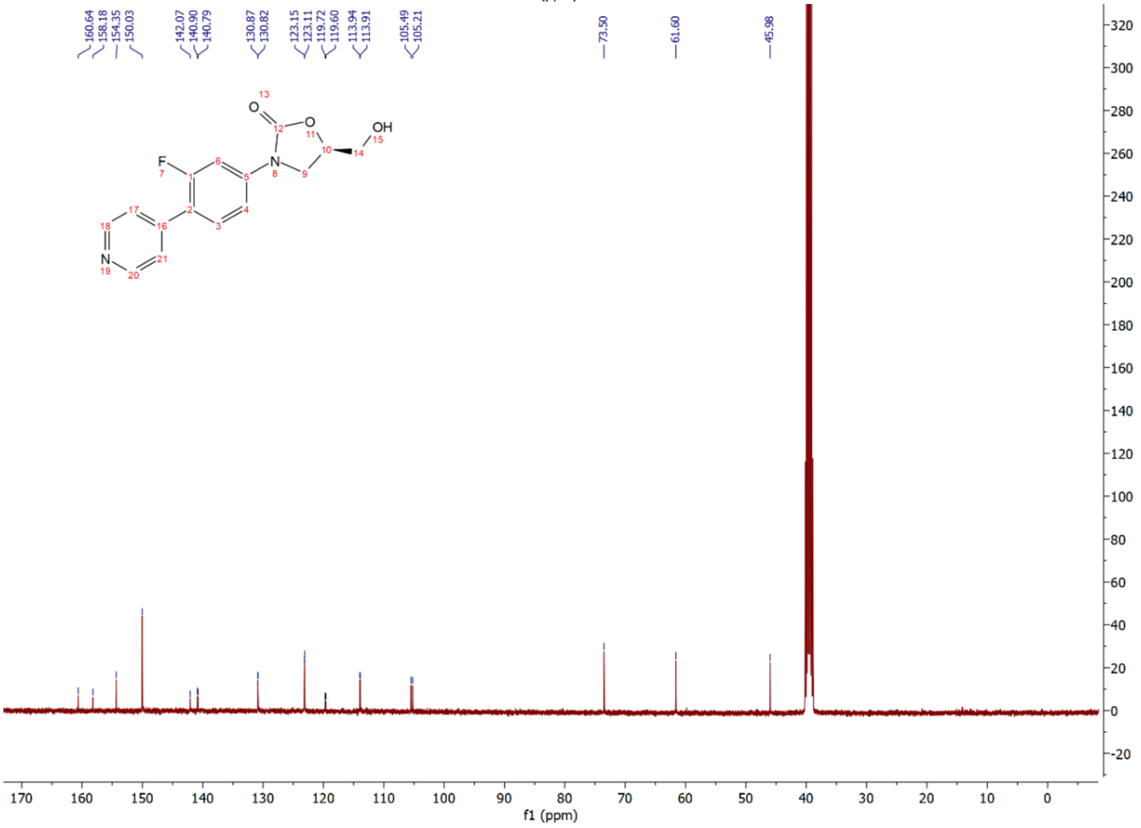

Compound **12g**: $^{1}$H and $^{13}$C NMR Spectrum

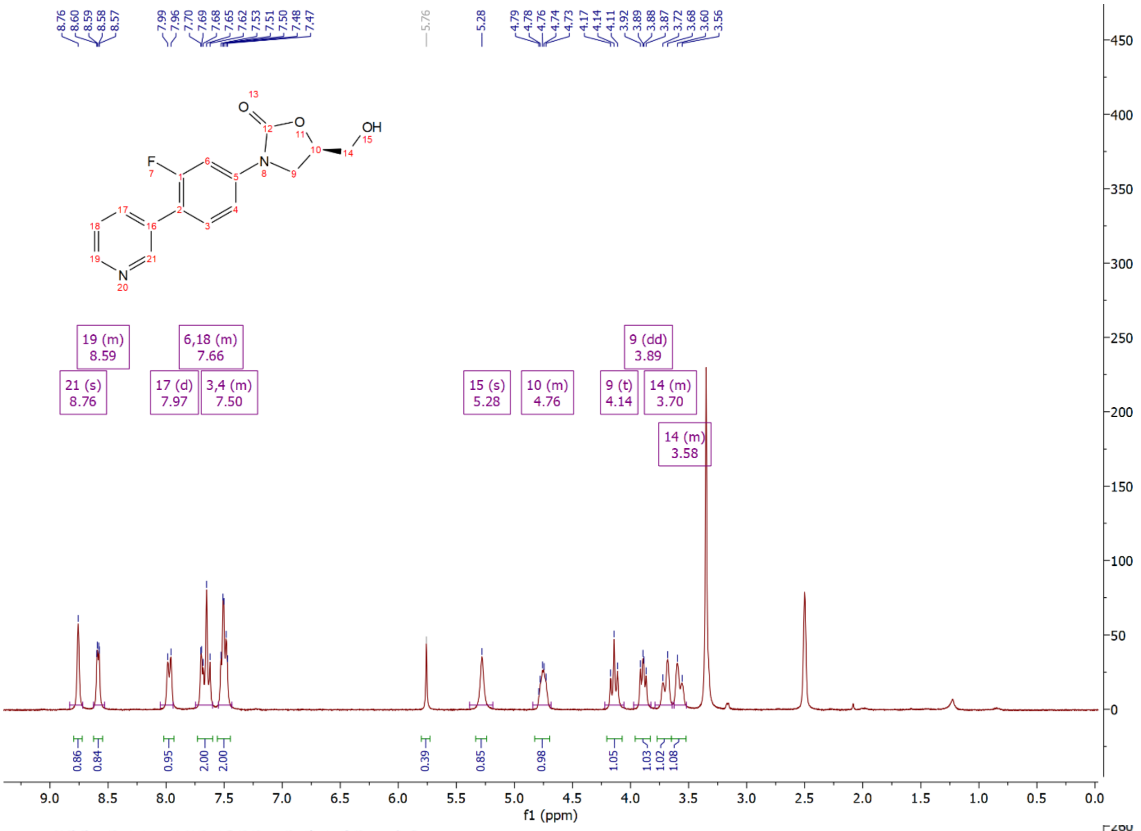

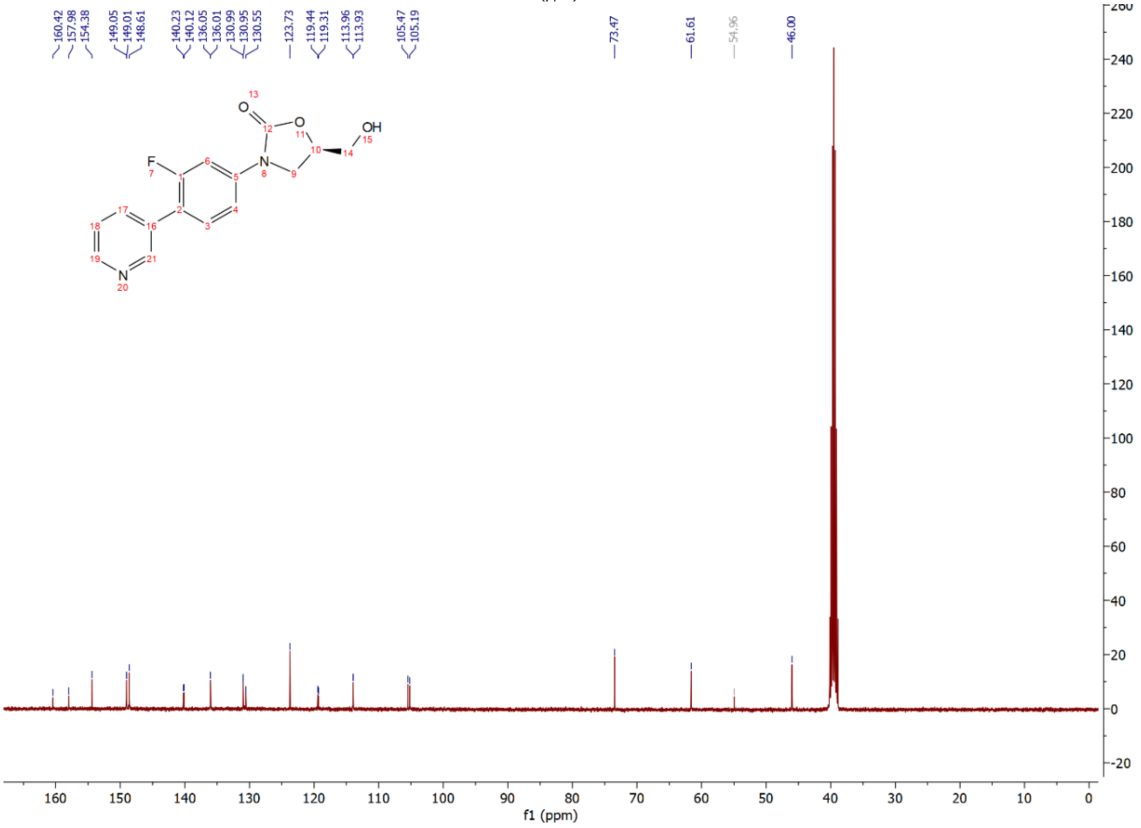

Compound **12h**: $^{1}$H and $^{13}$C NMR Spectrum

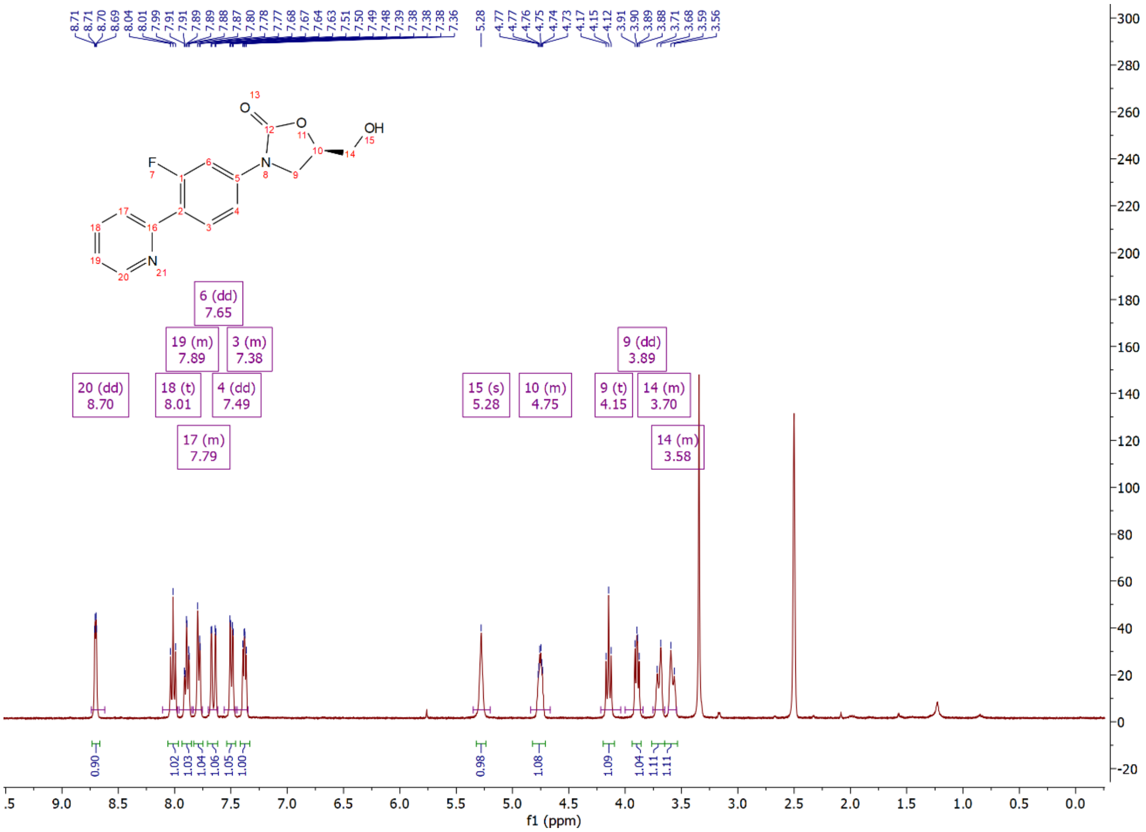

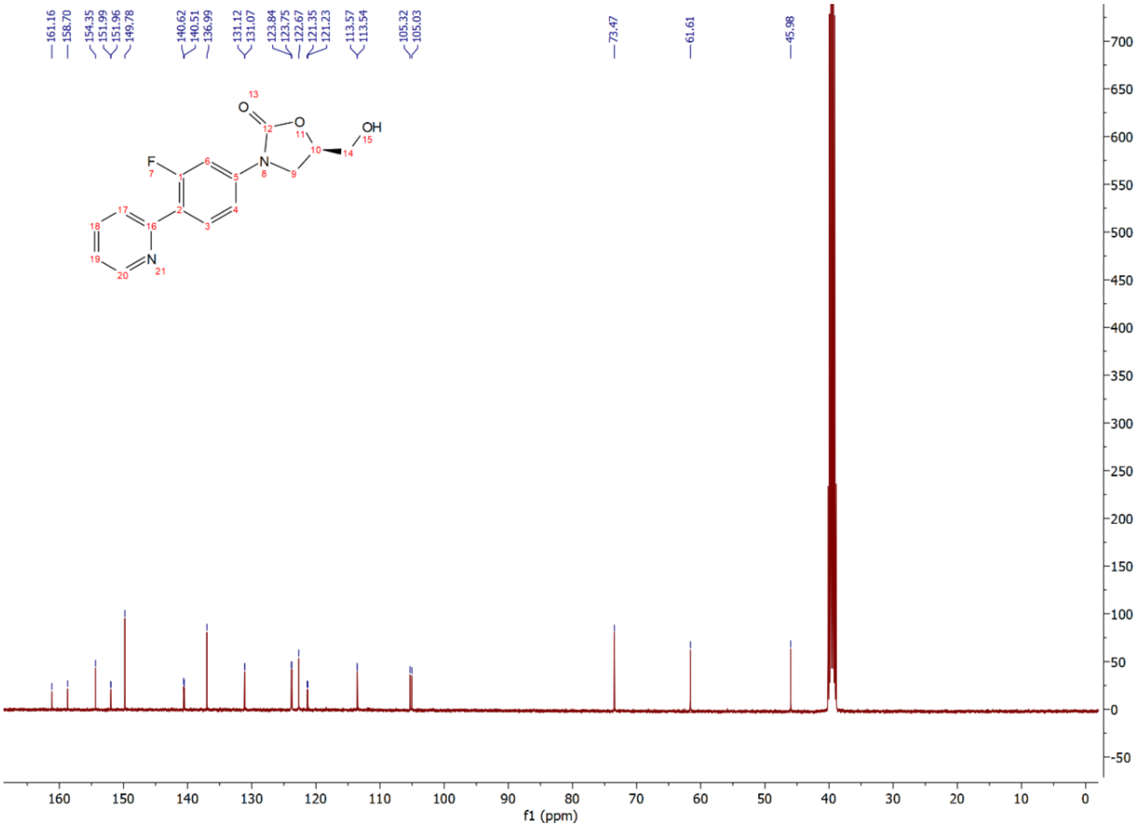

Compound **12i**: $^{1}$H and $^{13}$C NMR Spectrum

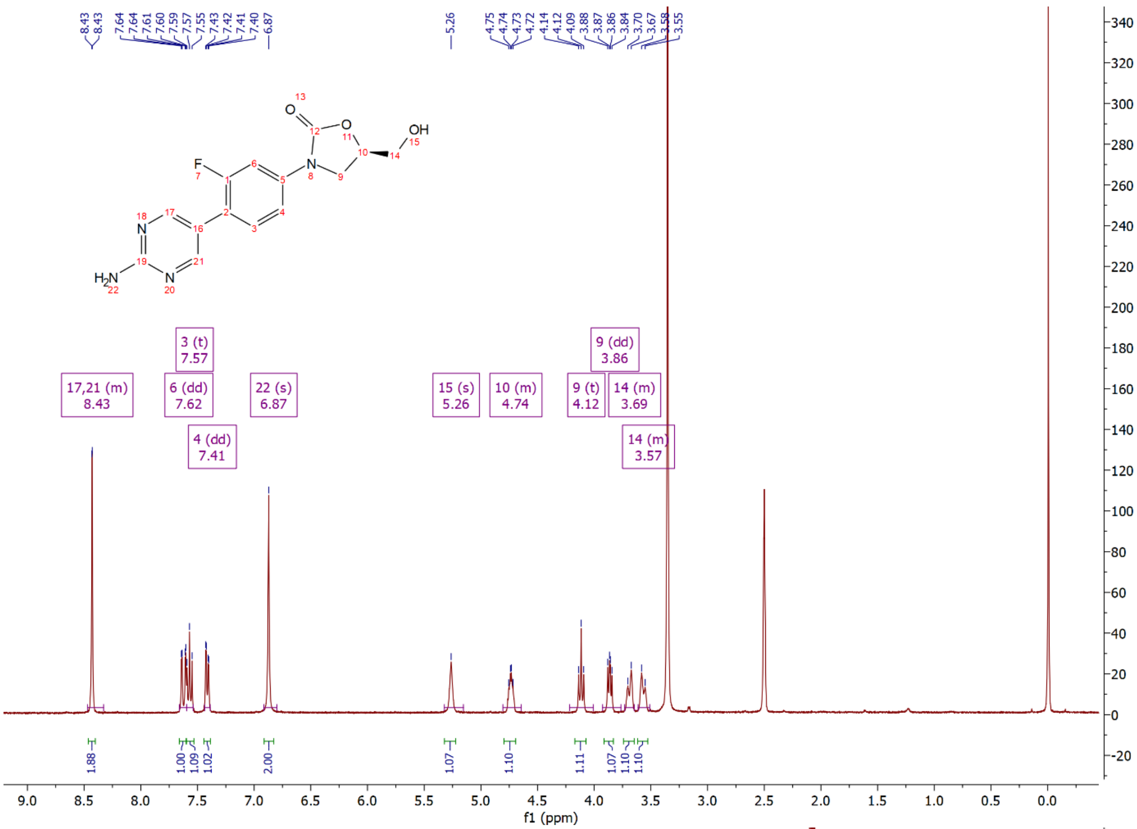

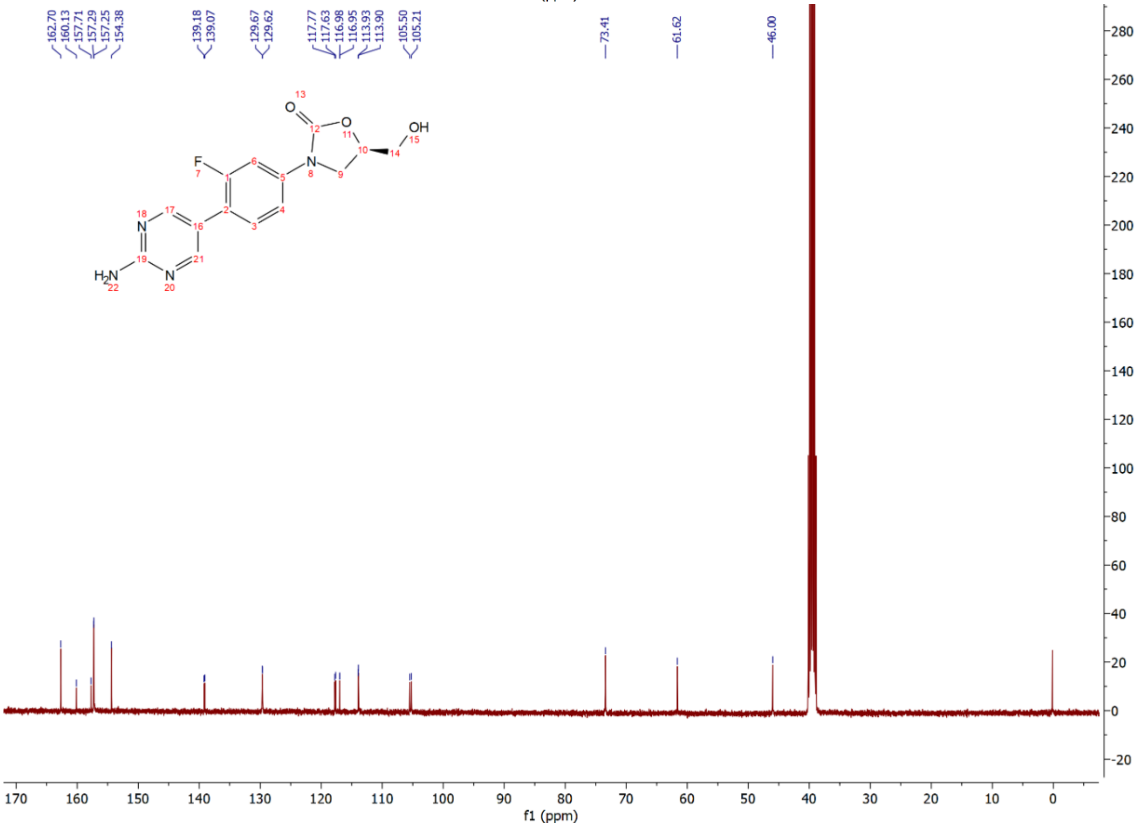

Compound **12j**: $^{1}H$ and $^{13}C$ NMR Spectrum

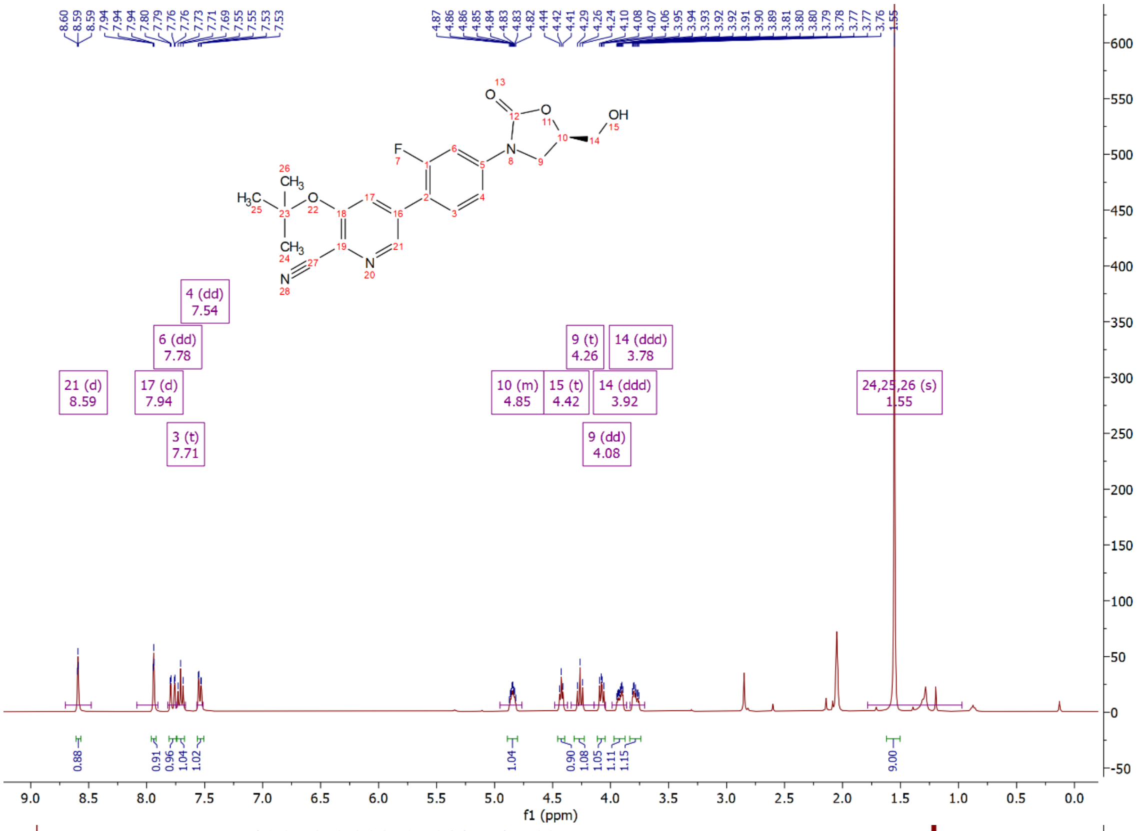

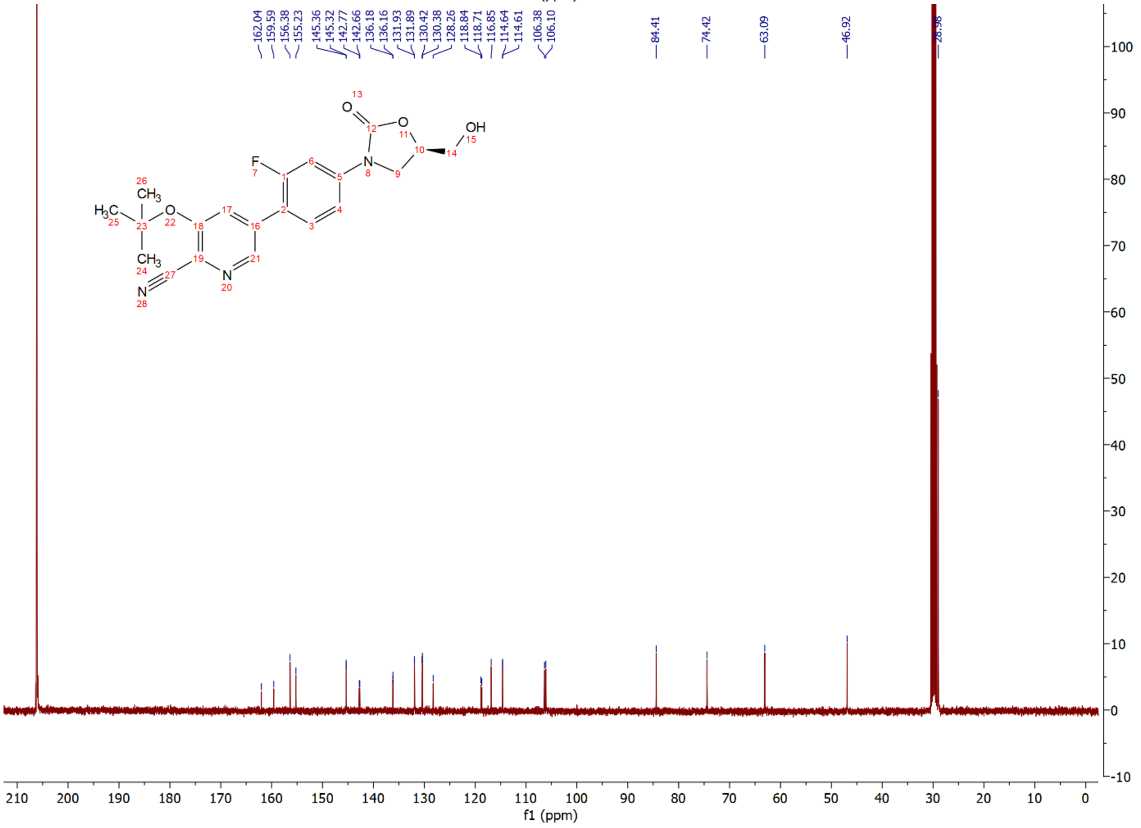

Compound **12k**: $^{1}$H and $^{13}$C NMR Spectrum

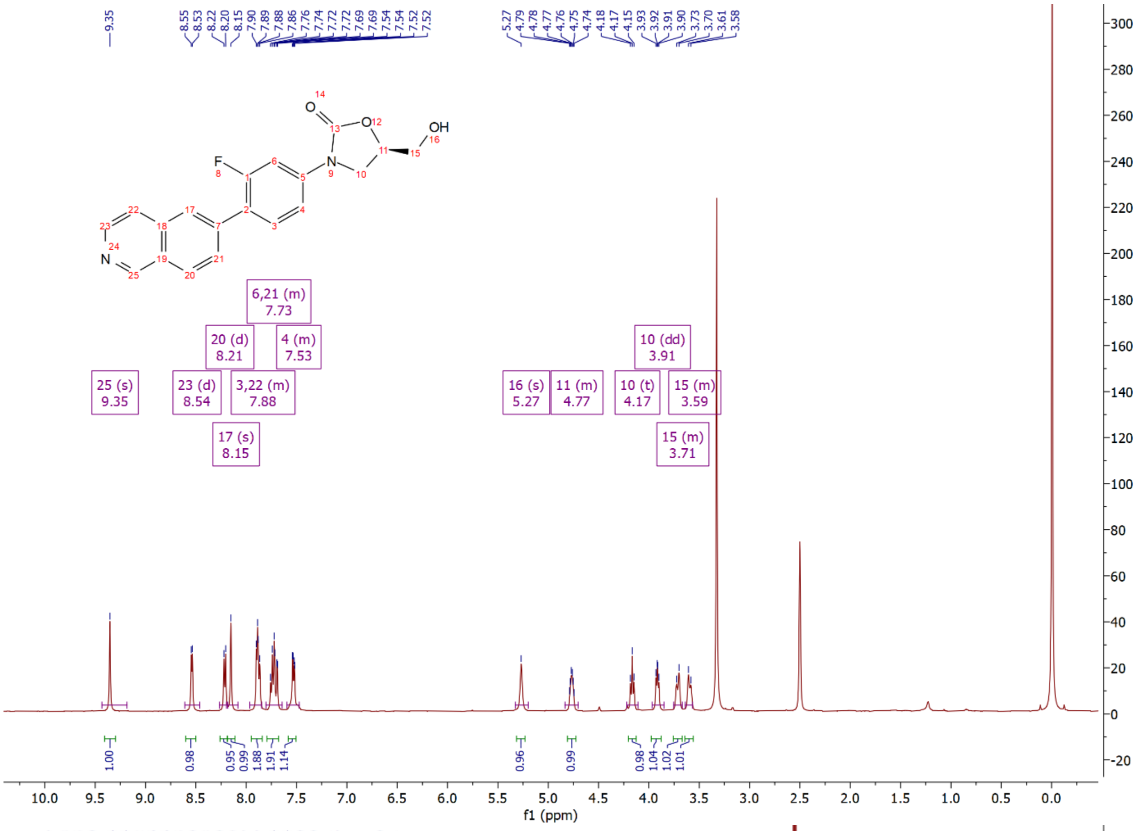


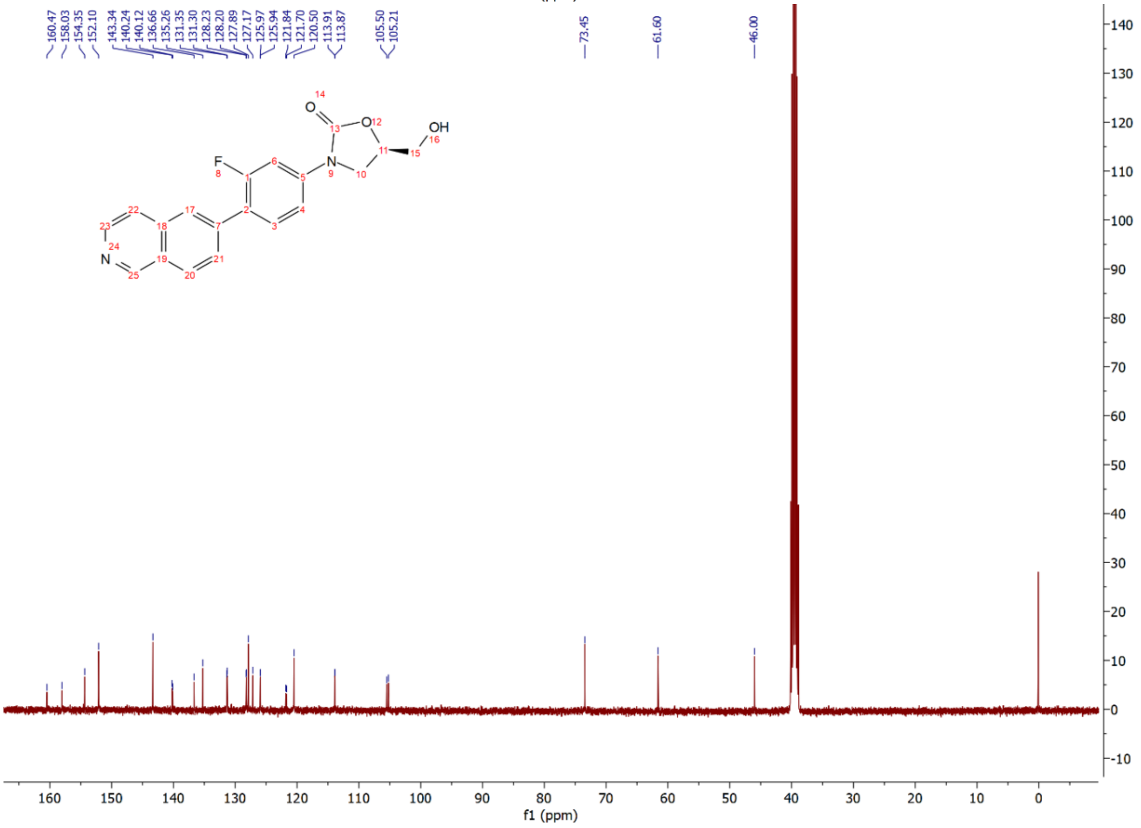

Compound **12l**: $^{1}$H and $^{13}$C NMR Spectrum

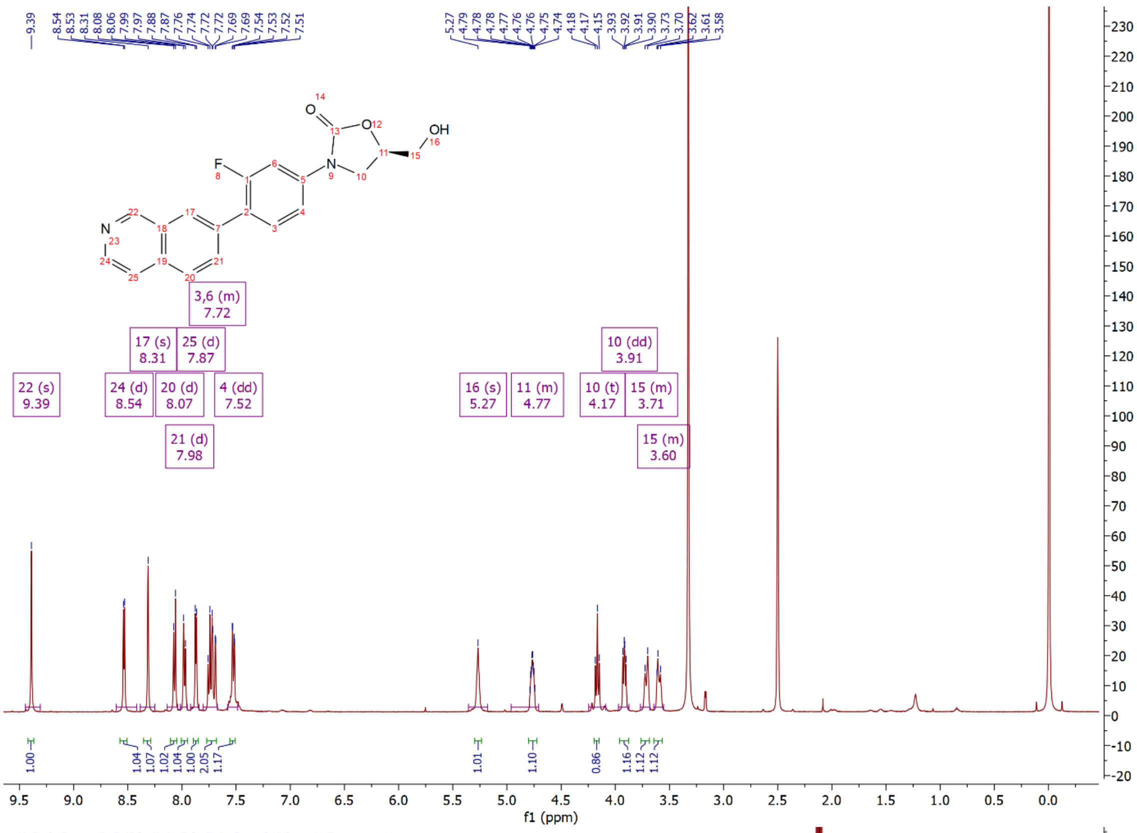

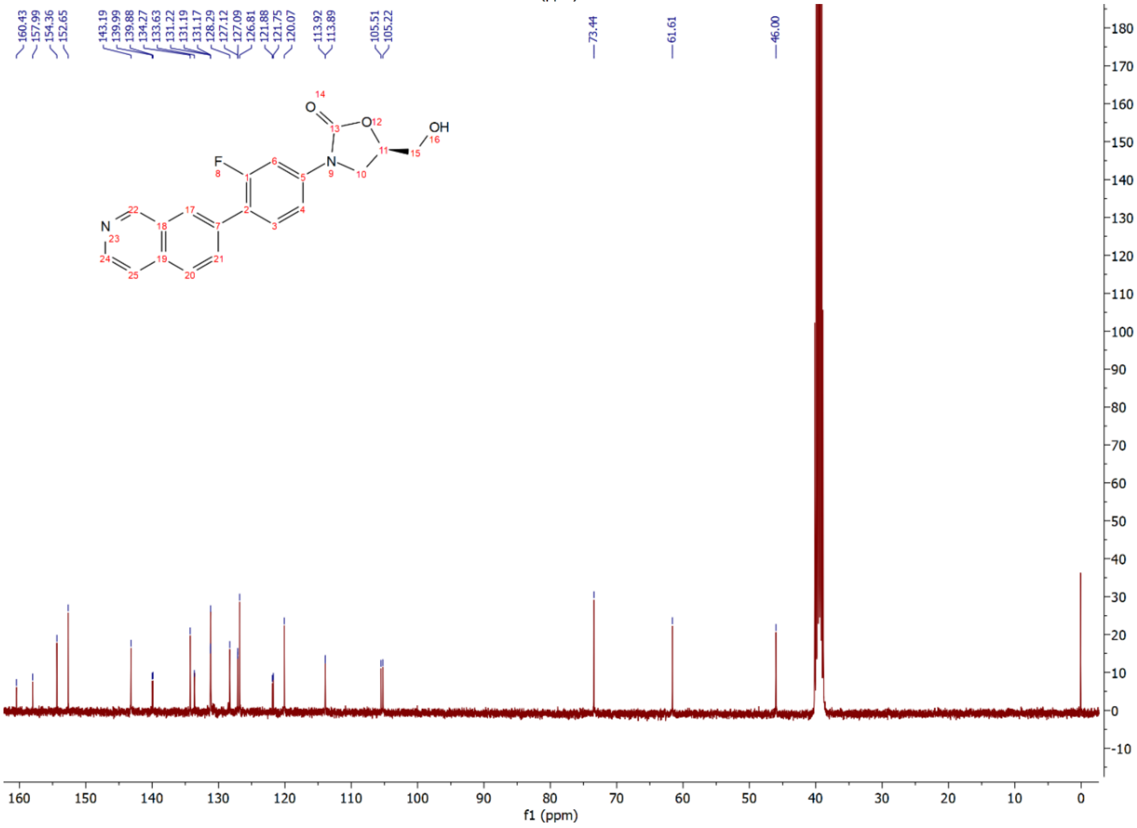

Compound **12m**: $^{1}H$ and $^{13}C$ NMR Spectrum

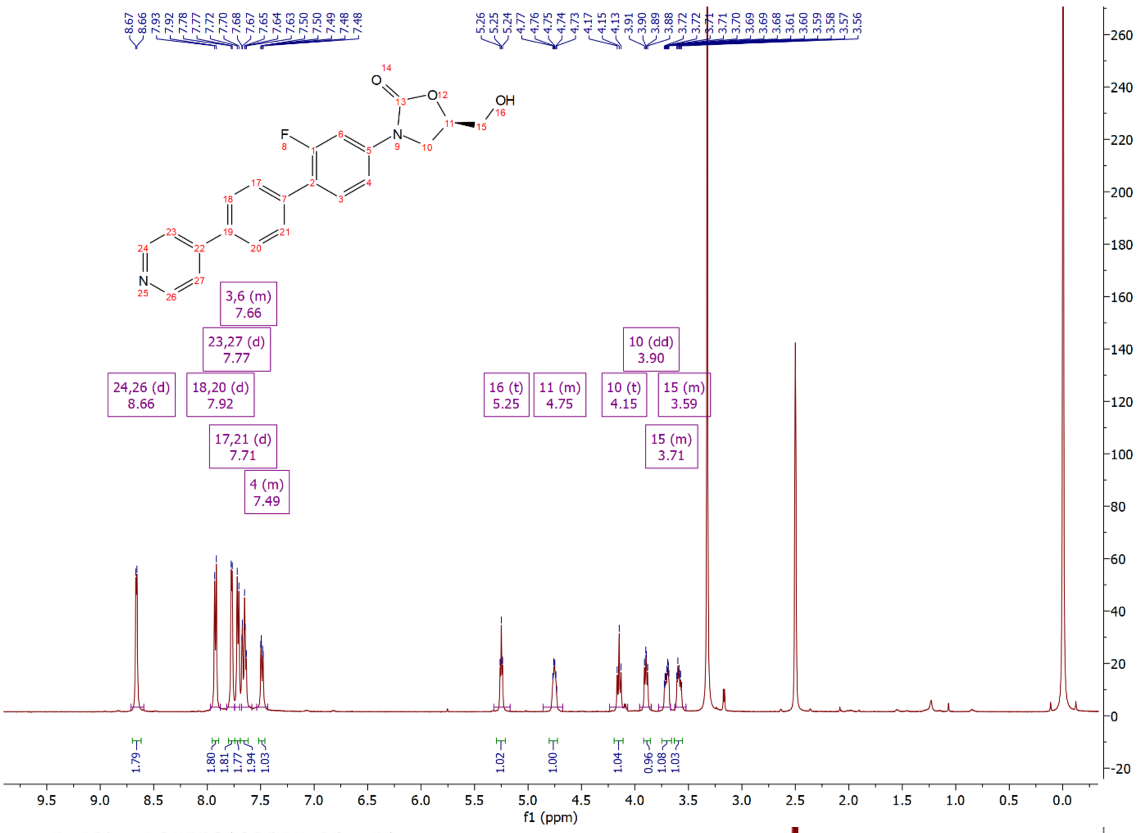

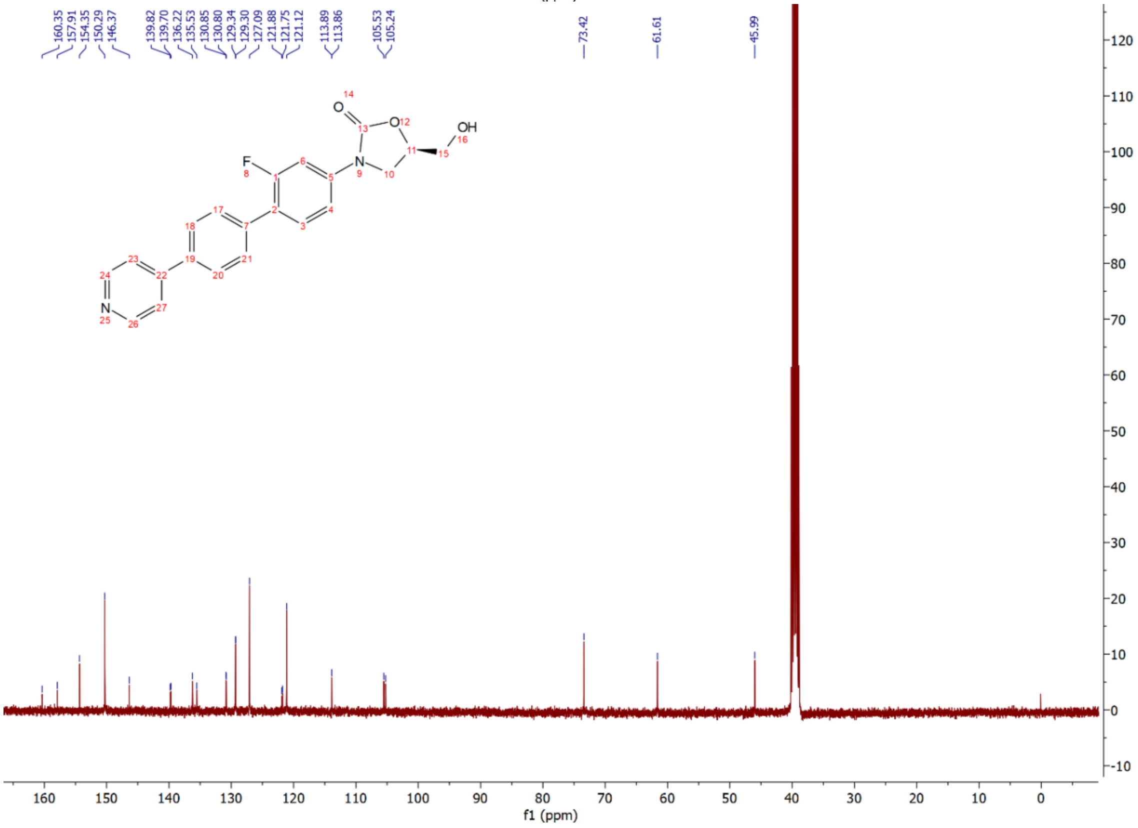

Compound **12n**: $^{1}$H and $^{13}$C NMR Spectrum

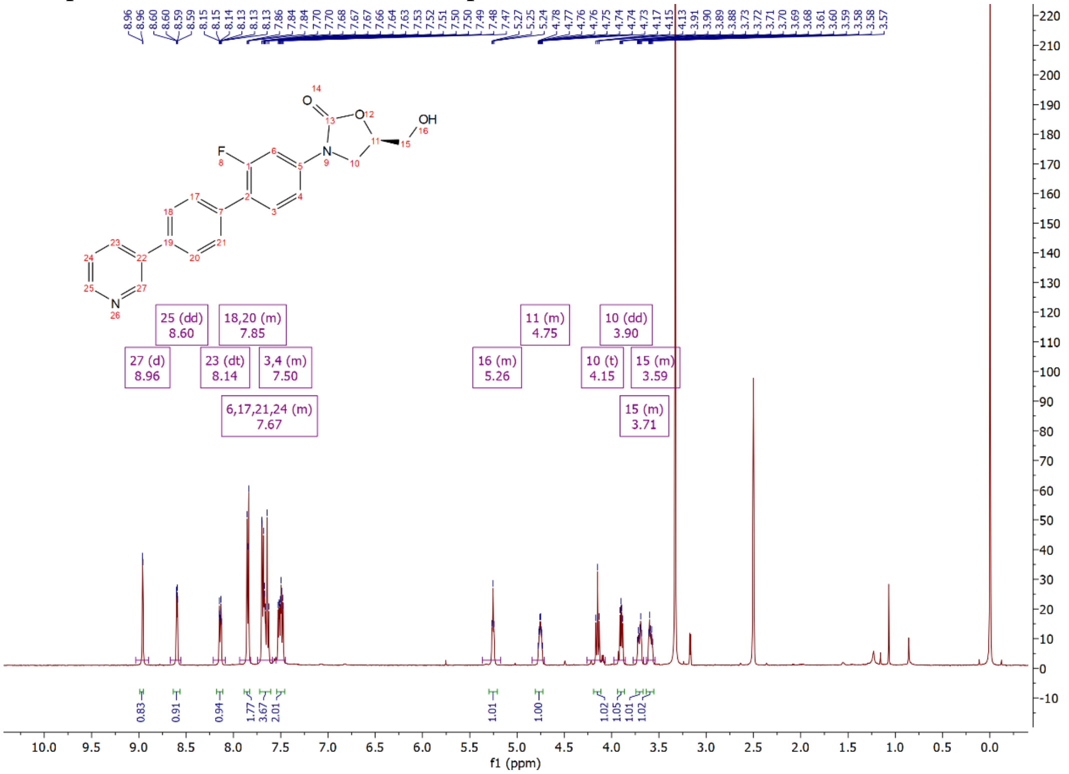

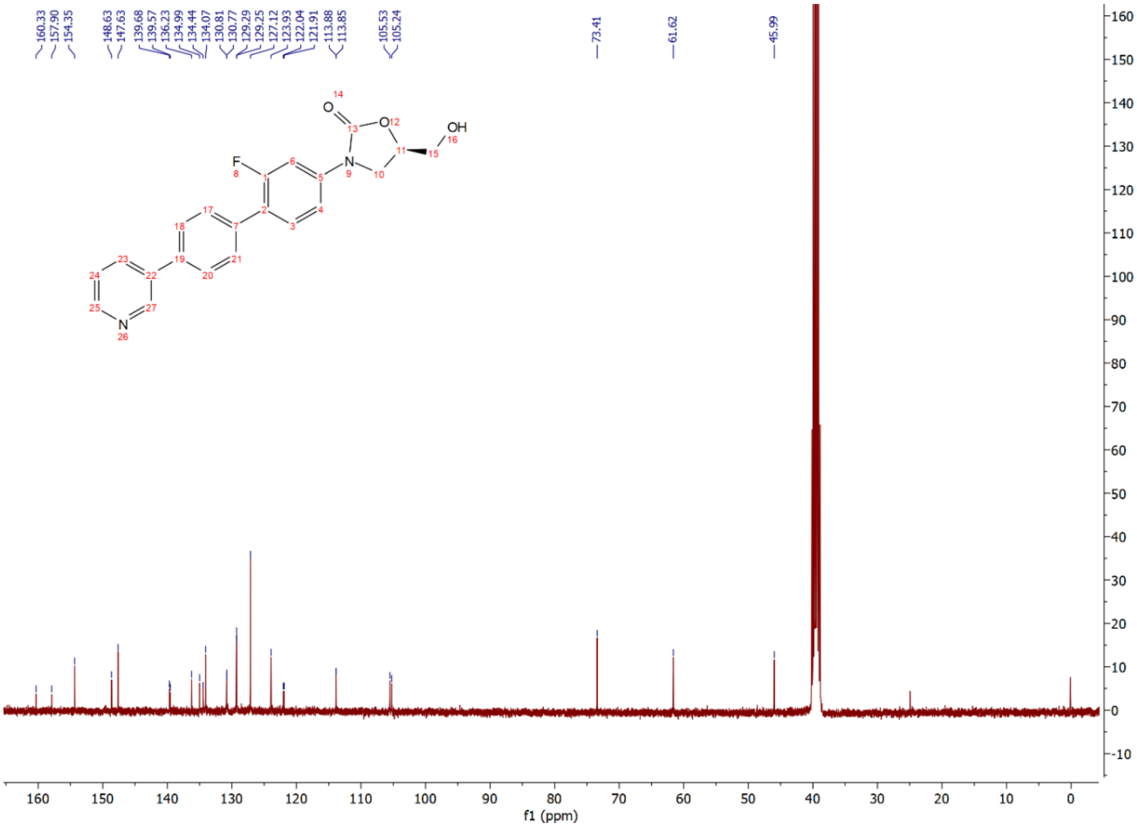

Compound **12o**: $^{1}$H and $^{13}$C NMR Spectrum

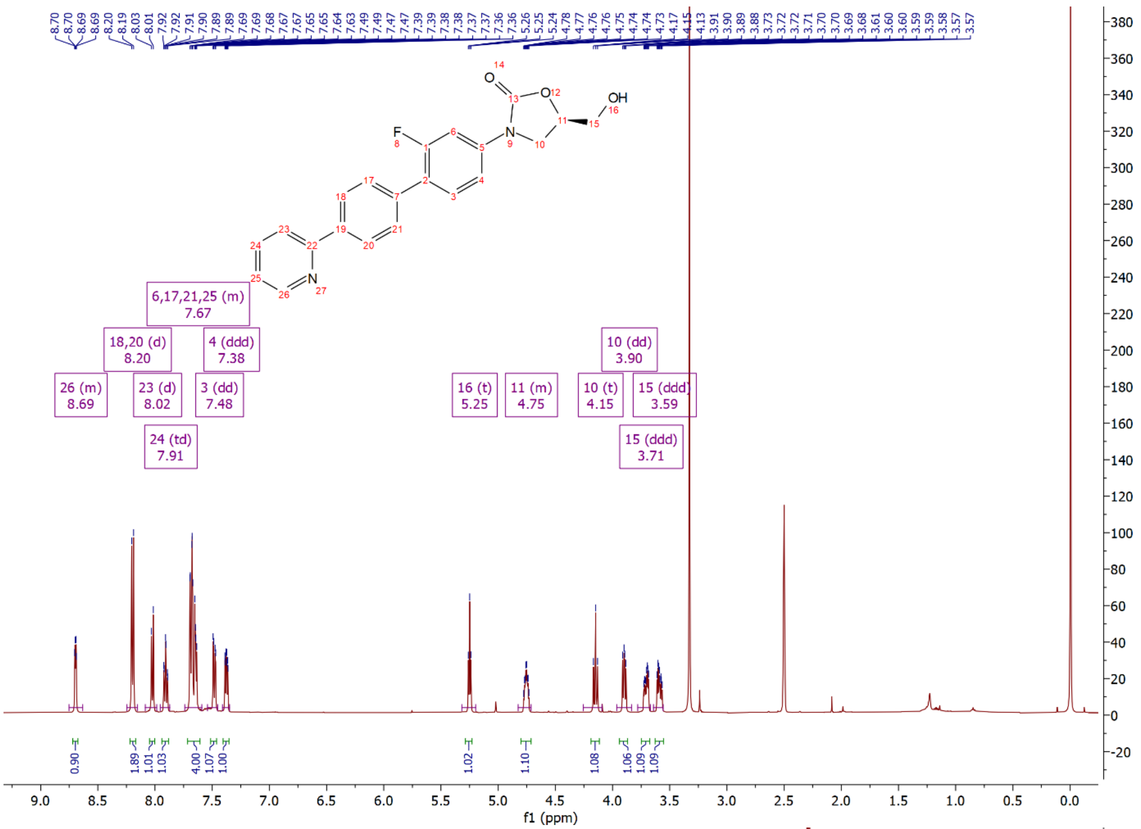

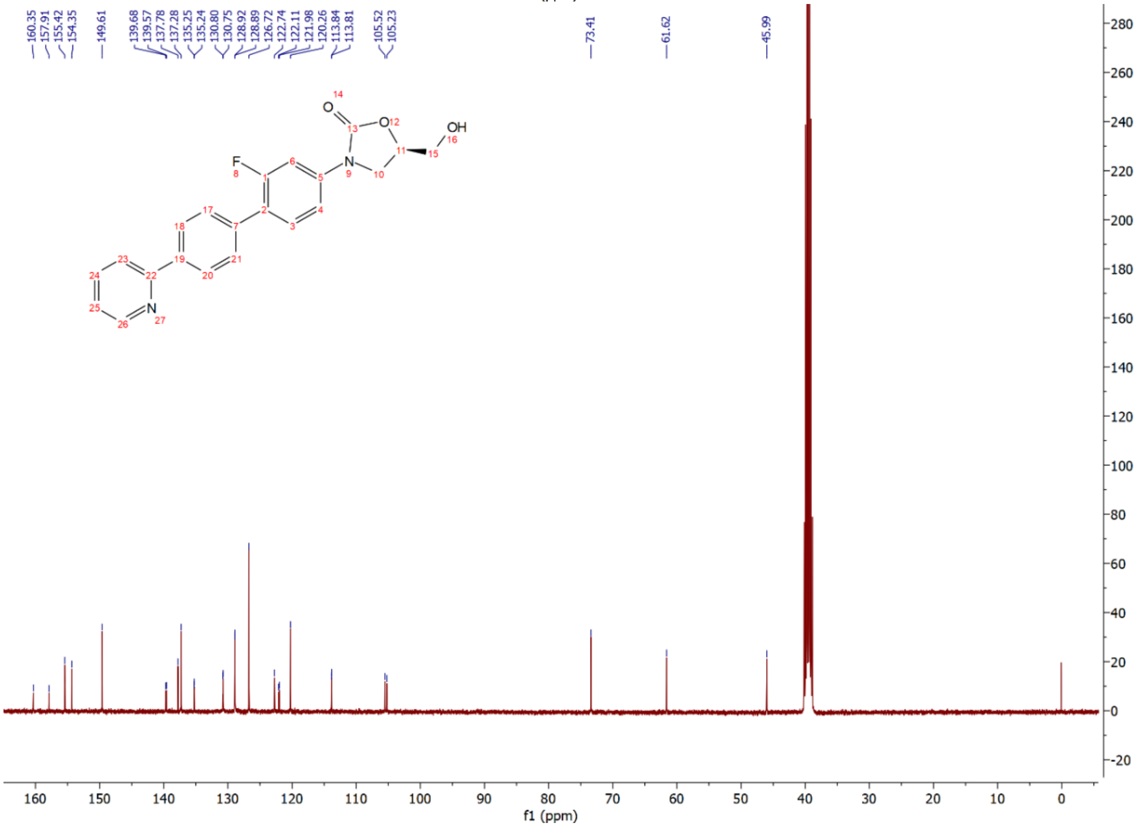

Compound **12p**: $^{1}$H and $^{13}$C NMR Spectrum

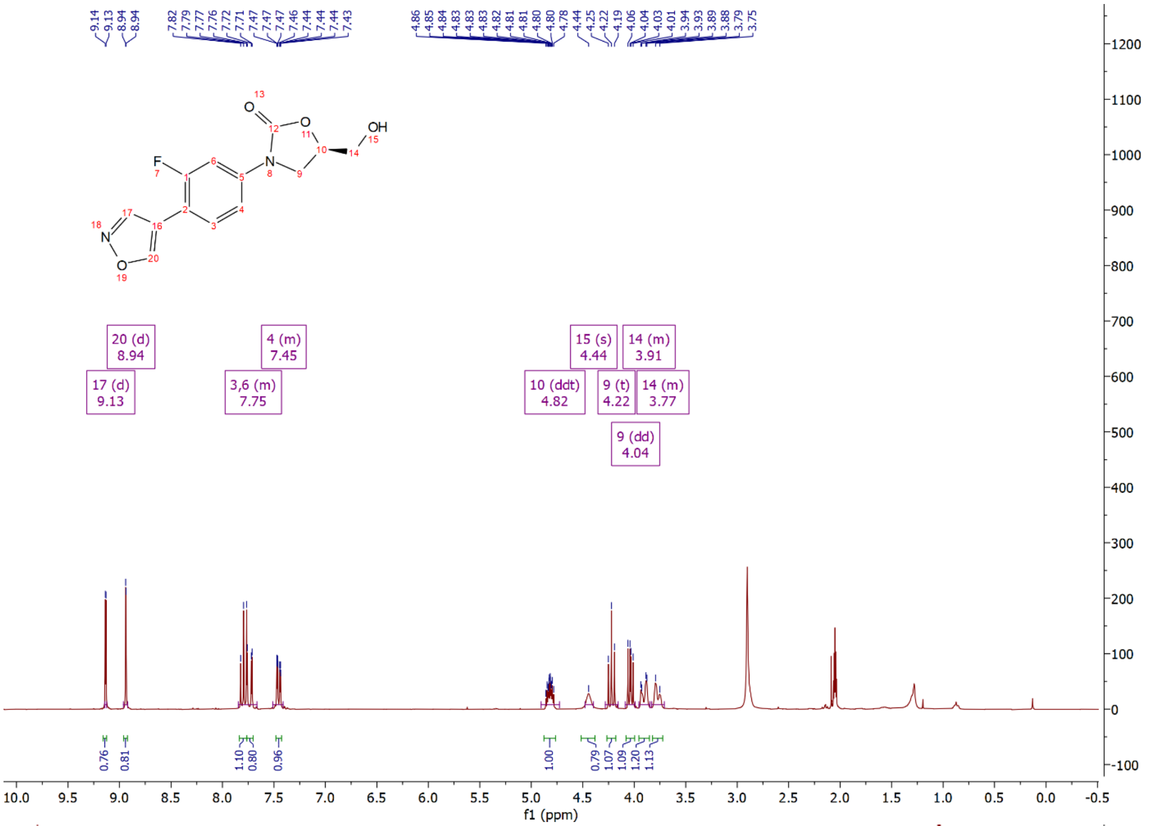

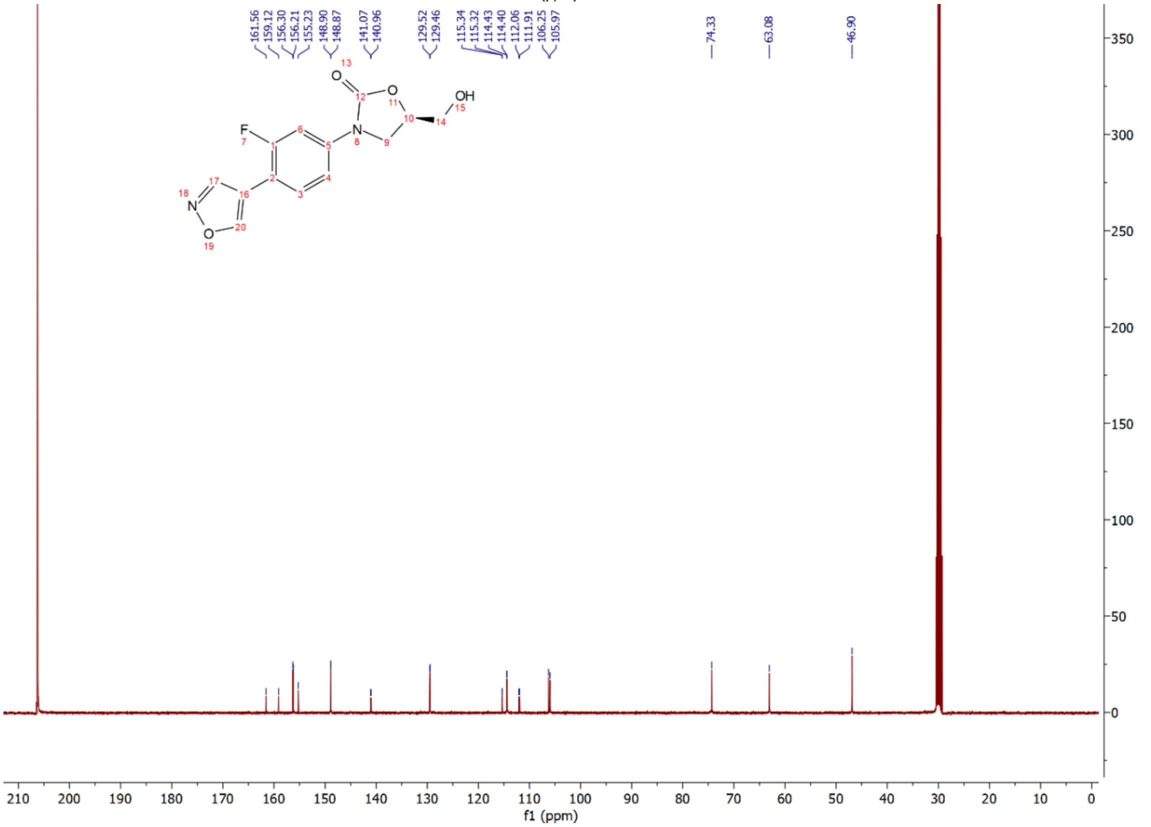

Compound **14**: $^{1}$H and $^{13}$C NMR Spectrum

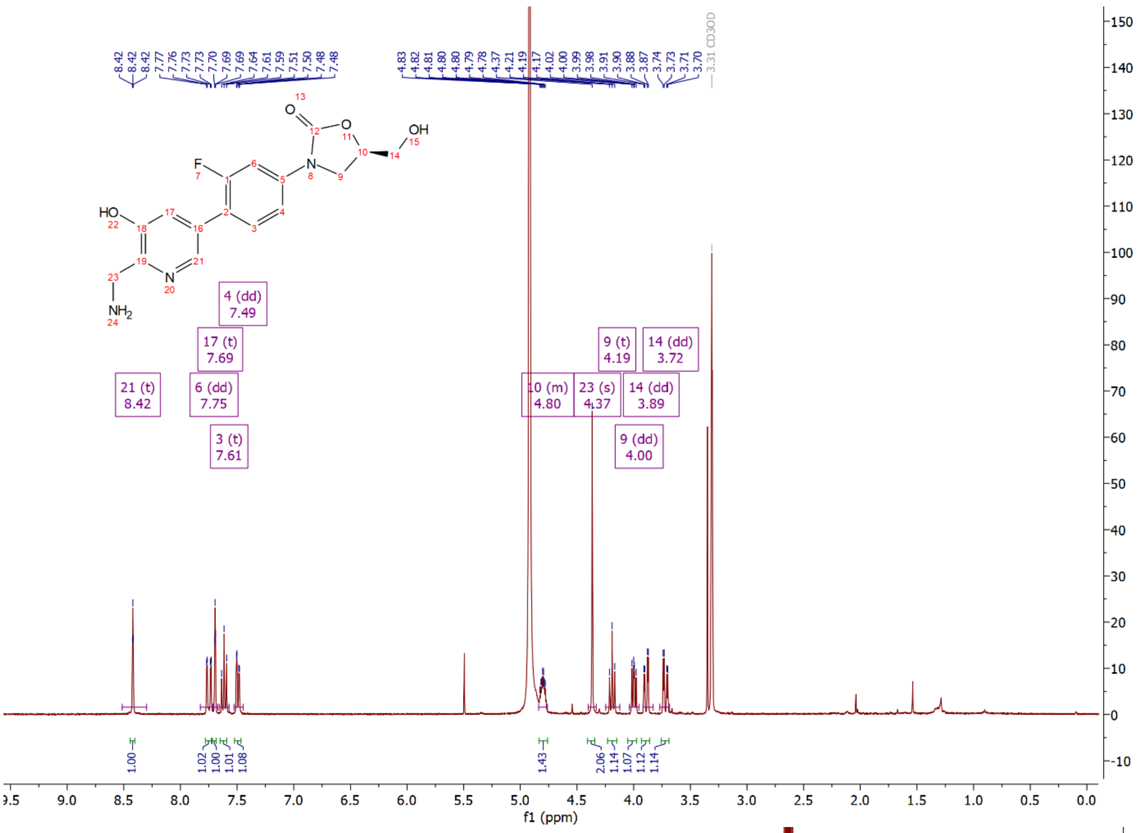

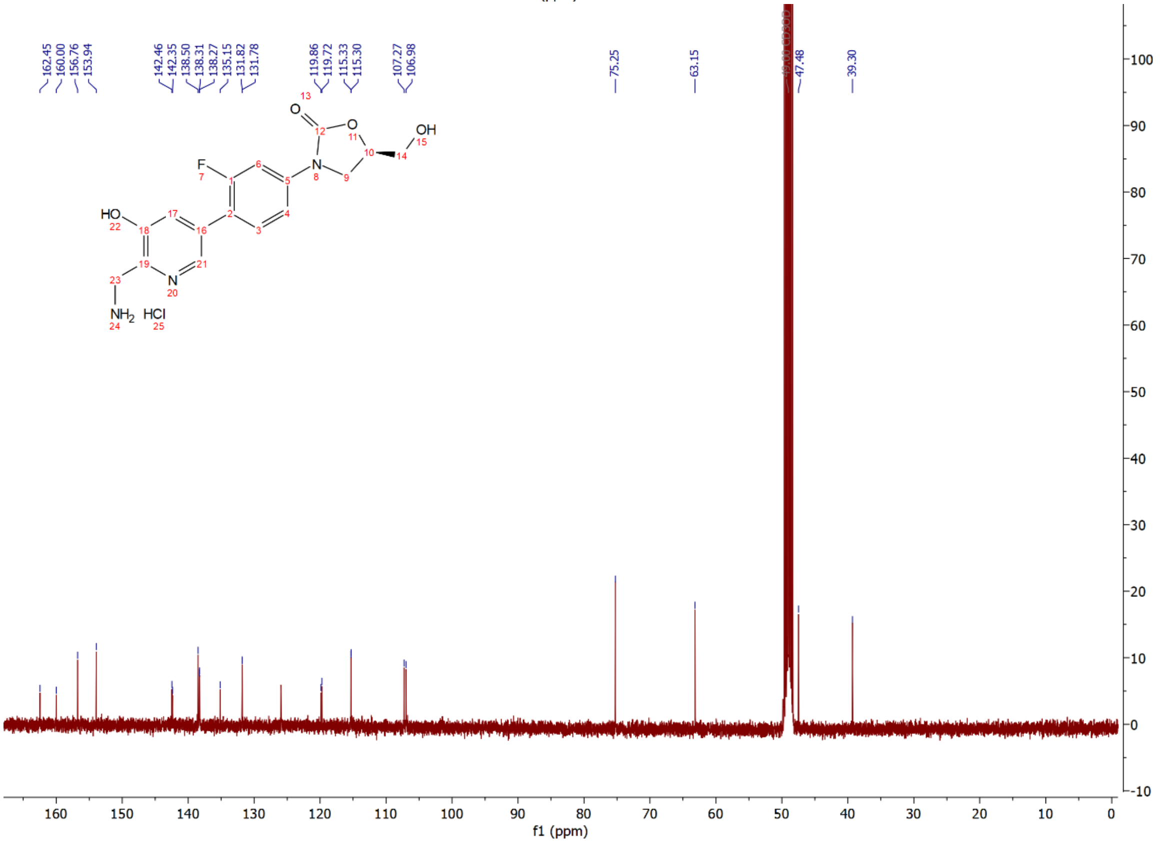

Compound **15a**: $^{1}$H and $^{13}$C NMR Spectrum

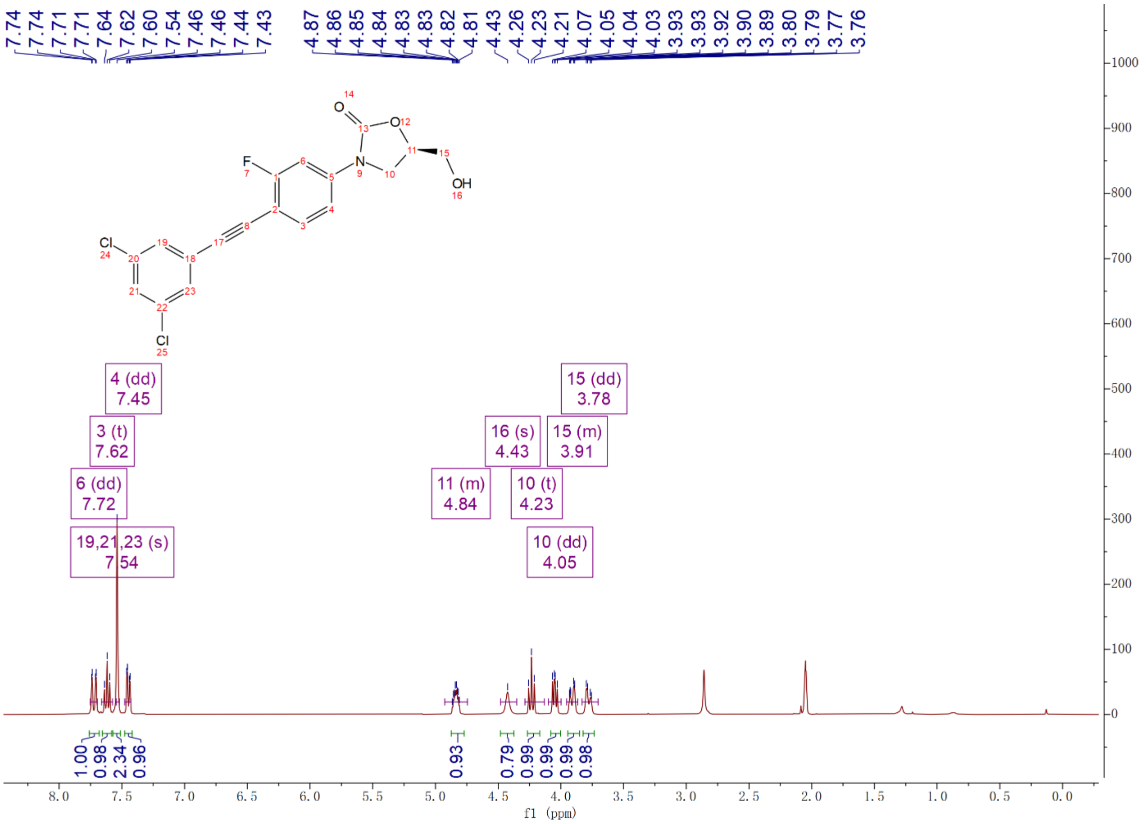


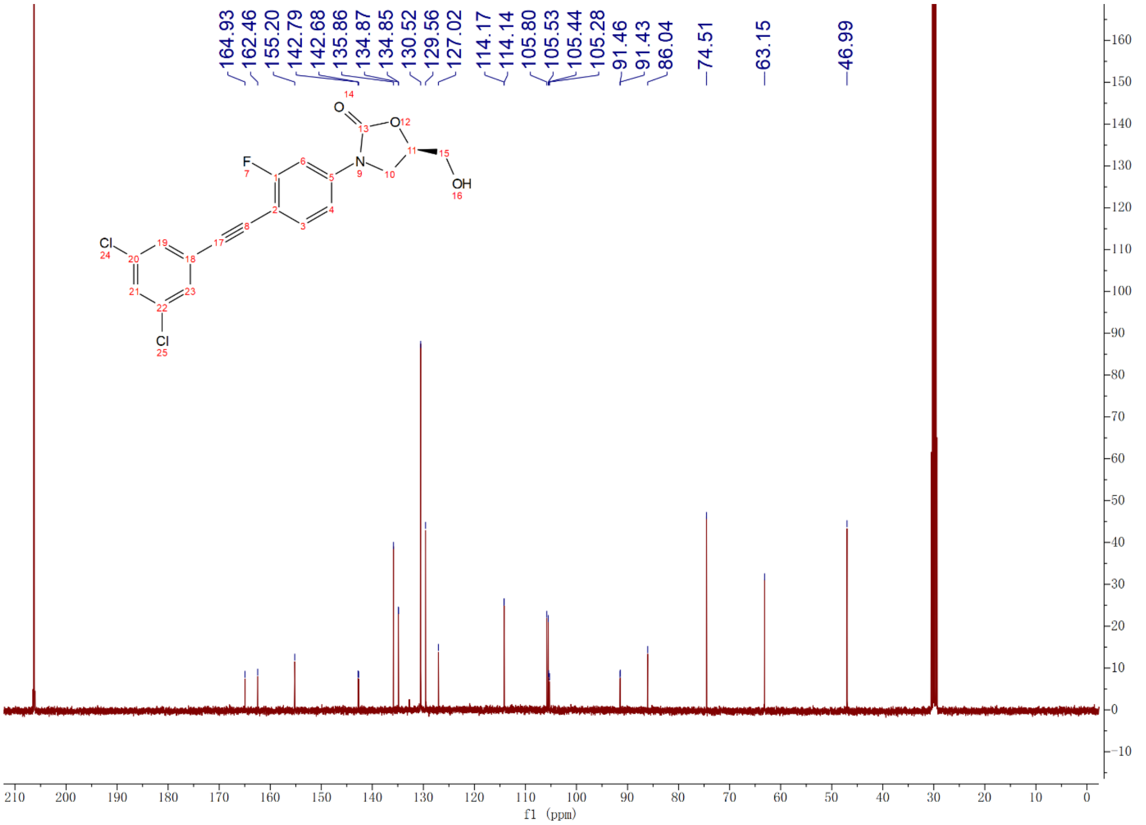

Compound **15b**: $^{1}$H and $^{13}$C NMR Spectrum

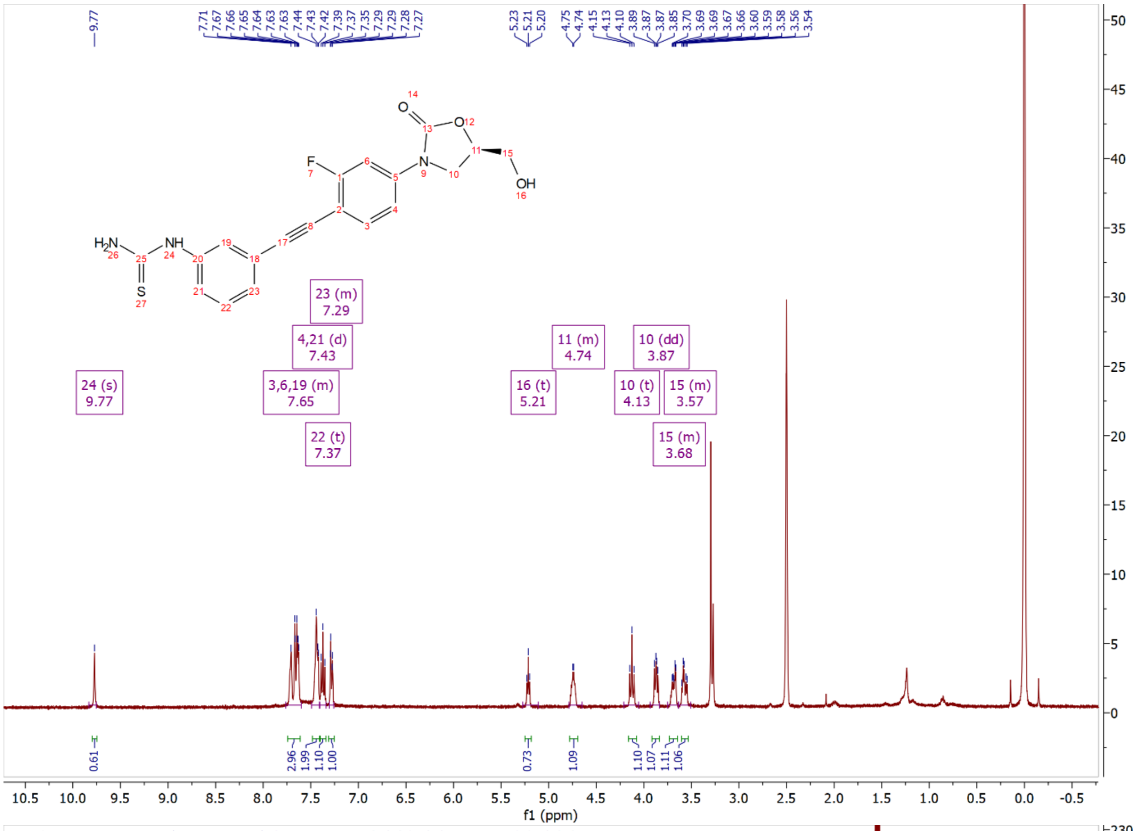

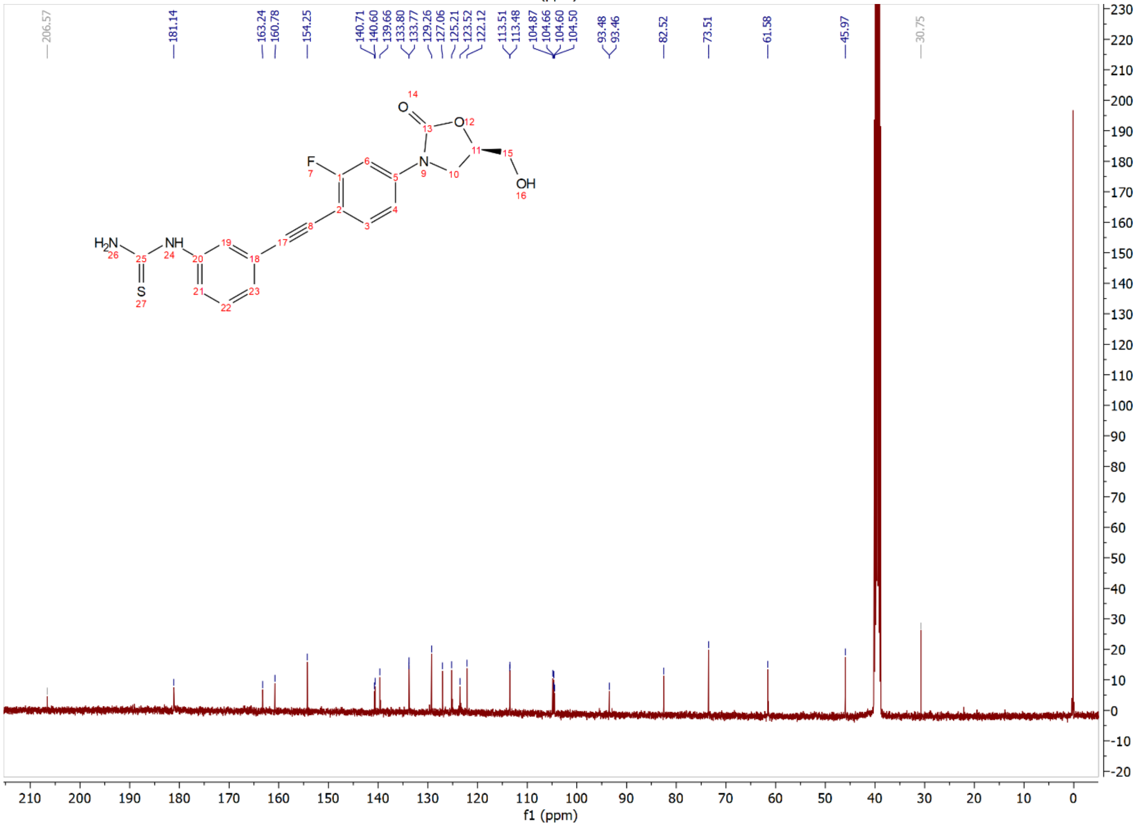

Compound **15c**: $^{1}$H and $^{13}$C NMR Spectrum

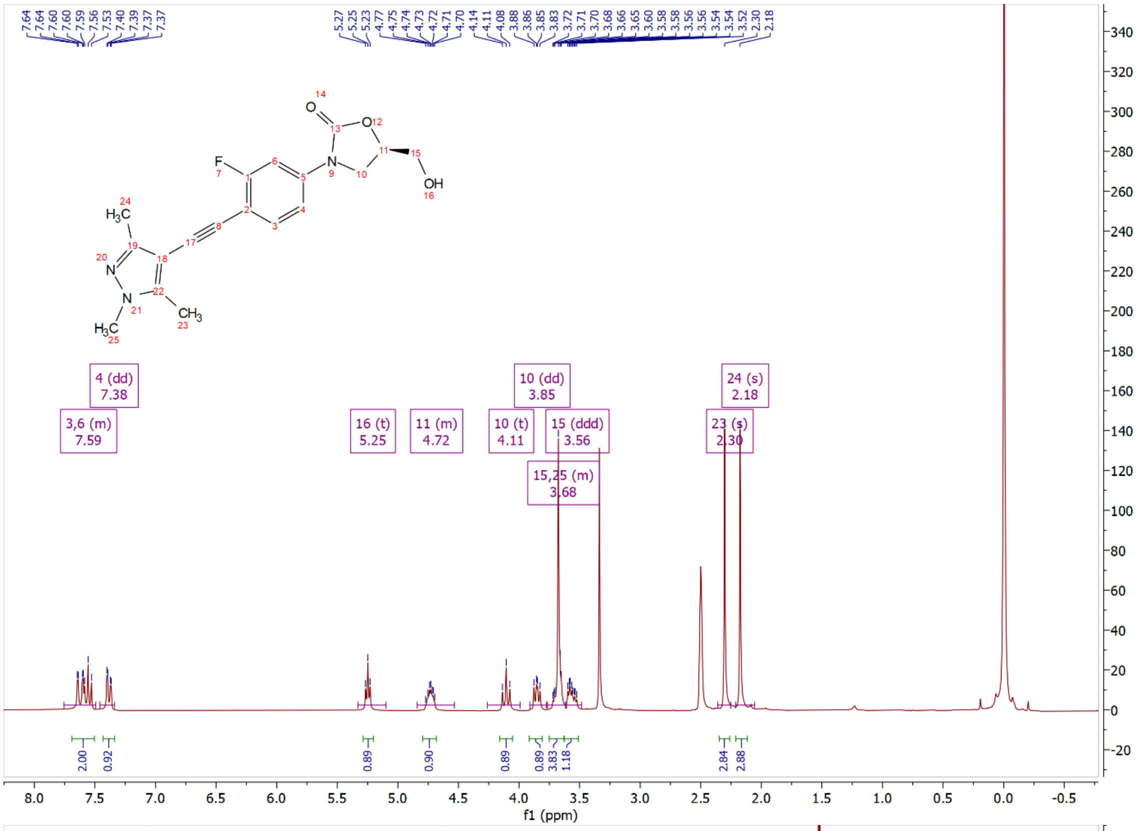

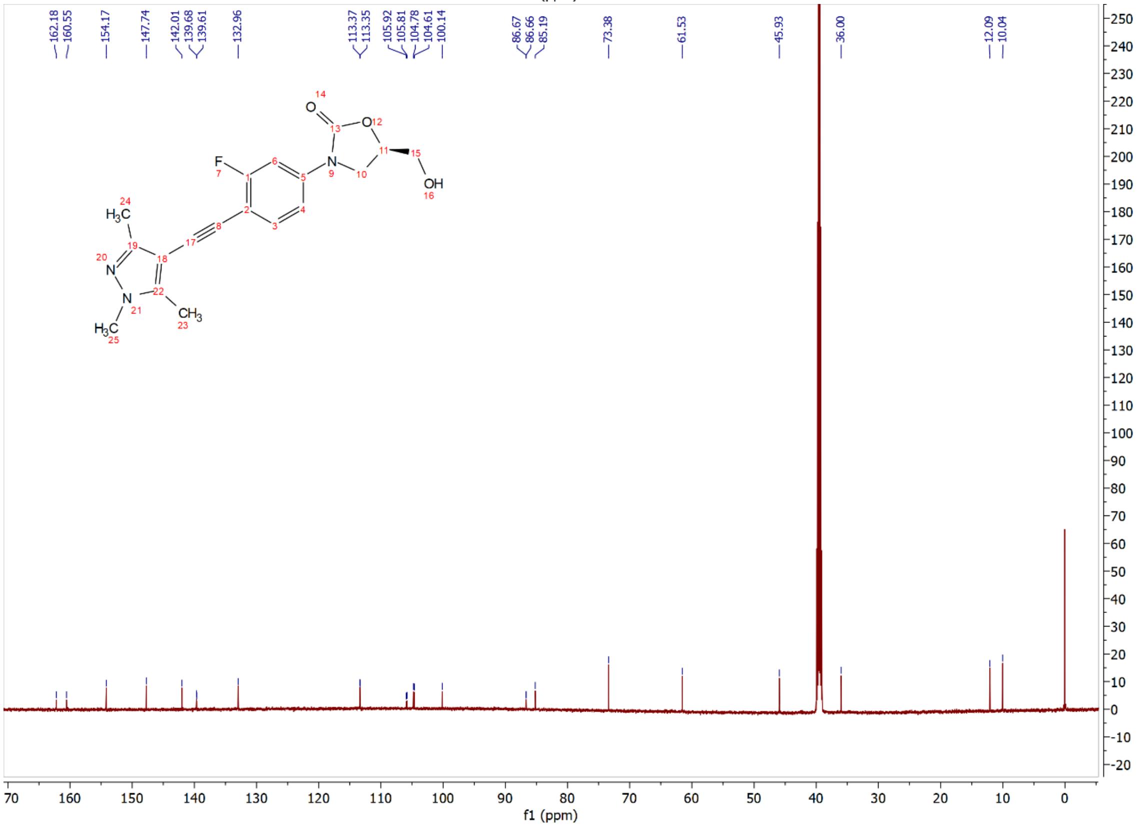

Compound **15d**: $^{1}$H and $^{13}$C NMR Spectrum

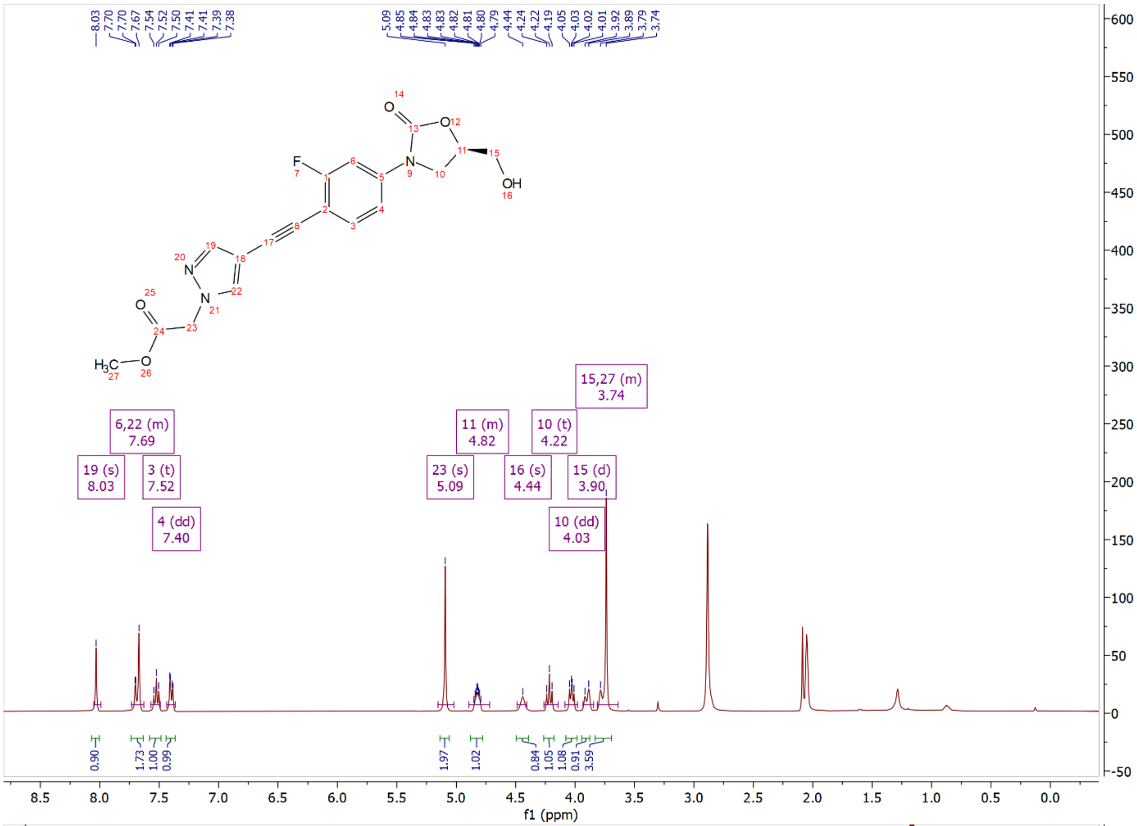

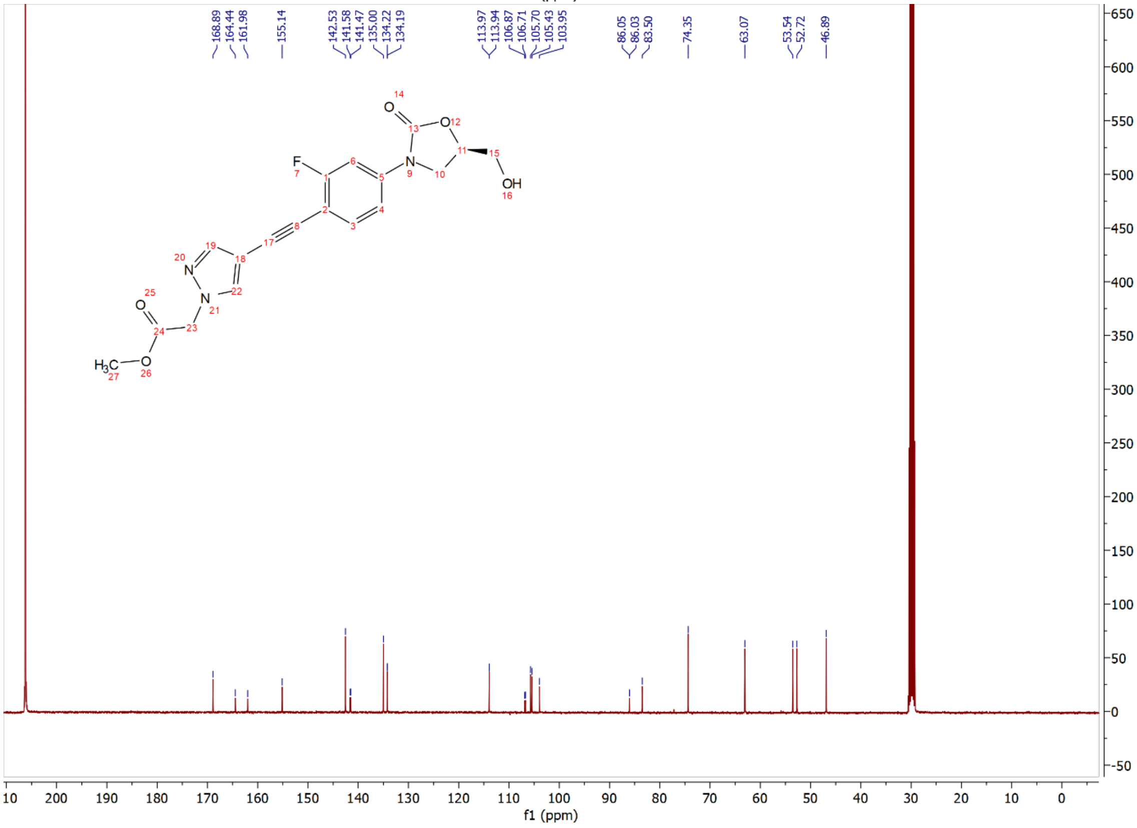

Compound **15e**: $^{1}$H and $^{13}$C NMR Spectrum

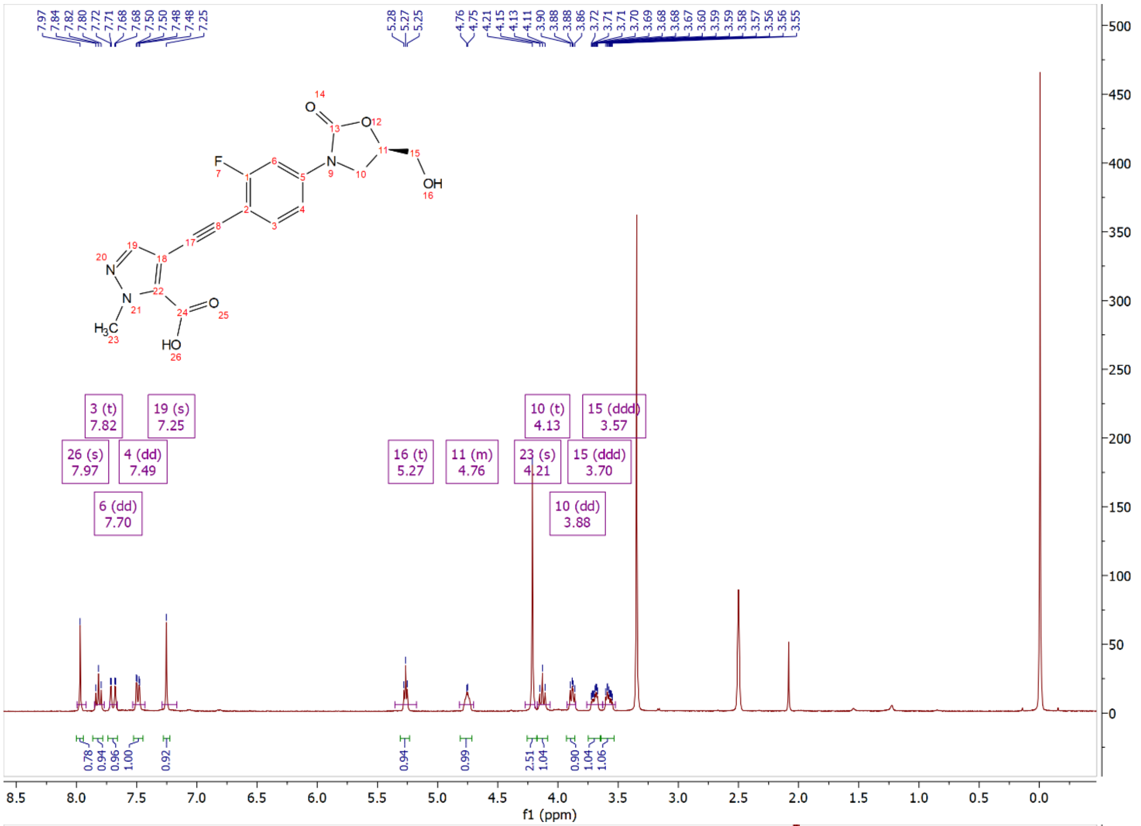

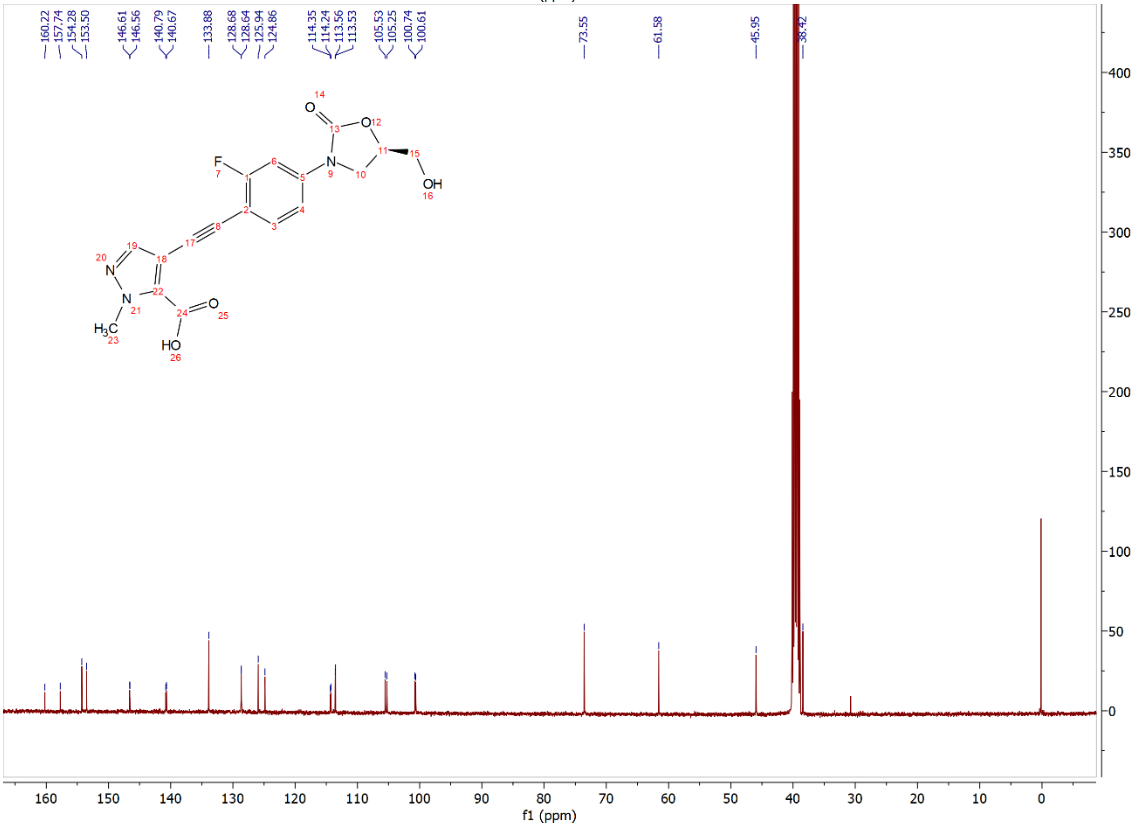

Compound **15f**: $^{1}$H and $^{13}$C NMR Spectrum

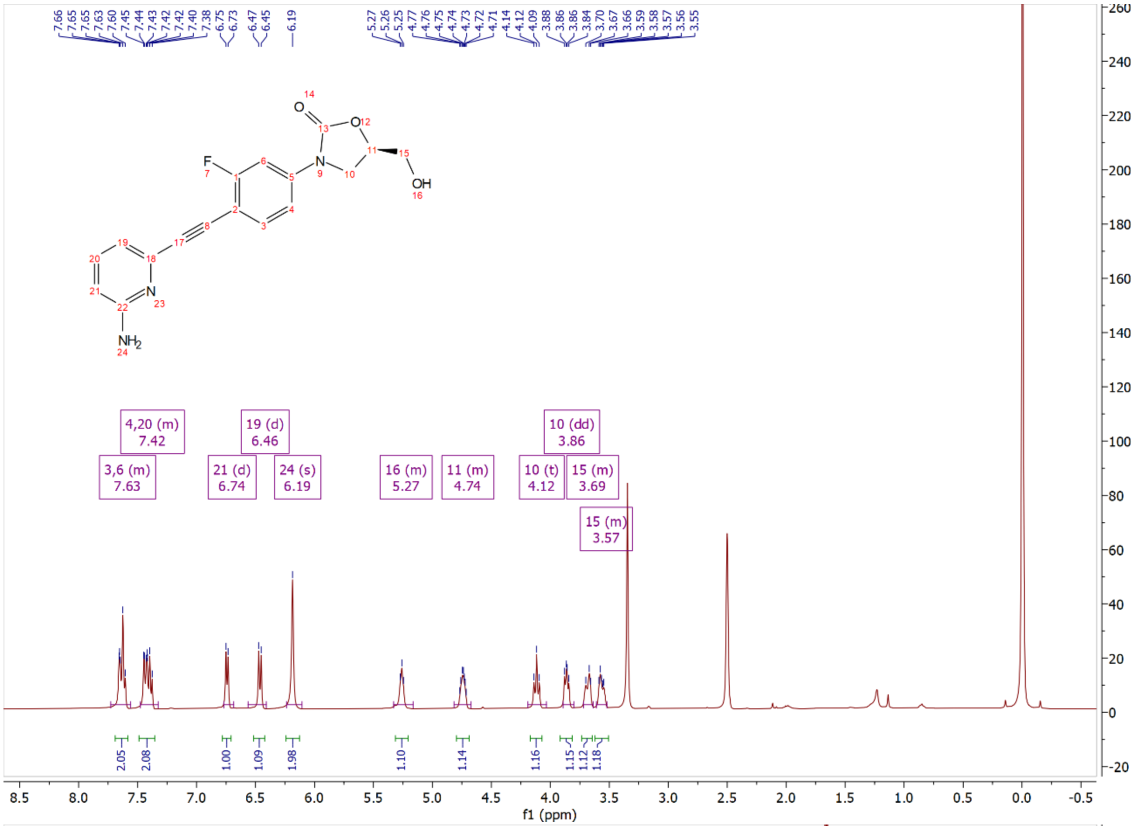

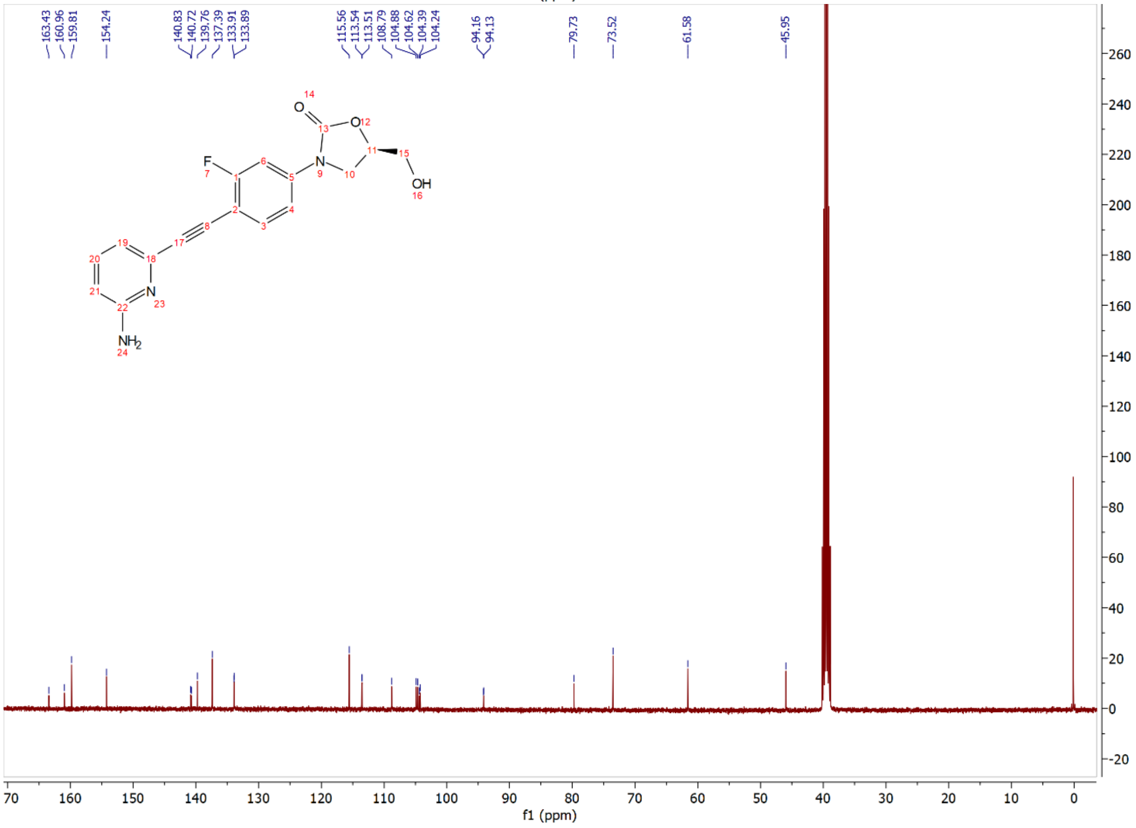

Compound **15g**: $^{1}$H and $^{13}$C NMR Spectrum

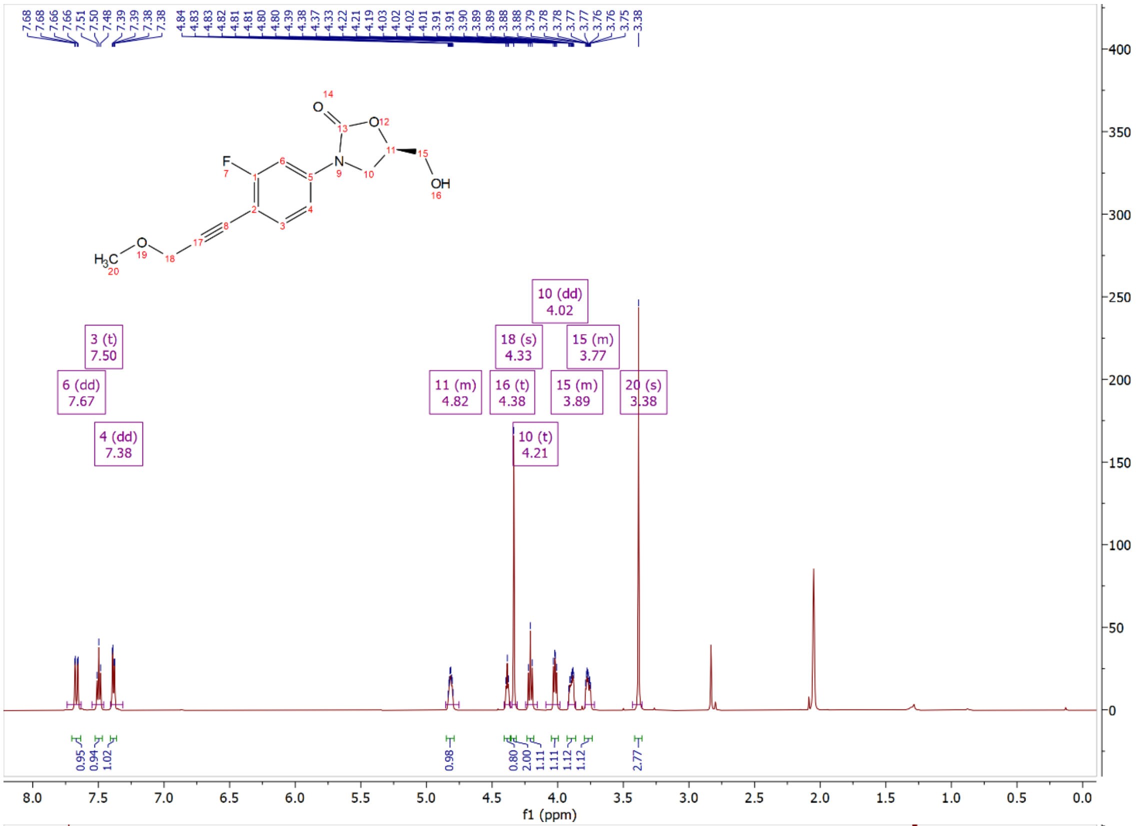

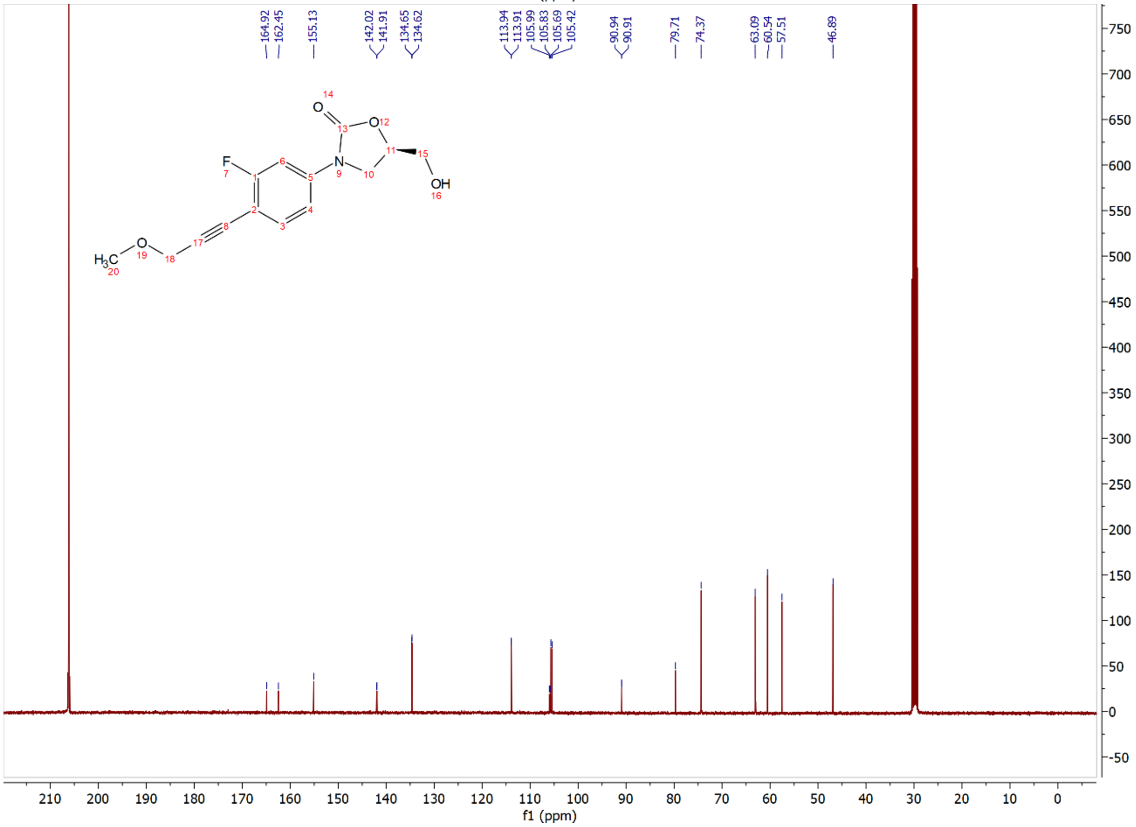

Compound **15h**: $^{1}$H and $^{13}$C NMR Spectrum

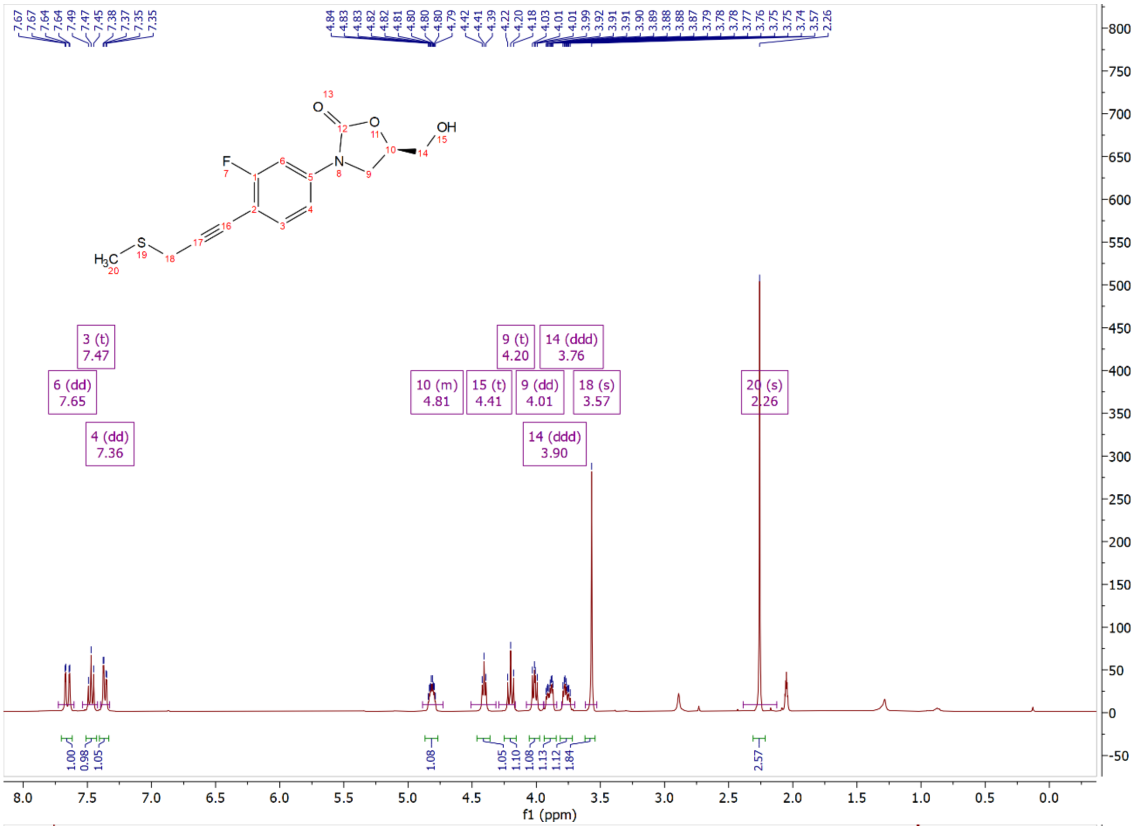


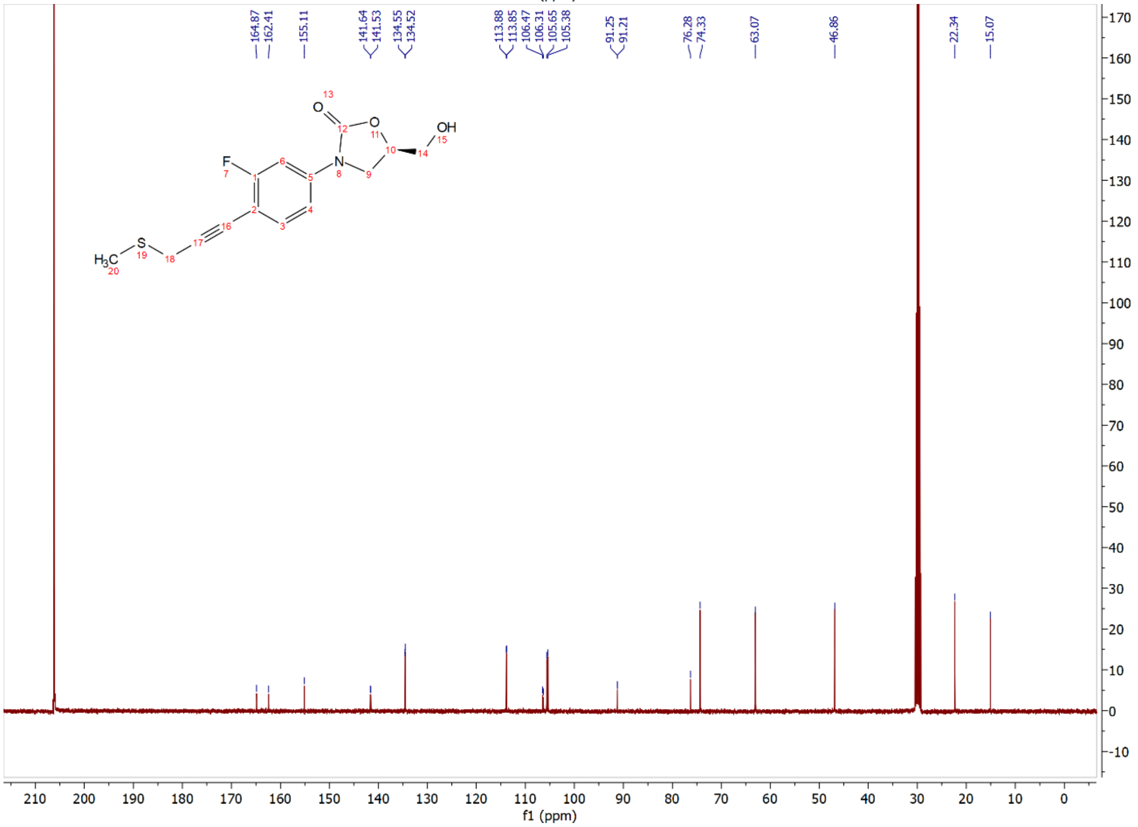

Compound **15i**: $^{1}$H and $^{13}$C NMR Spectrum

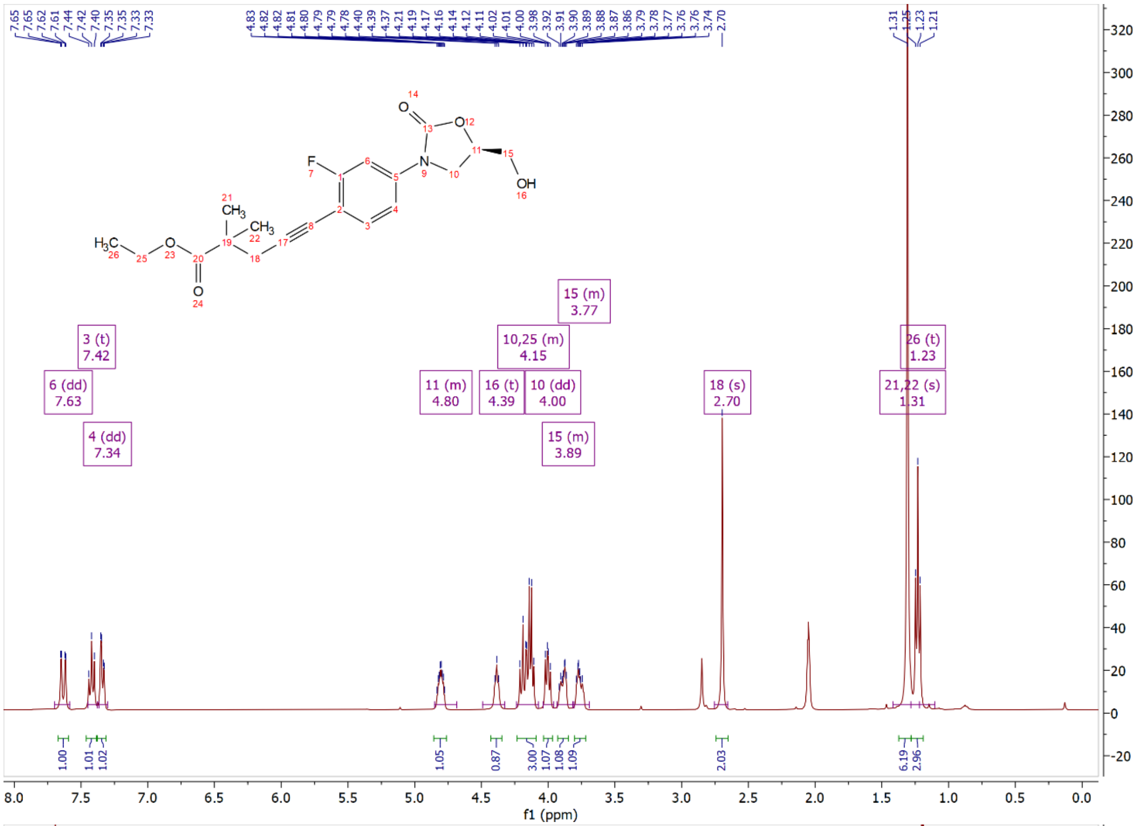

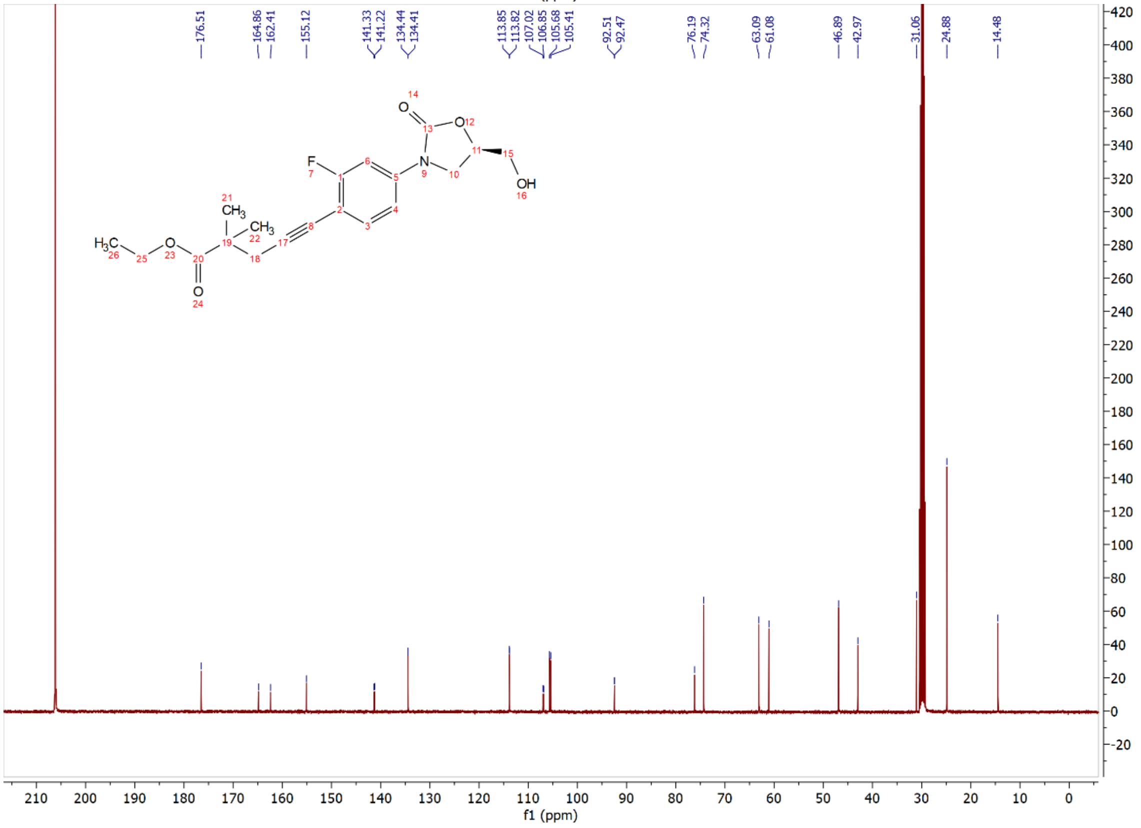

Compound **15j**: $^{1}H$ and $^{13}C$ NMR Spectrum

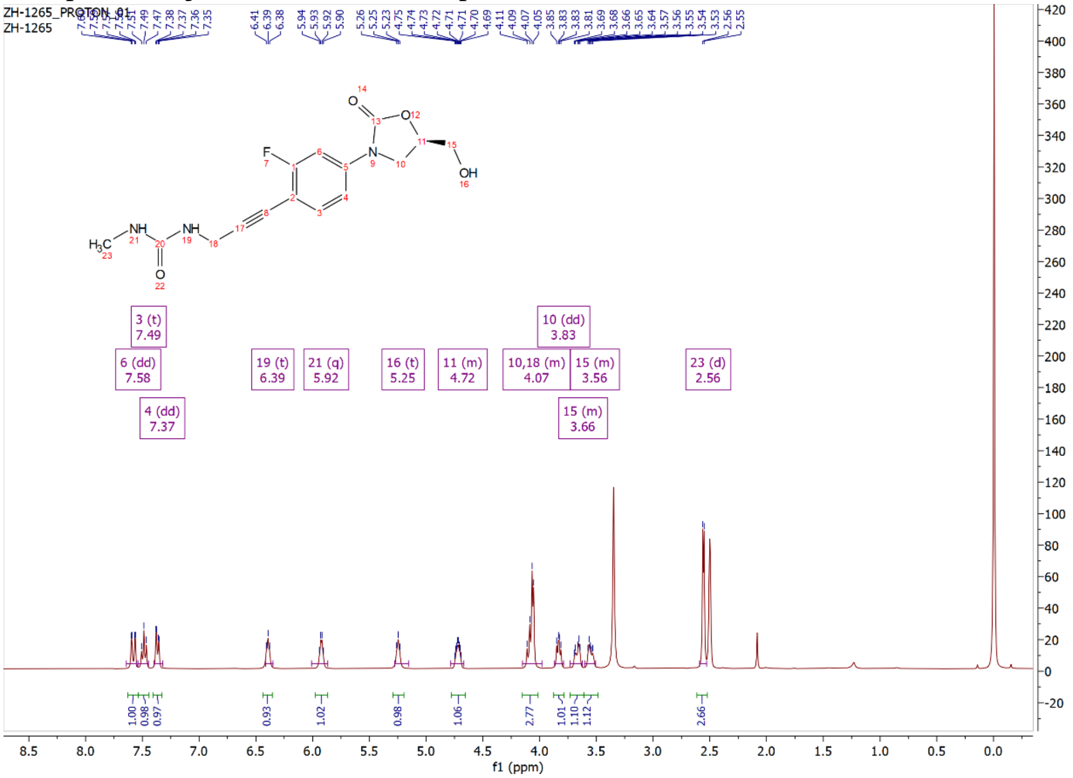

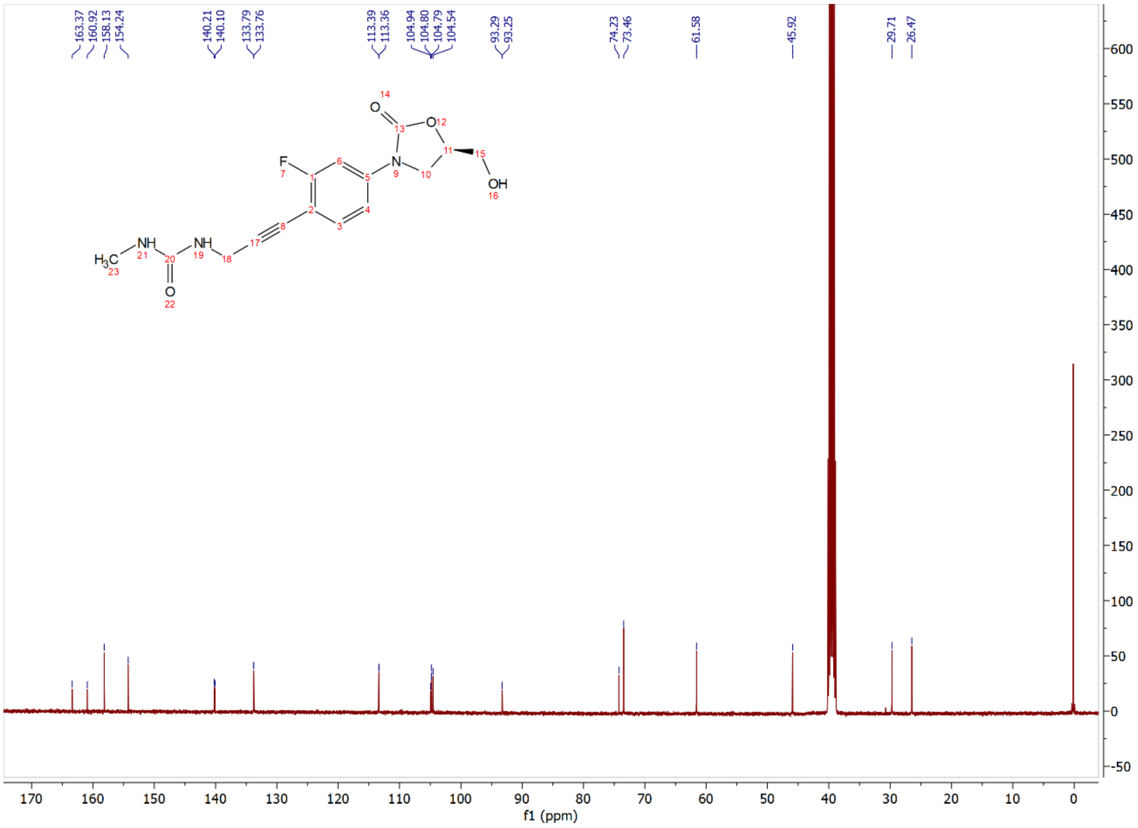

Compound **15k**: $^{1}$H and $^{13}$C NMR Spectrum

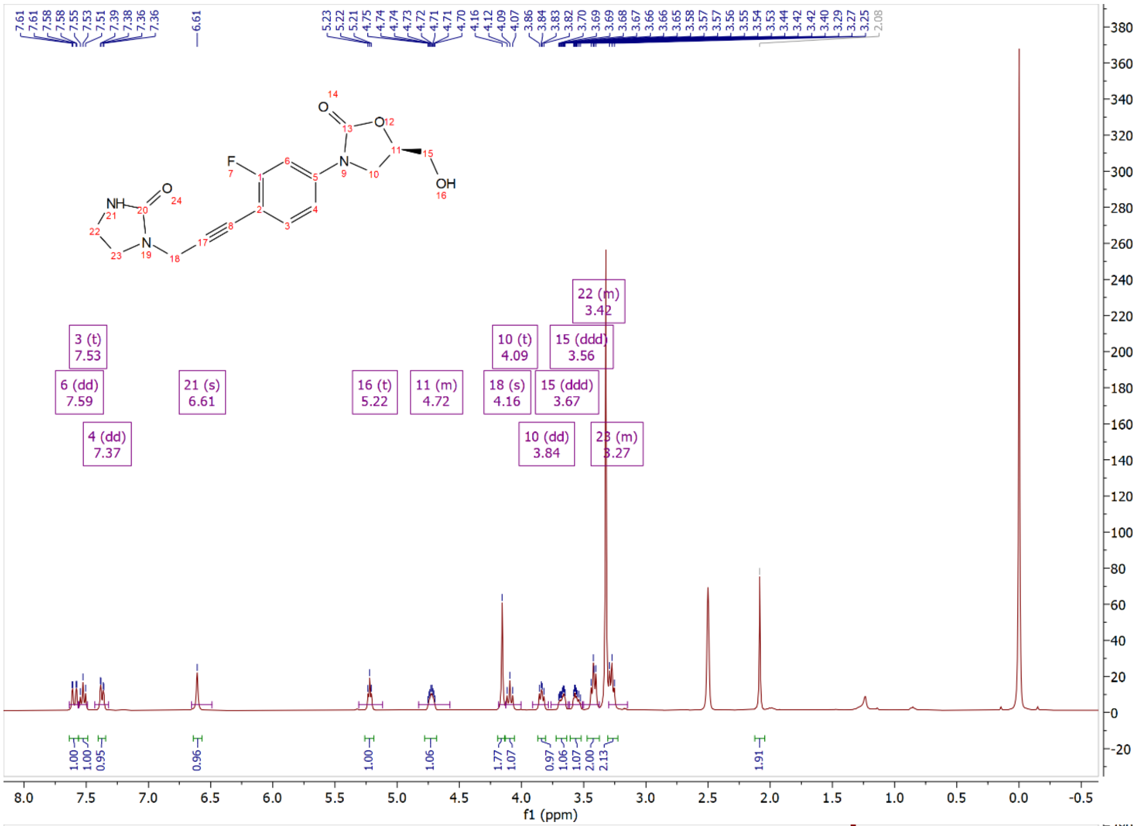

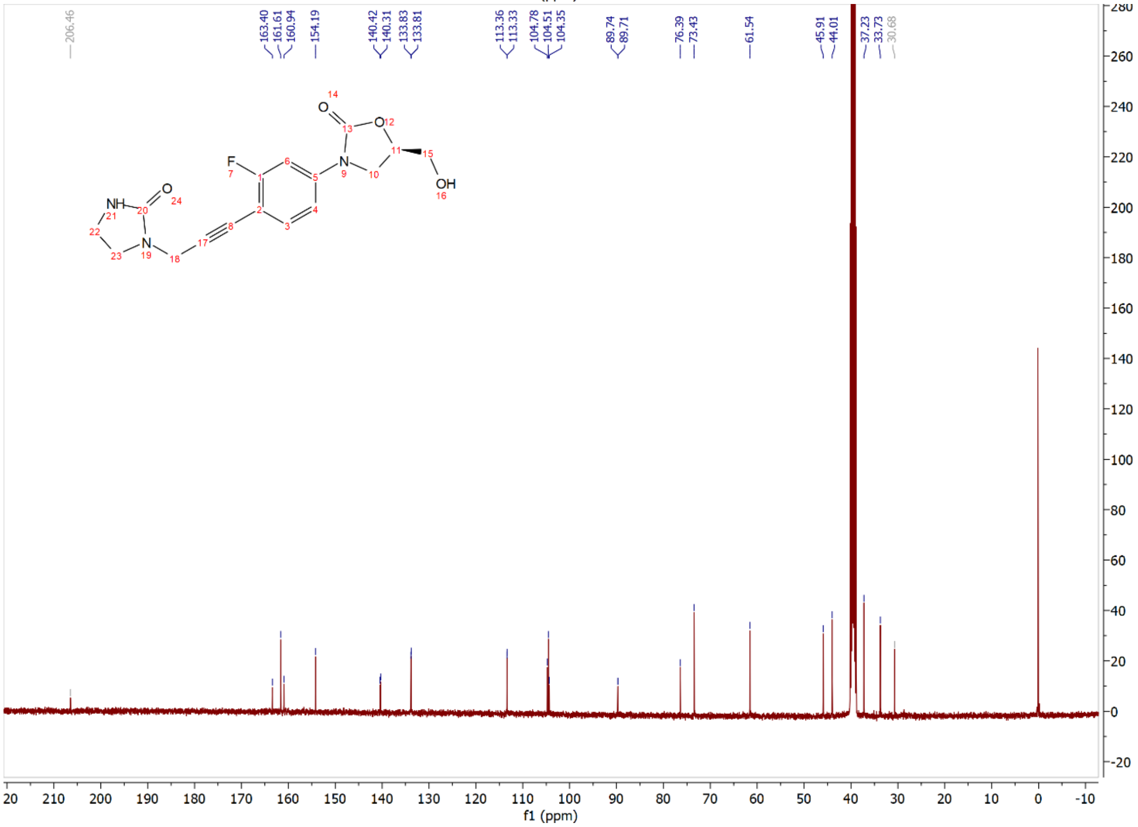

Compound **15l**: $^{1}$H and $^{13}$C NMR Spectrum

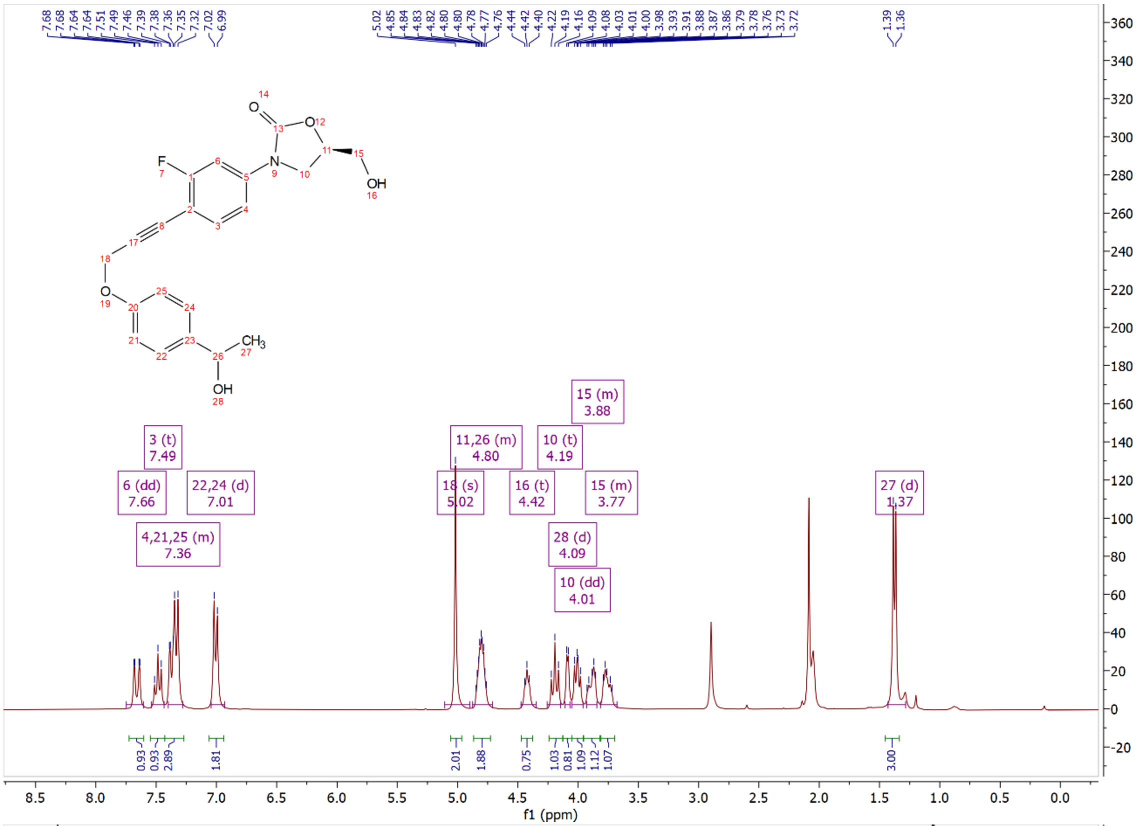

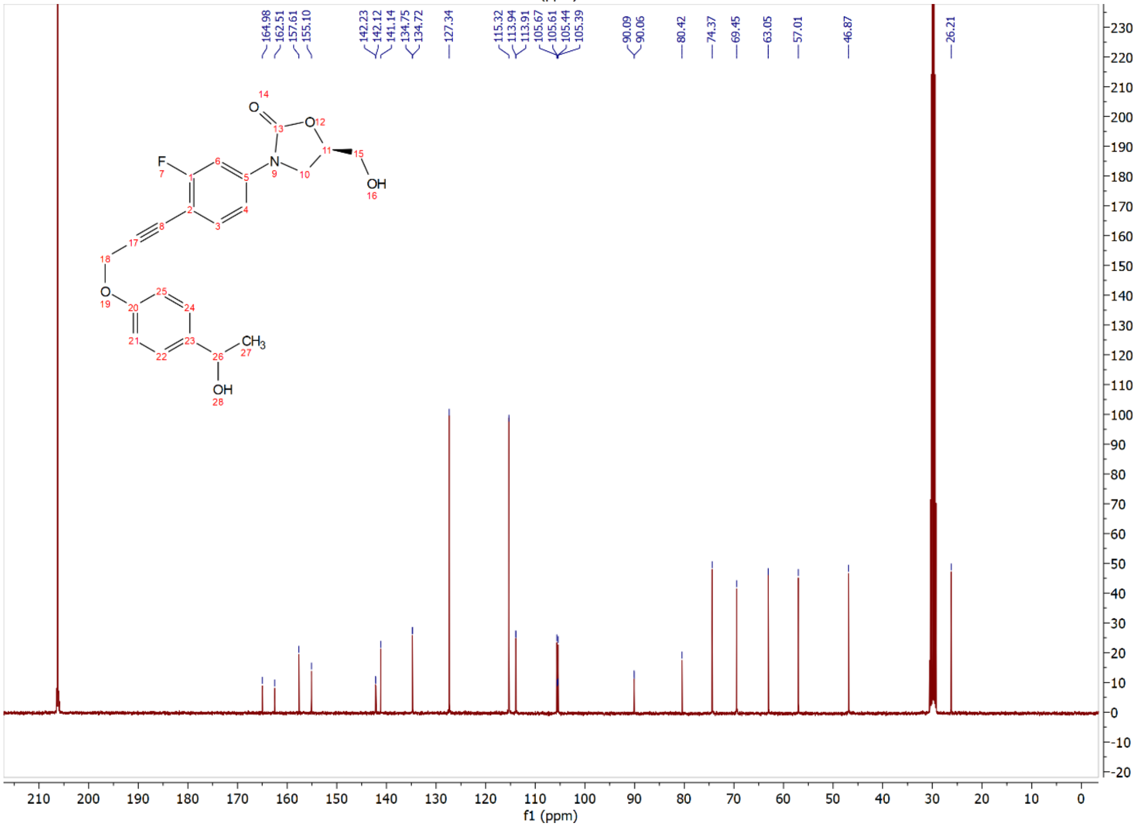

Compound **15m**: $^{1}$H and $^{13}$C NMR Spectrum

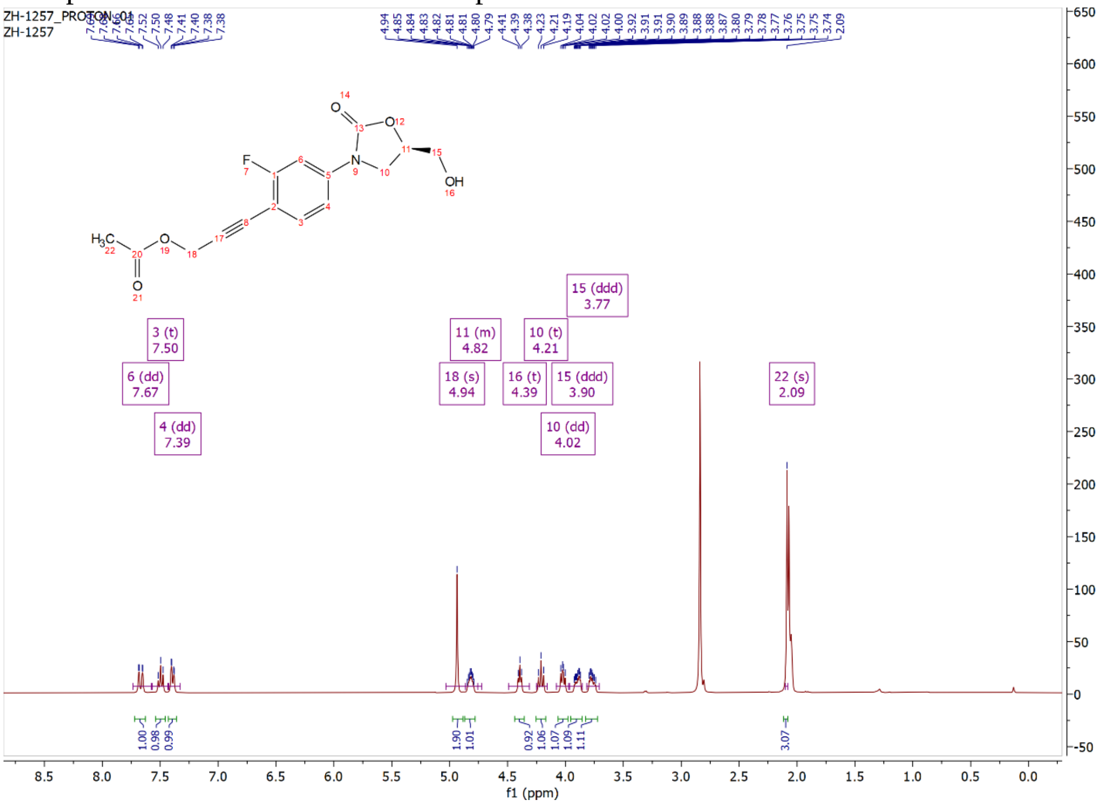

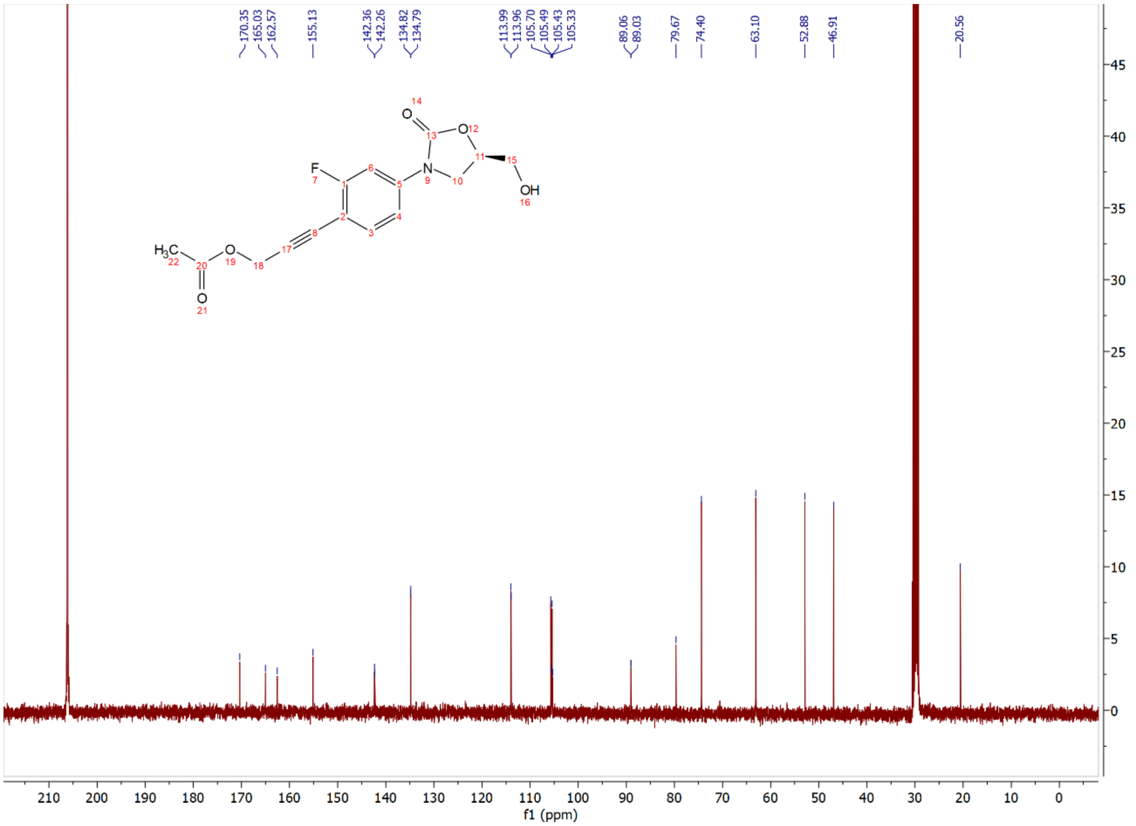

Compound **15ma** $^{1}$H and $^{13}$C NMR Spectrum

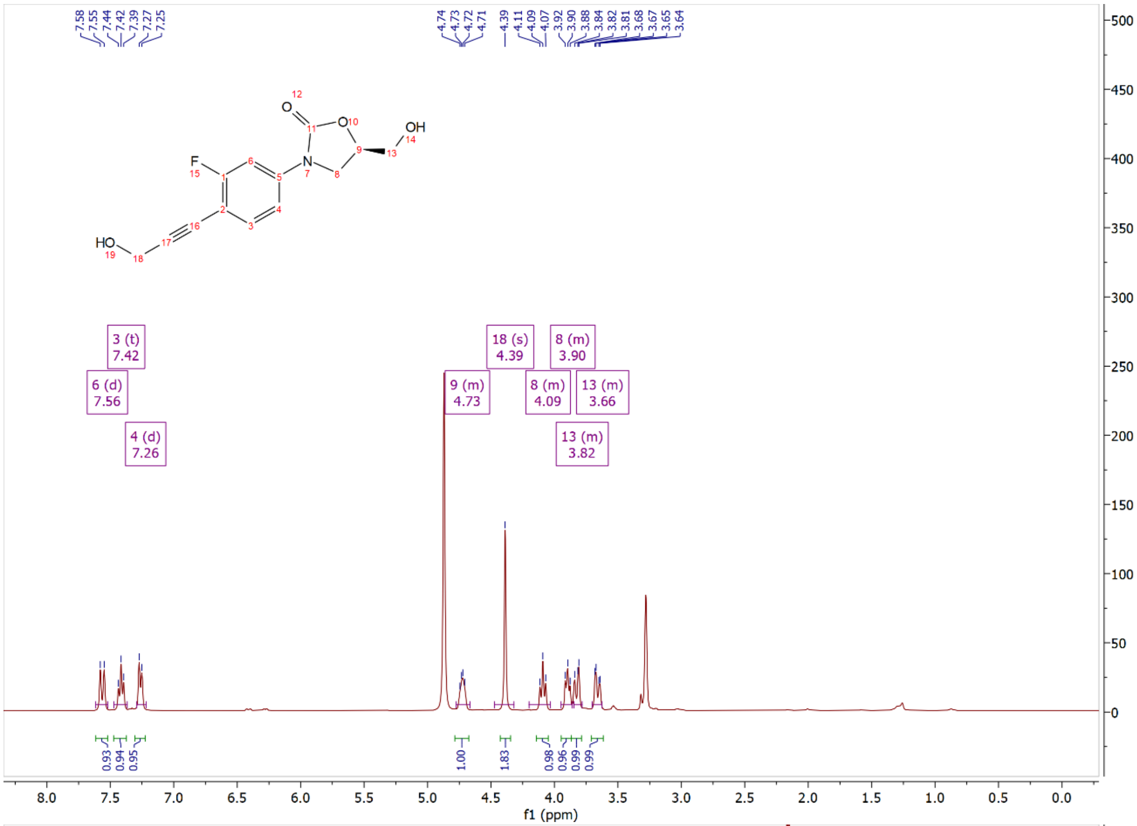


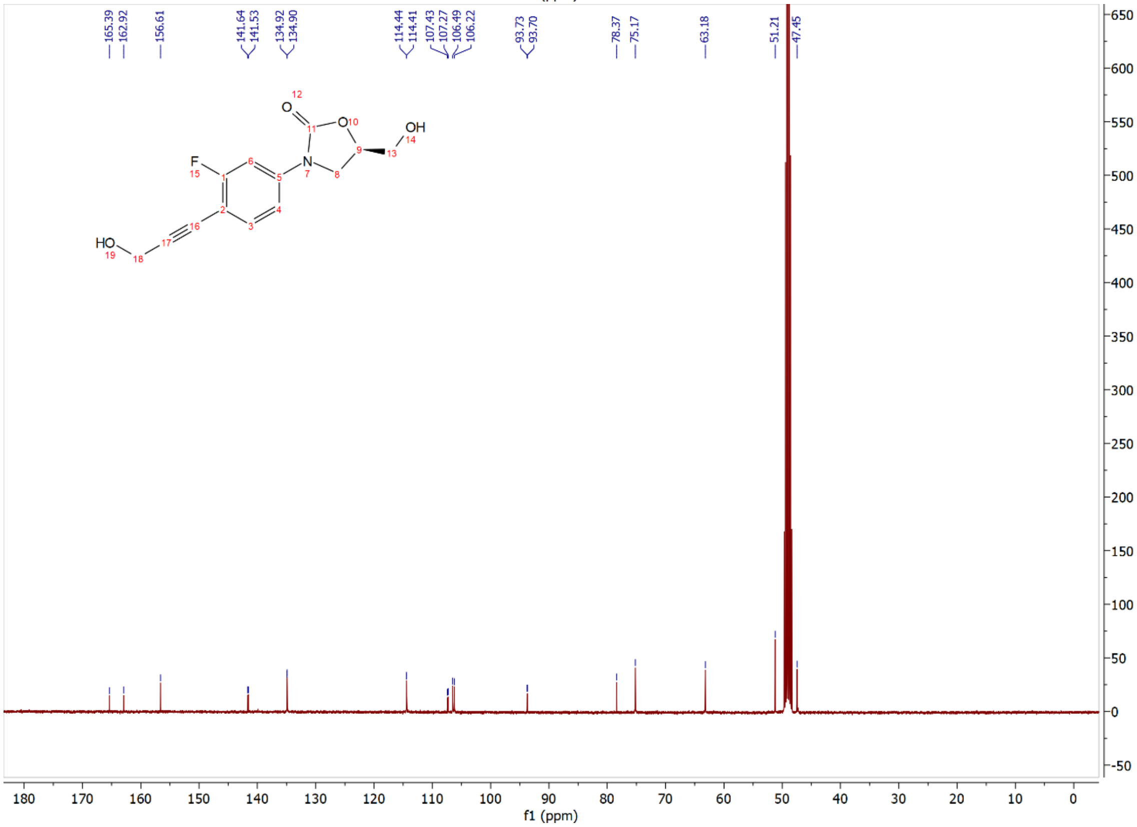

Compound **15n**: $^{1}$H and $^{13}$C NMR Spectrum

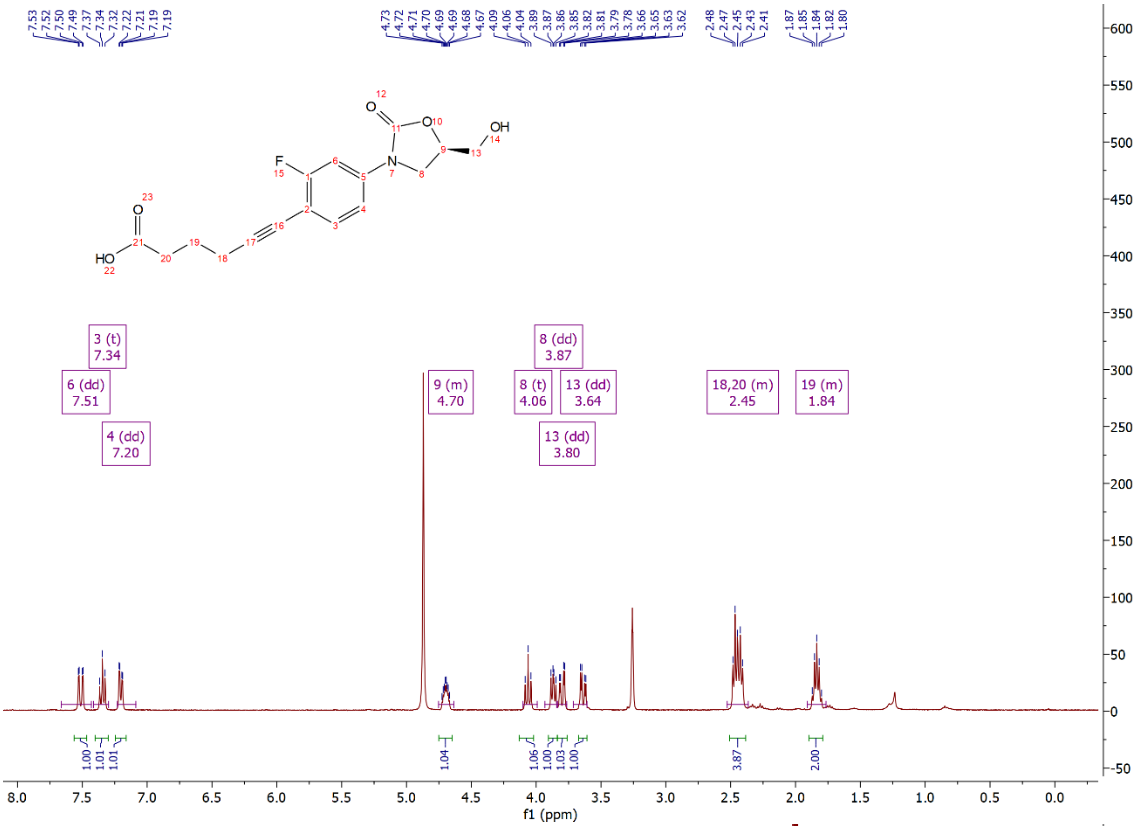

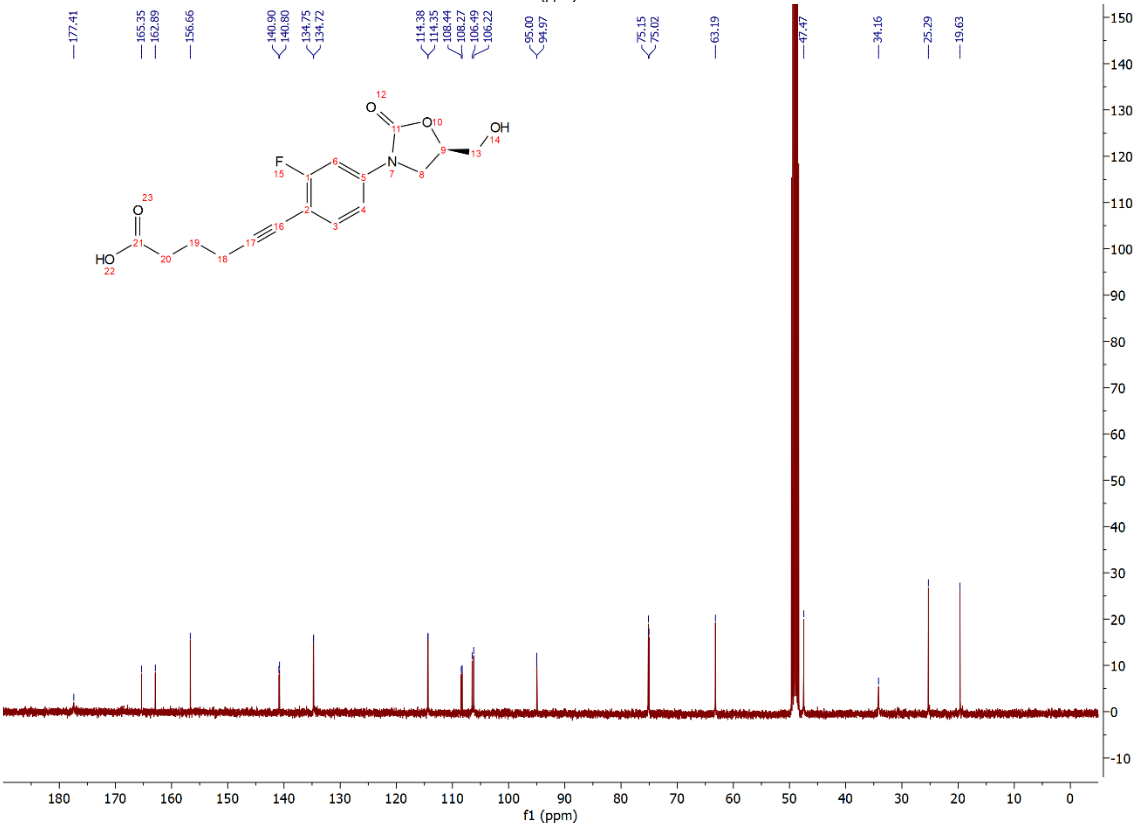

Compound **15o**: $^{1}H$ and $^{13}C$ NMR Spectrum

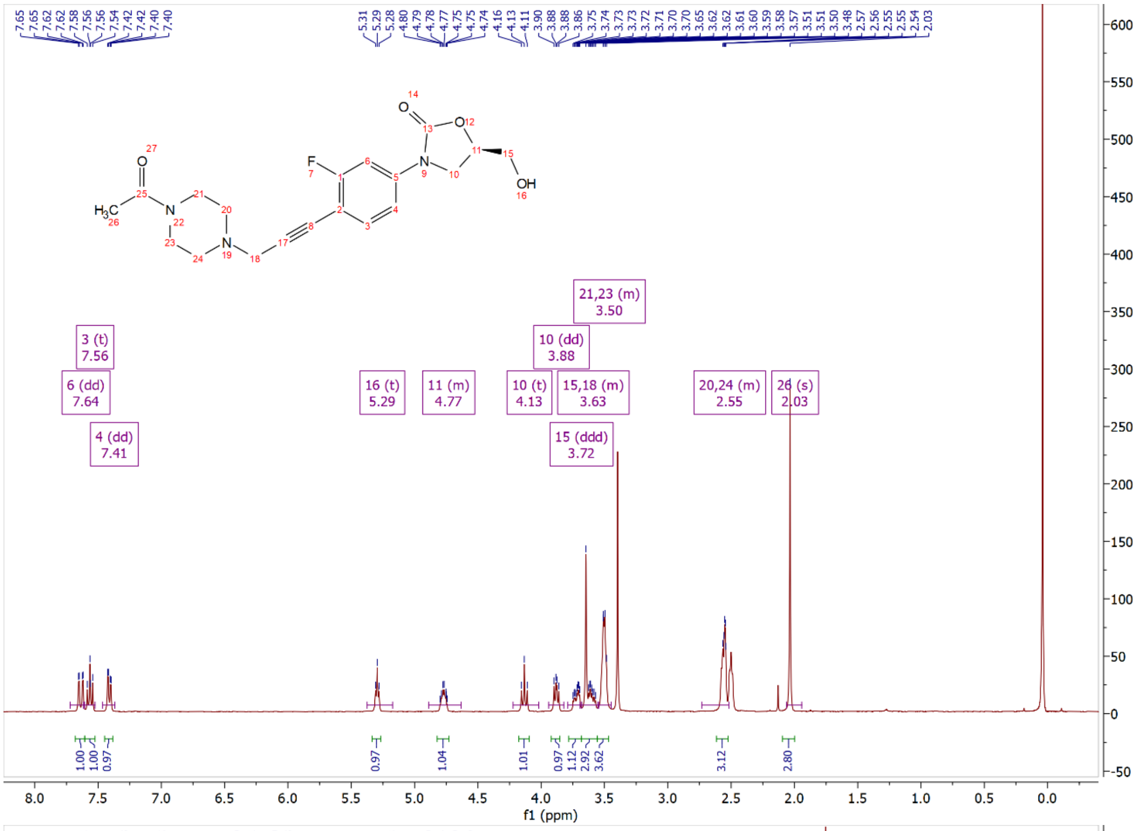

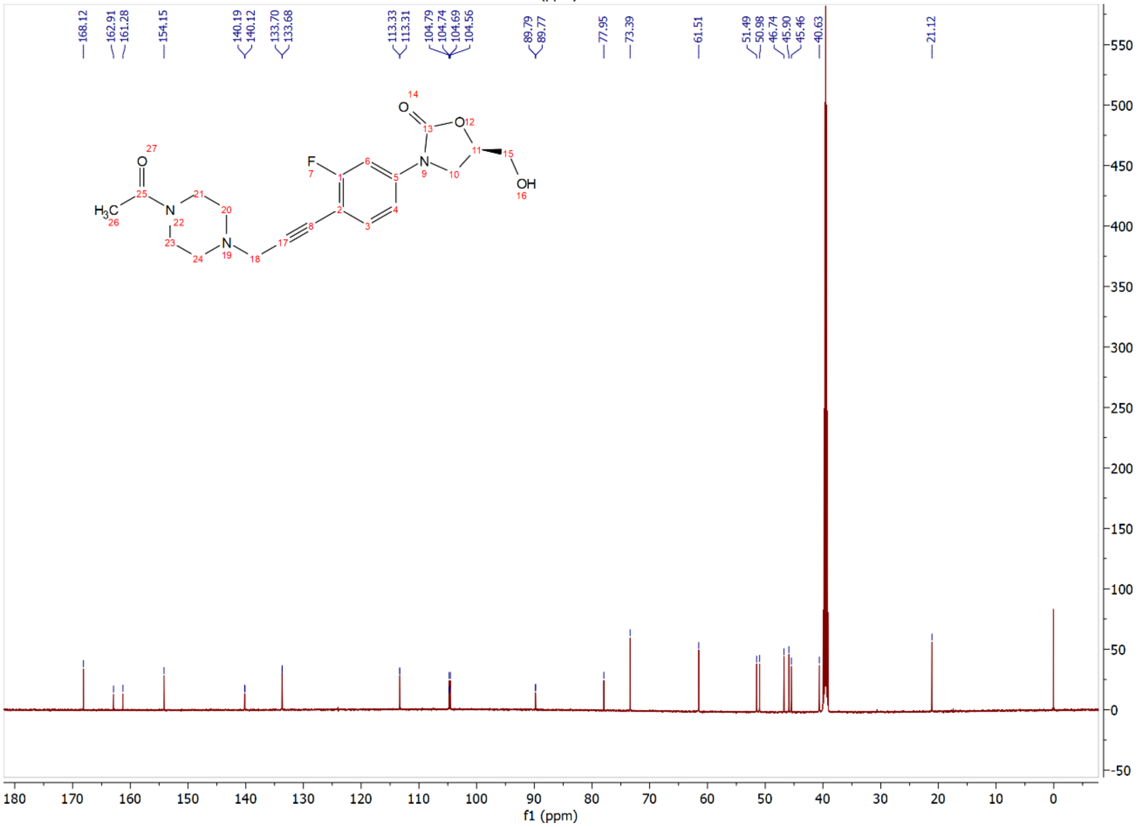

Compound **15p**: $^{1}H$ and $^{13}C$ NMR Spectrum

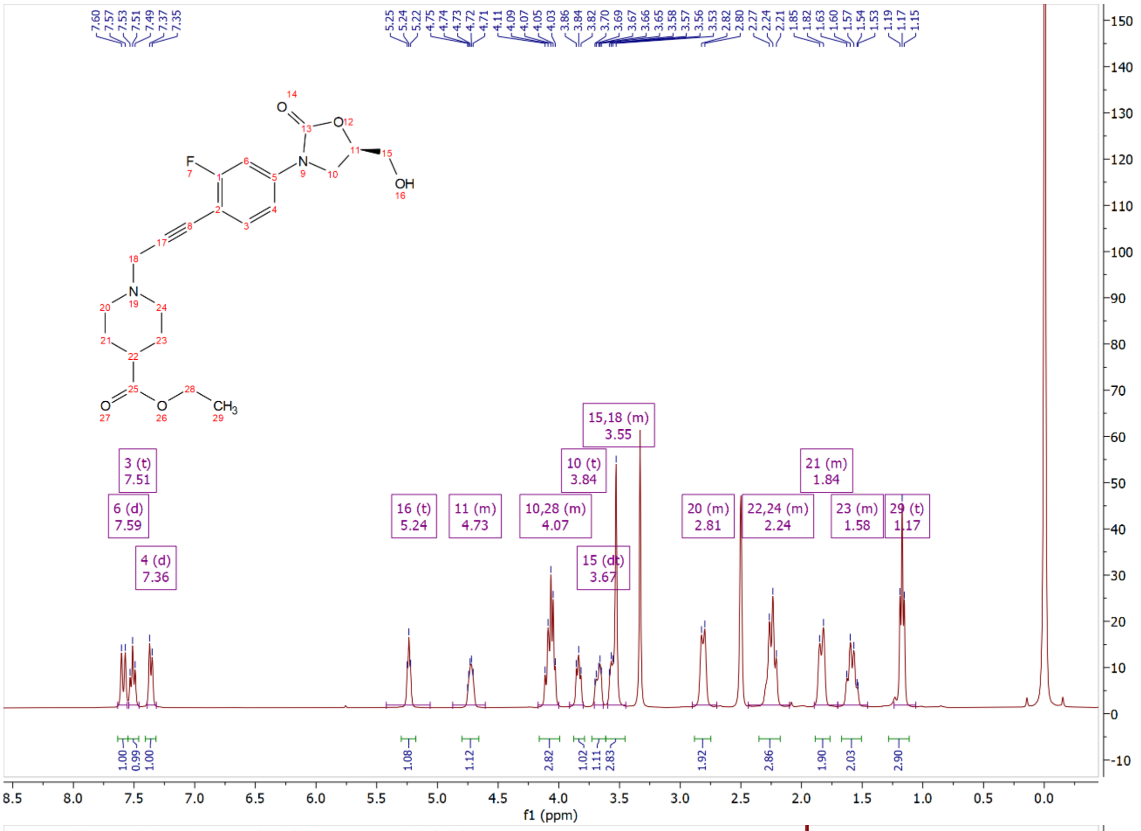

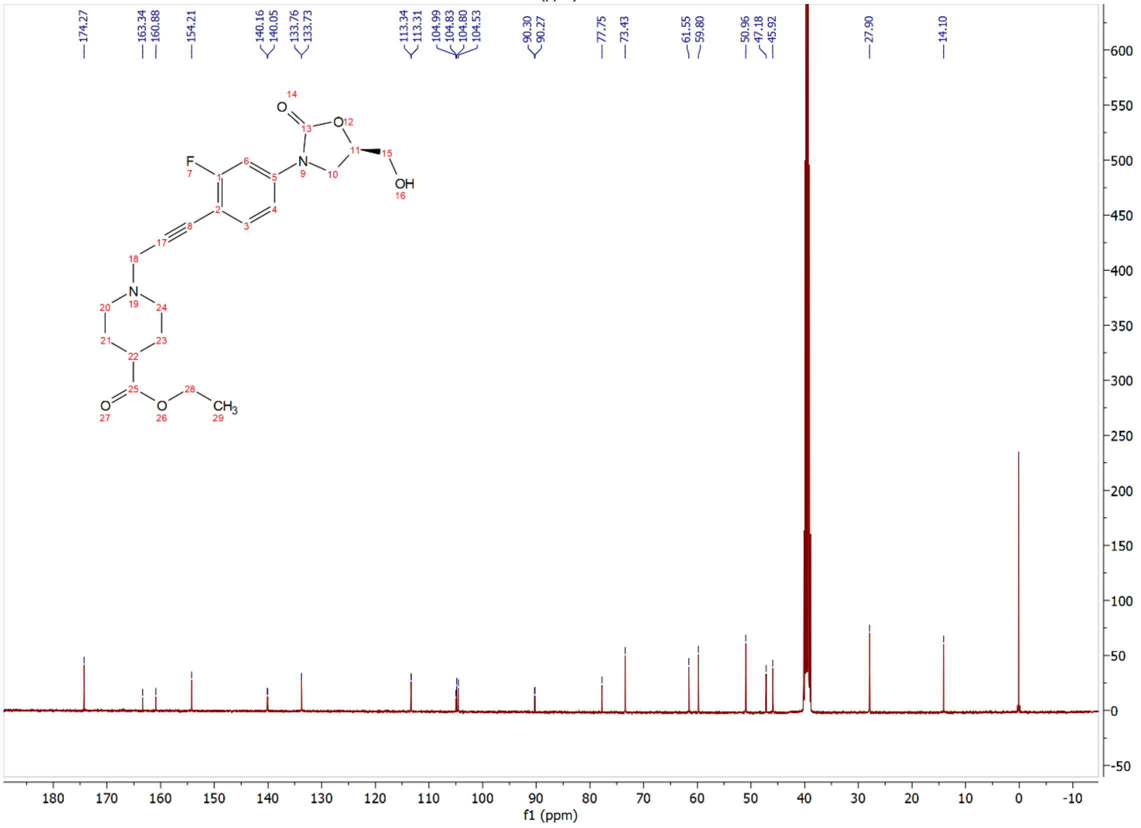

Compound **15q**: $^{1}$H and $^{13}$C NMR Spectrum

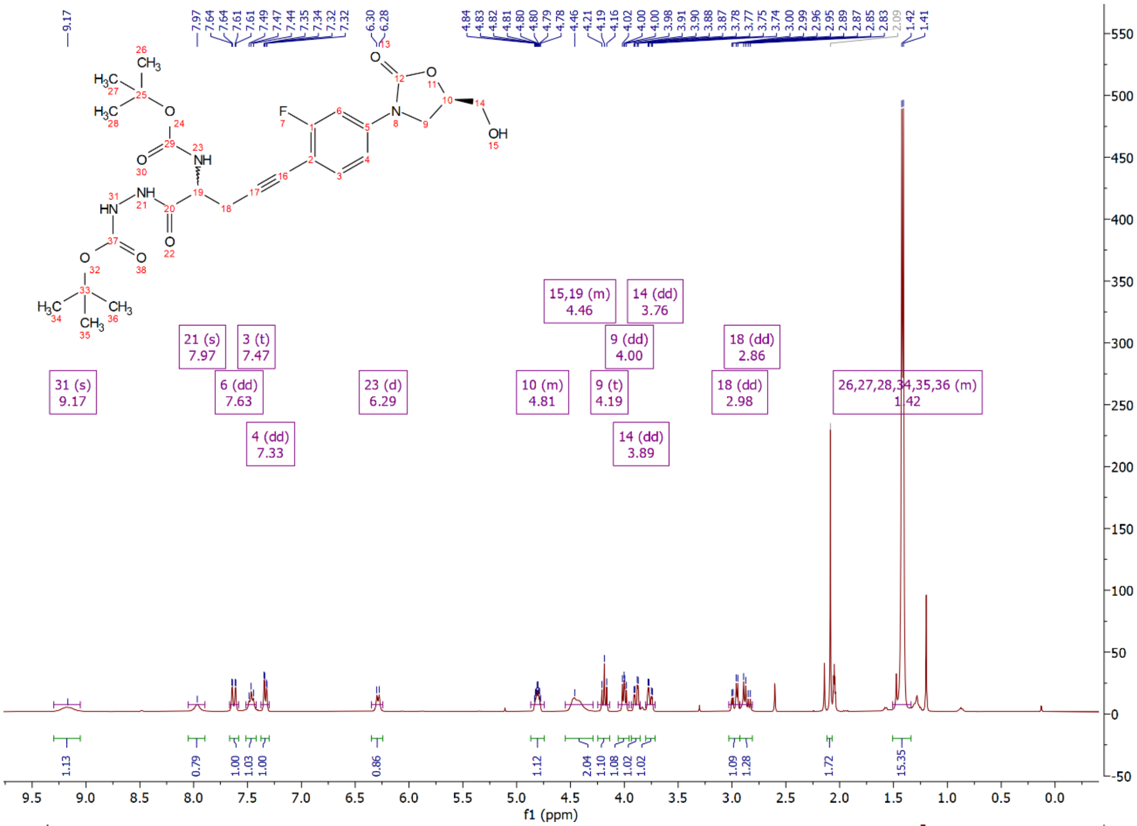

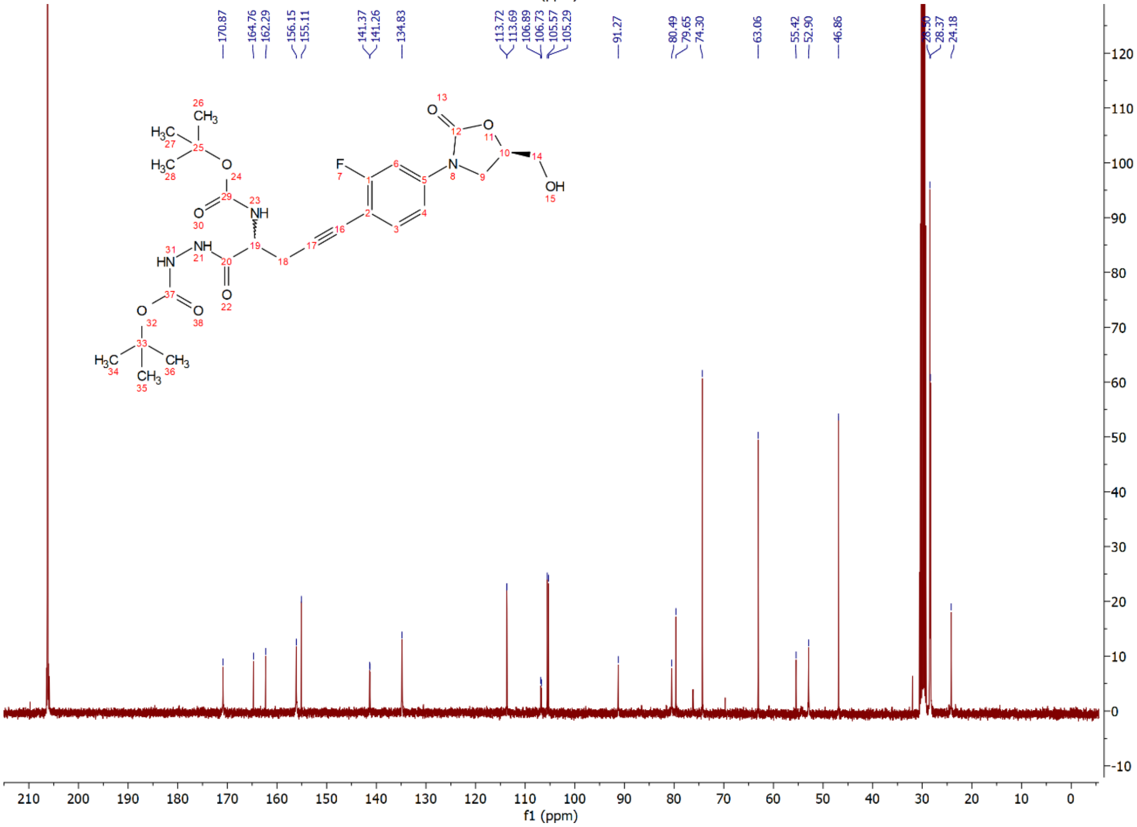

Compound **15r**: $^{1}$H and $^{13}$C NMR Spectrum

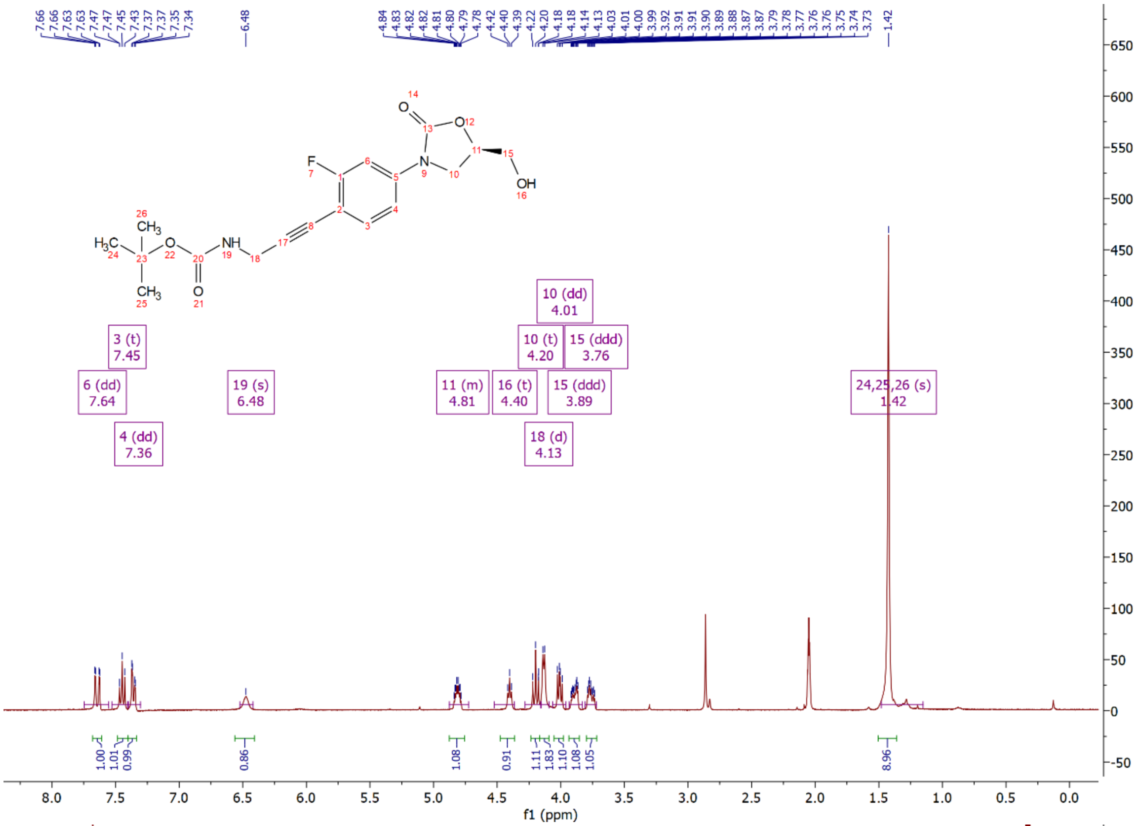

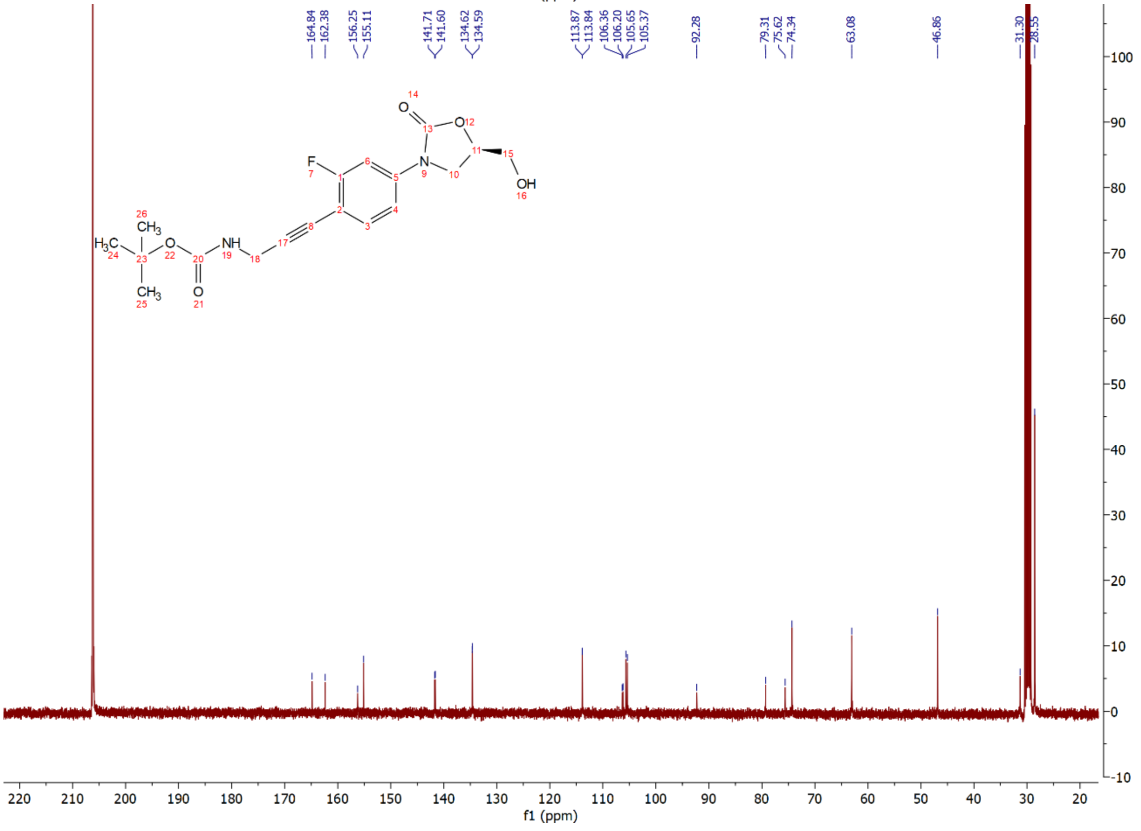

Compound **15ra**: $^{1}H$ and $^{13}C$ NMR Spectrum

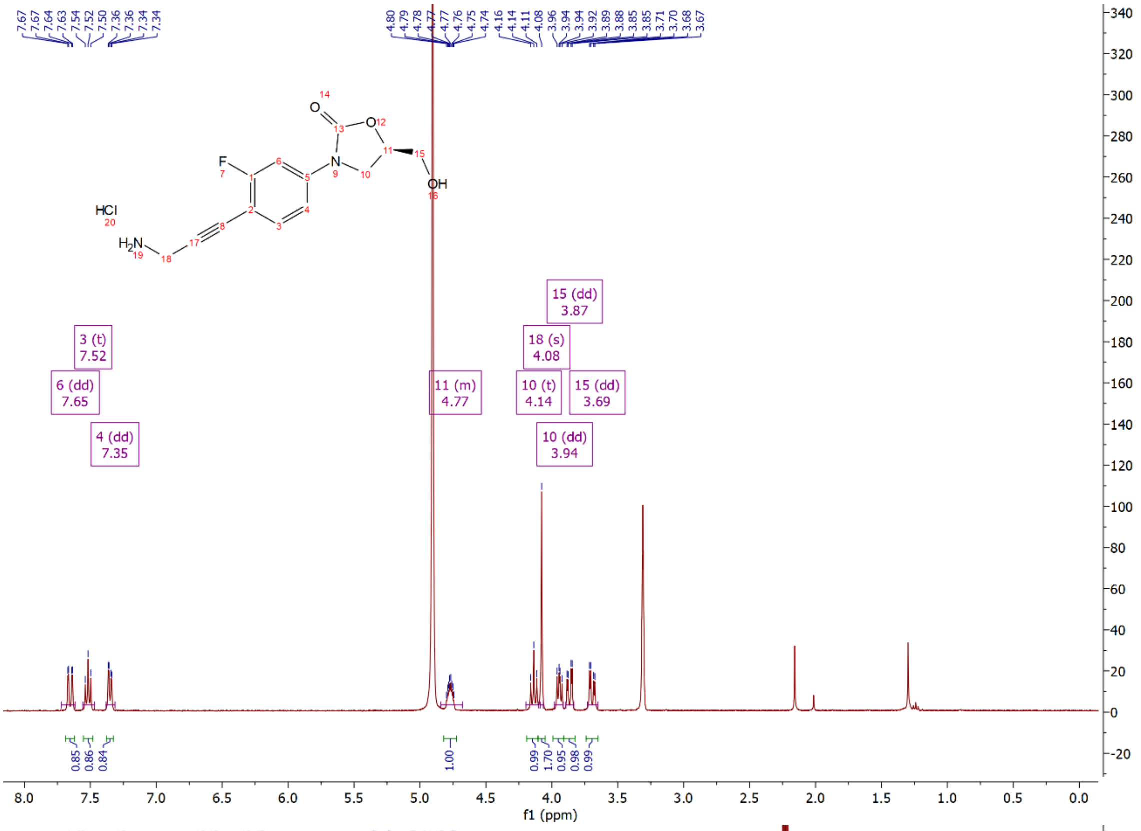

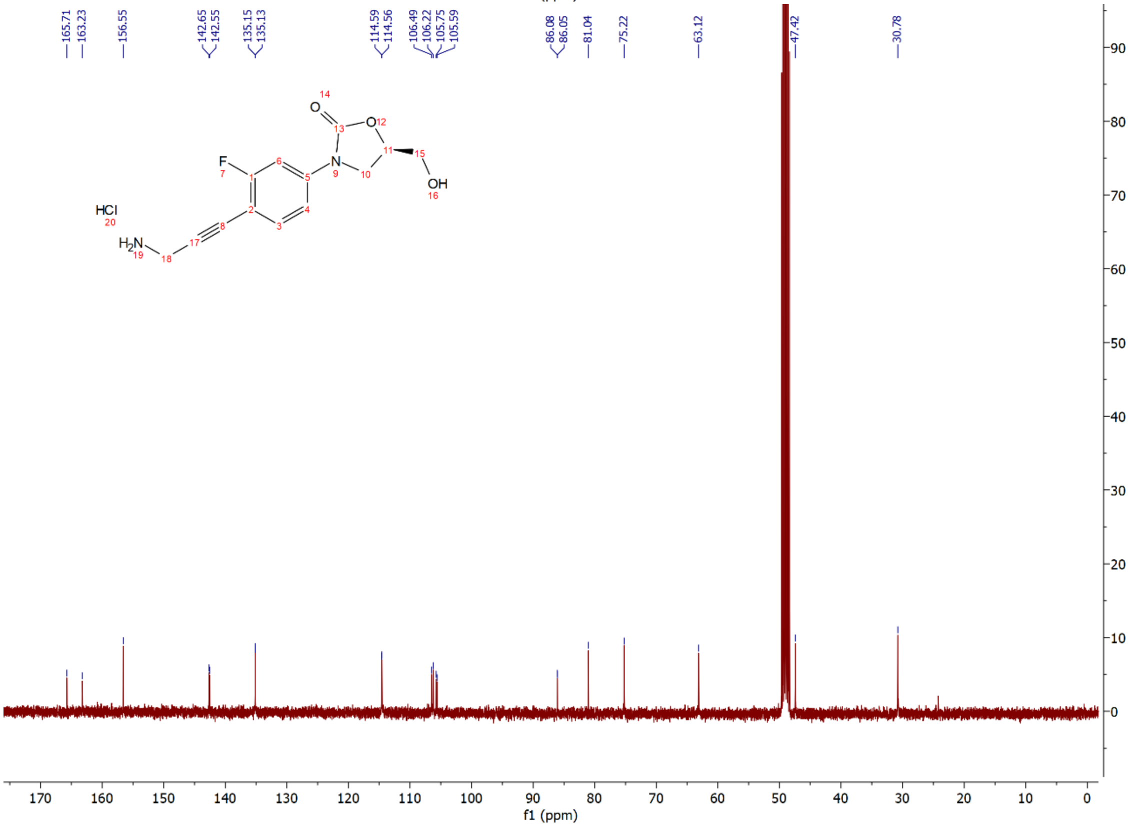

Compound **15s**: $^{1}$H and $^{13}$C NMR Spectrum

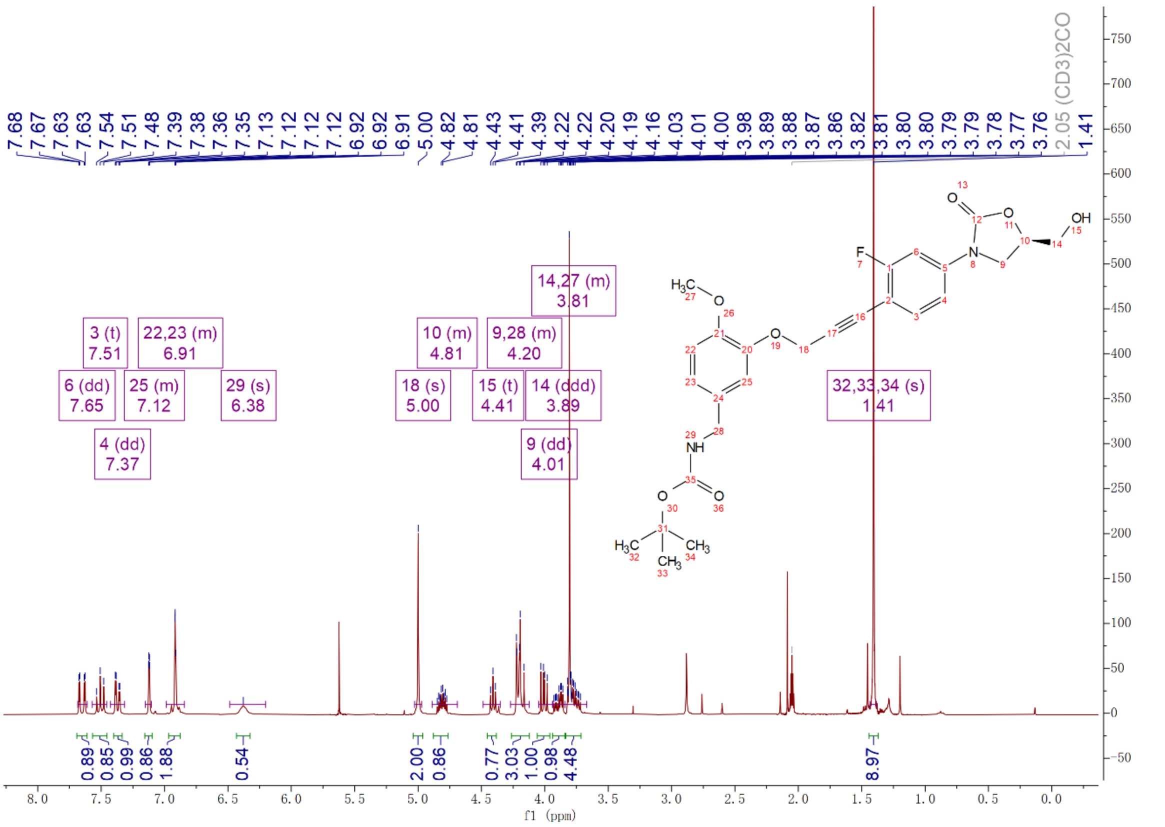

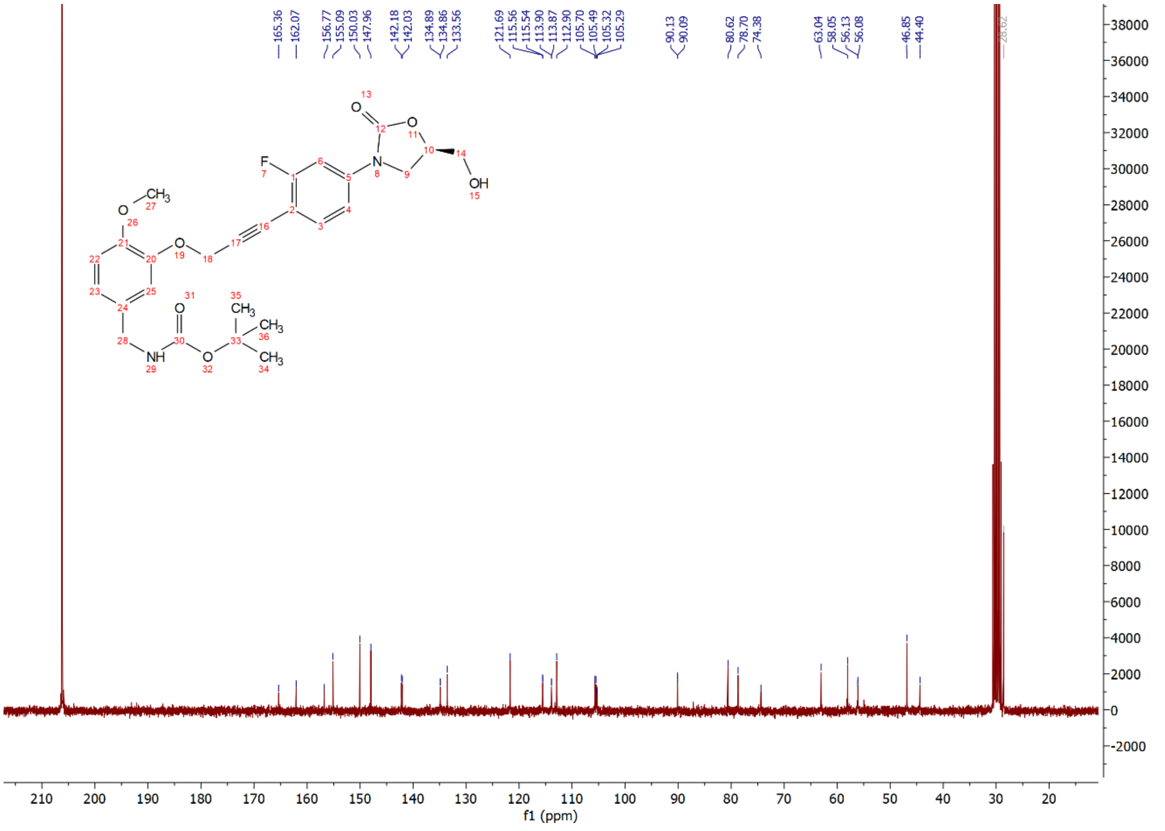

Compound **15sa**: $^{1}$H and $^{13}$C NMR Spectrum

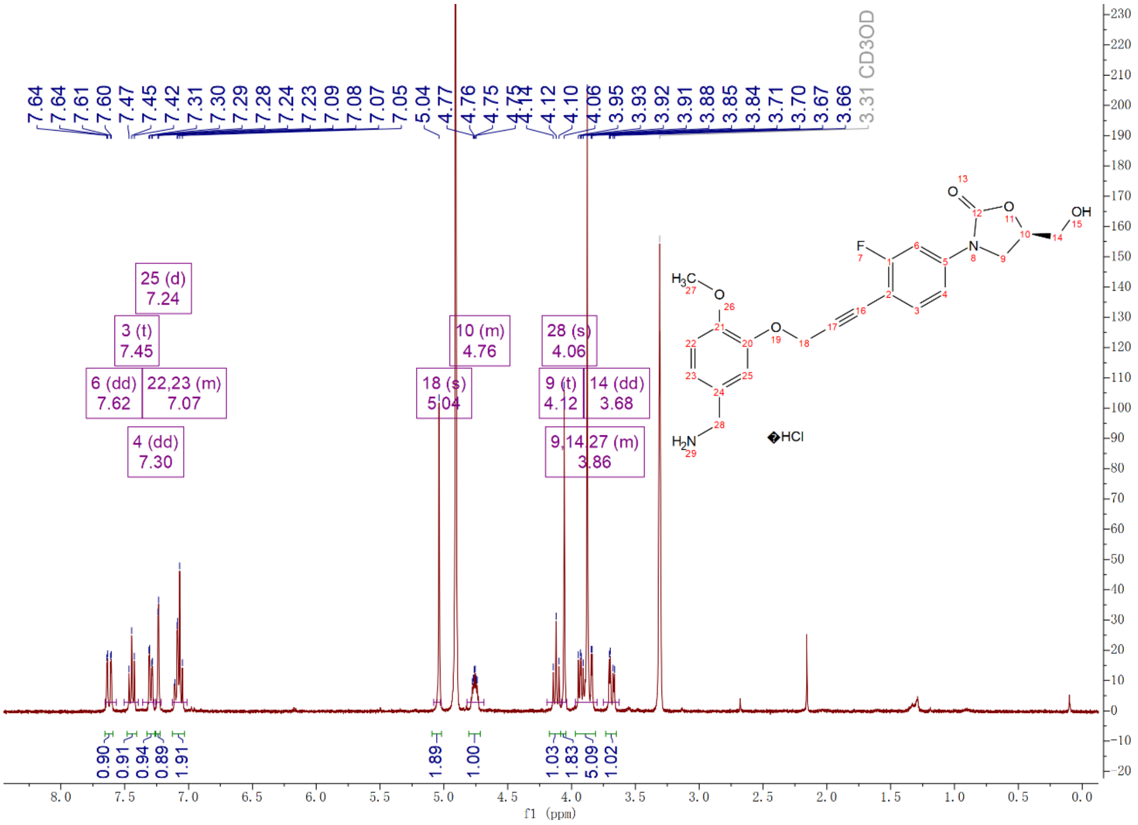

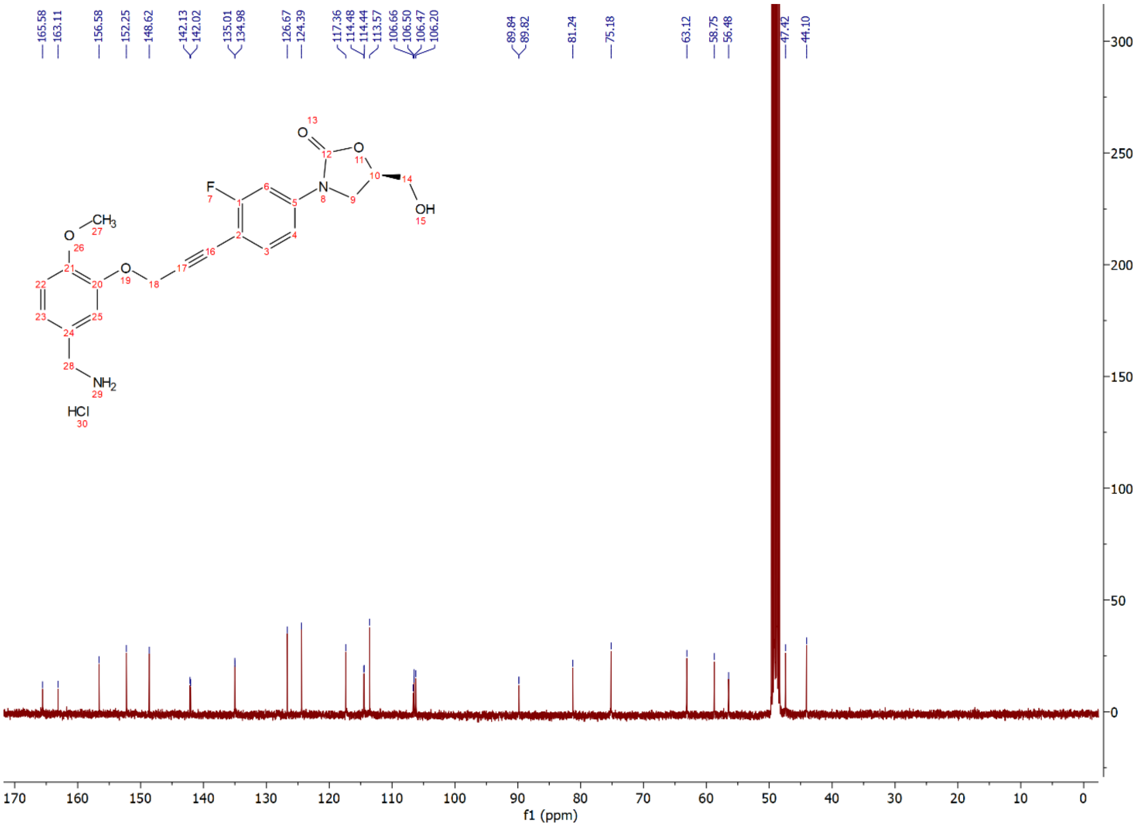

Compound **15t**: $^{1}H$ and $^{13}C$ NMR Spectrum

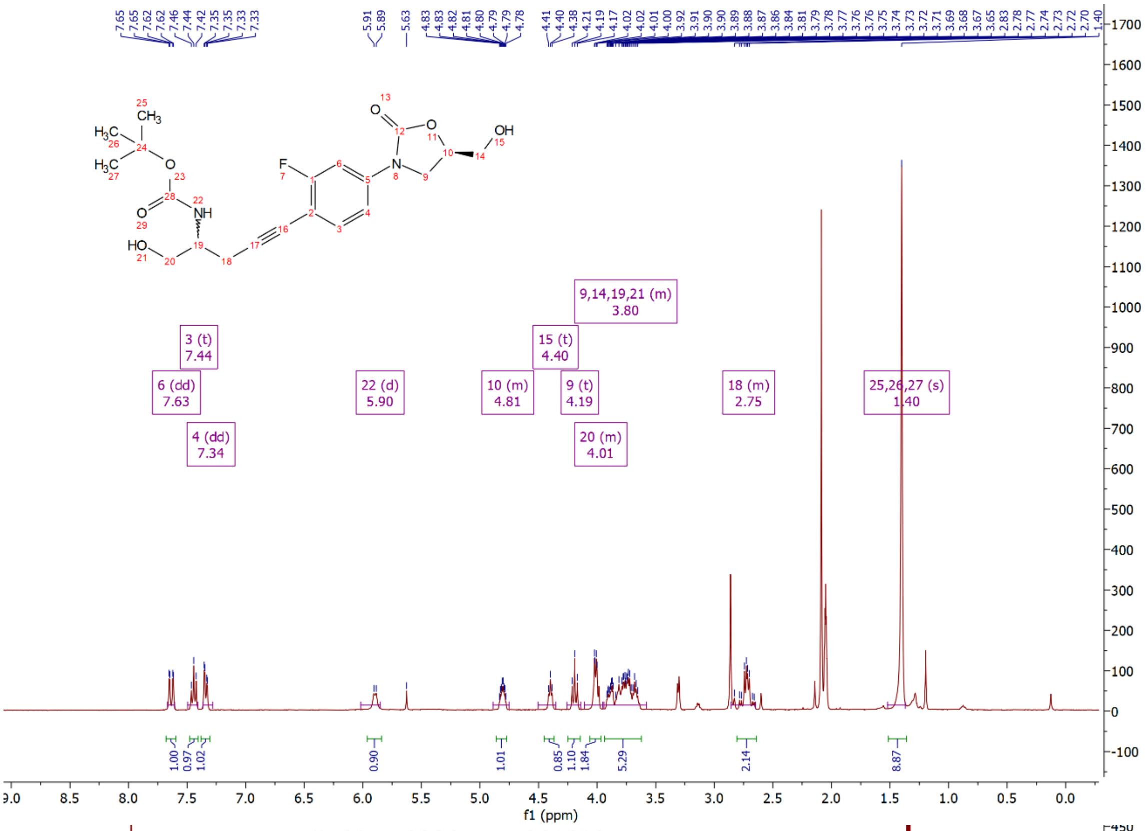

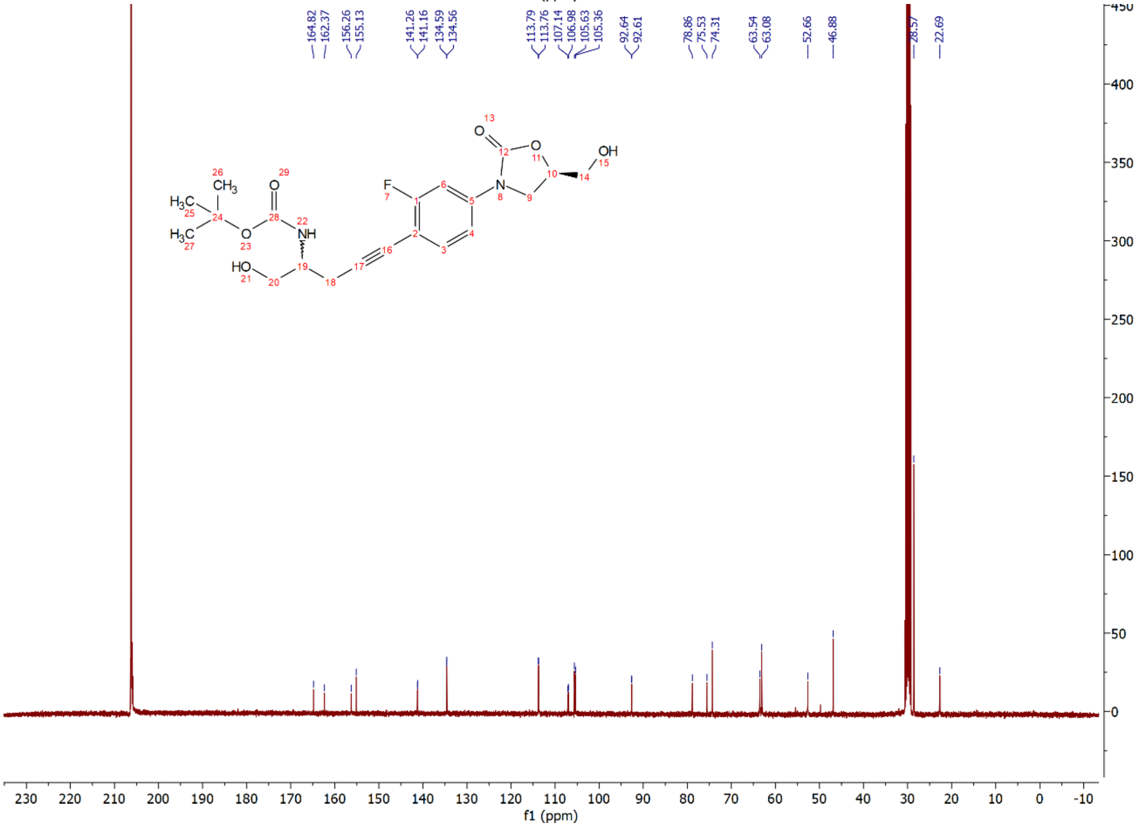

Compound **15ta**: $^{1}H$ and $^{13}C$ NMR Spectrum

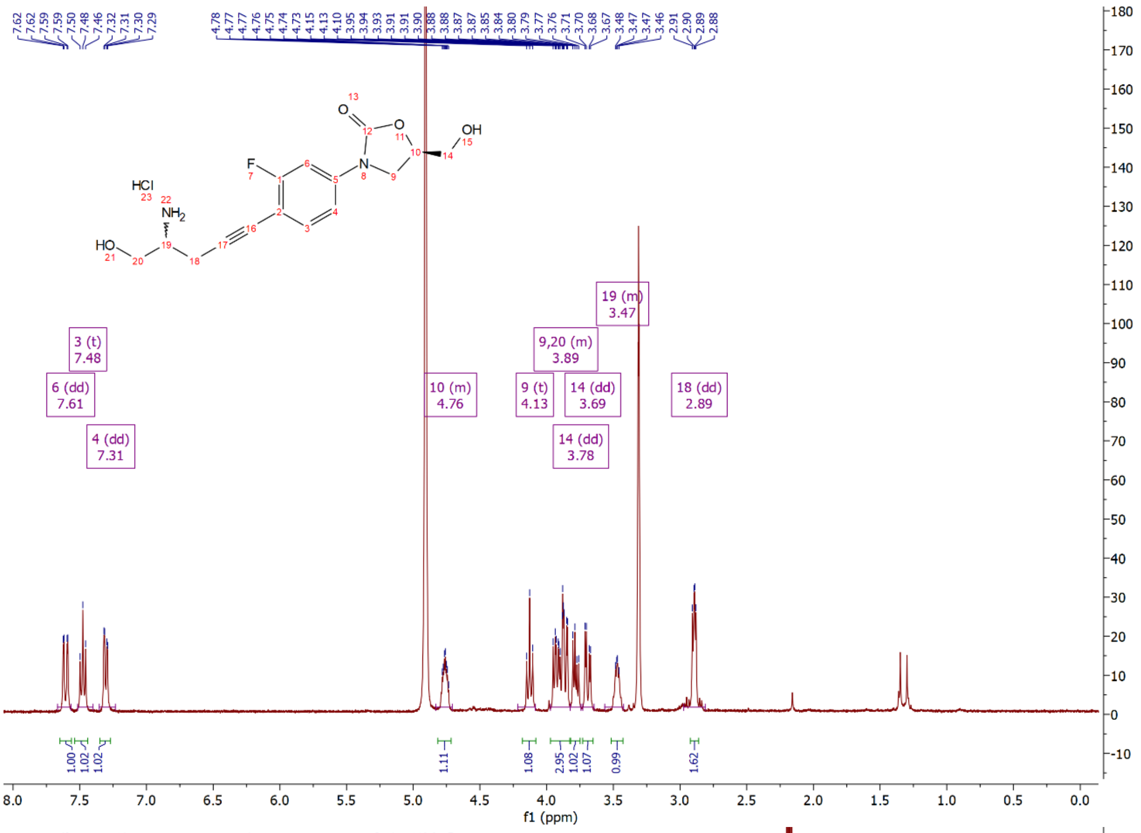

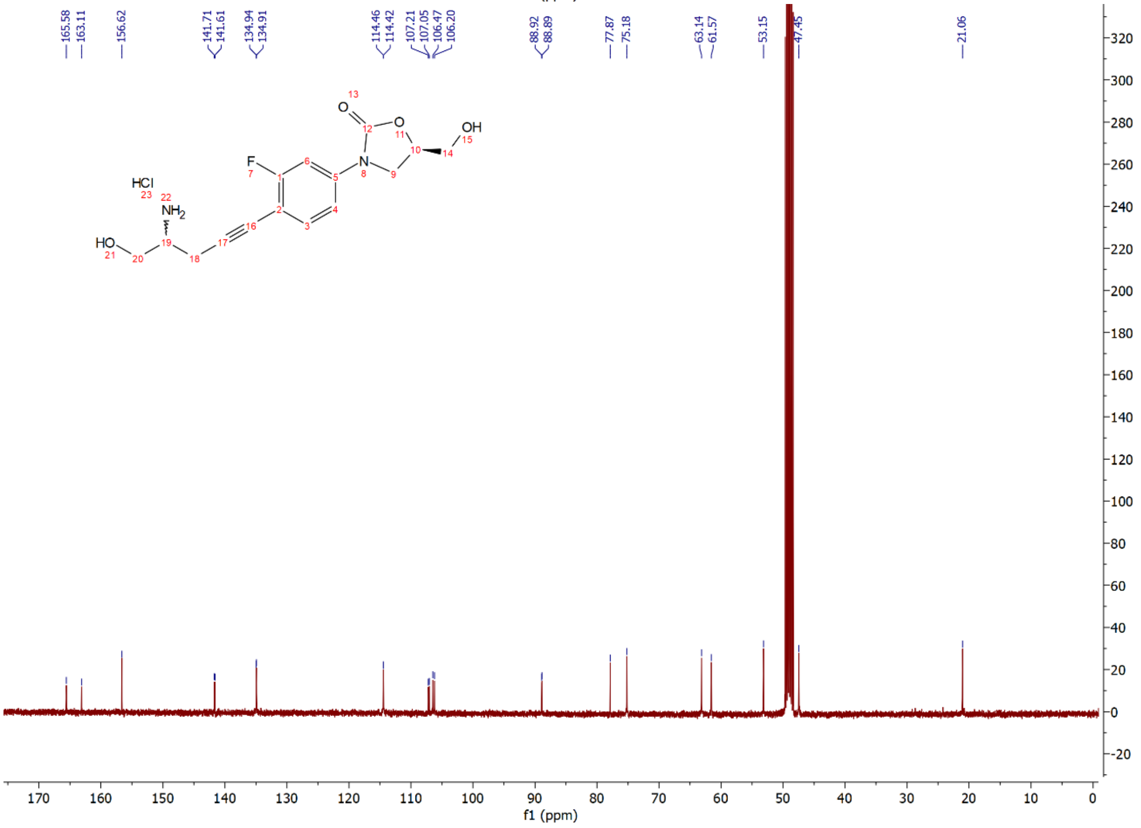

Compound **15u**: $^{1}$H and $^{13}$C NMR Spectrum

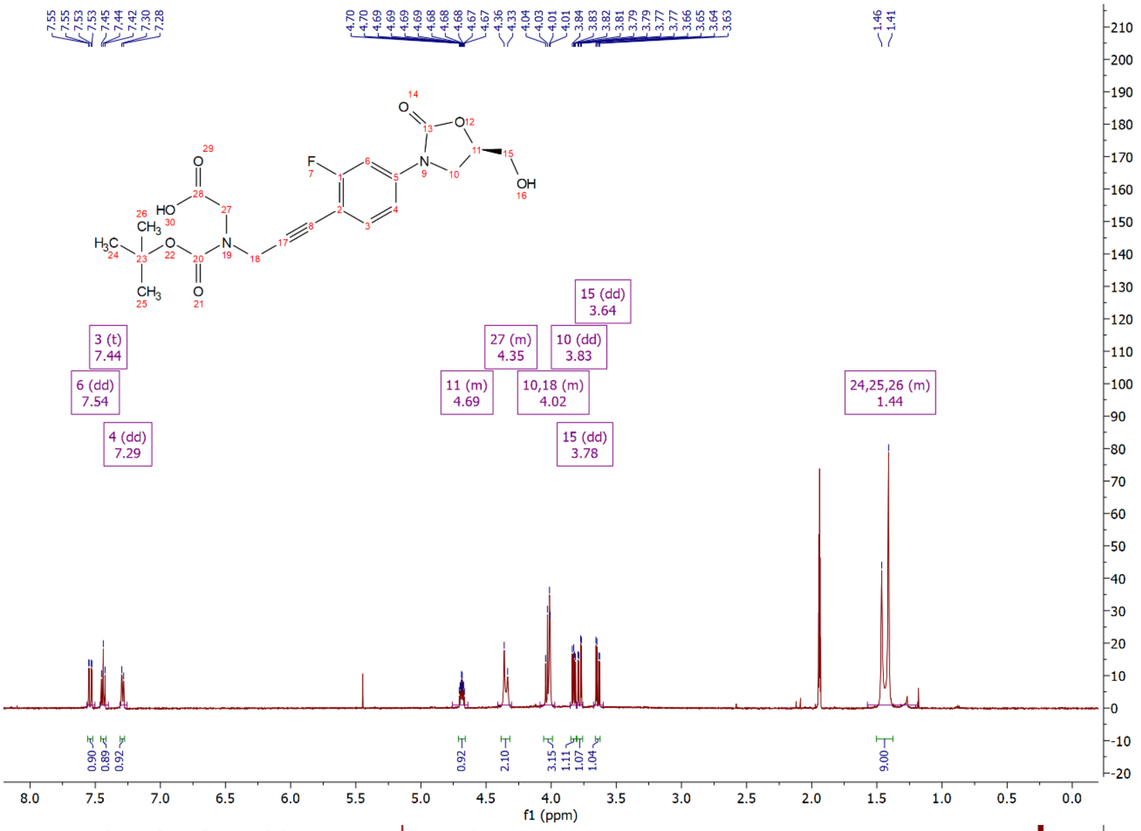

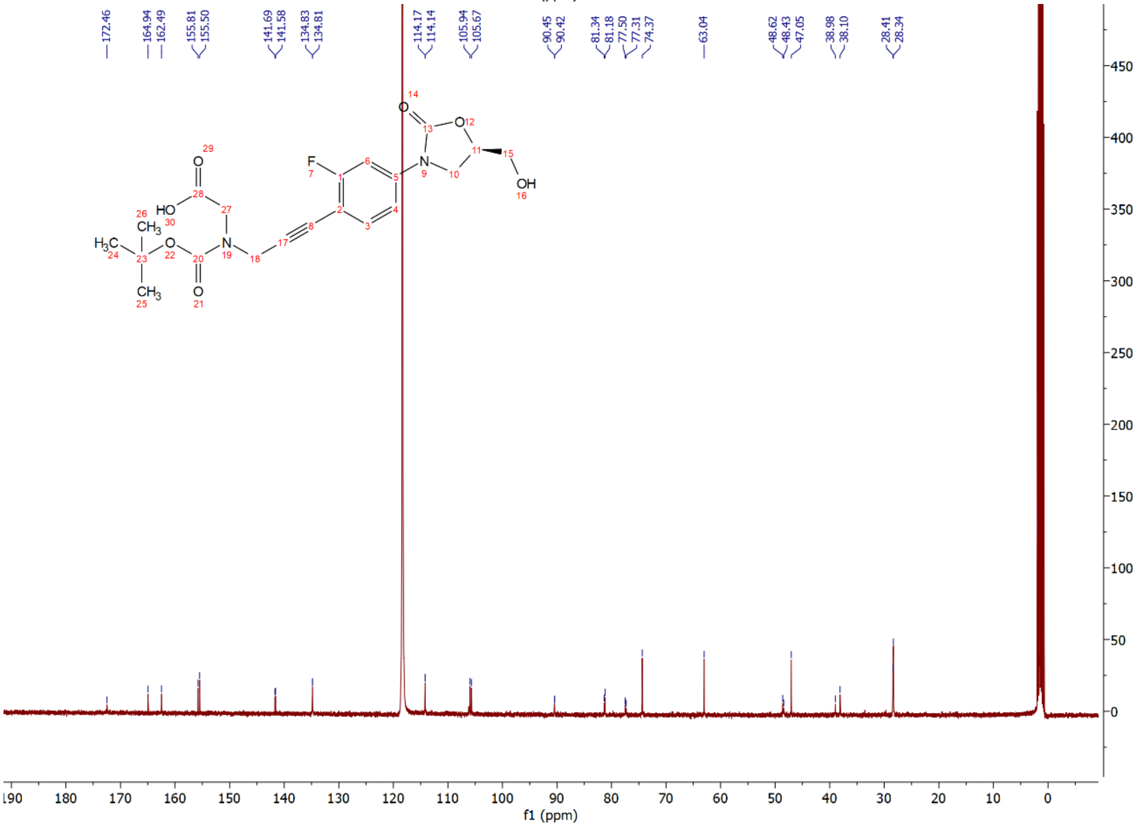

Compound **15ua**: $^{1}$H and $^{13}$C NMR Spectrum

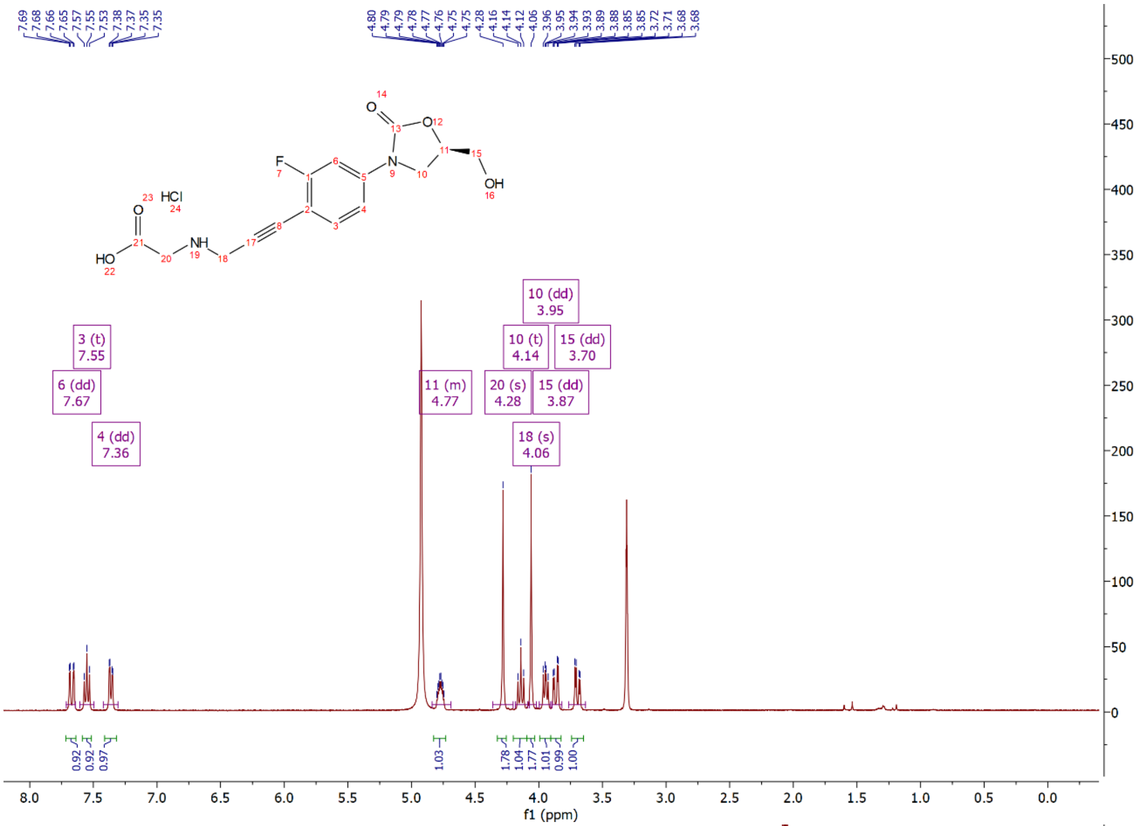


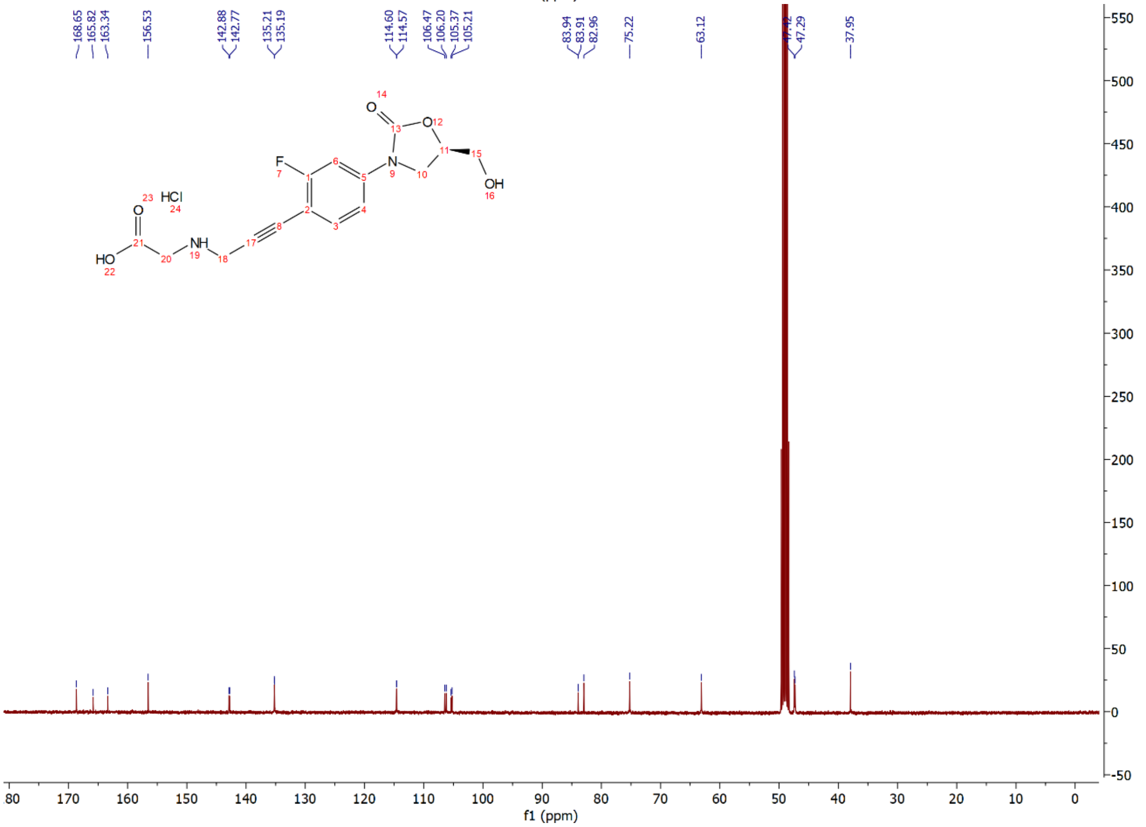

Compound **15v**: $^{1}H$ and $^{13}C$ NMR Spectrum

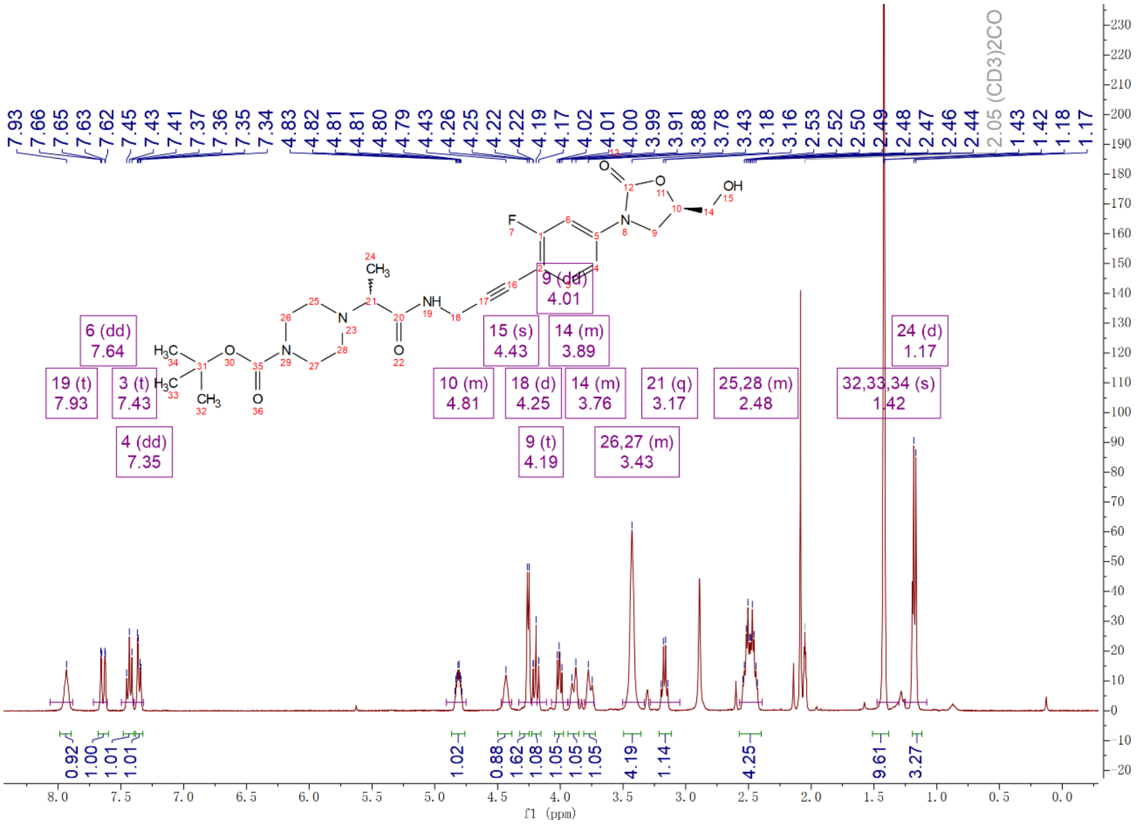

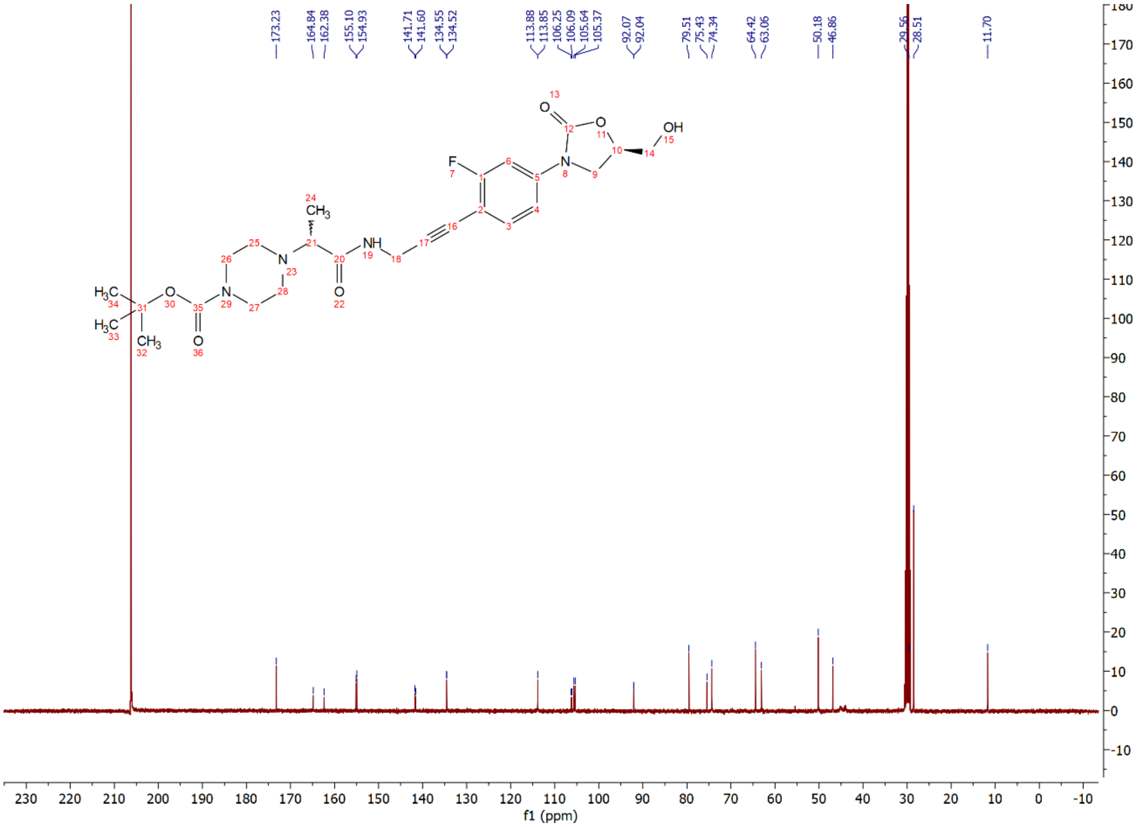

Compound **15va**: $^{1}$H and $^{13}$C NMR Spectrum

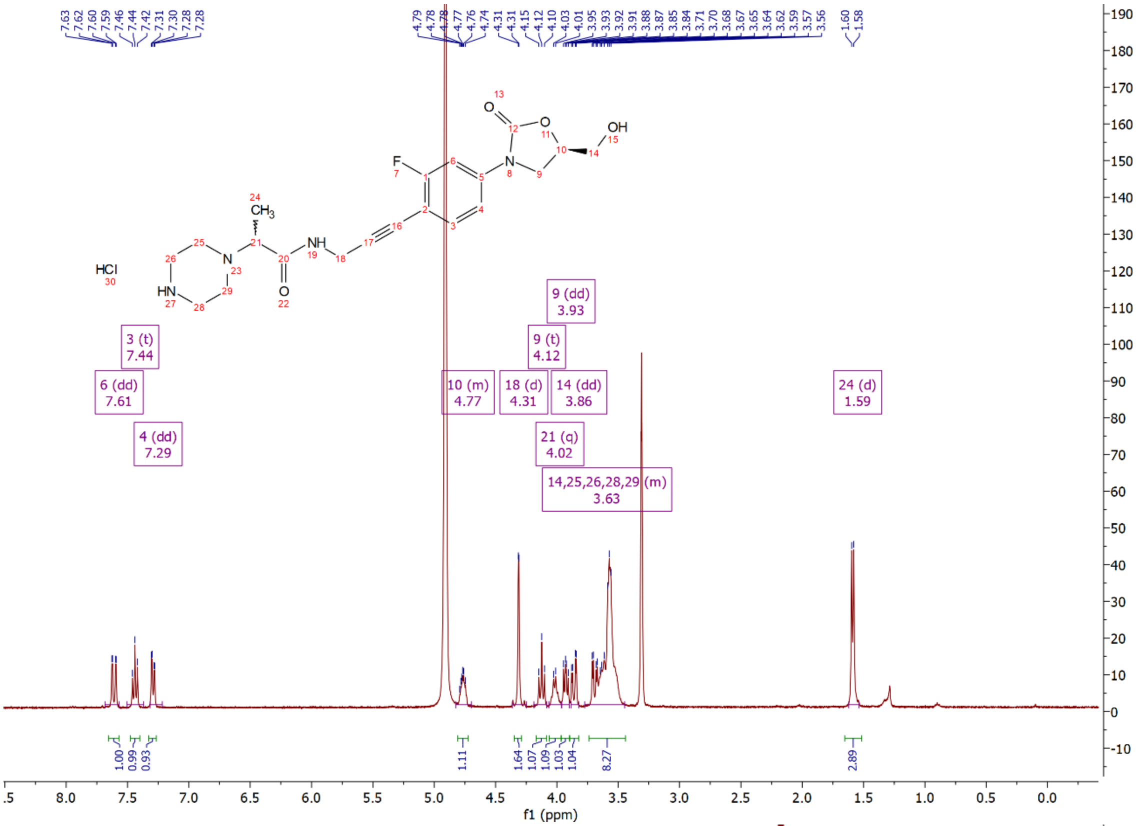

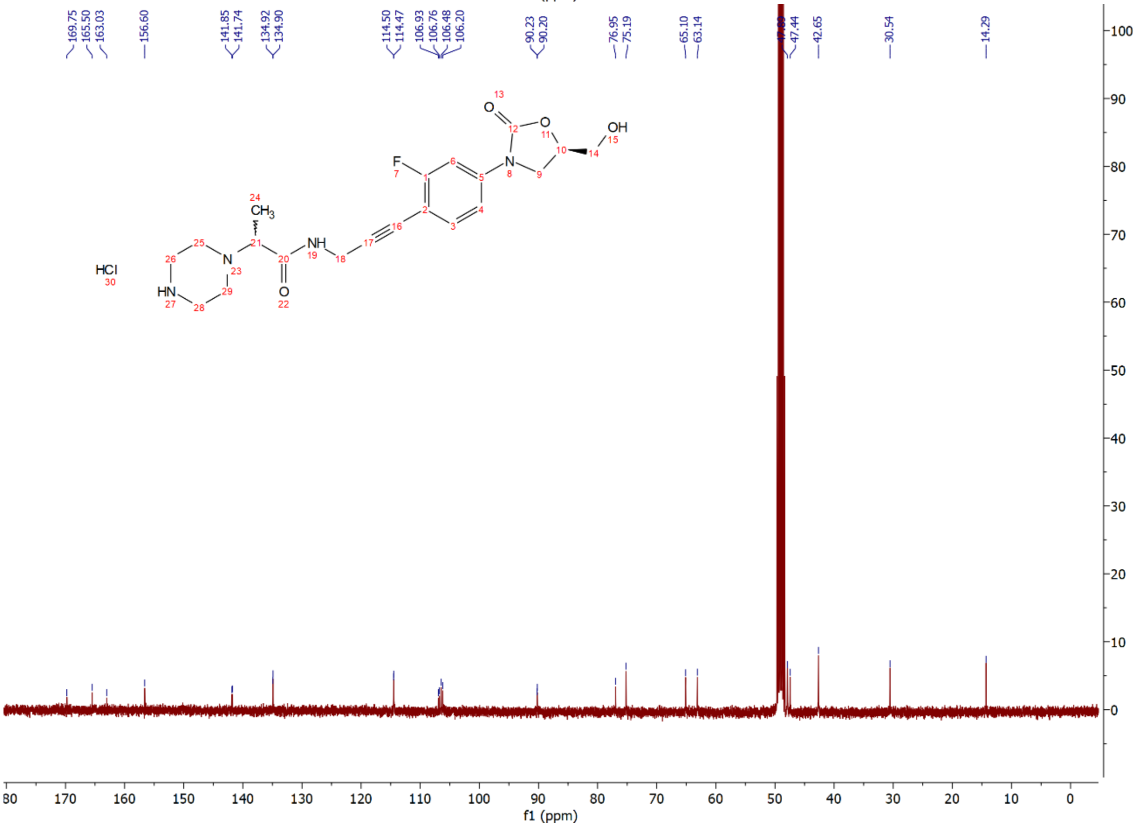

Compound **17a**: $^{1}$H and $^{13}$C NMR Spectrum

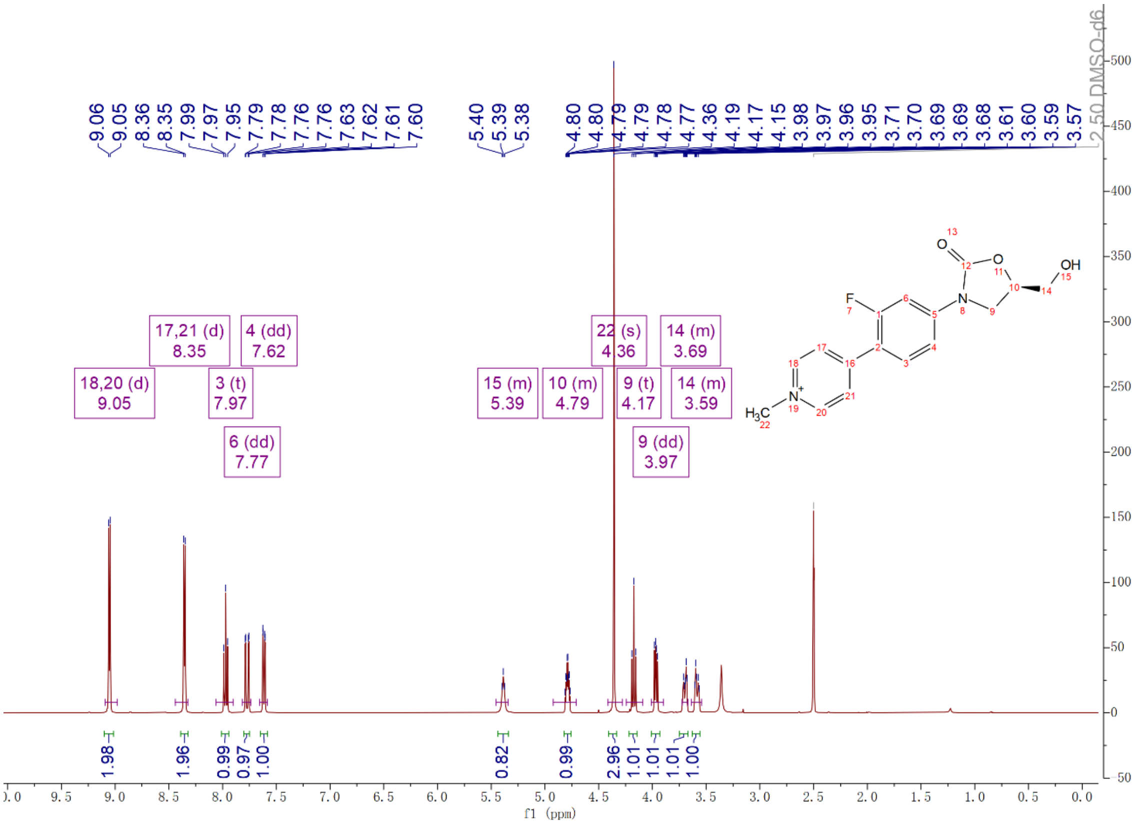

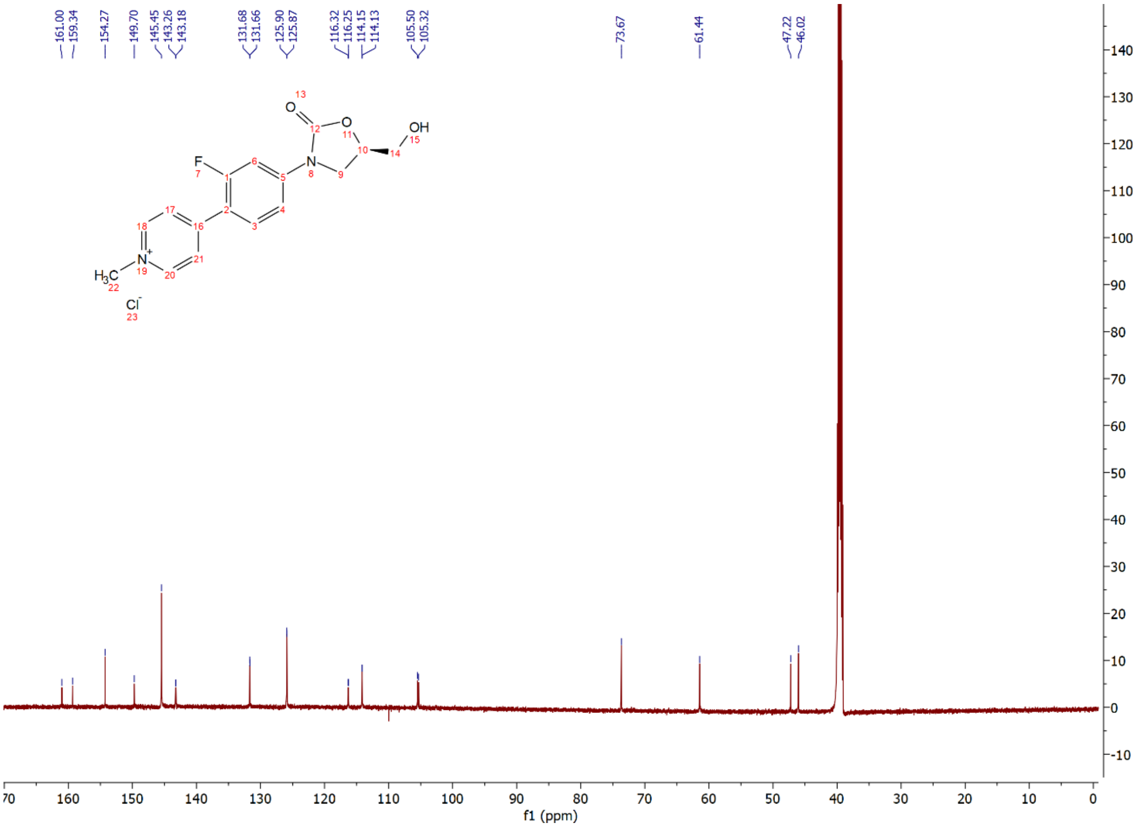

Compound **17b**: $^{1}$H and $^{13}$C NMR Spectrum

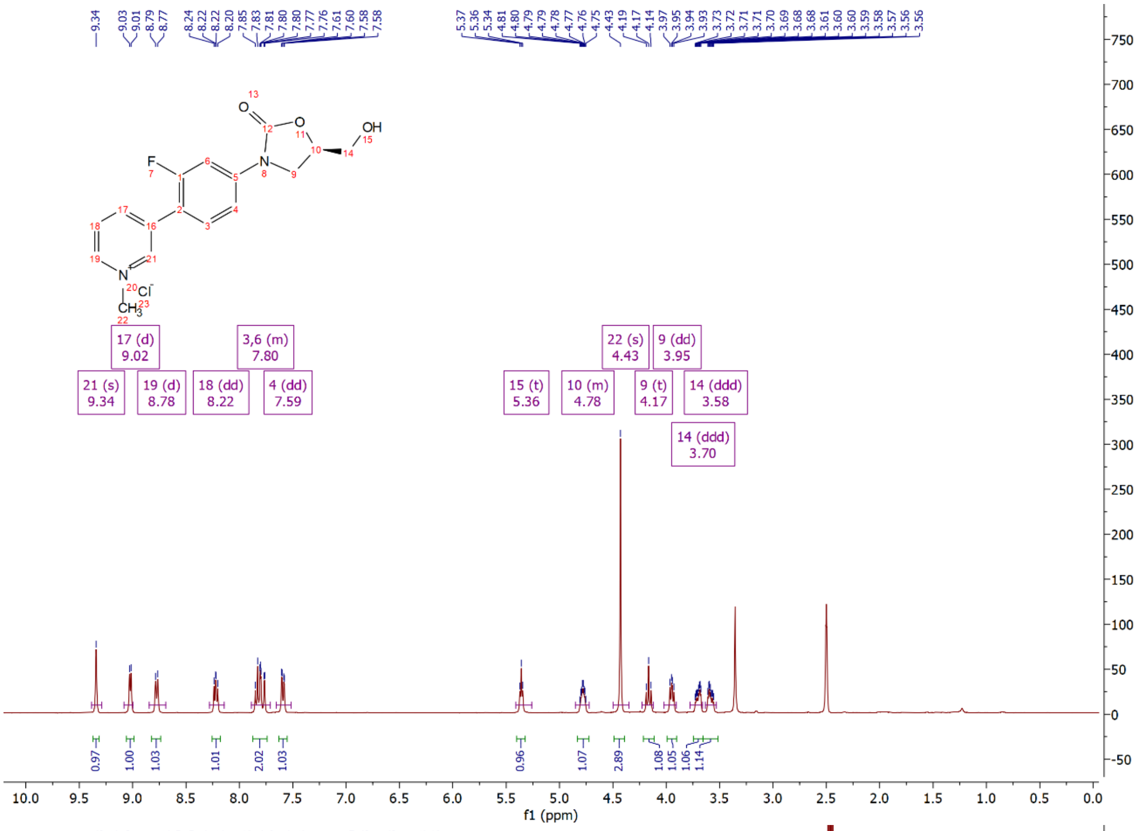

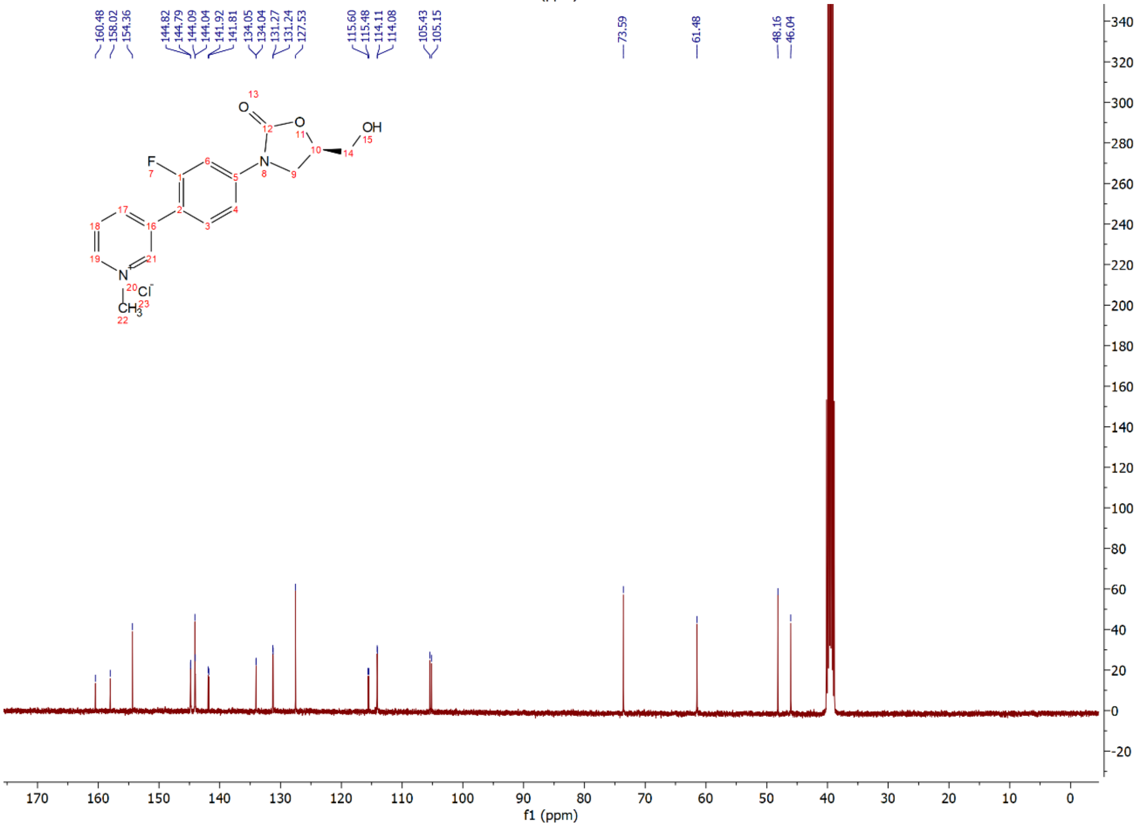

Compound **17c**: $^{1}$H and $^{13}$C NMR Spectrum

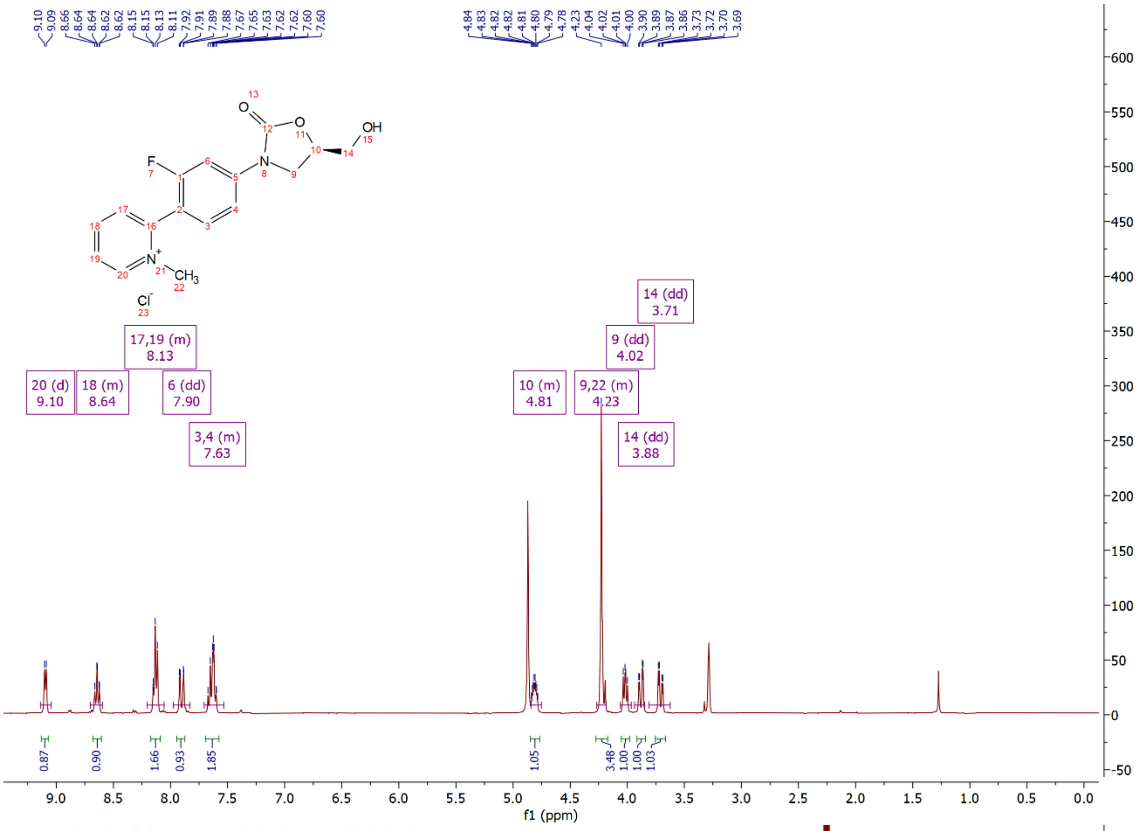

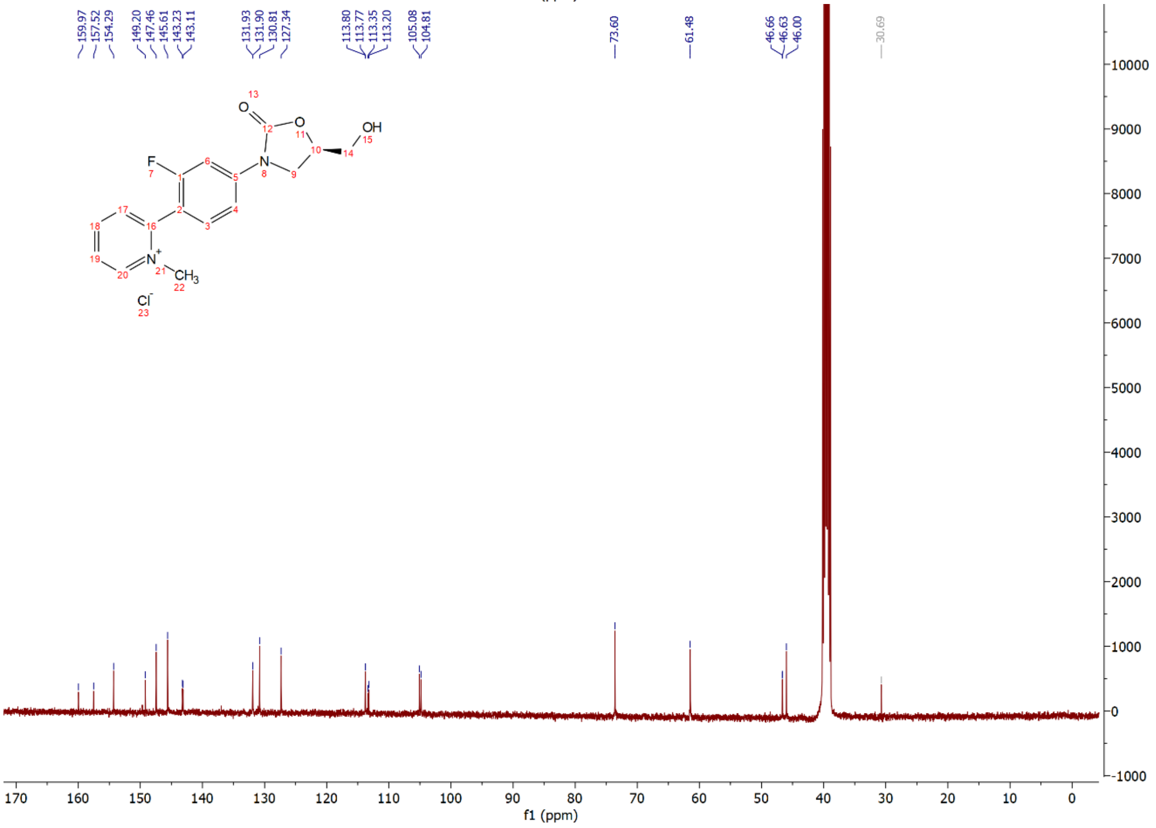

Compound **17d**: $^{1}$H and $^{13}$C NMR Spectrum

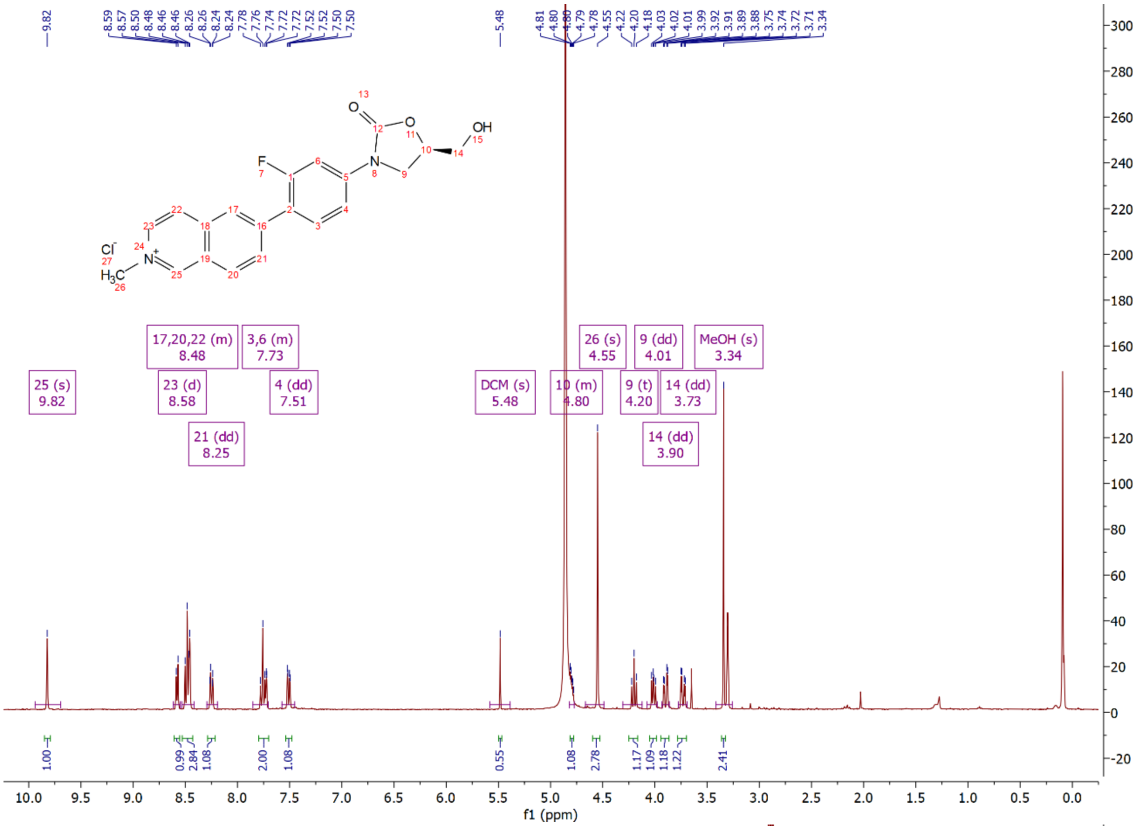

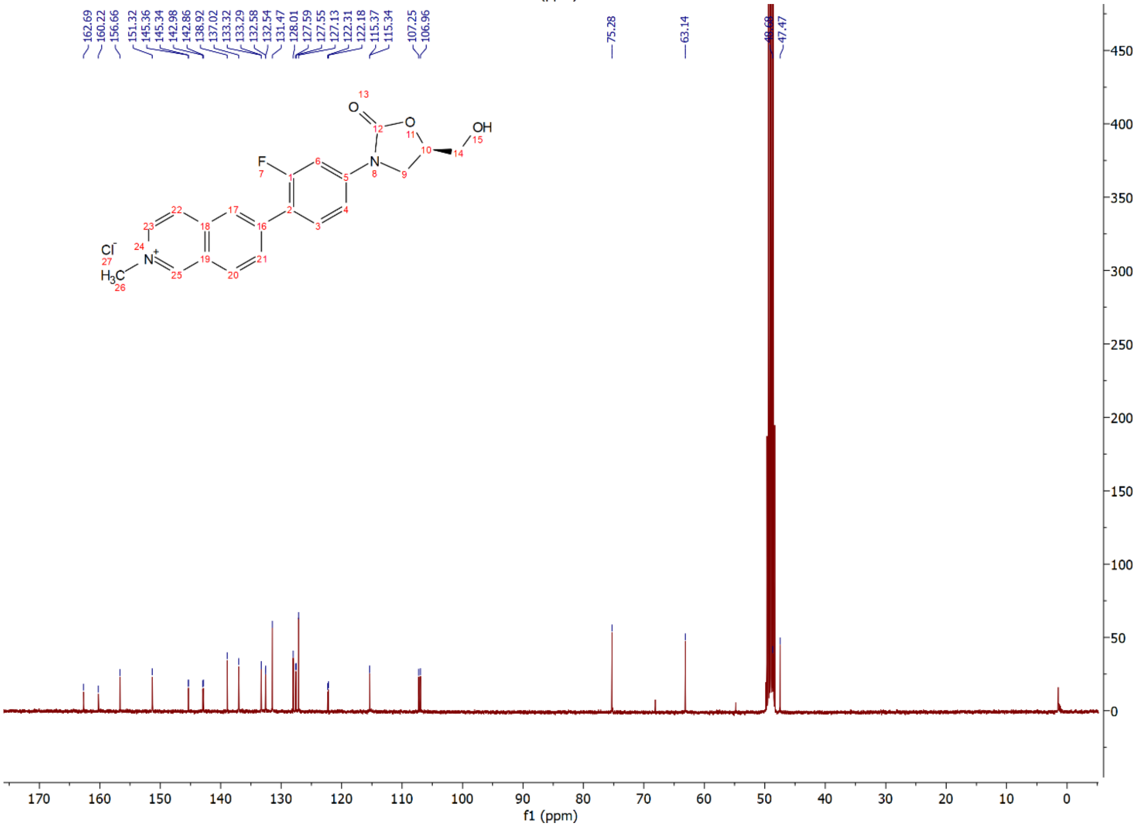

Compound **17e**: $^{1}$H and $^{13}$C NMR Spectrum

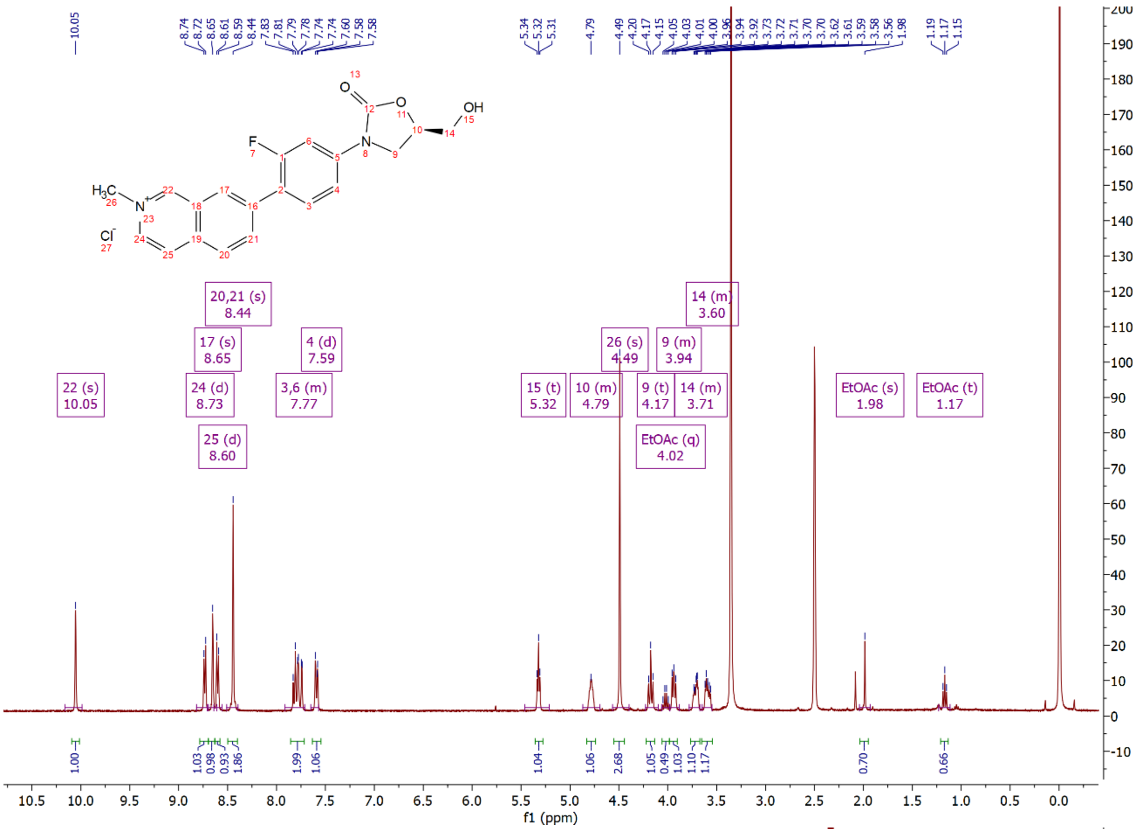

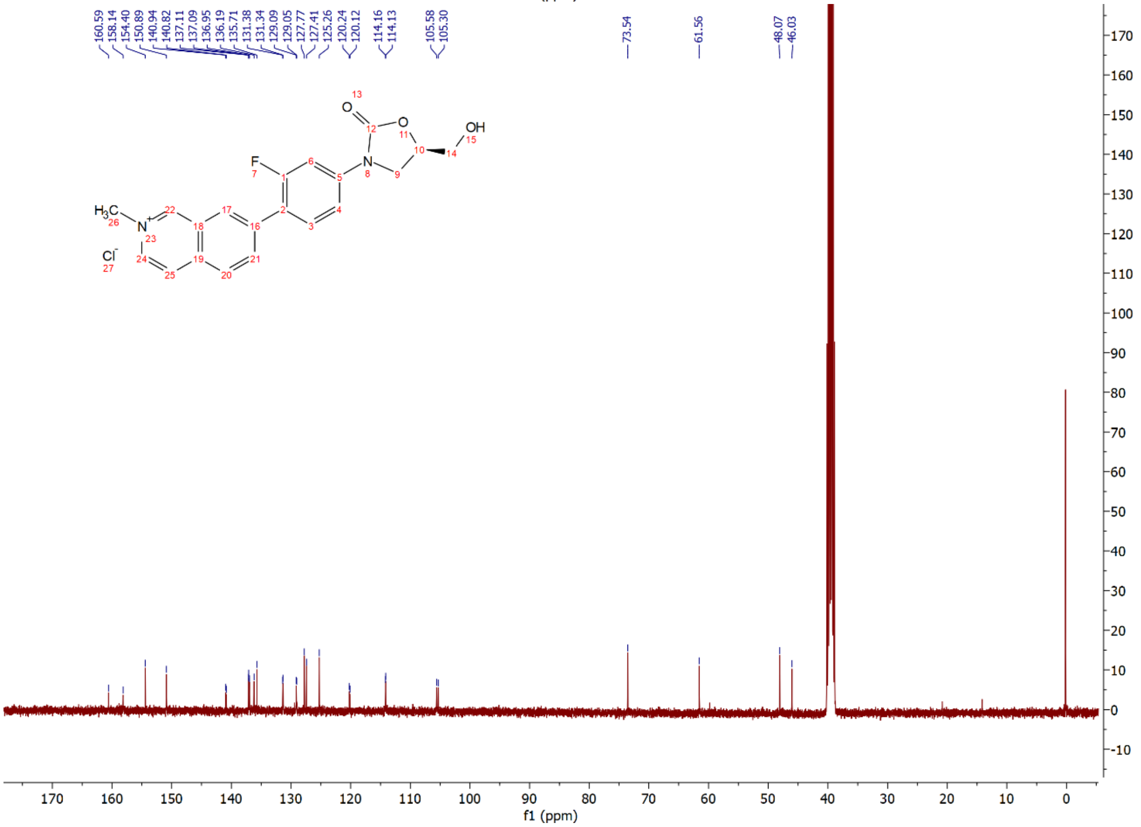

Compound **17f**: $^{1}H$ and $^{13}C$ NMR Spectrum

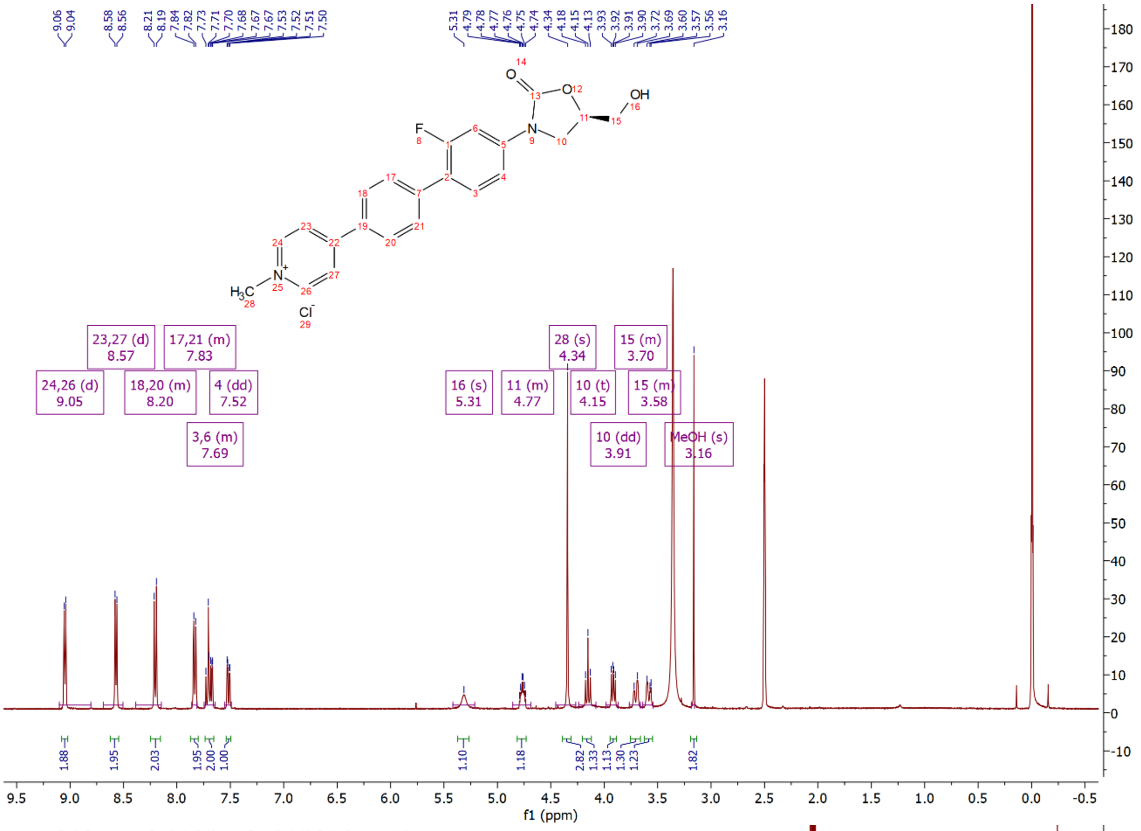

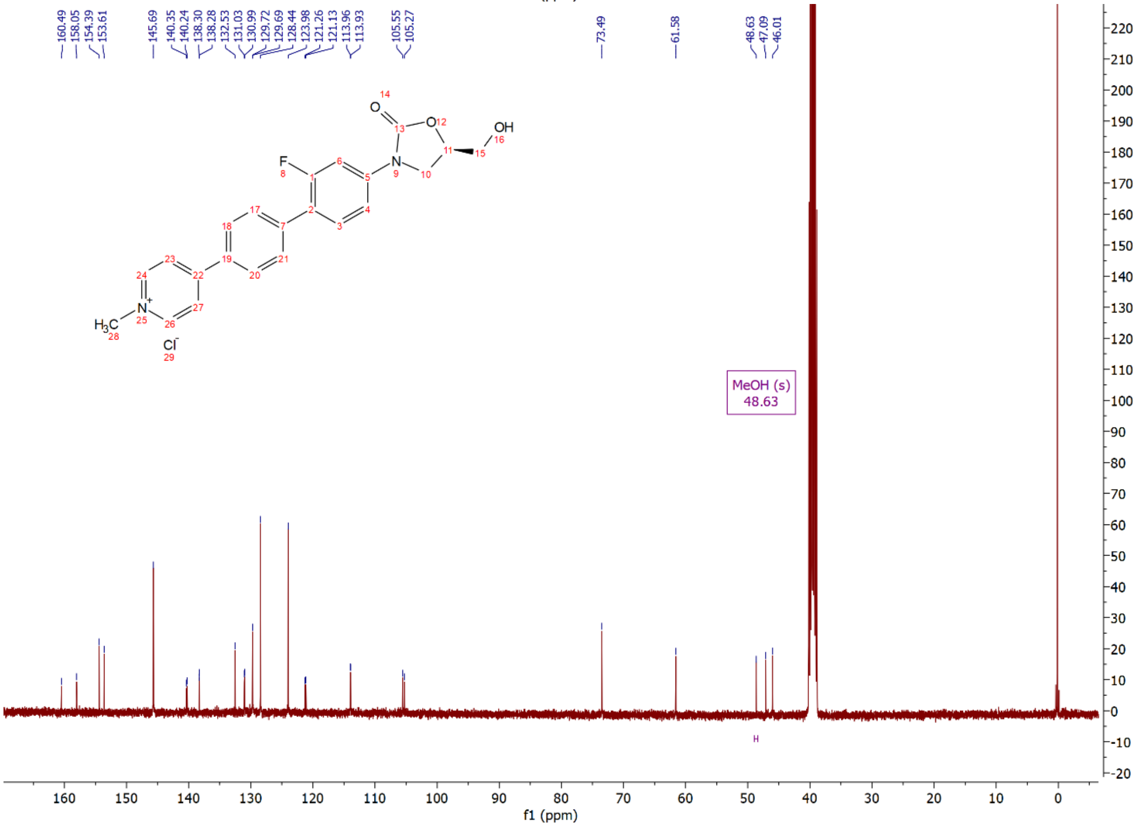

Compound **17g**: $^{1}H$ and $^{13}C$ NMR Spectrum

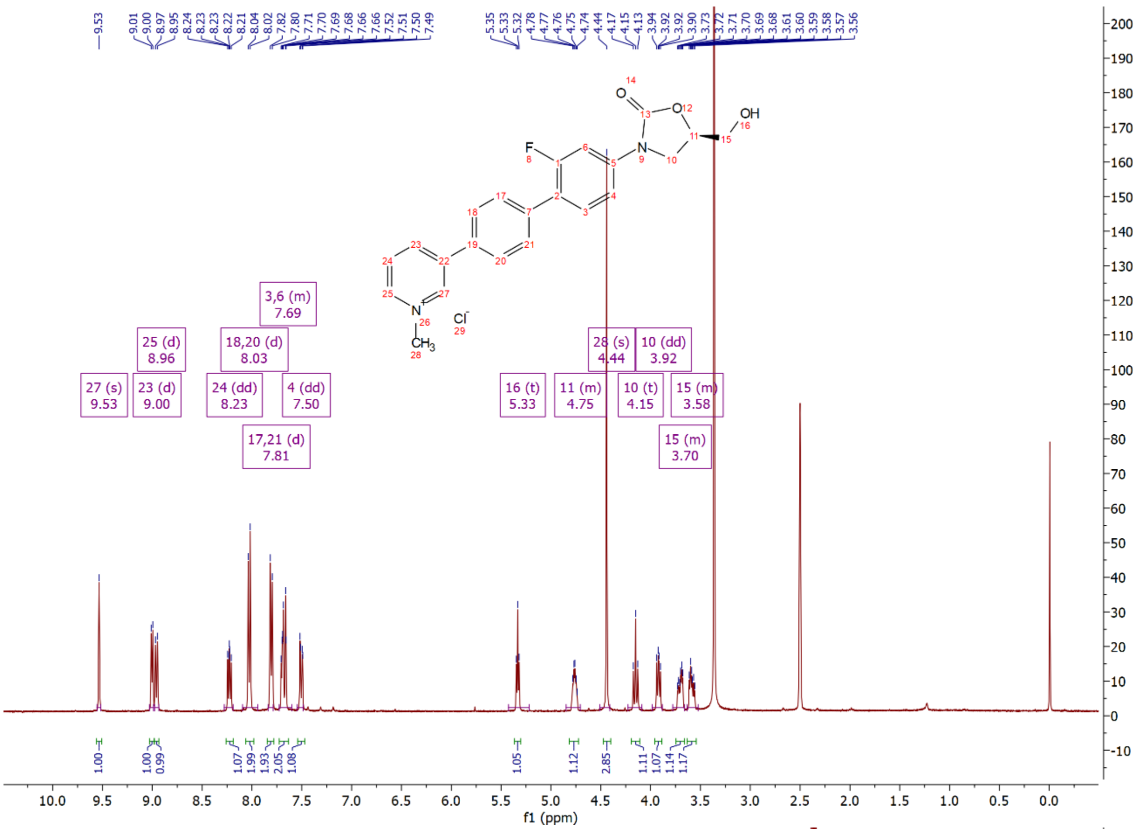

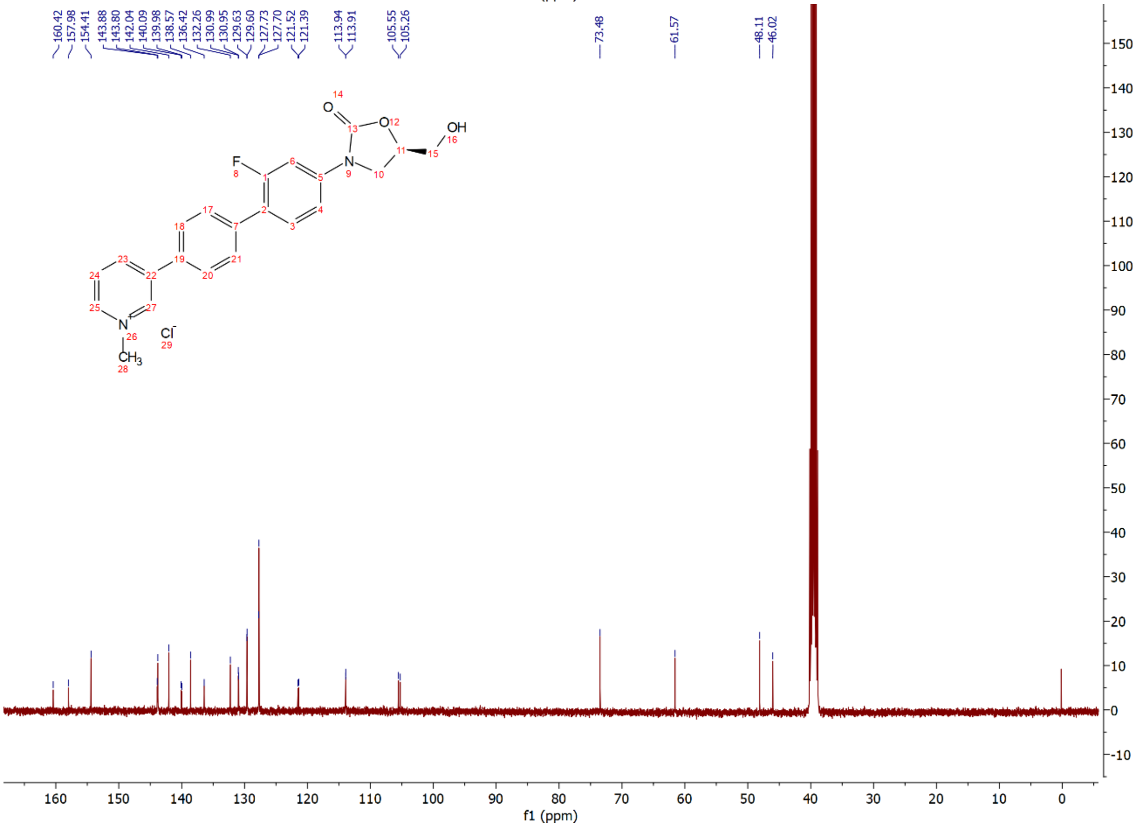

Compound **17h**: $^{1}$H and $^{13}$C NMR Spectrum

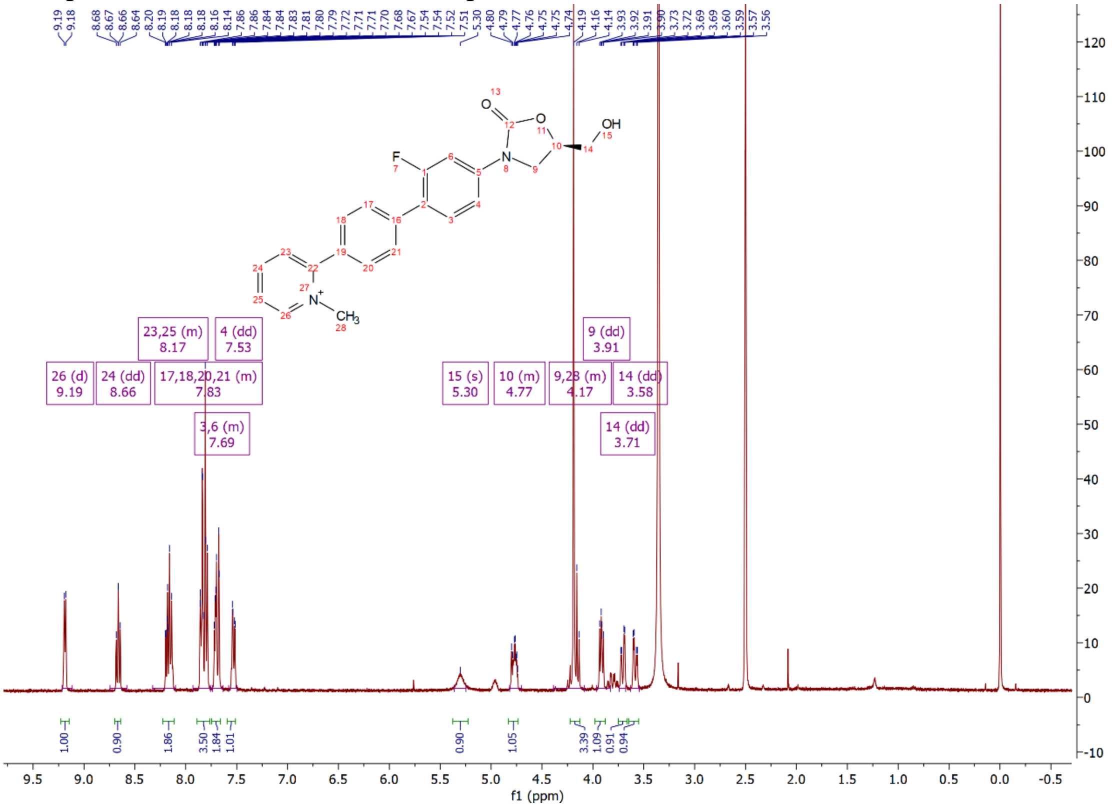

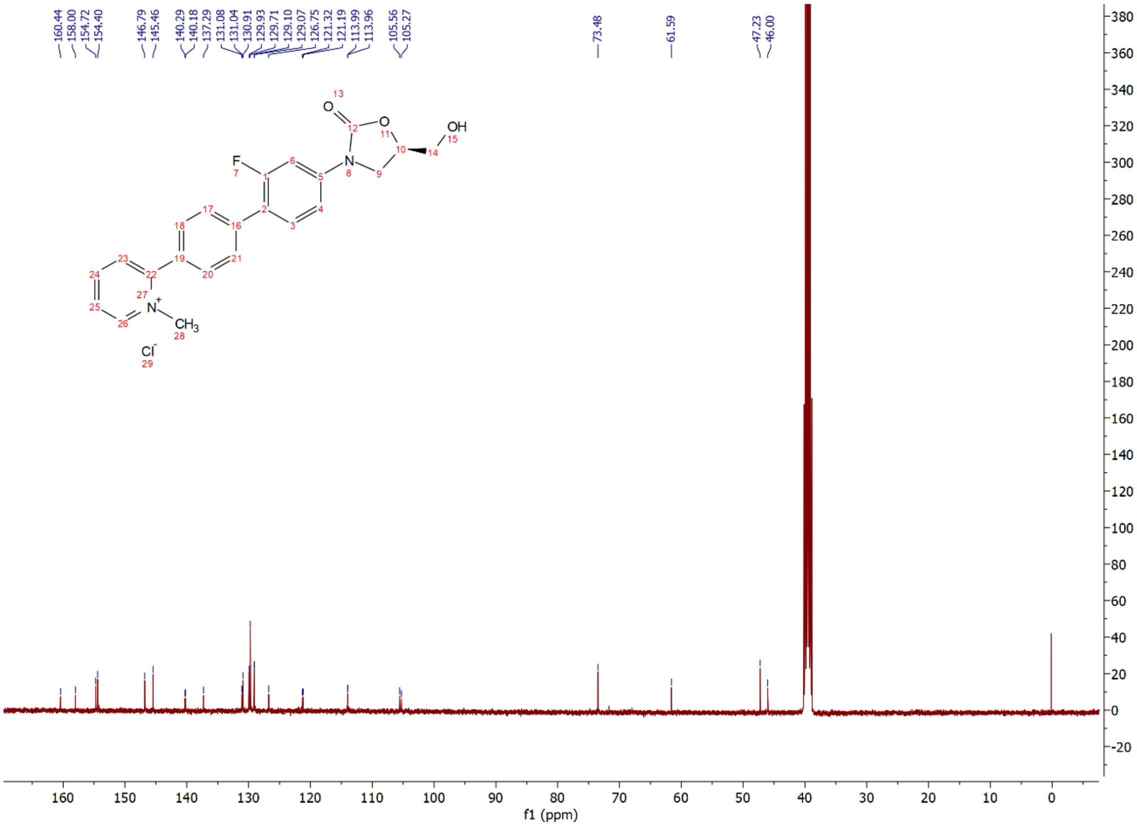

Compound **17i**: [1]H and [13]C NMR Spectrum

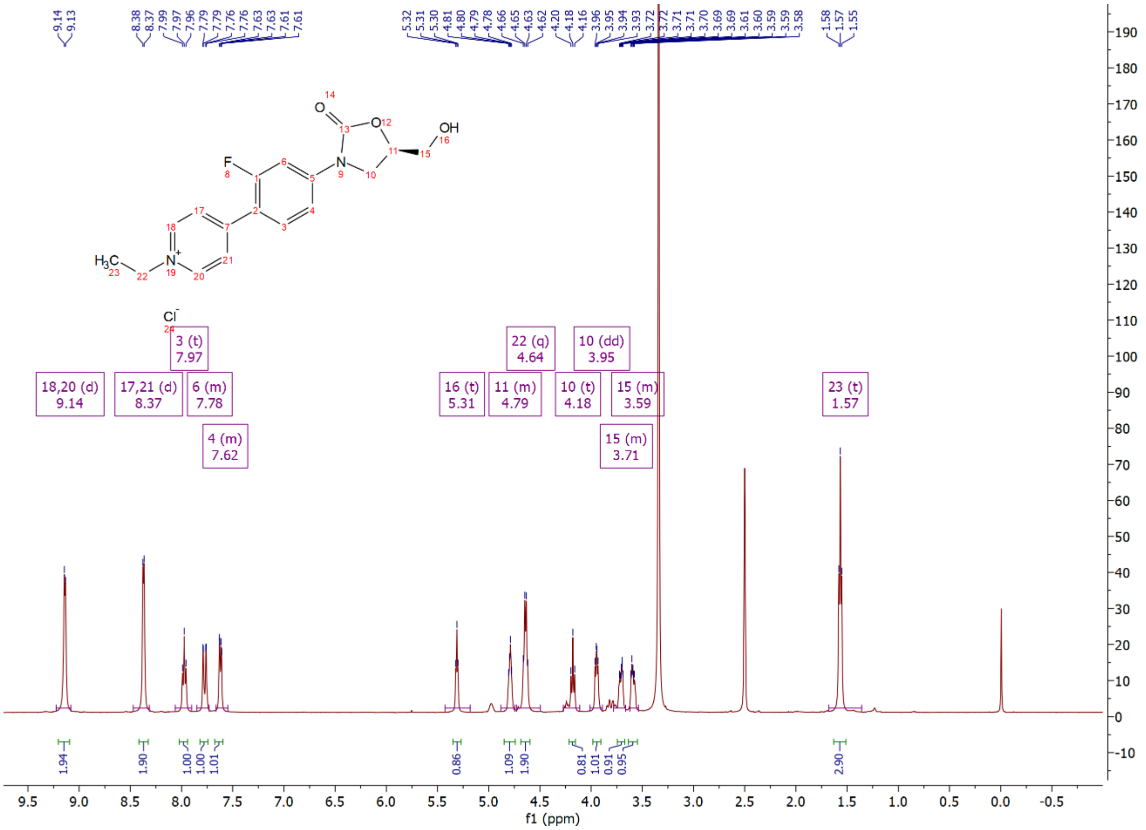

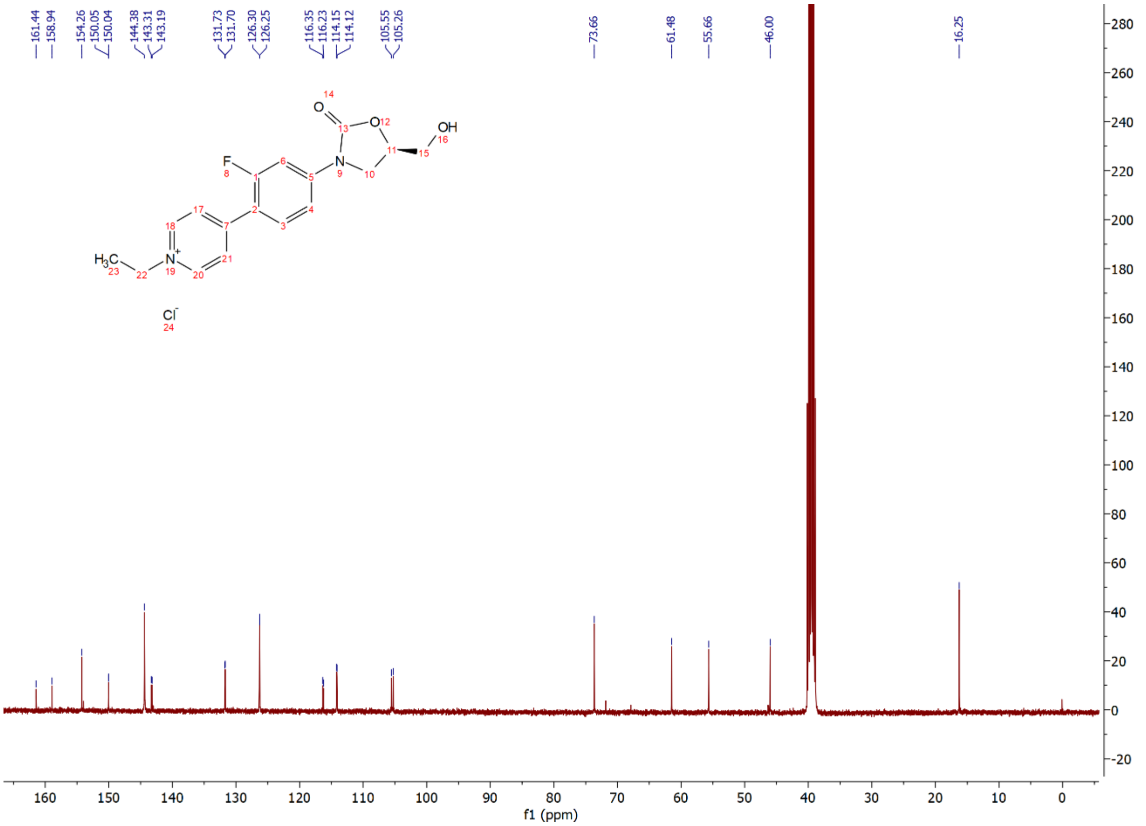

Compound **17j**: $^{1}$H and $^{13}$C NMR Spectrum

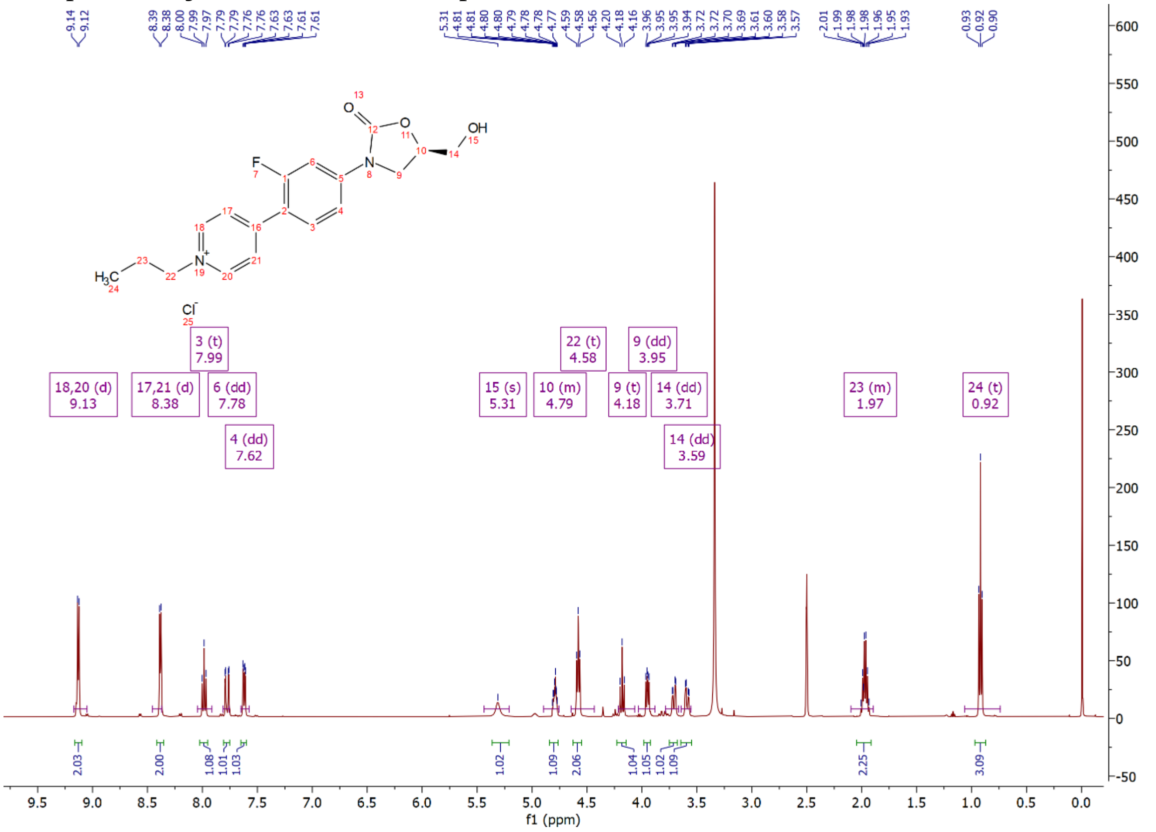

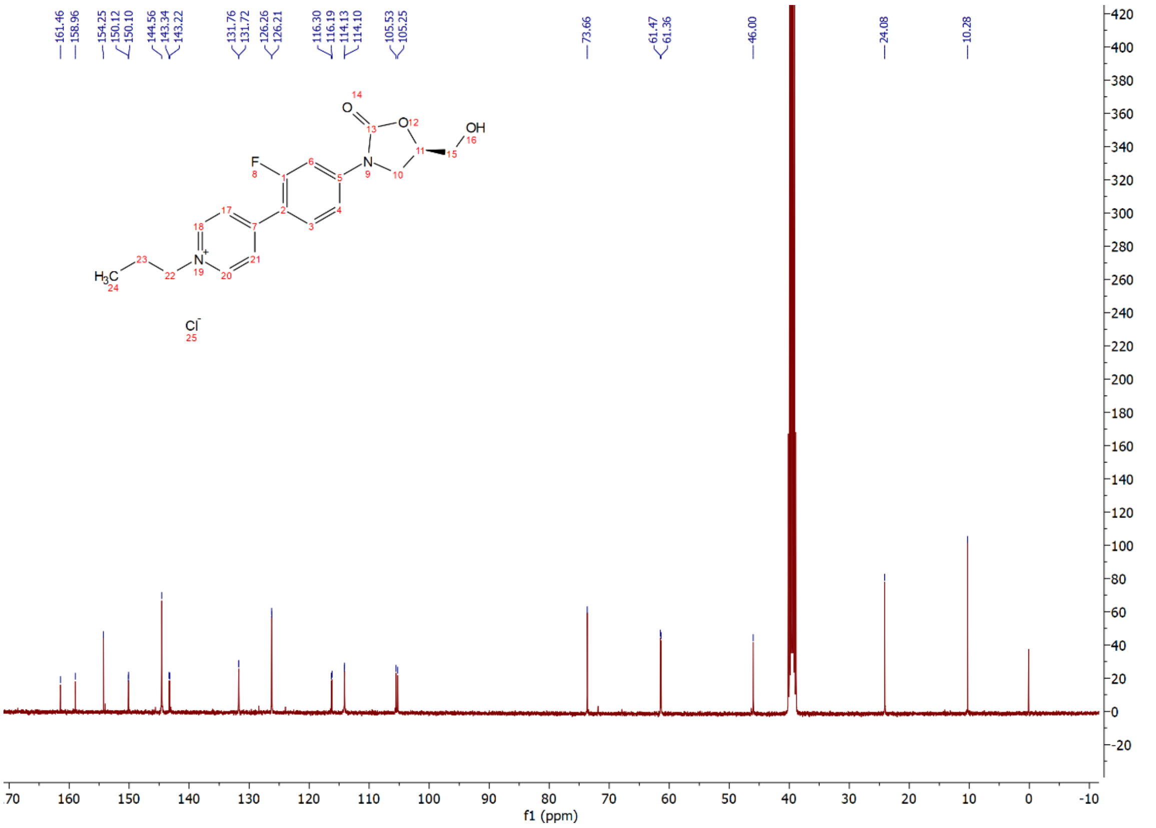

Compound **17k**: $^{1}$H and $^{13}$C NMR Spectrum

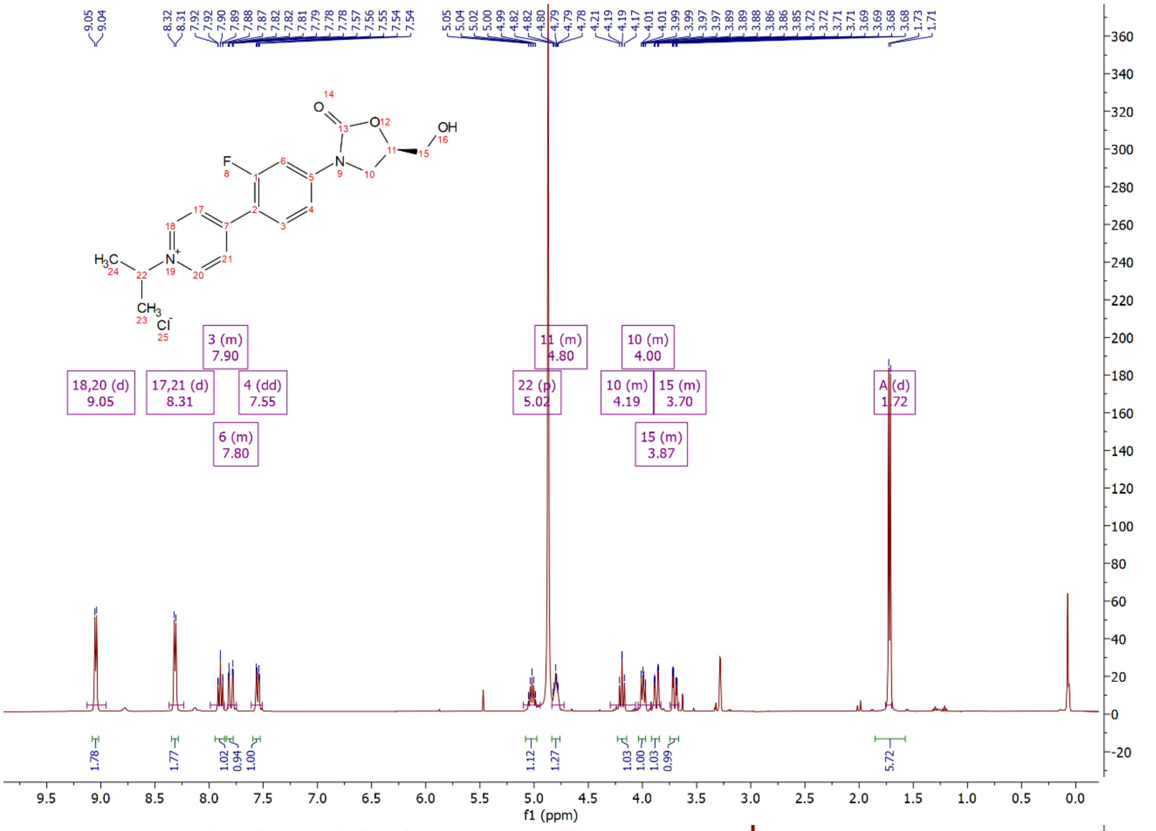

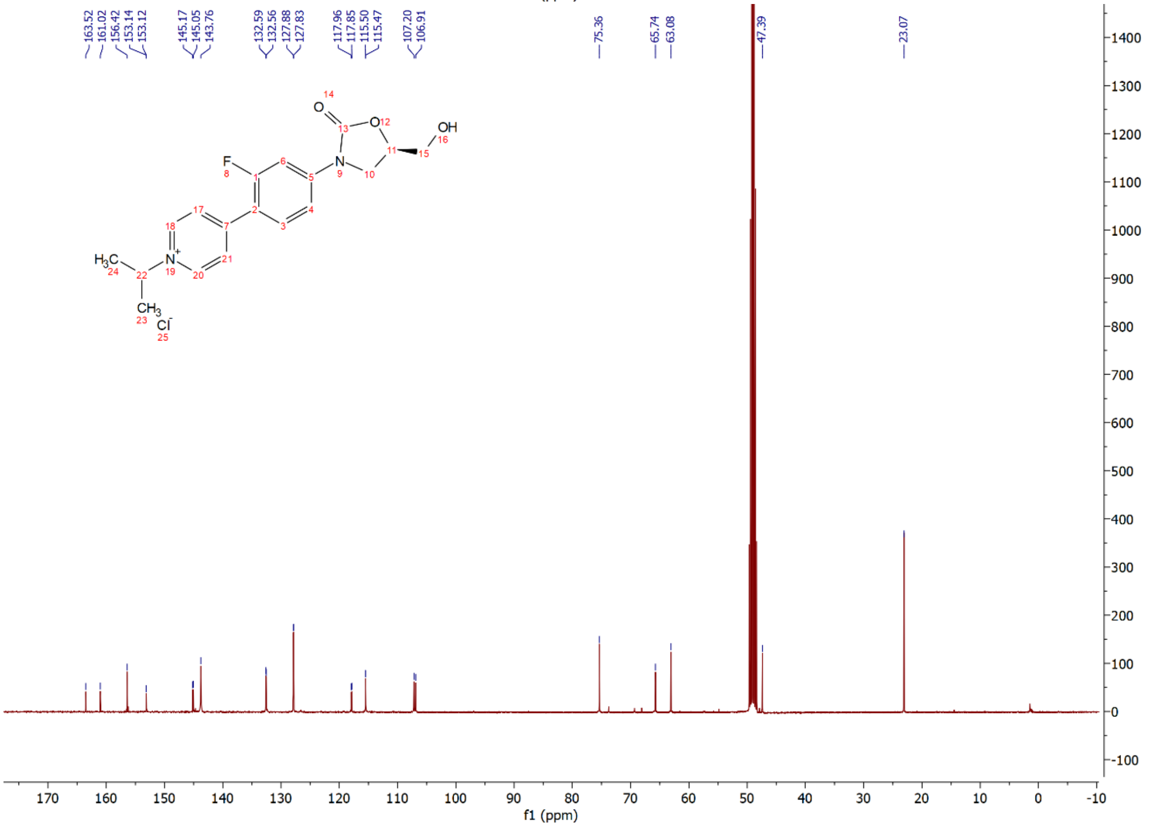

Compound **17l**: $^{1}$H and $^{13}$C NMR Spectrum

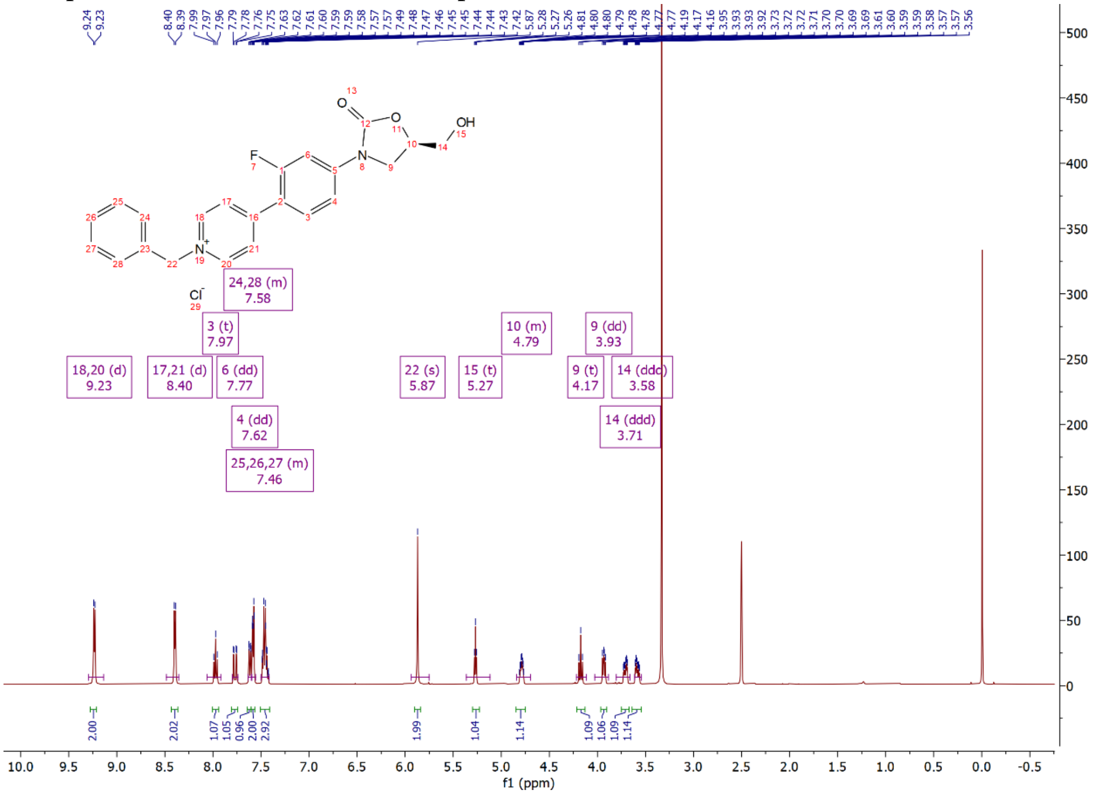

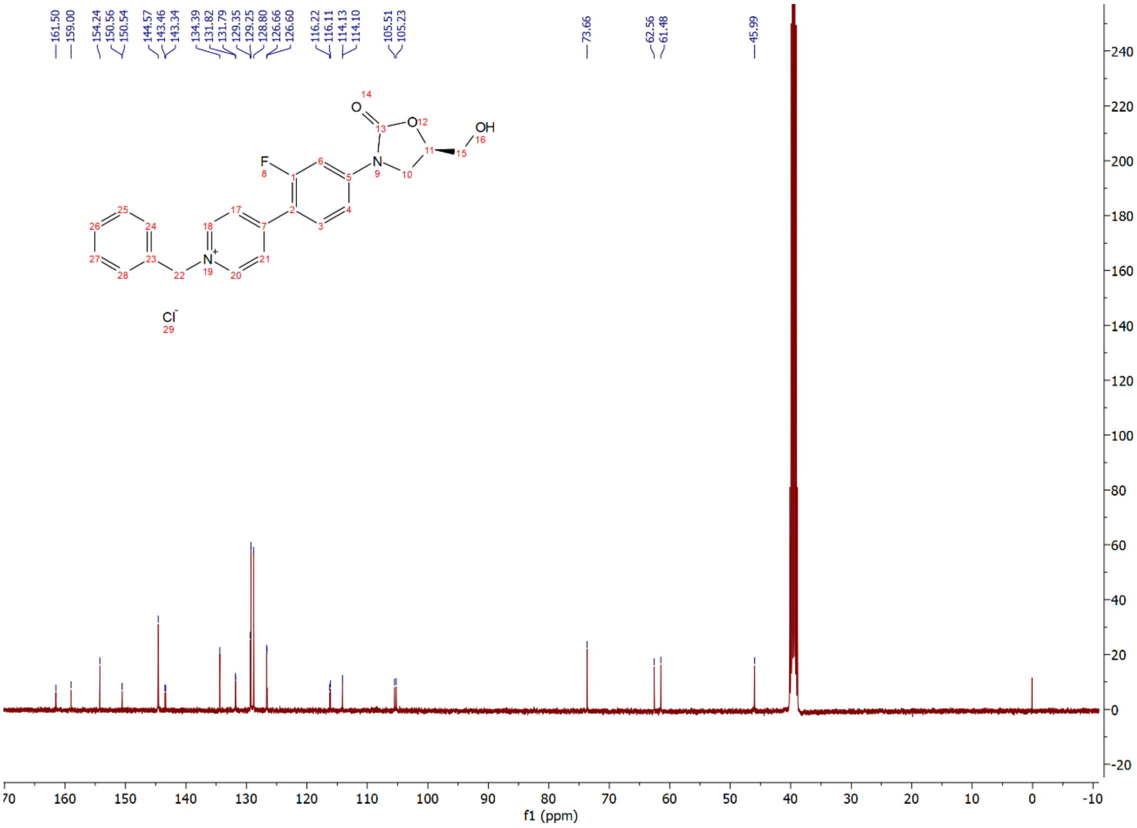
